# Comparative Transcriptomics and Genomics from Continuous Axenic Media Growth Identifies *Coxiella burnetii* Intracellular Survival Strategies

**DOI:** 10.1101/2023.02.06.527305

**Authors:** Archana Yadav, Melissa N. Brewer, Mostafa S. Elshahed, Edward I. Shaw

## Abstract

*Coxiella burnetii* (Cb) is an obligate intracellular pathogen in nature and the causative agent of acute Q fever as well as chronic diseases. In an effort to identify genes and proteins crucial to their normal intracellular growth lifestyle, we applied a “Reverse evolution” approach where the avirulent Nine Mile Phase II strain of Cb was grown for 67 passages in chemically defined ACCM-D media and gene expression patterns and genome integrity from various passages was compared to passage number one following intracellular growth. Transcriptomic analysis identified a marked downregulation of the structural components of the type 4B secretion system (T4BSS), the general secretory (sec) pathway, as well as 14 out of 118 previously identified genes encoding effector proteins. Additional downregulated pathogenicity determinants genes included several chaperones, LPS, and peptidoglycan biosynthesis. A general marked downregulation of central metabolic pathways was also observed, which was balanced by a marked upregulation of genes encoding transporters. This pattern reflected the richness of the media and diminishing anabolic and ATP-generation needs. Finally, genomic sequencing and comparative genomic analysis demonstrated an extremely low level of mutation across passages, despite the observed Cb gene expression changes following acclimation to axenic media.

## Introduction

*Coxiella burnetii* (Cb), the causative agent of acute and chronic Q fever (Arricau-Bouvery and Rodolakis 2005; Maurin and Raoult 1999; McQuiston, Childs and Thompson 2002; Miller, Shaw and Thompson 2006; Raoult, Marrie and Mege 2005; van Schaik, Chen, Mertens, *et al*. 2013), is an obligate intracellular pathogen that infects macrophages, and successfully propagates in a parasitophorous vacuole termed the *Coxiella* containing vacuole (CCV) (Heinzen, Hackstadt and Samuel 1999; Rudolf Toman 2012; Voth and Heinzen 2007). Cb has evolved multiple strategies to tolerate and thrive in the CCV, in spite of the prevailing low pH (≈ 4.5), low O2 content, oxygen radicals, and high level of degradative host factors such as acid hydrolases and defensins (Brennan, Russell, Zhang, *et al*. 2004; Hackstadt and Williams 1981; Heinzen, Hackstadt and Samuel 1999; Omsland, Cockrell, Howe, *et al*. 2009). Such remarkable ability has been the subject of a wide range of studies that employed a plethora of biochemical, genetic, imaging, and –omics-based approaches. Further, Cb employs a type 4B secretion system (T4BSS) to deliver effector proteins into the host throughout infection (Heinzen, Hackstadt and Samuel 1999; Rudolf Toman 2012; van Schaik, Chen, Mertens, *et al*. 2013; Voth and Heinzen 2007; Voth and Heinzen 2009). Cb effector proteins identified so far mediate a variety of biochemical activities and are known to target and modulate a broad array of host functions (Beare, Gilk, Larson, *et al*. 2011; Beare, Sandoz, Larson, *et al*. 2014; Crabill, Schofield, Newton, *et al*. 2018; Larson, Beare, Voth, *et al*. 2015; Larson and Heinzen 2017; Newton, Kohler, McDonough, *et al*. 2014; Newton, McDonough and Roy 2013; van Schaik, Chen, Mertens, *et al*. 2013). Prior studies have employed bioinformatic tools (Chen, Banga, Mertens, *et al*. 2010), transposon mutagenesis (Beare, Gilk, Larson, *et al*. 2011; Carey, Newton, Luhrmann, *et al*. 2011; Crabill, Schofield, Newton, *et al*. 2018; Martinez, Cantet, Fava, *et al*. 2014; Weber, Chen, Rowin, *et al*. 2013), microscopic localization studies (Chen, Banga, Mertens, *et al*. 2010; Howe, Melnicakova, Barak, *et al*. 2003; Morgan, Luedtke and Shaw 2010; Voth, Beare, Howe, *et al*. 2011) and cloning and infectivity testing to identify and characterize effector proteins. In addition, *Legionella pneumophila*, a close genetic neighbor of Cb with a very similar T4BSS (Nagai and Kubori 2011; Segal and Shuman 1999; Sexton and Vogel 2002), is known to use T4BSS-effector protein duality to infect its natural host cell, the amoeba. *L. pneumophila* has been extensively used as a proxy to identify putative effector proteins and propose molecular pathogenesis mechanisms in Cb (Pan, Lührmann, Satoh, *et al*. 2008; Segal, Feldman and Zusman 2005; Vogel 2004; Zamboni, McGrath, Rabinovitch, *et al*. 2003; Zusman, Yerushalmi and Segal 2003). Indeed, research on *L. pneumophila* has identified the structural features of the T4BSS, the nature of effector proteins secreted through the system, and possible function of some of these effectors.

Growth of Cb in an axenic media was first reported in 2009 using the undefined Acidified Citrate Cysteine Media (ACCM) media (Omsland, Cockrell, Howe, *et al*. 2009). Increased replication rates in the somewhat more defined ACCM-2 medium soon followed in 2011 (Omsland, Beare, Hill, *et al*. 2011). Subsequently, a nutritionally fully defined media (ACCM-D) with an even greater replication rate and physiologic parallels to intracellular bacteria was developed (Sanchez, Vallejo-Esquerra and Omsland 2018). Growing Cb in axenic media is opening new venues for investigating mechanisms of Cb molecular pathogenesis (Beare and Heinzen 2014; Beare, Jeffrey, Long, *et al*. 2018; Beare, Larson, Gilk, *et al*. 2012; Crabill, Schofield, Newton, *et al*. 2018; Martinez, Cantet and Bonazzi 2015; Rudolf Toman 2012; Sandoz, Beare, Cockrell, *et al*. 2016). Theoretically, when grown in axenic media, the expression of genes required for intracellular survival and host cell manipulation is no longer required for Cb viability. As such, continuous maintenance and passaging the bacterium for extended periods of times under axenic conditions could potentially remove the powerful selective pressure exerted by the host cell, thus potentially minimize/silence expression in such genes. As such, we posit that transcriptomic analysis of gene expression patterns as well as genomic identification of mutation and gene loss patterns in axenic grown versus Cb cultures derived from intracellular growth could be employed for identifying putative involvement of specific genes, as well as identification of novel genes necessary for Cb pathogenesis and survival in an intracellular environment. Similarly, continuous passaging could also lead to the propagation of mutations, DNA fragment losses, and rearrangements in genes/loci associated with intracellular survival, pathogenesis, and host cell manipulation. Such patterns could be regarded as “reverse evolution” i.e., the opposite of the natural evolution trajectory of Cb from a free-living ancestor to an obligate intracellular pathogen. Specifically, we hypothesized that: 1) changes in gene expression within the first few passages upon transition from intracellular to axenic media growth would be observed, and such differences would be more pronounced in genes involved in subverting and coopting host metabolism, as well as genes enabling general adaptation to physiological conditions prevalent in its intracellular vacuolar environment, and 2) Cb could acquire and accumulate DNA mutations upon transition from intracellular to axenic media growth after repetitive passages since certain bacterial genes/proteins are no longer required for successful growth.

In this study, we transitioned Cb Nine Mile phase II from cell cultures into axenic defined media ACCM-D and subcultured it into a long-term successive passage. We conducted transcriptomic and genomic sequencing on replicate samples at different time points (passages) to document temporal changes in gene expression patterns, and DNA mutations associated with adaptation to an axenic extracellular lifestyle.

## Materials and Methods

### Microorganism and growth conditions

*Coxiella burnetii* avirulent strain Nine Mile phase II (NMII), clone 4 (RSA439) was cultivated in rabbit epithelial RK13 cells (CCL-37; American Type Culture Collection) grown in Dulbecco’s modified Eagle medium DMEM (ThermoFisher Scientific) supplemented with 5% fetal bovine serum in T75 culture flasks. This method of collecting cells was adapted from (Coleman, Fischer, Howe, *et al*. 2004). Briefly, the infected cell line was split into multiple non-vented and capped T150 culture flasks that were incubated at 37°C in 5% CO_2_ for a week until confluent growth was observed. These flasks were then screwed tightly and left at room temperature for 2 weeks to induce cells to switch to the small cell variant (SCV) form. The cells were pelleted by ultracentrifugation (12,000 x g, 15 minutes) in 250 ml Nalgene round bottom tubes, scrapped off the round bottom tubes by using sterile 1X phosphate buffered saline (PBS) and then lysed by using Dounce homogenize. The lysed cells in PBS were then spun via centrifugation using Oakridge tubes in an ultracentrifuge at 12,000 x g for 15 minutes. The SCV pellets obtained were stored in SPG freezer media (0.7 M sucrose, 3.7 mM KH_2_PO_4_, 6.0mM K_2_HPO_4_, 0.15 M KCl, 5.0 mM glutamic acid, pH 7.4) at -80°C.

### Axenic growth in defined ACCM-D media

Cb cultures propagated intracellularly in rabbit epithelial RK13 cells were used to inoculate ACCM-D media (Sunrise Science Products, San Diego, CA). Approximately 10^6^ genome equivalents per mL was used as an inoculum (determined using the RT-PCR procedure as described (Brennan and Samuel 2003)). Cultures were grown in a T25 cell culture flasks at 5% O2, 5% CO_2_ and 37°C in a trigas incubator (Panasonic, MCO-170ML) for 7 days. Subsequent passages were achieved via a 1:1000 (6 μl into 6 ml) inoculum into freshly prepared ACCM-D media and incubation for 7 days. Axenically-grown Cb cultures were routinely (every five passages) subjected to contamination check by; inoculation into LB broth medium incubated under microaerophilic (5% O2, 5% CO_2_) conditions, LB broth medium incubated aerobically at 37°C, as well as ACCM-D medium incubated aerobically at 37°C.

### Measuring Growth and Host Cell Infectivity

To determine the infectivity of axenic- or intracellularly grown Cb; HeLa cells (CCL-2; American Type Culture Collection) were seeded onto 96 well culture plates at a density of 10^4^ in Roswell Park Memorial Institute (RPMI) medium containing 2% fetal bovine serum (FBS) for 16 hours. Cb cultures grown in ACCM-D were pelleted at 12000 x g at 4°C for 15 minutes. Serially passaged Cb were diluted in RPMI to normalize the number of genomes per volume, and 50 μl from various dilutions were inoculated onto the HeLa cell containing wells and centrifuged at 600 x g for 15 minutes at room temperature (Luedtke, Mahapatra, Lutter, *et al*. 2017). Immediately following centrifugation, the inoculating media was replaced with 200 μl of fresh RPMI containing 2% FBS. The plates were incubated at 37°C and 5% CO_2_ for 72 hours, fixed with ice-cold methanol for 10 minutes, then examined using indirect fluorescent antibody microscopy analysis as described previously (Luedtke, Mahapatra, Lutter, *et al*. 2017). Briefly, *C. burnetii* was stained using rabbit whole anti-*C. burnetii* NMII antibody diluted 1:1000 in PBS containing 3% bovine serum albumin (BSA) as a blocking agent. Primary antibodies were detected using Alexa Fluor 488 labeled goat anti-rabbit IgG antibodies diluted 1:1000 in PBS containing 3% BSA (Invitrogen). Total DNA was stained using 4’,6-diamidino-2- phenylindole (DAPI) diluted 1:10000 in PBS containing 3% BSA (Molecular Probes) to illuminate host cell nuclei. The methanol fixed and stained cultures were visualized on a Nikon Eclipse TE2000-S and the number of maturing CCVs were counted and calculations performed to ascertain the number of fluorescence forming units, which indicates the infectivity of the *C. burnetii* NMII in the ACCM-D samples.

### Transcriptomics

#### RNA extraction

Cells from axenic media growth passages 1, 3, 5, 10, 12, 16, 21, 31, 42, 51, 61 and 67 were harvested for transcriptomic analysis. RNA was extracted using a combination of hot Trizol treatment (Moormeier, Sandoz, Beare, *et al*. 2019) and the RNeasy Mini kit (Qiagen, Germany). Briefly, bacteria in 12 ml of ACCM-D culture (OD_600_ ∼ 0.3-0.4) were pelleted, resuspended in 700 μl of Trizol (ThermoFisher Scientific), boiled at 90°C for 10 min, and vortexed vigorously. 200 μl Chloroform was then added, followed by centrifugation at 12,000 x g at 4°C for 10 min. After separation, 300 μl of 100% ethanol was added to the aqueous phase, which was then quickly transferred to the spin column provided in the RNeasy mini kit. On-column DNA digestion was conducted by adding 80 μl (10 μl 1 Unit/μL RNase free DNase I, (ThermoFisher Scientific) in 70 μl reaction buffer from the Master Pure Yeast RNA Purification kit, Epicenter) of DNase preparation. The RNeasy mini kit’s protocol was followed for washing and eluting RNA. RNA quality was assessed visually on a gel as well as using RNA screen tape (Agilent) and RNA integrity number (RIN) value measurements using Tapestation and Bioanalyzer systems (Agilent).

#### Transcriptome sequencing and assembly

RNA sequencing (RNA-Seq) was conducted on the Illumina platform, using Nextseq 500 sequencer at Oklahoma State University Genomics and Proteomics core facility. Trimmomatic v0.38 (Bolger, Lohse and Usadel 2014) was used to process raw reads and remove Illumina adapter sequences. HISAT2 v2.1.0 (Kim, Paggi, Park, *et al*. 2019) was used to map the trimmed reads to the Chromosome (GenBank accession number: CP020616.1) and Plasmid (GenBank accession number: CP020617.1) of Cb NMII RSA 439. StringTie v2.1.4 (Kovaka, Zimin, Pertea, *et al*. 2019) was used to assemble reads alignments into potential transcript and to generate a non-redundant set of transcripts. The Python script prepDE.py supplemented with StringTie tool was used to convert transcripts per kilobase million (TPM) and fragments per kilobase million (FKPM) to gene level raw count matrix. The raw count table was imported to DESeq2 package (Love, Huber and Anders 2014) from Bioconductor in R programming language for further analysis.

#### Identification and analysis of Differentially Expressed Genes (DEGs)

The overall strategy for comparative transcriptomics analysis is outlined in Figure 1. DESeq2 was used to compute the fold change expression levels (reflected by logarithmic two-fold expression change i.e., L2fc) and its statistical significance (adjusted p-value, padj henceforth referred to as p-value) for every gene between passages when compared to passage one. DEseq2 tests the differential expression using negative binomial distribution and internally normalizes the counts by library size (Anders and Huber 2010). Genes with a p-value < 0.05 were labeled as significantly expressed. Only genes with TPM values > 10 in at least one passage were considered to minimize noise from minimally expressed genes. In most cases, differentially expressed genes in our temporal analysis were significantly expressed in more than one sampling point. In the few cases where differential expression was observed as a single spike in only one time point, a threshold of L2fc > 2 was considered as differentially expressed. Patterns of differential expression in DEGs was analyzed and visualized by constructing plot count graphs using the function “plotCounts” in DESeq2 package. The package “EnhancedVolcano” was used to visualize gene expression patterns as volcano plots. Differentially expressed patterns are classified into 1- Early up/downregulated, i.e., differential expression occurred in early (before passage 31) and the levels were sustained in subsequent late passages (see Figure 1). 2- Continuously up/downregulated, i.e., a constant/gradual increase in the magnitude of L2fc was observed throughout the sampling process (see Figure 1). 3- Late up/downregulated, i.e., differential expression was observed at or after passage 31. 4- Variable, i.e., expression levels were significantly higher than passage one in some timepoints and significantly lower than passage one in other time points (see Figure 1).

**Figure 1.**
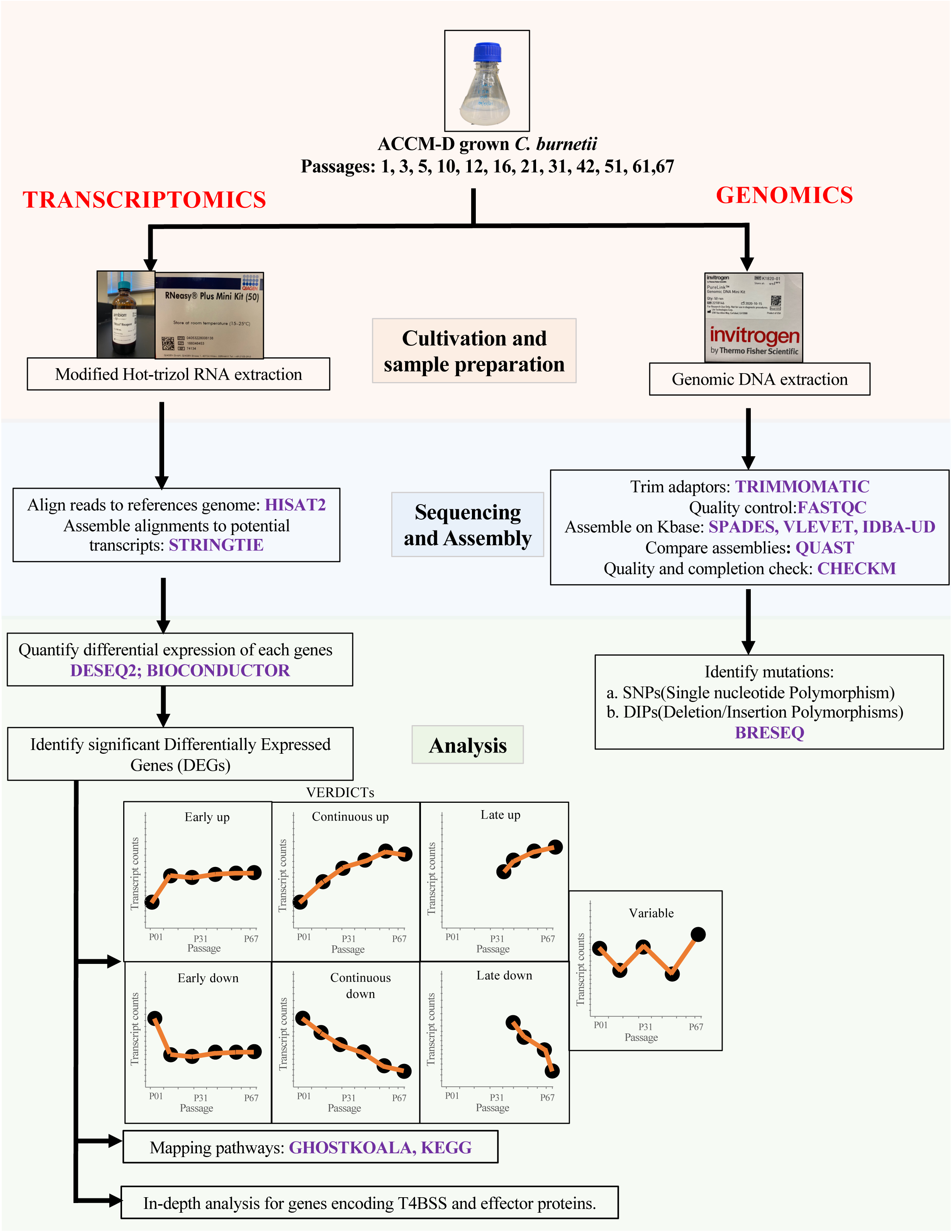
Flowchart representing the overall comparative transcriptomics and genomics strategy employed in this study.

#### Metabolic analysis and pathway mapping of DEGs

The subcellular protein localizations of proteins encoded by DEGs were predicted by using PSORTb (Yu, Wagner, Laird, *et al*. 2010). Transporter Classification Database (TCBD) was queried to find the putative transporter proteins. Pfam database (Mistry, Chuguransky, Williams, *et al*. 2020) was used to identify putative protein families for hypothetical proteins. BlastKOALA (Kanehisa, Sato and Morishima 2016) was used for functional annotation and assign KEGG Orthology (KO) numbers for the selected differentially expressed genes; and KEGG mapper (Kanehisa and Sato 2020) was then used to reconstruct metabolic pathway to visualize the differentially expressed genes in each pathway. The gene involvement in specific metabolic pathways were inferred from KEGG brite hierarchy file. Cluster of Orthologous genes (COGs) database (updated 2020) (Galperin, Wolf, Makarova, *et al*. 2021) downloaded from NCBI, was used to classify the effector proteins into functional categories.

### Genomics

#### DNA extraction and sequencing

8 ml of Cb cultures grown in ACCM-D for 7 days were pelleted by centrifugation at 12,000 x g and 4°C for 15 minutes. DNA extraction was conducted using Pure Link® Genomic DNA Kits (ThermoFisher Scientific) following the manufacturer’s instructions. Sequencing was conducted at Oklahoma State University Genomics and Proteomics core facility using Illumina’s NextSeq® 500 System. DNA quality was assessed visually on a gel as well as using DNA Screentape and Bioanalyzer systems (Agilent).

#### Genome assembly and quality control

The KBase platform (Allen, Drake, Harris, *et al*. 2017), which implements and integrates multiple bioinformatic tools, was used for DNA sequence data handling. Trimmomatic v 0.36 (Bolger, Lohse and Usadel 2014) was used to trim the Illumina adapter sequences. Quality check was done using FastQC v0.11.5 (Andrews 2010). Assembly of Illumina reads to contigs was attempted using four different assemblers (Spades v3.13.0, Velvet v1.2.10 and IDBA-UD v1.1.3 and Unicycler (Davis, Wattam, Aziz, *et al*. 2020)). The quality of genome assemblies from these four assemblers were assessed using QUAST v1.4 (Gurevich, Saveliev, Vyahhi, *et al*. 2013) and the best assemblies were selected using metrices such as total length, largest N50, lesser number of contigs and less Ns. CheckM (Parks, Imelfort, Skennerton, *et al*. 2015) was used to assess quality and completion of genomes (Figure 1).

#### Analysis of mutation frequencies

Breseq (Deatherage and Barrick 2014) was used to identify mutations/changes in the genome assemblies obtained, with Passage one used as a reference. The occurrence and frequency of both single nucleotide polymorphisms (SNPs) and deletion-insertion polymorphisms (DIPs) were examined (as outlined in Figure 1). Breseq was run in polymorphism mode, which identifies the mutations occurring in a fraction of a population in addition to consensus mutations in the entire population in a sample. This allows for the visualization of the propagation of a particular mutation as a frequency of evolved alleles and genetic diversity in the population.

#### Nucleotide sequences accession number

The whole-transcriptome and genome shotgun sequences were deposited in GenBank under the BioProject PRJNA796300 and BioSample accession numbers SAMN24840407-SAMN24840437 and SAMN24847762-SAMN24847773. The 31 transcriptomic assemblies were deposited in the SRA under project accession number SRX13723330-SRX13723360. Reads for 12 genomic assemblies can be found under SRA with accession SRX13726189- SRX13726200.

## Results

### Coxiella burnetii infectivity but not viability decreases with continuous passaging in axenic media

Following anecdotal observations, we sought to quantitatively assess whether serially passaged Cb infect cultured cells less readily than cell derived bacterial stocks. Using *C. burnetii* NMII serially passaged 1, 3, 5, and 10 times in ACCM-D, we initiated infections of Hela cells with bacterial dilutions normalized by the number of genomes in each sample. When the number of fluorescence forming units (FFU) per sample were calculated, they revealed a decrease in the number of *C. burnetii* filled vacuoles in tissue culture cells as the bacteria from subsequent passages were analyzed, respectively, resulting in a nearly two-log decrease between Passages 1 and 10 (Figure 2A). This indicated that there were fewer bacteria per genome that were capable of initiating a typical infection following multiple passages in axenic media. Next, we sought to determine if the decrease in infectivity of tissue culture cells was associated with a decrease in *in vitro* viability of the *C. burnetii* as measured by colony forming units on ACCM-D agar. To address this question, we plated dilutions of passages 1, 3, 5, and 10 on ACCM-D agar plates and performed colony counts. Contrary to the decrease in infectious units (Figure 2A), the colony counts indicated that there was no significant change in viable bacteria relative to genomes as the organism was serially passaged (Figure 2B). This indicated that the number of live and replicative bacteria did not change during axenic growth, and therefore bacterial death was not responsible for the decrease in Cb infectivity of the cultured eukaryotic cells observed.

**Figure 2.**
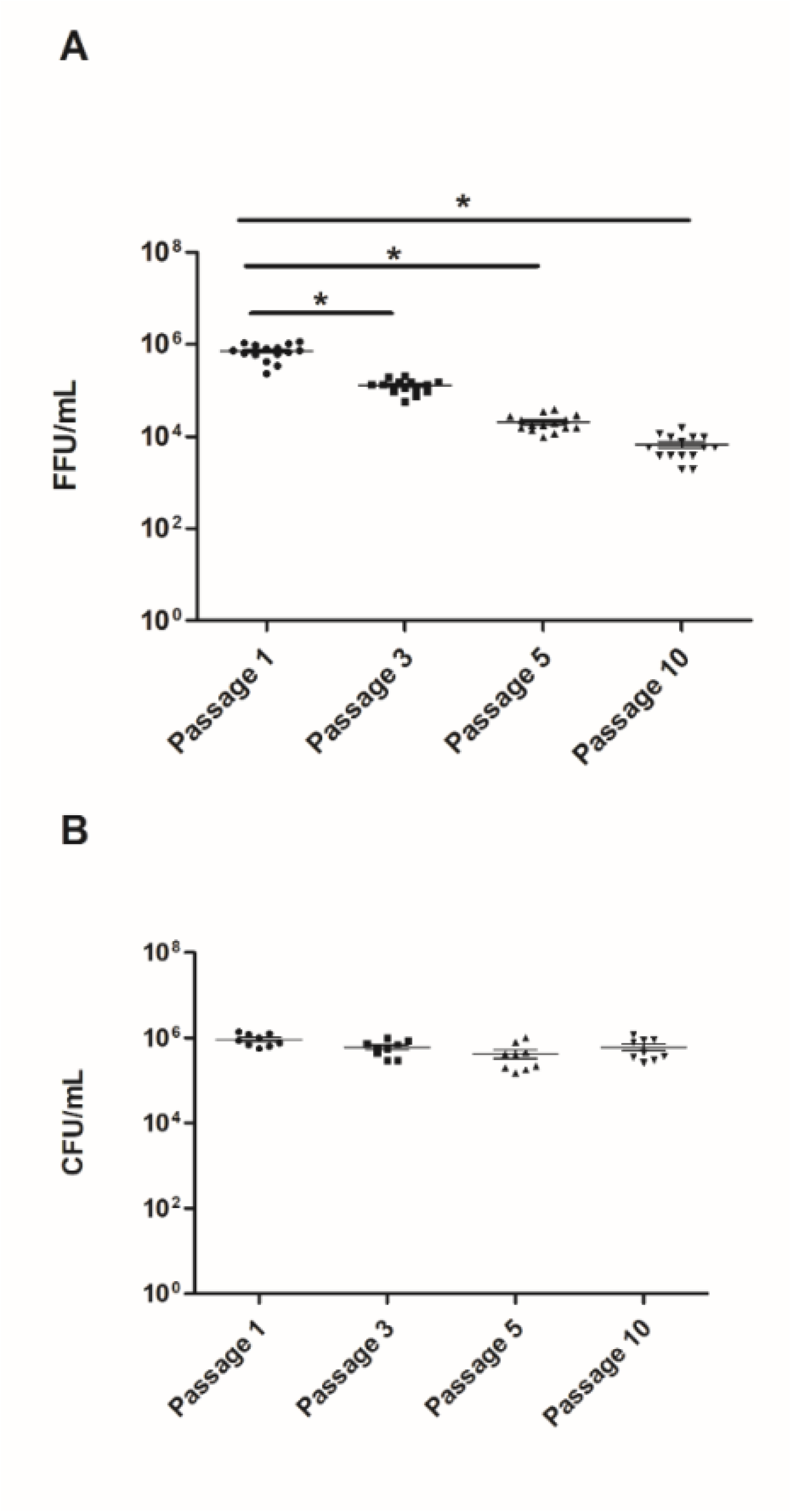
Intracellular vs Axenic Growth Following Serial Passage of *C. burnetii* NMII. Intracellular and axenic growth from 3 biological replicates of passaged ACCM-D growth **A)** fluorescence forming units (FFU) counts of infections of HeLa cells normalized by Cb genomes from passages 1, 3, 5, and 10. Significance between different passages are indicated by lines and *p<0.001. **B)** CFU enumeration of passages 1, 3, 5, and 10 Cb spread on ACCM-D plates normalized to genomes. No statistically significant difference was observed between groups.

### Transcriptional activity

RNA sequencing was conducted on 12 different passages (1, 3, 5, 10, 12, 16, 21, 31, 42, 51, 61, and 67). A total of 162.2 Gb data were obtained, with 6.05 – 22.57 million reads per sample (Average = 10.14 million reads). Transcripts representing each of the 2,217 genes in *C. burnetii* NMII strain (genome and plasmid) were identified in all samples, attesting to the depth of the sequencing effort conducted.

Expression level and overall pattern (Early up, Continuous up, Late up, Early down, Continuous down, Late down, Variable) for every gene in the Cb genome is shown in Table S1. A total of 845 genes were differentially expressed in at least one passage, with 464 upregulated and 371 downregulated (Figure 3a) genes. The number of differentially expressed genes (DEGs) per passage ranged between 25 and 807 (Figure 3b). A general pattern of an increasing number of differentially expressed genes per passage was observed through passage 51, after which the number of DEGs dropped in passage 61 and 67 (Figure 3B). The ratio of upregulated to downregulated genes in each passage ranged between 0.14 (in passage 5) and 1.26 (in passage 3). Of the 371 downregulated genes, 249 expressed early down pattern, 48 were continuous down, 43 were late down, and 31 were down in only one passage. Of the 464 upregulated genes, 288 were early up, 38 were continuous up, 85 were late up and 53 were up in only one passage (Figure 3C). Of the 845 DEGs, 81 were differentially regulated in 8-11 of the passages, 144 in 5- 7 of the passages, 526 in 2-4 of the passages and 84 in only one passage (Figure 3D).

**Figure 3.**
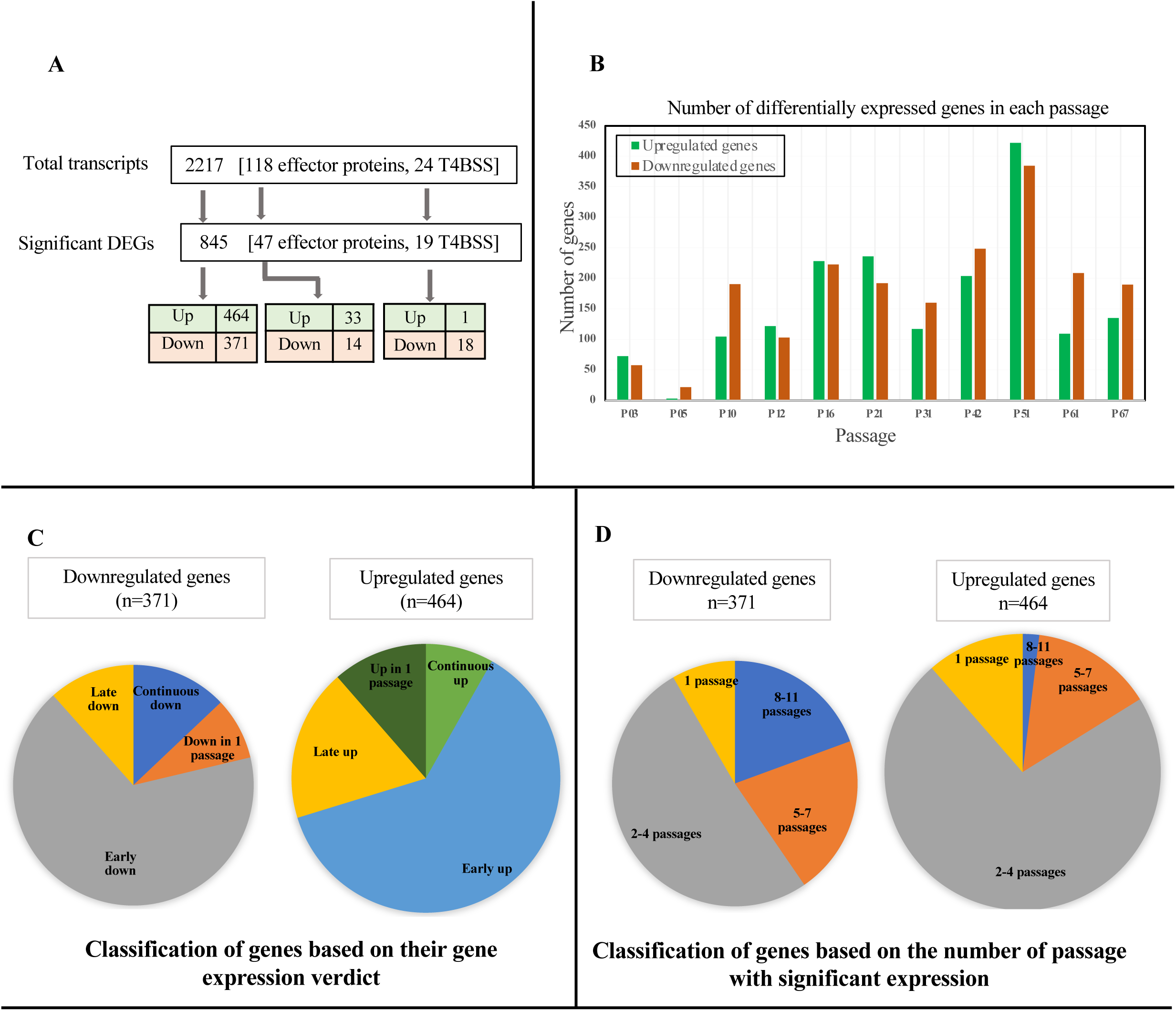
Overview for Differential Expression patterns in axenically-grown Cb. **A**) Summary for the number of total and differentially expressed genes, T4BSS and effector proteins in this experiment. **B**) Bar graph showing the number of upregulated (p-value<0.05 and L2fc > 0) and downregulated (p-value<0.05 and L2fc < 0) genes in each passage. **C**) Classification of genes based on their gene expression verdicts. **D**) Classification of genes based on the number of passages showing significant expression change.

Visualization of DEGs patterns using volcano plots was used to provide an overview of the overall level of expression changes (see Figure 4). Transcript expression levels from each passage were compared to passage one. The labeled boxes within each plot analysis represents the 10 highest differentially expressed transcripts (i.e., smallest p-value). Visual inspections demonstrate that chaperons and T4BSS machinery proteins consistently represent an important component of highly downregulated genes in all passages. Below, we provide a more detailed assessment on differential expression patterns for various genes and pathways.

**Figure 4.**
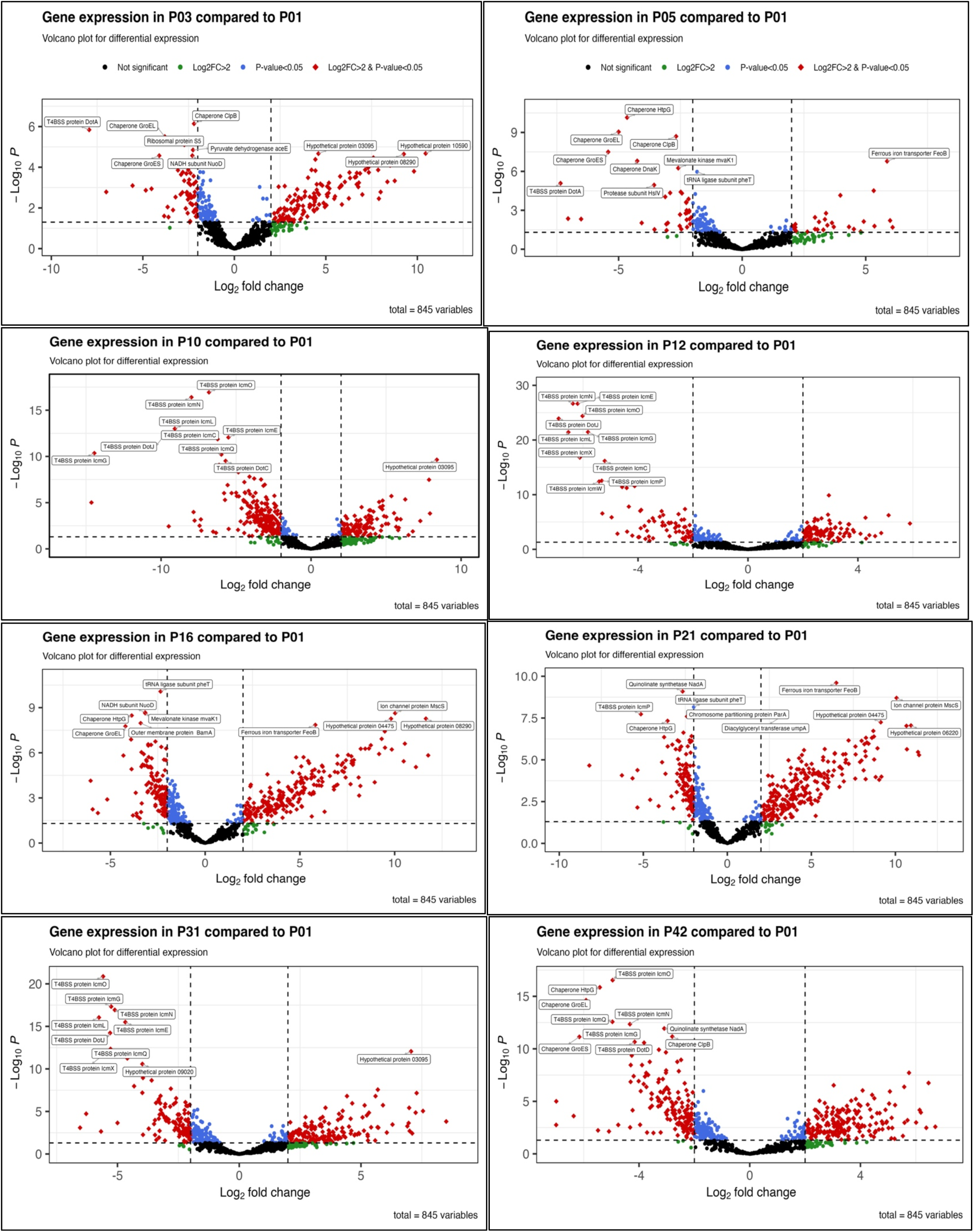
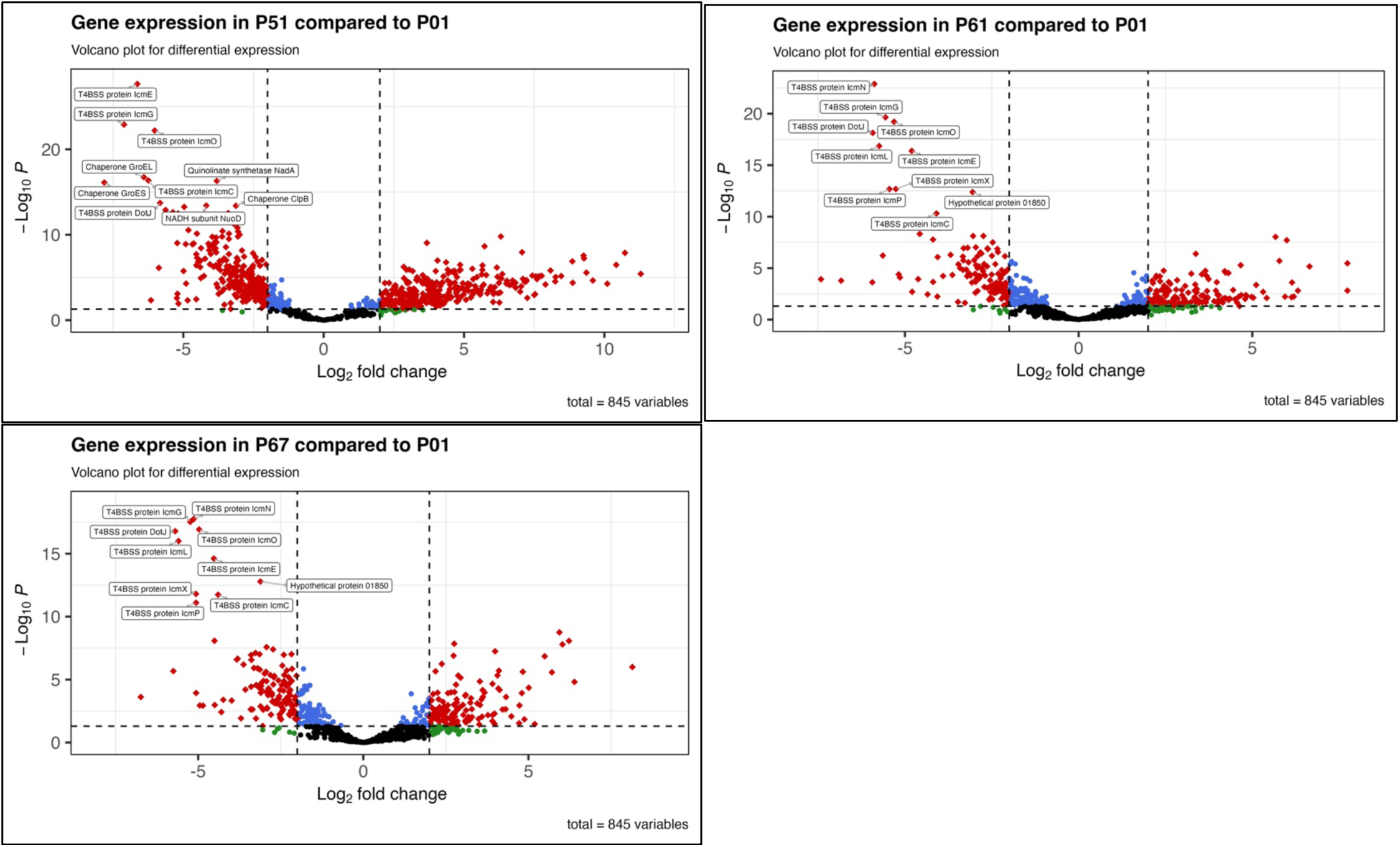
Volcano plots for 845 significant DEGs in different passages when compared to passage 01. Black dots represent non-significant DEGs, green dots represent non-DEGs with L2fc >2, blue dots represent significant DEGs with L2fc <2 whereas the red diamonds represent significant DEGs with L2fc 〾2. The 10 genes with the lowest p-value in every passage comparison are labeled in boxes.

### Secretory pathways are significantly downregulated in axenic growth media

The defective in organelle trafficking/intracellular multiplication (Dot/Icm) Type IVB secretion system (T4BSS) in Cb has been shown to secrete the effectors and other pathogenic determinants into the host cell, a process required for Cb intracellular growth and pathogenesis (Beare, Gilk, Larson, *et al*. 2011; Carey, Newton, Luhrmann, *et al*. 2011; Rudolf Toman 2012; van Schaik, Chen, Mertens, *et al*. 2013; Voth and Heinzen 2009). Interestingly, 19 out of the 24 components of the Cb T4BSS demonstrated significant differential expression (Figure 5A, Figure S1a, Table S1). Out of these, 18 genes were downregulated and only one gene was upregulated (Figure 5A, Figure S1a). Indeed, T4BSS encoded gene transcripts were some of the most significantly downregulated across the passages (see Figure 4).

**Figure 5.**
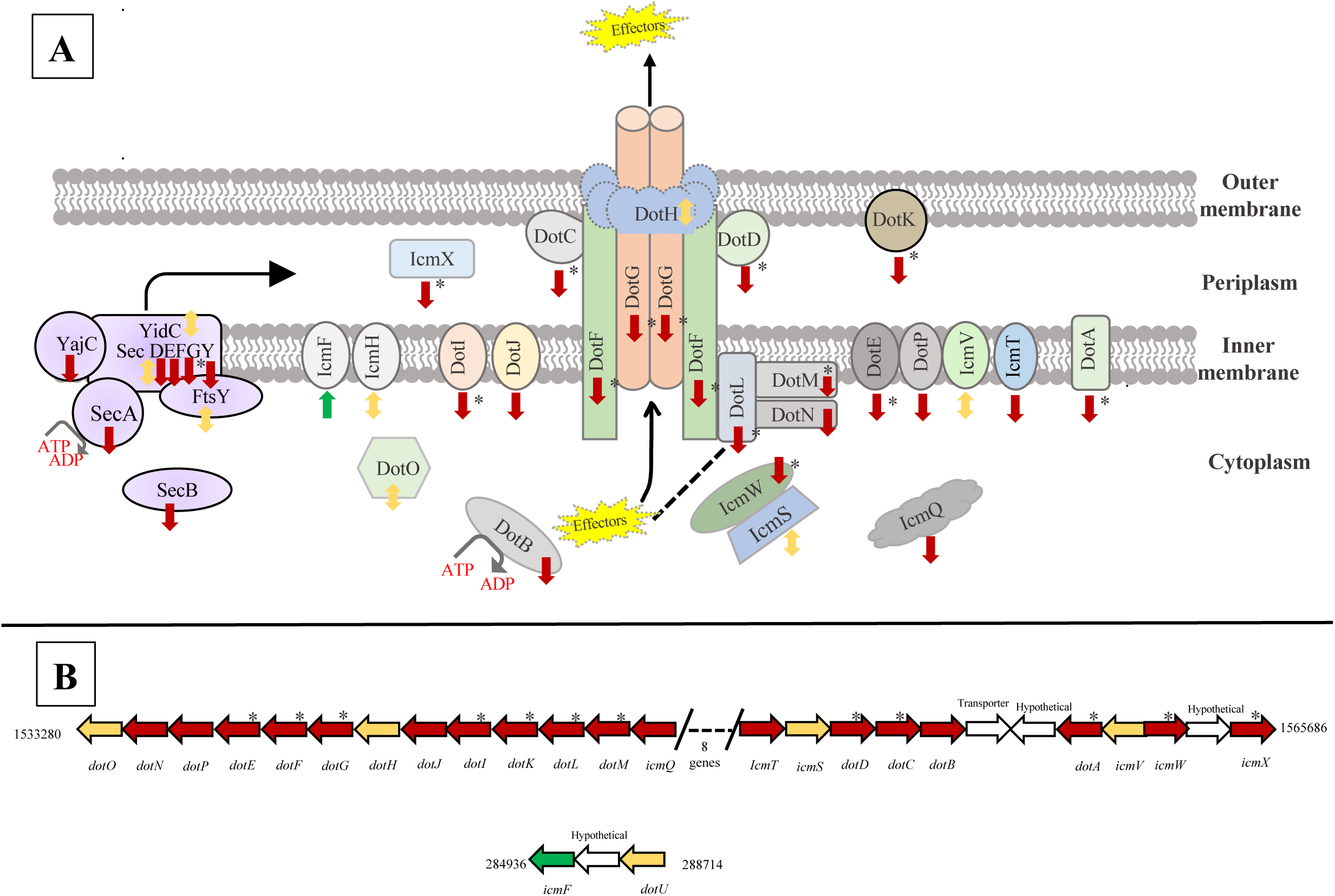
Cb T4BSS machinery and Sec expression changes during axenic passaging. **A**) Membrane complex model with gene transcript expression profiles for components of the Cb T4BSS (right side) and Sec protein export pathway (left side). DEG patterns are denoted by arrows (downregulated in maroon, no significant expression changes in yellow, and upregulated in green). * Represents genes that were highly downregulated (L2fc ≤ -3). **B**) Gene locus map for all the components in Cb NMII strain [CP020616.1]. The arrows are colored filled according to DEGs patterns as (downregulated in maroon, no significant expression change in yellow, and upregulated in green). The arrows not filled are genes not related to the T4BSS pathway.

Within the T4BSS core transport complex, transcripts for genes *dotC*, *dotD*, *dotF*, and *dotG* were early downregulated, while only *dotH* indicated no significant change in gene expression in all passages. In addition, expression changes in genes encoding components of the T4BSS coupling protein complex (*dotL*, *dotM* and *icmW*) demonstrated early down or continuous down (*dotN*), whereas *icmS* was the only component with no significant gene expression changes. Transcripts of the gene *dotB* was continuous downregulated whereas *dotA* and *icmX* were found to be early downregulated (Figure 4 and Figure S1a). Besides the two main complexes, other components of the Cb T4BSS that were transcriptionally downregulated during continuous axenic media passaging includes genes *dotE, dotP, dotK, dotI, dotJ, icmT* and *icmQ* (Figure 5a). *icmF*, located in a separate locus than the majority of the T4BSS genes (Figure 5b), was the only component that was transcriptionally upregulated in the system (Figure S1a).

Transcript expression of genes within additional secretory pathways in Cb were also analyzed. Genes of the general secretary (Sec) pathway revealed a general trend of downregulation (Figure S1b, Table S1). The Sec pathway provides a channel for polypeptide movement across the bacterial inner membrane (Green and Mecsas 2016). It is comprised of the proteins SecY, SecE and SecG and an ATPase (SecA) that drives protein movement (Green and Mecsas 2016; Tsirigotaki, De Geyter, Šoštarić, *et al*. 2017). This pathway is known to secrete proteins from the cytosol through the cytoplasmic membrane (Mori and Ito 2001). We identified all of the Cb Sec pathway components, as shown in Figure 5a. Transcripts for expression of the inner membrane proteins SecA, SecF and SecE were early down and SecY and YajC were continuously down, whereas the targeting proteins SecB and SecG were down at later passages.

### Expression patterns of T4BSS effector proteins previously implicated in Cb pathogenesis and intracellular survival

The differential expression in all 118 genes encoding T4BSS effector proteins previously identified in Cb through a variety of effector screens (Carey, Newton, Luhrmann, *et al*. 2011; Chen, Banga, Mertens, *et al*. 2010; Larson, Martinez, Beare, *et al*. 2016; Voth, Beare, Howe, *et al*. 2011; Weber, Chen, Rowin, *et al*. 2013) was examined. Forty-seven effector proteins were differentially expressed (column “Effector proteins” in Table S1, Figure 3a). Interestingly, more genes were upregulated (n=33) than downregulated (n= 14) (Figure 3a).

The 14 genes encoding effector proteins that were transcriptionally downregulated indicated an up to four-fold expression change, with the expression changes primarily beginning from passage 10 (Figure S2). These genes fell into COG functional groups of signal transduction mechanisms (*ankG*, *ankK* and *ankD*), carbohydrate transport and metabolism (B7L74_09020), posttranslational modification, protein turnover and chaperones (*cpeH*), replication, recombination, and repair (*cig57*), lipid transport and metabolism (B7L74_03275) and mobilome: prophages and transposons (B7L74_08400) (Table S2, Figure 6a, Figure 6b). Genes B7L74_08200 and *cpeF* were predicted as general function categories whereas there were no functional homologies for B7L74_07850, *cig2*, *cirC* and B7L74_03065 in the COG database (Table S2).

**Figure 6.**
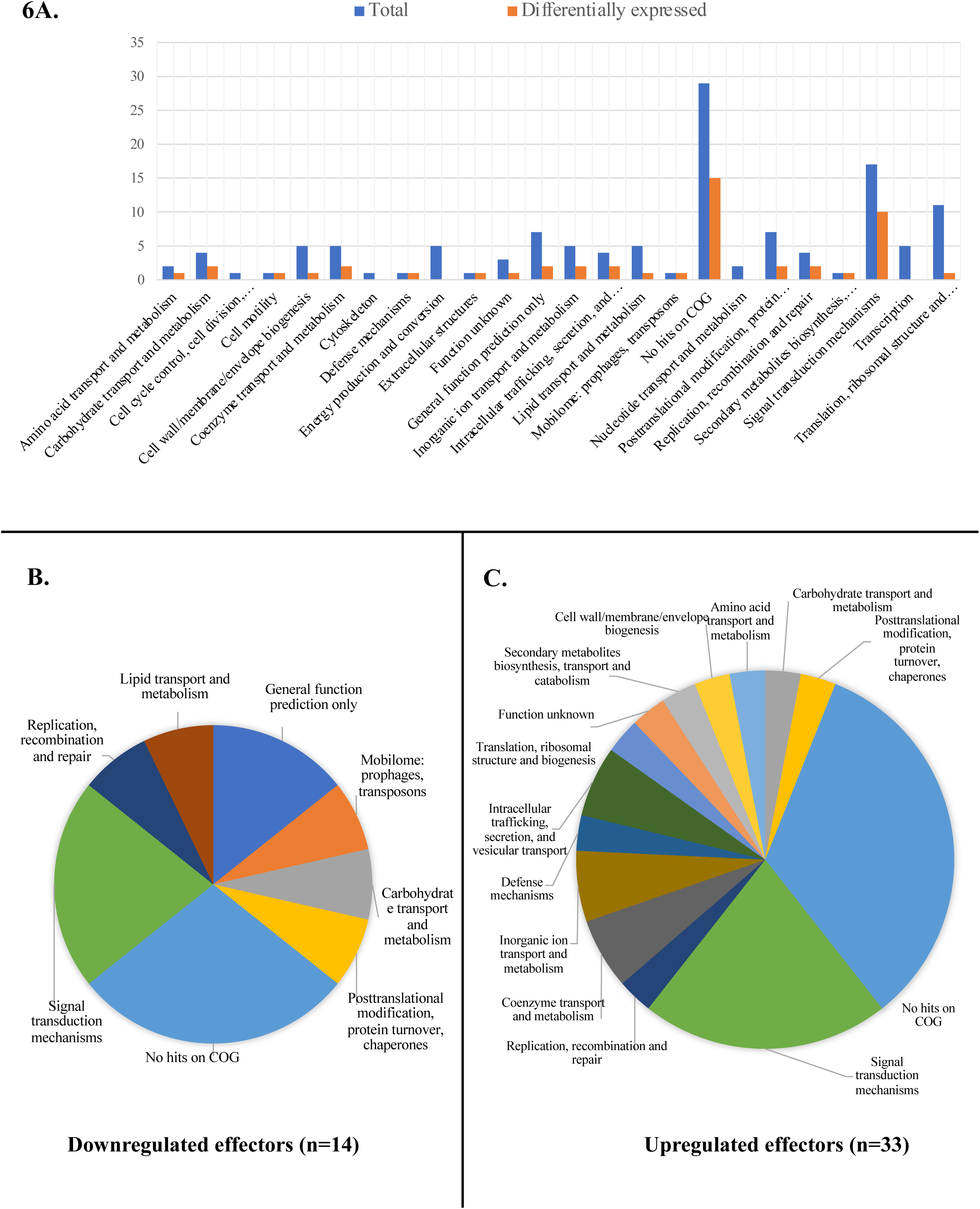
Transcript expression changes for T4BSS effector proteins during continuous axenic passaging. **A)** COG classification graph for 118 identified effector proteins. The blue bars represent total effector proteins in each COG functional group (x-axis) and orange bars represents the effectors that showed differential gene expression. **B**) Pie chart showing COG classification for 14 downregulated effector proteins. **C**) Pie chart showing COG classification for 33 upregulated effector proteins.

### Expression patterns of additional pathogenic determinants in Cb

Expression patterns of additional pathogenic determinants unrelated to effector proteins were examined. Of these, we noted three interesting patterns. First, the general downregulation of a wide range of chaperone proteins. Chaperone proteins primarily function as protein folding catalyst, but many are considered virulence factors for many intracellular pathogens given that they encounter stress related to phagosome acidification and phagosome fusion with lysosomes (Neckers and Tatu 2008). Amongst the 16 genes annotated as chaperons in the Cb genome, 10 were transcriptionally downregulated during continuous *in vitro* passaging (Figure 7a) (Table S1). Notable downregulated chaperones include glutaredoxins (*grxC* and *grxD)* that have been shown to be involved in CCV detoxification (Beare, Unsworth, Andoh, *et al*. 2009) and genes encoding heat shock proteins classes such as *dnaK, hptG, groEL* and *dnaJ*. These proteins are known to help bacteria adapt to stressful conditions (Arnold, Jackson, Waterfield, *et al*. 2007; Genevaux, Georgopoulos and Kelley 2007) (Figure S3). DnaK has been shown to be critical for survival of pathogenic bacteria inside the macrophage (Takaya, Tomoyasu, Matsui, *et al*. 2004) and is induced in Cb in high acid condition, the condition similar to the phagolysosome (Macellaro, Tujulin, Hjalmarsson, *et al*. 1998).

**Figure 7.**
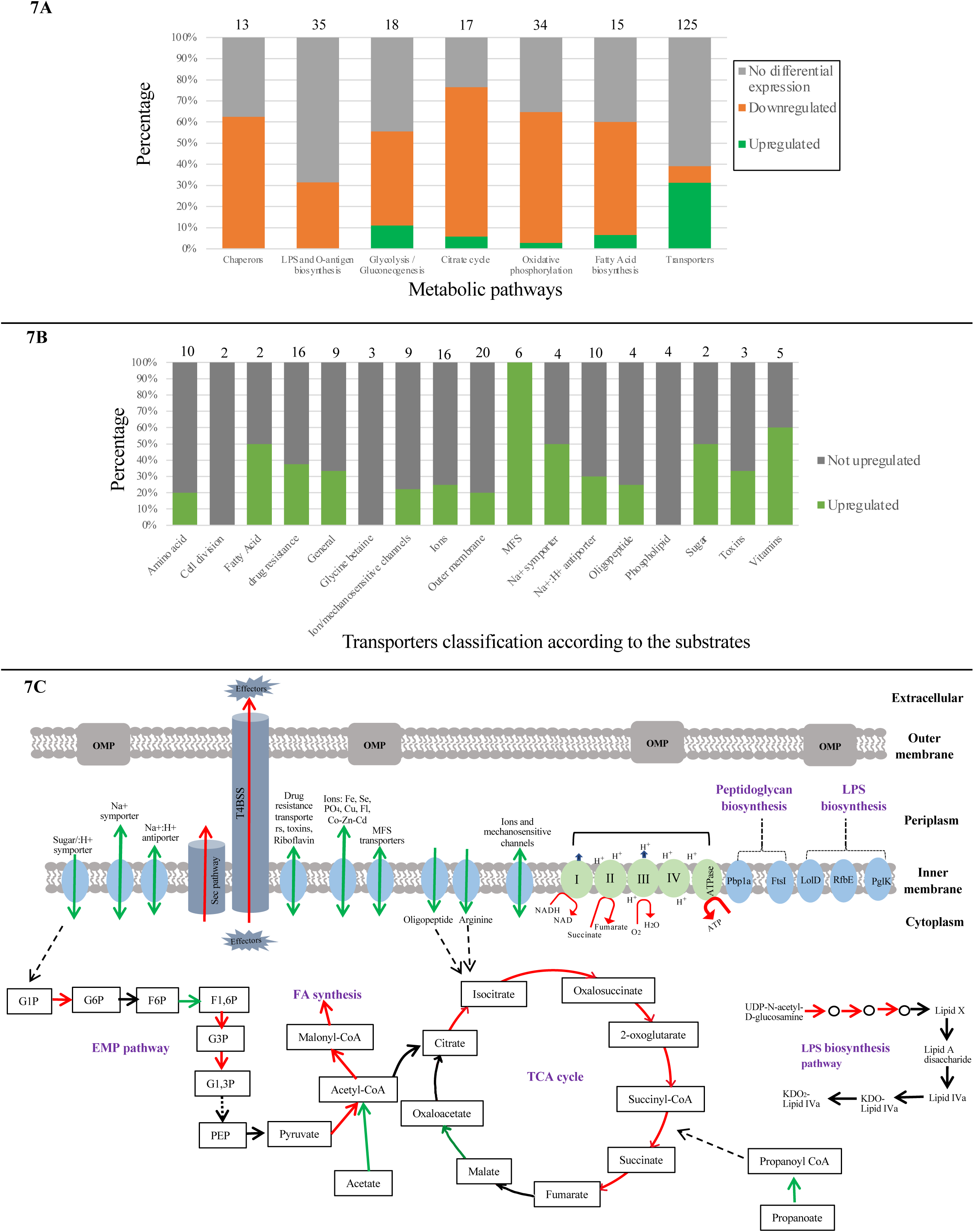
Gene expression changes in central metabolic pathways. **A**) Bar graph showing numbers of DEGs in each central metabolic pathway. The numbers on top of the bars represent the total number of genes classified into that metabolic category by KEGG. **B**) Broad classification of differentially expressed transporters according to their substrates. The number in parenthesis for each category represents the proportion of upregulated genes **C**) Graphical representation of all metabolic pathways discussed in the manuscript text.

The second observation is the downregulation of several genes involved in lipopolysaccharide biosynthesis. The Lipopolysaccharide (LPS) layer has long been known as a pathogenic determinant and important for the host interaction in *C. burnetii* (Gajdosova, Kovacova, Toman, *et al*. 1994; Hussein, Kovacova and Toman 2001; Williams and Waag 1991). Out of 35 genes related to LPS synthesis and O-antigen nucleotide sugar biosynthesis, 11 were downregulated (Figure 7a) (Column “LPS and O-antigen biosynthesis” in Table S1). Out of 11 downregulated genes, 4 genes are involved in KDO2-lipid IVA Wbp pathway for LPS biosynthesis whereas 8 genes are involved in O-antigen nucleotide sugar biosynthesis (Table S1). Genes involved in the first 3 steps i.e., UDUDP-N-acetylglucosamine acyltransferase (*lpxA),* UDP-3-O-[3-hydroxymyristoyl] N-acetylglucosamine deacetylase *(lpxC)* and P-3- O-[3-hydroxymyristoyl] glucosamine N-acyltransferase *(lpxD)* and the gene D-glycero-D-manno-heptose 1,7-bisphosphate phosphatase (*gmhB)* in LPS biosynthesis pathway are early down or continuously down (Figure S3) (Figure 7C). In addition, 3 transporters related to LPS synthesis i.e., a lipoprotein releasing system ATP-binding protein (*lolD)*, a lipid flippase important in cell membrane formation (*pglK)* and a probable O-antigen/lipopolysaccharide transport ATP-binding protein (*rfbE*) were also early down (Figure 7C). 8 downregulated genes including *wbpW, gmhB, galE, wbpD, galE, wbpI, cap1J* and *glmU* are involved in O-antigen nucleotide sugar biosynthesis pathway, a 14 genes pathway which is the first step in O-antigen biosynthesis where nucleotide sugars are assembled and activated by adding NTP (Samuel and Reeves 2003).

Finally, 3 of the 15 genes involved in peptidoglycan layer biosynthesis were downregulated (Table S1). These genes are penicillin-binding protein PBP3/*ftsI*, penicillin binding protein PBP1A/*mrcA* and undecaprenyl diphosphate synthase *uppS*. The peptidoglycan layer in Cb is an immunogenicity determinant and thickens substantially during LCV to SCV transition to help in environment resistance (Amano, Williams, McCaul, *et al*. 1984; Sandoz, Popham, Beare, *et al*. 2016). *ftsI* and *mrcA* are the only two genes encoding penicillin binding proteins in this genome that are involved in peptide cross linking. This suggests that the peptidoglycan layer may be even thinner than is in common intracellularly and could indicate a reduced requirement during axenic growth since the thick SCV peptidoglycan layer in Cb has been correlated to bacteria being more infectious (Sandoz, Popham, Beare, *et al*. 2016).

### Downregulation of multiple hypothetical proteins could suggest novel pathogenicity determinants

We posited that downregulated hypothetical proteins could represent previously unrecognized pathogenicity determinants, and that such identification could be a useful starting point for subsequent experimental validation. We identified 30 Cb genes encoding hypothetical proteins that were downregulated in more than 4 separate passages. Subcellular localization analysis software predicts that 13 are cytoplasmic, 8 inner membrane, 1 extracellular, 1 periplasmic and 7 with unknown localization (Table S4). Of the cytoplasmic proteins, Cig28 and an AMP binding protein (CBU_0787) possess a regulatory element recognized by PmrA, a sequence related to T4BSS expression and translocation and thus a potential predictor of effector proteins (Beare, Sandoz, Larson, *et al*. 2014), although they were subsequently shown not to be translocated by the Cb T4BSS (Beare, Sandoz, Larson, *et al*. 2014; Zusman, Aloni, Halperin, *et al*. 2007). Similarly, an uncharacterized protein, CBU_1234, has been shown to have a glutamate-rich C-terminal secretion signal (E-block), which is also a predictor of effector proteins (Weber 2014). Two Glycosyltransferase family 1 proteins (i.e. CBU_0839 and CBU_0841) have previously been linked to LPS mutations that lead to phase transitions (Beare, Jeffrey, Long, *et al*. 2018). The eight proteins localized in the inner membrane included Cig3, an immunoreactive peptidase CBU_0215 (Weber 2014) previously shown to contain a regulatory element recognized by PmrA but not translocated by Dot/Icm system (Beare, Sandoz, Larson, *et al*. 2014; Zusman, Aloni, Halperin, *et al*. 2007), an immunoreactive protein CBU_1865 (Beare, Chen, Bouman, *et al*. 2008) and a DUF3971 domain-containing protein CBU_1468 that has been shown to be important for intracellular replication (Newton, Kohler, McDonough, *et al*. 2014) (Table S4).

The one differentially expressed hypothetical protein identified by the localization software as an extracellular protein was CBU_0962; a predicted short chain dehydrogenase with a yet unknown specific function (Bewley 2015). Lastly, proteins with unknown localization included an exported protein Cig40 (Weber 2014) with a regulatory element recognized by PmrA, a hypothetical surface antigen Com1(Chen, Banga, Mertens, *et al*. 2010) and a hypothetical protein CBUA0012 located in an ORF containing other plasmid effectors but has shown to be not secreted by Dot system (Voth, Beare, Howe, *et al*. 2011) (Table S4).

### Transcriptional patterns of central metabolic pathways

Analysis of gene expression patterns of central metabolic pathways demonstrated a general trend of down-regulation in genes encoding enzymes in central catabolic, amphibolic, and anabolic pathways, coupled with a broad upregulation in genes encoding transporters. An overall pattern of downregulation of glycolysis genes (8/18 genes) was observed (Figure 7A), with several enzymes such as pyruvate dehydrogenases (*pdhC, pdhD*), fructose-bisphosphate aldolase (*fbaA*), glyceraldehyde 3-phosphate dehydrogenase (*gapA*), and phosphoenolpyruvate carboxykinase (*pckA*) downregulated early in the passaging (Figure S3, Table S1). Gene *pmm-pgm,* which encodes the enzyme phosphomannomutase/ phosphoglucomutase and is involved in the first step of glycolysis, was down early as well (Figure S3). Similarly, analysis of genes encoding enzymes of the tricarboxylic acid (TCA) cycle, demonstrated an overall downregulation (12/17 genes), with a downregulation of ∼2 fold in isocitrate dehydrogenase (*IDH2*), the rate-limiting enzyme in the TCA cycle (Figure S3). As well, the downregulations in multiple carbohydrate dehydrogenases (e.g., pyruvate dehydrogenases *pdhC* and *pdhD*), succinate dehydrogenases (*sdhA, sdhB, sdhD*), as well as genes that are necessary for oxidation of glycolytic and TCA cycle sugar intermediates was observed. Finally, genes encoding components of the electron transport chain (ETC) were also downregulated. These include Complex I: NADH-quinone oxidoreductase (*nuoB, nuoD, nuoE, nuoF, nuoH, nuoI, nuoK, nuoL, nuoM, nuoN*), Complex II:Succinate dehydrogenase (*sdhA, sdhB, sdhD*), Complex III:Cytochrome oxidoreductase (*cyoC, cyoD, cydA, cydB, cydX*), and Complex V:F-type ATPase (*atpA, atpB, atpE, atpD, atpF, atpG*) (Figure 7A, Figure S3). The downregulations in two complexes (*cyoC* and *cyoD*) of cytochrome o oxidase, which is known to be induced in oxygen rich growth conditions in bacteria, (Cotter, Chepuri, Gennis, *et al*. 1990) suggests a decreased affinity and/or competition for oxygen in the cell-free growth environment, as previously suggested (Kuley, Bossers-deVries, Smith, *et al*. 2015). Cytochrome d oxidase, which is shown to be expressed more in oxidative and nitrosative stress conditions (Cotter, Chepuri, Gennis, *et al*. 1990), also has expression of two of its components (i.e., *cydA* and *cydX*) early down and late down, respectively. Finally, an overall downregulation of fatty acids biosynthesis genes transcription (8/15) was also observed (Figure 7A, Figure S3).

In contrast to the general trend of downregulation of the central metabolic machinery of Cb, a marked upregulation of genes encoding transporters was observed. Out of 125 general transporters, the transcription of 39 were upregulated and 10 were downregulated (Column “Transporters” in Table S1) (Figure 7a). Transporters that were upregulated have double fold expression change (L2fc) ranging from 3-10 (Figure S3). Upregulated primary transporters included transporters for the amino acid arginine, oligopeptides, fatty acids, and vitamins such as riboflavin and thiamin (Figure S1) (Figure 7b) as well as a small number of transporters (4 out of 20 present) related to synthesis and maintenance of outer membrane (Table S1). On the other hand, upregulated secondary transporters included MFS transporters, symporters, antiporters, and mechanosensitive ion channels. Of these, a notable observation was made where all six MFS transporters and two out of four Na+ symporters found in Cb genome were found to be early upregulated (Table S1). These MFS transporters transports various compounds such as monosaccharides, oligosaccharides, amino acids, peptides, vitamins, cofactors, drugs, nucleobases, nucleosides, and organic and inorganic anions and cations. Also, a large proportion of transporters related to drug resistance (6 /16 present in the genome) and ion transporters mediating the uptake of ions such as copper, iron, fluoride, selenite, cobalt-cadmium-zinc, and phosphate were also upregulated (Table S1) (Figure 7b). In addition to transporters mediating substrate transport, transporters involved in pH homeostasis such as ions/mechanosensitive channels and Na+:H+ antiporter were also upregulated. (Figure 7b) (Table S1). The Na+:H+ antiporter functions to utilizes the proton motive force to efflux intracellular sodium ions for intracellular pH homeostasis (Ito, Morino and Krulwich 2017) and these antiporters along with ion/mechanosensitive channels have been proposed to play an important role in pH homeostasis and survival within the acidic PL (Seshadri, Paulsen, Eisen, *et al*. 2003). Lastly, 3 out of 9 transporters classified under general or unknown functions were upregulated as well (Figure 7b).

### Genomics reveals a stable Cb genome

For all 12 passages analyzed, genomes with 100% completeness (assessed by identifying all 265 housekeeping marker genes specific for the Proteobacteria (Parks, Imelfort, Skennerton, *et al*. 2015)) were obtained. N50 of genomic assemblies ranged between 49,903 and 75,629, N90 ranged between of 15,966 and 20,406, and the number of contigs per genome ranged between 56 and 64 (Table S5). Using Passage one as a reference, we identified 842 unique single nucleotide polymorphisms (SNPs) and 118 unique deletions/insertion polymorphisms (DIPs) (Table S5) (Figure 8).

**Figure 8.**
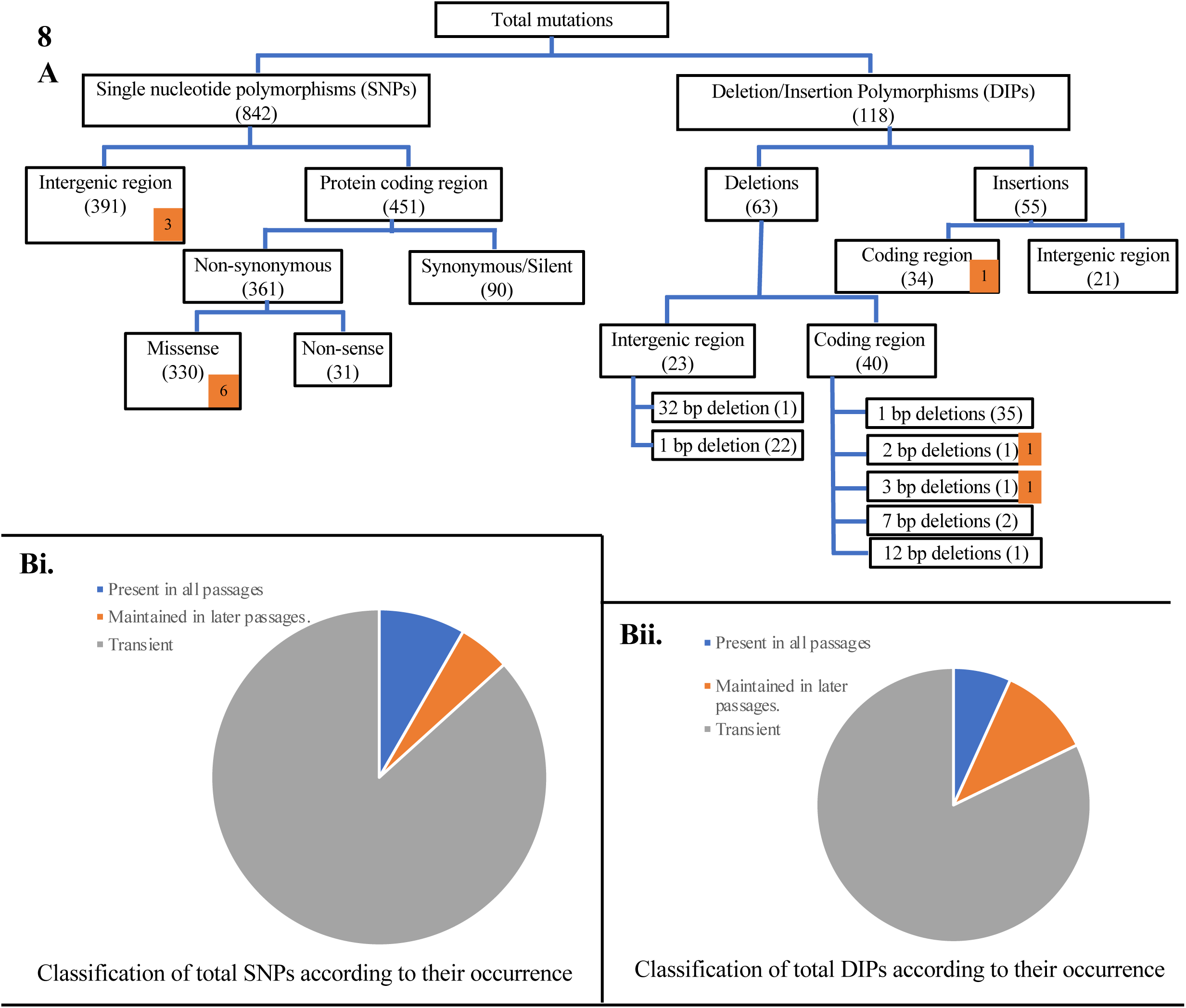
DEseq2 Analysis of Genomes from Passages. **A**) Flowchart for classification of different types of Single nucleotide polymorphisms (SNPs) and Deletion/Insertion polymorphisms (DIPs) found in this experiment. Numbers in the orange blocks at the tip of some boxes represents the number of those mutations that occurred in 100 % of the passages. **Bi**) Classification of SNPs according to its occurrence in number of passages. **Bii**) Classification of DIPs according to the occurrence in number of passages.

Of 842 unique SNPs, only 9 were identified in consensus mode (i.e., present in 100% of population in one or more passages) while the remaining 833 SNPs were identified in population mode, (i.e., occurring in a fraction of the community) when sequenced (Figure 8). Further, only 69 unique SNPs were identified to occur in all (i.e., passage 3-61), and only 43 SNPs were maintained in later passages (Figure 8b). More importantly, only one consensus mutation occurred in a gene that was downregulated in transcriptomic analysis. This gene GTP pyrophosphokinase SpoT (B7L74_01590) had a one amino acid (aa) substitution (T to A) at position 262, which propagates to 100% population in the last 9 passages analyzed and is also noticeably early down in gene expression. SpoT is a signal transduction component and transcriptional regulator with a role in helping *Coxiella* cope with the low-nutrient and high stress condition (Minnick and Raghavan 2012).

For the 118 unique DIPs, only 3 DIPs were identified in consensus mode and 115 in population mode (Figure 8). Lengths of insertions and deletions were always very minor with 93% of DIPs representing an insertions or deletions of a single bp (Table S3). The multi base pair deletions included deletions of 2, 3, 7 and 12 bp that occurred in coding region and the longest 32 base pair deletion occurring in an intergenic region. However, none of these genes appeared significantly affected transcriptionally by the deletion as there were no significant transcriptomic changes. Of the genes that were downregulated, 7 had DIPs mutations but all of them being in only a fraction of Cb populations within a passage (mostly 5-10% populations).

An interesting observation was the numerous mutations (SNPs and DIPs) over several passages in two genes i.e., *lapA* and *lapB*. In LapA, a 97 aa long protein has a non-sense mutation at the 85^th^ position in passage 13. LapB, 389 aa long protein, on the other hand has two missense mutations in a large proportion of cells, with one mutation propagating to 100% of the population at passage 67. It also has numerous insertions in the coding region but the noticeable one is a 3 bp deletion in the coding region that propagates to later passages (Table S3). Although these genes didn’t show any change in gene transcriptional expression modulated by mutations, it is possible that these genes are en route to simplifying the LPS and O-antigen layer in accordance with the absence of a environment, as seen in some bacteria (Maldonado, Sa-Correia and Valvano 2016).

Collectively, the low levels of DNA mutations within the passage populations and possibly the lack of effects, suggest a very stable and minor level in genomic mutations in modulating transcriptional levels.

## Discussion

Here, we attempted to identify genes and proteins crucial to Cb intracellular growth lifestyle using a “reverse evolution” approach paired with RNAseq and DNAseq comparative transcriptomics and genomics, respectively. We transitioned Cb Nine Mile phase II from cell cultures into the axenic defined media ACCM-D and subcultured it in a long-term successive passage model. Temporal changes in gene expression patterns, and DNA mutations associated with adaptation to an axenic extracellular lifestyle were identified. In general, we observe a significant number of differential expression (464 up, 371 down, 38% of overall Cb genes) through 67 passages. It is interesting to note that the majority (288 upregulated and 249 downregulated) of differentially expressed genes expressed an “early up” or “early down” expression pattern (Figure 3, Table S1), suggestive of a relatively rapid adaptation (within 31 passages out of 67 total passages) into this new axenic environment.

Differentially expressed genes identified in this study could be grouped into multiple structural and functional categories (secretory apparatus, effector proteins, other pathogenicity determinants, hypothetical proteins, and central metabolic pathways). In general, a broad (19/24 genes) encoding T4BSS components showed significant expression change, with 18 genes showing a decrease in the expression whereas only one gene that was upregulated. T4BSS is the most crucial conduit for pathogenicity and effector proteins in Cb (Carey, Newton, Luhrmann, *et al*. 2011; van Schaik, Chen, Mertens, *et al*. 2013; Voth and Heinzen 2009). Components of the T4BSS span both membranes and the periplasm and are bridged by the core transport complex comprising proteins DotC, DotD, DotF, DotG and DotH, which are predicted to provide a channel for export of effector substrates (Figure 5a) (Vincent, Friedman, Jeong, *et al*. 2006). The coupling protein complex provides a link between substrates and transport complex and includes DotL, DotM, DotN, IcmS and IcmW (Vincent, Friedman, Jeong, *et al*. 2012). DotB is an essential cytoplasmic protein with an ATPase activity and unknown function, but its mutation has been linked to failure in secreting effector proteins during the infection of host cells (Beare, Larson, Gilk, *et al*. 2012). DotA and IcmX has been shown to be released from the bacteria (Luedtke, Mahapatra, Lutter, *et al*. 2017). Besides these, other components of the T4BSS includes DotO localized in the cytoplasm, IcmX in periplasmic space, DotK in outer membrane whereas IcmF, IcmH, DotI, DotJ, DotA, DotE, DotP, IcmV and IcmT in inner membrane. (Figure 5a). The genes involved in the T4BSS in Cb are clustered in a single locus made up of two regions, with the exceptionof *icmF* and *dotU,* which are part of a separate operon (Figure 5b). This is similar to the gene rearrangement shown in the original *Coxiella burnetii* sequence (Seshadri, Paulsen, Eisen, *et al*. 2003). Gene *icmF*, which has been shown to be involved in intra-macrophage replication and inhibition of phagosome-lysosome fusion in *L. pneumophila* (VanRheenen, Duménil and Isberg 2004; Zusman, Feldman, Halperin, *et al*. 2004) and stabilization of the secretion complex (Sexton, Miller, Yoneda, *et al*. 2004) was the only T4BSS component that showed transcriptional upregulation.

The observed downregulation of this experimentally verified central intracellular pathogenic determinant makes biological sense and provides a general overall credence that gene downregulation under the experimental setting employed in this study could be regarded as a reasonable proxy for requirement for intracellular survival in cell-cultures. In addition to T4BSS, other secretory pathway such as the general secretory (sec) pathway and a component of type I secretary pathway (i.e., TolC) also exhibited a general trend of overall downregulation (Figure 5A, Figure S1a, Figure S1b, Table S1). In general, we interpret such overall lower expression of structural secretory apparatuses as a reflection of less need for these systems during interaction between Cb and the environment in a relatively rich axenic setting when compared to the organisms environmentally “normal” intracellular setting.

Interestingly, while genes encoding the production of secretary pathways were downregulated, expression patterns of predicted T4BSS effector proteins were mixed, with 33 upregulated and 14 downregulated. Of the 14 downregulated genes (all of which were early downregulated), nine have been experimentally verified based on experimental evidence of their translocation by the Dot/Icm system (Carey, Newton, Luhrmann, *et al*. 2011; Chen, Banga, Mertens, *et al*. 2010; Lifshitz, Burstein, Peeri, *et al*. 2013; Maturana, Graham, Sharma, *et al*. 2013; Voth, Beare, Howe, *et al*. 2011; Voth, Howe, Beare, *et al*. 2009; Weber, Chen, Rowin, *et al*. 2013), three genes containing ankyrin repeat domains (*ankG*, *ankD* and *ankK*) were considered effectors based on the presence of eukaryotic like domains and subsequently shown to be translocated by the Dot/Icm system(Voth, Howe, Beare, *et al*. 2009), gene *cirC (Coxiella* effector for intracellular replication) was verified as effector by transposon insertion mutation studies where its mutation was associated with a defect in Coxiella containing vacuole (CCV) biogenesis (Weber, Chen, Rowin, *et al*. 2013), and lastly a hypothetical protein B7L74_09020 was verified to be an effector based on loss-of-function mutation where its mutation was related to a smaller CCV phenotype (Crabill, Schofield, Newton, *et al*. 2018)(Table S2).

AnkG, AnkD and AnkK are ankyrin repeat-containing effector proteins in Cb (Seshadri, Paulsen, Eisen, *et al*. 2003). The eukaryotic type Ank domain in this protein family might have a role in host-cell attachment and allows the interaction of bacteria with a spectrum of host cell proteins and thus are particularly important in the pathogenic process (Batrukova, Betin, Rubtsov, *et al*. 2000; Cordsmeier, Rinkel, Jeninga, *et al*. 2022; Pechstein, Schulze-Luehrmann, Bisle, *et al*. 2020; Voth, Howe, Beare, *et al*. 2009). AnkD has both eukaryotic like domain and F-box domain, but the function is not yet clear (Voth, Howe, Beare, *et al*. 2009). AnkG has been shown to localize at the host microtubules and interferes with the host apoptosis pathway by interacting with the host protein gC1qR (p32) (Luhrmann, Nogueira, Carey, *et al*. 2010; Voth, Howe, Beare, *et al*. 2009). AnkK has been shown to have an important role for bacterial growth inside of macrophages (Habyarimana, Al-Khodor, Kalia, *et al*. 2008), although it is not delivered to the host cell via T4BSS (Voth, Howe, Beare, *et al*. 2009) (Table S2). *Coxiella* <u>p</u>lasmid <u>e</u>ffector proteins (CpeF and CpeH) are in the plasmid T4BSS effector family of proteins, and important for disrupting host cell mechanisms. CpeF specifically localized in host cell during infection, and has shown to cause a growth defect when mutated (Martinez, Cantet, Fava, *et al*. 2014; Voth, Beare, Howe, *et al*. 2011) whereas CpeH localizes to the host cell’s cytoplasm (Maturana, Graham, Sharma, *et al*. 2013). Cig57 mutation has been linked to an intracellular replication defect for Cb, whereas a Cig2 mutation causes both a growth defect and a CCV fusion defect (Newton, Kohler, McDonough, *et al*. 2014). These two proteins are early downregulated in 9 and 7 passages, respectively. CirC has been shown to be important for CCV biogenesis (Weber, Chen, Rowin, *et al*. 2013) and it is early downregulated in 4 passages. Of the remaining 6 downregulated effectors, 4 are hypothetical proteins with unknown functions, Cbu1752 has been shown to be important for vacuole biogenesis, and Cbu0635 appears important for host cell secretion (Table S2).

Of the 33 upregulated effector proteins, the majority fall into COG categories of unclassified (n=11), signal transduction mechanisms (n=7), transportation and metabolism of coenzyme and inorganic ions (n=6) (Figure 6, Table S1). These upregulated, and no expression change effector proteins (n=71), could also be involved in mediating general survival functions or other cellular functions besides their involvement in directly association with the intracellular pathogenesis process. This could be one of the explanations behind their upregulation, or no expression change, in this particular setting.

Such a pattern, where genes encoding the formation of the structural conduits (i.e., secretary pathways) are downregulated, but numerous genes encoding proteins thought to be secreted through these conduits (i.e., effector proteins) are upregulated is puzzling. We put forth the possibility that the expression of these effector proteins is controlled by Cb intracellular conditions, where high concentrations of intracellular metabolites (amino acids, inorganic salts, ATP/ADP ratio) regulate their expression. Under this scenario, high level of intracellular precursors in LCV Cb is associated with growth inside the cell, and possibly in the “rich” axenic media state of LCV growth. It remains to be seen whether translation of these effector transcripts to protein products and subsequent secretion occurs in Cb grown in axenic media. Limited reports suggest that Cb T4BSS effectors have not been observed during axenic media growth (Stead, Omsland, Beare, *et al*. 2013; Shaw, Unpublished data)

Multiple additional pathogenic determinants were also downregulated in axenic media. Specifically, chaperons, LPS, and peptidoglycan synthesis. Amongst the 16 genes annotated as chaperons in the Cb genome, 10 were transcriptionally downregulated starting at early passages (Figure 7a) (Table S1). The downregulation could be explained by the fact that chaperons play important roles for withstanding stress associated with intracellular survival (e.g., CCV detoxification) (Beare, Unsworth, Andoh, *et al*. 2009), survival inside the macrophage (Takaya, Tomoyasu, Matsui, *et al*. 2004) and low pH within the CCV (Macellaro, Tujulin, Hjalmarsson, *et al*. 1998). . Downregulations in the genes involved in synthesis of Lipid A and O-antigen biosynthesis as well as peptide cross linking during peptidoglycan layer suggests the possible further reduction in the virulence or need for these functions of *Coxiella burnetii* in axenic media, even within the CB NMII strain used in this study. These changes in the bacterial cell coverings could be tied to role/importance of these genes in intracellular host cell manipulation and bacterial survival and growth inside the host cell. These assumptions will certainly need to be verified with more experimental evidence. Here, we need to keep in mind that the strain we are using is the *Coxiella burnetii* avirulent strain Nine Mile phase II (NMII). These variants have a truncated LPS, due to a genomic deletion of about 25 Kbp of sequences that encodes all the three sugars i.e., virenose, dihydrohydroxystreptose, and galactosaminuronyl-α(1,6)-glucosamine that comprises the LPS O-antigen biosynthesis (Amano, Williams, Missler, *et al*. 1987; Denison, Massung and Thompson 2007). Thus, further downregulations in some of the remaining genes involved in LPS and O-antigen synthesis pathways could be due to their already altered functionality in the cell.

Analysis of central metabolic pathways showed a clear trend of downregulation of multiple catabolic (e.g., glycolysis and electron transport chain), amphibolic (e.g., citric acid cycle) and anabolic (e.g., FA synthesis) pathways, with a parallel upregulation of genes encoding transporters (Figure 7C). Such pattern could readily be explained by the nutrient rich growth environment (ACCM-D media) where Cb is grown in our continuous passage model. This media is a defined axenic medium for Cb growth and contains all 20 amino acids, salts (sodium phosphate and sodium bicarbonate), vitamins, minerals, and trace elements (Sandoz, Beare, Cockrell, *et al*. 2016). As such, the need for expression of genes encoding enzymes that are components of these biosynthetic pathways decreases and subsequently, the overall need for ATP generation for biosynthetic purposes (hence a decrease in respiratory activity). Likewise, the upregulation in several structural genes encoding transporters could be explained by the bacteria increasing production of more channels/transporters to accommodate the increased presence of metabolites/substrates from the media as opposed to needing to produce them via metabolic pathways in the bacterial cytoplasm. Finally, 30 hypothetical proteins were downregulated, and their predicted localization, predicted role in pathogenesis and their general analysis provide some possible explanations for such pattern (see results section). Regardless, we suggest that these could be important, hitherto untested possible pathogenicity determinants. Future biochemical and genetic efforts to test such assumptions represent a ready avenue for future research directions.

We also hypothesized that continuous passaging could also lead to the propagation of mutations, DNA fragment loses, and rearrangements in genes involved in intracellular survival, pathogenesis, and host cell manipulations could occur. While this is a well-known process, it doesn’t appear that within the timeframe of the experiments here (67 passages) that this is the case. The careful study of each of these mutations, their corresponding gene expression changes, number of passages affected by the mutations and their transiency, and overall fraction of population affected by these mutations showed that these mutations do not appear to be significant at the gene or transcript expression level. Out of 960 unique mutations (SNPs and DIPs collectively) observed at different spots within the genomes of various passages, only one mutation (i.e., a SNP) in GTP pyrophosphokinase SpoT, a signal transduction component and transcriptional regular with a role in helping *Coxiella* cope with the low-nutrient and high stress conditions in the CCV, seems to have effect on gene transcript expression (Minnick and Raghavan 2012). Early downregulation observed in this gene could be attributed to the fact that these cells are less stressed due to the growth condition in rich ACCM-D media, and this mutation could have caused the decrease in transcript expression of this gene. Overall, these findings suggest that Cb gene expression changes significantly following acclimation to axenic media, although extensive genomic rearrangement does not occur. Genomics reveals a relatively stable Cb NMII genome over the 67 passages that were analyzed here.

In conclusion, we present a detailed temporal analysis on how Cb transition from intracellular growth, (where a wide range of cellular processes is thought to be required to maintain survival and growth), to a defined, rich axenic media (where many of such processes are theoretically, no longer needed). As any genome-wide transcriptomics survey, the approach is useful for uncovering patterns, confirming prior observations, and generating new insights and hypothesis. We stress that while transcriptional downregulation in axenic media compared to cell culture could broadly be associated with a genes importance for survival in cell cultures, the precise nature of such correlation is yet unclear, and that differential expression patterns could further be modified on the translational and post translational levels. Experimental assessments and validation of many of the observed patterns may well open new avenues of Cb research. Nevertheless, our analysis is beneficial in providinge information on how specific genes and pathways in Cb may be important for this unusual organisms intracellular survival, as well as to identify putatively novel pathogenicity determinants in this naturally intracellular pathogen.

## Acknowledgements

**Acknowledgments.** This work was supported by the NIH grant number 5R03AI149144-02.

