## Supplementary document for "Comparative Transcriptomics and Genomics from Continuous Axenic Media Growth Identifies *Coxiella burnetii* Intracellular Survival Strategies"

**Figure S1. A)** Plot count graphs for all 19 DEG T4BSS genes. **B)** Plot count graphs for all 7 DEGs in the sec. pathway. The y-axis represents log of counts normalized by estimated size factors and x-axis represents the passage. Dots represent replicates in a passage, pink dots represent passages in which L2Fc was found to be significant, and the line(red) represents the mean of the counts in each passage.

A.

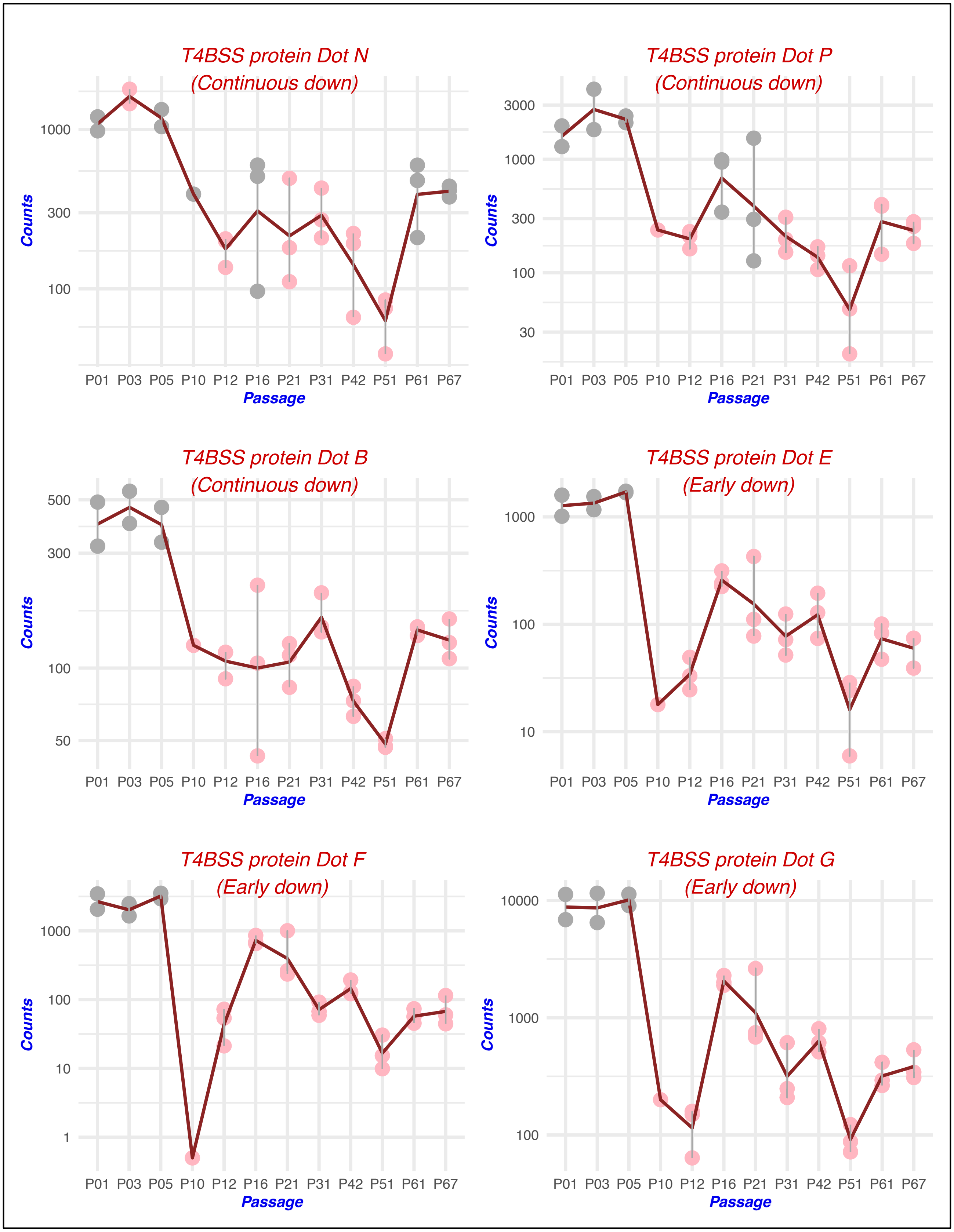
T4BSS protein Dot N
(Continuous down)
T4BSS protein Dot P
(Continuous down)
T4BSS protein Dot B
(Continuous down)
T4BSS protein Dot E
(Early down)
T4BSS protein Dot F
(Early down)
T4BSS protein Dot G
(Early down)
Counts
Passage
P01 P03 P05 P10 P12 P16 P21 P31 P42 P51 P61 P67

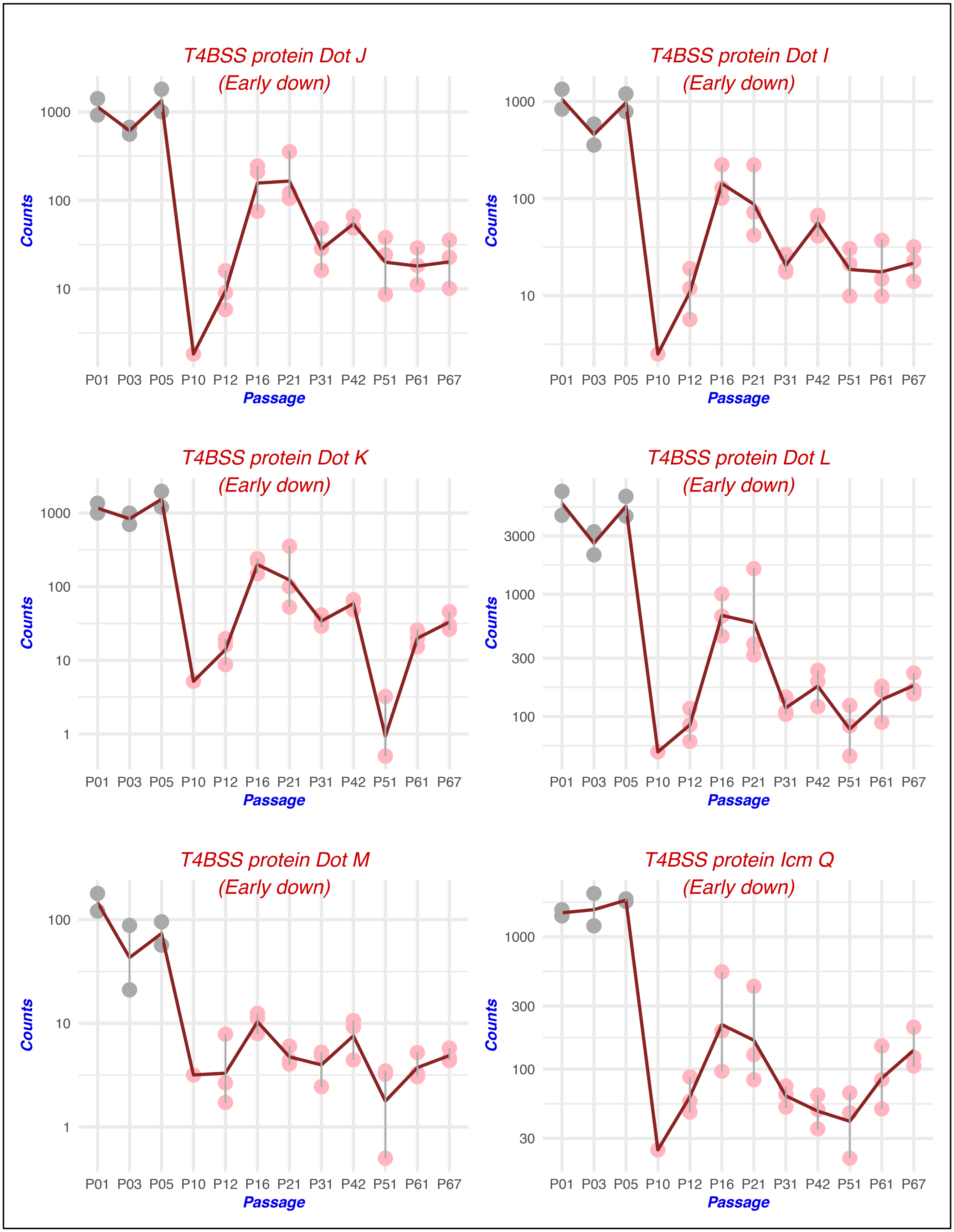

T4BSS protein Dot J
(Early down)
T4BSS protein Dot I
(Early down)
T4BSS protein Dot K
(Early down)
T4BSS protein Dot L
(Early down)
T4BSS protein Dot M
(Early down)
T4BSS protein Icm Q
(Early down)
Counts
Passage
P01 P03 P05 P10 P12 P16 P21 P31 P42 P51 P61 P67
1000
100
10
1
3000
300
30

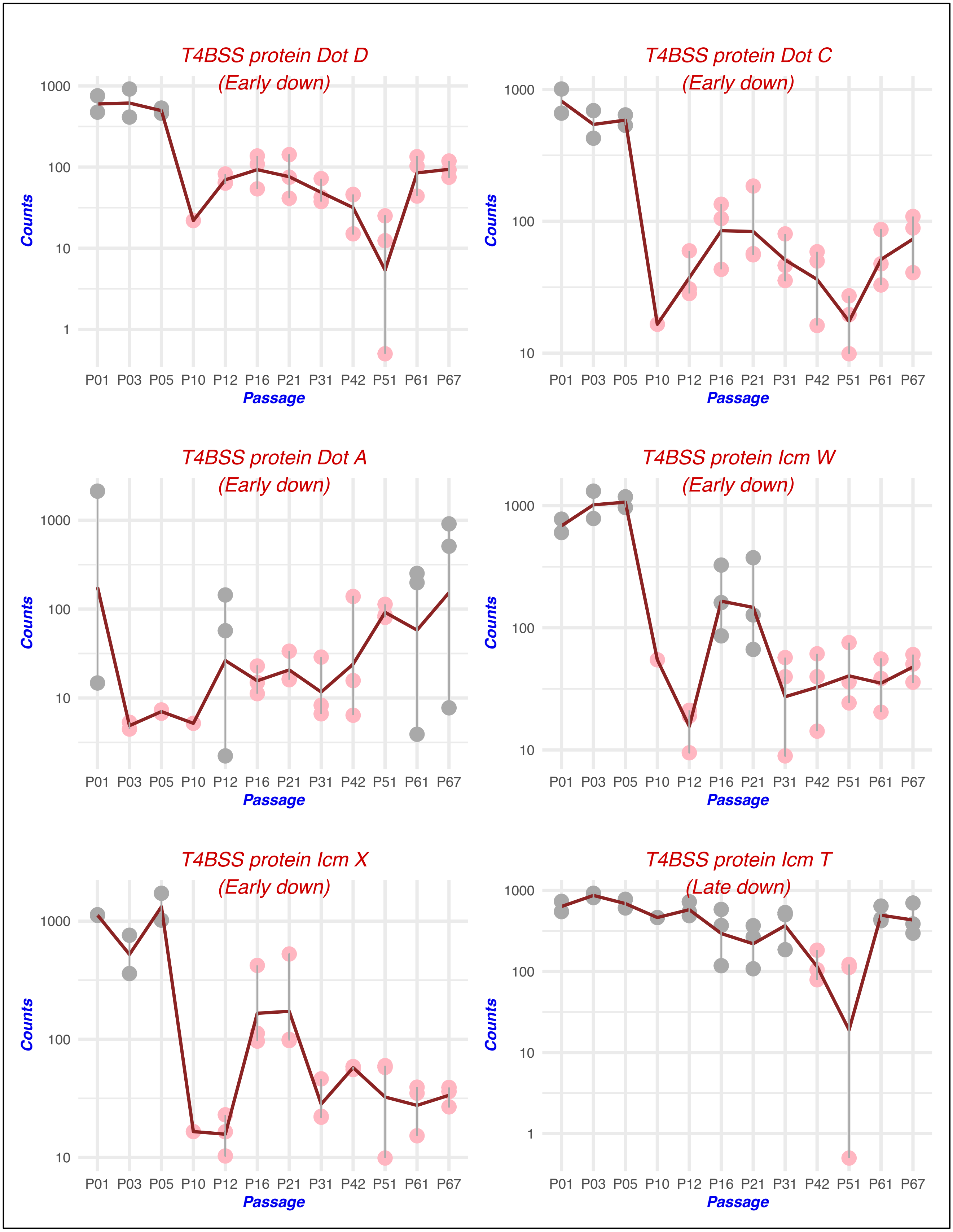

T4BSS protein Dot D
(Early down)
T4BSS protein Dot C
(Early down)
T4BSS protein Dot A
(Early down)
T4BSS protein Icm W
(Early down)
T4BSS protein Icm X
(Early down)
T4BSS protein Icm T
(Late down)
Counts
Passage
P01 P03 P05 P10 P12 P16 P21 P31 P42 P51 P61 P67
1000
100
10
1

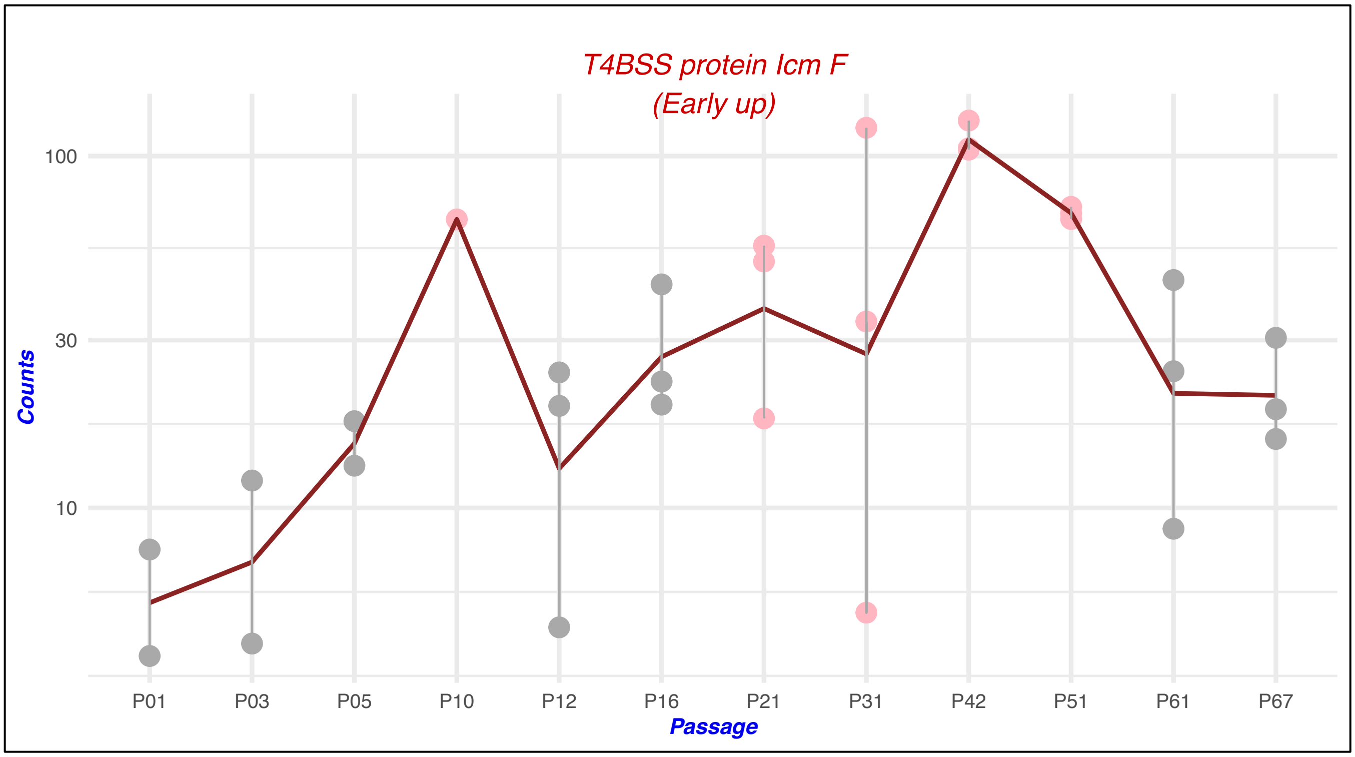
T4BSS protein Icm F
(Early up)
Counts
100
30
10
P01
P03
P05
P10
P12
P16
P21
P31
P42
P51
P61
P67
Passage

B.

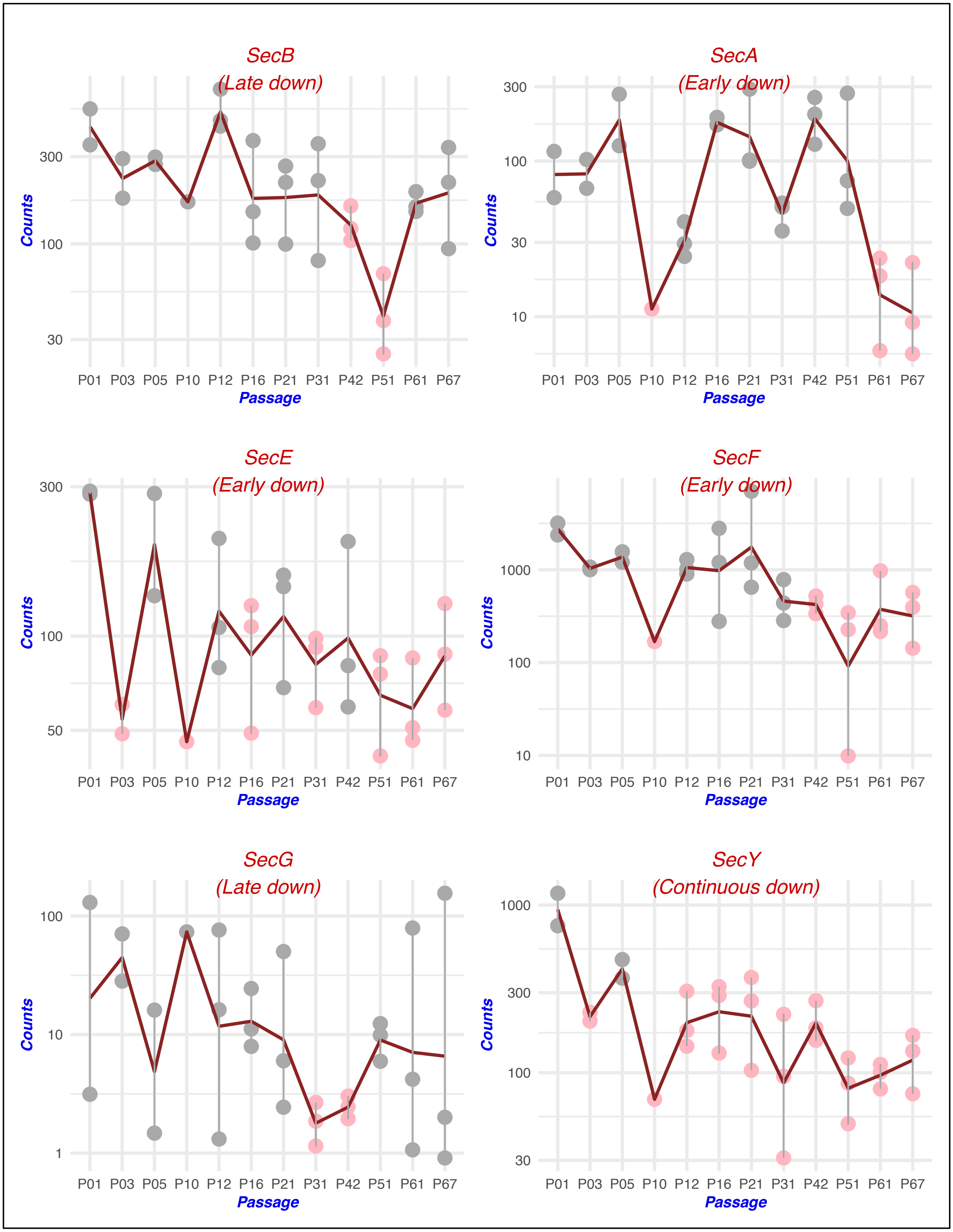
SecB
(Late down)
SecA
(Early down)
SecE
(Early down)
SecF
(Early down)
SecG
(Late down)
SecY
(Continuous down)
Counts
Passage
P01 P03 P05 P10 P12 P16 P21 P31 P42 P51 P61 P67
30
100
300
10
50
1000
1

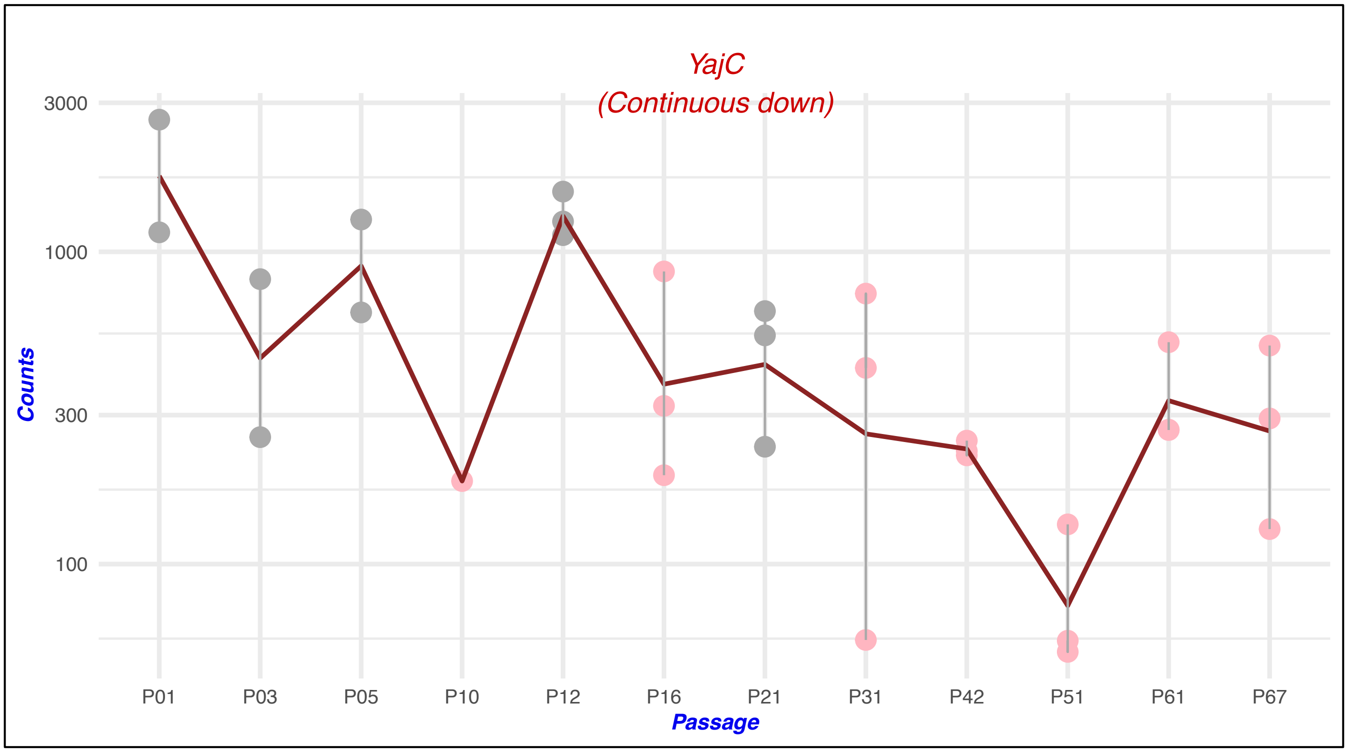

YajC
(Continuous down)
3000
1000
300
100
Counts
P01
P03
P05
P10
P12
P16
P21
P31
P42
P51
P61
P67
Passage

**Figure S2:** Plot count graphs for 14 downregulated effector proteins. The y-axis represents log of counts normalized by estimated size factors and x-axis represents the passage. Dots represent replicates in a passage, pink dots represent passages in which L2Fc was found to be significant, and the red line represents the mean of the counts in each passage.

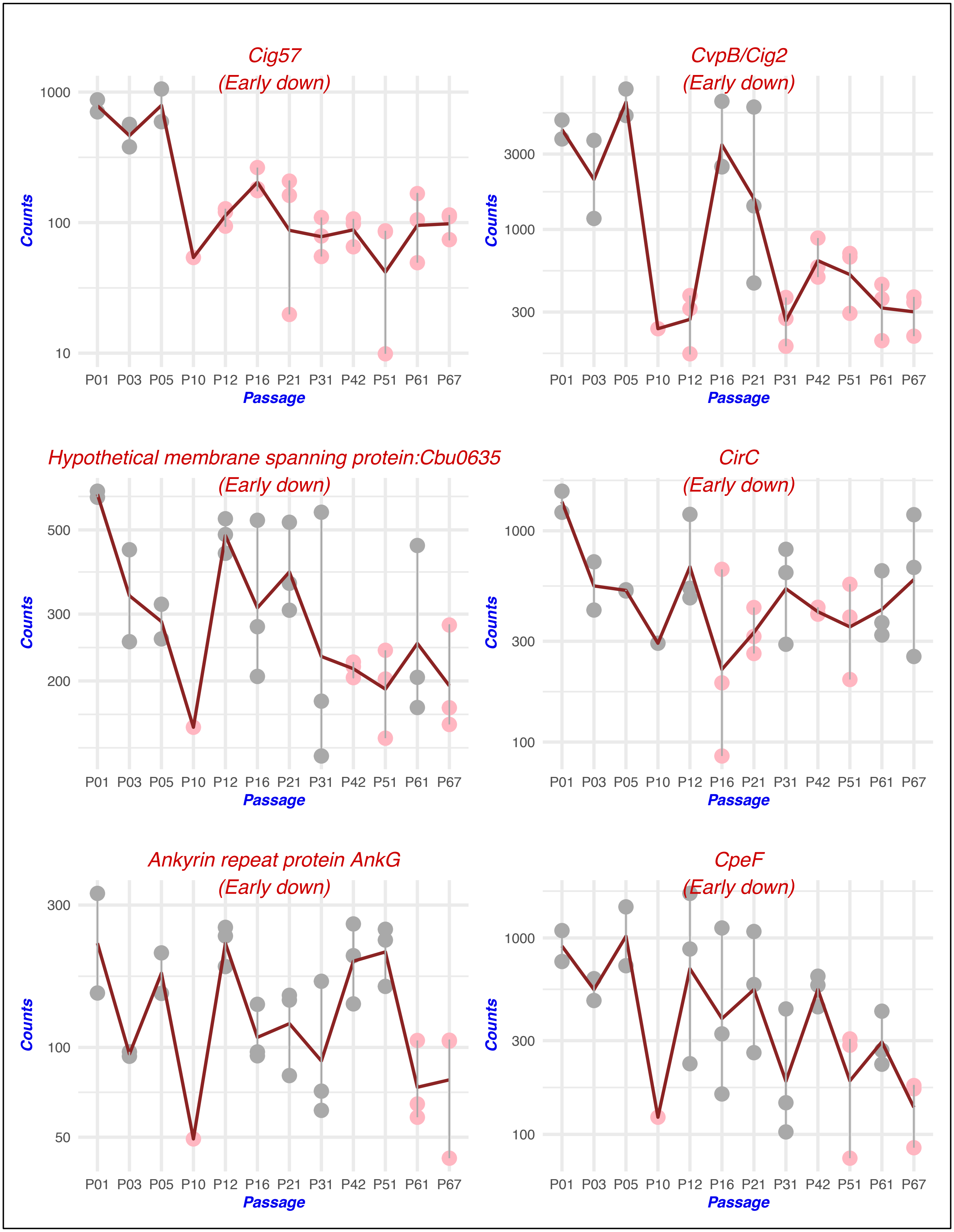
Cig57
(Early down)
CvpB/Cig2
(Early down)
Hypothetical membrane spanning protein:Cbu0635
(Early down)
CirC
(Early down)
Ankyrin repeat protein AnkG
(Early down)
CpeF
(Early down)
Counts
Passage
P01 P03 P05 P10 P12 P16 P21 P31 P42 P51 P61 P67
10
100
1000
300
3000
200
500
50

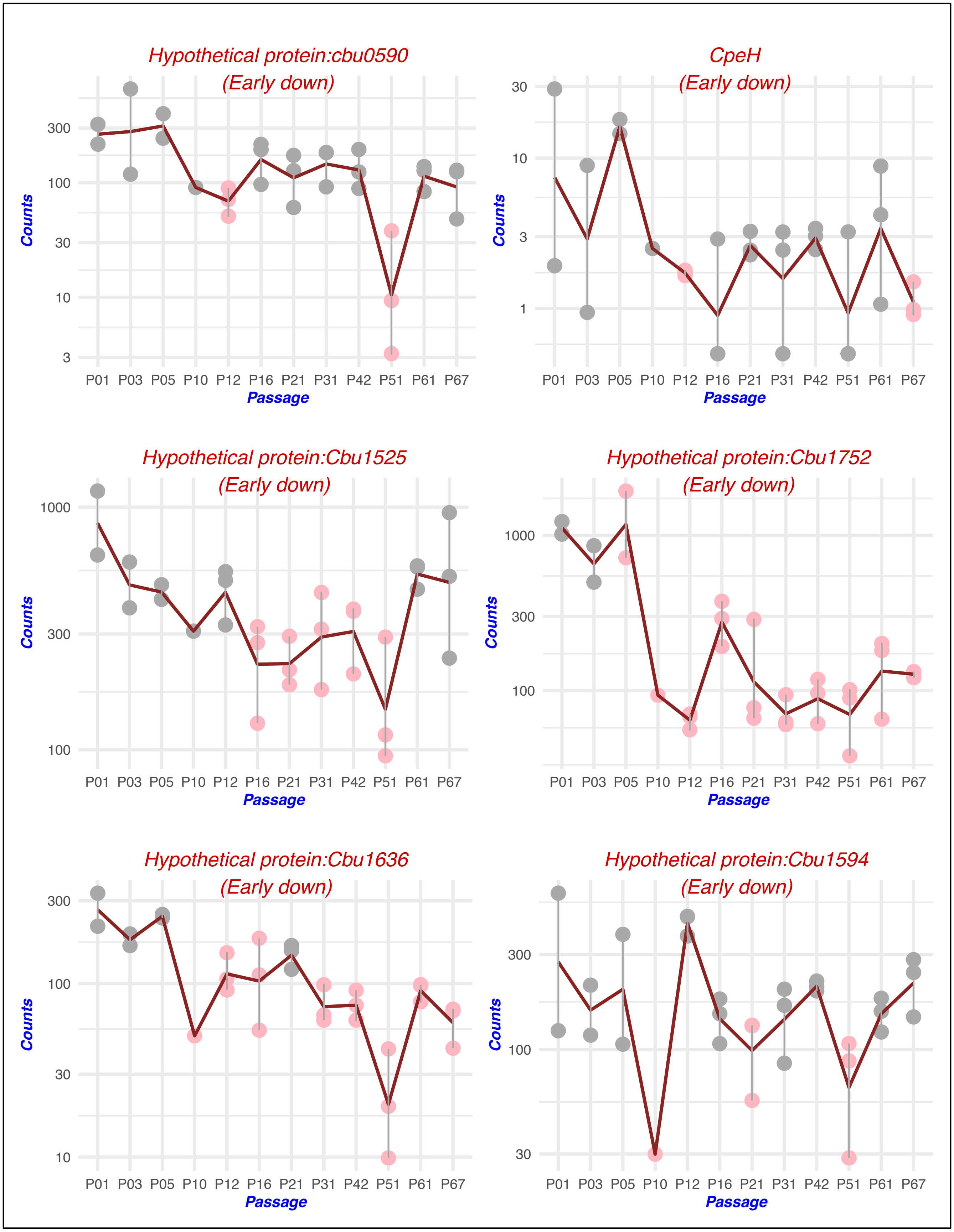

Hypothetical protein:cbu0590
(Early down)
Counts
300
100
30
10
3
P01 P03 P05 P10 P12 P16 P21 P31 P42 P51 P61 P67
Passage
CpeH
(Early down)
Counts
30
10
3
1
P01 P03 P05 P10 P12 P16 P21 P31 P42 P51 P61 P67
Passage
Hypothetical protein:Cbu1525
(Early down)
Counts
1000
300
100
P01 P03 P05 P10 P12 P16 P21 P31 P42 P51 P61 P67
Passage
Hypothetical protein:Cbu1752
(Early down)
Counts
1000
300
100
P01 P03 P05 P10 P12 P16 P21 P31 P42 P51 P61 P67
Passage
Hypothetical protein:Cbu1636
(Early down)
Counts
300
100
30
10
P01 P03 P05 P10 P12 P16 P21 P31 P42 P51 P61 P67
Passage
Hypothetical protein:Cbu1594
(Early down)
Counts
300
100
30
P01 P03 P05 P10 P12 P16 P21 P31 P42 P51 P61 P67
Passage

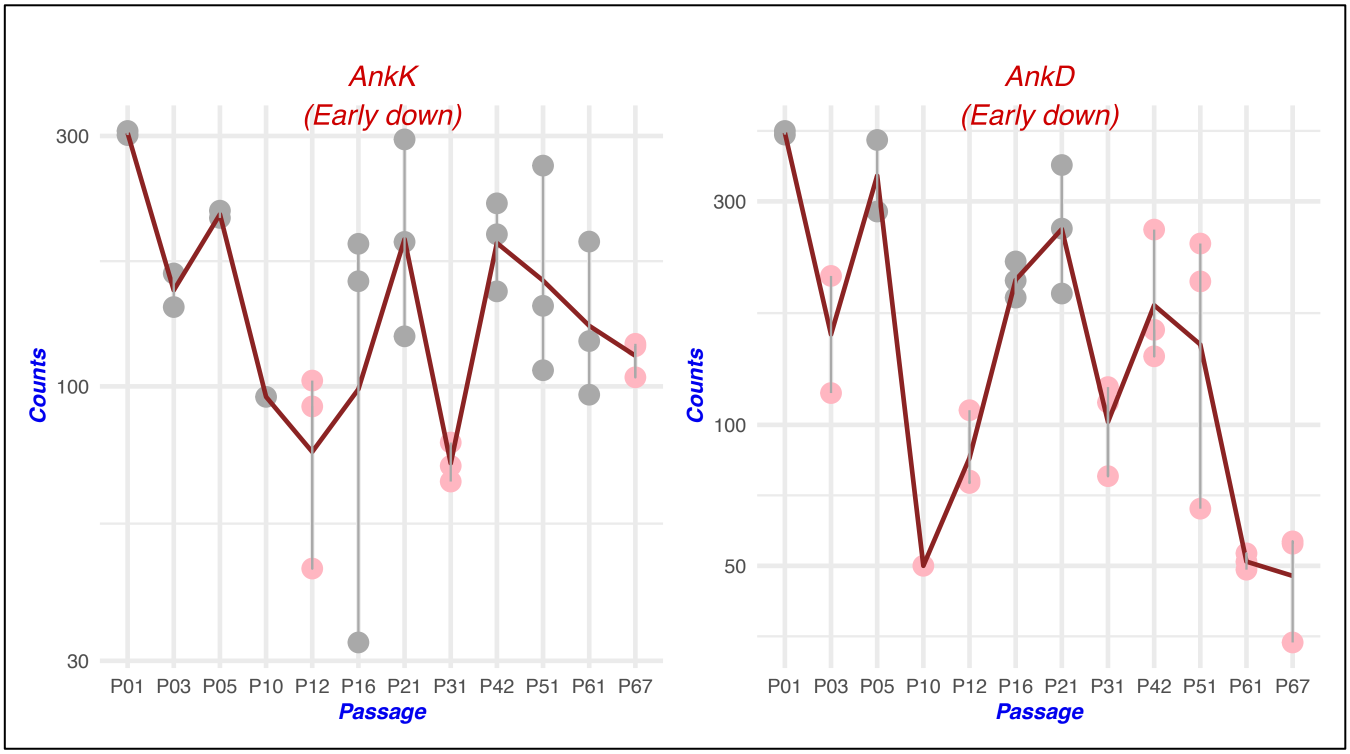

AnkK
(Early down)
Counts
300
100
30
P01 P03 P05 P10 P12 P16 P21 P31 P42 P51 P61 P67
Passage
AnkD
(Early down)
Counts
300
100
50
P01 P03 P05 P10 P12 P16 P21 P31 P42 P51 P61 P67
Passage

**Figure S3:** Heatmap for DEGs involved in central metabolic pathways and transporters. The L2fc values used to plot differential expression of the genes ranges from -8 to 10 over different passage.

| Pathway | Gene | Annotation | Verdict | Differential expression (L2FC value compared to P01) | | | | | | | | | | |
|---|---|---|---|---|---|---|---|---|---|---|---|---|---|---|
| | | | | P03 | P05 | P10 | P12 | P16 | P21 | P31 | P42 | P51 | P61 | P67 |
| Glycolysis / Gluconeogenesis | B7L74_01545 | *pmm-pgm* | Early down | | | | | | | | | | | |
| | B7L74_01780 | *pfkA* | Early up | | | | | | | | | | | |
| | B7L74_02360 | *aceE, pdhC* | Continuous down | | | | | | | | | | | |
| | B7L74_02365 | *aceF, pdhC* | Down in P42 | | | | | | | | | | | |
| | B7L74_02370 | *lpd, pdhD* | Continuous down | | | | | | | | | | | |
| | B7L74_06780 | *acs* | Late up | | | | | | | | | | | |
| | B7L74_09150 | *fbaA* | Down in P51 | | | | | | | | | | | |
| | B7L74_09170 | *gapA* | Early down | | | | | | | | | | | |
| | B7L74_06520 | phosphatase | Early down | | | | | | | | | | | |
| | B7L74_10840 | *pckA* | Early down | | | | | | | | | | | |
| Citrate cycle (TCA) | B7L74_02360 | *aceE* | Continuous down | | | | | | | | | | | |
| | B7L74_02365 | *aceF, pdhC* | Down in P42 | | | | | | | | | | | |
| | B7L74_02370 | *lpd, pdhD* | Continuous down | | | | | | | | | | | |
| | B7L74_06175 | *IDH2, icd* | Early down | | | | | | | | | | | |
| | B7L74_06385 | *mdh* | Up in P12 | | | | | | | | | | | |
| | B7L74_07200 | *sucD* | Early down | | | | | | | | | | | |
| | B7L74_07210 | *sucB* | Down in P42 | | | | | | | | | | | |
| | B7L74_07215 | *sucA* | Continuous down | | | | | | | | | | | |
| | B7L74_07220 | *sdhB, frdB* | Continuous down | | | | | | | | | | | |
| | B7L74_07225 | *sdhA, frdA* | Early down | | | | | | | | | | | |
| | B7L74_07230 | *sdhD, frdD* | Early down | | | | | | | | | | | |
| | B7L74_08855 | *acnA* | Late down | | | | | | | | | | | |
| | B7L74_10840 | *pckA* | Early down | | | | | | | | | | | |
| Oxidative phosphorylation | B7L74_03245 | *ppa* | Continuous down | | | | | | | | | | | |
| | B7L74_04955 | *cydA* | Early down | | | | | | | | | | | |
| | B7L74_04960 | *cydB* | Late up | | | | | | | | | | | |
| | B7L74_04965 | *cydX* | Late down | | | | | | | | | | | |
| | B7L74_05340 | *cyoD* | Early down | | | | | | | | | | | |
| | B7L74_05345 | *cyoC* | Early down | | | | | | | | | | | |
| | B7L74_07390 | *nuoN* | Continuous down | | | | | | | | | | | |
| | B7L74_07395 | *nuoM* | Early down | | | | | | | | | | | |
| | B7L74_07400 | *nuoL* | Early down | | | | | | | | | | | |
| | B7L74_07405 | *nuoK* | Early down | | | | | | | | | | | |
| | B7L74_07415 | *nuoI* | Early down | | | | | | | | | | | |
| | B7L74_07420 | *nuoH* | Continuous down | | | | | | | | | | | |
| | B7L74_07430 | *nuoF* | Early down | | | | | | | | | | | |
| | B7L74_07435 | *nuoE* | Continuous down | | | | | | | | | | | |
| | B7L74_07440 | *nuoD* | Continuous down | | | | | | | | | | | |
| | B7L74_07450 | *nuoB* | Early down | | | | | | | | | | | |
| | B7L74_10020 | *atpB* | Early down | | | | | | | | | | | |
| | B7L74_10025 | *atpE* | Early down | | | | | | | | | | | |
| | B7L74_10030 | *atpF* | Early down | | | | | | | | | | | |
| | B7L74_10040 | *atpA* | Early down | | | | | | | | | | | |
| | B7L74_10045 | *atpG* | Continuous down | | | | | | | | | | | |
| | B7L74_10050 | *atpD* | Early down | | | | | | | | | | | |
| Fatty Acid biosynthesis | B7L74_00185 | *acpP* | Early down | | | | | | | | | | | |
| | B7L74_00190 | *fabB* | Early down | | | | | | | | | | | |
| | B7L74_00195 | *fabZ* | Early up | | | | | | | | | | | |
| | B7L74_00200 | *fabA* | Early down | | | | | | | | | | | |
| | B7L74_00210 | *fabH* | Early down | | | | | | | | | | | |
| | B7L74_02540 | *fabD* | Early down | | | | | | | | | | | |
| | B7L74_02545 | *fabG* | Early down | | | | | | | | | | | |
| | B7L74_02550 | *acpP* | Early down | | | | | | | | | | | |
| | B7L74_02555 | *fabF* | Continuous down | | | | | | | | | | | |
| LPS and O-antigen biosynthesis | B7L74_00735 | *lpxC* | Early down | | | | | | | | | | | |
| | B7L74_03170 | *lpxD* | Continuous down | | | | | | | | | | | |
| | B7L74_03180 | *lpxA* | Early down | | | | | | | | | | | |
| | B7L74_03475 | *wbpW* | Early down | | | | | | | | | | | |
| | B7L74_03485 | *gmhB* | Early down | | | | | | | | | | | |
| | B7L74_03505 | *galE* | Down in P10 | | | | | | | | | | | |
| | B7L74_04220 | *wbpD* | Early down | | | | | | | | | | | |
| | B7L74_04225 | *galE* | Early down | | | | | | | | | | | |
| | B7L74_04290 | *wbpI* | Down in P10 | | | | | | | | | | | |
| | B7L74_04300 | *cap1J* | Late down | | | | | | | | | | | |
| | B7L74_10060 | *glmU* | Early down | | | | | | | | | | | |

10
6
3
0
-1
-4
-8

| | | | |
|---|---|---|---|
| Transporters | B7L74_02485 | *artQ* | Early up |
| | B7L74_03540 | *wzm, rfbA* | Early up |
| | B7L74_03545 | *wzt, rbfB* | Early down |
| | B7L74_05125 | *lolD* | Early down |
| | B7L74_05130 | *lolC_E* | Early up |
| | B7L74_09820 | *ftsE* | Early down |
| | B7L74_10570 | *lapB* | Early up |
| | B7L74_00460 | *pal* | Down in P51 |
| | B7L74_00480 | *yrbG* | Early up |
| | B7L74_01455 | MarC family | Early up |
| | B7L74_01835 | *czcD, zitB* | Late down |
| | B7L74_01880 | *PHO84* | Continuous up |
| | B7L74_01910 | *bcr, tcaB* | Early up |
| | B7L74_02215 | MFS transporter | Early up |
| | B7L74_02490 | *artM* | Early up |
| | B7L74_02665 | MFS transporter | Early up |
| | B7L74_02695 | *dedA* | Early up |
| | B7L74_03280 | EamA family | Down in P10 |
| | B7L74_03345 | *perM* | Up in P16 |
| | B7L74_03565 | *perM* | Late up |
| | B7L74_03840 | *bamD* | Up in P12 |
| | B7L74_03905 | Multidrug transport | Early up |
| | B7L74_04030 | MFS transporter | Late up |
| | B7L74_04070 | *emrA* | Early down |
| | B7L74_04245 | *pglK* | Early down |
| | B7L74_04625 | MFS transporter | Early up |
| | B7L74_04765 | *ABC-2.A* | Late up |
| | B7L74_05540 | *ynaI, mscMJ* | Early up |
| | B7L74_05815 | Oligopeptide transp | Early up |
| | B7L74_06065 | *cmlA, cmlB, floR* | Late up |
| | B7L74_06390 | *fadL* | Late up |
| | B7L74_06400 | MFS transporter | Early up |
| | B7L74_06620 | MFS transporter | Late up |
| | B7L74_06940 | *glcP* | Late up |
| | B7L74_07280 | *omp31* | Early up |
| | B7L74_07620 | *ahpC* | Early up |
| | B7L74_07760 | *copA, ctpA* | Early down |
| | B7L74_08085 | MFS transporter | Early up |
| | B7L74_08145 | *mnhF, mrpF* | Down in P10 |
| | B7L74_08165 | *mnhB, mrpB* | Early up |
| | B7L74_08170 | *mnhA, mrpA* | Continuous up |
| | B7L74_08255 | *mscK, kefA* | Early up |
| | B7L74_08895 | *pnuC* | Early up |
| | B7L74_08915 | Na+ symporter;NS | Up in P21 |
| | B7L74_08920 | Na+ symporter;NS | Continuous up |
| | B7L74_09100 | *feoB* | Early up |
| | B7L74_09260 | *yhhQ* | Early up |
| | B7L74_09330 | *TC.OOP;OmpA-O* | Early up |
| | B7L74_10175 | Bcr/CflA family ef | Continuous up |
| Chaperons | B7L74_06645 | *dnaJ* | Early down |
| | B7L74_06650 | *dnaK, HSPA9* | Early down |
| | B7L74_03255 | *fklB* | Early down |
| | B7L74_08845 | *groEL, HSPD1* | Continuous down |
| | B7L74_08850 | *groES* | Continuous down |
| | B7L74_07830 | *grxC* | Late down |
| | B7L74_03030 | *grxD* | Early down |
| | B7L74_10420 | *hslU* | Continuous down |
| | B7L74_01615 | *htpG, HSP90A* | Continuous down |
| | B7L74_10235 | *surA* | Late down |

Table S1. Mastertable with all the genes and their expression values. Gene annotations were obtained from NCBI annotation of the reference genome (Genbank accession number:CP020616.7). Verdicts are made for 845 Differentially expressed genes (DEGs). Column 4-14 indicates the L2fc values for each genes when compared to passage 1. The cells with significant L2fc values (pvalue<0.05) are color coded, where green celles representing upregulation (L2fc>0) and orange cells representing downregulation (L2fc<0). Column 16-25 indicates if the gene is involved in these specific pathways and where green "Yes" means they are present in differentially expressed genes (DEGs) list. For Column 23 (Transporters), classification of transporters by substrates is shown inside the brackets.

| TRANSCRIPT/ REF_GENE_ ID | GENE ANNOTATION | Involvement in Pathogenesis/Pathway | 2FOLD CHANGE COMPARED TO P1 (Color coded Adj Pval < | | | | | | | | | | | Verdict | INVOLVED IN? | | | | | | | | | |
|---|---|---|---|---|---|---|---|---|---|---|---|---|---|---|---|---|---|---|---|---|---|---|---|---|
| | | | P03 | P05 | P10 | P12 | P16 | P21 | P31 | P42 | P51 | P61 | P67 | | T4BSS | Effector proteins | Chaperons | Glycolysis/ Gluconeogenesis | TCA cycle | Respiration | FA biosynthesis | Transporters | LPS and O-antigen biosynthesis | Peptidoglycan synthesis |
| B7L74_00005 | hypothetical protein | Effector protein | -0.958 | 0.265 | -3.077 | -0.471 | 0.589 | -0.453 | -0.844 | 0.329 | -0.414 | -0.406 | -1.026 | | | Yes | | | | | | | | |
| B7L74_00010 | chromosomal replication initiator protein DnaA | Two-component system | -1.885 | -1.008 | -3.377 | -0.233 | -2.561 | -2.197 | -2.253 | -2.119 | -4.474 | -1.038 | -1.042 | Early down | | | | | | | | | | |
| B7L74_00015 | DNA polymerase III subunit beta | DNA replication | -1.569 | -1.045 | -2.243 | -4.960 | -1.076 | -0.790 | -1.281 | -2.117 | -0.033 | -1.439 | -2.796 | Early down | | | | | | | | | | |
| B7L74_00020 | DNA replication and repair protein RecF | DNA Homologous recombination | -0.525 | -0.845 | -1.542 | -0.321 | -1.502 | -1.660 | -1.943 | -1.709 | -2.365 | -1.126 | -1.213 | Late down | | | | | | | | | | |
| B7L74_00025 | DNA gyrase subunit B | | -0.164 | -0.725 | 1.568 | 0.802 | -1.407 | -1.227 | 0.392 | 0.072 | 0.484 | 0.715 | 0.980 | | | | | | | | | | | |
| B7L74_00030 | IS110 family transposase | | -0.253 | -1.007 | 1.732 | -0.064 | -1.408 | -1.110 | 1.747 | -1.267 | -0.685 | 1.144 | 1.562 | | | | | | | | | | | |
| B7L74_00035 | hypothetical protein | | 2.904 | -0.771 | 3.559 | -1.085 | 2.906 | 0.406 | 0.857 | 1.372 | 1.433 | 0.830 | 0.340 | | | | | | | | | | | |
| B7L74_00040 | antitoxin | | -1.237 | -0.145 | 0.502 | -1.028 | -0.060 | -0.442 | -0.783 | 0.111 | -0.254 | 0.471 | 0.524 | | | | | | | | | | | |
| B7L74_00045 | hypothetical cytosolic protein | | -0.177 | 1.806 | 3.671 | 1.361 | 0.025 | 1.859 | 2.930 | 1.530 | 3.820 | 0.836 | 2.836 | | | | | | | | | | | |
| B7L74_00050 | hypothetical protein | | -0.303 | 0.528 | -0.396 | 0.742 | -0.713 | -1.220 | -0.510 | -0.518 | 0.251 | 0.173 | 1.341 | | | | | | | | | | | |
| B7L74_00055 | hypothetical protein | | 2.234 | 0.495 | 0.679 | 0.622 | 3.661 | 2.979 | 0.236 | -1.065 | -2.072 | 1.582 | 0.438 | | | | | | | | | | | |
| B7L74_00060 | D-alanyl-D-alanine carboxypeptidase/D-alanyl-D-alani | Peptidoglycan syn | 1.743 | 0.347 | -0.606 | -0.452 | 2.146 | 4.251 | 1.116 | -0.858 | 3.330 | 0.190 | -0.136 | Early up | | | | | | | | | | Yes |
| B7L74_00065 | shikimate dehydrogenase | | 1.001 | 2.443 | -0.627 | 3.178 | 2.040 | 2.280 | 0.965 | 0.892 | 2.623 | 4.104 | 1.959 | | | | | | | | | | | |
| B7L74_00070 | hypothetical protein | | 2.833 | 1.863 | 0.975 | -1.483 | 3.752 | 4.453 | 0.778 | 2.511 | 1.195 | -0.350 | -1.358 | | | | | | | | | | | |
| B7L74_00075 | hypothetical protein | | -0.092 | 0.012 | -0.768 | -0.569 | 0.313 | 0.295 | -1.154 | -2.278 | -1.728 | -1.355 | -1.521 | Late down | | | | | | | | | | |
| B7L74_00080 | anion permease | | -0.068 | 0.469 | -0.021 | 1.036 | -0.635 | -0.665 | -0.665 | -0.787 | -1.917 | 0.135 | 0.073 | | | | | | | | | Yes (Ions/Phosphate) | | |
| B7L74_00085 | TIGR00153 family protein | | -0.860 | 1.059 | -0.420 | -0.330 | 0.070 | -0.974 | -0.783 | 0.389 | 0.044 | 1.315 | 0.407 | | | | | | | | | | | |
| B7L74_00090 | purine-nucleoside phosphorylase | Purine metabolism | -0.570 | 0.184 | -1.390 | -0.466 | -1.296 | -1.654 | -1.501 | -1.954 | -2.581 | -0.674 | -0.906 | Late down | | | | | | | | | | |
| B7L74_00095 | 2-deoxyribose-5-phosphate aldolase | | 0.949 | 0.880 | -0.262 | -1.525 | 0.634 | 1.073 | -0.075 | -0.331 | 1.636 | -1.689 | -1.593 | | | | | | | | | | | |
| B7L74_00100 | hypothetical protein, Hypothetical cytosolic protein | | -1.821 | 0.203 | -1.376 | -1.260 | -0.859 | -1.084 | -2.017 | 0.019 | -1.066 | -1.161 | -1.900 | Early down | | | | | | | | | | |
| B7L74_00105 | hypothetical protein | | 0.541 | 1.614 | 1.105 | -1.702 | 1.699 | 0.822 | 0.158 | 1.470 | 2.184 | -1.176 | -0.165 | | | | | | | | | | | |
| B7L74_00110 | ribosomal subunit interface protein | Ribosomal proteins | 0.469 | 1.913 | 4.018 | -1.230 | 2.362 | 1.783 | 1.866 | 2.565 | 2.997 | -0.463 | 0.304 | Early up | | | | | | | | | | |
| B7L74_00115 | hypothetical protein(cvpB/cig2 in Table 1 of RightonQ | Effector protein | -0.841 | 0.586 | -4.204 | -3.919 | -0.179 | -0.732 | -3.980 | -2.729 | -2.967 | -3.686 | -3.805 | Early down | | Yes | | | | | | | | |
| B7L74_00120 | hypothetical cytosolic protein | | -0.319 | -0.068 | 0.491 | 1.414 | -0.094 | -0.276 | -0.245 | 0.430 | -0.157 | -0.458 | -0.788 | | | | | | | | | | | |
| B7L74_00125 | hypothetical cytosolic protein | | -0.117 | -0.283 | 0.741 | 1.475 | -0.877 | -1.441 | -0.058 | -0.336 | -0.958 | 0.153 | -0.282 | | | | | | | | | | | |
| B7L74_00130 | carbon storage regulator | Signal transduction | -1.731 | -0.861 | -0.370 | 1.179 | -1.180 | 0.161 | 0.268 | 0.156 | -1.328 | -0.984 | -1.041 | | | | | | | | | | | |
| B7L74_00135 | hypothetical protein | | 0.663 | 0.953 | 1.418 | 3.062 | -0.010 | 1.212 | 1.396 | 2.676 | 1.183 | 0.603 | 1.384 | Early up | | | | | | | | | | |
| B7L74_00140 | ribose-5-phosphate isomerase | | -0.619 | -0.504 | 1.049 | 1.305 | 0.210 | -0.243 | 0.609 | -0.203 | 0.285 | 0.223 | 0.057 | | | | | | | | | | | |
| B7L74_00145 | acyltransferase | | -0.253 | -1.506 | -1.218 | 0.439 | -0.620 | -0.892 | -0.470 | 0.829 | -0.271 | 0.333 | 0.220 | | | | | | | | | | | |
| B7L74_00150 | FAD-binding oxidoreductase | | 0.385 | -1.618 | -0.518 | 0.440 | 1.079 | 0.083 | -0.248 | -0.881 | 0.726 | 0.026 | 0.112 | | | | | | | | | | | |
| B7L74_00155 | 1-acyl-sn-glycerol-3-phosphate acyltransferase | | -0.484 | -0.892 | 0.248 | 1.082 | -0.896 | -0.328 | -0.085 | 0.416 | -0.176 | 0.292 | 0.510 | | | | | | | | | | | |
| B7L74_00160 | hydroxymethylglutaryl-CoA reductase | | -0.736 | -0.432 | 1.323 | 2.328 | 0.801 | 0.257 | 1.992 | 1.130 | 1.408 | 2.029 | 2.227 | | | | | | | | | | | |
| B7L74_00165 | short-chain dehydrogenase | | -1.213 | -1.215 | -0.491 | 0.827 | -1.602 | -0.930 | -0.463 | -0.518 | -1.323 | -0.278 | -0.167 | | | | | | | | Yes | | | |
| B7L74_00170 | hypothetical protein | | 0.116 | -0.209 | -0.685 | -1.239 | -1.506 | -1.325 | 0.751 | 0.120 | -1.347 | -1.348 | -0.611 | | | | | | | | | | | |
| B7L74_00175 | hypothetical cytosolic protein | | -2.365 | 1.763 | 1.497 | -2.604 | 1.052 | 2.008 | 2.965 | 1.528 | 1.796 | 0.153 | 0.159 | | | | | | | | | | | |
| B7L74_00180 | 4'-phosphopantetheinyl transferase | | -1.250 | -1.330 | -3.041 | -0.302 | -1.739 | -1.272 | -0.358 | -0.508 | -1.409 | -0.592 | -1.064 | Down in P10 | | | | | | | | | | |
| B7L74_00185 | acyl carrier protein | Fatty acid biosynthesis | -2.189 | -1.289 | -3.375 | -0.598 | -1.306 | -2.069 | -2.064 | -2.406 | -2.656 | -1.211 | -1.785 | Early down | | | | | | | Yes | | | |
| B7L74_00190 | beta-ketoacyl-ACP synthase | Fatty acid biosynthesis | -0.380 | -1.101 | -0.454 | 0.614 | -0.952 | -1.622 | -0.083 | -1.900 | -2.860 | -0.103 | 0.794 | Early down | | | | | | | Yes | | | |
| B7L74_00195 | beta-hydroxyacyl-ACP dehydratase | Fatty acid biosynthesis | 0.093 | 1.510 | -0.034 | -1.758 | 4.085 | 5.975 | 0.503 | 3.545 | 5.242 | -0.719 | 0.834 | Early up | | | | | | | Yes | | | |
| B7L74_00200 | beta-hydroxydecanoyl-ACP dehydratase | Fatty acid biosynthesis | -1.290 | -1.377 | -2.824 | -0.646 | -3.097 | -2.985 | -2.004 | -2.960 | -2.416 | -1.710 | -1.597 | Early down | | | | | | | Yes | | | |
| B7L74_00205 | hypothetical protein | | 0.090 | 0.848 | 2.408 | 1.628 | 3.893 | 2.589 | 0.527 | 1.496 | 3.724 | 0.823 | 0.049 | | | | | | | | | | | |
| B7L74_00210 | ketoacyl-ACP synthase III | Fatty acid biosynthesis | -1.357 | -1.492 | -3.539 | 0.214 | -2.249 | -2.705 | -1.553 | -0.895 | -3.315 | -1.713 | -1.437 | Early down | | | | | | | Yes | | | |
| B7L74_00215 | oligopeptidase A | Peptidases and inhibitors | -2.399 | -1.833 | -3.034 | -0.767 | -2.577 | -2.225 | -2.195 | -2.159 | -3.768 | -2.324 | -2.085 | Early down | | | | | | | | | | |
| B7L74_00220 | IS110 family transposase | | 0.461 | 0.303 | 1.861 | -0.622 | -0.204 | -0.303 | 0.414 | -0.089 | -1.357 | 1.171 | 1.103 | | | | | | | | | | | |
| B7L74_00225 | hypothetical protein | | -1.847 | -1.111 | -2.000 | -0.283 | -1.671 | -1.819 | -2.197 | -1.950 | -0.835 | -2.194 | -2.852 | | | | | | | | | | | |
| B7L74_00230 | hypothetical protein(cirA in Table 1 of RightonQ pape | Effector protein | 0.929 | -0.147 | -0.008 | -0.666 | 1.258 | 0.992 | 0.035 | -0.227 | 1.403 | 1.130 | 0.847 | | | Yes | | | | | | | | |
| B7L74_00235 | ferrochelatase | | -1.724 | -1.257 | -0.932 | 0.269 | -0.788 | -0.896 | -1.274 | -0.870 | -0.009 | -0.534 | 0.181 | | | | | | | | | | | |
| B7L74_00240 | non-canonical purine NTP pyrophosphatase | Purine metabolism | 0.554 | -0.012 | 1.803 | 2.112 | -0.441 | -0.723 | 0.590 | -0.135 | -1.332 | 0.315 | 0.414 | Up in P12 | | | | | | | | | | |
| B7L74_00245 | hypothetical exported protein | | -1.061 | -1.169 | -0.214 | 1.788 | -1.299 | -1.214 | -0.502 | -1.339 | -2.299 | -0.390 | -0.002 | Down in P51 | | | | | | | | | | |
| B7L74_00250 | lipoprotein | | -1.170 | -0.903 | -0.531 | 1.975 | -1.516 | -1.735 | -0.226 | -1.003 | -1.875 | 0.419 | 0.534 | | | | | | | | | | | |
| B7L74_00255 | mannose-1-phosphate guanyltransferase | | -0.848 | -0.525 | 1.613 | -0.054 | -2.438 | -1.681 | 0.231 | -0.616 | -0.362 | 0.247 | 0.585 | | | | | | | | | Yes (General) | | |
| B7L74_00260 | multidrug ABC transporter ATP-binding protein | Transporter | 0.400 | 1.010 | 1.426 | 1.135 | 1.278 | -0.241 | 0.571 | 0.809 | 1.282 | 1.494 | 1.365 | | | | | | | | | Yes (drug resistance) | | |
| B7L74_00265 | hemolysin D | | -1.421 | -0.437 | 1.503 | -0.663 | 0.047 | -0.370 | 0.467 | 0.995 | 2.947 | -0.194 | 0.278 | | | | | | | | | | | |
| B7L74_00270 | hypothetical protein | | -0.156 | 0.209 | 1.009 | 0.970 | -0.796 | -1.023 | -0.353 | 0.561 | -0.167 | 1.223 | 1.200 | | | | | | | | | | | |
| B7L74_00275 | hypothetical cytosolic protein | | -0.512 | -1.004 | -0.465 | -0.625 | -1.257 | -2.104 | -0.302 | -0.991 | -1.580 | 0.083 | -0.835 | | | | | | | | | | | |
| B7L74_00280 | hypothetical protein | | -1.086 | -0.244 | -1.024 | -0.720 | -0.793 | -1.219 | -0.780 | -0.461 | -0.737 | -0.071 | -0.586 | | | | | | | | | | | |
| B7L74_00285 | L%2CD-transpeptidase, Enhanced entry protein | Entry | 0.589 | 0.483 | 2.079 | -1.991 | 0.493 | 0.200 | 0.809 | 0.059 | 1.347 | 0.225 | 0.063 | | | | | | | | | | | |
| B7L74_00290 | chorismate--pyruvate lyase | | -0.918 | -0.806 | -0.266 | 0.565 | -1.088 | -1.999 | -1.064 | -1.011 | -0.733 | 0.052 | -0.022 | | | | | | | | | | | |
| B7L74_00295 | 4-hydroxybenzoate octaprenyltransferase | | 1.423 | 1.058 | 1.485 | 1.394 | 2.187 | 2.671 | 1.572 | 1.338 | 1.678 | -0.767 | 2.049 | | | | | | | | | | | |
| B7L74_00300 | channel protein TolCType I secretion outer membrane | Secretion system I beta-Lactam resistance | -0.544 | -0.780 | -0.403 | -0.992 | -0.834 | -0.613 | -0.793 | -1.254 | -2.194 | -1.240 | -1.216 | Late down | | | | | | | | | | |
| B7L74_00305 | DUF1658 domain-containing protein | | 1.164 | -0.437 | 0.495 | 0.229 | 1.276 | 0.695 | 1.025 | -1.654 | -0.902 | -0.012 | -0.484 | | | | | | | | | | | |
| B7L74_00310 | hypothetical protein | | 2.502 | -0.371 | 0.960 | 1.384 | 1.633 | 2.831 | 1.109 | 0.490 | 1.788 | 2.079 | 2.202 | | | | | | | | | | | |
| B7L74_00315 | hypothetical protein | | 3.132 | 0.727 | 1.559 | 1.128 | 3.394 | 3.162 | 1.960 | 1.877 | 2.941 | 2.583 | 2.719 | Early up | | | | | | | | | | |
| B7L74_00320 | molecular chaperone DnaJ | Effector protein | 6.290 | 2.090 | 3.029 | 1.531 | 6.981 | 7.036 | 4.077 | 1.618 | 5.601 | 3.467 | 2.690 | Early up | | Yes | | | | | | | | |
| B7L74_00325 | 3-deoxy-D-manno-octulosonic acid transferase | Lipopolysaccharide biosynthesis | 1.362 | 0.848 | 2.408 | -1.750 | 3.885 | 2.589 | 1.611 | 2.458 | 2.628 | 5.002 | 0.049 | | | | | | | | | | Yes | |
| B7L74_00330 | DNA topoisomerase IV subunit B | DNA replication proteins | -0.368 | -1.696 | -0.172 | -0.012 | -1.829 | -1.731 | -0.408 | -1.604 | -1.936 | 0.358 | 0.416 | Early down | | | | | | | | | | |
| B7L74_00335 | sulfurtransferase | | -0.922 | -1.279 | -0.929 | 0.024 | -1.455 | -2.436 | -0.574 | -2.119 | -1.676 | -0.282 | 0.107 | Early down | | | | | | | | | | |
| B7L74_00340 | hypothetical protein | | -0.371 | -0.059 | -0.058 | 0.126 | -0.874 | -0.721 | 0.159 | -0.032 | 0.052 | 1.242 | 0.378 | | | | | | | | | | | |
| B7L74_00345 | 5-formyltetrahydrofolate cyclo-ligase | | 1.345 | 1.217 | 3.456 | -0.341 | 1.139 | 0.476 | 1.059 | 1.956 | -0.795 | 1.686 | 1.422 | | | | | | | | | | | |
| B7L74_00350 | 6S RNA (gene=ssrS) | rRNA | 1.180 | 1.629 | 4.778 | -0.810 | 2.283 | 1.548 | 3.425 | 3.249 | 2.007 | 0.625 | 0.855 | Early up | | | | | | | | | | |
| B7L74_00355 | cell division protein ZapA | | -0.280 | -0.741 | 0.627 | 1.468 | -0.485 | -1.593 | 0.229 | -1.090 | -1.057 | 1.803 | 0.607 | | | | | | | | | | | |
| B7L74_00360 | TIGR02449 family protein | | -1.210 | -0.205 | 1.785 | 2.052 | 1.776 | -0.816 | 1.059 | 0.345 | -0.151 | 2.141 | 0.637 | | | | | | | | | | | |
| B7L74_00365 | hypothetical protein | | 1.898 | 0.538 | 1.650 | -0.909 | 1.337 | 0.710 | 1.124 | 1.343 | 1.621 | -0.537 | 0.624 | | | | | | | | | | | |
| B7L74_00370 | hypothetical protein | Effector protein | 0.787 | -0.160 | 0.634 | -0.346 | 1.212 | 1.052 | 0.417 | -0.472 | 0.289 | 1.022 | 1.089 | | | Yes | | | | | | | | |
| B7L74_00375 | hypothetical protein(ankA in Table 1 of RightonQ pap | Effector protein | 1.823 | -0.359 | 2.719 | 2.338 | 1.361 | 1.818 | 2.398 | -0.460 | 1.297 | 3.370 | 3.536 | Late up | | Yes | | | | | | | | |
| B7L74_00380 | Xaa-Pro aminopeptidase | Peptidases and inhibitors | -1.934 | -1.252 | -2.789 | -0.368 | -2.087 | -2.468 | -2.071 | -2.228 | -3.069 | -2.088 | -2.073 | Early down | | | | | | | | | | |
| B7L74_00385 | hypothetical protein | | -0.561 | -0.342 | -1.930 | -1.286 | -1.450 | -1.893 | -1.134 | -2.227 | -2.811 | -1.721 | -1.226 | | | | | | | | | | | |
| B7L74_00390 | 2-octaprenyl-3-methyl-6-methoxy-1%2C4-benzoquino | | -10.131 | -0.689 | -1.509 | -0.504 | -2.878 | -0.477 | 0.061 | -1.496 | -1.445 | -0.382 | -0.384 | | | | | | | | | | | |
| B7L74_00395 | ubiquinone biosynthesis protein UbiH | | -1.279 | 1.816 | -0.035 | 0.371 | 1.712 | 1.754 | 1.484 | 2.742 | 4.038 | 0.892 | -0.803 | | | | | | | | | | | |
| B7L74_00400 | hypothetical membrane spanning protein | Effector protein | 0.370 | -0.081 | -1.173 | 0.315 | 0.597 | 1.264 | -0.939 | -1.161 | -0.439 | -0.337 | -0.125 | | | Yes | | | | | | | | |
| B7L74_00405 | heavy metal transporter | Transporter | 4.132 | 1.018 | 1.082 | 1.406 | 5.268 | 5.592 | 1.659 | -0.436 | 3.911 | 1.993 | 1.702 | Early up | | | | | | | | | | |
| B7L74_00410 | proline--tRNA ligase | | 0.925 | 1.028 | 1.246 | -0.012 | 0.024 | -0.321 | 0.870 | 1.206 | 0.745 | 0.741 | 1.149 | | | | | | | | | | | |
| B7L74_00415 | hypothetical protein | | 1.675 | 1.384 | 2.544 | 0.800 | 0.577 | 0.308 | 1.693 | 1.465 | 1.166 | 1.383 | 2.249 | | | | | | | | | | | |
| B7L74_00420 | sulfatase | | -0.602 | 0.086 | -2.126 | -3.083 | -1.745 | -2.497 | -2.036 | -1.279 | -1.212 | -1.893 | -1.535 | Early down | | | | | | | | | | |
| B7L74_00425 | entericidin B | | -0.682 | 0.028 | -0.425 | 0.635 | -0.192 | -1.640 | -0.937 | -0.880 | -1.050 | -1.296 | -1.253 | | | | | | | | | Yes (toxins) | | |
| B7L74_00430 | amidohydrolase | | -0.274 | -0.306 | -1.019 | -0.147 | -0.539 | -1.138 | -1.096 | -0.322 | -0.723 | -1.998 | -1.571 | | | | | | | | | | | |
| B7L74_00435 | phosphoglycerol transferase | | 0.249 | 1.242 | 0.086 | -1.434 | 1.687 | 1.679 | -0.680 | 1.313 | 2.334 | -0.613 | -0.940 | Early up | | | | | | | | | | |
| B7L74_00440 | protein TolA | | 0.459 | 0.299 | 0.582 | -1.975 | 0.769 | 0.645 | -0.582 | -1.024 | -0.866 | -0.658 | -1.445 | | | | | | | | | Yes (ion channels/mechanosensitive channel) | | |
| B7L74_00445 | hypothetical protein | | -0.724 | -0.752 | -0.777 | -0.507 | -0.361 | -0.786 | -1.416 | -1.096 | -1.424 | -1.376 | -1.573 | | | | | | | | | | | |
| B7L74_00450 | hypothetical protein | | 0.108 | 1.329 | 5.355 | 0.017 | 2.570 | 1.883 | 1.740 | 1.803 | 1.938 | -0.236 | 0.053 | Early up | | | | | | | | | | |
| B7L74_00455 | Tol-Pal system beta propeller repeat protein TolB | | -0.660 | -0.500 | 0.347 | -1.098 | -0.630 | -0.817 | -1.245 | -1.137 | -1.434 | -0.692 | -0.519 | | | | | | | | | Yes (ion channels/mechanosensitive channel) | | |
| B7L74_00460 | peptidoglycan-associated lipoprotein | Transporter | -1.504 | -0.504 | -0.219 | -0.176 | -0.385 | -0.630 | -1.317 | -1.178 | -2.378 | -1.313 | -1.076 | Down in P51 | | | | | | | | Yes (Outer membrane/Lipoprotein) | | |
| B7L74_00465 | tol-pal system protein YbgF | | -0.536 | 0.247 | -0.629 | -2.385 | 2.418 | 1.028 | 0.936 | 1.727 | 4.317 | -1.864 | -1.500 | | | | | | | | | | | |
| B7L74_00470 | 7-carboxy-7-deazaguanine synthase QueE | Folate biosynthesis | -0.581 | -0.689 | -2.259 | -1.226 | -0.822 | -1.378 | -2.096 | -1.367 | -1.746 | -1.365 | -1.008 | Early down | | | | | | | | | | |
| B7L74_00475 | chaperone protein ClpB | | -2.204 | -2.674 | -1.199 | -1.129 | -2.056 | -1.704 | -1.519 | -2.799 | -3.126 | -1.232 | -1.323 | Early down | | | | | | | | | | |
| B7L74_00480 | sodium:calcium exchanger | Transporter | 2.996 | 3.362 | 7.926 | 1.822 | 4.830 | 2.321 | 5.713 | 5.593 | 5.829 | 2.991 | 4.484 | Early up | | | | | | | | Yes (Na+:H+ antiporter ) | | |
| B7L74_00485 | cardiolipin synthase B | Eucaryotic-like ORFs. | 1.373 | 0.848 | -0.034 | -0.078 | 3.847 | 3.363 | 1.507 | 3.098 | 3.562 | 0.465 | 0.052 | | | | | | | | | | | |
| B7L74_00490 | acetyltransferase | | 1.373 | 0.848 | -0.034 | -0.078 | 3.848 | 3.363 | 1.507 | 2.458 | 3.562 | 0.856 | 0.052 | | | | | | | | | | | |
| B7L74_00495 | CDP-diacylglycerol--serine O-phosphatidyltransferase | | 1.751 | -0.372 | 2.313 | 1.474 | 2.384 | 2.440 | 1.748 | 0.345 | 1.889 | 2.551 | 2.034 | | | | | | | | | | | |
| B7L74_00500 | nicotinate-nucleotide diphosphorylase (carboxylating) | | -0.964 | -0.164 | 1.551 | -0.126 | 1.636 | 0.620 | -0.170 | 0.486 | 2.650 | -0.002 | -0.314 | | | | | | | | | | | |
| B7L74_00505 | hypothetical protein | | 1.566 | 1.024 | 3.926 | 1.861 | 1.758 | 2.457 | 1.827 | 1.215 | 1.200 | 3.648 | 2.549 | Early up | | | | | | | | | | |
| B7L74_00510 | tRNA-Glu | tRNA | 1.224 | 1.192 | 2.380 | 2.696 | 0.683 | 0.360 | 3.649 | 1.730 | -0.432 | 1.051 | 1.536 | | | | | | | | | | | |
| B7L74_00515 | tRNA-Ala | tRNA | 1.388 | 0.779 | 2.232 | 2.633 | 1.254 | 0.195 | 1.007 | 1.466 | -0.701 | 1.187 | 1.886 | Up in P12 | | | | | | | | | | |
| B7L74_00520 | hypothetical protein | | -0.064 | -1.891 | 0.616 | 2.947 | -3.594 | -0.921 | 0.763 | -0.132 | 0.352 | 2.443 | 2.409 | Up in P12 | | | | | | | | | | |
| B7L74_00525 | phosphohistidine phosphatase SixA | | 0.906 | 1.599 | -2.384 | 1.258 | 2.367 | -0.427 | 0.683 | -0.491 | 0.608 | -1.753 | -0.934 | | | | | | | | | | | |
| B7L74_00530 | lipoprotein | | 0.335 | -0.376 | 1.352 | 1.367 | -0.527 | -0.103 | 0.824 | -0.244 | -0.217 | 2.448 | 1.927 | Late up | | | | | | | | | | |
| B7L74_00535 | L-aspartate oxidase | Alanine, aspartate and glutamate m | 0.869 | -4.065 | -4.456 | -2.461 | -0.191 | -4.086 | -1.536 | -3.710 | -4.847 | -2.118 | -3.250 | Early down | | | | | | | | | | |
| B7L74_00540 | hypothetical protein | | -0.002 | -0.809 | -1.021 | 0.617 | 0.057 | -1.587 | -0.804 | -0.296 | -0.671 | 0.759 | 0.377 | | | | | | | | | | | |
| B7L74_00545 | peptidase M20 | | 0.397 | -1.274 | -0.125 | 2.438 | 0.480 | 0.897 | 0.078 | -1.208 | -0.307 | 0.365 | 0.452 | | | | | | | | | | | |
| B7L74_00550 | hypothetical protein | | 6.653 | 1.738 | 3.962 | 0.908 | 7.850 | 7.222 | 4.165 | 2.422 | 6.705 | 4.608 | 2.541 | Early up | | | | | | | | | | |
| B7L74_00555 | hypothetical protein | | -0.294 | 0.393 | 2.147 | 0.279 | 1.602 | 1.072 | 1.588 | 2.270 | 2.222 | 1.483 | 0.908 | | | | | | | | | | | |
| B7L74_00560 | hypothetical protein | | 0.148 | -0.043 | -2.477 | -1.703 | 1.969 | 0.146 | -1.916 | 0.911 | 2.154 | -0.606 | -0.231 | | | | | | | | | | | |
| B7L74_00565 | methionine import ATP-binding protein MetN | | -0.404 | -0.213 | 0.145 | -0.005 | -1.753 | -2.201 | -0.690 | 0.007 | -0.619 | 0.336 | 0.332 | | | | | | | | | Yes (Amino acid/Methionine) | | |
| B7L74_00570 | methionine ABC transporter permease | | 2.523 | 2.565 | 2.288 | -1.778 | 3.427 | 5.268 | 2.368 | 1.916 | 1.877 | 3.577 | 3.779 | | | | | | | | | Yes (Amino acid/Methionine) | | |
| B7L74_00575 | hypothetical protein methionine-binding protein | | 0.033 | 0.395 | 1.471 | -0.133 | 0.222 | -0.393 | -0.150 | -0.297 | 0.034 | 0.048 | 0.484 | | | | | | | | | Yes (Amino acid/Methionine) | | |
| B7L74_00580 | hypothetical exported protein | Exported | 0.567 | 1.121 | 3.747 | -0.505 | 1.310 | 1.127 | 1.046 | 1.205 | 1.315 | 1.110 | 1.445 | | | | | | | | | | | |
| B7L74_00585 | RNase P RNA component class A (gene=rnpB) | | 1.865 | 0.752 | 3.044 | 0.634 | 3.042 | 1.990 | 4.640 | 4.001 | 4.266 | 0.367 | 0.989 | Continuous up | | | | | | | | | | |
| B7L74_00590 | glycine C-acetyltransferase | Glycine, serine and threonine meta | -0.757 | -1.137 | -1.052 | -0.305 | -1.715 | -2.355 | -1.001 | -1.846 | -2.833 | -1.113 | -1.002 | Continuous down | | | | | | | | | | |
| B7L74_00595 | L-threonine 3-dehydrogenase | Glycine, serine and threonine meta | -0.489 | -1.247 | -0.655 | 0.241 | -1.594 | -2.458 | -0.969 | -2.105 | -2.468 | -1.016 | -0.490 | Continuous down | | | | | | | | | | |
| B7L74_00600 | hypothetical protein | Effector protein | 0.111 | 0.359 | 2.384 | 2.026 | 0.467 | -0.237 | 1.248 | 0.218 | 0.701 | 1.063 | 1.821 | | | Yes | | | | | | | | |
| B7L74_00605 | nucleotide-binding protein | Signaling proteins | -0.893 | -1.202 | -0.534 | 2.110 | -0.912 | -1.377 | -0.430 | -0.841 | -1.542 | -0.850 | -0.547 | Up in P12 | | | | | | | | | | |
| B7L74_00610 | division/cell wall cluster transcriptional repressor MraZ | | -1.117 | -1.872 | 4.750 | 0.933 | -0.374 | -0.453 | 1.397 | -0.157 | 2.449 | 1.554 | 2.473 | Early up | | | | | | | | | | |
| B7L74_00615 | ribosomal RNA small subunit methyltransferase H | | -1.729 | -1.877 | 2.104 | -0.762 | -1.631 | -1.484 | -0.963 | -1.293 | -0.133 | 0.075 | 0.206 | | | | | | | | | | | |
| B7L74_00620 | cell division protein FtsL | | -0.905 | -1.458 | 2.220 | -1.224 | -0.510 | -0.319 | 0.209 | 0.156 | -0.310 | -0.778 | -0.934 | | | | | | | | | | | |
| B7L74_00625 | penicillin-binding protein | Peptidoglycan synbeta-Lactam resi | -1.342 | -0.383 | 3.047 | -0.967 | -0.149 | -3.681 | 1.710 | 0.363 | -2.839 | -0.944 | -1.949 | Early down | | | | | | | | | | Yes |
| B7L74_00630 | hypothetical protein | | 2.990 | -0.360 | 1.143 | 0.449 | 3.295 | 2.897 | 1.849 | 1.139 | 3.229 | 2.029 | 1.918 | Early up | | | | | | | | | | |
| B7L74_00635 | hypothetical protein | | 0.612 | 0.963 | -2.025 | -1.859 | 0.997 | 0.047 | 0.289 | 1.073 | 1.548 | -0.506 | -0.347 | | | | | | | | | | | |
| B7L74_00640 | hypothetical membrane associated protein | Effector protein | 0.246 | 1.720 | 2.220 | 0.996 | 3.092 | 1.169 | 2.640 | 1.292 | 2.526 | 1.603 | 2.497 | Early up | | Yes | | | | | | | | |
| B7L74_00645 | UDP-N-acetylmuramoyl-L-alanyl-D-glutamate--2%2C | | -0.095 | -0.790 | -0.193 | 0.741 | -0.072 | -1.192 | -0.371 | -0.612 | -1.415 | 0.243 | 0.234 | | | | | | | | | | | Yes |
| B7L74_00650 | UDP-N-acetylmuramoyl-tripeptide--D-alanyl-D-alanin | | 1.152 | 0.750 | -1.856 | 0.860 | 0.926 | 0.327 | 2.029 | -1.191 | 0.983 | 1.305 | -0.022 | | | | | | | | | | | Yes |
| B7L74_00655 | phospho-N-acetylmuramoyl-pentapeptide-transferase | | -0.494 | -1.219 | -2.350 | -0.481 | -0.561 | -1.083 | -0.078 | -0.925 | -0.302 | -0.696 | -0.348 | | | | | | | | | | | Yes |
| B7L74_00660 | hypothetical protein | | 0.326 | -0.807 | 0.035 | -0.104 | 0.185 | -0.137 | 0.233 | -0.720 | 0.140 | -0.658 | -0.472 | | | | | | | | | | | |
| B7L74_00665 | hypothetical protein | | 0.171 | -0.216 | -0.171 | 2.255 | 0.125 | -0.770 | 0.020 | -0.371 | -0.117 | 1.597 | 1.074 | Up in P12 | | | | | | | | | | |
| B7L74_00670 | hypothetical protein | | 0.033 | 0.836 | -0.034 | -1.812 | 3.065 | 1.620 | 0.441 | 1.485 | 2.540 | -1.284 | -0.913 | | | | | | | | | | | |
| B7L74_00675 | hypothetical protein | Hypothetical | 3.514 | -1.570 | 1.683 | 0.699 | 2.459 | 2.732 | 2.260 | 0.896 | 1.508 | 1.250 | 0.299 | Up in P03 only | | | | | | | | | | |
| B7L74_00680 | UDP-N-acetylmuramoyl-L-alanine--D-glutamate ligase | | 0.027 | -0.035 | 0.462 | -0.175 | -0.612 | -0.536 | -0.049 | -0.334 | 0.144 | 0.351 | -0.065 | | | | | | | | | | | Yes |
| B7L74_00685 | putative lipid II flippase FtsW | | -0.547 | 0.258 | -0.611 | -2.391 | 2.052 | 1.042 | -0.139 | 1.286 | 3.111 | -1.863 | -1.491 | | | | | | | | | | | |
| B7L74_00690 | hypothetical protein | | -0.206 | -0.201 | 0.030 | 0.757 | -0.400 | -0.563 | -0.194 | -0.977 | -0.646 | 0.652 | 0.313 | | | | | | | | | | | |
| B7L74_00695 | hypothetical protein | | -3.810 | 0.052 | -1.238 | 1.529 | -0.152 | 1.171 | 0.059 | 2.177 | 1.167 | 1.410 | 0.968 | | | | | | | | | | | |
| B7L74_00700 | undecaprenyldiphospho-muramoylpentapeptide beta-N | | 0.146 | -0.041 | 1.415 | 0.320 | 0.046 | 0.050 | 0.613 | -0.094 | -0.100 | 0.920 | 1.201 | | | | | | | | | | | Yes |
| B7L74_00705 | UDP-N-acetylmuramate--L-alanine ligase | | 0.745 | -0.478 | 1.124 | 0.850 | -0.634 | -0.436 | 0.783 | -1.408 | -0.677 | 1.710 | 1.367 | | | | | | | | | | | Yes |
| B7L74_00710 | UDP-N-acetylenolpyruvoylglucosamine reductase | Peptidoglycan syn | 0.990 | 0.163 | 1.338 | 1.128 | 1.162 | 0.737 | 1.117 | 0.712 | 1.270 | 2.229 | 1.623 | Late up | | | | | | | | | | Yes |
| B7L74_00715 | cell division protein FtsQ | Cell division | 0.165 | 0.544 | 2.156 | 1.238 | 1.951 | 0.568 | 1.670 | 0.814 | 0.969 | 2.807 | 1.795 | Up in P61 | | | | | | | | | | |
| B7L74_00720 | carbonate dehydratase | | -2.906 | -3.425 | -4.308 | -3.930 | 0.740 | -2.646 | -3.746 | -2.777 | -0.578 | -3.812 | -5.170 | | | | | | | | | | | |
| B7L74_00725 | cell division protein FtsA | | 0.245 | -0.174 | 0.890 | 0.165 | -0.827 | -1.094 | 0.084 | -0.741 | -1.630 | 0.925 | 0.641 | | | | | | | | | | | |
| B7L74_00730 | cell division protein FtsZ | Cell division | -0.693 | -0.749 | -0.705 | -0.258 | -1.561 | -1.370 | -0.663 | -1.835 | -2.594 | -0.418 | -0.449 | Continuous down | | | | | | | | | | |
| B7L74_00735 | UDP-3-O-[3-hydroxymyristoyl] N-acetylglucosamine | Lipopolysaccharide biosynthesis | -0.372 | -0.648 | -1.775 | -0.137 | -1.926 | -2.036 | -0.181 | -1.936 | -1.757 | 0.316 | -0.044 | Early down | | | | | | | | | Yes | |
| B7L74_00740 | hypothetical protein protein with ankyrin repeats/AnkB | Effector protein | -0.611 | -1.634 | 1.201 | 1.203 | -1.022 | -2.062 | 1.061 | 0.419 | -2.205 | 1.662 | 2.846 | Up in P67 | | Yes | | | | | | | | |
| B7L74_00745 | hypothetical protein protein with AnkB repeats/ | Effector protein | 1.341 | 1.466 | -0.098 | -0.661 | 0.931 | 0.623 | -0.253 | 0.168 | 1.266 | -0.066 | 1.588 | | | Yes | | | | | | | | |
| B7L74_00750 | hypothetical protein | | -0.082 | -1.362 | 0.325 | 0.973 | -2.526 | -2.605 | 0.403 | -2.253 | -1.410 | 0.928 | 1.537 | Early down | | | | | | | | | | |
| B7L74_00755 | preprotein translocase subunit SecA | Protein export/sec pathway | -0.031 | 1.185 | -3.010 | -1.474 | 1.039 | 0.923 | -0.908 | 1.169 | 0.528 | -2.459 | -2.865 | Early down | | | | | | | | | | |
| B7L74_00760 | 7%2C8-dihydro-8-oxoguanine-triphosphatase | DNA | 0.802 | 3.046 | -0.034 | -1.795 | 3.766 | 2.026 | 1.054 | 2.466 | 4.437 | -1.272 | -0.906 | | | | | | | | | | | |
| B7L74_00765 | DUF1658 domain-containing protein | | 0.035 | 0.837 | -0.034 | -1.810 | 2.635 | 1.620 | 0.443 | 1.486 | 2.542 | -1.283 | -0.912 | | | | | | | | | | | |
| B7L74_00770 | DUF1658 domain-containing protein | | 0.527 | 0.379 | -0.634 | -0.778 | 1.501 | 0.716 | 0.777 | 0.262 | 1.186 | -0.582 | -1.181 | | | | | | | | | | | |
| B7L74_00775 | cell division protein ZapD | Cell division | 0.421 | 0.554 | 2.906 | 0.994 | 0.666 | -0.190 | 1.135 | 0.648 | 0.188 | 1.695 | 1.167 | Early up | | | | | | | | | | |
| B7L74_00780 | NADPH-dependent 7-cyano-7-deazaguanine reductase | | -0.391 | 0.062 | 1.832 | 1.474 | -0.216 | -0.152 | 0.064 | -0.363 | -0.887 | 0.461 | 0.268 | | | | | | | | | | | |
| B7L74_00785 | dephospho-CoA kinase | Pantothenate and CoA biosynthesi | 2.200 | -0.036 | 0.468 | -0.347 | 3.329 | 3.898 | 0.977 | -0.349 | 3.014 | 1.224 | -0.130 | Early up | | | | | | | | | | |
| B7L74_00790 | prepilin peptidase, PilD | Pilus | 0.022 | 0.424 | -0.410 | 0.450 | -0.122 | -0.121 | 0.026 | -0.114 | 0.648 | 0.604 | 0.339 | | | | | | | | | | | |
| B7L74_00795 | type 4 fimbrial biosynthesis protein. pilC | Secretion system, Pilus | 3.364 | 1.796 | 1.406 | 0.514 | 3.994 | 4.001 | 2.204 | 2.418 | 3.495 | 2.762 | 2.724 | Early up | | | | | | | | | | |
| B7L74_00800 | type 4 pili biogenesis protein (nuleotide-binding protei | Pilus | -0.264 | 0.765 | 0.213 | -0.267 | -0.189 | -0.308 | 0.030 | 1.976 | -0.870 | -0.162 | 0.025 | | | | | | | | | | | |
| B7L74_00805 | pilus assembly protein PilA | Pilus | 0.077 | 1.374 | -0.034 | 1.094 | 1.972 | 0.970 | 2.132 | 3.445 | 2.995 | -0.192 | 1.241 | | | | | | | | | | | |
| B7L74_00810 | tRNA-Thr | tRNA | 0.961 | 2.327 | 2.440 | 2.611 | 1.866 | -0.052 | 2.738 | 1.889 | 1.590 | 2.356 | 2.466 | | | | | | | | | | | |
| B7L74_00815 | integrase | | -0.919 | 0.496 | 0.614 | 1.036 | -0.012 | -1.025 | -0.581 | 0.203 | -1.056 | 0.393 | 0.557 | | | | | | | | | | | |
| B7L74_00820 | hypothetical protein | | 1.099 | 1.074 | 1.281 | 1.417 | 0.571 | 2.020 | 1.300 | 3.975 | 0.665 | 1.823 | 0.722 | | | | | | | | | | | |
| B7L74_00825 | type I-F CRISPR-associated endoribonuclease Cas6/C | CRISPR/Cas | -2.432 | 1.739 | -0.136 | -2.395 | 1.412 | -0.812 | -0.093 | -0.019 | 0.175 | -1.893 | -0.845 | | | | | | | | | | | |
| B7L74_00830 | CRISPR-associated protein Csy3 | | -0.324 | 1.308 | 3.112 | 0.216 | 0.886 | 1.487 | -0.298 | 1.533 | 1.218 | 0.348 | 0.680 | | | | | | | | | | | |
| B7L74_00835 | type I-F CRISPR-associated protein Csy2 | CRISPR/Cas | 0.078 | 1.782 | -0.255 | -0.312 | 1.991 | 0.694 | 0.655 | 1.397 | 2.421 | 0.870 | 0.752 | | | | | | | | | | | |
| B7L74_00840 | type I-F CRISPR-associated protein Csy1 | CRISPR/Cas | 0.025 | 1.380 | -0.034 | -0.887 | 3.073 | 1.618 | 0.822 | 2.395 | 3.481 | -1.289 | -0.548 | | | | | | | | | | | |
| B7L74_00845 | hypothetical protein | | -1.304 | 1.067 | -0.439 | -0.433 | -1.567 | 0.036 | -1.168 | 1.396 | 2.053 | 0.036 | 0.119 | | | | | | | | | | | |
| B7L74_00850 | hypothetical protein | | 3.039 | 1.493 | 3.773 | 1.349 | 4.637 | 3.092 | 1.391 | 2.959 | 2.927 | 2.468 | 0.376 | | | | | | | | | | | |
| B7L74_00855 | hypothetical protein | | -0.419 | 1.362 | -0.609 | -0.184 | -0.994 | -0.080 | -1.209 | 1.214 | -0.058 | -0.626 | -0.637 | | | | | | | | | | | |
| B7L74_00860 | subtype I-F CRISPR-associated endonuclease Cas1 | CRISPR/Cas | -1.627 | 2.277 | -2.846 | -0.330 | 1.716 | -0.239 | 0.561 | 0.788 | 0.522 | -0.281 | -1.666 | | | | | | | | | | | |
| B7L74_00865 | hypothetical protein | | -0.476 | 1.285 | 0.058 | 0.333 | 0.170 | 0.448 | 0.645 | 1.616 | 0.922 | 0.913 | 0.745 | | | | | | | | | | | |
| B7L74_00870 | hypothetical protein | | -0.977 | 1.595 | 0.253 | -0.299 | 0.745 | 0.340 | 0.303 | 2.599 | 0.541 | 0.296 | 0.112 | | | | | | | | | | | |
| B7L74_00875 | enoyl-CoA hydratase | Polyketide biosynthesis | 3.937 | 4.793 | 2.976 | 0.133 | 3.126 | 2.644 | 3.185 | 6.705 | 6.036 | 2.093 | 0.128 | Late up | | | | | | | | | | |
| B7L74_00880 | acyl carrier protein | | 1.801 | 1.408 | -0.052 | 0.172 | 3.739 | 3.484 | -0.418 | 1.467 | 1.408 | 1.880 | 0.666 | Early up | | | | | | | | | | |
| B7L74_00885 | hypothetical protein | | 0.698 | 2.038 | 0.543 | 0.210 | 2.513 | 1.877 | 1.314 | 2.176 | 1.409 | 1.406 | 0.088 | | | | | | | | | | | |
| B7L74_00890 | hypothetical protein | | -1.809 | 1.076 | -3.004 | -2.965 | -0.271 | 1.194 | 2.293 | 1.712 | 0.010 | -1.061 | -3.867 | | | | | | | | | | | |
| B7L74_00895 | hypothetical membrane spanning protein | | 1.750 | 0.786 | 0.881 | -0.551 | -0.521 | -0.207 | 1.264 | 2.778 | 2.075 | 1.137 | 1.292 | Late up | | | | | | | | | | |
| B7L74_00900 | hypothetical protein | | 0.568 | 0.672 | 4.537 | 1.305 | 0.480 | 0.598 | 2.357 | 2.997 | 2.258 | 1.922 | 2.456 | Early up | | | | | | | | | | |
| B7L74_00905 | serine/threonine protein kinase | Effector protein | 4.120 | 1.333 | 2.770 | 2.074 | 5.999 | 5.216 | 3.291 | 2.829 | 5.078 | 3.348 | 2.337 | Early up | | Yes | | | | | | | | |
| B7L74_00910 | serine protease | | 2.507 | 1.123 | 1.062 | -0.268 | 0.670 | -0.107 | 1.851 | 3.185 | 1.212 | 2.112 | 1.740 | | | | | | | | | | | |
| B7L74_00915 | quaternary ammonium transporter | | -1.194 | -0.176 | -1.461 | -1.070 | -1.542 | -1.097 | -1.656 | -0.965 | -1.785 | -1.562 | -1.752 | | | | | | | | | Yes (Glycine betaine) | | |
| B7L74_00920 | glycine/betaine ABC transporter ATP-binding protein | ATP | -0.072 | -1.403 | -2.863 | 0.825 | -0.432 | -0.908 | 0.707 | -0.581 | 0.351 | -3.872 | -0.747 | | | | | | | | | Yes (Glycine betaine) | | |
| B7L74_00925 | anhydro-N-acetylmuramic acid kinase | | 0.262 | 0.434 | -1.336 | -0.767 | -1.234 | -1.470 | -0.248 | 0.196 | -1.520 | -0.576 | -0.837 | | | | | | | | | | | |
| B7L74_00930 | peptidase M23 | | 1.247 | -0.099 | -0.397 | -0.854 | 1.933 | 2.467 | -0.171 | -0.997 | 1.225 | -0.151 | -1.151 | | | | | | | | | | | |
| B7L74_00935 | tyrosine--tRNA ligase | Aminoacyl-tRNA synthase | -0.881 | 0.104 | -2.015 | -1.315 | -1.237 | -0.854 | -1.341 | -1.194 | -1.870 | -2.099 | -1.953 | Late down | | | | | | | | | | |
| B7L74_00940 | hypothetical protein | | 2.889 | 0.872 | 1.441 | -0.198 | 0.368 | -0.515 | 3.121 | 1.032 | 1.485 | 3.687 | 3.018 | Late up | | | | | | | | | | |
| B7L74_00945 | 16S ribosomal RNA | rRNA | 1.061 | 1.753 | 5.140 | 2.307 | 5.061 | 4.706 | 5.903 | 6.461 | 7.068 | 1.058 | 1.838 | Continuous up | | | | | | | | | | |
| B7L74_00950 | tRNA-Ile | tRNA | 0.062 | 0.842 | -0.034 | -1.409 | 1.973 | 0.975 | 0.469 | 1.491 | 1.135 | -1.268 | -0.904 | | | | | | | | | | | |
| B7L74_00955 | tRNA-Ala | tRNA | 0.040 | 0.838 | -0.034 | -1.126 | 1.973 | 1.620 | 0.447 | 1.487 | 2.547 | -1.280 | -0.911 | | | | | | | | | | | |
| B7L74_00960 | 23S ribosomal RNA | rRNA | -0.772 | 0.851 | 1.263 | 3.297 | -2.237 | -1.757 | 6.979 | 4.936 | 5.294 | -0.566 | -0.034 | Late up | | | | | | | | | | |
| B7L74_00965 | 5S ribosomal RNA (gene=rrf) | rRNA | 0.745 | 1.082 | 5.706 | 1.282 | 2.274 | 1.461 | 5.239 | 3.008 | 3.418 | 1.678 | 2.512 | Early up | | | | | | | | | | |
| B7L74_00970 | hypothetical protein | Effector protein | 2.164 | 0.587 | 1.557 | -0.116 | 1.449 | 0.805 | 2.531 | -7.453 | 2.270 | 2.602 | 3.650 | | | Yes | | | | | | | | |
| B7L74_00975 | hypothetical membrane associated protein | | 0.306 | 1.103 | 0.739 | -0.056 | -1.826 | 1.576 | -0.553 | -6.362 | -5.230 | 1.010 | 0.666 | Late down | | | | | | | | | | |
| B7L74_00980 | hypothetical protein | | 1.892 | 0.677 | 0.417 | -0.510 | 2.557 | 3.674 | 0.259 | 1.210 | 2.434 | 1.936 | 1.434 | | | | | | | | | | | |
| B7L74_00985 | membrane-associated protein | | 2.804 | 0.774 | 0.886 | 0.025 | 4.054 | 4.385 | 1.531 | 1.150 | 3.633 | 1.753 | 1.163 | Early up | | | | | | | | | | |
| B7L74_00990 | hypothetical protein | Hypothetical | 0.554 | 0.019 | -2.154 | -0.731 | 2.489 | 3.358 | -2.363 | 0.434 | 1.731 | 1.101 | -0.302 | Up in P21 | | | | | | | | | | |
| B7L74_00995 | CPBP family intramembrane metalloprotease domain-c | | 0.784 | 1.531 | -0.056 | 0.127 | 2.412 | 2.424 | 0.732 | 1.807 | 1.522 | 1.257 | 0.813 | Early up | | | | | | | | | | |
| B7L74_01000 | diacylglyceryl transferase | | 1.323 | 1.787 | 1.452 | 2.912 | 2.386 | 3.301 | 1.764 | 3.034 | 3.676 | 1.085 | 1.005 | Early up | | | | | | | | | | |
| B7L74_01005 | hypothetical protein, Hypothetical membrane associate | Membrane | 1.887 | 1.020 | 1.887 | -0.473 | 1.208 | 1.238 | 1.513 | 1.987 | 2.482 | 2.017 | 1.612 | | | | | | | | | | | |
| B7L74_01010 | L-serine ammonia-lyase | | -0.182 | -0.664 | 0.436 | 0.494 | -0.919 | -0.656 | 0.022 | -0.487 | -1.392 | -0.182 | 0.166 | | | | | | | | | | | |
| B7L74_01015 | MFS transporter | Transporter | 1.563 | 0.811 | 1.923 | 1.946 | 0.816 | 1.123 | 1.437 | 1.625 | 1.435 | 1.957 | 2.753 | Continuous up | | | | | | | | | | |
| B7L74_01020 | hypothetical protein | | 0.161 | 0.346 | 0.083 | -0.431 | 0.371 | 1.122 | 0.871 | 1.055 | 0.664 | -0.398 | 0.349 | | | | | | | | | | | |
| B7L74_01025 | DUF490 domain-containing protein | | -0.208 | -0.474 | -0.747 | -0.862 | -0.292 | 0.368 | -0.178 | -0.440 | -0.036 | -0.266 | -0.544 | | | | | | | | | Yes (ion channels/mechanosensitive channel) | | |
| B7L74_01030 | outer membrane protein assembly factor | Membrane | 1.514 | -2.937 | -1.345 | 0.994 | 0.453 | 1.247 | 0.611 | 0.790 | 0.380 | 0.817 | -0.140 | | | | | | | | | Yes (ion channels/mechanosensitive channel) | | |
| B7L74_01035 | type I pantothenate kinase | | 0.587 | -0.324 | -0.853 | 0.216 | 2.390 | 2.297 | -0.020 | -0.269 | 1.560 | 0.360 | -0.168 | | | | | | | | | | | |
| B7L74_01040 | aldose epimerase | | 0.027 | -0.170 | 0.247 | 0.997 | -0.432 | -0.159 | -0.150 | -0.678 | -0.698 | 0.502 | 0.352 | | | | | | | | | | | |
| B7L74_01045 | hypothetical proteinprotein=ankyrin repeat protein | Effector protein | -0.314 | 0.250 | 0.948 | 1.413 | -1.117 | -1.336 | 0.335 | 0.606 | -0.646 | 0.736 | 0.491 | | | Yes | | | | | | | | |
| B7L74_01050 | hypothetical protein | | -0.483 | -0.333 | 1.162 | -0.079 | -0.353 | 0.080 | 1.228 | 0.376 | 0.152 | 0.480 | 0.419 | | | | | | | | | | | |
| B7L74_01055 | glutamate--tRNA ligase | | -0.422 | -1.166 | -0.512 | -0.055 | -0.598 | -0.704 | -0.361 | -0.599 | -0.853 | -0.147 | -0.029 | | | | | | | | | | | |
| B7L74_01060 | hypothetical membrane spanning protein | | 1.974 | 0.741 | 1.842 | 0.412 | 0.489 | -0.134 | 1.539 | 1.053 | 0.507 | 1.850 | 1.248 | | | | | | | | | | | |
| B7L74_01065 | tRNA-Thr | tRNA | 2.753 | 0.358 | 0.840 | 0.348 | -0.363 | 0.036 | 2.122 | 1.535 | 0.832 | 2.213 | 1.997 | Early up | | | | | | | | | | |
| B7L74_01070 | hypothetical protein | | 0.631 | 0.012 | 0.103 | 0.474 | -0.277 | -1.186 | -0.257 | 1.673 | 0.569 | 1.366 | 0.663 | | | | | | | | | | | |
| B7L74_01075 | EamA family transporter | Transporter | -0.299 | 0.568 | -0.953 | 0.133 | -0.030 | 0.959 | -0.567 | 0.907 | 0.285 | 0.421 | -0.121 | | | | | | | | | | | |
| B7L74_01080 | hypothetical protein | DotF interacting protein | 0.558 | 0.136 | 0.472 | 0.589 | 0.092 | 0.920 | 0.817 | 1.159 | 1.246 | 1.382 | 1.319 | | | | | | | | | | | |
| B7L74_01085 | hypothetical protein | | 0.055 | 0.841 | -0.034 | -2.477 | 2.860 | 0.977 | 0.462 | 1.490 | 2.562 | -1.272 | -0.906 | | | | | | | | | | | |
| B7L74_01090 | hypothetical protein | | -0.226 | -0.979 | -0.042 | 0.501 | -0.877 | -0.952 | 0.442 | -1.434 | -1.093 | 0.334 | 1.311 | | | | | | | | | | | |
| B7L74_01095 | hypothetical protein | | -0.046 | -0.511 | -0.849 | -0.443 | 0.396 | 0.272 | -0.013 | -0.464 | 1.529 | -1.471 | -1.089 | | | | | | | | | | | |
| B7L74_01100 | hypothetical proteinintegral membrane protein | | -0.743 | -0.786 | -1.503 | 1.531 | -0.950 | -1.406 | -0.654 | -1.035 | -1.717 | -0.785 | -0.468 | | | | | | | | | | | |
| B7L74_01105 | NlpC-P60 family protein | | -1.872 | -0.752 | -3.066 | -1.246 | -0.845 | -0.482 | -2.245 | -1.647 | -2.335 | -2.398 | -2.393 | Early down | | | | | | | | | | |
| B7L74_01110 | hypothetical protein | | 0.839 | 0.157 | -0.886 | -0.074 | 1.494 | 0.990 | 0.455 | 0.267 | 1.457 | 0.396 | 0.626 | | | | | | | | | | | |
| B7L74_01115 | DUF1658 domain-containing protein | | 0.899 | 2.497 | -0.034 | 0.749 | 2.763 | 3.577 | 0.108 | 2.539 | 3.044 | -0.080 | -0.171 | | | | | | | | | | | |
| B7L74_01120 | hypothetical protein | | -0.258 | 1.053 | -2.394 | -1.866 | 0.758 | 1.411 | -0.420 | 0.435 | 0.506 | -1.903 | -1.341 | | | | | | | | | | | |
| B7L74_01125 | cytochrome d terminal oxidase subunit 1 | Respiration/cyt bd | 1.595 | 0.791 | 0.778 | -0.187 | 1.983 | 2.613 | 1.394 | 1.460 | 2.389 | 0.641 | 0.349 | Early up | | | | | | Yes | | | | |
| B7L74_01130 | cytochrome d ubiquinol oxidase subunit II | Respiration/cyt bd | -1.641 | 0.608 | -1.092 | -1.628 | 0.033 | 0.676 | -0.783 | -0.014 | 0.786 | -0.236 | -0.302 | | | | | | | Yes | | | | |
| B7L74_01135 | hypothetical protein | | -0.678 | 0.839 | -1.361 | -1.016 | -1.243 | -0.290 | -1.502 | -0.479 | -1.675 | -1.821 | -1.803 | | | | | | | | | | | |
| B7L74_01140 | phosphopantetheine-protein transferase | | -0.961 | -1.257 | -2.497 | -2.781 | 0.104 | -0.447 | -0.989 | -2.308 | -1.151 | -2.484 | -2.179 | | | | | | | | | | | |
| B7L74_01145 | tRNA-Tyr | tRNA | -0.242 | 0.606 | -1.468 | -0.483 | -0.532 | -0.334 | -1.458 | -0.380 | -1.467 | -1.876 | -1.784 | | | | | | | | | | | |
| B7L74_01150 | hypothetical protein | | 0.078 | 0.977 | -2.931 | -0.555 | -2.084 | -0.613 | -1.102 | -0.763 | -2.902 | -1.542 | -1.131 | | | | | | | | | | | |
| B7L74_01155 | tRNA-Gly | tRNA | -0.182 | -0.122 | -3.766 | -0.674 | -1.139 | -1.165 | -1.452 | -1.013 | -3.358 | -1.629 | -1.579 | Early down | | | | | | | | | | |
| B7L74_01160 | tRNA-Thr | tRNA | 0.095 | 1.572 | -2.742 | -0.795 | -0.805 | -0.333 | -0.116 | -0.704 | -4.185 | -1.738 | -0.891 | | | | | | | | | | | |
| B7L74_01165 | elongation factor Tu | | -0.333 | -0.705 | -1.713 | 0.433 | -1.033 | -1.198 | -0.099 | -1.513 | -2.124 | -0.697 | 0.999 | | | | | | | | | | | |
| B7L74_01170 | tRNA-Trp | tRNA | 0.298 | 0.300 | -1.769 | -0.597 | -0.689 | -0.727 | -1.068 | -1.389 | -2.753 | -1.905 | -1.526 | Down in P51 | | | | | | | | | | |
| B7L74_01175 | preprotein translocase subunit SecE | Protein export/sec pathway | -2.400 | -0.449 | -2.655 | -1.142 | -1.627 | -1.229 | -1.798 | -1.356 | -2.110 | -2.251 | -1.663 | Early down | | | | | | | | | | |
| B7L74_01180 | transcription termination/antitermination protein NusG | Transcription machinery | -1.645 | -0.311 | -3.717 | -1.176 | -1.642 | -1.123 | -2.177 | -1.668 | -2.480 | -1.964 | -0.941 | Early down | | | | | | | | | | |
| B7L74_01185 | 50S ribosomal protein L11 | Ribosomal proteins | -1.357 | -0.453 | -3.591 | -1.012 | -1.890 | -1.017 | -2.293 | -1.520 | -2.224 | -2.535 | -2.226 | Early down | | | | | | | | | | |
| B7L74_01190 | 50S ribosomal protein L1 | Ribosomal proteins | 1.844 | 3.973 | -0.860 | -2.282 | 0.939 | 0.234 | -0.167 | 1.667 | 2.697 | -1.686 | -1.350 | Early up | | | | | | | | | | |
| B7L74_01195 | 50S ribosomal protein L10 | Ribosomal proteins | -1.601 | -0.443 | -2.861 | -1.827 | -2.076 | -1.506 | -2.354 | -1.669 | -1.850 | -2.627 | -2.200 | Early down | | | | | | | | | | |
| B7L74_01200 | 50S ribosomal protein L7/L12 | Ribosomal proteins | -2.223 | -0.742 | -3.411 | -1.580 | -1.915 | -1.403 | -2.660 | -1.785 | -1.757 | -3.070 | -2.457 | Early down | | | | | | | | | | |
| B7L74_01205 | hypothetical protein | | -1.072 | 0.014 | -3.327 | -2.257 | -0.608 | -0.439 | -2.166 | -1.071 | -0.881 | -3.209 | -3.102 | Early down | | | | | | | | | | |
| B7L74_01210 | DNA-directed RNA polymerase subunit beta | RNA polymerase | -2.592 | -0.877 | -4.812 | -1.762 | -2.119 | -1.618 | -3.059 | -1.971 | -2.472 | -3.461 | -2.937 | Early down | | | | | | | | | | |
| B7L74_01215 | DNA-directed RNA polymerase subunit beta' | RNA polymerase | -1.529 | -0.929 | -4.083 | -1.636 | -1.195 | -0.755 | -1.940 | -2.277 | -1.984 | -2.155 | -1.005 | Early down | | | | | | | | | | |
| B7L74_01220 | 30S ribosomal protein S12 | Ribosomal proteins | -0.870 | -0.360 | -2.104 | 0.161 | -0.960 | -0.373 | -2.061 | -0.636 | -2.217 | -2.135 | -1.375 | Late down | | | | | | | | | | |
| B7L74_01225 | 30S ribosomal protein S7 | Ribosomal proteins | -1.700 | -0.218 | -2.223 | 0.075 | -1.273 | -0.603 | -1.574 | -0.925 | -1.754 | -1.932 | -1.334 | Early down | | | | | | | | | | |
| B7L74_01230 | elongation factor G | Translation factors | -1.143 | -0.235 | -3.036 | -0.268 | -0.815 | -0.620 | -1.773 | -0.964 | -1.626 | -2.265 | -0.986 | Early down | | | | | | | | | | |
| B7L74_01235 | elongation factor Tu | | -0.213 | -0.450 | -1.657 | 0.218 | -0.834 | -1.083 | -0.242 | -1.364 | -1.942 | -0.759 | 0.808 | | | | | | | | | | | |
| B7L74_01240 | 30S ribosomal protein S10 | Ribosomal proteins | -0.749 | -0.201 | -3.297 | -0.831 | -0.335 | -1.472 | -2.031 | -1.742 | -2.521 | -2.221 | -1.563 | Early down | | | | | | | | | | |
| B7L74_01245 | 50S ribosomal protein L3 | Ribosomal proteins | -1.717 | -0.676 | -3.567 | -1.378 | -1.318 | -1.300 | -2.107 | -1.827 | -2.657 | -2.705 | -2.349 | Early down | | | | | | | | | | |
| B7L74_01250 | 50S ribosomal protein L4 | Ribosomal proteins | -0.493 | -0.570 | -3.735 | -1.630 | -1.840 | -1.393 | -2.053 | -1.917 | -2.423 | -2.221 | -1.642 | Early down | | | | | | | | | | |
| B7L74_01255 | 50S ribosomal protein L23 | Ribosomal proteins | -3.351 | -3.057 | -3.926 | -3.671 | -1.269 | -0.377 | -3.459 | -2.408 | 0.058 | -5.180 | -3.986 | Early down | | | | | | | | | | |
| B7L74_01260 | 50S ribosomal protein L2 | Ribosomal proteins | -1.211 | -0.635 | -3.289 | -0.878 | -1.596 | -1.527 | -2.069 | -2.250 | -2.875 | -2.826 | -2.401 | Early down | | | | | | | | | | |
| B7L74_01265 | 30S ribosomal protein S19 | Ribosomal proteins | -1.450 | -0.854 | -3.094 | -0.992 | -1.716 | -1.708 | -1.870 | -2.270 | -2.261 | -2.547 | -1.376 | Early down | | | | | | | | | | |
| B7L74_01270 | 50S ribosomal protein L22 | Ribosomal proteins | -1.974 | -1.025 | -3.543 | -1.834 | -1.489 | -1.606 | -2.441 | -2.302 | -3.469 | -3.009 | -2.302 | Early down | | | | | | | | | | |
| B7L74_01275 | 30S ribosomal protein S3 | Ribosomal proteins | -4.839 | 0.602 | -3.440 | -4.397 | -1.816 | -2.207 | -3.295 | -3.480 | -0.576 | -5.145 | -5.063 | Early down | | | | | | | | | | |
| B7L74_01280 | 50S ribosomal protein L16 | Ribosomal proteins | -2.078 | -0.980 | -3.627 | -1.411 | -1.376 | -1.366 | -2.501 | -2.552 | -2.014 | -3.329 | -2.883 | Early down | | | | | | | | | | |
| B7L74_01285 | 50S ribosomal protein L29 | Ribosomal proteins | -1.622 | -0.785 | -3.175 | -1.521 | -1.180 | -1.495 | -2.408 | -2.583 | -2.211 | -2.490 | -2.002 | Early down | | | | | | | | | | |
| B7L74_01290 | 30S ribosomal protein S17 | Ribosomal proteins | -2.282 | -1.041 | -3.451 | -1.771 | -2.207 | -2.150 | -2.892 | -2.448 | -3.138 | -3.150 | -2.539 | Early down | | | | | | | | | | |
| B7L74_01295 | 50S ribosomal protein L14 | Ribosomal proteins | -2.420 | -0.953 | -4.093 | -1.876 | -2.217 | -2.261 | -2.608 | -2.782 | -3.839 | -2.854 | -2.007 | Continuous down | | | | | | | | | | |
| B7L74_01300 | 50S ribosomal protein L24 | Ribosomal proteins | -1.597 | -1.061 | -2.723 | -0.496 | -1.873 | -1.929 | -1.118 | -2.435 | -2.659 | -1.540 | 0.223 | Late down | | | | | | | | | | |
| B7L74_01305 | 50S ribosomal protein L5 | Ribosomal proteins | -0.574 | -1.372 | -1.405 | 0.612 | -0.501 | -0.546 | -0.595 | -2.215 | -1.540 | -0.270 | 0.971 | | | | | | | | | | | |
| B7L74_01310 | 30S ribosomal protein S14 | Ribosomal proteins | -0.368 | -1.260 | -1.809 | 0.557 | -0.290 | -0.948 | -0.264 | -2.381 | -1.897 | -0.272 | 0.962 | Late down | | | | | | | | | | |
| B7L74_01315 | 30S ribosomal protein S8 | Ribosomal proteins | 0.055 | 0.841 | -0.034 | -1.793 | 4.979 | 1.622 | 0.462 | 1.490 | 2.562 | -1.271 | -0.906 | Up in P16 | | | | | | | | | | |
| B7L74_01320 | 50S ribosomal protein L6 | Ribosomal proteins | -2.215 | -1.390 | -3.046 | -0.917 | -1.648 | -1.721 | -2.109 | -2.799 | -3.450 | -2.274 | -1.134 | Early down | | | | | | | | | | |
| B7L74_01325 | 50S ribosomal protein L18 | Ribosomal proteins | 0.052 | 0.840 | -0.034 | -1.796 | 2.655 | 1.622 | 1.889 | 1.489 | 2.559 | -1.273 | -0.907 | | | | | | | | | | | |
| B7L74_01330 | 30S ribosomal protein S5 | Ribosomal proteins | -2.732 | -1.497 | -4.521 | -1.295 | -2.068 | -1.910 | -2.694 | -2.555 | -3.482 | -2.960 | -2.213 | Early down | | | | | | | | | | |
| B7L74_01335 | 50S ribosomal protein L30 | Ribosomal proteins | -2.709 | -2.148 | -4.067 | -1.736 | -2.834 | -2.386 | -3.231 | -3.068 | -4.817 | -3.832 | -3.262 | Continuous down | | | | | | | | | | |

| | | | | | | | | | | | | | | | | | | | | | | | |
|---|---|---|---|---|---|---|---|---|---|---|---|---|---|---|---|---|---|---|---|---|---|---|---|
| B7L74_01340 | 50S ribosomal protein L15 | Ribosomal proteins | -0.980 | -1.025 | -2.996 | -2.623 | -1.342 | -1.397 | -2.603 | -2.438 | -2.737 | -3.176 | -3.142 | Early down | | | | | | | | | |
| B7L74_01345 | preprotein translocase subunit SecY | Translocase<br>Protein export/sec pathway | -2.159 | -1.202 | -3.806 | -2.205 | -1.962 | -1.969 | -3.037 | -2.252 | -3.545 | -3.311 | -2.946 | Continuous down | | | | | | | | | |
| B7L74_01350 | 50S ribosomal protein L36 | Ribosomal proteins | -2.837 | -1.978 | -4.245 | -2.954 | -3.041 | -2.175 | -3.303 | -3.534 | -3.393 | -4.059 | -3.396 | Early down | | | | | | | | | |
| B7L74_01355 | hypothetical protein | | -1.520 | -1.393 | -4.475 | -2.652 | -1.768 | -1.840 | -2.636 | -3.181 | -3.367 | -3.319 | -3.031 | Continuous down | | | | | | | | | |
| B7L74_01360 | hypothetical protein | | 0.035 | 0.837 | -0.034 | -1.810 | 2.635 | 1.620 | 0.443 | 1.486 | 2.542 | -1.283 | -0.912 | | | | | | | | | | |
| B7L74_01365 | 30S ribosomal protein S13 | Ribosomal proteins | -1.898 | -1.388 | -2.682 | -0.359 | -1.834 | -1.419 | -1.734 | -2.579 | -3.115 | -1.832 | -1.142 | Early down | | | | | | | | | |
| B7L74_01370 | 30S ribosomal protein S11 | Ribosomal proteins | -2.017 | -1.604 | -3.082 | -0.987 | -1.685 | -1.381 | -2.209 | -2.293 | -3.852 | -2.316 | -2.061 | Continuous down | | | | | | | | | |
| B7L74_01375 | 30S ribosomal protein S4 | Ribosomal proteins | -2.466 | -1.693 | -3.185 | -1.472 | -2.116 | -1.498 | -2.517 | -2.546 | -3.202 | -2.770 | -2.524 | Early down | | | | | | | | | |
| B7L74_01380 | DNA-directed RNA polymerase subunit alpha | RNA polymerase | -2.520 | -1.727 | -4.447 | -1.956 | -2.446 | -1.622 | -2.835 | -2.909 | -3.893 | -3.194 | -2.706 | Continuous down | | | | | | | | | |
| B7L74_01385 | 50S ribosomal protein L17 | Ribosomal proteins | -0.948 | -1.063 | -4.058 | -1.056 | -1.220 | -0.924 | -1.851 | -2.287 | -3.678 | -2.454 | -1.918 | Early down | | | | | | | | | |
| B7L74_01390 | glucose/galactose MFS transporter | Transporter | -1.758 | -1.357 | -1.412 | -1.103 | -0.719 | -0.262 | -1.593 | -0.933 | 0.346 | -0.045 | -0.716 | | | | | | | | | Yes (Sugar/Fucose) | |
| B7L74_01395 | hypothetical protein | | -0.582 | -1.497 | -4.817 | -1.187 | -0.368 | 1.032 | -2.562 | 1.457 | 1.771 | -1.865 | 0.506 | | | | | | | | | | |
| B7L74_01400 | hypothetical protein | | -0.540 | -2.006 | -0.416 | -1.017 | 0.870 | 0.786 | -2.189 | 2.070 | 2.407 | -0.291 | 0.789 | | | | | | | | | | |
| B7L74_01405 | hypothetical protein | | 1.157 | -0.207 | -0.034 | -0.697 | 2.592 | 1.894 | 0.604 | 0.830 | 1.872 | -1.881 | -0.893 | | | | | | | | | | |
| B7L74_01410 | trans-2-enoyl-CoA reductase | Effector protein | 0.536 | 0.619 | 0.312 | 1.253 | 0.698 | 0.490 | 0.595 | -0.204 | -0.428 | 0.445 | 0.321 | | | Yes | | | | | | | |
| B7L74_01415 | single-stranded DNA-binding protein | DNA | 0.551 | 0.935 | 0.520 | 1.189 | 0.616 | 0.355 | 0.068 | -0.369 | -1.335 | 0.083 | -0.120 | | | | | | | | | | |
| B7L74_01420 | MFS transporter | Transporter | -0.078 | 0.932 | -0.989 | 0.462 | 0.194 | 0.539 | -1.212 | 0.181 | -0.487 | -0.579 | -0.685 | | | | | | | | | | |
| B7L74_01425 | hypothetical protein | | 0.554 | 0.334 | -1.723 | -1.194 | -0.869 | -0.742 | -1.184 | -1.049 | -1.237 | -0.885 | -1.176 | | | | | | | | | | |
| B7L74_01430 | excinuclease ABC subunit A | | -0.644 | -0.525 | -0.338 | 0.093 | -0.747 | -0.689 | -0.817 | -1.205 | -1.971 | -0.849 | -0.973 | | | | | | | | | | |
| B7L74_01435 | hypothetical protein | | 0.681 | 1.730 | 3.957 | -1.273 | 1.891 | 1.434 | 2.206 | 1.745 | 2.150 | 0.681 | 0.570 | Early up | | | | | | | | | |
| B7L74_01440 | uroporphyrinogen decarboxylase | | 0.280 | 0.242 | 0.800 | 0.111 | -0.326 | -0.632 | 0.131 | -0.245 | -0.380 | -0.524 | -0.411 | | | | | | | | | | |
| B7L74_01445 | pyridine nucleotide-disulfide oxidoreductase | PV detoxification | -0.507 | 0.268 | -0.609 | -1.981 | 3.613 | 1.517 | 0.982 | 3.261 | 2.000 | -1.840 | -1.477 | | | | | | | | | | |
| B7L74_01450 | hypothetical exported protein | | 0.933 | 0.263 | 0.261 | 0.440 | 0.245 | 1.465 | 0.898 | 0.229 | 0.598 | 0.385 | 0.811 | | | | | | | | | | |
| B7L74_01455 | hypothetical proteinMarC family integral membrane pr | | 4.623 | 1.076 | 2.460 | 2.058 | 5.596 | 5.495 | 3.253 | 2.081 | 4.465 | 2.948 | 1.979 | Early up | | | | | | | | Yes (drug resistance) | |
| B7L74_01460 | amidinotransferase | | -0.192 | 0.211 | 0.141 | 1.581 | -1.101 | -0.119 | 0.649 | 1.587 | 0.410 | 0.675 | 0.947 | | | | | | | | | | |
| B7L74_01465 | DNA polymerase IV | DNA | 0.732 | 0.781 | 1.533 | 0.094 | 0.618 | 1.045 | 0.987 | 0.311 | -0.296 | 0.313 | 0.471 | | | | | | | | | | |
| B7L74_01470 | catalase | | 0.828 | 0.653 | 1.154 | 1.784 | 0.169 | 0.002 | 1.078 | 0.249 | -0.264 | 0.884 | 0.708 | | | | | | | | | | |
| B7L74_01475 | transcriptional regulator | | -0.685 | 0.526 | 2.241 | -0.379 | 0.223 | -0.100 | 2.584 | 0.788 | 1.634 | 2.158 | 2.002 | | | | | | | | | | |
| B7L74_01480 | hypothetical protein | | -0.748 | -0.448 | 1.028 | -0.281 | -0.338 | 0.054 | 0.143 | -0.798 | -1.048 | 1.498 | 1.906 | | | | | | | | | | |
| B7L74_01485 | hypothetical cytosolic protein | | 0.055 | 0.841 | -0.034 | -1.794 | 2.659 | 1.622 | 2.289 | 1.490 | 3.436 | 3.144 | 3.499 | | | | | | | | | | |
| B7L74_01490 | hypothetical protein | | -1.331 | -1.011 | -0.717 | -0.219 | -1.242 | -0.153 | -0.148 | -1.323 | -1.321 | 0.527 | 0.561 | | | | | | | | | | |
| B7L74_01495 | tRNA-Glu | tRNA | -0.054 | -0.773 | 0.180 | 0.621 | -0.208 | -0.480 | 0.315 | 0.747 | -0.452 | 0.113 | 0.386 | | | | | | | | | | |
| B7L74_01500 | polynucleotide adenylyltransferase PcnB | | -0.584 | -1.844 | -0.562 | 0.257 | -1.140 | -0.935 | -0.808 | -0.398 | 0.010 | 0.412 | 0.356 | | | | | | | | | | |
| B7L74_01505 | 2-amino-4-hydroxy-6-hydroxymethyldihydropteridine | Folate biosynthesis | 0.653 | 0.201 | 1.113 | 1.240 | 0.658 | 0.889 | 1.275 | -0.075 | 1.693 | 1.798 | 2.944 | Late up | | | | | | | | | |
| B7L74_01510 | phosphopantetheine adenylyltransferase | | 1.059 | -0.655 | 0.125 | 1.328 | 1.200 | 1.385 | 0.086 | -2.728 | 1.250 | 0.818 | 1.026 | | | | | | | | | | |
| B7L74_01515 | ferredoxin | | -0.490 | 0.597 | 1.051 | 2.518 | 0.702 | -0.431 | 0.542 | 1.208 | 2.129 | 2.640 | 3.991 | Continuous up | | | | | | | | | |
| B7L74_01520 | 50S ribosomal protein L33 | Ribosomal proteins | -1.090 | -0.168 | -2.564 | 2.877 | -0.331 | -0.259 | -0.203 | 0.324 | 0.191 | -0.191 | -0.272 | Up in P12 | | | | | | | | | |
| B7L74_01525 | 50S ribosomal protein L28 | Ribosomal proteins | -0.571 | -0.181 | -0.441 | 3.183 | 0.315 | 0.326 | 0.333 | -0.054 | -0.453 | 0.212 | 0.892 | Up in P12 | | | | | | | | | |
| B7L74_01530 | hypothetical protein | | 0.874 | 0.268 | 2.496 | -0.290 | 0.823 | 1.216 | 1.937 | 1.241 | 2.454 | 0.934 | 1.799 | | | | | | | | | | |
| B7L74_01535 | hypothetical protein | | 0.402 | 0.648 | 3.157 | -0.183 | 1.016 | 0.686 | 2.032 | 1.248 | 1.436 | -0.611 | -0.025 | | | | | | | | | | |
| B7L74_01540 | deoxyuridine 5'-triphosphate nucleotidohydrolase | | -0.230 | -0.124 | -0.608 | 1.135 | -0.559 | -0.186 | -0.459 | -0.155 | -0.656 | 0.269 | 0.173 | | | | | | | | | | |
| B7L74_01545 | phosphomannomutase/phosphoglucomutase | Glycolysis / Gluconeogenesis | 1.438 | -0.436 | -1.372 | -3.764 | 1.522 | 1.088 | 1.424 | 1.075 | 0.953 | -2.909 | -2.914 | Early down | | | | Yes | | | | | |
| B7L74_01550 | hypothetical protein | Effector protein | 4.226 | -0.636 | -1.518 | 1.021 | 6.115 | 6.495 | -0.956 | 2.809 | 1.145 | 1.066 | 1.564 | | | Yes | | | | | | | |
| B7L74_01555 | orotate phosphoribosyltransferase | | 0.542 | -0.628 | -1.511 | 0.932 | 2.380 | 4.645 | -0.949 | 2.120 | 1.152 | -0.131 | -1.425 | | | | | | | | | | |
| B7L74_01560 | exodeoxyribonuclease III | Base excision repair | -0.232 | 0.291 | -0.640 | -0.368 | -0.373 | -0.332 | -0.669 | -0.485 | -2.177 | -0.841 | -0.940 | Down in P51 | | | | | | | | | |
| B7L74_01565 | glutamate racemase | | 0.561 | 0.302 | 0.917 | 0.923 | 0.000 | -0.082 | 0.382 | -0.126 | -0.355 | 0.904 | 1.608 | | | | | | | | | | |
| B7L74_01570 | ribonuclease PH | | -1.132 | 1.993 | 1.962 | -1.376 | 1.957 | 1.886 | 0.205 | 1.297 | -0.603 | -1.346 | -0.774 | | | | | | | | | | |
| B7L74_01575 | YicC family protein | | -0.608 | -0.930 | -1.451 | -0.683 | -1.366 | -1.016 | -1.354 | -1.015 | -1.522 | -1.041 | -1.183 | Early down | | | | | | | | | |
| B7L74_01580 | guanylate kinase | | 0.035 | 0.836 | -0.034 | -1.811 | 2.634 | 3.290 | 0.442 | 2.766 | 3.484 | -1.283 | -0.912 | | | | | | | | | | |
| B7L74_01585 | DNA-directed RNA polymerase subunit omega | RNA polymerase | 1.157 | 0.910 | 0.212 | 0.328 | 0.580 | 0.149 | -0.418 | -0.326 | -2.490 | -0.804 | -0.809 | Down in P51 | | | | | | | | | |
| B7L74_01590 | GTP pyrophosphokinase, SpoT | Signal transduction<br>Purine metabolism | -1.164 | -0.310 | -2.046 | -0.300 | -0.960 | -0.932 | -1.983 | -1.272 | -3.049 | -2.294 | -2.324 | Early down | | | | | | | | | |
| B7L74_01595 | reactive intermediate/imine deaminase | | -1.959 | -0.112 | -2.939 | -0.638 | -1.404 | -1.295 | -2.244 | -1.467 | -3.375 | -2.623 | -2.451 | Early down | | | | | | | | | |
| B7L74_01600 | DNA helicase RecG | Homologous recombination | -1.400 | -0.332 | -3.333 | -1.020 | -0.888 | -0.783 | -2.110 | -1.445 | -3.193 | -2.340 | -1.774 | Early down | | | | | | | | | |
| B7L74_01605 | hypothetical protein | | -0.048 | 0.585 | -1.477 | 0.937 | -0.330 | -0.544 | -1.367 | -0.264 | -2.356 | -2.023 | -1.047 | Late down | | | | | | | | | |
| B7L74_01610 | hypothetical protein Outer membrane protein | Membrane | 0.750 | -0.006 | -0.530 | 1.734 | -0.265 | -2.244 | -0.260 | -0.405 | -1.889 | -1.811 | -0.467 | | | | | | | | | | |
| B7L74_01615 | molecular chaperone HtpG | Chaperones and folding catalysts | -2.864 | -4.656 | -3.304 | -1.924 | -3.875 | -3.556 | -2.310 | -5.417 | -4.967 | -2.332 | -2.204 | Continuous down | | | Yes | | | | | | |
| B7L74_01620 | hypothetical proteinouter membrane porin P1 | | 0.432 | 0.189 | 1.414 | 1.338 | 0.159 | -0.626 | 0.510 | -0.376 | -1.269 | 1.098 | 0.488 | | | | | | | | | Yes (General/Hypothetical) | |
| B7L74_01625 | bifunctional methylenetetrahydrofolate dehydrogenase | Carbon fixation pathways in proka | 0.774 | -0.202 | 0.971 | 2.278 | -0.098 | -0.597 | 0.617 | 0.362 | 0.979 | 1.940 | 1.981 | Early up | | | | | | | | | |
| B7L74_01630 | DUF1658 domain-containing protein | | 0.108 | 0.627 | -1.533 | -1.228 | -0.527 | -0.392 | -1.132 | -0.394 | -1.794 | -1.651 | -0.398 | | | | | | | | | | |
| B7L74_01635 | DUF1658 domain-containing protein | | 0.050 | -0.335 | 0.628 | -0.630 | -0.274 | -0.353 | 1.133 | -0.182 | -0.309 | -0.497 | 0.360 | | | | | | | | | | |
| B7L74_01640 | lytic transglycosylase | | -0.363 | -0.139 | 2.022 | 0.424 | -0.205 | -0.032 | 0.630 | 1.518 | 1.604 | 0.448 | 0.827 | | | | | | | | | | |
| B7L74_01645 | hydroxyacylglutathione hydrolase | Detox | 1.258 | -0.011 | 2.416 | -0.950 | 0.470 | 0.636 | 0.876 | -0.029 | -0.275 | 1.358 | 0.544 | | | | | | | | | | |
| B7L74_01650 | SAM-dependent methyltransferase | | -0.040 | -1.419 | 0.399 | 0.498 | -0.833 | -0.173 | 0.762 | 0.351 | -0.288 | 0.791 | 0.836 | | | | | | | | | | |
| B7L74_01655 | ribonuclease HI | | -0.700 | -2.191 | 1.061 | 0.326 | -1.246 | -1.269 | 0.191 | -0.753 | -0.849 | 0.108 | 0.506 | | | | | | | | | | |
| B7L74_01660 | DUF1658 domain-containing protein | CB specific protein family | -0.612 | -0.803 | -0.629 | -1.401 | -0.866 | -0.689 | -0.274 | -1.414 | -2.297 | -1.099 | -0.631 | Early down | | | | | | | | | |
| B7L74_01665 | DNA polymerase III subunit epsilon | DNA | 1.602 | 0.465 | 2.600 | -1.581 | 1.446 | 1.313 | 1.407 | 0.050 | 1.818 | 1.226 | 1.567 | | | | | | | | | | |
| B7L74_01670 | L%2CD-transpeptidase, Enhanced entry protein | Entry | 1.479 | 0.599 | 4.038 | -1.012 | 1.165 | 1.482 | 2.040 | 1.396 | 3.068 | 1.729 | 2.208 | | | | | | | | | | |
| B7L74_01675 | hypothetical protein | | 1.010 | 0.335 | 2.493 | -1.119 | 0.744 | 1.471 | 1.774 | 2.138 | 2.599 | 1.298 | 1.805 | | | | | | | | | | |
| B7L74_01680 | hypothetical protein IcmF | T4BSS, Secretion system VI | 0.606 | 1.514 | 3.659 | 1.603 | 2.328 | 2.967 | 3.300 | 4.423 | 3.723 | 2.284 | 2.030 | Early up | Yes | | | | | | | | |
| B7L74_01685 | hypothetical protein | | -0.201 | -1.493 | 0.362 | 0.660 | -1.074 | -0.200 | 0.773 | 0.827 | 0.725 | 0.740 | 0.856 | | | | | | | | | | |
| B7L74_01690 | Type IV secretion protein IcmH | T4BSS | 0.002 | 1.614 | 3.005 | 1.333 | 2.192 | 1.568 | 0.261 | -1.292 | 2.028 | 1.535 | 2.009 | | Yes | | | | | | | | |
| B7L74_01695 | hypothetical protein | | 0.486 | -0.103 | 0.811 | 1.050 | -0.272 | 0.396 | 1.081 | 0.859 | 1.450 | 0.364 | 0.068 | | | | | | | | | | |
| B7L74_01700 | acyl carrier protein | | 0.057 | 0.841 | -0.034 | -1.792 | 1.973 | 0.147 | 0.464 | 2.617 | 2.564 | -1.270 | -0.905 | | | | | | | | Yes | | |
| B7L74_01705 | acyl-CoA synthetase | | -0.340 | -0.020 | 0.635 | 0.232 | 0.291 | 0.576 | 0.022 | 0.623 | 0.703 | -0.100 | -0.088 | | | | | | | | | | |
| B7L74_01710 | phosphoribosylamine--glycine ligase | Purine metabolism | -1.281 | -1.501 | -1.198 | 0.816 | -1.429 | -1.399 | -0.784 | -1.738 | -2.266 | -1.080 | -1.065 | Late down | | | | | | | | | |
| B7L74_01715 | hypothetical protein | | -0.351 | -0.221 | -1.534 | -1.189 | 2.250 | 2.158 | -1.173 | 0.278 | 1.053 | -0.989 | -1.646 | | | | | | | | | | |
| B7L74_01720 | hypothetical protein | | 4.518 | 0.877 | 2.787 | -0.467 | 6.134 | 7.030 | 2.556 | 1.084 | 5.570 | 3.140 | 0.537 | Early up | | | | | | | | | |
| B7L74_01725 | phosphomethylpyrimidine synthase | | 0.172 | -1.712 | -1.251 | -0.510 | 0.667 | 0.440 | 0.196 | -0.448 | 0.857 | -0.787 | -0.686 | | | | | | | | | | |
| B7L74_01730 | glycine oxidase ThiO | | -1.929 | -1.119 | -1.755 | -1.077 | -1.939 | 0.853 | -1.827 | 0.183 | 0.954 | -2.894 | -2.169 | | | | | | | | | | |
| B7L74_01735 | thiamine biosynthesis protein ThiS | | 0.555 | 0.845 | -0.034 | -0.746 | 1.207 | 1.625 | -0.961 | 1.966 | 1.877 | -1.261 | -0.900 | | | | | | | | | | |
| B7L74_01740 | thiazole synthase | | -1.303 | -1.300 | -0.948 | -2.075 | -2.442 | -0.008 | -0.088 | 0.408 | 0.464 | -1.476 | -0.643 | | | | | | | | | | |
| B7L74_01745 | thiamine-phosphate diphosphorylase | Thiamine metabolism | -0.625 | 0.709 | 1.163 | -1.913 | 3.946 | 3.212 | 1.856 | 0.824 | 4.131 | -0.377 | 0.884 | Early up | | | | | | | | | |
| B7L74_01750 | acid phosphatase | | -0.434 | -0.476 | -0.120 | 0.685 | -0.202 | 0.452 | 0.833 | 0.790 | -0.193 | 0.125 | | | | | | | | | | | |
| B7L74_01755 | bifunctional phosphoribosylaminoimidazolecarboxami | | 0.459 | -0.832 | -0.612 | 1.957 | -0.797 | -0.462 | 0.435 | -0.537 | -0.881 | 1.019 | 1.201 | | | | | | | | | | |
| B7L74_01760 | Fis family transcriptional regulator | | -0.121 | -0.172 | -0.052 | 3.103 | 2.615 | 1.305 | -0.147 | 0.950 | 0.858 | -1.074 | -1.184 | | | | | | | | | | |
| B7L74_01765 | aminopeptidase N | | -1.573 | -0.939 | -1.106 | 0.535 | -1.342 | 0.320 | -0.229 | 0.315 | 0.263 | -0.708 | -0.242 | | | | | | | | | | |
| B7L74_01770 | hypothetical membrane associated protein | | 0.673 | 0.605 | 1.603 | 1.257 | 2.100 | 2.701 | 1.701 | 2.148 | 3.328 | 1.749 | 2.308 | Early up | | | | | | | | | |
| B7L74_01775 | hypothetical cytosolic protein | | -0.437 | -0.450 | 3.056 | 1.934 | -0.760 | 0.296 | 1.590 | 0.042 | 1.485 | 1.571 | 2.113 | | | | | | | | | | |
| B7L74_01780 | 6-phosphofructokinase | RNA degradation | 2.239 | 0.565 | 1.725 | 1.360 | 3.889 | 3.296 | 1.136 | -0.339 | 3.264 | 1.701 | 1.193 | Early up | | | | Yes | | | | | |
| B7L74_01785 | hypothetical protein | | 0.090 | 0.848 | -0.035 | -0.787 | 2.737 | 1.627 | 0.527 | 1.496 | 2.628 | 1.288 | -0.897 | | | | | | | | | | |
| B7L74_01790 | hypothetical protein | | 1.385 | 1.001 | 2.035 | 0.735 | 1.471 | 2.730 | 1.846 | 2.157 | 2.410 | 2.203 | 1.798 | Early up | | | | | | | | | |
| B7L74_01795 | hypothetical protein | | 1.077 | 0.896 | 3.955 | 0.108 | 4.081 | 1.956 | 2.358 | 2.485 | 5.084 | 1.709 | 2.232 | | | | | | | | | | |
| B7L74_01800 | hypothetical protein | | 1.923 | 1.566 | 3.071 | 0.699 | 1.583 | 1.991 | 2.712 | 2.799 | 2.936 | 2.738 | 2.615 | Late up | | | | | | | | | |
| B7L74_01805 | xylose isomerase | Pentose and glucuronate interconv | 0.561 | -0.024 | 1.255 | -0.025 | 2.342 | 2.536 | 1.072 | 1.210 | 2.116 | 1.218 | 1.152 | Early up | | | | | | | | | |
| B7L74_01810 | xylulokinase | | 1.243 | -1.890 | -0.027 | 1.111 | 0.878 | 2.581 | 0.059 | -0.262 | 0.570 | -0.020 | 0.141 | | | | | | | | | | |
| B7L74_01815 | MFS transporter, D-xylose-proton symporter | Transporter | 0.218 | -1.296 | 0.232 | 0.324 | 0.379 | 1.856 | 0.049 | -0.932 | 0.076 | -0.079 | -0.090 | | | | | | | | | | |
| B7L74_01820 | phosphoglycolate phosphatase%2C bacterial | | 0.453 | -0.119 | -1.961 | 0.417 | -0.521 | -0.811 | -0.846 | -1.469 | -1.438 | -1.152 | -0.894 | | | | | | | | | | |
| B7L74_01825 | bifunctional 3-demethylubiquinone 3-O-methyltransfer | Ubiquinone and other terpenoid-qu | -0.289 | -0.429 | -2.504 | -0.547 | -1.084 | -1.157 | -1.172 | -2.200 | -2.891 | -1.359 | -1.409 | Early down | | | | | | | | | |
| B7L74_01830 | histidine kinase | | 4.417 | 2.926 | 3.135 | 3.047 | 2.915 | 0.629 | 3.646 | 1.248 | 2.730 | 1.978 | 3.253 | | | | | | | | | | |
| B7L74_01835 | cation transporter | Transporter | -0.747 | -0.709 | -0.897 | 0.011 | -1.229 | -1.142 | -1.251 | -1.834 | -2.731 | -1.726 | -1.453 | Late down | | | | | | | | Yes (Ions/cobalt-zinc-cadmium) | |
| B7L74_01840 | protease SohB | | 2.133 | -0.029 | 0.368 | 2.871 | 2.865 | 3.372 | 0.774 | -1.358 | 1.765 | 0.998 | 1.925 | | | | | | | | | | |
| B7L74_01845 | amino acid permease | Transporter | 0.732 | 0.840 | 0.306 | -0.531 | 1.886 | 0.984 | 0.027 | -0.029 | -0.419 | -2.370 | -1.677 | Variable | | | | | | | | Yes (Amino acid/Aspartate, Glutamate) | |
| B7L74_01850 | hypothetical protein | Effector protein | -1.384 | -0.285 | -3.084 | -2.290 | -1.038 | -0.635 | -2.036 | -1.180 | -1.227 | -3.052 | -3.115 | Early down | | Yes | | | | | | | |
| B7L74_01855 | hypothetical protein | | 0.244 | 0.532 | -1.703 | -2.512 | -0.142 | 0.279 | -1.119 | -0.062 | -1.228 | -2.150 | -1.854 | Early down | | | | | | | | | |
| B7L74_01860 | hypothetical protein | | 1.046 | -0.644 | -1.535 | -0.278 | 2.635 | 2.710 | 1.001 | 3.241 | 4.236 | 0.317 | 0.455 | Early up | | | | | | | | | |
| B7L74_01865 | hypothetical protein | | 2.496 | -1.136 | 2.858 | 1.478 | 3.574 | 5.117 | 2.948 | 5.251 | 6.169 | 2.053 | 2.666 | Continuous up | | | | | | | | | |
| B7L74_01870 | amidohydrolase | | -0.506 | -0.774 | 0.560 | 1.805 | -0.469 | 0.157 | 0.128 | 0.814 | 1.157 | 0.520 | 0.581 | | | | | | | | | | |
| B7L74_01875 | peptidase | | -0.932 | -0.157 | -1.035 | -2.782 | 3.189 | 1.620 | 0.937 | 1.624 | 0.878 | -0.700 | -1.902 | | | | | | | | | | |
| B7L74_01880 | MFS transporter | Transporter | 4.702 | 1.375 | 2.727 | 2.137 | 5.036 | 5.706 | 3.846 | 3.241 | 5.790 | 3.340 | 3.218 | Continuous up | | | | | | | | Yes (Ions/Phosphate) | |
| B7L74_01885 | two-component sensor histidine kinase, PhoR | two component system PhoB-PhoR | -1.171 | -0.936 | -1.508 | -0.117 | -1.519 | -1.353 | -1.269 | -0.152 | -0.698 | -0.787 | -0.918 | | | | | | | | | | |
| B7L74_01890 | DNA-binding response regulator, PhoB | Signal transductiontwo component | 0.327 | 3.633 | 3.468 | 3.693 | 5.051 | 3.693 | 3.863 | 1.926 | 4.937 | 2.329 | 1.276 | Continuous up | | | | | | | | | |
| B7L74_01895 | glucose sorbosone dehydrogenase | | -0.016 | -0.643 | 0.609 | 1.145 | -1.181 | -0.278 | 0.292 | 0.073 | 0.324 | 0.554 | 0.568 | | | | | | | | | | |
| B7L74_01900 | hypothetical protein | | 2.647 | 1.404 | 0.940 | 1.994 | 1.922 | -0.302 | 2.047 | 0.285 | 1.205 | 2.397 | 2.738 | | | | | | | | | | |
| B7L74_01905 | hypothetical membrane associated protein | | 0.882 | 0.708 | 2.098 | 0.893 | 0.764 | 0.364 | 2.169 | 1.118 | 1.776 | 1.633 | 2.160 | Early up | | | | | | | | | |
| B7L74_01910 | MFS transporter | Transporter | 5.147 | 0.921 | 1.972 | 0.496 | 6.140 | 5.937 | 3.470 | 0.883 | 4.985 | 2.369 | 1.161 | Early up | | | | | | | | Yes (drug resistance) | |
| B7L74_01915 | cell filamentation protein Fic | Effector protein | -0.437 | 0.164 | -0.375 | -0.784 | 1.112 | 1.332 | -0.681 | 0.486 | 1.463 | -0.957 | -0.981 | | | Yes | | | | | | | |
| B7L74_01920 | dehydratase | | 1.115 | -2.148 | 0.911 | -0.049 | 1.511 | 2.027 | 0.670 | 2.210 | 2.581 | 0.008 | 0.701 | | | | | | | | | | |
| B7L74_01925 | CoA ester lyase | Two-component system | 0.052 | 1.489 | 3.425 | 2.609 | 1.973 | 4.359 | 2.029 | 3.560 | 4.369 | 2.220 | 2.658 | Early up | | | | | | | | | |
| B7L74_01930 | hypothetical protein | | 2.446 | 0.010 | -0.678 | 2.120 | 3.111 | 2.099 | 0.469 | 0.606 | 2.324 | 0.794 | 0.938 | | | | | | | | | | |
| B7L74_01935 | hypothetical protein | | 1.365 | -0.469 | -0.678 | 1.754 | 1.265 | 0.621 | 0.458 | 0.432 | 1.847 | 0.901 | 1.352 | | | | | | | | | | |
| B7L74_01940 | hypothetical protein | | 0.585 | -1.985 | -0.072 | 0.926 | -0.156 | 0.903 | 0.667 | 1.389 | 2.193 | 0.177 | 1.650 | | | | | | | | | | |
| B7L74_01945 | lipoprotein | | -0.663 | -0.681 | -1.505 | 1.712 | -0.293 | 0.041 | -1.219 | 0.631 | 0.825 | -1.421 | -1.277 | | | | | | | | | | |
| B7L74_01950 | nucleoside transporter | | 0.203 | -0.851 | -1.260 | 0.412 | -0.726 | -0.709 | 0.144 | -0.998 | -0.774 | -0.068 | 0.081 | | | | | | | | | | |
| B7L74_01955 | hypothetical protein | | -1.665 | -1.010 | -1.503 | 0.717 | -1.117 | -0.051 | -1.685 | 0.263 | 0.434 | -0.649 | -1.095 | | | | | | | | | | |
| B7L74_01960 | hypothetical proteinprotein=glutamate-rich protein | | -0.206 | -0.775 | -0.612 | 0.184 | -0.636 | -0.419 | -0.284 | -0.669 | -0.819 | -0.710 | -1.293 | | | | | | | | | | |
| B7L74_01965 | hypothetical protein | | 0.264 | 1.208 | 2.827 | 0.421 | 1.615 | 1.362 | 1.271 | 1.188 | 3.041 | 0.715 | 0.148 | Early up | | | | | | | | | |
| B7L74_01970 | tRNA-Pro | tRNA | -0.693 | -0.707 | 0.268 | 0.665 | -2.403 | -1.145 | 0.589 | 0.310 | -1.539 | -0.431 | -0.106 | | | | | | | | | | |
| B7L74_01975 | octaprenyl diphosphate synthase | | -0.753 | -1.194 | -0.924 | 0.384 | -0.931 | -0.952 | -0.308 | -0.106 | 0.485 | -0.130 | -0.094 | | | | | | | | | | |
| B7L74_01980 | DNA-3-methyladenine glycosylase I | | -0.252 | -0.450 | -0.004 | 0.963 | -0.399 | 0.902 | 0.149 | 1.194 | 0.761 | 0.424 | 0.350 | | | | | | | | | | |
| B7L74_01985 | hypothetical protein | | 2.176 | 2.149 | 4.003 | 1.364 | 1.968 | 2.713 | 4.425 | 3.390 | 2.829 | 2.296 | 1.625 | Early up | | | | | | | | | |
| B7L74_01990 | IS110 family transposase | Replication and repair | 1.376 | 0.844 | 2.370 | 1.806 | 1.260 | 4.514 | 2.899 | 3.159 | 3.792 | 2.846 | 2.115 | Early up | | | | | | | | | |
| B7L74_01995 | hypothetical protein | | 0.090 | 0.848 | -0.035 | -0.788 | 2.737 | 1.627 | 0.527 | 1.496 | 3.555 | -0.331 | 0.097 | | | | | | | | | | |
| B7L74_02000 | hypothetical protein | | -0.420 | -2.338 | -0.878 | -1.079 | -0.204 | 0.666 | 0.387 | -0.089 | 1.106 | 1.143 | 0.650 | | | | | | | | | | |
| B7L74_02005 | 50S ribosomal protein L21 | Ribosomal proteins | -0.315 | -0.148 | -0.882 | 2.959 | -0.280 | -0.174 | 0.126 | -0.508 | -1.173 | 0.188 | 0.887 | Up in P12 | | | | | | | | | |
| B7L74_02010 | 50S ribosomal protein L27 | Ribosomal proteins | 0.814 | 0.842 | 0.966 | -1.789 | 1.973 | 1.623 | 2.789 | 1.491 | 2.568 | 0.760 | 1.665 | | | | | | | | | | |
| B7L74_02015 | GTPase ObgE | | 0.481 | -0.329 | -0.256 | 1.289 | -0.351 | -0.177 | 0.495 | -0.080 | -0.830 | 0.760 | 1.263 | | | | | | | | | | |
| B7L74_02020 | hypothetical protein | Effector protein | 5.745 | 2.353 | 2.610 | 0.614 | 7.430 | 6.707 | 3.530 | 1.213 | 6.006 | 2.952 | 1.217 | Continuous up | | Yes | | | | | | | |
| B7L74_02025 | 30S ribosomal protein S20 | Ribosomal proteins | -1.930 | -0.973 | 0.492 | 2.659 | -0.550 | -0.209 | 0.082 | 0.442 | 0.054 | 0.786 | 0.950 | Up in P12 | | | | | | | | | |
| B7L74_02030 | lipid II flippase MurJ | | -0.593 | -1.225 | -0.961 | -0.174 | -0.536 | -0.498 | -0.304 | -1.175 | -0.743 | 0.005 | 0.081 | | | | | | | | | | |
| B7L74_02035 | bifunctional riboflavin kinase/FMN adenylyltransferase | Riboflavin metabolism | 3.262 | 3.706 | 4.656 | -1.712 | 3.299 | 4.342 | 0.770 | 2.496 | 3.970 | 1.380 | 1.392 | Early up | | | | | | | | | |
| B7L74_02040 | hypothetical protein | | -0.449 | 0.070 | 0.456 | 1.086 | 0.968 | 1.891 | 0.309 | 2.140 | 2.068 | 1.453 | 0.883 | Late up | | | | | | | | | |
| B7L74_02045 | hypothetical protein | | 1.307 | -0.125 | 0.674 | -1.102 | 2.652 | 2.781 | 0.785 | 0.494 | 1.762 | 0.417 | 0.177 | | | | | | | | | | |
| B7L74_02050 | lipoprotein | | -0.141 | -0.799 | 0.522 | 2.345 | -0.364 | -0.634 | 1.337 | -1.943 | -1.406 | 1.413 | 1.171 | | | | | | | | | | |
| B7L74_02055 | isoleucine--tRNA ligase | | -1.639 | -1.534 | -0.490 | -0.570 | -1.173 | -0.526 | 0.024 | 0.163 | -0.224 | -0.617 | -0.280 | | | | | | | | | | |
| B7L74_02060 | signal peptidase II | Protein export/signal peptidase | 3.137 | 0.166 | 1.681 | -0.134 | 6.193 | 4.590 | 2.359 | 1.573 | 5.481 | 2.412 | 0.055 | Early up | | | | | | | | | |
| B7L74_02065 | hypothetical membrane spanning protein | | -0.429 | 0.577 | 2.437 | 0.996 | -0.019 | 0.085 | 1.152 | 1.290 | 1.445 | 1.031 | 1.283 | | | | | | | | | | |
| B7L74_02070 | hypothetical protein | | -0.292 | 0.439 | 1.600 | 2.462 | 0.198 | -0.451 | 0.840 | 2.165 | 1.057 | 1.185 | 1.467 | | | | | | | | | | |
| B7L74_02075 | hypothetical protein | | 3.201 | 2.228 | 5.498 | -1.992 | 3.090 | 2.577 | 1.712 | 1.378 | 1.795 | 3.161 | 1.959 | | | | | | | | | | |
| B7L74_02080 | acetyltransferase | | 0.558 | 2.040 | 0.966 | 3.042 | 1.972 | 1.626 | 0.493 | 4.666 | 4.280 | 4.681 | 0.446 | Late up | | | | | | | | | |
| B7L74_02085 | acetyltransferase | | 0.556 | 0.845 | -0.034 | 2.595 | 1.972 | 0.147 | 0.485 | 3.193 | 3.253 | 3.342 | 0.706 | | | | | | | | | | |
| B7L74_02090 | aminoglycoside resistance protein | | -1.077 | -0.744 | -3.367 | -0.157 | 0.031 | 0.211 | -0.145 | 1.215 | 2.047 | 1.213 | 0.615 | | | | | | | | | | |
| B7L74_02095 | hypothetical protein | | 0.549 | 0.544 | -1.855 | 0.653 | 1.196 | 2.039 | 2.114 | 3.019 | 3.797 | 1.692 | 1.348 | Late up | | | | | | | | | |
| B7L74_02100 | hypothetical protein | | 1.471 | 2.855 | 3.334 | 2.211 | 2.142 | 2.945 | 0.343 | 2.709 | 1.903 | 1.304 | 1.765 | | | | | | | | | | |
| B7L74_02105 | hypothetical protein | | -0.193 | 0.450 | -1.366 | -1.185 | 0.148 | 1.806 | 0.769 | 1.301 | 2.034 | -0.774 | -0.440 | | | | | | | | | | |
| B7L74_02110 | hypothetical membrane spanning protein | Effector protein | -0.373 | 0.670 | -1.639 | -2.105 | 1.932 | 1.279 | -1.225 | 0.275 | 1.417 | -1.328 | -2.373 | | | Yes | | | | | | | |
| B7L74_02115 | type IV pilin, pilE | Secretion system, Pilus | 0.857 | 1.372 | 3.283 | 0.741 | -0.543 | -0.230 | 1.899 | 3.600 | 4.419 | 1.957 | 2.464 | Late up | | | | | | | | | |
| B7L74_02120 | tRNA-Lys | tRNA | 0.887 | 1.213 | 3.879 | 0.327 | 1.631 | 0.817 | 2.311 | 1.812 | 1.483 | 0.369 | 0.469 | Early up | | | | | | | | | |
| B7L74_02125 | hypothetical protein | | -1.233 | 1.380 | 1.641 | 0.565 | 1.274 | 1.676 | 0.220 | 3.203 | 2.151 | 0.768 | -0.305 | Late up | | | | | | | | | |
| B7L74_02130 | hypothetical protein | | 0.063 | -0.199 | -0.034 | -1.119 | 2.063 | 3.566 | 0.849 | 1.961 | 1.879 | -0.839 | -0.903 | | | | | | | | | | |
| B7L74_02135 | tRNA-specific adenosine deaminase | Purine metabolism | 0.268 | 0.083 | 2.228 | -1.280 | 0.327 | 1.453 | 1.622 | 1.943 | 2.295 | 1.030 | 1.318 | Late up | | | | | | | | | |
| B7L74_02140 | hypothetical protein | | 0.417 | 0.469 | 2.418 | 0.688 | -0.932 | 0.702 | 1.334 | 0.885 | 1.948 | 1.748 | 2.187 | Late up | | | | | | | | | |
| B7L74_02145 | hypothetical protein | | 0.092 | -0.350 | 0.629 | -0.510 | 0.804 | 0.966 | 0.630 | 0.807 | 0.766 | 0.084 | 0.091 | | | | | | | | | | |
| B7L74_02150 | 4-amino-4-deoxychorismate lyase | Cysteine and methionine metabolis | 3.380 | 0.661 | 2.375 | 3.266 | 4.849 | 5.797 | 2.209 | 3.597 | 5.460 | 3.340 | 2.850 | Early up | | | | | | | | | |
| B7L74_02155 | polysaccharide deacetylase | Metabolism | 3.257 | 1.420 | 3.113 | 1.704 | 3.045 | 0.502 | 4.056 | 5.064 | 3.840 | 3.157 | 2.494 | Continuous up | | | | | | | | | |
| B7L74_02160 | siderophore biosynthesis protein%2C IucA/IucC famil | | -0.108 | -0.149 | 0.238 | -0.060 | 0.616 | 0.929 | -0.098 | 1.631 | 2.949 | -0.101 | 0.940 | Late up | | | | | | | | | |
| B7L74_02165 | aspartate 1-decarboxylase | beta-Alanine metabolism | -0.247 | 0.491 | 2.568 | 1.072 | 1.361 | 0.610 | 3.323 | 1.470 | 3.298 | 0.876 | 1.506 | Early up | | | | | | | | | |
| B7L74_02170 | pantothenate synthetase | beta-Alanine metabolism | 2.959 | -1.426 | -2.836 | -4.209 | 0.217 | 0.315 | -1.667 | -1.310 | 0.382 | -4.067 | -3.704 | Early down | | | | | | | | | |
| B7L74_02175 | 3-methyl-2-oxobutanoate hydroxymethyltransferase | | 0.185 | -0.577 | -0.766 | 1.145 | -0.719 | 0.282 | -0.324 | 0.380 | -0.064 | -0.138 | -0.088 | | | | | | | | | | |
| B7L74_02180 | hypothetical protein(cirB in Table 1 of RightonQ pape | Effector protein | 4.392 | 1.109 | 2.353 | 0.353 | 5.091 | 4.682 | 2.316 | 2.276 | 5.701 | 3.408 | 1.348 | Early up | | Yes | | | | | | | |
| B7L74_02185 | amino acid permease | Transporter | -0.035 | 0.049 | 1.508 | -0.647 | 0.226 | 0.148 | 0.923 | 0.376 | 0.582 | 1.155 | 0.934 | | | | | | | | | | |
| B7L74_02190 | hypothetical protein | | -2.415 | -0.463 | -3.352 | -3.901 | 1.100 | 2.925 | -1.575 | -0.523 | 0.924 | -4.618 | -4.240 | Variable | | | | | | | | | |
| B7L74_02195 | peptide chain release factor 2 | Translation factors | -7.000 | -6.517 | -6.336 | -1.003 | -3.011 | -5.351 | -6.538 | -5.482 | -5.173 | -6.837 | -6.738 | Early down | | | | | | | | | |
| B7L74_02200 | aminoacyl-tRNA hydrolase | | -0.738 | -0.211 | -0.034 | -1.733 | 1.323 | 0.957 | -0.232 | 0.825 | 1.130 | -1.873 | -0.887 | | | | | | | | | | |
| B7L74_02205 | lysine--tRNA ligase | | -0.462 | -0.858 | -0.489 | 0.597 | -1.211 | -0.786 | -0.114 | -0.915 | -0.793 | 0.484 | 0.543 | | | | | | | | | | |
| B7L74_02210 | glycine cleavage system protein T | | -7.816 | -7.058 | -7.941 | -1.905 | -2.169 | -2.263 | -6.293 | -3.182 | -2.105 | -8.235 | -8.803 | | | | | | | | | | |
| B7L74_02215 | MFS transporter | Transporter | 1.382 | 0.507 | 1.730 | 0.588 | -0.098 | 0.572 | 1.332 | 1.366 | 0.978 | 1.589 | 1.451 | Early up | | | | | | | | Yes (MFS) | |
| B7L74_02220 | mechanosensitive ion channel protein MscS | | -0.670 | -0.857 | -1.081 | -0.703 | -0.613 | -0.015 | -0.470 | -1.019 | 0.437 | -0.323 | -0.759 | | | | | | | | | Yes (ion channels/mechanosensitive channel) | |
| B7L74_02225 | DUF1658 domain-containing protein | | 0.676 | -0.015 | 2.645 | -1.149 | 0.423 | 0.612 | 1.672 | 0.293 | 0.563 | 0.026 | 0.260 | | | | | | | | | | |
| B7L74_02230 | hypothetical protein | | 0.877 | 0.242 | 2.218 | -0.110 | 0.637 | 0.595 | 1.302 | -0.505 | -0.891 | 0.678 | 0.655 | | | | | | | | | | |
| B7L74_02235 | hypothetical protein | | 1.589 | 2.434 | 1.966 | 2.007 | 0.295 | 2.909 | 3.108 | 3.968 | 4.755 | 2.624 | 1.537 | Late up | | | | | | | | | |
| B7L74_02240 | hypothetical protein | | 6.993 | 3.617 | -14.640 | 4.143 | 8.191 | 8.231 | 3.984 | 6.379 | 8.872 | 5.942 | 4.717 | Variable | | | | | | | | | |
| B7L74_02245 | hypothetical membrane spanning protein | | 1.518 | -0.317 | 1.095 | -1.640 | 2.263 | 4.273 | 1.795 | 1.874 | 3.360 | 0.530 | -2.090 | Up in P21 | | | | | | | | | |
| B7L74_02250 | hypothetical protein | | 4.385 | 0.727 | 2.373 | 2.309 | 5.371 | 4.553 | 3.382 | -0.460 | 3.238 | 2.592 | 3.101 | Early up | | | | | | | | | |
| B7L74_02255 | hypothetical protein | | 0.837 | -1.200 | 0.476 | 0.893 | 0.529 | 0.890 | 0.099 | -1.025 | -0.250 | 0.963 | 1.821 | | | | | | | | | | |
| B7L74_02260 | hypothetical protein | | -0.034 | 1.381 | 2.691 | 2.473 | 3.560 | 1.032 | 1.816 | 1.596 | 2.375 | -1.844 | 1.393 | | | | | | | | | | |
| B7L74_02265 | methylated-DNA--[protein]-cysteine S-methyltransfera | | -0.215 | -1.165 | -0.942 | 0.320 | 0.212 | 0.446 | -0.842 | 0.177 | 0.877 | -0.703 | -0.162 | | | | | | | | | | |
| B7L74_02270 | 50S ribosomal protein L19 | Ribosomal proteins | -0.167 | -0.514 | -0.559 | 1.396 | -0.604 | -0.634 | -0.019 | 0.139 | 0.059 | 0.646 | 0.639 | | | | | | | | | | |
| B7L74_02275 | tRNA (guanosine(37)-N1)-methyltransferase TrmD | Transfer RNA biogenesis | 1.603 | 2.883 | 2.287 | 1.334 | 3.597 | 5.221 | 2.681 | 4.082 | 4.150 | 0.876 | -0.412 | Up in P21 | | | | | | | | | |
| B7L74_02280 | ribosome maturation factor RimM | Ribosome biogenesis | -0.214 | 0.084 | 2.393 | 2.077 | -0.120 | 0.028 | 0.263 | 1.243 | 1.586 | 0.900 | 1.053 | Early up | | | | | | | | | |
| B7L74_02285 | 30S ribosomal protein S16 | Ribosomal proteins | 0.429 | 0.133 | -0.401 | 2.863 | -0.245 | 0.101 | 0.204 | 0.434 | -0.419 | 0.568 | 0.845 | Up in P12 | | | | | | | | | |
| B7L74_02290 | hypothetical protein | | -1.811 | -0.513 | -1.476 | -0.296 | -0.090 | -0.330 | -1.505 | 0.092 | -0.136 | 0.006 | -0.777 | | | | | | | | | | |
| B7L74_02295 | hypothetical protein(ankF in Table 1 of RightonQ pape | Effector protein | -0.278 | -0.826 | 0.760 | 2.021 | -0.947 | -1.196 | 0.334 | -1.497 | 0.113 | 1.413 | 1.951 | Early up | | Yes | | | | | | | |
| B7L74_02300 | hypothetical protein | | 2.487 | 0.846 | 2.215 | -1.923 | 2.641 | 2.704 | 1.984 | 1.225 | 1.473 | 2.268 | 0.728 | | | | | | | | | | |
| B7L74_02305 | 3'%2C5'-cyclic-nucleotide phosphodiesterase | | 5.193 | 1.425 | 2.877 | 0.569 | 6.266 | 6.351 | 3.132 | 2.036 | 5.651 | 3.649 | 2.040 | Early up | | | | | | | | | |
| B7L74_02310 | signal recognition particle protein | | -1.846 | -0.015 | -1.894 | 0.611 | -3.025 | 0.631 | -0.965 | 0.585 | 0.376 | -1.382 | -1.163 | | | | | | | | | | |
| B7L74_02315 | cytochrome C assembly protein | Cytochrome | -0.332 | 0.218 | 0.549 | 0.196 | -0.794 | 0.069 | -0.296 | 0.761 | 0.602 | 0.827 | 0.600 | | | | | | | | | | |
| B7L74_02320 | magnesium/cobalt efflux protein | Transporter | 1.241 | 1.864 | 3.666 | 1.460 | 4.899 | 3.988 | 3.366 | 0.978 | 3.659 | -0.819 | 1.193 | | | | | | | | | | |
| B7L74_02325 | prepilin-type cleavage/methylation domain-containing | Pilus | -0.815 | -0.532 | 0.039 | 0.952 | 0.313 | 1.606 | -0.663 | 0.725 | 1.899 | 0.945 | 0.803 | | | | | | | | | | |
| B7L74_02330 | adenylate kinase | | -0.997 | -0.257 | -0.434 | 0.143 | -1.014 | -0.370 | -0.670 | -1.139 | -1.244 | -0.313 | -0.391 | | | | | | | | | | |
| B7L74_02335 | thiol reductase thioredoxin | PV detoxification | 0.010 | -0.046 | 1.285 | 1.069 | -0.072 | -0.163 | 0.781 | -0.770 | -0.400 | 0.703 | 1.538 | | | | | | | | | | |
| B7L74_02340 | histone-like protein Hq1 | | 1.073 | 0.387 | 3.394 | 1.750 | 0.329 | 0.048 | 1.672 | 0.020 | -0.056 | 1.207 | 1.410 | | | | | | | | | | |
| B7L74_02345 | hypothetical protein | | -0.557 | -0.500 | -0.501 | -0.557 | -1.731 | -1.169 | -0.881 | -1.481 | -1.725 | -0.195 | 0.176 | | | | | | | | | | |
| B7L74_02350 | sodium:proton antiporter | Transporter | 0.037 | 0.837 | -0.034 | -1.809 | 3.230 | 2.629 | 0.444 | 1.948 | 5.775 | -1.282 | -0.911 | | | | | | | | | | |
| B7L74_02355 | hypothetical protein, Beta-lactamase induction protein | Resistance, Signaling proteins | -1.876 | -0.838 | -2.404 | -1.418 | -1.848 | -1.774 | -2.171 | -2.290 | -2.911 | -2.323 | -2.406 | Continuous down | | | | | | | | | |
| B7L74_02360 | pyruvate dehydrogenase (acetyl-transferring)%2C hom | Pyruvate metabolism | -2.260 | -1.333 | -3.002 | -1.917 | -2.590 | -2.022 | -3.046 | -3.029 | -3.396 | -2.743 | -1.833 | Continuous down | | | | Yes | Yes | | | | |
| B7L74_02365 | dihydrolipoyllysine-residue acetyltransferase | Pyruvate metabolism | 0.651 | -1.099 | -2.354 | -0.052 | 0.862 | 1.076 | -1.185 | -3.218 | -0.200 | -1.159 | -0.144 | Down in P42 | | | | Yes | Yes | | | | |
| B7L74_02370 | dihydrolipoyl dehydrogenase | TCA cycle onlyPyruvate to Acetyl | -1.363 | -0.766 | -3.360 | -1.708 | -1.992 | -1.881 | -2.189 | -3.228 | -4.104 | -1.882 | -0.721 | Continuous down | | | | Yes | Yes | | | | |
| B7L74_02375 | amidophosphoribosyltransferase | | -0.601 | -0.603 | -2.685 | -1.823 | -0.545 | -0.939 | -1.382 | -2.689 | -1.339 | -1.408 | -0.347 | Early down | | | | | | | | | |
| B7L74_02380 | hypothetical protein | | -0.459 | -2.025 | -5.367 | 0.140 | -1.701 | -2.743 | -1.135 | -3.836 | -2.704 | -3.120 | 0.722 | | | | | | | | | | |
| B7L74_02385 | DUF1658 domain-containing protein | | -0.118 | -0.179 | -0.604 | -1.067 | -0.823 | -0.577 | -0.395 | -0.439 | -1.795 | -1.323 | -0.390 | | | | | | | | | | |
| B7L74_02390 | DUF1658 domain-containing protein | | -0.136 | 0.427 | -1.696 | -1.633 | -0.420 | -0.505 | -0.425 | 0.427 | -1.156 | -2.047 | -1.203 | | | | | | | | | | |
| B7L74_02395 | malonyl-[acyl-carrier protein] O-methyltransferase Bio | | -0.078 | -0.043 | -0.442 | 0.908 | -0.472 | -1.144 | -0.074 | 1.198 | 0.051 | -0.075 | 0.701 | | | | | | | | | | |
| B7L74_02400 | exodeoxyribonuclease 7 small subunit | Mismatch repair | 0.918 | 1.141 | 0.931 | 2.408 | 1.190 | -0.073 | -0.102 | 1.643 | 0.481 | 0.588 | 0.855 | Early up | | | | | | | | | |
| B7L74_02405 | hypothetical protein | Effector protein | 0.796 | 1.288 | 1.378 | 0.610 | 0.840 | -0.226 | 0.109 | 1.041 | 0.954 | -0.688 | -0.514 | | | Yes | | | | | | | |
| B7L74_02410 | phosphoribosyl transferase | | 0.561 | 1.176 | 1.876 | -0.466 | 0.990 | 0.560 | 0.140 | 2.098 | 1.460 | 0.271 | 1.261 | | | | | | | | | | |
| B7L74_02415 | hypothetical protein | | 3.286 | 1.007 | 1.816 | -1.358 | 5.488 | 4.148 | 1.885 | 1.303 | 2.989 | 1.910 | 0.982 | Early up | | | | | | | | | |
| B7L74_02420 | hypothetical protein | | 1.365 | 1.509 | 0.247 | 0.045 | 1.865 | 1.527 | 0.991 | 2.405 | 1.714 | 0.348 | 0.953 | Early up | | | | | | | | | |
| B7L74_02425 | DNA helicase RecQ | | 0.159 | 0.533 | 0.150 | 1.746 | 0.932 | 1.093 | -0.121 | 0.529 | 1.145 | 0.066 | 0.539 | | | | | | | | | | |
| B7L74_02430 | DNA-binding protein | DNA repair and recombination pr | -0.791 | -0.490 | 1.009 | 3.010 | -0.043 | -0.441 | 0.831 | -0.300 | -0.312 | 1.171 | 1.843 | Up in P12 | | | | | | | | | |
| B7L74_02435 | hypothetical protein | | -0.067 | 0.070 | -0.397 | 1.292 | -0.145 | -0.096 | -0.034 | -0.840 | -1.495 | -0.026 | 0.320 | | | | | | | | | | |
| B7L74_02440 | hypothetical protein | | 0.966 | 0.362 | -0.166 | 0.505 | 0.234 | 0.210 | 0.248 | -0.917 | -2.742 | -0.502 | -1.036 | Down in P51 | | | | | | | | | |
| B7L74_02445 | hypothetical protein | | -0.637 | -0.577 | -0.712 | 0.881 | -0.217 | -0.058 | -0.402 | 0.419 | 0.421 | -0.320 | -0.160 | | | | | | | | | | |
| B7L74_02450 | hypothetical protein | Effector<br>AnkF. Eucaryotic-like ORFs. | 0.730 | 1.223 | 2.025 | 0.068 | 1.261 | 1.274 | 0.766 | 2.016 | 1.108 | 0.124 | 0.601 | | | Yes | | | | | | | |
| B7L74_02455 | hypothetical protein | | 0.122 | -1.632 | 1.513 | 1.146 | 0.257 | -0.853 | -1.953 | 2.978 | 4.323 | 1.034 | 2.248 | | | | | | | | | | |
| B7L74_02460 | hypothetical proteinprotein=conserved cytosolic protei | | 0.065 | -0.764 | 0.622 | 1.079 | -1.175 | -0.537 | 0.680 | 0.582 | 0.613 | 1.816 | 1.682 | | | | | | | | | | |
| B7L74_02465 | 3-deoxy-manno-octulosonate cytidylyltransferase | Lipopolysaccharide biosynthesis | 1.581 | 1.899 | 1.538 | 2.942 | 2.157 | 0.824 | 0.646 | 1.076 | 3.143 | 0.380 | 1.773 | | | | | | | | | | Yes |
| B7L74_02470 | arginine repressor | Transcriptional regulators | 0.089 | 0.847 | -0.035 | -0.789 | 5.785 | 4.858 | 0.527 | 1.496 | 2.628 | 2.122 | -0.897 | | | | | | | | | | |
| B7L74_02475 | arginine ABC transporter ATP-binding protein | ABC transporters | 0.250 | -1.421 | -0.106 | -0.594 | -2.229 | -1.808 | 0.002 | -0.647 | 0.136 | 1.589 | 1.725 | Variable | | | | | | | | Yes (Amino acid/Arginine) | |
| B7L74_02480 | arginine ABC transporter substrate-binding protein | Transporter | -1.138 | -1.916 | -1.833 | -1.569 | -2.867 | -0.961 | -1.147 | -1.654 | -1.042 | -0.158 | -0.218 | | | | | | | | | Yes (Amino acid/Arginine) | |
| B7L74_02485 | arginine ABC transporter permease | ABC transporters | 1.907 | 0.233 | 0.478 | -0.346 | 4.066 | 4.616 | 0.476 | 0.711 | 3.963 | 2.220 | 0.697 | Early up | | | | | | | | Yes (Amino acid/Arginine) | |
| B7L74_02490 | arginine transporter permease subunit ArtM | Transporter | 2.999 | 1.268 | 4.074 | 1.275 | 6.983 | 8.236 | 3.121 | 2.356 | 6.291 | 4.980 | 2.581 | Early up | | | | | | | | Yes (Amino acid/Arginine) | |
| B7L74_02495 | hypothetical membrane spanning protein | | 0.994 | -0.732 | 0.267 | 0.877 | 1.093 | 0.911 | 0.776 | -0.747 | 0.196 | 0.548 | 0.912 | | | | | | | | | | |
| B7L74_02500 | ribonuclease E/G | | 0.076 | -0.840 | -0.796 | -0.923 | -0.645 | -0.906 | -0.803 | -0.929 | -1.229 | -1.484 | -0.952 | | | | | | | | | | |
| B7L74_02505 | 23S rRNA pseudouridine(955/2504/2580) synthase | | 1.407 | -0.569 | 0.932 | 0.988 | 0.153 | 1.641 | 1.185 | 0.767 | 0.757 | 0.801 | 1.037 | | | | | | | | | | |
| B7L74_02510 | serine/threonine protein phosphatase | Eucaryotic-like ORFs. | 0.548 | 1.321 | 0.405 | -1.100 | 1.403 | 0.748 | 0.708 | 1.383 | 0.863 | 0.349 | 0.144 | | | | | | | | | | |
| B7L74_02515 | phospholipase A(1) | | -0.128 | 0.412 | -0.855 | -0.122 | 0.348 | -0.267 | -0.625 | 0.764 | 0.828 | 0.494 | 0.061 | | | | | | | | | | |
| B7L74_02520 | DNA polymerase/3'-5' exonuclease PolX | DNA | -0.118 | 0.168 | 0.769 | -1.180 | 0.663 | 0.747 | -0.020 | 0.308 | 0.679 | -0.008 | -0.198 | | | | | | | | | | |
| B7L74_02525 | 50S ribosomal protein L32 | Ribosomal proteins | -0.843 | -0.474 | -0.925 | 1.102 | -1.246 | -0.737 | -0.268 | -0.027 | -0.033 | -0.962 | -0.642 | | | | | | | | | | |
| B7L74_02530 | phosphate acyltransferase | Glycerolipid metabolism | -1.259 | -1.500 | -2.899 | -0.277 | -3.022 | -1.478 | -1.941 | -1.439 | -1.259 | -2.073 | -1.718 | Early down | | | | | | | | | |
| B7L74_02535 | 3-oxoacyl-ACP synthase III | | -0.321 | -0.366 | -0.391 | -0.756 | -0.732 | 0.291 | -1.380 | -2.095 | -0.473 | -1.978 | -0.314 | | | | | | | | Yes | | |
| B7L74_02540 | malonyl CoA-acyl carrier protein transacylase | Fatty acid biosynthesis | -0.981 | -1.856 | -2.661 | -0.286 | -1.751 | -2.040 | -1.113 | -2.220 | -2.214 | -1.590 | -0.266 | Early down | | | | | | | Yes | | |
| B7L74_02545 | beta-ketoacyl-ACP reductase(gene=fabG) | Fatty acid biosynthesis | -1.830 | -2.169 | -1.978 | -1.263 | -2.237 | -2.211 | -1.884 | -3.021 | -2.574 | -2.213 | -1.627 | Early down | | | | | | | Yes | | |
| B7L74_02550 | acyl carrier protein | Fatty acid biosynthesis | -0.799 | -1.672 | -0.538 | 0.939 | -1.482 | -1.618 | -0.910 | -1.527 | -2.410 | -0.270 | 0.196 | Early down | | | | | | | Yes | | |
| B7L74_02555 | beta-ketoacyl-[acyl-carrier-protein] synthase II | Fatty acid biosynthesis | -0.210 | -1.389 | -1.667 | 0.018 | -1.985 | -1.915 | -0.993 | -2.074 | -2.515 | -0.663 | -0.495 | Continuous down | | | | | | | Yes | | |
| B7L74_02560 | hypothetical exported protein | | 0.627 | 0.690 | 1.793 | -0.252 | 0.710 | 1.328 | 0.658 | 0.463 | 1.000 | 1.034 | 0.898 | | | | | | | | | | |
| B7L74_02565 | dTMP kinase | | -0.290 | -0.546 | 0.359 | -0.303 | -0.235 | -0.604 | 0.215 | 0.011 | -0.427 | 0.978 | 0.730 | | | | | | | | | | |
| B7L74_02570 | DNA polymerase III subunit delta' | DNA | -0.415 | 0.164 | -1.350 | -1.356 | 2.157 | 0.304 | -0.874 | 1.340 | 2.354 | -1.277 | -1.764 | | | | | | | | | | |
| B7L74_02575 | hypothetical protein | | 0.250 | 0.101 | 1.421 | -0.180 | 0.358 | 0.705 | 0.933 | 0.184 | -0.278 | 1.249 | 0.751 | | | | | | | | | | |
| B7L74_02580 | TatD family deoxyribonuclease | | 0.008 | -0.562 | 0.827 | 0.907 | -0.970 | -0.273 | 0.146 | 0.263 | 0.422 | 1.234 | 0.945 | | | | | | | | | | |
| B7L74_02585 | hypothetical protein | | -0.582 | -0.626 | -0.531 | 0.046 | 0.475 | 0.254 | -0.211 | 0.518 | -0.329 | -0.524 | -0.438 | | | | | | | | | | |
| B7L74_02590 | glutamine synthetase | | -0.269 | -0.943 | -0.482 | 0.376 | -0.834 | -0.963 | -0.366 | -1.959 | -1.300 | -0.366 | -0.104 | | | | | | | | | | |
| B7L74_02595 | MFS transporter, Di-/tripeptide transporter | Transporter | -0.040 | -0.348 | -0.211 | -0.500 | -0.633 | -0.165 | 0.097 | 0.026 | 0.207 | 0.289 | 0.578 | | | | | | | | | Yes (oligopeptide) | |
| B7L74_02600 | N-acetyltransferase<br>Ribosomal-proteinalanine acetyltransferase | Ribosomal | 0.603 | 0.256 | -0.938 | -1.183 | -0.748 | -0.385 | -1.171 | -0.827 | -0.573 | -1.517 | -1.764 | | | | | | | | | | |
| B7L74_02605 | single-stranded-DNA-specific exonuclease | DNA | 0.507 | 0.039 | 0.707 | 1.371 | -0.094 | -0.174 | 0.234 | 0.197 | 0.162 | 1.239 | 0.514 | | | | | | | | | | |
| B7L74_02610 | hypothetical cytosolic protein | | 0.202 | 0.173 | 3.259 | 2.654 | 0.357 | 1.032 | 1.743 | -0.113 | 1.442 | 1.860 | 2.551 | Early up | | | | | | | | | |
| B7L74_02615 | hypothetical membrane spanning protein | | 5.848 | 2.173 | 3.367 | 0.218 | 6.842 | 7.059 | 3.730 | 0.836 | 6.096 | 3.665 | 1.537 | Continuous up | | | | | | | | | |
| B7L74_02620 | Trp operon repressor | Transcriptional regulators | 0.736 | 0.080 | 1.994 | 2.118 | 0.570 | 0.973 | 1.004 | -0.868 | 0.162 | 1.992 | 2.808 | Early up | | | | | | | | | |
| B7L74_02625 | hypothetical protein | | 1.685 | -0.545 | 0.999 | 1.945 | 2.048 | 2.186 | 0.967 | -1.535 | 1.047 | 1.707 | 2.225 | | | | | | | | | | |
| B7L74_02630 | DUF1658 domain-containing protein | | 1.119 | -0.411 | 0.747 | 1.547 | 0.967 | 0.644 | 0.974 | -1.479 | -0.068 | 1.300 | 2.153 | | | | | | | | | | |
| B7L74_02635 | DUF1658 domain-containing protein | | 0.113 | -0.140 | 0.211 | 1.485 | 0.271 | 0.000 | 0.461 | -0.442 | -1.249 | 0.543 | 1.859 | | | | | | | | | | |
| B7L74_02640 | MFS transporter | Transporter | 2.180 | 0.595 | 1.964 | 2.444 | 2.640 | 4.144 | 2.349 | 3.937 | 4.837 | 2.409 | 2.318 | Early up | | | | | | | | | |
| B7L74_02645 | peptidase M14 | | 0.300 | 0.922 | 0.606 | 2.943 | 0.707 | 0.368 | -0.200 | 1.776 | 0.995 | 0.425 | 0.623 | Early up | | | | | | | | | |
| B7L74_02650 | fructose 1%2C6-bisphosphatase | Effector protein | 1.129 | 0.365 | 1.813 | 0.562 | -0.002 | 1.828 | 1.184 | 1.326 | 0.659 | 1.344 | 0.661 | | | Yes | | Yes | | | | | |
| B7L74_02655 | hypothetical membrane spanning protein | | 1.213 | 2.366 | -1.035 | -0.482 | 8.447 | 2.466 | 4.078 | 1.970 | 7.563 | -0.649 | -0.084 | Early up | | | | | | | | | |
| B7L74_02660 | hypothetical protein | | -0.199 | 0.004 | 2.828 | 0.314 | -0.216 | 0.020 | 1.378 | 1.610 | 1.148 | 0.958 | 1.095 | | | | | | | | | | |
| B7L74_02665 | MFS transporter | Transporter | 4.341 | 0.834 | 1.911 | 0.585 | 5.191 | 4.765 | 2.714 | 1.340 | 4.048 | 2.372 | 1.411 | Early up | | | | | | | | Yes (MFS) | |
| B7L74_02670 | hypothetical protein | | 1.470 | 0.086 | -0.825 | -1.529 | 1.928 | 2.020 | -0.807 | -0.675 | 1.015 | -0.953 | -1.403 | | | | | | | | | | |
| B7L74_02675 | hypothetical protein | | -0.286 | 0.675 | 1.449 | 1.240 | 0.155 | -0.518 | 1.737 | 0.047 | 0.051 | 1.065 | 1.800 | | | | | | | | | | |
| B7L74_02680 | tRNA-Asn | tRNA | 0.570 | 0.886 | 2.441 | -0.948 | 1.342 | 1.007 | 1.801 | 0.686 | 0.832 | -0.412 | -0.331 | Early up | | | | | | | | | |
| B7L74_02685 | aspartate aminotransferase | | -0.043 | 0.053 | -0.087 | -0.238 | -0.924 | -0.814 | -0.321 | -0.276 | -1.180 | 0.145 | 0.066 | | | | | | | | | | |
| B7L74_02690 | excinuclease ABC subunit B | | 1.033 | 0.610 | -2.499 | -2.175 | 2.274 | 3.253 | -1.958 | 1.545 | 3.039 | 0.600 | -1.666 | | | | | | | | | | |
| B7L74_02695 | hypothetical protein, DedA family protein - cell membr | Membrane | 4.362 | 1.280 | 2.727 | 0.117 | 5.671 | 6.022 | 2.758 | 0.005 | 5.977 | 3.900 | 0.985 | Early up | | | | | | | | Yes (Ions/Selenite) | |
| B7L74_02700 | hydroxymethylglutaryl-CoA lyase | Peroxisome | -1.699 | -0.633 | -2.126 | -0.448 | -2.338 | -1.491 | -2.008 | -0.997 | -1.299 | -1.395 | -1.333 | Early down | | | | | | | | | |
| B7L74_02705 | N-ethylammeline chlorohydrolase | | -3.842 | 0.407 | -5.032 | 1.363 | -0.427 | 0.755 | -0.742 | -1.546 | -2.004 | -6.241 | -1.523 | | | | | | | | | | |

| | | | | | | | | | | | | | | | | | | | | | | | | |
|---|---|---|---|---|---|---|---|---|---|---|---|---|---|---|---|---|---|---|---|---|---|---|---|---|
| B7L74_02710 | hypothetical protein | | 6.804 | 2.877 | 2.760 | -1.424 | 6.793 | 7.895 | 3.567 | 1.174 | 6.053 | 4.010 | 0.782 | Early up | | | | | | | | | | |
| B7L74_02715 | hypothetical protein | | 2.563 | 0.044 | 1.285 | -0.704 | 1.685 | 2.787 | 0.881 | -0.498 | 1.326 | 1.082 | 1.287 | | | | | | | | | | | |
| B7L74_02720 | IS110 family transposase | | 0.101 | -2.229 | -0.669 | -3.865 | 2.297 | 3.986 | -1.604 | 0.018 | -0.449 | -0.858 | 0.696 | | | | | | | | | | | |
| B7L74_02725 | DNA gyrase subunit A | | -0.197 | 0.731 | -0.639 | -1.153 | -0.487 | -0.240 | -0.745 | 0.186 | -1.730 | -0.498 | -0.822 | | | | | | | | | | | |
| B7L74_02730 | phosphoserine aminotransferase | Methane metabolism | -1.079 | 0.091 | -3.623 | -1.418 | -1.155 | -0.503 | -1.760 | -0.932 | -2.233 | -1.231 | -1.465 | Early down | | | | | | | | | | |
| B7L74_02735 | 3-phosphoshikimate 1-carboxyvinyltransferase | Phenylalanine, tyrosine and tryptop | -1.005 | -0.530 | -2.596 | -0.540 | -1.873 | -1.544 | -1.502 | -1.642 | -2.546 | -1.207 | -0.809 | Early down | | | | | | | | | | |
| B7L74_02740 | cytidylate kinase | Pyrimidine metabolism | -0.611 | -0.973 | -2.481 | -3.603 | -1.627 | -1.890 | -2.441 | -2.625 | -0.668 | -2.966 | -3.103 | Early down | | | | | | | | | | |
| B7L74_02745 | 30S ribosomal protein S1 | Ribosomal proteins | -1.270 | -1.449 | -1.071 | -0.474 | -1.766 | -1.982 | -0.999 | -2.464 | -1.716 | -0.856 | -0.322 | Early down | | | | | | | | | | |
| B7L74_02750 | Membrane associated protein LapA | Glycan metabolism | -1.517 | -0.470 | -1.542 | -0.725 | -1.220 | -0.104 | -1.151 | -0.989 | -1.276 | -0.808 | -0.668 | | | | | | | | | | Yes | |
| B7L74_02755 | lipopolysaccharide assembly protein LapB | Glycan metabolism | -2.800 | -1.043 | -0.857 | -2.171 | -0.425 | 0.568 | -0.538 | -0.505 | 2.109 | -1.583 | -1.223 | | | | | | | | | | Yes | |
| B7L74_02760 | orotidine 5'-phosphate decarboxylase | | -1.021 | -0.526 | -3.365 | -2.550 | -0.084 | -0.471 | -1.284 | -1.406 | 0.338 | -1.685 | -2.379 | | | | | | | | | | | |
| B7L74_02765 | competence protein ComEA | Pseudogenes | -0.714 | 0.742 | 1.103 | 0.950 | 1.963 | -0.154 | -0.133 | 0.421 | -0.925 | -0.569 | -1.096 | | | | | | | | | | | |
| B7L74_02770 | undecaprenyl-phosphate alpha-N-acetylglucosaminyl 1 | O-Antigen repeat unit biosynthesis | -0.159 | -0.632 | -0.562 | 0.661 | -0.505 | -0.658 | -0.507 | -0.875 | -1.242 | 0.166 | 0.031 | | | | | | | | | | Yes | |
| B7L74_02775 | hypothetical membrane associated protein | Effector protein | 3.833 | 1.043 | 1.063 | 0.359 | 4.114 | 4.144 | 1.893 | 0.931 | 2.818 | 2.443 | 1.444 | Early up | | Yes | | | | | | | | |
| B7L74_02780 | sporulation protein | | 0.378 | -0.803 | -0.068 | 0.196 | -1.165 | -0.839 | 0.018 | -1.325 | -1.782 | 0.202 | -0.024 | | | | | | | | | | | |
| B7L74_02785 | hypothetical protein | | 0.937 | 0.657 | 2.392 | -0.258 | 0.807 | 0.452 | 1.489 | -0.146 | 0.313 | 0.803 | 3.204 | | | | | | | | | | | |
| B7L74_02790 | DUF1658 domain-containing protein | | 1.692 | 1.491 | 2.352 | -0.756 | 1.253 | 0.725 | 1.076 | 0.519 | 0.743 | 0.799 | 1.154 | | | | | | | | | | | |
| B7L74_02795 | hypothetical protein | | 1.917 | 1.158 | 2.629 | -1.617 | 1.368 | 1.056 | 2.561 | 1.069 | 0.043 | 0.475 | 0.985 | | | | | | | | | | | |
| B7L74_02800 | DUF1658 domain-containing protein | CB specific protein family | 1.580 | 1.363 | 3.866 | -0.536 | 1.032 | 0.959 | 3.326 | 1.253 | 1.366 | 0.640 | 1.577 | Early up | | | | | | | | | | |
| B7L74_02805 | MFS transporter (gene=tppB) | Transporter | -0.234 | -0.345 | -0.546 | -0.886 | -0.387 | -0.214 | -0.676 | -1.095 | -0.983 | -1.478 | -1.165 | | | | | | | | | Yes (oligopeptide) | | |
| B7L74_02810 | chromosome partition protein Smc | | -0.985 | -1.216 | -0.771 | 0.530 | -1.497 | -1.520 | -0.373 | -1.682 | -1.667 | -0.044 | 0.888 | | | | | | | | | | | |
| B7L74_02815 | cell division protein ZipA | Cell division | -1.116 | -0.931 | -2.852 | -1.578 | -1.514 | -1.736 | -1.537 | -2.235 | -2.050 | -1.284 | -0.985 | Early down | | | | | | | | | | |
| B7L74_02820 | DNA ligase (NAD(+)) (gene=ligA) | DNA replication | -0.841 | -1.507 | -2.170 | -1.451 | -1.535 | -0.945 | -1.372 | -1.893 | -1.506 | -0.983 | -0.583 | Early down | | | | | | | | | | |
| B7L74_02825 | hypothetical membrane spanning protein | | 6.693 | 2.881 | 3.633 | 0.426 | 8.210 | 8.249 | 3.937 | 1.831 | 7.098 | 4.258 | 2.631 | Early up | | | | | | | | | | |
| B7L74_02830 | hypothetical protein | | 0.508 | 0.727 | 1.700 | 2.177 | 0.307 | 0.716 | 0.762 | 1.016 | 0.510 | 1.019 | 1.258 | Up in P12 | | | | | | | | | | |
| B7L74_02835 | LemA family protein | | -1.015 | 0.504 | -0.096 | 0.617 | 0.873 | 0.556 | -1.199 | 0.031 | -0.181 | -0.845 | -1.580 | | | | | | | | | | | |
| B7L74_02840 | zinc metalloprotease HtpX | | -0.336 | 0.151 | -0.499 | 0.131 | 0.206 | 0.257 | -0.743 | -0.757 | -1.110 | -1.046 | -1.024 | | | | | | | | | | | |
| B7L74_02845 | hypothetical protein tetratricopeptide repeat family prot | Eucaryotic-like ORFs. | -0.816 | -1.094 | 0.217 | 0.211 | -1.351 | -1.008 | -0.544 | 0.057 | -0.427 | -0.167 | -0.271 | | | | | | | | | | | |
| B7L74_02850 | lytic murein transglycosylase B | | 0.038 | 0.837 | -0.034 | -1.808 | 2.638 | 1.620 | 0.445 | 1.486 | 4.229 | -0.778 | -0.470 | | | | | | | | | | | |
| B7L74_02855 | rod shape-determining protein RodA | Chromosome and associated protei | 1.831 | -0.550 | -0.710 | -1.116 | 2.947 | 3.353 | -0.203 | -0.849 | 2.221 | 0.011 | -0.985 | Up in P21 | | | | | | | | | | |
| B7L74_02860 | penicillin-binding protein | | -0.419 | -0.911 | -0.678 | -1.146 | -0.888 | -0.791 | -0.560 | -0.674 | -1.113 | -0.704 | -0.736 | | | | | | | | | | | Yes |
| B7L74_02865 | 23S rRNA (pseudouridine(1915)-N(3))-methyltransfer | | -0.451 | -0.424 | 0.796 | 2.540 | -2.347 | 0.773 | -1.623 | 2.352 | 2.211 | 0.064 | 0.988 | | | | | | | | | | | |
| B7L74_02870 | ribosome silencing factor RsfS | | -1.208 | 0.563 | -0.333 | 3.424 | -1.004 | 1.610 | 1.275 | 1.480 | 0.575 | -0.902 | -0.128 | | | | | | | | | | | |
| B7L74_02875 | IS110 family transposase | | 0.454 | 0.339 | 1.908 | -0.647 | -0.091 | -0.025 | 0.478 | 0.051 | -0.996 | 1.252 | 1.162 | | | | | | | | | | | |
| B7L74_02880 | hypothetical protein | | 0.104 | -0.081 | -0.034 | -1.746 | 1.315 | 0.961 | -0.229 | -0.946 | 1.871 | -1.244 | -0.891 | | | | | | | | | | | |
| B7L74_02885 | hypothetical protein | | 1.122 | 0.088 | 2.601 | -0.999 | 1.168 | 0.538 | 1.544 | 0.198 | 1.244 | 2.968 | 2.919 | Late up | | | | | | | | | | |
| B7L74_02890 | nicotinate-nicotinamide nucleotide adenylyltransferase | | -1.816 | -1.442 | -0.868 | -1.247 | -2.499 | -1.144 | -0.878 | -0.857 | -0.194 | -0.126 | -0.233 | | | | | | | | | | | |
| B7L74_02895 | DNA polymerase III subunit delta | DNA | 1.755 | -0.751 | -0.049 | 2.608 | 3.679 | 2.834 | -0.455 | 2.687 | 2.086 | -1.816 | -1.153 | | | | | | | | | | | |
| B7L74_02900 | hypothetical proteinrare lipoprotein B precursor | | 1.594 | -0.609 | -1.178 | -0.927 | 1.477 | 1.819 | -0.049 | -0.715 | 1.272 | -0.902 | -0.551 | | | | | | | | | Yes (Outer membrane/LPS) | | |
| B7L74_02905 | leucine--tRNA ligase | | -1.717 | -4.317 | -6.490 | -1.358 | -0.637 | -0.365 | -4.338 | -1.416 | -3.207 | -1.956 | -3.676 | | | | | | | | | | | |
| B7L74_02910 | hypothetical protein, Hypothetical cytosolic protein | | 0.915 | -0.028 | 2.439 | 1.333 | 1.519 | 0.953 | 1.355 | -0.067 | 0.979 | 1.099 | 1.789 | Early up | | | | | | | | | | |
| B7L74_02915 | phytochelatin synthase | | 0.868 | 1.415 | 0.782 | 1.336 | 2.113 | 1.540 | 1.044 | 2.452 | 2.555 | 0.416 | 0.445 | Early up | | | | | | | | | | |
| B7L74_02920 | ATP-binding protein | ATP | -0.867 | 0.356 | -1.768 | -1.330 | -0.625 | -0.686 | -0.871 | -0.296 | -0.227 | 0.175 | -0.510 | | | | | | | | | | | |
| B7L74_02925 | hypothetical protein | | 0.290 | 1.369 | 2.438 | 0.797 | 1.084 | 0.559 | 0.694 | 0.981 | 1.790 | 0.523 | 0.192 | | | | | | | | | | | |
| B7L74_02930 | hypothetical protein | | 0.035 | 0.837 | -0.034 | -1.810 | 2.635 | 1.620 | 0.443 | 1.486 | 2.542 | -1.283 | -0.912 | | | | | | | | | | | |
| B7L74_02935 | apolipoprotein N-acyltransferase | | -0.564 | -0.348 | -0.970 | -0.132 | -0.416 | -0.392 | -0.392 | -0.147 | -0.210 | -0.305 | -0.462 | | | | | | | | | | | |
| B7L74_02940 | magnesium/cobalt efflux protein | | 2.860 | 2.632 | -0.611 | 3.126 | 3.606 | 2.581 | 3.140 | 3.945 | 2.968 | 2.712 | 2.678 | | | | | | | | | Yes (Ions/Magnesium, cobalt) | | |
| B7L74_02945 | MFS transporter | Transporter | 0.274 | -0.341 | 0.859 | -0.671 | -0.720 | -0.130 | 0.477 | -0.150 | 0.491 | 0.924 | 0.972 | | | | | | | | | | | |
| B7L74_02950 | endoribonuclease YbeY | | -0.635 | -0.751 | 0.003 | 0.419 | -1.259 | -1.299 | -1.152 | -1.356 | -1.797 | 0.092 | -0.553 | | | | | | | | | | | |
| B7L74_02955 | phosphate starvation-inducible protein PhoH | | 2.556 | -0.473 | 1.859 | -0.941 | 0.700 | 1.510 | 2.113 | 1.174 | 3.080 | 2.946 | 2.379 | | | | | | | | | | | |
| B7L74_02960 | tRNA (N6-isopentenyl adenosine(37)-C2)-methylthiot | Transfer RNA biogenesis | -1.755 | -1.278 | -4.105 | -2.418 | -2.853 | -2.451 | -2.609 | -3.027 | -2.657 | -3.039 | -2.929 | Continuous down | | | | | | | | | | |
| B7L74_02965 | amino acid permease | Transporter | -0.408 | -1.314 | -1.834 | -1.151 | 0.482 | 0.139 | -2.093 | -2.168 | 0.989 | -0.820 | -1.750 | | | | | | | | | | | |
| B7L74_02970 | hypothetical exported protein | | 1.308 | 0.111 | 1.155 | 0.033 | 0.885 | 0.255 | 0.561 | 1.164 | 0.908 | 0.885 | 1.176 | | | | | | | | | | | |
| B7L74_02975 | leucyl aminopeptidase | | 0.042 | -0.416 | -0.933 | -0.057 | -0.390 | 0.347 | -0.737 | -1.616 | -1.403 | -1.069 | -0.673 | | | | | | | | | | | |
| B7L74_02980 | acyl-CoA dehydrogenase(gene=fadE) | Fatty acid degradation | -0.335 | -0.490 | -0.106 | -0.113 | -1.208 | -1.353 | -0.290 | -2.127 | -2.142 | -0.089 | 1.071 | Late down | | | | | | | | | | |
| B7L74_02985 | DUF1658 domain-containing protein | | 0.877 | 0.271 | 0.842 | 0.213 | 0.059 | -0.472 | 0.667 | -1.388 | -1.675 | 0.338 | 1.559 | | | | | | | | | | | |
| B7L74_02990 | acetyl-CoA acetyltransferase | Pyruvate metabolism | -1.094 | -0.802 | -1.162 | -1.934 | -1.226 | -1.444 | -1.574 | -2.239 | -1.856 | -1.403 | -1.471 | Early down | | | | | | | | | | |
| B7L74_02995 | crotonase | | 0.290 | -0.298 | -0.729 | -1.410 | 2.678 | 2.736 | -1.167 | -1.601 | 1.854 | 0.137 | -1.106 | | | | | | | | | | | |
| B7L74_03000 | hypothetical membrane spanning protein | | 6.772 | 1.577 | 4.741 | -0.508 | 9.177 | 6.676 | 6.957 | 2.921 | 10.732 | 2.718 | 1.653 | Continuous up | | | | | | | | | | |
| B7L74_03005 | glycosyltransferase | | -0.739 | -0.794 | -2.295 | -0.447 | -0.794 | -0.400 | -2.675 | -0.852 | -0.033 | -1.388 | -1.814 | | | | | | | | | | | |
| B7L74_03010 | UDP-phosphomannose--protein mannosyltransferase | | 0.495 | 0.861 | -1.140 | 0.014 | 2.150 | 2.106 | 0.252 | 0.169 | -1.329 | -1.007 | -0.934 | | | | | | | | | | | |
| B7L74_03015 | hypothetical protein | | -0.694 | -0.482 | -1.126 | 0.594 | -1.018 | -0.695 | -1.186 | -0.822 | -0.891 | -0.915 | -1.166 | | | | | | | | | | | |
| B7L74_03020 | ferredoxin | | 0.090 | 0.848 | -0.035 | 5.721 | 3.861 | 1.627 | 2.184 | 1.496 | 2.628 | 3.015 | 3.156 | | | | | | | | | | | |
| B7L74_03025 | BolA family transcriptional regulator | Transcriptional regulators | -2.401 | -1.340 | -1.135 | 0.423 | -2.057 | -2.194 | -1.416 | -2.004 | -3.239 | -1.453 | -1.359 | Early down | | | | | | | | | | |
| B7L74_03030 | monothiol glutaredoxin%2C Grx4 family | PV detoxification | -2.040 | -1.290 | -2.679 | 0.240 | -1.951 | -2.459 | -1.696 | -2.328 | -3.635 | -1.905 | -2.001 | Early down | | | Yes | | | | | | | |
| B7L74_03035 | hypothetical protein | | -0.799 | -0.989 | -2.087 | 1.173 | -1.561 | -2.160 | -0.823 | -1.587 | -2.817 | -0.378 | -0.514 | Early down | | | | | | | | | | |
| B7L74_03040 | hypothetical protein | | -0.488 | -0.811 | 0.177 | 1.811 | -0.541 | -0.804 | 0.226 | -1.242 | -1.461 | 0.510 | 0.397 | | | | | | | | | | | |
| B7L74_03045 | pyridine nucleotide-disulfide oxidoreductase | | -0.302 | -0.701 | 0.499 | -1.264 | -0.769 | -0.452 | -0.444 | -1.836 | -1.245 | -0.301 | -0.765 | | | | | | | | | | | |
| B7L74_03050 | thiosulfate sulfurtransferase | Sulfur metabolism | -0.864 | -1.417 | 0.443 | 0.448 | -1.681 | -1.881 | -0.567 | -2.213 | -2.533 | -0.375 | -0.525 | Continuous down | | | | | | | | | | |
| B7L74_03055 | quinolinate synthetase | Nicotinate and nicotinamide metab | -1.092 | -1.898 | -2.035 | -0.807 | -2.390 | -2.661 | -1.391 | -3.089 | -3.806 | -1.972 | -1.776 | Continuous down | | | | | | | | | | |
| B7L74_03060 | ferredoxin | | -0.336 | -0.547 | -2.402 | -2.120 | -1.164 | -1.493 | -1.328 | -1.535 | -3.616 | -1.925 | -1.969 | Early down | | | | | | | | | | |
| B7L74_03065 | hypothetical protein | Effector protein | 0.519 | 0.254 | -1.577 | -1.935 | -0.646 | -1.170 | -0.816 | -0.990 | -4.509 | -1.213 | -1.430 | Early down | | Yes | | | | | | | | |
| B7L74_03070 | hypothetical cytosolic protein | | -1.703 | -1.624 | -2.244 | -1.458 | -2.173 | -2.245 | -1.887 | -2.350 | -3.055 | -1.792 | -2.022 | Early down | | | | | | | | | | |
| B7L74_03075 | hypothetical membrane spanning protein | | -0.852 | -1.124 | -2.187 | -0.760 | -1.686 | -1.735 | -1.709 | -2.589 | -3.090 | -1.037 | -1.529 | Continuous down | | | | | | | | | | |
| B7L74_03080 | hypothetical protein | | 0.089 | 0.847 | -0.035 | -0.096 | 2.736 | 4.363 | 0.526 | 1.496 | 8.154 | 3.503 | 0.096 | | | | | | | | | | | |
| B7L74_03085 | hypothetical protein | | 0.090 | 0.847 | -0.035 | -0.096 | 2.736 | 2.589 | 0.527 | 1.496 | 3.555 | -0.331 | 0.097 | | | | | | | | | | | |
| B7L74_03090 | MBL fold hydrolase | | 0.170 | 0.197 | 1.527 | 1.813 | -2.307 | 2.115 | 1.220 | 3.013 | 2.434 | 1.851 | 1.814 | Late up | | | | | | | | | | |
| B7L74_03095 | hypothetical protein | | 4.593 | 3.393 | 8.383 | 2.994 | 2.794 | 1.695 | 7.056 | 4.502 | 6.312 | 5.669 | 5.938 | Early up | | | | | | | | | | |
| B7L74_03100 | coproporphyrinogen III oxidase | | 0.974 | 0.607 | 1.573 | 1.962 | 1.433 | 1.071 | 1.149 | 2.285 | 2.725 | 1.810 | 2.409 | Late up | | | | | | | | | | |
| B7L74_03105 | ADP compounds hydrolase NudE | | 0.184 | -0.471 | 1.069 | 1.455 | -0.170 | 0.172 | 0.419 | -0.057 | -0.449 | 0.755 | 0.841 | | | | | | | | | | | |
| B7L74_03110 | 3'(2')%2C5'-bisphosphate nucleotidase CysQ | | 0.038 | 0.837 | -0.034 | -1.808 | 3.702 | 1.620 | 0.446 | 1.948 | 4.640 | -1.281 | -0.911 | | | | | | | | | | | |
| B7L74_03115 | hypothetical protein | | 7.148 | 3.314 | 4.285 | 1.092 | 8.742 | 8.603 | 4.357 | 2.070 | 7.679 | 4.830 | 2.615 | Early up | | | | | | | | | | |
| B7L74_03120 | hypothetical protein | | 7.460 | 3.251 | 4.136 | 1.041 | 9.479 | 8.404 | 4.847 | 2.013 | 7.213 | 4.839 | 2.850 | Early up | | | | | | | | | | |
| B7L74_03125 | hypothetical protein | | 0.421 | 0.193 | 1.716 | 2.183 | -0.999 | 1.179 | 0.519 | 1.624 | 2.059 | 0.177 | 0.575 | | | | | | | | | | | |
| B7L74_03130 | hypothetical protein | | 0.962 | -0.115 | -0.461 | 2.127 | 0.841 | 2.240 | 0.897 | 2.892 | 3.015 | -0.026 | 0.706 | Late up | | | | | | | | | | |
| B7L74_03135 | hypothetical protein | | 0.107 | -0.290 | -0.435 | 1.306 | -0.569 | 1.577 | -0.155 | 2.449 | 2.751 | 0.647 | 0.392 | Late up | | | | | | | | | | |
| B7L74_03140 | dihydrofolate reductase | Eucaryotic-like ORFs. | 0.119 | -2.176 | -1.810 | -1.525 | 1.686 | 1.424 | -1.381 | -1.574 | 1.194 | -0.976 | -1.380 | | | | | | | | | | | |
| B7L74_03145 | mevalonate kinase | | 0.042 | 0.838 | 2.966 | -1.805 | 2.643 | 4.257 | 1.028 | 3.109 | 2.391 | -1.279 | -0.910 | | | | | | | | | | | |
| B7L74_03150 | mevalonate kinase | Kinase , Peroxisome | -1.354 | -2.586 | -0.852 | -0.890 | -3.161 | -2.186 | -1.342 | -1.964 | -1.511 | -0.551 | -0.427 | Early down | | | | | | | | | | |
| B7L74_03155 | hydroxymethylglutaryl-CoA reductase (NADPH) | | -1.399 | -0.964 | -5.429 | -0.782 | -1.438 | 0.273 | -0.378 | -1.706 | 0.066 | -0.825 | -0.021 | | | | | | | | | | | |
| B7L74_03160 | outer membrane protein assembly factor BamA | Membrane | -2.226 | -2.262 | -2.712 | -1.197 | -3.402 | -2.406 | -2.108 | -2.449 | -2.526 | -1.773 | -1.745 | Early down | | | | | | | | | | |
| B7L74_03165 | hypothetical proteinouter membrane protein | Structural proteins | -2.936 | -2.230 | -2.285 | 0.555 | -2.621 | -2.628 | -1.602 | -3.601 | -4.535 | -1.925 | -1.928 | Continuous down | | | | | | | | | | |
| B7L74_03170 | UDP-3-O-(3-hydroxymyristoyl)glucosamine N-acyltra | Lipopolysaccharide biosynthesis | -1.117 | -2.129 | -2.341 | 0.318 | -2.360 | -2.311 | -0.973 | -3.786 | -4.079 | -1.249 | -0.874 | Continuous down | | | | | | | | | Yes | |
| B7L74_03175 | beta-hydroxyacyl-ACP dehydratase | | 0.035 | 0.837 | -0.034 | -1.810 | 2.635 | 1.620 | 0.443 | 1.486 | 2.542 | -1.283 | -0.912 | | | | | | | | Yes | | | |
| B7L74_03180 | acyl-[acyl-carrier-protein]--UDP-N-acetylglucosamine | Lipopolysaccharide biosynthesis | -1.041 | -2.231 | -2.157 | -0.433 | -2.629 | -1.533 | -1.444 | -1.884 | -1.304 | -1.162 | -1.067 | Early down | | | | | | | | | Yes | |
| B7L74_03185 | hydrolase | | 0.055 | 0.841 | 1.551 | -0.358 | 2.173 | 0.147 | -0.219 | 3.803 | 3.525 | -0.922 | -0.435 | | | | | | | | | | | |
| B7L74_03190 | hypothetical membrane spanning protein | | 7.311 | 3.108 | 4.560 | 0.771 | 8.644 | 8.725 | 4.933 | 2.249 | 7.767 | 5.310 | 2.791 | Early up | | | | | | | | | | |
| B7L74_03195 | acyltransferase | | 5.849 | 2.093 | 3.953 | 0.602 | 7.272 | 7.622 | 4.579 | 2.745 | 6.729 | 4.201 | 2.776 | Continuous up | | | | | | | | | | |
| B7L74_03200 | hypothetical exported protein | | -0.429 | -1.420 | -1.188 | -1.589 | -2.284 | -0.826 | -0.407 | -0.411 | -1.362 | -0.443 | -0.787 | | | | | | | | | | | |
| B7L74_03205 | lipid-A-disaccharide synthase | Lipopolysaccharide biosynthesis | -0.171 | -0.526 | -0.701 | 0.522 | -0.198 | -0.017 | -0.688 | -1.107 | -0.889 | -0.366 | -0.564 | | | | | | | | | | Yes | |
| B7L74_03210 | UDP-N-acetyl-D-glucosamine dehydrogenase | | 0.230 | -0.388 | -1.874 | 1.287 | -2.223 | -0.639 | 0.162 | -1.792 | -3.942 | -0.214 | -0.152 | | | | | | | | | | | |
| B7L74_03215 | tRNA-specific adenosine deaminase | | 0.160 | -0.112 | 0.794 | -0.253 | -0.416 | -0.246 | 0.371 | 0.496 | 0.158 | 0.658 | -0.177 | | | | | | | | | | | |
| B7L74_03220 | hypothetical protein | | 0.298 | 0.796 | -0.429 | 0.155 | 0.442 | 1.223 | -0.235 | 1.560 | 1.442 | 0.535 | 0.043 | | | | | | | | | | | |
| B7L74_03225 | hypothetical protein | | -1.097 | 1.427 | 0.380 | 1.552 | -1.291 | -2.401 | -1.348 | 1.605 | 1.672 | 1.024 | 0.278 | | | | | | | | | | | |
| B7L74_03230 | hypothetical protein | | 0.029 | -0.188 | 0.680 | -0.473 | -0.659 | -1.404 | -0.346 | -1.271 | -1.292 | 0.039 | -0.024 | | | | | | | | | | | |
| B7L74_03235 | hypothetical protein | Effector protein | 0.043 | -0.489 | -1.430 | -1.522 | -0.898 | -1.047 | -1.191 | -1.265 | -0.799 | -1.053 | -1.067 | | | Yes | | | | | | | | |
| B7L74_03240 | hypothetical protein | | 1.246 | -0.967 | 0.246 | -0.154 | -1.663 | -2.263 | -0.051 | -3.534 | -2.976 | -0.387 | 0.481 | Early down | | | | | | | | | | |
| B7L74_03245 | inorganic pyrophosphatase | Oxidative phosphorylation | -0.555 | -1.669 | -0.069 | 0.651 | -2.324 | -2.480 | -0.133 | -2.684 | -2.784 | -0.256 | 0.973 | Continuous down | | | | | | Yes | | | | |
| B7L74_03250 | bifunctional proline dehydrogenase/L-glutamate gamm | Alanine, aspartate and glutamate m | -0.914 | -1.347 | -2.528 | -0.629 | -2.066 | -2.577 | -1.695 | -3.307 | -4.914 | -2.172 | -1.087 | Continuous down | | | | | | | | | | |
| B7L74_03255 | peptidyl-prolyl cis-trans isomerase | Chaperones and folding catalysts | -3.213 | -1.402 | -0.823 | 2.004 | -0.846 | -2.269 | -1.801 | -2.289 | -3.611 | -2.970 | -2.521 | Early down | | | Yes | | | | | | | |
| B7L74_03260 | phosphoribosylformylglycinamidine synthase | | -2.875 | -0.394 | -2.924 | -2.214 | 1.479 | 1.480 | 0.958 | 0.982 | 0.301 | -1.164 | -1.140 | | | | | | | | | | | |
| B7L74_03265 | ferritin | | -0.715 | -0.725 | -0.138 | 1.519 | -0.280 | -0.516 | -0.416 | -1.357 | -2.981 | -1.123 | -0.788 | Down in P51 | | | | | | | | | | |
| B7L74_03270 | hybrid sensor histidine kinase/response regulator | Kinase | -1.011 | -1.087 | -1.363 | -0.340 | -0.551 | -0.337 | -1.319 | -1.180 | -1.190 | -0.426 | -1.131 | | | | | | | | | | | |
| B7L74_03275 | hypothetical membrane spanning protein | Effector protein | -0.831 | -1.109 | -2.043 | -0.356 | -0.864 | -0.643 | -1.108 | -1.529 | -1.654 | -1.168 | -1.626 | Early down | | Yes | | | | | | | | |
| B7L74_03280 | EamA family transporter | Transporter | -1.186 | -1.687 | -3.441 | -0.416 | -1.223 | -0.879 | -0.677 | -1.259 | -0.587 | -0.929 | -1.396 | Down in P10 | | | | | | | | Yes (Vitamins/Riboflavin) | | |
| B7L74_03285 | pyrroloquinoline quinone biosynthesis protein PqqC | Effector protein | 1.354 | -1.161 | -0.599 | 2.642 | 1.183 | 1.133 | 0.791 | 0.510 | 1.076 | 2.483 | 1.069 | Early up | | Yes | | | | | | | | |
| B7L74_03290 | branched-chain alpha-keto acid dehydrogenase subuni | Propanoate metabolism | -0.952 | -3.113 | -3.250 | -1.125 | -3.332 | -2.591 | -1.416 | -3.281 | -3.058 | -1.008 | -1.082 | Early down | | | | | | | | | | |
| B7L74_03295 | alpha-ketoacid dehydrogenase subunit beta | | -0.515 | 0.778 | -0.630 | 0.792 | 3.953 | 3.278 | 1.377 | 3.147 | 2.653 | -0.418 | 1.404 | | | | | | | | | | | |
| B7L74_03300 | pyruvate dehydrogenase (acetyl-transferring) E1 comp | Propanoate metabolism | -1.319 | -3.579 | -2.504 | 0.156 | -2.112 | -1.857 | -1.151 | -1.792 | -1.114 | -0.859 | -0.005 | Early down | | | | | | | | | | |
| B7L74_03305 | leucine dehydrogenase | Valine, leucine and isoleucine deg | 2.616 | 2.441 | 1.787 | 3.690 | 4.082 | 3.951 | 3.166 | 2.853 | 1.782 | 2.343 | 1.588 | Early up | | | | | | | | | | |
| B7L74_03310 | glycerophosphoryl diester phosphodiesterase | | -0.478 | -0.970 | 0.272 | 0.313 | -1.418 | -0.954 | -0.321 | 0.172 | 0.414 | 0.452 | 0.572 | | | | | | | | | | | |
| B7L74_03315 | bifunctional diaminohydroxyphosphoribosylaminopyri | | 2.301 | 0.150 | 0.880 | -0.180 | 2.239 | 2.500 | 1.725 | -0.701 | 1.619 | -0.576 | 0.562 | | | | | | | | | | | |
| B7L74_03320 | plasmid stabilization protein | | -1.266 | 1.559 | 0.569 | -0.610 | 2.068 | 3.369 | -0.469 | 1.671 | 2.005 | 1.708 | 1.211 | | | | | | | | | | | |
| B7L74_03325 | prevent-host-death protein | | -0.010 | 1.164 | 1.822 | 0.063 | 1.631 | 0.713 | 0.958 | 1.475 | 1.731 | 1.834 | 1.475 | | | | | | | | | | | |
| B7L74_03330 | riboflavin synthase | Riboflavin metabolism | -1.415 | -1.045 | -1.877 | -0.157 | -1.703 | -1.852 | -1.508 | -1.535 | -2.152 | -1.150 | -0.950 | Continuous down | | | | | | | | | | |
| B7L74_03335 | bifunctional 3%2C4-dihydroxy-2-butanone-4-phospha | Riboflavin metabolism | -1.238 | -1.109 | -1.511 | -0.045 | -2.115 | -1.952 | -1.336 | -2.329 | -3.480 | -0.768 | -0.631 | Continuous down | | | | | | | | | | |
| B7L74_03340 | 6%2C7-dimethyl-8-ribityllumazine synthase | | 0.031 | -1.115 | -0.815 | 0.217 | -1.097 | 1.550 | -0.467 | -1.797 | 1.412 | 0.321 | -0.223 | | | | | | | | | | | |
| B7L74_03345 | AI-2E family transporterRiboflavin transporter | Transporter | 3.133 | -1.251 | -2.811 | -1.702 | 5.510 | 3.753 | -0.958 | -1.959 | 2.910 | 0.480 | -1.409 | Up in P16 | | | | | | | | Yes (Vitamins/Riboflavin) | | |
| B7L74_03350 | hypothetical protein | | 1.279 | 0.265 | 1.565 | 1.596 | 0.644 | -0.354 | 1.943 | 1.741 | 1.040 | 1.402 | 2.414 | Early up | | | | | | | | | | |
| B7L74_03355 | hypothetical protein | | 1.095 | 1.174 | 2.199 | 1.450 | 5.191 | 3.688 | 2.060 | 4.351 | 4.515 | 2.793 | 3.286 | Early up | | | | | | | | | | |
| B7L74_03360 | hypothetical protein | | 1.265 | 0.919 | -1.641 | -1.385 | 1.776 | 2.410 | -2.315 | 2.548 | 3.254 | -0.309 | 0.032 | | | | | | | | | | | |
| B7L74_03365 | hypothetical protein | | 0.761 | -1.571 | 0.401 | -0.480 | -0.120 | 1.079 | 0.537 | 1.236 | 1.583 | -0.213 | 0.018 | | | | | | | | | | | |
| B7L74_03370 | hypothetical protein | | 3.371 | 0.846 | -0.034 | -0.707 | 4.241 | 2.592 | 1.972 | 1.495 | 2.976 | 0.761 | -0.057 | | | | | | | | | | | |
| B7L74_03375 | hypothetical protein | | 8.513 | 2.561 | 2.665 | 1.387 | 9.848 | 8.737 | 3.754 | 2.649 | 8.249 | 3.819 | 1.681 | Early up | | | | | | | | | | |
| B7L74_03380 | YbaB/EbfC family nucleoid-associated protein | | -1.411 | -1.488 | -0.765 | 2.409 | 0.273 | -0.302 | -0.154 | 0.092 | 0.094 | 0.930 | 0.462 | | | | | | | | | | | |
| B7L74_03385 | recombination protein RecR | | 2.033 | 1.058 | 1.337 | 0.285 | 0.751 | -0.315 | 1.034 | -0.060 | -1.773 | -0.202 | -0.445 | | | | | | | | | | | |
| B7L74_03390 | hypothetical protein | | -0.259 | 0.373 | 1.895 | 0.517 | -0.581 | -0.664 | -0.587 | 0.173 | -2.704 | -0.913 | -0.719 | Down in P51 | | | | | | | | | | |
| B7L74_03395 | DNA polymerase III subunit gamma/tau | DNA replication | -1.087 | -1.181 | -0.441 | -1.326 | -2.227 | -1.388 | -1.367 | -2.136 | -1.141 | -0.211 | -0.212 | Early down | | | | | | | | | | |
| B7L74_03400 | signal recognition particle sRNA small type (gene=ffs) | rRNA | 1.115 | 0.761 | 3.639 | -0.628 | 2.127 | 1.347 | 4.576 | 3.081 | 1.557 | -0.808 | 0.385 | Early up | | | | | | | | | | |
| B7L74_03405 | tRNA-Ser | tRNA | 1.678 | 0.952 | -2.035 | -1.781 | 1.082 | 0.010 | 0.324 | -0.089 | 0.554 | -2.025 | -0.788 | | | | | | | | | | | |
| B7L74_03410 | hypothetical protein | | 3.440 | -0.179 | 2.396 | 0.668 | 2.941 | 3.427 | 1.505 | 4.007 | 4.503 | 2.518 | 2.484 | Early up | | | | | | | | | | |
| B7L74_03415 | hypothetical protein | | 1.656 | 2.237 | 2.408 | 0.684 | 4.681 | 4.911 | 3.256 | 3.940 | 5.045 | 0.490 | 1.548 | | | | | | | | | | | |
| B7L74_03420 | hypothetical protein | | 4.943 | 0.759 | 0.986 | -1.324 | 5.104 | 5.464 | 3.523 | 2.438 | 5.068 | 3.348 | 1.134 | Early up | | | | | | | | | | |
| B7L74_03425 | ISAs1 family transposase | | 2.244 | 0.881 | 1.259 | 0.785 | 2.707 | 4.269 | 1.979 | 3.800 | 4.582 | 1.523 | 1.946 | Continuous up | | | | | | | | | | |
| B7L74_03430 | hypothetical protein(CypA in Table 1 of RightonQ par | Effector protein | 1.088 | 0.580 | -2.210 | 0.298 | 2.179 | 2.985 | 0.991 | 3.349 | 3.646 | 0.667 | 0.257 | Early up | | Yes | | | | | | | | |
| B7L74_03435 | succinyl-diaminopimelate desuccinylase | Lysine biosynthesis | -1.050 | -1.517 | -1.888 | 0.107 | -1.198 | -1.080 | -0.998 | -0.849 | -0.682 | -0.994 | -1.040 | Early down | | | | | | | | | | |
| B7L74_03440 | 2%2C3%2C4%2C5-tetrahydropyridine-2%2C6-dicarb | | -0.962 | -0.956 | 0.502 | 1.290 | -1.013 | -1.336 | -0.693 | 0.264 | 0.018 | 0.492 | 0.282 | | | | | | | | | | | |
| B7L74_03445 | hypothetical protein | | 0.266 | 1.797 | 2.628 | 0.475 | -0.633 | 0.009 | 1.027 | 2.721 | 2.008 | 2.979 | 0.943 | Late up | | | | | | | | | | |
| B7L74_03450 | hypothetical protein | | 0.090 | 0.848 | 2.408 | -0.078 | 3.847 | 2.589 | 0.527 | 3.373 | 2.628 | 1.261 | 1.582 | | | | | | | | | | | |
| B7L74_03455 | tRNA-Pro | tRNA | 0.160 | -1.314 | 2.453 | -0.034 | -0.078 | -0.585 | 5.064 | 1.122 | 0.583 | 0.734 | 2.771 | Up in P31 | | | | | | | | | | |
| B7L74_03460 | ATP-dependent RNA helicase RhlE | | 0.414 | -0.619 | -1.091 | 2.071 | 1.456 | 0.330 | 1.058 | 2.269 | 2.863 | 0.514 | 0.722 | | | | | | | | | | | |
| B7L74_03465 | tRNA-Arg | tRNA | 0.440 | -0.283 | -0.263 | -1.895 | 0.149 | -0.436 | 0.230 | -1.095 | -0.502 | -0.830 | -0.389 | | | | | | | | | | | |
| B7L74_03470 | tRNA-His | tRNA | -0.458 | -0.583 | 1.451 | 0.431 | -0.622 | -0.740 | 0.714 | -1.262 | -0.179 | -0.170 | 0.665 | | | | | | | | | | | |
| B7L74_03475 | mannose-1-phosphate guanylyltransferase/mannose-6- | O-Antigen nucleotide sugar biosyn | -1.560 | -2.438 | -1.749 | -0.873 | -1.972 | -1.055 | -1.305 | -1.072 | -0.664 | -1.400 | -1.037 | Early down | | | | | | | | | Yes | |
| B7L74_03480 | hypothetical protein | | -2.433 | -0.949 | -5.492 | -0.352 | -2.534 | -6.273 | -5.000 | 0.768 | 1.155 | -4.190 | -3.408 | Early down | | | | | | | | | | |
| B7L74_03485 | histidinol phosphate phosphatase | Lipopolysaccharide biosynthesis | -3.834 | -1.263 | -7.431 | -0.942 | -2.744 | -8.212 | -1.723 | -1.057 | 0.058 | -4.793 | -3.043 | Early down | | | | | | | | | Yes | |
| B7L74_03490 | sugar isomerase | Lipopolysaccharide biosynthesis | -0.471 | 1.170 | 1.977 | -0.621 | 2.738 | -1.389 | 4.906 | 1.564 | 2.420 | -0.225 | 0.083 | | | | | | | | | | Yes | |
| B7L74_03495 | transaldolase | | -1.167 | -1.785 | -2.359 | -0.144 | -1.593 | -0.251 | -1.977 | -0.701 | 0.504 | -1.892 | -2.506 | | | | | | | | | | | |
| B7L74_03500 | UDP-glucose 4-epimerase | | 0.794 | -1.814 | -1.945 | -0.668 | 2.373 | 3.228 | -1.190 | -0.843 | 2.391 | -0.509 | -2.268 | | | | | | | | | | | |
| B7L74_03505 | NAD-dependent dehydratase | O-Antigen nucleotide sugar biosyn | -0.903 | -1.692 | -3.181 | -0.310 | -1.276 | -1.827 | -1.581 | -1.470 | -0.869 | -2.096 | -1.452 | Down in P10 | | | | | | | | | Yes | |
| B7L74_03510 | ADP-heptose synthase | LPS phase II / virenose biosynthesis | -5.561 | -3.549 | -7.346 | -4.748 | -5.167 | -4.594 | -3.404 | -4.751 | -4.538 | -3.473 | -4.298 | Early down | | | | | | | | | | |
| B7L74_03515 | hypothetical protein | | 0.321 | 0.125 | 1.887 | 1.774 | 0.176 | -0.475 | 1.152 | 2.354 | 0.905 | 0.043 | 0.144 | Early up | | | | | | | | | | |
| B7L74_03520 | sulfotransferase | | 0.422 | -0.634 | 0.370 | 1.613 | 0.600 | 0.212 | 0.806 | -0.225 | 0.769 | 0.530 | 0.916 | | | | | | | | | | | |
| B7L74_03525 | adenylyltransferase | | -0.529 | -1.291 | -1.306 | -0.350 | -1.876 | -1.167 | -0.550 | -0.454 | -0.066 | -0.715 | -0.947 | | | | | | | | | | | |
| B7L74_03530 | 3'(2')%2C5'-bisphosphate nucleotidase CysQ | | -0.077 | -0.600 | 1.932 | -6.129 | 0.254 | -0.559 | 0.877 | -2.594 | -1.769 | -0.685 | -3.589 | | | | | | | | | | | |
| B7L74_03535 | short-chain dehydrogenase | | -0.559 | -0.861 | -0.676 | 1.041 | -2.933 | -0.943 | -0.368 | 0.795 | 1.366 | -0.185 | -0.306 | | | | | | | | Yes | | | |
| B7L74_03540 | sugar ABC transporter permease | ABC transporters | 0.988 | -1.286 | -0.310 | -1.461 | 3.497 | 3.969 | -0.014 | 1.119 | 4.092 | 1.048 | -0.783 | Early up | | | | | | | | Yes (Outer membrane/LPS) | | |
| B7L74_03545 | ABC transporter ATP-binding protein | Transporter | -1.193 | -2.244 | -4.629 | -1.468 | -2.003 | -1.114 | -1.474 | 0.004 | 0.399 | -1.934 | -1.985 | Early down | | | | | | | | Yes (Outer membrane/LPS) | | |
| B7L74_03550 | tRNA-Lys | tRNA | 0.704 | 0.677 | 3.293 | 2.089 | 2.442 | 1.851 | 3.052 | 2.961 | 3.533 | 1.552 | 1.697 | Late up | | | | | | | | | | |
| B7L74_03555 | hypothetical protein | | 0.321 | 1.292 | 0.130 | 1.769 | 1.039 | 2.263 | 0.312 | 4.264 | 4.877 | 0.424 | 1.228 | Late up | | | | | | | | | | |
| B7L74_03560 | hypothetical protein | | -0.072 | 0.368 | 2.079 | 1.460 | 0.888 | 0.676 | 0.572 | 1.710 | 1.538 | 0.802 | 1.570 | Early up | | | | | | | | | | |
| B7L74_03565 | permease | Transporter | 1.835 | 1.801 | 1.678 | 1.542 | 2.545 | 2.649 | 3.038 | 4.695 | 5.327 | 1.274 | 3.186 | Late up | | | | | | | | Yes (General) | | |
| B7L74_03570 | hypothetical protein | | 1.379 | 2.459 | -0.034 | -0.072 | 2.737 | 2.589 | 0.527 | 3.960 | 4.293 | 0.470 | 1.167 | | | | | | | | | | | |
| B7L74_03575 | transposase | Replication and repair | -0.671 | -0.662 | -2.162 | -0.482 | -0.210 | 0.850 | -0.587 | 3.187 | 3.663 | -1.173 | 0.003 | Late up | | | | | | | | | | |
| B7L74_03580 | hypothetical protein | | 0.551 | 1.376 | 1.551 | 0.131 | 3.416 | 2.683 | 2.074 | 2.767 | 3.693 | -1.269 | 1.288 | | | | | | | | | | | |
| B7L74_03585 | hypothetical protein | | 0.360 | 0.123 | 1.306 | 1.630 | 0.947 | 0.390 | 1.069 | 2.470 | 2.694 | 0.027 | 1.498 | Late up | | | | | | | | | | |
| B7L74_03590 | hypothetical protein | Purine metabolism | -0.410 | 0.818 | 2.726 | 2.830 | 1.199 | -0.190 | 0.655 | 2.126 | 3.376 | 0.085 | 1.455 | Early up | | | | | | | | | | |
| B7L74_03595 | hypothetical protein | | 0.159 | 1.269 | 4.398 | 3.084 | 2.396 | 1.070 | 2.661 | 2.451 | 4.860 | 1.564 | 2.811 | Early up | | | | | | | | | | |
| B7L74_03600 | DNA-binding response regulator, GacA.1 | Transcriptional regulators two component system GacA-GacS" | -0.965 | 0.607 | -0.571 | 0.145 | -0.782 | -1.040 | -1.273 | -0.025 | -1.521 | -1.434 | -1.611 | | | | | | | | | | | |
| B7L74_03605 | MFS transporter | Transporter | 1.055 | 0.308 | 0.315 | -0.570 | 0.767 | 0.863 | 0.064 | 0.217 | 0.621 | -0.309 | -0.313 | | | | | | | | | Yes (Ions/Phosphate) | | |
| B7L74_03610 | hypothetical protein | | -1.024 | 0.426 | -0.894 | -0.181 | 0.136 | -0.165 | -1.203 | -0.125 | -0.836 | -1.852 | -1.809 | | | | | | | | | | | |
| B7L74_03615 | IS30 family transposase | Replication and repair | 1.679 | 1.364 | 0.365 | -0.100 | 2.382 | 2.804 | 0.640 | 1.502 | 2.381 | 0.580 | 0.063 | Early up | | | | | | | | | | |
| B7L74_03620 | thioredoxin reductase | | 0.264 | 1.061 | 1.030 | 0.013 | 1.386 | 0.562 | 0.873 | 0.819 | 1.974 | 0.704 | 0.954 | | | | | | | | | | | |
| B7L74_03625 | hypothetical protein | | 0.846 | 1.281 | -0.431 | 0.201 | 0.286 | -0.356 | -0.785 | 0.244 | -1.293 | -0.470 | -0.901 | | | | | | | | | | | |
| B7L74_03630 | hypothetical protein | | 0.627 | 0.813 | 0.916 | 0.734 | 0.619 | 0.961 | -0.021 | 2.174 | 1.437 | -0.443 | -0.101 | | | | | | | | | | | |
| B7L74_03635 | hypothetical protein | | 0.594 | 0.770 | 0.982 | 0.868 | 2.565 | 2.447 | 0.148 | 3.278 | 2.484 | -2.253 | 0.468 | | | | | | | | | | | |
| B7L74_03640 | hypothetical protein, Hypothetical membrane associate | Membrane | 1.009 | 2.120 | 3.816 | -0.933 | 2.325 | 1.984 | 1.449 | 1.890 | 2.732 | 1.095 | 0.849 | Early up | | | | | | | | | | |
| B7L74_03645 | hypothetical protein | | 1.307 | 0.838 | -0.034 | -1.805 | 2.643 | 1.621 | 0.449 | 1.487 | 2.549 | -1.279 | -0.910 | | | | | | | | | | | |
| B7L74_03650 | agmatinase | | -0.220 | 0.458 | -0.135 | -1.348 | -0.611 | -0.703 | -1.206 | -0.765 | 0.332 | -0.963 | -0.740 | | | | | | | | | | | |
| B7L74_03655 | deoxyhypusine synthase | | -0.922 | -0.142 | -1.018 | -2.393 | 0.989 | 1.850 | 1.492 | 2.608 | 3.173 | -2.251 | -1.888 | | | | | | | | | | | |
| B7L74_03660 | ornithine decarboxylase | | -1.560 | -0.529 | -2.239 | -0.608 | -3.024 | -1.611 | -1.330 | -1.105 | -1.399 | -0.487 | 0.707 | | | | | | | | | | | |
| B7L74_03665 | IS30 family transposase | Replication and repair | 2.552 | 0.823 | 0.770 | -0.491 | 3.581 | 4.144 | 1.440 | 1.937 | 3.090 | 0.763 | -0.041 | Early up | | | | | | | | | | |
| B7L74_03670 | hypothetical protein | | -1.380 | -0.418 | -2.513 | -0.116 | -0.559 | 0.547 | -1.037 | 0.537 | 0.763 | -1.564 | -0.915 | | | | | | | | | | | |
| B7L74_03675 | hypothetical protein | | -3.480 | 0.284 | -4.073 | 0.215 | -2.067 | -3.071 | -1.435 | -0.320 | -1.022 | -3.005 | -0.136 | | | | | | | | | | | |
| B7L74_03680 | ABC transporter permease | Transporter | 0.200 | -0.509 | -1.408 | -0.393 | -0.718 | -1.363 | -0.976 | -1.364 | -2.017 | -1.500 | -1.507 | | | | | | | | | Yes (Phospholipid) | | |
| B7L74_03685 | ABC transporter ATP-binding protein | Transporter | -0.012 | -0.009 | -0.915 | -0.325 | -0.553 | -0.714 | -1.083 | -0.821 | -1.421 | -1.178 | -1.255 | | | | | | | | | Yes (Phospholipid) | | |
| B7L74_03690 | ABC transporter substrate-binding protein | Transporter | -1.331 | -1.175 | -0.252 | 0.248 | -0.524 | -0.385 | -0.773 | -0.672 | -1.710 | -0.113 | -1.388 | | | | | | | | | Yes (Phospholipid) | | |
| B7L74_03695 | hypothetical protein | | 0.917 | 0.882 | 1.654 | -1.907 | 2.111 | 1.137 | 0.894 | 0.704 | 1.132 | -0.154 | -0.710 | | | | | | | | | Yes (FA/Cholesterol) | | |
| B7L74_03700 | hypothetical exported protein | | 2.530 | 2.859 | 5.036 | -1.064 | -3.549 | 2.075 | 3.794 | 3.302 | 2.420 | 1.960 | 3.896 | | | | | | | | | | | |
| B7L74_03705 | hypothetical protein | | 1.815 | 0.720 | 0.401 | -0.418 | 3.432 | 2.787 | 0.694 | 0.628 | 2.458 | 1.126 | 1.283 | | | | | | | | | | | |
| B7L74_03710 | hypothetical protein | | -0.098 | 0.385 | -1.005 | -1.272 | 0.148 | 0.021 | 0.155 | 1.028 | 0.348 | -0.713 | -0.312 | | | | | | | | | | | |
| B7L74_03715 | uridine phosphorylase | | 2.201 | 0.847 | -0.035 | 4.467 | 3.924 | 1.627 | 0.526 | 4.274 | 3.747 | 4.191 | 1.426 | | | | | | | | | | | |
| B7L74_03720 | hypothetical exported protein | | -0.544 | 0.015 | -1.244 | 0.167 | -0.983 | -0.806 | -0.687 | 0.113 | -0.608 | -1.246 | -1.073 | | | | | | | | | | | |
| B7L74_03725 | tRNA-Leu | tRNA | 1.132 | 0.904 | 1.525 | -1.231 | -0.043 | 0.077 | 1.154 | -0.075 | -1.177 | 0.252 | 0.373 | | | | | | | | | | | |
| B7L74_03730 | trigger factor | | -0.851 | -0.118 | 0.141 | -0.251 | -0.236 | -0.344 | -0.935 | -0.764 | -0.989 | -1.160 | -0.943 | | | | | | | | | | | |
| B7L74_03735 | ATP-dependent Clp protease proteolytic subunit | | -1.008 | -0.185 | -0.244 | 0.354 | -0.456 | -0.433 | -0.999 | -0.763 | -1.480 | -1.158 | -1.221 | | | | | | | | | | | |
| B7L74_03740 | ATP-dependent Clp protease ATP-binding subunit Cl | ATP | -1.962 | -1.023 | -2.350 | -1.856 | -0.855 | -0.857 | -2.328 | -1.686 | -2.146 | -2.246 | -2.382 | Late down | | | | | | | | | | |
| B7L74_03745 | endopeptidase La | Cell cycle - Caulobacter | -0.989 | -0.463 | -1.867 | -2.184 | -0.443 | -0.484 | -1.918 | -1.771 | -1.820 | -2.075 | -2.152 | Early down | | | | | | | | | | |
| B7L74_03750 | metalloprotease PmbA | Peptidases and inhibitors | -0.515 | -0.024 | -2.444 | -1.093 | -0.717 | -1.188 | -1.714 | -1.131 | -2.054 | -1.403 | -1.809 | Early down | | | | | | | | | | |
| B7L74_03755 | hypothetical protein | | -0.989 | -0.686 | -2.851 | -0.255 | 0.073 | -0.505 | -2.258 | -1.223 | -3.743 | -1.690 | -3.355 | Late down | | | | | | | | | | |
| B7L74_03760 | phosphocarrier protein HPr | | -0.448 | 0.481 | 0.924 | 2.877 | -1.484 | -0.749 | 1.496 | 0.795 | 1.017 | 0.648 | 1.365 | | | | | | | | | | | |
| B7L74_03765 | HPr kinase/phosphorylase | | 0.334 | 2.138 | 2.339 | 3.078 | 2.183 | 2.011 | 2.830 | 2.786 | 0.805 | 1.640 | 1.686 | | | | | | | | | | | |
| B7L74_03770 | ribosomal subunit interface protein | | -0.314 | 0.831 | 0.702 | 1.735 | 0.856 | 0.885 | 0.164 | 1.478 | -2.419 | 0.629 | -0.698 | | | | | | | | | | | |
| B7L74_03775 | ABC transporter ATP-binding protein | Transporter | -0.431 | -1.116 | 0.521 | 1.043 | -0.943 | -0.552 | 0.083 | -0.682 | -1.125 | 0.403 | 0.299 | | | | | | | | | Yes (Outer membrane/LPS) | | |
| B7L74_03780 | lipopolysaccharide transport periplasmic protein LptA | | 0.204 | -0.863 | 1.559 | 0.956 | -0.493 | -0.409 | 0.567 | -0.645 | -0.718 | 1.321 | 1.110 | | | | | | | | | Yes (Outer membrane/LPS) | | |
| B7L74_03785 | LPS export ABC transporter periplasmic protein LptC | | 0.437 | -0.463 | 0.314 | 0.952 | -0.568 | -0.437 | 1.030 | -0.493 | -0.196 | 1.114 | 0.561 | | | | | | | | | Yes (Outer membrane/LPS) | | |
| B7L74_03790 | phenylphosphate carboxylase subunit delta | Lipopolysaccharide biosynthesis | 0.420 | -1.043 | -0.367 | 1.660 | -0.749 | -0.790 | 0.414 | -1.158 | -0.746 | 0.819 | 1.495 | | | | | | | | | | Yes | |
| B7L74_03795 | arabinose-5-phosphate isomerase | Lipopolysaccharide biosynthesis | 0.544 | -1.194 | -0.137 | 0.941 | -1.371 | -0.728 | 0.500 | -0.267 | -0.227 | 1.066 | 0.872 | | | | | | | | | | Yes | |
| B7L74_03800 | UDP-N-acetylglucosamine 1-carboxyvinyltransferase | Peptidoglycan syn | 3.086 | 1.135 | 3.110 | 0.933 | 3.341 | 3.549 | 2.799 | 2.106 | 3.043 | 3.398 | 3.028 | Early up | | | | | | | | | | Yes |
| B7L74_03805 | alpha/beta hydrolase | | 7.898 | 3.276 | 5.896 | -1.666 | 11.121 | 11.408 | 5.584 | 1.514 | 10.104 | 3.258 | 1.842 | Continuous up | | | | | | | | | | |
| B7L74_03810 | DUF1658 domain-containing protein | | 0.047 | 0.839 | -0.034 | -1.800 | 2.649 | 1.621 | 0.454 | 1.488 | 1.156 | -1.276 | -0.908 | | | | | | | | | | | |
| B7L74_03815 | multidrug transporter AcrB | Transporter | -0.963 | -0.308 | -0.848 | -3.581 | 4.476 | 3.032 | 0.837 | 0.957 | 6.192 | -2.051 | -2.234 | Early up | | | | | | | | | | |
| B7L74_03820 | MexH family multidrug efflux RND transporter peripl | beta-Lactam resistance | -1.639 | -0.661 | -4.094 | -2.165 | -2.792 | -2.860 | -2.220 | -2.258 | -5.217 | -2.386 | -2.741 | Early down | | | | | | | | | | |
| B7L74_03825 | endopeptidase | | -0.439 | 0.150 | -1.814 | -1.380 | -2.003 | -2.170 | -1.743 | -1.493 | -3.012 | -0.987 | -1.481 | Early down | | | | | | | | | | |
| B7L74_03830 | polyphenol oxidase | | 0.080 | 0.578 | 2.498 | -0.057 | 0.911 | 0.220 | 3.883 | 4.566 | 1.672 | 1.063 | 3.490 | Early up | | | | | | | | | | |
| B7L74_03835 | 23S rRNA pseudouridine(1911/1915/1917) synthase | | -0.712 | 0.068 | -0.106 | 0.248 | -1.390 | -1.443 | -0.441 | -0.697 | -1.648 | -1.424 | -1.119 | | | | | | | | | | | |
| B7L74_03840 | outer membrane protein assembly factor BamD | Membrane Transporter | 1.905 | 0.767 | -0.846 | 2.775 | 0.783 | -0.197 | 1.673 | 1.402 | 1.092 | 2.108 | 1.343 | Up in P12 | | | | | | | | Yes (Outer membrane/Lipoprotein) | | |
| B7L74_03845 | hypothetical protein | | 0.840 | 0.845 | 1.966 | 4.626 | 2.687 | 1.625 | 5.416 | 6.183 | 3.991 | 2.313 | 2.806 | Continuous up | | | | | | | | | | |
| B7L74_03850 | hypothetical protein | | 0.072 | 0.844 | -0.034 | -1.778 | 0.294 | 0.972 | 0.480 | 1.493 | 1.877 | -1.262 | -0.901 | | | | | | | | | | | |
| B7L74_03855 | response regulator, GacS | Signal transductiontwo component | -1.875 | 0.539 | -1.301 | 1.529 | 0.693 | -0.445 | -0.803 | -0.230 | 0.581 | -0.276 | 1.504 | | | | | | | | | | | |
| B7L74_03860 | hypothetical protein | | 2.431 | 0.247 | 1.248 | 2.998 | -0.097 | -0.770 | 2.549 | 0.027 | -0.346 | 3.068 | 4.063 | Continuous up | | | | | | | | | | |
| B7L74_03865 | transcriptional regulator | | 1.269 | -0.289 | -0.736 | 1.898 | -0.984 | -2.430 | 0.576 | -1.998 | -1.916 | 1.420 | 2.315 | | | | | | | | | | | |
| B7L74_03870 | hypothetical membrane spanning protein | | 0.037 | 0.837 | -0.034 | -0.389 | 1.973 | 1.620 | 0.444 | 1.486 | 2.544 | -1.282 | -0.911 | | | | | | | | | | | |
| B7L74_03875 | hypothetical protein | Fatty acid degradation | 0.619 | 0.407 | 0.612 | 1.147 | 0.521 | 1.391 | 1.148 | 2.119 | 2.215 | 1.771 | 1.337 | Late up | | | | | | | | | | |
| B7L74_03880 | long-chain-fatty-acid--CoA ligase | Fatty acid biosynthesis | 0.100 | 0.850 | -0.034 | -1.751 | 2.506 | 5.000 | 4.732 | 4.520 | 4.011 | -1.247 | -0.411 | Early up | | | | | | | | | | |
| B7L74_03885 | acetoacetyl-CoA synthetase | | -0.448 | 0.329 | -1.218 | -0.907 | -0.290 | 0.316 | -1.029 | 0.058 | -0.027 | -0.941 | -0.937 | | | | | | | | | | | |
| B7L74_03890 | tRNA-Val | tRNA | -1.209 | 0.158 | -0.331 | 0.118 | -2.587 | 0.449 | 0.321 | 1.169 | -0.484 | -1.892 | -1.457 | | | | | | | | | | | |
| B7L74_03895 | hypothetical protein | | 0.862 | 0.066 | -1.575 | 0.214 | -0.214 | 1.147 | -3.456 | 2.910 | 2.762 | -2.501 | -1.580 | | | | | | | | | | | |
| B7L74_03900 | hypothetical protein | | -1.453 | 0.482 | -5.978 | -1.386 | 0.225 | 0.195 | -1.454 | 0.724 | 1.551 | -1.374 | -2.723 | | | | | | | | | | | |
| B7L74_03905 | multidrug transporter | Transporter | 0.188 | 1.270 | 0.150 | 0.119 | 2.450 | 3.292 | 0.919 | 3.572 | 3.943 | -0.191 | -0.024 | Early up | | | | | | | | Yes (drug resistance) | | |
| B7L74_03910 | GMP synthase | | -0.144 | 0.205 | -0.373 | 0.394 | 0.449 | 0.376 | 0.022 | 0.218 | 0.593 | -1.274 | -0.689 | | | | | | | | | | | |
| B7L74_03915 | ATPase | ATP | -0.996 | -0.215 | -1.608 | -0.195 | -1.458 | -0.119 | -1.168 | 0.602 | 1.031 | -0.918 | -1.051 | | | | | | | | | | | |
| B7L74_03920 | methylisocitrate lyase | | -0.731 | 0.285 | -1.226 | -0.076 | -0.587 | -0.224 | -1.536 | -0.558 | -1.314 | -1.995 | -1.569 | | | | | | | | | | | |
| B7L74_03925 | citrate synthase/methylcitrate synthase | Propanoate metabolism | -4.527 | -0.551 | -2.574 | 0.029 | -1.373 | -1.110 | -3.718 | -1.619 | -1.190 | -3.902 | -2.534 | Late down | | | | | | | | | | |
| B7L74_03930 | glyoxalase | Effector protein | -1.056 | 0.414 | -0.401 | 0.949 | -0.701 | -0.381 | -1.275 | -0.483 | -1.486 | -1.069 | -1.077 | | | Yes | | | | | | | | |
| B7L74_03935 | hypothetical proteinpspC (PspC domain containing pro | Transcriptional regulators | -1.092 | 0.467 | 3.023 | 2.107 | 0.195 | 0.485 | 1.129 | 0.828 | 2.558 | 1.708 | 2.093 | Early up | | | | | | | | | | |
| B7L74_03940 | GntR family transcriptional regulator | Transcriptional regulators | -0.700 | -0.033 | 2.058 | 1.682 | -0.026 | 0.107 | 0.465 | 1.095 | 2.145 | 1.376 | 1.429 | | | | | | | | | | | |
| B7L74_03945 | ABC transporter ATP-binding protein | Transporter | -1.154 | -0.170 | -1.546 | -0.852 | -0.421 | -0.432 | -1.418 | -0.049 | 0.038 | -1.412 | -1.463 | | | | | | | | | Yes (General) | | |
| B7L74_03950 | ABC transporter permease | Transporter | -0.013 | 0.456 | 0.282 | -1.268 | -0.742 | 0.357 | -0.699 | -1.377 | -1.259 | -1.456 | -2.182 | | | | | | | | | Yes (Outer membrane/Membrane) | | |
| B7L74_03955 | protease | | -0.666 | 0.066 | -1.025 | -0.535 | 0.459 | -0.031 | -1.567 | -1.111 | -0.352 | -0.926 | -0.290 | | | | | | | | | | | |
| B7L74_03960 | hypothetical protein | | 4.878 | 0.746 | 1.803 | 0.327 | 5.716 | 6.002 | 2.145 | 2.919 | 5.305 | 2.542 | 1.156 | Early up | | | | | | | | | | |
| B7L74_03965 | DNA-binding response regulator (gene=sirA) | Transcriptional regulators | -1.113 | -0.324 | -0.483 | 0.687 | -1.454 | -1.755 | -1.263 | -1.166 | -1.365 | -0.998 | -0.864 | | | | | | | | | | | |
| B7L74_03970 | hypothetical protein ankyrin repeat protein AnkG | Effector protein | -1.342 | -0.419 | -2.295 | -0.094 | -1.106 | -0.941 | -1.254 | -0.258 | -0.156 | -1.663 | -1.512 | Early down | | Yes | | | | | | | | |
| B7L74_03975 | hypothetical protein | | -0.676 | -0.362 | -1.894 | -1.059 | -0.526 | 0.401 | -0.691 | 1.673 | 1.860 | -1.508 | -1.267 | | | | | | | | | | | |
| B7L74_03980 | hypothetical protein | beta-Lactam resistance | 1.344 | -0.508 | -3.268 | -0.222 | 2.193 | 2.561 | -0.075 | 2.532 | 3.259 | 0.019 | -0.276 | Early up | | | | | | | | | | |
| B7L74_03985 | hypothetical protein | | 0.713 | 0.655 | 0.028 | -1.383 | 2.334 | 1.815 | 0.260 | 0.774 | 1.768 | 0.575 | -0.314 | | | | | | | | | | | |
| B7L74_03990 | hypothetical protein | | -1.570 | 0.020 | 0.409 | -0.135 | -0.951 | -0.386 | -0.354 | -0.492 | 0.245 | -0.483 | 0.395 | | | | | | | | | | | |
| B7L74_03995 | CoA transferase | | 0.092 | 1.494 | 2.184 | 0.240 | 0.813 | -0.019 | 0.165 | 1.189 | 0.616 | 0.376 | 0.663 | | | | | | | | | | | |
| B7L74_04000 | AMP-binding protein | | -1.562 | -0.298 | -2.711 | -2.323 | -0.321 | -0.233 | -2.591 | -0.625 | -1.331 | -2.496 | -2.261 | Early down | | | | | | | | | | |
| B7L74_04005 | polyketide synthase | | -1.919 | 0.289 | -2.803 | -2.852 | 0.976 | 1.811 | -0.395 | 1.024 | 1.952 | -2.077 | -3.029 | Early down | | | | | | | | | | |
| B7L74_04010 | hybrid sensor histidine kinase/response regulator | Kinase, Two-component system | -1.157 | -0.338 | -0.569 | -0.174 | -0.665 | -0.791 | -1.556 | -0.047 | -0.996 | -2.092 | -2.470 | Late down | | | | | | | | | | |
| B7L74_04015 | hypothetical protein | | 0.278 | 0.595 | 0.569 | -0.568 | -0.196 | 1.052 | -0.388 | 2.076 | 2.277 | -1.445 | -0.430 | | | | | | | | | | | |
| B7L74_04020 | hypothetical protein | | -0.444 | 0.352 | -1.316 | -2.176 | 0.375 | 1.355 | -0.937 | 1.906 | 2.508 | -1.868 | -1.253 | | | | | | | | | | | |
| B7L74_04025 | hypothetical protein | | 3.450 | 0.794 | 1.576 | -0.288 | 4.752 | 4.796 | 2.125 | 3.383 | 4.662 | 0.828 | 0.395 | Early up | | | | | | | | | | |
| B7L74_04030 | MFS transporter | Transporter | 0.232 | 0.153 | 0.159 | 0.478 | 0.339 | 2.450 | 0.886 | 3.756 | 3.892 | -0.423 | 0.361 | Late up | | | | | | | | Yes (MFS) | | |
| B7L74_04035 | hypothetical protein | | -0.615 | 0.837 | 1.164 | 1.912 | 0.944 | 3.193 | 1.355 | 4.048 | 4.157 | 0.501 | 0.568 | Late up | | | | | | | | | | |
| B7L74_04040 | hypothetical protein | | 1.352 | -0.276 | -1.316 | -2.242 | -0.313 | -2.135 | 0.481 | 2.768 | 2.026 | -2.741 | -2.195 | | | | | | | | | | | |
| B7L74_04045 | hypothetical protein | Effector protein | -0.678 | 0.474 | -0.630 | -0.400 | 0.796 | 0.727 | 0.698 | 1.949 | 1.931 | -0.303 | -0.096 | | | Yes | | | | | | | | |
| B7L74_04050 | tRNA-Leu | tRNA | -0.540 | 0.089 | -1.391 | 0.300 | -1.061 | -0.910 | -0.469 | -1.387 | -1.569 | -0.812 | -0.504 | | | | | | | | | | | |
| B7L74_04055 | GTP cyclohydrolase I FolE | Folate biosynthesis | -1.114 | -1.995 | -1.087 | 0.154 | -2.106 | -1.884 | -1.034 | -1.959 | -1.830 | -0.861 | -0.311 | Early down | | | | | | | | | | |
| B7L74_04060 | histidine triad nucleotide-binding protein | | 0.783 | 0.193 | 1.991 | 0.858 | 1.111 | 0.467 | 1.602 | 1.853 | 1.490 | -0.480 | -1.326 | | | | | | | | | | | |
| B7L74_04065 | MFS transporter Multidrug resistance protein B | Resistance | -1.995 | -2.153 | -2.326 | -1.236 | -3.406 | -1.815 | -1.676 | -1.506 | -0.941 | -1.623 | -1.174 | | | | | | | | | Yes (drug resistance) | | |
| B7L74_04070 | multidrug resistance protein A | Transporter | -1.417 | -1.994 | -2.244 | -0.427 | -3.295 | -2.670 | -1.555 | -2.247 | -2.727 | -1.392 | -1.226 | Early down | | | | | | | | Yes (drug resistance) | | |
| B7L74_04075 | hypothetical protein | | 0.660 | -0.032 | 0.410 | 0.070 | -2.594 | -1.093 | -0.067 | 0.704 | -0.017 | 0.806 | 1.522 | | | | | | | | | | | |
| B7L74_04080 | hypothetical protein | | -0.744 | -1.513 | -0.862 | -0.041 | -1.327 | -0.609 | -0.318 | -0.419 | -1.281 | -0.524 | -0.572 | | | | | | | | | | | |
| B7L74_04085 | ribosomal-protein-alanine N-acetyltransferase RimI | Effector protein | -2.102 | -1.323 | -2.200 | -0.635 | 3.026 | 0.287 | -1.317 | -0.674 | 1.150 | -0.025 | -0.437 | | | Yes | | | | | | | | |

| | | | | | | | | | | | | | | | | | | | | | | | |
|---|---|---|---|---|---|---|---|---|---|---|---|---|---|---|---|---|---|---|---|---|---|---|---|
| B7L74_04090 | hypothetical protein | | -1.465 | -0.652 | -2.474 | -1.130 | -1.766 | -1.833 | -1.778 | -1.711 | -1.933 | 0.306 | -0.669 | | | | | | | | | | |
| B7L74_04095 | MexH family multidrug efflux RND transporter peripla | beta-Lactam resistance | -0.775 | -0.627 | -1.325 | 0.869 | -1.747 | -1.578 | -1.129 | -1.846 | -2.571 | -1.300 | -0.849 | Early down | | | | | | | | | |
| B7L74_04100 | multidrug transporter AcrB | Transporter | -2.102 | -1.490 | -3.701 | -0.527 | -2.006 | -1.452 | -2.113 | -1.903 | -1.655 | -2.799 | -2.344 | Early down | | | | | | | | | |
| B7L74_04105 | hypothetical protein | | 0.620 | 0.280 | -0.138 | -1.509 | 0.238 | -0.362 | 0.113 | -0.078 | -0.690 | 0.748 | 0.048 | | | | | | | | | | |
| B7L74_04110 | hypothetical protein | | 0.742 | 0.923 | -0.713 | -0.754 | -0.273 | -0.840 | -0.459 | 0.879 | -0.815 | 0.671 | 0.742 | | | | | | | | | | |
| B7L74_04115 | serine hydrolase (B-lactamase) | | 0.699 | -0.454 | -0.182 | 1.333 | -0.345 | -0.369 | 0.437 | 0.003 | -0.058 | 0.492 | 0.925 | | | | | | | | | | |
| B7L74_04120 | valine--tRNA ligase | | -1.031 | -1.247 | -1.542 | -0.389 | -1.915 | -0.951 | -0.563 | -0.491 | -0.404 | -0.120 | 0.185 | | | | | | | | | | |
| B7L74_04125 | LPS export ABC transporter permease LptF | | 0.117 | -1.164 | -0.316 | 0.905 | -2.571 | -0.407 | 0.307 | -0.487 | -0.266 | 0.493 | 1.651 | | | | | | | | | Yes (Outer membrane/LPS) | |
| B7L74_04130 | LPS export ABC transporter permease LptG | | 0.551 | 0.251 | 0.955 | -1.710 | 3.642 | 3.811 | 1.779 | 3.820 | 1.776 | -1.324 | -1.493 | | | | | | | | | Yes (Outer membrane/LPS) | |
| B7L74_04135 | peptide chain release factor 3 | Translation factors | -1.047 | -0.584 | -1.916 | -0.514 | -3.203 | -0.588 | -1.196 | -0.409 | -0.173 | -1.510 | -1.313 | Early down | | | | | | | | | |
| B7L74_04140 | MATE family efflux transporter | | -0.704 | -0.363 | 0.098 | -0.921 | -0.348 | 0.237 | 0.492 | 1.964 | 1.171 | -1.160 | -0.887 | | | | | | | | | Yes (drug resistance) | |
| B7L74_04145 | DUF4145 domain-containing protein | | -1.088 | -0.450 | -1.861 | -0.553 | 0.409 | 1.687 | -1.991 | 0.746 | 1.469 | 0.176 | -0.933 | | | | | | | | | | |
| B7L74_04150 | hypothetical protein | | 0.239 | -0.491 | -1.375 | -0.062 | 2.480 | 1.689 | 0.919 | 2.550 | 3.760 | 0.715 | 0.151 | | | | | | | | | | |
| B7L74_04155 | hypothetical protein | | 0.563 | -0.208 | -2.477 | -1.749 | 1.971 | 0.936 | 2.740 | 2.650 | 3.433 | -1.245 | 1.038 | | | | | | | | | | |
| B7L74_04160 | hypothetical protein | | 0.105 | -1.375 | -3.760 | 0.600 | 0.571 | 1.902 | -2.229 | 2.978 | 3.894 | -1.120 | -2.179 | | | | | | | | | | |
| B7L74_04165 | N-acetyltransferase | | 0.956 | 0.951 | 0.508 | -0.285 | 3.131 | 2.914 | 0.110 | 1.120 | 1.973 | 0.644 | -0.008 | Early up | | | | | | | | | |
| B7L74_04170 | TetR family transcriptional regulator | Transcriptional regulators | -0.875 | 0.114 | -0.082 | -0.614 | -1.694 | -0.684 | -0.819 | -0.446 | -2.050 | -0.141 | -0.150 | Down in P51 | | | | | | | | | |
| B7L74_04175 | glutathione S-transferase | Glutathione metabolism | -2.655 | 0.522 | -0.855 | 0.040 | -1.348 | -0.603 | -1.321 | -0.426 | -2.805 | -0.684 | -0.817 | Down in P51 | | | | | | | | | |
| B7L74_04180 | hypothetical protein | | -1.642 | -0.229 | -1.825 | 0.220 | -1.061 | -0.507 | -0.758 | 0.105 | -0.544 | -0.635 | -0.478 | | | | | | | | | | |
| B7L74_04185 | cell filamentation protein Fic | | -1.069 | -2.218 | -0.653 | -0.309 | -2.539 | -1.585 | -0.319 | -0.092 | 0.265 | -0.123 | -0.749 | | | | | | | | | | |
| B7L74_04190 | hypothetical protein | | -1.282 | -0.508 | -4.653 | -2.548 | -1.882 | -1.191 | -4.091 | 2.262 | 2.764 | -2.146 | -1.045 | | | | | | | | | | |
| B7L74_04195 | NAD-dependent malic enzyme | | -0.583 | -1.161 | -1.748 | -0.793 | -1.235 | -0.907 | -1.554 | -0.781 | -0.393 | -1.694 | -1.178 | | | | | | | | | | |
| B7L74_04200 | adenylosuccinate lyase | Purine metabolism | -1.281 | -1.591 | -1.774 | -0.955 | -1.692 | -1.086 | -1.038 | -0.759 | -0.582 | -1.586 | -1.443 | Early down | | | | | | | | | |
| B7L74_04205 | aminotransferase DegT | O-Antigen nucleotide sugar biosyn | 0.133 | -0.873 | -0.313 | -0.197 | -1.023 | -0.692 | -0.581 | -0.480 | -0.838 | -0.588 | 0.146 | | | | | | | | | | Yes |
| B7L74_04210 | transcription/translation regulatory transformer protein | | 0.049 | 0.839 | -0.034 | -1.798 | 1.973 | 2.084 | 0.456 | 1.489 | 3.985 | -1.275 | -0.908 | | | | | | | | | | |
| B7L74_04215 | hypothetical protein | | -1.698 | -2.733 | -0.967 | 0.133 | -2.283 | -0.534 | -0.565 | 0.031 | 0.093 | 0.500 | -0.022 | | | | | | | | | | |
| B7L74_04220 | oxidoreductase | O-Antigen nucleotide sugar biosyn | -0.884 | -1.729 | -1.463 | -0.778 | -2.195 | -1.715 | -0.987 | -1.688 | -1.433 | -1.081 | -0.653 | Early down | | | | | | | | | Yes |
| B7L74_04225 | NAD-dependent dehydratase | O-Antigen nucleotide sugar biosyn | -0.849 | -1.267 | -2.840 | -1.216 | -1.932 | -2.085 | -1.666 | -2.278 | -2.813 | -1.750 | -1.480 | Early down | | | | | | | | | Yes |
| B7L74_04230 | perosamine synthetase | O-Antigen nucleotide sugar biosyn | 0.772 | 1.630 | 1.371 | 1.867 | 2.418 | 2.296 | 1.995 | 1.319 | 2.348 | -0.828 | 0.807 | | | | | | | | | | Yes |
| B7L74_04235 | asparagine synthetase B | Ligase Alanine, aspartate and gluta | -2.034 | -1.277 | -4.074 | -1.722 | -1.839 | -1.302 | -2.028 | -1.584 | -1.430 | -2.017 | -2.018 | Down in P10 | | | | | | | | | |
| B7L74_04240 | acetyltransferase | | -0.888 | -0.940 | -2.686 | -1.563 | -2.073 | -1.540 | -1.165 | -1.885 | -1.338 | -1.524 | -0.584 | Early down | | | | | | | | | |
| B7L74_04245 | ABC transporter permease | ABC transporters | -0.925 | -0.588 | -4.284 | -2.322 | 0.934 | 0.249 | -1.560 | -1.260 | -0.450 | -2.485 | -3.110 | Early down | | | | | | | | Yes (Outer membrane/Membrane) | |
| B7L74_04250 | SAM-dependent methyltransferase | Methylation | -2.538 | -1.092 | -3.906 | -1.964 | -1.693 | -1.581 | -2.489 | -2.350 | -2.083 | -2.714 | -2.754 | Early down | | | | | | | | | |
| B7L74_04255 | hypothetical protein | | -1.185 | -1.186 | -3.845 | -1.679 | -1.754 | -1.604 | -1.501 | -2.116 | -2.309 | -1.934 | -1.339 | Early down | | | | | | | | | |
| B7L74_04260 | radical SAM protein | | 0.114 | -1.195 | -2.985 | -2.098 | 0.525 | 0.405 | -1.722 | -2.737 | -0.072 | -1.938 | -2.000 | | | | | | | | | | |
| B7L74_04265 | hypothetical membrane spanning protein | | 3.230 | 0.898 | 0.602 | -1.783 | 5.740 | 5.777 | 0.534 | 1.458 | 4.521 | 1.255 | -0.860 | Early up | | | | | | | | | |
| B7L74_04270 | lipopolysaccharide N-acetylglucosaminyltransferase | | 1.707 | -0.232 | -1.286 | -1.027 | 3.162 | 1.909 | 0.702 | -0.550 | 1.088 | -0.685 | -0.422 | | | | | | | | | | |
| B7L74_04275 | glycosyl transferase | | -1.386 | -0.862 | -2.799 | -1.827 | -1.001 | -0.871 | -1.668 | -0.814 | -0.898 | -2.189 | -2.261 | Early down | | | | | | | | | |
| B7L74_04280 | asparagine synthetase B | | -0.786 | -0.731 | -2.297 | -2.070 | -0.065 | -0.285 | -2.475 | -1.158 | 2.030 | -1.633 | -2.691 | | | | | | | | | | |
| B7L74_04285 | glycosyltransferase family 1 protein | | -1.484 | -0.897 | -4.948 | -2.498 | -1.151 | -0.855 | -2.118 | -1.254 | -0.759 | -2.977 | -2.767 | Early down | | | | | | | | | |
| B7L74_04290 | UDP-N-acetylglucosamine 2-epimerase (non-hydrolyz | O-Antigen nucleotide sugar biosyn | -1.755 | -0.531 | -4.695 | -2.080 | -1.012 | -0.581 | -2.261 | -0.417 | -0.760 | -2.575 | -2.488 | Down in P10 | | | | | | | | | Yes |
| B7L74_04295 | glycosyl transferase | | -0.664 | -0.513 | -1.350 | -1.492 | -0.774 | -0.631 | -0.570 | -0.628 | 0.094 | -1.561 | -1.078 | | | | | | | | | | |
| B7L74_04300 | NAD-dependent epimerase | O-Antigen nucleotide sugar biosyn | -0.321 | -0.860 | -2.589 | -1.025 | -1.122 | -1.119 | -1.651 | -1.791 | -2.024 | -2.170 | -1.678 | Late down | | | | | | | | | Yes |
| B7L74_04305 | GDP-mannose dehydrogenase | O-Antigen nucleotide sugar biosyn | 1.334 | 0.623 | -1.513 | 0.022 | 0.657 | -0.435 | 2.336 | 0.773 | 2.941 | -0.466 | -0.209 | | | | | | | | | | Yes |
| B7L74_04310 | UDP-glucose 6-dehydrogenase | O-Antigen nucleotide sugar biosyn | -1.068 | -0.299 | -1.534 | -0.816 | -0.814 | -0.417 | -1.026 | -0.488 | -0.903 | -0.729 | -0.776 | | | | | | | | | | Yes |
| B7L74_04315 | 3-hydroxyacyl-CoA dehydrogenase | | -2.181 | -1.257 | -4.190 | -2.263 | -2.132 | -0.859 | -1.776 | -1.442 | -1.306 | -2.442 | -2.241 | Early down | | | | | | | | | |
| B7L74_04320 | glucose-6-phosphate isomerase | | 0.882 | -0.417 | -0.878 | -0.873 | 1.796 | 2.127 | 0.110 | -1.132 | 1.263 | -0.600 | -0.921 | | | | | Yes | | | | | |
| B7L74_04325 | UTP--glucose-1-phosphate uridylyltransferase | O-Antigen nucleotide sugar biosyn | 0.742 | 0.246 | -0.629 | 0.143 | 2.067 | 1.785 | 1.301 | 1.726 | 1.970 | -1.865 | 0.535 | | | | | | | | | | Yes |
| B7L74_04330 | hypothetical membrane spanning protein | | 0.724 | -0.615 | 1.268 | -0.389 | 0.253 | 0.797 | 1.344 | 1.695 | 1.979 | 1.791 | 1.328 | | | | | | | | | | |
| B7L74_04335 | 30S ribosomal protein S15 | Ribosomal proteins | -0.675 | -2.462 | -0.170 | 3.657 | -3.619 | 0.620 | 0.816 | 0.843 | 1.048 | 0.618 | 1.765 | Up in P12 | | | | | | | | | |
| B7L74_04340 | polyribonucleotide nucleotidyltransferase | RNA degradation | -0.812 | -1.402 | -2.730 | 2.164 | -1.778 | -0.604 | -0.803 | -0.354 | -0.061 | -1.289 | -0.188 | Variable | | | | | | | | | |
| B7L74_04345 | methylglyoxal synthase | Propanoate metabolism | 0.090 | -0.888 | 0.906 | 2.675 | -0.480 | 0.426 | 0.915 | 0.481 | -0.228 | 1.262 | 1.309 | Up in P12 | | | | | | | | | |
| B7L74_04350 | hypothetical protein | | -0.903 | 1.471 | -3.018 | -1.288 | 0.824 | -1.356 | -2.456 | 2.830 | 2.642 | -0.569 | -0.408 | | | | | | | | | | |
| B7L74_04355 | hypothetical protein | Secretion system | 1.132 | -1.015 | 0.960 | 1.606 | 1.402 | 1.441 | 1.225 | 2.455 | 3.505 | 2.186 | 1.691 | Late up | | | | | | | | | |
| B7L74_04360 | hypothetical protein | Secretion system | 1.557 | 0.968 | 0.959 | -0.142 | 4.047 | 4.608 | 2.341 | 3.347 | 4.713 | 1.834 | 1.331 | Continuous up | | | | | | | | | |
| B7L74_04365 | DNA internalization-related competence protein ComE | | -0.340 | -1.132 | -0.802 | 1.869 | -0.713 | -0.225 | 1.144 | -0.321 | 0.059 | 0.979 | 2.285 | | | | | | | | | | |
| B7L74_04370 | lipid A export permease/ATP-binding protein MsbA | | 0.088 | -1.103 | -0.492 | 1.344 | -0.466 | -0.349 | 0.570 | -0.855 | -0.727 | 0.355 | 1.190 | | | | | | | | | Yes (Outer membrane/LPS) | |
| B7L74_04375 | tetraacyldisaccharide 4'-kinase | Lipopolysaccharide biosynthesis | -0.180 | -1.010 | -1.926 | -0.453 | 0.733 | -0.591 | -1.126 | -0.276 | 1.264 | -0.918 | -2.042 | | | | | | | | | | Yes |
| B7L74_04380 | NAD+ synthase | Nicotinate and nicotinamide metab | -3.599 | -1.683 | -1.271 | -3.732 | -3.811 | -1.886 | -2.288 | -1.547 | -0.684 | -4.348 | -4.855 | Late down | | | | | | | | | |
| B7L74_04385 | hypothetical membrane spanning protein | | -0.834 | -1.183 | 1.458 | 1.027 | -1.740 | -0.457 | 0.599 | 0.206 | 0.522 | -0.278 | 0.151 | | | | | | | | | | |
| B7L74_04390 | hypothetical protein | | 2.527 | 1.295 | 2.271 | 1.444 | 2.721 | 3.829 | 3.533 | 4.146 | 4.375 | 1.862 | 2.144 | Early up | | | | | | | | | |
| B7L74_04395 | hypothetical protein | | 0.840 | -0.447 | 1.041 | 1.134 | 1.324 | 2.435 | 1.852 | 2.295 | 3.053 | 1.191 | 2.081 | Early up | | | | | | | | | |
| B7L74_04400 | hypothetical protein | | -1.073 | -0.310 | -2.254 | 2.046 | -1.505 | -0.467 | -0.313 | -0.331 | -0.774 | -0.656 | -0.073 | | | | | | | | | | |
| B7L74_04405 | 30S ribosomal protein S6 | Ribosomal proteins | -0.140 | 0.004 | 3.025 | 3.586 | 2.672 | 0.973 | 1.587 | 2.871 | 0.096 | 0.426 | 0.345 | Early up | | | | | | | | | |
| B7L74_04410 | 30S ribosomal protein S18 | Ribosomal proteins | -1.161 | -0.792 | -2.531 | 1.405 | -0.736 | -0.974 | -0.829 | -0.726 | -0.525 | -1.524 | -0.342 | | | | | | | | | | |
| B7L74_04415 | hypothetical membrane spanning protein | | -1.239 | -0.538 | -1.958 | 0.951 | -1.210 | -0.602 | -1.139 | -0.151 | -0.480 | -0.613 | -0.013 | | | | | | | | | | |
| B7L74_04420 | 50S ribosomal protein L9 | Ribosomal proteins | -1.166 | -1.222 | -1.664 | 0.901 | -2.266 | -1.252 | -0.653 | -1.886 | -1.357 | -0.926 | -0.471 | Early down | | | | | | | | | |
| B7L74_04425 | intein-containing replicative DNA helicase | | -0.386 | -1.928 | -0.127 | 0.475 | -1.927 | -1.434 | 0.283 | -1.245 | -0.893 | 0.349 | 0.835 | | | | | | | | | | |
| B7L74_04430 | alanine racemase | | 0.575 | 0.974 | 1.182 | -1.628 | 1.343 | 0.716 | 1.419 | 1.419 | 2.994 | 0.638 | 1.469 | | | | | | | | | | |
| B7L74_04435 | succinate dehydrogenase assembly factor 2 family prot | | 0.541 | 1.045 | 0.437 | -2.023 | 1.284 | 1.282 | 0.681 | 0.396 | 2.854 | 0.873 | 1.386 | | | | | | | | | | |
| B7L74_04440 | deacylase | | -4.683 | -2.390 | -18.284 | -3.962 | -4.271 | -3.304 | -2.832 | 0.607 | -0.352 | -0.875 | 1.071 | | | | | | | | | | |
| B7L74_04445 | uridine kinase | Pyrimidine metabolism | 1.038 | -0.693 | 1.540 | 2.079 | -0.182 | 1.723 | 1.808 | 2.031 | 2.558 | 2.346 | 2.874 | Early up | | | | | | | | | |
| B7L74_04450 | DNA mismatch repair protein MutS | DNA repair and recombination pro | 2.282 | 3.285 | 2.966 | 3.113 | 4.139 | 2.962 | 2.280 | 4.774 | 3.473 | 4.036 | 4.309 | Early up | | | | | | | | | |
| B7L74_04455 | chorismate synthase | Phenylalanine, tyrosine and tryptop | 0.598 | -1.280 | 2.402 | 1.885 | 0.396 | -0.003 | 1.805 | 0.631 | 1.040 | 1.934 | 2.733 | Early up | | | | | | | | | |
| B7L74_04460 | aspartate-semialdehyde dehydrogenase | | 0.575 | -0.423 | 0.509 | 1.782 | 0.193 | 0.612 | 1.123 | 1.059 | 1.457 | 1.190 | 1.583 | | | | | | | | | | |
| B7L74_04465 | thiamine biosynthesis protein ThiF | | 0.234 | -0.586 | 0.401 | 0.847 | -0.738 | 1.050 | 0.598 | 1.238 | 1.503 | 0.266 | 0.083 | | | | | | | | | | |
| B7L74_04470 | hypothetical protein | | 9.816 | 6.087 | 6.495 | 2.194 | 11.768 | 11.348 | 7.105 | 5.233 | 11.295 | 7.738 | 5.190 | Early up | | | | | | | | | |
| B7L74_04475 | hypothetical protein | | 7.602 | 4.322 | 4.562 | 4.297 | 9.808 | 9.130 | 4.909 | 2.839 | 8.081 | 4.810 | 3.512 | Early up | | | | | | | | | |
| B7L74_04480 | hypothetical cytosolic protein | | 4.350 | 0.697 | -0.027 | 1.639 | 4.572 | 5.068 | 1.702 | 2.800 | 4.018 | 0.855 | 0.594 | Early up | | | | | | | | | |
| B7L74_04485 | hypothetical protein | | -0.616 | -0.607 | -0.929 | -0.536 | -0.406 | -0.080 | -0.403 | -1.344 | 0.196 | -0.953 | -1.237 | | | | | | | | | | |
| B7L74_04490 | hypothetical cytosolic protein | Effector protein | 7.800 | 3.604 | 4.237 | 1.542 | 9.314 | 8.207 | 6.295 | 3.440 | 7.599 | 4.503 | 2.100 | Early up | | Yes | | | | | | | |
| B7L74_04495 | hypothetical protein | | 2.651 | 0.840 | -0.124 | 0.651 | 2.741 | 1.819 | 1.005 | -0.695 | 1.375 | 0.296 | 0.140 | | | | | | | | | | |
| B7L74_04500 | transposase | | -0.594 | -0.380 | 0.371 | 0.677 | -0.823 | -1.015 | -0.745 | -0.817 | -0.930 | 0.516 | 0.222 | | | | | | | | | | |
| B7L74_04505 | helix-turn-helix domain-containing protein | | 0.041 | 0.838 | -0.034 | -1.805 | 2.642 | 2.081 | 0.448 | 1.487 | 1.883 | -1.279 | -0.910 | | | | | | | | | | |
| B7L74_04510 | hypothetical protein | | -0.926 | -0.791 | 1.136 | 0.591 | -0.906 | -1.104 | -0.623 | -1.051 | -0.513 | 0.664 | 1.045 | | | | | | | | | | |
| B7L74_04515 | translational GTPase TypA | Host-cell attachment | -0.105 | -1.151 | 1.033 | 0.834 | -0.572 | 0.019 | 1.253 | 0.329 | 0.595 | 0.359 | 1.463 | | | | | | | | | | |
| B7L74_04520 | hypothetical cytosolic protein | Effector protein | -0.582 | -1.073 | -1.329 | -0.150 | 0.642 | 0.840 | 0.636 | 0.408 | 1.089 | -0.305 | 0.543 | | | Yes | | | | | | | |
| B7L74_04525 | bifunctional 4'-phosphopantothenoylcysteine decarbox | | -0.673 | -1.605 | 0.976 | 1.334 | -1.729 | -0.473 | 0.521 | 0.305 | 0.906 | 1.471 | 1.846 | | | | | | | | | | |
| B7L74_04530 | patatin | | 0.299 | -2.149 | -1.042 | 0.480 | 0.082 | 0.984 | 2.475 | 1.822 | 2.544 | 0.881 | 0.918 | | | | | | | | | | |
| B7L74_04535 | disulfide formation protein | | -1.323 | -1.625 | -1.512 | 0.846 | -2.764 | -3.737 | -1.047 | -0.772 | -0.990 | -1.171 | -0.873 | | | | | | | | | | |
| B7L74_04540 | dihydromonapterin reductase | | -1.225 | -0.938 | -1.823 | 0.762 | 1.738 | 2.897 | -0.877 | -0.290 | 2.656 | -1.619 | -1.585 | | | | | | | | | | |
| B7L74_04545 | Zn-dependent hydrolase | | -0.904 | -1.484 | -0.921 | 0.715 | -1.548 | -1.737 | 0.020 | 0.163 | 0.412 | -0.080 | 0.239 | | | | | | | | | | |
| B7L74_04550 | hypothetical exported membrane associated protein | | 0.134 | -1.662 | -1.617 | 1.261 | 0.129 | 0.849 | -0.089 | -1.245 | -0.263 | -0.391 | -0.107 | | | | | | | | | | |
| B7L74_04555 | tRNA pseudouridine(38-40) synthase TruA | | -0.287 | -1.615 | 0.894 | 1.181 | -1.642 | -1.375 | 0.178 | -0.902 | -0.508 | 1.075 | 1.500 | | | | | | | | | | |
| B7L74_04560 | acetyl-CoA carboxylase carboxyl transferase subunit b | | -0.013 | -0.301 | 1.372 | 0.439 | 0.598 | 0.348 | 0.698 | -0.362 | 1.696 | 1.355 | 1.666 | | | | | | | | | | |
| B7L74_04565 | bifunctional tetrahydrofolate synthase/dihydrofolate syn | | -0.342 | -1.114 | -0.013 | 1.107 | -0.743 | -0.777 | 0.080 | 0.065 | -0.098 | 0.308 | 0.273 | | | | | | | | | | |
| B7L74_04570 | SPOR domain-containing protein | | -2.550 | -3.068 | -4.869 | -5.298 | -1.398 | -3.218 | -3.272 | -2.603 | -1.909 | -5.638 | -5.753 | Early down | | | | | | | | | |
| B7L74_04575 | colicin V synthesis protein | | -1.087 | -1.914 | -1.699 | -0.408 | -1.685 | -0.692 | -1.026 | -0.799 | -0.653 | -0.998 | -0.991 | | | | | | | | | | |
| B7L74_04580 | amidophosphoribosyltransferase | | -1.055 | -1.615 | -1.581 | 0.147 | -1.980 | -1.532 | -0.856 | -1.424 | -1.465 | -0.917 | -1.002 | | | | | | | | | | |
| B7L74_04585 | hypothetical proteinthyroglobulin type-1 repeat domain | Eucaryotic-like ORFs. | 2.933 | 0.602 | 2.791 | 4.094 | 1.242 | 0.572 | 1.512 | 0.859 | -0.546 | 2.883 | 2.417 | Up in P12 | | | | | | | | | |
| B7L74_04590 | hypothetical protein | | -1.080 | -1.309 | -2.572 | 0.052 | -1.744 | -1.786 | -0.678 | -1.376 | -1.113 | -1.225 | -1.405 | | | | | | | | | | |
| B7L74_04595 | hypothetical protein | | 0.961 | -0.793 | -0.629 | 2.103 | 1.478 | 0.978 | 0.591 | 4.347 | 4.143 | 0.531 | 0.598 | Late up | | | | | | | | | |
| B7L74_04600 | hypothetical protein | | 3.142 | -1.755 | -4.016 | 1.506 | 2.221 | 3.261 | 1.410 | 6.011 | 6.212 | 0.296 | 0.577 | Late up | | | | | | | | | |
| B7L74_04605 | hypothetical protein | | -0.579 | -1.478 | 0.450 | -2.289 | 1.880 | 3.101 | 1.198 | 1.194 | 1.669 | -0.139 | 0.511 | | | | | | | | | | |
| B7L74_04610 | hypothetical protein | | 4.217 | 0.830 | 3.664 | 2.297 | 4.803 | 4.411 | 3.684 | 3.045 | 4.681 | 4.231 | 4.090 | Early up | | | | | | | | | |
| B7L74_04615 | tRNA dihydrouridine(20/20a) synthase DusA | | -2.582 | -3.830 | -1.266 | -2.613 | -2.197 | -3.513 | -1.292 | -2.456 | -2.638 | -4.052 | -1.260 | | | | | | | | | | |
| B7L74_04620 | 50S ribosomal protein L31 | Ribosomal proteins | 1.168 | -1.024 | 0.044 | 4.276 | 0.483 | 0.852 | 1.325 | 2.251 | 2.863 | 1.176 | 1.454 | Early up | | | | | | | | | |
| B7L74_04625 | MFS transporter | Transporter | 1.707 | -1.636 | 0.545 | 1.863 | 1.580 | 2.726 | 0.796 | 1.955 | 3.057 | 1.335 | 1.308 | Early up | | | | | | | | Yes (MFS) | |
| B7L74_04630 | hypothetical protein=YciL | | 0.220 | -0.459 | 1.173 | 0.889 | -0.626 | 0.578 | 1.987 | 1.946 | 1.865 | 1.477 | 1.132 | | | | | | | | | | |
| B7L74_04635 | intracellular septation protein A | Cell growth | 0.315 | 0.627 | 0.203 | 0.988 | 0.748 | 1.587 | 1.704 | 3.017 | 2.788 | 2.272 | 1.299 | Late up | | | | | | | | | |
| B7L74_04640 | threonylcarbamoyl-AMP synthase | DNA | 1.367 | 0.283 | 2.617 | -0.025 | 0.613 | 0.418 | 2.241 | 0.945 | 1.588 | 1.469 | 1.124 | | | | | | | | | | |
| B7L74_04645 | hypothetical protein, Hypothetical membrane associate | Membrane | 1.268 | 0.028 | 0.652 | 1.669 | 0.762 | 1.063 | 1.279 | 2.402 | 3.170 | 1.455 | 1.380 | Late up | | | | | | | | | |
| B7L74_04650 | hypothetical protein | | 1.008 | -0.106 | -0.846 | 1.898 | -0.549 | 0.932 | 0.004 | 3.315 | 4.024 | 1.738 | 1.792 | Late up | | | | | | | | | |
| B7L74_04655 | 2-methylcitrate dehydratase | Propanoate metabolism | 0.783 | -1.190 | 0.522 | 2.267 | 0.535 | 0.889 | 1.304 | 2.493 | 2.861 | 1.321 | 1.592 | Early up | | | | | | | | | |
| B7L74_04660 | thioesterase | Terpenoid synthesis | 4.502 | 0.222 | 2.157 | 2.544 | 4.783 | 5.154 | 3.680 | 3.380 | 4.466 | 3.058 | 2.772 | Early up | | | | | | | | | |
| B7L74_04665 | hypothetical membrane spanning protein | | 0.152 | 0.859 | -0.350 | -0.612 | 0.284 | 0.172 | -0.624 | 0.585 | 0.600 | -0.804 | -1.375 | | | | | | | | | | |
| B7L74_04670 | enhanced entry protein | | -0.350 | 0.623 | 0.338 | 0.239 | 0.078 | 0.090 | -0.781 | 0.446 | 0.642 | -0.399 | -1.162 | | | | | | | | | | |
| B7L74_04675 | protein=endonuclease/exonuclease/phosphatase family | | 0.677 | 0.981 | 1.601 | 0.177 | 0.514 | -0.147 | 2.246 | 1.858 | 2.310 | 0.485 | 1.407 | | | | | | | | | | |
| B7L74_04680 | cyclic nucleotide-binding protein | | -0.090 | 0.463 | -0.690 | 1.600 | -0.152 | -0.582 | -1.000 | 1.997 | 2.069 | -1.236 | -1.317 | Late up | | | | | | | | | |
| B7L74_04685 | hypothetical protein | | 3.726 | 0.091 | -2.673 | -2.351 | 4.472 | 3.974 | -0.787 | 0.059 | 3.002 | -0.528 | -1.665 | Early up | | | | | | | | | |
| B7L74_04690 | hypothetical protein | | 0.022 | 0.834 | -0.034 | -1.430 | 2.619 | 1.618 | 0.430 | 1.863 | 3.234 | 0.865 | -0.916 | | | | | | | | | Yes (Ions/Potassium) | |
| B7L74_04695 | fatty acid desaturase | Eucaryotic-like ORFs. | -2.370 | -1.069 | -1.390 | -0.437 | -3.156 | -2.251 | -1.764 | -1.201 | -2.045 | -2.593 | -1.475 | | | | | | | | | | |
| B7L74_04700 | hypothetical protein, Hypothetical cytosolic protein | | 0.726 | 1.060 | 1.038 | -0.774 | -1.205 | -1.251 | 0.226 | 1.759 | 0.503 | 0.221 | 2.589 | | | | | | | | | | |
| B7L74_04705 | MFS transporter | Transporter | 2.534 | 1.261 | 1.956 | -0.184 | 2.076 | 1.415 | 1.298 | 1.265 | 1.977 | 0.982 | 2.984 | | | | | | | | | | |
| B7L74_04710 | AI-2E family transporter | Transporter, Structural proteins | 4.502 | 1.424 | 4.988 | 3.482 | 4.456 | 3.743 | 3.994 | 2.836 | 4.490 | 5.780 | 5.711 | Early up | | | | | | | | | |
| B7L74_04715 | O-methyltransferase | | 1.333 | 0.731 | 2.165 | -0.122 | 1.220 | 0.744 | 0.724 | 0.264 | 0.931 | 1.082 | 0.392 | | | | | | | | | | |
| B7L74_04720 | lytic transglycosylaseMembrane-bound lytic murein tra | Membrane | 2.491 | 0.963 | 2.081 | -1.606 | 1.806 | 2.375 | 1.713 | 1.278 | 0.322 | 0.750 | 0.443 | | | | | | | | | | |
| B7L74_04725 | 3-hydroxyisobutyrate dehydrogenase | | 0.230 | -0.400 | 1.184 | -0.406 | -0.806 | -0.297 | 0.693 | 0.989 | 1.088 | 0.962 | 1.059 | | | | | | | | | | |
| B7L74_04730 | methylmalonate-semialdehyde dehydrogenase (CoA ac | | -0.369 | -0.666 | -1.929 | -0.273 | -1.046 | -0.785 | -0.778 | -0.274 | 0.301 | -0.878 | -0.956 | | | | | | | | | | |
| B7L74_04735 | pyridoxine/pyridoxamine 5'-phosphate oxidase | | -0.855 | -0.730 | -1.632 | -0.288 | -0.910 | -0.655 | -0.861 | -0.484 | -0.306 | -0.522 | -0.829 | | | | | | | | | | |
| B7L74_04740 | short-chain dehydrogenase | | -0.360 | -0.306 | -1.120 | -0.531 | -1.045 | -0.760 | -0.590 | 0.691 | 0.481 | -0.325 | -0.588 | | | | | | | | | | |
| B7L74_04745 | 3-methyladenine DNA glycosylase | | -0.006 | -0.823 | 0.814 | 0.372 | -0.448 | -0.218 | 0.688 | 0.213 | 1.272 | 0.675 | 1.368 | | | | | | | | | | |
| B7L74_04750 | glycerol-3-phosphate dehydrogenase | | 0.305 | 0.129 | 1.782 | 0.876 | 0.414 | -0.891 | 2.000 | 0.473 | 1.244 | 1.678 | 2.656 | Early up | | | | | | | | | |
| B7L74_04755 | glycerol kinase | Glycerolipid metabolism | 1.631 | 0.013 | 0.822 | -0.242 | 2.397 | 3.426 | 1.005 | 0.941 | 2.424 | 0.742 | 0.395 | Early up | | | | | | | | | |
| B7L74_04760 | ABC transporter permease | Transporter | 4.067 | 0.849 | 2.148 | 0.070 | 3.817 | 1.365 | 2.199 | 1.909 | 3.552 | 2.594 | 1.977 | | | | | | | | | Yes (Phospholipid) | |
| B7L74_04765 | ABC transporter ATP-binding protein | Transporter | 0.406 | -0.691 | -0.285 | 0.594 | -0.244 | 0.221 | 0.559 | 1.984 | 2.395 | 0.680 | 0.694 | Late up | | | | | | | | Yes (General) | |
| B7L74_04770 | RNA-binding protein | | -0.248 | -0.661 | -0.746 | 0.474 | -1.311 | -0.892 | -0.225 | 1.739 | 2.148 | 0.719 | 0.063 | | | | | | | | | | |
| B7L74_04775 | hypothetical proteinOmpA-like transmembrane domain | | -0.115 | 0.264 | 0.864 | 1.185 | -0.570 | -1.019 | -0.577 | 0.434 | -0.058 | -0.155 | -0.229 | | | | | | | | | | |
| B7L74_04780 | hypothetical protein(cirC in Table 1 of RightonQ paper | Effector protein | -1.284 | -1.404 | -2.237 | -0.906 | -2.118 | -2.035 | -1.249 | -1.740 | -1.841 | -1.635 | -0.970 | Early down | | Yes | | | | | | | |
| B7L74_04785 | tRNA (guanosine(46)-N7)-methyltransferase TrmB | | 0.197 | -0.084 | 0.148 | -0.219 | 0.169 | 0.483 | 0.922 | 1.246 | 2.151 | -0.032 | 0.407 | | | | | | | | | | |
| B7L74_04790 | tRNA-Gly | tRNA | -0.207 | -0.312 | -0.959 | -0.435 | -0.833 | -1.358 | 0.489 | -0.064 | -1.327 | -1.133 | -0.383 | | | | | | | | | | |
| B7L74_04795 | AsmA family protein | | -1.042 | -0.921 | -1.802 | -0.398 | -0.805 | -0.627 | -1.071 | 0.191 | 0.265 | -1.356 | -1.117 | | | | | | | | | | |
| B7L74_04800 | A/G-specific adenine glycosylase | | 0.465 | -0.618 | 0.077 | 0.386 | -0.059 | 0.452 | 0.367 | 1.349 | 1.629 | -0.123 | -0.034 | | | | | | | | | | |
| B7L74_04805 | Fe(2+)-trafficking protein | | -0.796 | -0.504 | 0.624 | 2.764 | 1.905 | 1.512 | 1.015 | 0.809 | 1.877 | -0.031 | 0.625 | | | | | | | | | | |
| B7L74_04810 | hypothetical exported protein | | 2.523 | 1.983 | 0.989 | 2.642 | 2.348 | 1.928 | 2.135 | 3.932 | 3.863 | 1.803 | 2.227 | Early up | | | | | | | | | |
| B7L74_04815 | rhodanese related sulfurtransferase | Transferase | 1.466 | 0.406 | 1.602 | 0.808 | 0.205 | -0.393 | 1.566 | 0.463 | 0.906 | 1.889 | 1.129 | | | | | | | | | | |
| B7L74_04820 | tRNA-Phe | tRNA | 0.889 | 0.524 | 0.976 | -0.202 | 0.457 | 0.260 | 1.303 | 0.150 | 0.332 | 0.803 | 0.374 | | | | | | | | | | |
| B7L74_04825 | hypothetical protein | | -0.475 | -0.335 | -1.463 | -0.704 | 0.846 | 0.965 | -0.044 | 1.849 | 2.374 | -0.226 | -0.065 | | | | | | | | | | |
| B7L74_04830 | hypothetical membrane associated protein | | -1.713 | 0.129 | -2.418 | -1.073 | -0.007 | -0.213 | -2.219 | 0.202 | 0.670 | -1.047 | -1.910 | | | | | | | | | | |
| B7L74_04835 | death-on-curing protein | | -0.391 | -0.290 | -0.325 | 1.359 | -0.496 | 0.079 | 0.072 | 1.472 | 1.872 | 0.586 | 0.692 | | | | | | | | | | |
| B7L74_04840 | hypothetical protein | | 1.117 | 0.622 | 1.504 | 0.785 | 0.138 | 2.019 | 1.352 | 3.245 | 3.547 | 0.533 | 0.924 | Late up | | | | | | | | | |
| B7L74_04845 | hypothetical protein | | -0.927 | 0.510 | -0.708 | 1.386 | 0.562 | 0.748 | -0.226 | 2.356 | 1.874 | 0.173 | -0.425 | | | | | | | | | | |
| B7L74_04850 | hypothetical cytosolic protein | Prokaryotic defense system | 2.384 | 3.348 | 2.148 | 3.521 | 3.259 | 2.992 | 1.918 | 5.582 | 5.658 | 1.624 | 0.586 | Continuous up | | | | | | | | | |
| B7L74_04855 | type I-F CRISPR-associated helicase Cas3 | CRISPR/Cas, Prokaryotic defense | 0.375 | -0.462 | 0.066 | 0.956 | -0.529 | 1.417 | 0.378 | 2.711 | 3.049 | -0.532 | -0.150 | Late up | | | | | | | | | |
| B7L74_04860 | subtype I-F CRISPR-associated endonuclease Cas1 | CRISPR/Cas | -0.496 | 0.271 | 0.391 | 0.575 | 3.256 | 1.764 | 2.066 | 1.568 | 2.013 | -1.833 | 0.426 | | | | | | | | | | |
| B7L74_04865 | hypothetical exported protein | | -1.567 | -0.720 | -0.070 | 1.814 | -0.603 | -1.061 | -0.316 | -0.799 | -1.231 | -1.487 | -0.833 | | | | | | | | | | |
| B7L74_04870 | amino acid permease | Transporter | -0.672 | -0.531 | -1.582 | -0.628 | -1.623 | 0.225 | -0.771 | 1.536 | 1.071 | -1.208 | -0.519 | | | | | | | | | Yes (Amino acid) | |
| B7L74_04875 | NADH pyrophosphatase | | -0.274 | -0.637 | -1.863 | 0.233 | -0.366 | -0.140 | -1.202 | 0.841 | 1.368 | -0.535 | -1.032 | | | | | | | | | | |
| B7L74_04880 | DNA-binding response regulator (gene=sirA) | Transcriptional regulators | -0.166 | 0.242 | 0.305 | 0.171 | -0.452 | -0.565 | -0.988 | -0.703 | -0.235 | -0.312 | -0.244 | | | | | | | | | | |
| B7L74_04885 | hypothetical membrane associated protein | | 0.216 | 1.078 | 2.317 | -1.592 | 1.446 | 1.290 | 0.518 | 0.783 | 3.120 | 0.267 | 1.961 | | | | | | | | | | |
| B7L74_04890 | hypothetical protein | | 0.847 | 1.637 | 1.362 | -1.894 | 2.174 | 1.686 | 0.787 | 2.280 | 3.527 | 0.756 | 0.566 | Early up | | | | | | | | | |
| B7L74_04895 | hypothetical protein | | -2.353 | -1.595 | -0.034 | -1.738 | 0.294 | 0.147 | -0.963 | 3.089 | 2.628 | -1.875 | -0.889 | | | | | | | | | | |
| B7L74_04900 | hypothetical protein | | 0.333 | 0.383 | 0.191 | 0.141 | -0.541 | 0.383 | -0.013 | 2.522 | 2.853 | 0.255 | -0.071 | Late up | | | | | | | | | |
| B7L74_04905 | hypothetical protein | | 0.722 | 0.892 | 0.957 | 0.065 | 1.455 | 1.797 | 1.140 | 2.597 | 2.355 | 0.323 | -0.865 | | | | | | | | | | |
| B7L74_04910 | trehalose-phosphatase | | 0.314 | 0.393 | 0.511 | 0.503 | -0.015 | -0.513 | 0.259 | 0.670 | 0.420 | 0.710 | 0.423 | | | | | | | | | | |
| B7L74_04915 | Bcr/CflA family drug resistance efflux transporter | Transporter | -1.166 | -0.309 | 0.255 | 0.706 | -0.931 | -1.085 | -0.471 | -0.033 | 0.011 | 0.209 | -0.455 | | | | | | | | | Yes (drug resistance) | |
| B7L74_04920 | XRE family transcriptional regulator | | -0.583 | 0.347 | 3.478 | 0.775 | 0.083 | -0.817 | 0.664 | 0.534 | 1.316 | 1.133 | 1.208 | | | | | | | | | | |
| B7L74_04925 | hypothetical protein | | -0.539 | 0.120 | 1.077 | 0.814 | -0.744 | -1.295 | -1.018 | 0.313 | 0.105 | -0.388 | -1.377 | | | | | | | | | | |
| B7L74_04930 | short-chain dehydrogenase | | -1.923 | -1.498 | -2.235 | 0.019 | -2.287 | -2.054 | -1.643 | -2.671 | -3.045 | -2.001 | -1.592 | Continuous down | | | | | | | | | |
| B7L74_04935 | peroxiredoxin | PV detoxification | -0.826 | -0.232 | -2.844 | 0.981 | -0.739 | -1.999 | -0.500 | -4.554 | -2.876 | -1.622 | -1.536 | | | | | | | | | | |
| B7L74_04940 | hypothetical protein | | -1.290 | -1.504 | -1.751 | 0.684 | -1.203 | -1.403 | -0.302 | -0.119 | -0.079 | -0.902 | 0.250 | | | | | | | | | | |
| B7L74_04945 | hypothetical protein | | 0.283 | 0.564 | -0.313 | -0.137 | -0.334 | 0.353 | -0.002 | 0.939 | 1.220 | -1.540 | 0.335 | | | | | | | | | | |
| B7L74_04950 | hypothetical protein | | -1.370 | -1.586 | -2.597 | -0.078 | -1.958 | -1.410 | -1.739 | -0.752 | -0.855 | -1.173 | -1.491 | | | | | | | | | | |
| B7L74_04955 | cytochrome d terminal oxidase subunit 1 | Respiration/cyt bd | -1.750 | -1.584 | -2.924 | -0.855 | -2.120 | -1.659 | -1.922 | -0.884 | -0.526 | -2.041 | -1.861 | Early down | | | | | | Yes | | | |
| B7L74_04960 | cytochrome d ubiquinol oxidase subunit II | Respiration/cyt bd | 1.295 | -0.162 | -1.034 | -0.250 | 1.646 | 1.800 | -0.549 | 3.565 | 3.248 | -1.924 | -0.281 | Late up | | | | | | Yes | | | |
| B7L74_04965 | cytochrome bd-I oxidase subunit CydX | Respiration/cyt bd | -1.136 | 0.184 | -1.881 | -1.438 | 0.089 | 0.297 | -2.040 | -0.818 | -0.825 | -2.973 | -3.626 | Late down | | | | | | Yes | | | |
| B7L74_04970 | phospholipase | | 1.341 | 0.910 | 1.190 | 0.694 | 3.029 | 3.113 | 0.456 | 0.265 | 1.953 | 0.245 | -0.580 | Early up | | | | | | | | | |
| B7L74_04975 | hypothetical exported membrane associated protein | | -0.846 | -0.226 | -0.408 | 0.185 | -0.061 | 0.554 | -0.969 | 0.419 | 0.578 | -1.175 | -1.196 | | | | | | | | | | |
| B7L74_04980 | dihydroorotate dehydrogenase (quinone) | | 5.518 | 4.183 | 3.921 | 2.208 | 3.251 | 0.991 | 4.287 | 3.523 | 5.661 | 4.841 | 3.039 | | | | | | | | | | |
| B7L74_04985 | hypothetical cytosolic protein | | 1.145 | 0.999 | 1.978 | 0.082 | 2.378 | 1.159 | 2.334 | 3.021 | 3.337 | 2.767 | -0.361 | | | | | | | | | | |
| B7L74_04990 | isovaleryl-CoA dehydrogenase | | -0.523 | -0.746 | -2.309 | 0.094 | -1.938 | -1.197 | -0.739 | 0.086 | 0.375 | -1.026 | -0.781 | | | | | | | | | | |
| B7L74_04995 | acetyl-CoA acetyltransferase | | 0.931 | 0.639 | -2.645 | 0.090 | 2.539 | 1.263 | 0.329 | 0.230 | 0.971 | -2.229 | 0.385 | | | | | | | | | | |
| B7L74_05000 | methylcrotonoyl-CoA carboxylase | Valine, leucine and isoleucine deg | -1.239 | -1.294 | -2.960 | -0.934 | -1.649 | -1.339 | -1.499 | 0.046 | 0.399 | -2.042 | -1.784 | Early down | | | | | | | | | |
| B7L74_05005 | methylglutaconyl-CoA hydratase | Valine, leucine and isoleucine deg | -1.594 | -0.785 | -1.282 | -1.776 | 0.173 | -0.289 | -1.381 | 0.944 | 1.995 | -2.192 | -2.152 | Variable | | | | | | | | | |
| B7L74_05010 | biotin carboxylase | Valine, leucine and isoleucine deg | -1.333 | -0.415 | -2.100 | -0.928 | -1.465 | -0.625 | -1.088 | 1.830 | 2.075 | -1.916 | -1.487 | Variable | | | | | | | | | |
| B7L74_05015 | hypothetical membrane associated protein | Effector protein | 0.506 | 0.551 | 1.166 | 1.173 | -0.322 | -0.617 | 0.314 | 1.142 | 1.070 | 1.131 | 0.555 | | | Yes | | | | | | | |
| B7L74_05020 | lipoprotein | | 1.456 | 1.270 | 2.279 | 0.368 | 0.372 | -0.081 | 1.261 | 0.636 | -0.973 | 0.940 | 0.389 | | | | | | | | | | |
| B7L74_05025 | lipoprotein | Membrane | 1.703 | 1.670 | 3.860 | -0.302 | 1.841 | 1.146 | 2.445 | 1.877 | 3.238 | 2.170 | 2.048 | Early up | | | | | | | | | |
| B7L74_05030 | hypothetical protein | | 0.450 | 0.317 | 0.020 | -0.136 | 1.231 | 0.238 | 0.305 | 1.452 | 1.961 | -0.001 | 0.927 | | | | | | | | | | |
| B7L74_05035 | 3-deoxy-7-phosphoheptulonate synthase | | 0.039 | 0.056 | 0.346 | -1.032 | -0.248 | -0.167 | -0.033 | 0.261 | 0.124 | -1.103 | -1.309 | | | | | | | | | | |
| B7L74_05040 | chorismate mutase | | -0.946 | -0.079 | -0.844 | -0.589 | -1.197 | -0.667 | -1.274 | 0.861 | 0.754 | -1.628 | -1.607 | | | | | | | | | | |
| B7L74_05045 | prephenate dehydratase | | -0.599 | -0.483 | -1.446 | -1.369 | -1.596 | -0.357 | -1.044 | 1.204 | 1.667 | -1.579 | -1.930 | | | | | | | | | | |
| B7L74_05050 | prephenate dehydrogenase | | -0.908 | -0.152 | -1.035 | -2.759 | 1.507 | 2.125 | 4.053 | 3.409 | 4.496 | -2.251 | -1.415 | | | | | | | | | | |
| B7L74_05055 | hypothetical cytosolic protein | | -1.322 | -0.593 | -1.193 | -0.985 | -1.321 | -1.077 | -1.565 | -0.454 | -0.119 | -0.765 | -0.840 | | | | | | | | | | |
| B7L74_05060 | 23S rRNA (guanosine(2251)-2'-O)-methyltransferase I | | 0.382 | 0.502 | 0.964 | -0.868 | 0.060 | 0.151 | 0.454 | -0.133 | 0.433 | -0.502 | -0.799 | | | | | | | | | | |
| B7L74_05065 | septal ring lytic transglycosylase RlpA family lipoprote | | -0.344 | 0.231 | 2.067 | -0.738 | 0.182 | -0.080 | 0.290 | -0.206 | 0.461 | 0.574 | -0.541 | | | | | | | | | | |
| B7L74_05070 | uracil-DNA glycosylase | DNA | 0.477 | -0.156 | 0.317 | 0.732 | -0.701 | -0.368 | 0.182 | -0.703 | -0.428 | -0.119 | -0.375 | | | | | | | | | | |
| B7L74_05075 | hypothetical protein | | -0.403 | 1.191 | -0.534 | -0.098 | -0.051 | 0.655 | -0.875 | 1.225 | 0.688 | -0.941 | -1.469 | | | | | | | | | | |
| B7L74_05080 | hypothetical protein | | 1.209 | 1.598 | 2.164 | 0.665 | 1.776 | 1.456 | 1.630 | 3.512 | 3.676 | 0.659 | 0.408 | Late up | | | | | | | | | |
| B7L74_05085 | tRNA-Leu | tRNA | 1.649 | 1.777 | 2.888 | 0.325 | 0.753 | 1.846 | 3.543 | 2.452 | 1.936 | 1.510 | 1.838 | Early up | | | | | | | | | |
| B7L74_05090 | peptide deformylase | | 0.667 | 0.393 | 1.261 | 1.089 | -0.051 | -0.082 | 0.787 | 0.960 | 0.372 | 0.532 | 1.142 | | | | | | | | | | |
| B7L74_05095 | hypothetical protein | | 2.294 | 0.516 | 1.223 | 2.116 | 1.206 | 1.600 | 1.713 | 2.834 | 2.237 | 1.183 | 1.811 | | | | | | | | | | |
| B7L74_05100 | cytochrome B | Cytochrome | -0.499 | -0.830 | -1.236 | 0.918 | -1.074 | 0.360 | 0.128 | 1.476 | 1.551 | 0.379 | 0.116 | | | | | | | Yes | | | |
| B7L74_05105 | protoheme IX farnesyltransferase | | -0.928 | 1.988 | -2.331 | -1.153 | 0.998 | 0.323 | -0.099 | 0.896 | 1.551 | -0.565 | -0.905 | | | | | | | Yes | | | |
| B7L74_05110 | lipoprotein | | -0.765 | -1.477 | -2.296 | 0.949 | -0.346 | -1.635 | -0.676 | -0.487 | 0.494 | -0.724 | -0.869 | | | | | | | | | | |
| B7L74_05115 | adenylosuccinate synthetase | Purine metabolism | -1.111 | -0.678 | -2.400 | -0.221 | -1.516 | -0.937 | -0.901 | -0.629 | -0.256 | -0.943 | -0.234 | Early down | | | | | | | | | |
| B7L74_05120 | GTP-binding protein | | -0.306 | -0.211 | -0.773 | 0.554 | -0.723 | -0.086 | -0.127 | 0.273 | 0.075 | -0.162 | 0.445 | | | | | | | | | | |
| B7L74_05125 | lipoprotein-releasing system ATP-binding protein LolD | ABC transporters | -0.954 | -0.331 | -2.153 | 0.277 | -1.845 | -0.911 | -1.348 | -0.181 | -0.162 | -1.132 | -0.945 | Early down | | | | | | | | Yes (Outer membrane/Lipoprotein) | |
| B7L74_05130 | ABC transporterLipoprotein releasing system transmem | ABC transporters | -0.796 | 1.520 | 0.643 | 3.680 | 0.506 | 1.454 | 1.736 | 2.764 | 2.095 | 2.214 | 2.232 | Early up | | | | | | | | Yes (Outer membrane/Lipoprotein) | |
| B7L74_05135 | bifunctional biotin--[acetyl-CoA-carboxylase] syntheta | Biotin metabolism | 0.349 | 0.076 | 0.383 | 0.152 | 0.037 | 0.883 | 0.335 | 2.146 | 2.434 | -0.067 | -0.264 | Late up | | | | | | | | | |
| B7L74_05140 | dethiobiotin synthase | Biotin metabolism | 1.029 | 2.372 | -1.483 | -0.816 | 3.629 | 5.025 | 1.436 | 5.150 | 3.536 | -0.908 | 0.082 | Early up | | | | | | | | | |
| B7L74_05145 | malonyl-[acyl-carrier protein] O-methyltransferase Bio | | 0.015 | 2.843 | -2.496 | -1.830 | 3.678 | 3.190 | 0.423 | 4.766 | 2.522 | -1.921 | -0.932 | | | | | | | | | | |
| B7L74_05150 | carboxylesterase | Biotin metabolism | 1.307 | 2.844 | -2.499 | -0.455 | 4.982 | 4.455 | 0.438 | 4.806 | 2.538 | -1.919 | -0.929 | Early up | | | | | | | | | |
| B7L74_05155 | 8-amino-7-oxononanoate synthase | Biotin metabolism | -1.213 | 1.541 | -6.529 | -3.453 | 0.814 | 2.292 | 0.074 | 1.822 | 0.338 | -5.939 | -4.950 | Late down | | | | | | | | | |
| B7L74_05160 | biotin synthase | | 1.728 | 2.804 | -3.092 | -1.827 | 2.488 | 2.863 | 1.685 | 2.572 | 3.037 | -2.507 | -1.518 | | | | | | | | | | |
| B7L74_05165 | DUF1658 domain-containing protein | | 1.361 | 0.844 | 1.551 | -3.230 | 2.069 | 1.530 | 0.478 | 1.860 | 2.750 | -1.891 | 0.143 | | | | | | | | | | |
| B7L74_05170 | adenosylmethionine--8-amino-7-oxononanoate aminot | Biotin metabolism | -0.094 | 0.432 | 0.736 | -0.131 | 0.759 | 0.681 | 1.115 | 3.544 | 3.776 | 0.304 | 0.411 | Late up | | | | | | | | | |
| B7L74_05175 | hypothetical protein | | -0.026 | 1.108 | 1.574 | 1.114 | 0.126 | -0.160 | 1.865 | 1.432 | 1.614 | 1.737 | 1.342 | | | | | | | | | Yes (Ions/Fluoride) | |
| B7L74_05180 | hypothetical protein | Transporter | -0.550 | -0.992 | -1.770 | 0.359 | -1.690 | 0.815 | -3.953 | 3.378 | 3.256 | -0.582 | -0.954 | Variable | | | | | | | | Yes (Ions/Fluoride) | |
| B7L74_05185 | hypothetical protein | | 1.797 | 0.274 | 0.392 | -1.285 | 0.630 | 1.342 | 3.220 | 2.971 | 3.692 | -0.876 | -0.891 | | | | | | | | | | |
| B7L74_05190 | beta-carotene 15%2C15'-monooxygenase | | 0.487 | 0.502 | 0.863 | 0.788 | 0.727 | 1.480 | 0.018 | 3.556 | 3.170 | -0.786 | -0.837 | Late up | | | | | | | | | |
| B7L74_05195 | hypothetical protein | | -2.140 | 0.642 | -1.324 | 0.143 | 0.738 | 0.662 | -1.737 | 1.659 | 0.965 | -3.413 | -3.246 | Late down | | | | | | | | | |
| B7L74_05200 | hypothetical protein | | 3.642 | 1.689 | 1.765 | -0.330 | 7.549 | 5.985 | 2.567 | 1.449 | 5.581 | 2.563 | 0.546 | Early up | | | | | | | | | |
| B7L74_05205 | tRNA-Cys | tRNA | 0.718 | 0.272 | 0.565 | 0.578 | -0.799 | -0.298 | 0.275 | -0.332 | -0.597 | -0.317 | 0.079 | | | | | | | | | | |
| B7L74_05210 | tRNA-Leu | tRNA | 1.027 | -0.345 | 0.804 | 0.532 | -0.566 | -0.770 | 0.564 | 0.262 | 0.897 | 0.089 | 0.256 | | | | | | | | | | |
| B7L74_05215 | integrase | | 3.001 | 2.114 | 0.942 | 0.981 | 3.280 | 3.449 | 2.203 | 6.139 | 6.525 | 1.024 | 1.997 | Late up | | | | | | | | | |
| B7L74_05220 | hypothetical membrane associated protein | | 1.037 | 1.350 | 0.034 | -1.290 | 2.035 | 2.717 | -1.485 | 2.854 | 4.418 | 0.209 | -0.213 | Early up | | | | | | | | | |
| B7L74_05225 | hypothetical cytosolic protein | | 0.552 | 0.667 | 2.339 | 0.145 | 0.381 | 0.297 | 0.977 | 2.655 | 2.688 | 1.227 | 1.438 | Late up | | | | | | | | | |
| B7L74_05230 | hypothetical protein | | 1.757 | 1.428 | 1.536 | -0.371 | 3.444 | 3.930 | 1.245 | 3.266 | 4.124 | 1.895 | 1.756 | Early up | | | | | | | | | |
| B7L74_05235 | ATPase | ATP | -0.868 | -0.925 | -1.305 | -1.422 | -1.493 | -0.209 | -0.601 | 0.563 | 1.086 | 0.736 | 0.233 | | | | | | | | | | |
| B7L74_05240 | DUF4440 domain-containing protein | Uncharacterized CB specific prote | 0.772 | 1.050 | -0.744 | -2.052 | 2.548 | 3.034 | -1.431 | 0.759 | 1.877 | 0.623 | -0.088 | Early up | | | | | | | | | |
| B7L74_05245 | hypothetical protein | | 2.986 | -0.291 | 2.183 | -0.029 | 3.753 | 4.019 | 1.274 | 1.070 | 2.850 | 1.641 | 1.617 | Early up | | | | | | | | | |
| B7L74_05250 | hypothetical protein | | 4.122 | 3.046 | 0.997 | -0.488 | 3.844 | 3.314 | 2.008 | 4.063 | 5.216 | 2.207 | 2.266 | Early up | | | | | | | | | |
| B7L74_05255 | nucleotidyltransferase | | -0.143 | 1.228 | 0.319 | 0.327 | 0.560 | 0.065 | -0.087 | 1.535 | 0.807 | 0.012 | -0.290 | | | | | | | | | | |
| B7L74_05260 | nucleotidyltransferase | | 0.040 | 0.838 | -0.034 | -1.806 | 2.640 | 2.732 | 0.447 | 1.487 | 2.547 | -1.280 | -0.910 | | | | | | | | | | |
| B7L74_05265 | quinone oxidoreductase | | 0.763 | -0.001 | 1.325 | 0.662 | 0.560 | 1.422 | 1.515 | 3.249 | 3.584 | 0.167 | 1.002 | Late up | | | | | | | | | |
| B7L74_05270 | hypothetical protein | | 1.546 | 1.457 | -0.274 | 1.868 | 2.050 | 2.551 | 0.890 | 4.412 | 4.687 | -0.295 | -0.024 | Late up | | | | | | | | | |
| B7L74_05275 | hypothetical protein | Effector protein | -0.660 | 0.779 | -0.252 | 1.177 | 1.019 | 0.015 | -0.558 | 1.561 | 2.418 | -0.249 | -0.048 | | | Yes | | | | | | | |
| B7L74_05280 | ATPase | ATP | -0.982 | 0.482 | -0.387 | -0.453 | 0.413 | 0.626 | -1.011 | 0.603 | -0.672 | -1.396 | -1.686 | | | | | | | | | | |
| B7L74_05285 | hypothetical protein | | 1.491 | 1.305 | 2.192 | 1.401 | 2.121 | 1.607 | 1.203 | 1.267 | 1.292 | 0.235 | 0.349 | | | | | | | | | | |
| B7L74_05290 | non-ribosomal peptide synthetase module-containing p | | 2.163 | 1.829 | 2.590 | 0.113 | 3.402 | 2.910 | 3.107 | 3.188 | 3.034 | 1.360 | 0.903 | Early up | | | | | | | | | |
| B7L74_05295 | hypothetical protein | | 0.274 | 1.786 | 1.748 | 0.081 | 0.593 | 0.428 | 0.252 | 3.602 | 2.242 | -0.221 | 0.288 | | | | | | | | | | |
| B7L74_05300 | lysozyme | | -0.040 | 0.312 | 1.426 | 2.153 | 0.678 | 0.169 | 0.302 | 1.150 | 1.258 | 0.698 | 0.653 | Early up | | | | | | | | | |
| B7L74_05305 | hypothetical protein | | 0.243 | 0.598 | 0.492 | -0.283 | 0.082 | 1.613 | 0.889 | 3.404 | 3.959 | -0.585 | 0.251 | Late up | | | | | | | | | |
| B7L74_05310 | tRNA-Gly | tRNA | 1.304 | 1.086 | 3.015 | 0.202 | 1.278 | 1.236 | 2.705 | 2.764 | 1.280 | 0.566 | 1.317 | Early up | | | | | | | | | |
| B7L74_05315 | CDP-diacylglycerol--glycerol-3-phosphate 3-phosphati | Glycerophospholipid metabolism | 3.752 | 1.440 | 2.974 | 1.546 | 5.288 | 5.696 | 2.397 | 1.940 | 5.125 | 3.086 | 1.730 | Early up | | | | | | | | | |
| B7L74_05320 | nicotinate phosphoribosyltransferase | | 0.057 | 0.218 | 0.117 | -0.083 | 0.556 | 0.409 | 0.047 | 0.159 | 0.894 | 0.369 | 0.133 | | | | | | | | | | |
| B7L74_05325 | isochorismatase | Nicotinate and nicotinamide metab | 0.971 | 0.724 | 1.405 | 2.391 | 0.160 | 1.387 | 1.883 | 3.551 | 3.725 | 1.393 | 1.700 | Late up | | | | | | | | | |
| B7L74_05330 | acriflavine resistance protein B | | -0.825 | 1.229 | -0.595 | -1.004 | 0.026 | 3.447 | -0.245 | 1.419 | 1.944 | -0.842 | -0.953 | | | | | | | | | Yes (drug resistance) | |
| B7L74_05335 | tRNA-Arg | tRNA | 0.894 | 0.360 | 1.066 | 2.031 | 0.815 | 1.867 | 2.105 | 1.532 | 1.351 | 0.918 | 1.441 | | | | | | | | | | |
| B7L74_05340 | cytochrome o ubiquinol oxidase subunit IV | Respiration/Cyt C | -2.100 | -1.304 | -5.736 | -1.471 | -2.625 | -1.958 | -2.441 | -2.218 | -1.971 | -1.344 | -1.817 | Early down | | | | | | Yes | | | |
| B7L74_05345 | cytochrome o ubiquinol oxidase subunit III | Respiration/Cyt C | -1.329 | -2.063 | -2.204 | -1.767 | -2.955 | -2.015 | -1.482 | -2.208 | -2.069 | -1.406 | -0.482 | Early down | | | | | | Yes | | | |
| B7L74_05350 | cytochrome ubiquinol oxidase subunit I | Respiration/Cyt C, Ion Binding | 2.775 | 1.781 | 0.912 | 2.042 | 3.061 | 0.422 | 2.442 | 0.864 | 2.082 | 2.468 | -0.038 | | | | | | | Yes | | | |
| B7L74_05355 | cytochrome ubiquinol oxidase subunit II | Respiration/Cyt C | -1.208 | -1.262 | -1.203 | 0.479 | -2.465 | -1.259 | -0.269 | -0.709 | -0.081 | 0.921 | 2.227 | | | | | | | Yes | | | |
| B7L74_05360 | hypothetical protein | | -0.966 | -0.644 | -0.650 | 0.729 | -1.306 | -1.586 | -0.740 | -0.343 | -0.678 | 0.249 | -0.158 | | | | | | | | | | |
| B7L74_05365 | hypothetical membrane spanning protein | | 0.090 | 0.848 | -0.035 | -1.750 | 2.737 | 1.627 | 0.527 | 1.496 | 2.628 | 0.632 | -0.897 | | | | | | | | | | |
| B7L74_05370 | two-component system response regulator UvrY | Potential transcriptional regulators | 0.054 | -0.667 | 0.268 | 0.214 | -0.636 | -0.623 | 0.225 | -1.042 | -1.153 | -0.021 | 0.453 | | | | | | | | | | |
| B7L74_05375 | hypothetical protein | | 5.805 | 1.058 | 1.871 | 0.142 | 6.582 | 6.670 | 3.507 | 2.537 | 5.554 | 2.858 | 0.977 | Early up | | | | | | | | | |
| B7L74_05380 | hypothetical protein | | -0.485 | 0.289 | -3.032 | -0.001 | 2.468 | 2.710 | 1.186 | 2.286 | 1.707 | -0.719 | -1.952 | | | | | | | | | | |
| B7L74_05385 | hypothetical protein | | 0.222 | -0.490 | 0.078 | -0.119 | -1.107 | 0.406 | 0.434 | 1.105 | 1.260 | 0.677 | 0.649 | | | | | | | | | | |
| B7L74_05390 | hypothetical protein | | -2.438 | -0.542 | -1.530 | -4.287 | 1.023 | 1.371 | -0.768 | -1.006 | 0.069 | -0.996 | -0.098 | | | | | | | | | | |
| B7L74_05395 | hypothetical protein | | -3.726 | -1.324 | -3.850 | -4.603 | 1.549 | 1.065 | -3.289 | -2.320 | -0.137 | -3.351 | -4.713 | | | | | | | | | | |
| B7L74_05400 | hypothetical protein | | 0.110 | -0.153 | 2.235 | -0.823 | 0.761 | 1.142 | 1.258 | 1.955 | 2.520 | 2.033 | 2.282 | Late up | | | | | | | | | |
| B7L74_05405 | tRNA-Ser | tRNA | 0.925 | -0.228 | 2.112 | 1.234 | 0.474 | 1.219 | 2.299 | 1.383 | 1.402 | 2.016 | 3.506 | | | | | | | | | | |
| B7L74_05410 | carbon storage regulator (csrA) | Signal transduction, Two-compone | 1.356 | 1.764 | 4.161 | -1.028 | 1.538 | 1.792 | 2.979 | 1.891 | 2.695 | 1.864 | 2.450 | Early up | | | | | | | | | |
| B7L74_05415 | aspartate kinase | | -0.167 | -0.375 | -0.787 | -0.762 | -0.516 | -0.107 | -0.039 | -1.063 | -1.058 | 0.266 | 0.327 | | | | | | | | | | |
| B7L74_05420 | alanine--tRNA ligase | | -0.949 | -0.848 | -0.924 | -1.291 | -0.719 | 0.310 | -0.121 | -1.252 | -1.456 | -0.008 | 0.155 | | | | | | | | | | |
| B7L74_05425 | recombinase RecX | | 0.452 | -0.037 | 1.226 | -0.251 | 0.044 | 0.853 | 1.078 | -1.035 | -0.600 | 0.859 | 0.958 | | | | | | | | | | |
| B7L74_05430 | DNA recombination/repair protein RecA | DNA | -0.747 | -0.595 | 2.004 | 0.231 | -0.082 | 0.283 | 1.289 | -0.699 | -1.112 | 1.533 | 1.318 | | | | | | | | | | |
| B7L74_05435 | damage-inducible protein CinA | | -0.500 | -0.263 | 1.440 | -0.842 | 0.217 | 0.964 | 0.504 | 0.059 | 1.444 | 1.134 | 1.000 | | | | | | | | | | |
| B7L74_05440 | DNA mismatch repair protein MutS | DNA | -0.418 | 0.010 | 1.886 | -1.071 | 0.887 | 1.291 | 1.784 | 0.755 | 2.336 | 0.698 | 0.577 | | | | | | | | | | |
| B7L74_05445 | glycosyl transferase | | 0.095 | 2.940 | 5.848 | -0.337 | 7.807 | 5.941 | 7.547 | 3.585 | 6.033 | 0.641 | 1.696 | Continuous up | | | | | | | | | |
| B7L74_05450 | hypothetical protein, Hypothetical membrane spanning | Membrane | -0.045 | 0.032 | -0.399 | -1.104 | 0.288 | 0.933 | -0.003 | -0.031 | 1.253 | 0.508 | -0.008 | | | | | | | | | | |
| B7L74_05455 | 23S rRNA pseudouridylate synthase B | Ribosome biogenesis | -0.169 | -1.193 | -1.285 | 0.686 | -2.476 | -2.361 | 0.091 | -1.796 | -4.992 | -0.397 | -0.521 | Early down | | | | | | | | | |
| B7L74_05460 | SMC-Scp complex subunit ScpB. (segregation/conden | Chromosome and associated prote | -0.713 | -0.994 | -0.982 | -1.085 | -1.761 | -0.859 | -0.189 | -1.639 | -2.549 | -0.384 | -0.563 | Early down | | | | | | | | | |
| B7L74_05465 | segregation/condensation protein A | | -0.348 | -0.448 | 1.308 | 0.738 | -0.973 | -0.002 | 0.518 | -0.301 | -0.588 | 1.043 | 0.871 | | | | | | | | | | |
| B7L74_05470 | hypothetical membrane associated protein | | -0.209 | -1.660 | 0.248 | -1.348 | 0.420 | 2.735 | 0.859 | 3.863 | 4.207 | -0.605 | 0.753 | Early up | | | | | | | | | |
| B7L74_05475 | hypothetical protein | Effector protein | 1.610 | 0.439 | 0.511 | -0.114 | 3.333 | 3.987 | 0.144 | 0.898 | 2.852 | 0.905 | -0.005 | Early up | | Yes | | | | | | | |
| B7L74_05480 | hypothetical cytosolic protein | | -0.142 | -0.585 | 1.147 | 1.536 | -0.383 | 0.380 | 1.193 | 0.143 | -0.056 | 0.988 | 0.916 | | | | | | | | | | |

| | | | | | | | | | | | | | | | | | | | | | | | | |
|---|---|---|---|---|---|---|---|---|---|---|---|---|---|---|---|---|---|---|---|---|---|---|---|---|
| B7L74_05485 | 2'-5' RNA ligase | | -0.032 | 0.192 | 0.374 | 0.386 | -0.695 | 0.525 | 0.390 | 0.361 | 0.413 | 0.692 | 0.138 | | | | | | | | | | | |
| B7L74_05490 | SLC13 family permease | | 2.205 | -0.448 | 1.690 | 0.577 | 3.335 | 4.386 | 1.395 | 1.211 | 3.824 | 2.213 | 1.391 | Early up | | | | | | | | | | |
| B7L74_05495 | tRNA 2-thiocytidine(32) synthetase TtcA | | 2.575 | 3.545 | -0.035 | 4.531 | 5.563 | 4.235 | 1.476 | 1.496 | 2.628 | 6.764 | 2.141 | | | | | | | | | | | |
| B7L74_05500 | hypothetical proteinprotein=adenylate cyclase | | 1.063 | 0.536 | 2.904 | 1.725 | 1.400 | 2.590 | 2.055 | 2.966 | 3.323 | 3.244 | 2.561 | Early up | | | | | | | | | | |
| B7L74_05505 | hypothetical protein | Purine metabolism | 0.548 | 0.652 | 1.232 | 0.369 | -0.505 | 1.631 | 1.305 | 4.079 | 4.361 | 2.590 | 1.523 | Late up | | | | | | | | | | |
| B7L74_05510 | intracellular signaling protein | | 1.268 | 0.356 | 1.839 | -1.501 | 1.380 | 3.860 | 2.284 | 1.996 | 4.003 | -1.313 | 0.977 | Early up | | | | | | | | | | |
| B7L74_05515 | adenylate/guanylate cyclase domain-containing protein | Purine metabolism | 0.430 | 0.629 | 0.090 | 0.968 | -0.413 | 0.579 | 0.908 | 2.587 | 2.963 | 1.569 | 0.879 | Late up | | | | | | | | | | |
| B7L74_05520 | DoxX family protein | | 0.145 | 0.159 | 2.236 | 0.869 | 0.032 | 0.357 | 1.438 | 0.154 | -0.079 | 2.236 | 2.002 | Early up | | | | | | | | | | |
| B7L74_05525 | tRNA-guanine(34) transglycosylase | | -0.118 | -0.591 | -0.037 | -0.119 | -1.927 | -0.042 | 0.465 | 0.267 | 0.520 | 0.654 | 1.263 | | | | | | | | | | | |
| B7L74_05530 | hypothetical exported protein | | -0.369 | -0.039 | 0.797 | 0.819 | -0.220 | 0.571 | 0.337 | 0.407 | 0.450 | 1.258 | 0.649 | | | | | | | | | | | |
| B7L74_05535 | hypoxanthine-guanine phosphoribosyltransferase | Purine metabolism | 1.836 | -0.219 | 0.590 | 2.083 | 2.448 | 3.863 | 0.936 | -0.131 | 1.597 | 1.689 | 1.889 | Up in P21 | | | | | | | | | | |
| B7L74_05540 | mechanosensitive ion channel protein MscS | Transporter | 3.987 | -0.234 | 1.138 | -0.803 | 6.998 | 6.812 | 5.121 | 3.312 | 7.434 | 2.963 | 2.009 | Early up | | | | | | | | Yes (ion channels/mechanosensitive channel) | | |
| B7L74_05545 | hypothetical protein | | 3.172 | -0.060 | 1.828 | -0.690 | 2.270 | 3.410 | 1.323 | -0.126 | 2.139 | 1.276 | 1.328 | | | | | | | | | | | |
| B7L74_05550 | IS110 family transposase | | 1.178 | -0.584 | 5.500 | 4.864 | 1.305 | 0.196 | 3.254 | 4.692 | 4.807 | 4.938 | 2.937 | | | | | | | | | | | |
| B7L74_05555 | beta-N-acetylhexosaminidase | | -0.222 | -1.054 | -0.768 | 1.590 | -3.272 | 1.014 | 0.185 | 0.776 | 1.743 | -0.241 | -0.026 | | | | | | | | | | | |
| B7L74_05560 | hypothetical cytosolic protein | | 0.023 | -0.824 | -2.078 | 0.436 | 0.336 | 0.331 | -0.320 | -0.156 | 0.635 | -0.769 | -0.611 | | | | | | | | | | | |
| B7L74_05565 | hypothetical protein | Effector protein | 0.144 | -0.221 | 1.115 | 0.065 | -0.332 | -0.739 | 0.062 | -0.039 | 0.446 | -0.014 | 0.019 | | | Yes | | | | | | | | |
| B7L74_05570 | L%2CD-transpeptidase | | -0.422 | -0.641 | -0.291 | 1.397 | -0.934 | -0.641 | -0.249 | 0.208 | 0.226 | -0.477 | 0.445 | | | | | | | | | | | |
| B7L74_05575 | hypothetical exported protein | | -0.160 | 0.043 | 0.560 | 0.864 | -1.432 | -1.406 | 0.649 | -1.238 | -2.161 | 1.187 | 1.046 | Down in P51 | | | | | | | | | | |
| B7L74_05580 | tRNA (adenosine(37)-N6)-dimethylallyltransferase Mi | | -0.712 | 0.075 | -1.250 | -0.599 | -1.685 | 0.043 | -0.626 | 0.257 | 0.537 | -1.350 | -1.201 | | | | | | | | | | | |
| B7L74_05585 | DNA mismatch repair protein MutL | | -0.497 | -0.212 | -0.748 | 0.085 | -0.876 | -0.558 | -0.626 | -0.117 | -0.139 | -0.656 | -0.816 | | | | | | | | | | | |
| B7L74_05590 | hybrid sensor histidine kinase/response regulator | | 1.003 | -1.069 | -0.727 | -0.525 | 1.483 | 1.277 | 0.413 | -1.354 | 0.335 | 0.297 | -0.043 | | | | | | | | | | | |
| B7L74_05595 | N-acetylmuramoyl-L-alanine amidase | | 2.451 | 0.027 | 0.151 | -0.514 | 3.030 | 2.541 | 0.815 | -0.624 | 3.481 | 1.903 | 1.490 | | | | | | | | | | | |
| B7L74_05600 | IS30 family transposase | Replication and repair | 1.919 | 1.062 | 0.542 | -0.089 | 2.590 | 3.359 | 0.753 | 1.622 | 2.563 | 0.706 | 0.127 | Early up | | | | | | | | | | |
| B7L74_05605 | tRNA (adenosine(37)-N6)-threonylcarbamoyltransfera | ATP, Transfer RNA biogenesis | -1.269 | -0.840 | -7.763 | 0.270 | -3.382 | -0.450 | -1.873 | -0.682 | 0.356 | -1.011 | -1.133 | Early down | | | | | | | | | | |
| B7L74_05610 | bifunctional ADP-dependent NAD(P)H-hydrate dehyc | | 0.653 | 0.291 | 0.051 | 0.818 | -0.626 | -0.641 | 0.809 | -0.784 | -1.264 | 0.909 | 1.680 | | | | | | | | | | | |
| B7L74_05615 | hypothetical protein | | 0.758 | 0.425 | 1.557 | 2.494 | 0.462 | 0.319 | 1.276 | 0.430 | 0.566 | 1.513 | 1.573 | Up in P12 | | | | | | | | | | |
| B7L74_05620 | IS110 family transposase | | -0.409 | -1.066 | 1.692 | 0.007 | -0.830 | 0.236 | 0.617 | -0.650 | -0.848 | 1.071 | 1.521 | | | | | | | | | | | |
| B7L74_05625 | hypothetical protein | | 2.530 | 0.841 | -0.034 | 0.314 | 2.172 | 1.622 | 1.129 | 1.599 | 1.136 | 1.538 | 0.448 | | | | | | | | | | | |
| B7L74_05630 | ribonuclease R | | -0.571 | 0.242 | -1.536 | 0.448 | -0.744 | -0.409 | -0.412 | -0.493 | -0.941 | -1.012 | -0.410 | | | | | | | | | | | |
| B7L74_05635 | lipoprotein | | 0.917 | 0.943 | 0.998 | -0.120 | -0.586 | -0.804 | 0.543 | 0.570 | -0.338 | 1.771 | 1.126 | | | | | | | | | | | |
| B7L74_05640 | multidrug efflux protein | | -0.640 | -0.142 | -2.216 | -1.142 | -0.816 | -0.172 | -1.482 | -1.500 | -0.994 | -1.481 | -1.897 | | | | | | | | | | | |
| B7L74_05645 | MexH family multidrug efflux RND transporter peripl | | -1.015 | 0.332 | 0.299 | -0.413 | -1.383 | -2.498 | -0.666 | -2.401 | -4.104 | -2.165 | -1.718 | | | | | | | | | | | |
| B7L74_05650 | tRNA-Arg | tRNA | 1.670 | 1.421 | 0.941 | -0.427 | 1.896 | 1.986 | 1.228 | 1.298 | 0.684 | -1.208 | -0.297 | | | | | | | | | | | |
| B7L74_05655 | hypothetical exported protein | | -0.465 | 0.792 | 1.131 | -0.904 | 0.937 | 0.387 | -0.125 | 0.270 | 1.371 | -0.688 | -0.298 | | | | | | | | | | | |
| B7L74_05660 | class II fumarate hydratase | | -0.197 | 0.497 | -0.029 | -0.944 | 0.083 | 0.075 | -1.066 | -0.637 | -0.405 | -1.049 | -1.132 | | | | | | Yes | | | | | |
| B7L74_05665 | hypothetical protein | | 0.411 | 0.114 | 1.967 | 0.880 | 0.117 | -0.222 | 0.841 | -0.057 | 0.073 | 0.462 | 1.162 | | | | | | | | | | | |
| B7L74_05670 | acetolactate decarboxylase | | -0.972 | 0.476 | -1.034 | -2.816 | 1.988 | 1.011 | -0.564 | 0.484 | 1.535 | -2.287 | -1.914 | | | | | | | | | | | |
| B7L74_05675 | hypothetical cytosolic protein | | -1.023 | 0.771 | -1.554 | -1.991 | 0.094 | 0.214 | -3.007 | -0.425 | -2.369 | -4.574 | -4.506 | Late down | | | | | | | | | | |
| B7L74_05680 | S26 family signal peptidase | | -0.061 | -0.178 | -1.078 | -0.150 | -0.524 | 0.479 | 0.163 | 0.258 | 0.430 | -0.721 | -0.718 | | | | | | | | | | | |
| B7L74_05685 | hypothetical exported protein | | -0.698 | -0.179 | 0.510 | 0.480 | -0.091 | -0.070 | 2.105 | 0.784 | 0.684 | 0.127 | -0.055 | | | | | | | | | | | |
| B7L74_05690 | hypothetical protein | | -0.325 | 0.704 | 0.262 | -0.530 | 0.819 | 1.418 | 0.727 | 0.671 | 1.968 | -0.599 | -0.632 | | | | | | | | | | | |
| B7L74_05695 | hypothetical protein | | 5.876 | 2.224 | 4.163 | -1.839 | 8.097 | 8.110 | 3.591 | 0.933 | 7.900 | 3.546 | 1.188 | Early up | | | | | | | | | | |
| B7L74_05700 | lytic transglycosylase | | 3.163 | -1.277 | -3.750 | -1.152 | 0.694 | 2.299 | -2.167 | -2.220 | -1.088 | -3.243 | -3.662 | | | | | | | | | | | |
| B7L74_05705 | IS30 family transposase | Replication and repair | 1.532 | 1.152 | 0.729 | 0.060 | 1.941 | 2.725 | 0.597 | 1.435 | 2.075 | 0.732 | 0.271 | Early up | | | | | | | | | | |
| B7L74_05710 | hypothetical protein | | 0.519 | 0.857 | 2.028 | 0.442 | -1.609 | 1.173 | 1.738 | 1.819 | 2.142 | 1.124 | 1.504 | | | | | | | | | | | |
| B7L74_05715 | hypothetical protein | | -1.868 | 1.182 | -1.154 | -1.288 | 0.949 | 1.574 | -0.251 | 2.203 | 2.140 | -1.112 | -1.572 | | | | | | | | | | | |
| B7L74_05720 | hypothetical protein | | 1.926 | 0.040 | -0.270 | 1.089 | 3.874 | 2.753 | 1.599 | 1.449 | 2.644 | 1.033 | 0.940 | Early up | | | | | | | | | | |
| B7L74_05725 | murein transglycosylase | | -0.815 | -1.409 | 0.950 | 0.577 | -0.959 | -0.797 | -0.702 | -1.139 | -0.505 | -0.746 | -0.706 | | | | | | | | | | | |
| B7L74_05730 | hypothetical protein with endo/excinuclease domain | Replication and repair | 2.376 | 0.602 | 4.286 | 3.361 | 3.677 | 2.520 | 2.679 | -1.354 | 2.056 | 3.210 | 3.974 | Early up | | | | | | | | | | |
| B7L74_05735 | hypothetical protein | | 0.058 | -0.181 | 0.445 | -0.564 | -0.404 | 0.021 | 0.339 | 0.246 | 0.100 | 0.060 | 0.245 | | | | | | | | | | | |
| B7L74_05740 | hypothetical protein | | -0.860 | -0.007 | 0.499 | 2.204 | -0.216 | -0.280 | 1.790 | 0.103 | 0.410 | 1.572 | 2.525 | | | | | | | | | | | |
| B7L74_05745 | alanine dehydrogenase | Alanine, aspartate and glutamate m | 0.118 | -1.172 | -1.303 | -0.563 | -1.852 | -1.763 | -0.363 | -1.830 | -2.233 | -0.859 | -0.621 | Early down | | | | | | | | | | |
| B7L74_05750 | electron transfer flavoprotein subunit alpha | Exosome | -1.746 | -1.942 | -2.652 | -1.081 | -3.836 | -1.540 | -1.251 | -1.754 | -1.972 | -1.840 | -1.590 | Early down | | | | | | | | | | |
| B7L74_05755 | electron transfer flavoprotein subunit beta | | -0.659 | -0.906 | -1.624 | 0.850 | -2.361 | -1.471 | -0.612 | -0.685 | -1.115 | 0.007 | 0.049 | | | | | | | | | | | |
| B7L74_05760 | carboxymethylenebutenolidase | | -0.741 | -1.351 | 1.704 | 2.968 | 1.287 | -0.185 | -0.790 | -1.280 | 0.027 | 0.402 | -0.203 | Up in P12 | | | | | | | | | | |
| B7L74_05765 | electron-transferring-flavoprotein dehydrogenase | | 0.397 | -0.025 | 0.473 | -0.352 | -0.320 | 0.287 | 0.231 | -0.442 | -0.751 | -0.012 | -0.402 | | | | | | | | | | | |
| B7L74_05770 | hypothetical protein | | 0.090 | 0.848 | -0.035 | -0.093 | 5.090 | 1.627 | 0.527 | 1.496 | 3.732 | 0.455 | -0.897 | | | | | | | | | | | |
| B7L74_05775 | L%2CD-transpeptidase, EnhAnced entry protein, Enh | Entry | -0.577 | -0.241 | 0.246 | 0.834 | -0.578 | 0.495 | 1.252 | 0.855 | 0.842 | 0.320 | 0.742 | | | | | | | | | | | |
| B7L74_05780 | curved DNA-binding protein | | 0.298 | -0.410 | 0.772 | 1.370 | 0.992 | 1.298 | 0.642 | 0.162 | 0.596 | 1.318 | 1.068 | | | | Yes | | | | | | | |
| B7L74_05785 | chaperone modulatory protein CbpM | | -0.450 | -0.949 | 0.448 | 1.994 | -1.709 | 0.379 | 0.624 | 0.282 | 0.326 | 1.117 | 1.059 | | | | | | | | | | | |
| B7L74_05790 | hypothetical protein | Amino sugar and nucleotide sugar | 1.000 | -0.817 | 0.557 | 2.057 | 0.304 | 2.121 | 1.524 | 2.761 | 3.303 | 2.835 | 1.975 | Continuous up | | | | | | | | | | |
| B7L74_05795 | hypothetical protein | | 0.761 | -0.305 | -1.184 | -0.721 | -1.855 | 0.894 | 0.512 | 0.310 | 0.251 | 0.127 | -0.079 | | | | | | | | | | | |
| B7L74_05800 | HPP family protein | | 0.563 | 0.076 | -0.318 | 0.689 | -0.827 | -0.160 | 0.170 | -0.374 | -2.187 | 1.289 | 1.042 | | | | | | | | | | | |
| B7L74_05805 | hypothetical protein IscU | | -0.541 | 0.246 | -0.629 | -1.551 | 2.680 | 1.492 | 1.991 | 1.362 | 2.861 | -1.100 | -0.481 | | | | | | | | | | | |
| B7L74_05810 | IscS subfamily cysteine desulfurase | Thiamine metabolism | 0.764 | -0.758 | 0.781 | 0.696 | -1.011 | 2.866 | 0.494 | -0.619 | -0.447 | 0.903 | 0.499 | Up in P21 | | | | | | | | | | |
| B7L74_05815 | oligopeptide transporter%2C OPT family | Transporter | 2.016 | 0.363 | 0.314 | 0.159 | 4.411 | 3.733 | 0.388 | 0.045 | 4.303 | 1.408 | 1.183 | Early up | | | | | | | | Yes (Oligopeptide) | | |
| B7L74_05820 | tRNA (cytosine(32)/uridine(32)-2'-O)-methyltransferas | | 1.483 | 0.717 | 1.767 | 1.231 | -0.436 | -1.194 | 1.224 | 0.088 | -1.584 | 1.669 | 1.619 | | | | | | | | | | | |
| B7L74_05825 | hypothetical protein | | -1.850 | 0.408 | -0.454 | 1.983 | -2.317 | 1.277 | 0.862 | 0.698 | 1.986 | 0.606 | 0.598 | | | | | | | | | | | |
| B7L74_05830 | inositol monophosphatase | Inositol phosphate metabolism | 1.125 | -1.172 | -0.260 | -3.955 | 0.207 | 2.611 | -2.280 | 3.417 | 3.201 | -2.460 | 0.705 | Late up | | | | | | | | | | |
| B7L74_05835 | hypothetical membrane associated protein | | 0.543 | 0.158 | 1.228 | 1.870 | -0.542 | -0.329 | 1.057 | 0.051 | -0.519 | 0.703 | 0.578 | | | | | | | | | | | |
| B7L74_05840 | hypothetical exported protein | Exported | 0.904 | 0.722 | 1.990 | -0.364 | 0.124 | 0.526 | 0.543 | -0.068 | -0.196 | 0.846 | 0.196 | | | | | | | | | | | |
| B7L74_05845 | hypothetical protein, Enhanced entry protein enhC, tet | Entry | 4.429 | 0.180 | 3.002 | 0.861 | 0.947 | 1.153 | 3.386 | 1.432 | 2.309 | 4.205 | 3.621 | Early up | | | | | | | | | | |
| B7L74_05850 | enhanced entry protein | Entry | 0.035 | 0.837 | -0.034 | -1.810 | 2.635 | 1.620 | 0.443 | 1.486 | 2.542 | -1.283 | -0.912 | | | | | | | | | | | |
| B7L74_05855 | L%2CD-transpeptidase, Enhanced entry protein | Entry | 2.561 | 1.517 | 2.085 | -0.816 | 1.268 | -0.175 | 2.187 | 1.624 | 1.643 | 2.263 | 2.710 | | | | | | | | | | | |
| B7L74_05860 | hypothetical protein | | 0.613 | 0.802 | 0.495 | -1.106 | 0.385 | 1.074 | 0.594 | 2.159 | -0.803 | 0.706 | 0.309 | | | | | | | | | | | |
| B7L74_05865 | hypothetical membrane associated protein | | 3.902 | 0.808 | 3.557 | 1.537 | 2.296 | 3.498 | 4.641 | 5.057 | 4.464 | 3.660 | 3.904 | | | | | | | | | | | |
| B7L74_05870 | hypothetical protein | | 1.332 | 1.001 | 1.284 | 2.753 | 0.690 | 2.349 | 1.683 | 2.558 | 3.115 | 2.509 | 2.293 | Early up | | | | | | | | | | |
| B7L74_05875 | hypothetical protein | | 0.386 | 0.212 | -0.707 | -0.637 | -0.177 | 2.374 | -0.441 | 0.212 | 3.313 | -0.883 | -1.740 | | | | | | | | | | | |
| B7L74_05880 | protein translocase subunit SecF | Protein export/sec pathway | -1.424 | -1.005 | -4.047 | -1.377 | -0.959 | 0.089 | -2.468 | -2.695 | -3.817 | -2.533 | -2.912 | Early down | | | | | | | | | | |
| B7L74_05885 | protein translocase subunit SecD | | 1.764 | 3.200 | -1.035 | -1.719 | 2.021 | 3.283 | 0.559 | 3.396 | 2.974 | -0.230 | 0.695 | | | | | | | | | | | |
| B7L74_05890 | preprotein translocase subunit YajC | Protein export/sec pathway | -1.826 | -0.995 | -3.370 | -0.535 | -2.036 | -2.003 | -2.227 | -3.027 | -4.537 | -2.442 | -2.630 | Continuous down | | | | | | | | | | |
| B7L74_05895 | hypothetical membrane associated protein | | -0.131 | -0.475 | -1.592 | 0.096 | -0.658 | -0.450 | -0.267 | -2.034 | -2.615 | -0.613 | -1.088 | Late down | | | | | | | | | | |
| B7L74_05900 | LanM family lanthionine synthetase | | -0.933 | -0.315 | 0.637 | 0.800 | -0.532 | -0.073 | -0.754 | 1.085 | 0.660 | -0.426 | -0.960 | | | | | | | | | | | |
| B7L74_05905 | tRNA 2-thiouridine(34) synthase MnmA | | 0.046 | -0.258 | -1.155 | -0.511 | -1.231 | -0.432 | -1.248 | 0.162 | -0.677 | -0.731 | -1.010 | | | | | | | | | | | |
| B7L74_05910 | transcription-repair coupling factor | | -0.114 | -0.328 | -1.989 | -1.332 | 0.536 | 1.117 | -0.742 | -0.246 | 0.652 | -1.189 | -1.734 | | | | | | | | | | | |
| B7L74_05915 | hypothetical protein | | 3.318 | 0.232 | 1.713 | -0.025 | 3.501 | 3.495 | 1.175 | 2.696 | 3.183 | 1.225 | 0.675 | Early up | | | | | | | | | | |
| B7L74_05920 | HAD family hydrolase | | 0.001 | 0.786 | -1.067 | 1.781 | -0.466 | 0.562 | -0.654 | 1.134 | 0.597 | 0.078 | 0.031 | | | | | | | | | | | |
| B7L74_05925 | anthranilate synthase component I | Phenylalanine, tyrosine and tryptop | 0.237 | 0.642 | -1.467 | -0.828 | -0.611 | 1.118 | 0.780 | 2.418 | 2.219 | -1.019 | -0.784 | Late up | | | | | | | | | | |
| B7L74_05930 | aminodeoxychorismate/anthranilate synthase compone | | -3.269 | -0.102 | -2.386 | -3.147 | 0.569 | -0.415 | -1.985 | 1.199 | -0.759 | -2.759 | -3.403 | | | | | | | | | | | |
| B7L74_05935 | hypothetical protein | Phenylalanine, tyrosine and tryptop | 2.841 | 2.640 | 0.551 | 1.028 | 1.513 | 3.589 | 2.116 | 4.601 | 4.956 | -1.186 | 0.898 | Early up | | | | | | | | | | |
| B7L74_05940 | hypothetical protein | | -0.407 | 0.386 | -1.132 | -1.689 | 0.708 | 1.589 | -1.875 | 2.004 | 1.357 | -1.620 | -1.619 | | | | | | | | | | | |
| B7L74_05945 | hypothetical protein | Phenylalanine, tyrosine and tryptop | -0.935 | 1.118 | -1.995 | -1.695 | 0.983 | 0.618 | -1.366 | 0.317 | -0.653 | -2.954 | -3.133 | Late down | | | | | | | | | | |
| B7L74_05950 | indole-3-glycerol phosphate synthase | Phenylalanine, tyrosine and tryptop | -0.934 | 0.373 | -1.321 | -2.092 | -0.869 | -0.181 | -1.408 | -0.170 | -0.832 | -2.503 | -2.813 | Early down | | | | | | | | | | |
| B7L74_05955 | tryptophan synthase subunit beta | | 0.000 | -1.899 | -0.872 | -2.705 | -1.827 | -0.614 | -0.997 | 0.595 | 0.484 | -1.667 | -2.117 | | | | | | | | | | | |
| B7L74_05960 | tryptophan synthase subunit alpha | | -0.075 | 0.189 | -0.915 | -1.333 | -0.264 | 0.266 | -0.700 | -0.176 | -0.007 | -1.098 | -1.095 | | | | | | | | | | | |
| B7L74_05965 | lipoprotein | | 0.089 | 0.847 | -0.035 | -0.788 | 6.539 | 2.589 | 4.835 | 1.496 | 2.628 | -0.331 | -0.897 | | | | | | | | | | | |
| B7L74_05970 | 7-dehydrocholesterol reductase | Eucaryotic-like ORFs. | 1.303 | 0.134 | -0.202 | -1.044 | 1.155 | 2.122 | 0.244 | 1.056 | 2.484 | 0.312 | -0.770 | | | | | | | | | | | |
| B7L74_05975 | hypothetical cytosolic protein | Eucaryotic-like ORFs. | 3.605 | 0.225 | 1.480 | -0.685 | 4.911 | 4.794 | 0.639 | 0.007 | 4.046 | 1.889 | -0.977 | Early up | | | | | | | | | | |
| B7L74_05980 | DUF1658 domain-containing protein | CB specific protein family | -0.376 | 0.233 | -0.571 | -1.021 | -0.523 | -0.070 | -1.334 | -0.095 | -0.323 | -2.863 | -2.621 | Late down | | | | | | | | | | |
| B7L74_05985 | MFS transporter | Transporter | 0.006 | -0.184 | -1.857 | -1.075 | -0.672 | -0.340 | -0.475 | -0.164 | -0.596 | -1.431 | -1.704 | | | | | | | | | | | |
| B7L74_05990 | hypothetical protein | | -1.012 | -0.117 | -0.505 | -1.076 | -0.394 | 0.010 | -0.599 | -0.394 | -1.407 | -1.130 | -1.582 | | | | | | | | | | | |
| B7L74_05995 | hypothetical protein | | -0.281 | -0.363 | 0.080 | -0.556 | 0.084 | 1.275 | -0.169 | 1.430 | 0.333 | -0.537 | -0.745 | | | | | | | | | | | |
| B7L74_06000 | hypothetical protein | | 0.680 | 0.765 | -0.120 | -0.879 | 0.252 | 0.773 | 0.683 | 1.704 | 1.081 | -0.771 | 0.607 | | | | | | | | | | | |
| B7L74_06005 | tRNA-Arg | tRNA | 0.946 | 1.383 | 3.776 | -0.790 | 1.527 | 2.529 | 2.958 | 1.279 | 0.054 | -0.148 | 1.036 | Early up | | | | | | | | | | |
| B7L74_06010 | protein kinase | | -1.089 | 0.160 | -2.281 | -1.702 | -0.277 | 0.077 | -1.364 | 0.312 | 0.230 | -1.039 | -1.323 | | | | | | | | | | | |
| B7L74_06015 | heat-shock protein Hsp20 | Protein processing in endoplasmic | 0.287 | -0.198 | 3.167 | 0.138 | 1.882 | 0.967 | 1.318 | -0.066 | -0.105 | -0.247 | 0.657 | Early up | | | | | | | | | | |
| B7L74_06020 | DUF378 domain-containing protein | | 1.714 | 2.949 | 5.720 | -0.795 | 3.410 | 2.775 | 5.328 | 4.905 | 5.630 | 1.395 | 5.009 | Early up | | | | | | | | | | |
| B7L74_06025 | secondary thiamine-phosphate synthase | | 1.116 | 1.874 | 2.275 | -0.665 | 1.373 | 0.771 | 2.129 | 3.001 | 3.074 | 0.743 | 1.847 | Late up | | | | | | | | | | |
| B7L74_06030 | RNA-splicing ligase RtcB | | 0.949 | 1.120 | 1.659 | -1.093 | 0.540 | 0.233 | 0.558 | 0.291 | 0.098 | 0.387 | 0.276 | | | | | | | | | | | |
| B7L74_06035 | hypothetical protein | | 0.430 | 1.251 | 2.266 | -0.927 | 0.839 | 0.652 | 0.832 | 1.190 | 0.736 | 0.809 | 0.636 | | | | | | | | | | | |
| B7L74_06040 | archease | Transfer RNA biogenesis | 0.645 | 0.528 | 2.047 | 0.728 | 0.736 | -0.062 | 1.020 | 0.634 | 0.774 | 1.833 | 2.376 | Early up | | | | | | | | | | |
| B7L74_06045 | hypothetical membrane spanning protein | | -0.082 | 0.164 | 1.714 | 0.448 | 0.244 | 0.420 | 0.934 | 1.058 | 0.731 | 1.327 | 1.898 | | | | | | | | | | | |
| B7L74_06050 | deoxyribodipyrimidine photolyase | DNA repair and recombination pro | 2.454 | 0.576 | 1.120 | -0.153 | 3.576 | 4.392 | 1.391 | 1.375 | 2.891 | 1.475 | 1.236 | Early up | | | | | | | | | | |
| B7L74_06055 | hypothetical cytosolic protein | | 0.874 | 1.123 | 1.167 | -1.005 | 1.261 | 1.219 | 0.739 | 1.544 | 0.768 | -0.290 | -0.336 | | | | | | | | | | | |
| B7L74_06060 | hypothetical protein | | 0.132 | -1.017 | 1.320 | 2.186 | 0.287 | 1.585 | 0.563 | 2.031 | 2.565 | 1.284 | 0.721 | Early up | | | | | | | | | | |
| B7L74_06065 | Bcr/CflA family drug resistance efflux transporter | Transporter | 0.829 | -0.152 | 2.481 | 1.153 | 1.140 | 1.246 | 2.378 | 0.716 | 2.649 | 2.569 | 2.826 | Late up | | | | | | | | Yes (drug resistance) | | |
| B7L74_06070 | hypothetical protein | | 2.332 | 0.587 | 0.271 | 0.607 | 1.342 | 3.236 | -0.683 | 2.055 | 3.079 | 1.176 | -0.701 | | | | | | | | | | | |
| B7L74_06075 | tRNA 4-thiouridine(8) synthase ThiI | Thiamine metabolism | 1.204 | 0.146 | 1.796 | 2.519 | 2.205 | 3.638 | 2.193 | 3.667 | 3.993 | 1.833 | 1.543 | Continuous up | | | | | | | | | | |
| B7L74_06080 | class V aminotransferase | Thiamine metabolism | 3.015 | -0.333 | 2.940 | 2.621 | 3.928 | 4.643 | 2.607 | 2.226 | 2.843 | 3.050 | 3.219 | Early up | | | | | | | | | | |
| B7L74_06085 | RNA-binding protein | | 0.728 | -0.812 | 0.139 | 2.262 | -0.818 | -1.337 | 1.127 | -2.020 | -2.862 | 2.141 | 2.070 | Variable | | | | | | | | | | |
| B7L74_06090 | 1-acyl-sn-glycerol-3-phosphate acyltransferase | | 0.420 | -0.677 | 0.494 | 1.657 | -0.374 | -0.227 | 1.071 | 0.149 | -0.415 | 1.594 | 1.426 | | | | | | | | | | | |
| B7L74_06095 | excinuclease ABC subunit C | | 0.103 | -0.757 | 0.699 | 0.902 | -0.601 | 0.371 | 0.482 | 0.764 | 0.398 | 0.713 | 1.499 | | | | | | | | | | | |
| B7L74_06100 | tRNA-Ser | tRNA | 1.776 | 0.417 | 2.248 | 2.541 | 0.610 | 0.895 | 2.097 | 0.384 | -0.943 | 1.099 | 1.983 | Early up | | | | | | | | | | |
| B7L74_06105 | IS110 family transposase | | 0.072 | -0.504 | 2.330 | 0.285 | -0.895 | 1.334 | 1.359 | 0.503 | -0.847 | 1.727 | 2.125 | | | | | | | | | | | |
| B7L74_06110 | hypothetical protein | | -1.377 | 1.844 | 0.941 | 1.130 | 2.351 | 4.186 | 0.708 | 1.897 | 2.148 | -1.787 | -1.384 | | | | | | | | | | | |
| B7L74_06115 | hypothetical exported protein | Transposases harbored by IS1111 | 0.122 | -0.820 | 0.461 | 0.408 | -0.368 | -0.113 | 0.300 | 0.522 | 0.207 | -0.350 | -0.235 | | | | | | | | | | | |
| B7L74_06120 | serine--tRNA ligase | Transposases harbored by IS1111 | -1.042 | -1.563 | -4.128 | -2.214 | -3.455 | -2.309 | -2.457 | -6.991 | -5.208 | -2.285 | -2.600 | Late down | | | | | | | | | | |
| B7L74_06125 | replication-associated recombination protein A | Transposases harbored by IS1111 | -0.257 | -1.010 | -0.655 | -1.297 | -0.828 | -1.146 | -0.347 | -0.973 | -2.992 | -1.243 | -1.383 | Early down | | | | | | | | | | |
| B7L74_06130 | outer-membrane lipoprotein carrier protein LolA | Membrane, Structural proteins | 0.899 | 0.841 | -0.034 | 0.615 | 2.664 | 1.623 | 0.855 | 1.491 | 6.242 | -1.269 | -0.905 | Up in P51 | | | | | | | | | | |
| B7L74_06135 | DNA translocase FtsK | Translocase<br>Chromosome and associated proteins | -1.373 | -0.987 | -2.156 | -1.426 | -1.742 | -1.451 | -1.781 | -1.925 | -2.260 | -1.605 | -1.469 | Early down | | | | | | | | | | |
| B7L74_06140 | thioredoxin-disulfide reductase | Transposases harbored by IS1111 | -0.602 | -1.135 | -1.929 | -0.236 | -2.073 | -1.627 | -1.000 | -1.545 | -2.735 | -0.981 | -0.376 | Early down | | | | | | | | | | |
| B7L74_06145 | hypothetical membrane spanning protein | Transposases harbored by IS1111 | -2.084 | -1.180 | -1.639 | -1.557 | -2.200 | -1.257 | -0.412 | -1.010 | -1.867 | -1.079 | -1.376 | Early down | | | | | | | | | | |
| B7L74_06150 | translation initiation factor IF-1 | Transposases harbored by IS1111 | 1.226 | 0.788 | 0.816 | 2.440 | 0.584 | 0.508 | 0.342 | 0.361 | 0.066 | 1.372 | 1.147 | Up in P12 | | | | | | | | | | |
| B7L74_06155 | ATP-dependent Clp protease ATP-binding subunit Cl | ATP | 1.022 | 1.225 | 1.182 | -1.544 | 2.497 | 2.106 | 0.919 | 0.750 | 2.481 | 0.229 | 0.191 | | | | Yes | | | | | | | |
| B7L74_06160 | hypothetical membrane associated protein | Transposases harbored by IS1111 | 1.064 | 1.336 | 3.918 | 0.180 | 2.587 | 2.232 | 4.214 | 3.122 | 3.292 | 1.860 | 3.450 | Early up | | | | | | | | | | |
| B7L74_06165 | hypothetical cytosolic protein | Effector protein | 0.877 | 0.184 | 1.355 | 1.153 | 0.205 | -0.497 | 0.489 | 0.359 | -0.613 | 1.609 | 0.930 | | | Yes | | | | | | | | |
| B7L74_06170 | hypothetical protein | Transposases harbored by IS1111 | 0.511 | -0.306 | 0.192 | 0.055 | -0.881 | -0.215 | -0.566 | 0.572 | -0.331 | -0.339 | -0.320 | | | | | | | | | | | |
| B7L74_06175 | isocitrate dehydrogenase (NADP(+)) | Transposases harbored by IS1111 | -0.755 | -1.253 | -1.515 | 0.454 | -1.655 | -1.159 | -1.259 | -1.618 | -2.324 | -1.529 | -1.090 | Early down | | | | | Yes | | | | | |
| B7L74_06180 | tRNA preQ1(34) S-adenosylmethionine ribosyltransfe | Transposases harbored by IS1111 | -0.125 | 0.144 | -0.127 | 1.184 | 0.203 | 0.632 | 0.499 | 1.049 | -0.354 | 0.836 | 1.221 | | | | | | | | | | | |
| B7L74_06185 | hypothetical membrane spanning protein | Transposases harbored by IS1111 | -0.767 | -0.177 | 1.344 | 0.927 | -0.284 | 0.013 | -0.604 | 0.361 | 0.779 | -0.463 | -0.646 | | | | | | | | | | | |
| B7L74_06190 | FMN-binding glutamate synthase family protein | Transposases harbored by IS1111 | 0.023 | -0.999 | 0.110 | 0.917 | -0.960 | -0.332 | -0.048 | 0.212 | 0.040 | 0.005 | -0.478 | | | | | | | | | | | |
| B7L74_06195 | succinate-semialdehyde dehydrogenase | Transposases harbored by IS1111 | -0.043 | -0.597 | -0.709 | 0.392 | -0.934 | -0.520 | -0.192 | -0.548 | -1.479 | -0.499 | -0.992 | | | | | | | | | | | |
| B7L74_06200 | delta(24(24(1)))-sterol reductase | | -2.062 | -3.621 | 0.344 | -2.345 | 0.427 | 1.622 | 0.396 | 0.501 | 2.590 | -1.166 | 0.288 | | | | | | | | | | | |
| B7L74_06205 | hypothetical protein | Transposases harbored by IS1111 | -0.496 | -0.804 | 1.369 | -2.346 | 2.708 | 1.295 | -0.825 | 0.902 | 2.687 | -1.843 | 1.450 | | | | | | | | | | | |
| B7L74_06210 | MFS transporter | Transporter | 0.002 | 0.406 | 0.777 | 1.148 | -0.455 | 0.673 | 0.453 | 1.710 | 1.754 | 0.257 | 0.081 | | | | | | | | | Yes (drug resistance) | | |
| B7L74_06215 | hypothetical membrane spanning protein | | 3.435 | 1.736 | 2.637 | -0.392 | 5.345 | 4.563 | 1.670 | 1.818 | 4.657 | 0.600 | 1.193 | Early up | | | | | | | | | | |
| B7L74_06220 | hypothetical protein | | 9.127 | 5.353 | 6.779 | 0.100 | 10.538 | 10.927 | 6.061 | 3.444 | 10.422 | 5.413 | 2.923 | Continuous up | | | | | | | | | | |
| B7L74_06225 | hypothetical proteinprotein=ankyrin repeat protein An | Effector protein | -0.560 | 1.141 | 0.025 | -1.018 | 1.700 | 1.748 | 0.777 | 2.375 | 1.780 | -0.193 | 0.038 | | | Yes | | | | | | | | |
| B7L74_06230 | hypothetical cytosolic protein | | 1.414 | 1.292 | -0.035 | 0.281 | 0.287 | -1.815 | 0.057 | 2.716 | 2.250 | -0.341 | -2.377 | | | | | | | | | | | |
| B7L74_06235 | hypothetical protein | | -2.573 | -0.316 | 1.281 | -1.605 | 2.336 | 1.072 | 1.823 | 1.881 | 2.874 | 0.307 | -0.840 | | | | | | | | | | | |
| B7L74_06240 | hypothetical protein | | 0.419 | 0.280 | 1.352 | 0.615 | 0.257 | 0.622 | 1.104 | -0.060 | 1.180 | 1.318 | 2.147 | | | | | | | | | | | |
| B7L74_06245 | hypothetical protein | | 1.951 | 2.487 | -1.486 | 0.618 | 3.904 | 3.717 | 1.269 | 4.023 | 4.013 | 1.259 | 1.538 | | | | | | | | | | | |
| B7L74_06250 | hypothetical membrane spanning protein | Effector protein | 0.005 | -0.026 | 0.499 | -0.194 | 0.323 | 0.982 | 0.156 | 1.312 | 1.975 | -0.611 | -0.533 | | | Yes | | | | | | | | |
| B7L74_06255 | tRNA-Ser | tRNA | -0.082 | 0.497 | 3.243 | 0.850 | 0.855 | 0.109 | 1.978 | 0.848 | -0.042 | 0.629 | 0.479 | | | | | | | | | | | |
| B7L74_06260 | IS110 family transposase | | 0.064 | 0.810 | 1.931 | 0.173 | 0.239 | 1.702 | 1.108 | 0.862 | 0.641 | 0.944 | 0.679 | | | | | | | | | | | |
| B7L74_06265 | hypothetical protein | | 0.811 | -1.971 | -1.264 | -0.342 | -3.520 | -3.668 | -0.626 | -2.929 | -3.629 | -0.422 | -0.598 | | | | | | | | | | | |
| B7L74_06270 | hypothetical cytosolic protein | | -1.570 | -1.062 | -3.861 | -1.434 | -1.733 | -1.251 | -1.746 | -1.019 | -1.502 | -1.561 | -2.029 | Early down | | | | | | | | | | |
| B7L74_06275 | phosphoribosylaminoimidazolesuccinocarboxamide sy | Purine metabolism | -1.157 | -1.157 | -1.656 | 0.126 | -1.984 | -1.400 | -1.039 | -1.564 | -2.253 | -0.853 | -1.065 | Early down | | | | | | | | | | |
| B7L74_06280 | lipoprotein | | -0.532 | -0.383 | 0.993 | -1.238 | 1.936 | 0.187 | -0.399 | -0.008 | 0.511 | -3.701 | -1.121 | | | | | | | | | | | |
| B7L74_06285 | 4-hydroxy-tetrahydrodipicolinate synthase | | 0.149 | -0.567 | 1.140 | 1.348 | -0.723 | -0.861 | 0.615 | -0.988 | -0.270 | 1.186 | 1.495 | | | | | | | | | | | |
| B7L74_06290 | hypothetical protein | | 0.570 | 0.259 | -0.059 | 0.580 | -0.052 | 0.201 | -0.050 | -0.191 | -0.974 | 0.568 | 0.595 | | | | | | | | | | | |
| B7L74_06295 | 2-dehydro-3-deoxygluconokinase | | -9.058 | -8.300 | -5.739 | 1.210 | -2.034 | -2.966 | -3.637 | -7.651 | -6.519 | -9.476 | -4.650 | | | | | | | | | | | |
| B7L74_06300 | DNA recombination protein RmuC | DNA | 0.326 | -0.306 | -0.784 | -0.664 | 0.625 | 0.577 | -0.053 | -0.770 | -0.180 | -0.736 | -0.861 | | | | | | | | | | | |
| B7L74_06305 | hypothetical protein | | -0.625 | 0.389 | 2.523 | 1.500 | -0.353 | -0.495 | -0.294 | 0.112 | 0.237 | -0.106 | -0.123 | | | | | | | | | | | |
| B7L74_06310 | haloalkane dehalogenase | Chloroalkane and chloroalkene de | -0.928 | -1.020 | -0.887 | -1.075 | -2.105 | -2.165 | -0.895 | -1.864 | -2.170 | -0.642 | -0.497 | Early down | | | | | | | | | | |
| B7L74_06315 | glutamate dehydrogenase | | 3.211 | 3.863 | 2.889 | -2.466 | 4.281 | 4.440 | 1.391 | 2.566 | 2.574 | -4.995 | 1.057 | | | | | | | | | | | |
| B7L74_06320 | DNA-binding response regulator probably QseB(Pmr/ | Intracellular growthtwo componen | 0.104 | -0.293 | 1.244 | 0.542 | -0.246 | -0.243 | 0.406 | -0.886 | -0.367 | 0.022 | -0.221 | | | | | | | | | | | |
| B7L74_06325 | two-component sensor histidine kinase, QseC | two component system QseB-Qse | 0.033 | 0.836 | 0.966 | -1.812 | 2.632 | 2.387 | 0.440 | 1.485 | 2.540 | -1.284 | -0.913 | | | | | | | | | | | |
| B7L74_06330 | ATP-dependent helicase | | -0.194 | -0.770 | -0.105 | -0.328 | -0.434 | 0.369 | 0.038 | 0.862 | 0.805 | 0.025 | -0.078 | | | | | | | | | | | |
| B7L74_06335 | ATP-dependent helicase | | 0.395 | -0.659 | -2.519 | -2.384 | -1.059 | 0.554 | -0.480 | -0.376 | 0.335 | -1.174 | -0.075 | | | | | | | | | | | |
| B7L74_06340 | hypothetical membrane associated protein | | -1.031 | -0.428 | -1.294 | -0.146 | -0.721 | -0.925 | -0.312 | -0.234 | -0.578 | -0.492 | -0.125 | | | | | | | | | | | |
| B7L74_06345 | 5'-nucleotidase | | -0.220 | -0.208 | -0.588 | 0.682 | -0.083 | 0.239 | -0.448 | -0.717 | -0.687 | -1.251 | -1.648 | | | | | | | | | | | |
| B7L74_06350 | ATPase | ATP | -1.706 | -0.482 | -2.956 | -1.145 | -1.222 | -0.947 | -1.688 | -1.028 | -1.257 | -2.101 | -2.167 | Early down | | | | | | | | | | |
| B7L74_06355 | oligoribonuclease | Ribosome biogenesis in eukaryotes | 0.554 | 0.130 | 1.272 | 2.418 | 0.068 | 0.245 | 0.971 | 1.486 | 1.074 | 1.430 | 1.095 | Up in P12 | | | | | | | | | | |
| B7L74_06360 | hypothetical protein | | 3.024 | 2.180 | 1.564 | 1.982 | -0.692 | 0.953 | 1.785 | 2.942 | 3.392 | 1.710 | -0.198 | | | | | | | | | | | |
| B7L74_06365 | PIN domain nuclease | | -0.162 | -1.079 | 0.682 | 0.507 | -1.515 | -1.513 | 0.318 | 0.042 | 0.193 | 0.041 | 0.067 | | | | | | | | | | | |
| B7L74_06370 | CopG family transcriptional regulator | | -0.508 | -0.899 | 1.161 | 0.366 | -0.809 | -1.294 | 0.420 | 0.341 | 1.058 | 0.556 | 0.584 | | | | | | | | | | | |
| B7L74_06375 | acyl-phosphate glycerol 3-phosphate acyltransferase | | 0.542 | -0.121 | 1.810 | 1.041 | -0.459 | -0.252 | 0.909 | 0.111 | 1.520 | 1.106 | 0.884 | | | | | | | | | | | |
| B7L74_06380 | tRNA (adenosine(37)-N6)-threonylcarbamoyltransfera | | -0.073 | -0.628 | 0.387 | 0.955 | 0.348 | 0.541 | 0.685 | 1.631 | 1.300 | 0.680 | 0.723 | | | | | | | | | | | |
| B7L74_06385 | malate dehydrogenase | TCA cycle only | -0.224 | -1.132 | 0.345 | 2.540 | -0.416 | 0.425 | 0.849 | 0.130 | 0.435 | 0.715 | 1.350 | Up in P12 | | | | | Yes | | | | | |
| B7L74_06390 | long-chain fatty acid transporter | Transporter | 0.749 | 0.156 | 1.535 | 2.262 | -0.653 | 1.610 | 1.267 | 2.804 | 2.900 | 2.536 | 2.089 | Late up | | | | | | | | Yes (FA) | | |
| B7L74_06395 | exodeoxyribonuclease VII large subunit | | 0.162 | -1.119 | 0.782 | 0.551 | -0.042 | 2.337 | 0.595 | -0.034 | 0.181 | 1.579 | 0.333 | | | | | | | | | | | |
| B7L74_06400 | MFS transporter Multidrug resistance protein B | Resistance, Transporter | 6.686 | 0.299 | 2.171 | -1.132 | 7.412 | 5.379 | 3.584 | 3.861 | 6.840 | 1.173 | 1.979 | Early up | | | | | | | | Yes (MFS) | | |
| B7L74_06405 | ribosome biogenesis GTPase Der | Ribosome biogenesis | -1.071 | -1.829 | -1.636 | -1.183 | -2.047 | -1.330 | -0.914 | -1.376 | -1.495 | -1.034 | -1.075 | Early down | | | | | | | | | | |
| B7L74_06410 | hypothetical protein | | 0.090 | 0.848 | -0.035 | 1.139 | 3.899 | 3.832 | 0.527 | 1.496 | 2.628 | 1.002 | 0.095 | | | | | | | | | | | |
| B7L74_06415 | hypothetical protein | | -1.736 | -2.249 | 3.369 | -1.410 | -2.591 | -2.182 | -1.225 | -2.825 | -2.484 | -1.594 | -1.381 | Variable | | | | | | | | Yes (Outer membrane/Membrane) | | |
| B7L74_06420 | histidine--tRNA ligase | | -0.272 | -1.416 | 0.497 | 1.437 | -1.495 | -1.453 | 0.287 | -1.392 | -1.378 | 0.718 | 0.917 | | | | | | | | | | | |
| B7L74_06425 | transcriptional regulator | Membrane | 0.150 | -2.109 | 0.450 | 1.132 | -2.340 | -3.099 | 0.749 | -2.367 | -1.514 | 0.746 | 1.298 | | | | | | | | | | | |
| B7L74_06430 | hypothetical protein | | 0.808 | 0.573 | 1.199 | -0.336 | -0.158 | -0.373 | 1.312 | 1.053 | 0.070 | 1.567 | 1.294 | | | | | | | | | | | |
| B7L74_06435 | cell filamentation protein Fic | | -5.339 | -4.582 | -5.464 | 2.232 | 0.926 | 2.189 | -3.959 | -3.933 | 1.169 | 1.286 | -5.380 | | | | | | | | | | | |
| B7L74_06440 | bifunctional tRNA (adenosine(37)-C2)-methyltransfera | Ribosome biogenesis | -0.619 | -0.623 | -0.949 | -0.346 | -0.778 | -0.646 | -0.920 | -1.139 | -2.144 | -1.465 | -1.626 | Late down | | | | | | | | | | |
| B7L74_06445 | hypothetical protein | | -0.419 | -0.207 | -0.647 | 2.384 | -0.009 | -0.909 | -0.113 | 0.571 | -0.605 | -0.036 | -0.691 | Up in P12 | | | | | | | | | | |
| B7L74_06450 | hypothetical protein | Effector protein | 0.128 | -0.702 | 0.101 | 0.112 | -0.366 | 0.430 | 0.196 | 0.515 | 0.803 | 0.593 | 0.746 | | | Yes | | | | | | | | |
| B7L74_06455 | DUF1658 domain-containing protein | | 0.745 | 0.071 | 1.790 | 0.800 | 0.338 | 0.484 | 1.116 | -0.126 | 0.238 | 0.561 | 1.179 | | | | | | | | | | | |
| B7L74_06460 | polyprenyl synthetase | | -0.865 | -1.123 | 0.050 | 0.526 | -1.046 | -0.673 | -0.191 | -1.176 | -1.355 | -0.022 | -0.273 | | | | | | | | | | | |
| B7L74_06465 | hypothetical protein | | -0.883 | -1.573 | -1.828 | 1.182 | -1.607 | -0.915 | -0.747 | -0.689 | -3.273 | -0.559 | -0.495 | | | | | | | | | | | |
| B7L74_06470 | hypothetical cytosolic protein | | 2.043 | 2.234 | -0.111 | 2.533 | 3.333 | 0.241 | 0.953 | 2.384 | 2.819 | 0.863 | 1.439 | Early up | | | | | | | | | | |
| B7L74_06475 | nucleoside-diphosphate kinase | | 0.089 | 0.847 | -0.035 | 0.866 | 3.924 | 1.627 | 0.527 | 1.496 | 5.985 | 4.897 | -0.897 | | | | | | | | | | | |
| B7L74_06480 | Na+/H+ antiporter | Transporter | -0.523 | -1.061 | -0.458 | -0.045 | -1.023 | -1.093 | -0.486 | -0.724 | -0.990 | -0.070 | -0.186 | | | | | | | | | Yes (Na+:H+ antiporter ) | | |
| B7L74_06485 | hypothetical proteinUncharacterized protein CBU_126 | Invasin protein and Induce cellular | -1.044 | -1.170 | -0.406 | 1.847 | -1.166 | -0.316 | -0.140 | -0.018 | -0.005 | -0.094 | 0.652 | | | | | | | | | | | |
| B7L74_06490 | D-alanyl-D-alanine carboxypeptidase | | 0.103 | -1.873 | -1.050 | 0.014 | -0.861 | -0.071 | -0.528 | -1.706 | -1.212 | -0.621 | -0.709 | | | | | | | | | | | Yes |
| B7L74_06495 | DUF493 domain-containing protein | | 0.049 | 0.839 | -0.034 | 1.312 | 2.652 | 0.979 | 0.456 | 1.489 | 1.882 | -1.275 | 0.810 | | | | | | | | | | | |
| B7L74_06500 | hypothetical membrane associated protein | | 0.360 | -0.291 | 2.107 | 1.677 | 1.586 | 0.434 | 2.201 | -0.095 | 1.257 | 1.204 | 1.501 | | | | | | | | | | | |
| B7L74_06505 | hypothetical protein | | 0.834 | -0.753 | 1.394 | 1.416 | 0.604 | 2.507 | 1.871 | 3.024 | 3.126 | 1.731 | 1.584 | Early up | | | | | | | | | | |
| B7L74_06510 | octanoyltransferase | Lipoic acid metabolism | -0.647 | -2.261 | -0.616 | 0.478 | -2.054 | -1.723 | -0.656 | -1.255 | -1.611 | 0.051 | -0.052 | Early down | | | | | | | | | | |
| B7L74_06515 | lipoyl synthase | | -1.160 | -2.709 | -0.979 | -0.184 | -2.892 | 0.436 | -0.741 | -2.080 | -0.743 | -0.968 | -1.091 | | | | | | | | | | | |
| B7L74_06520 | Membrane-bound glucose-6-phosphatase | Glycolysis / Gluconeogenesis | -1.851 | -2.485 | -2.603 | -1.447 | -2.820 | -1.440 | -1.234 | -1.613 | -1.763 | -1.265 | -1.187 | Early down | | | | Yes | | | | | | |
| B7L74_06525 | protein ScvA involved in DNA binding | DNA binding | 3.024 | 2.947 | 3.549 | -2.376 | 2.414 | 2.347 | 3.495 | 2.241 | 3.083 | 2.383 | 3.912 | Early up | | | | | | | | | | |
| B7L74_06530 | pyridine nucleotide-disulfide oxidoreductase | Effector protein | -0.045 | -1.146 | 0.067 | 1.785 | -0.436 | -0.110 | 0.300 | 0.053 | -0.303 | 0.620 | 1.552 | | | Yes | | | | | | | | |
| B7L74_06535 | acyl-CoA thioesterase | Metabolism | -0.265 | -1.562 | 0.015 | 2.148 | -0.791 | -0.086 | 0.277 | -0.657 | -0.767 | 0.801 | 1.025 | Up in P12 | | | | | | | | | | |
| B7L74_06540 | hypothetical protein | | 0.453 | -0.521 | 1.763 | -0.243 | 0.134 | 0.351 | 0.500 | -1.205 | 0.092 | 1.359 | 1.483 | | | | | | | | | | | |
| B7L74_06545 | IS110 family transposase | | 0.910 | 0.438 | 1.542 | -0.418 | -0.871 | -0.528 | 0.238 | 0.008 | -1.420 | 1.094 | 1.635 | | | | | | | | | | | |
| B7L74_06550 | LanM family lanthionine synthetase | | 0.907 | 1.329 | 1.072 | -0.884 | 0.571 | 0.028 | 0.592 | 2.047 | 1.423 | -1.537 | -0.718 | | | | | | | | | | | |
| B7L74_06555 | hypothetical protein | | 1.004 | 2.069 | 2.658 | -1.227 | 1.293 | 0.905 | 2.032 | 3.170 | 2.062 | -1.289 | -0.489 | Late up | | | | | | | | | | |
| B7L74_06560 | 6-phosphofructokinase | | -0.051 | -0.400 | -0.682 | 0.425 | -1.144 | -1.067 | -0.453 | 0.291 | -0.337 | -0.727 | -0.750 | | | | | Yes | | | | | | |
| B7L74_06565 | hypothetical proteinInhibitor_I42 domaincontaining pr | Uncharacter ized | -0.467 | -0.182 | -1.060 | 0.048 | -1.235 | -0.690 | -0.538 | -0.436 | -0.788 | -0.577 | -0.432 | | | | | | | | | | | |
| B7L74_06570 | mandelate racemase | | -1.173 | -1.134 | -2.722 | -2.800 | 2.413 | -0.267 | 0.249 | -0.639 | -0.846 | -0.914 | -1.347 | | | | | | | | | | | |
| B7L74_06575 | short-chain dehydrogenase | | 0.629 | -0.152 | 0.248 | 1.897 | -0.208 | -0.092 | 0.800 | -0.319 | -0.777 | 1.030 | 1.894 | | | | | | | | | | | |
| B7L74_06580 | 2-dehydro-3-deoxyphosphooctonate aldolase | | 0.755 | -0.071 | -0.700 | 1.636 | -0.758 | 0.026 | 0.561 | 0.450 | 0.238 | 0.404 | 0.056 | | | | | | | | | | | |
| B7L74_06585 | thioredoxin family protein | | -0.601 | -0.185 | -1.192 | 0.219 | -0.823 | -0.092 | 0.159 | 0.735 | 0.385 | -0.600 | -0.762 | | | | | | | | | | | |
| B7L74_06590 | sugar transporter | Transporter | 0.122 | 0.484 | 0.533 | -2.791 | 5.205 | 4.562 | 1.714 | 1.448 | 3.139 | -0.740 | -0.548 | Early up | | | | | | | | | | |
| B7L74_06595 | DUF1658 domain-containing protein | | 1.270 | 1.073 | -0.107 | -0.936 | 1.242 | 0.546 | 0.552 | 0.916 | 0.348 | -0.853 | -0.836 | | | | | | | | | | | |
| B7L74_06600 | transcription elongation factor | | 0.311 | 1.366 | 0.961 | 1.356 | 0.325 | 0.039 | -0.264 | 1.469 | -0.897 | 0.469 | 0.203 | | | | | | | | | | | |
| B7L74_06605 | hypothetical protein | | 1.524 | 1.919 | 3.093 | 0.223 | 1.881 | 0.897 | 0.822 | 1.447 | 2.362 | 1.504 | 1.719 | | | | | | | | | | | |
| B7L74_06610 | carbamoyl phosphate synthase large subunit (gene=car | | 0.046 | -0.423 | -0.230 | -0.047 | 0.482 | 0.464 | -0.185 | -0.040 | 0.047 | -0.139 | 0.537 | | | | | | | | | | | |
| B7L74_06615 | carbamoyl phosphate synthase small subunit | | 0.176 | -0.137 | -1.430 | 1.091 | -1.288 | 0.760 | -0.117 | 0.603 | -0.097 | 0.052 | 0.469 | | | | | | | | | | | |
| B7L74_06620 | MFS transporter | Transporter | 0.414 | 0.480 | 0.789 | 0.731 | 0.716 | 1.600 | 1.461 | 2.120 | 2.423 | 1.090 | 1.074 | Late up | | | | | | | | Yes (drug resistance) | | |
| B7L74_06625 | ATPase | ATP | -0.258 | 0.138 | 0.694 | 0.481 | -0.781 | 0.354 | 0.325 | 2.061 | 1.861 | 0.035 | -0.183 | | | | | | | | | | | |
| B7L74_06630 | MFS transporter | Transporter | 1.577 | 1.012 | 1.959 | -1.048 | 2.556 | 2.685 | 0.566 | 0.046 | 2.382 | 0.040 | 0.579 | | | | | | | | | | | |
| B7L74_06635 | ATPase | ATP | -0.517 | -0.272 | -0.625 | -0.043 | -0.040 | 0.468 | 0.003 | 1.552 | 1.504 | 0.058 | -0.501 | | | | | | | | | | | |
| B7L74_06640 | hypothetical protein | | -0.158 | 0.487 | -0.643 | -0.030 | 0.692 | 1.064 | -0.309 | 1.419 | 2.180 | -0.364 | -1.076 | Late up | | | | | | | | | | |
| B7L74_06645 | molecular chaperone DnaJ | Chaperones and folding catalysts | -1.556 | -2.915 | -3.015 | -1.340 | -3.108 | -2.234 | -2.312 | -3.892 | -3.348 | -1.486 | -1.662 | Early down | | | Yes | | | | | | | |
| B7L74_06650 | molecular chaperone DnaK | RNA degradation | -3.083 | -4.247 | -3.344 | -2.126 | -3.909 | -3.549 | -2.997 | -4.717 | -4.369 | -2.544 | -2.191 | Early down | | | Yes | | | | | | | |
| B7L74_06655 | acetyl-CoA carboxylase biotin carboxyl carrier protein | subunit | -0.482 | -0.420 | -2.055 | -1.609 | -1.602 | -1.792 | -1.564 | -1.640 | -1.810 | -1.993 | -1.935 | Early down | | | | | | | | | | |
| B7L74_06660 | hypothetical proteinprotein=ankyrin repeat protein | Effector<br>AnkK. Eucaryotic-like ORFs. | -0.992 | -0.515 | -1.675 | -1.932 | -1.172 | -0.579 | -2.089 | -0.683 | -0.893 | -1.168 | -1.410 | Early down | | Yes | | | | | | | | |
| B7L74_06665 | nucleotide exchange factor GrpE | Protein Folding | -1.095 | -0.689 | -0.776 | 0.660 | -1.806 | -1.533 | -1.199 | -1.098 | -1.562 | -0.871 | -0.126 | | | | Yes | | | | | | | |
| B7L74_06670 | PIN domain nuclease | | 0.051 | 0.231 | 1.158 | -0.277 | -0.499 | -0.513 | -0.295 | 0.324 | -0.409 | 0.720 | 0.634 | | | | | | | | | | | |
| B7L74_06675 | hypothetical membrane associated protein | | 2.857 | 3.728 | 4.476 | 0.892 | 3.179 | 3.153 | 2.331 | 2.130 | 1.475 | 0.667 | -0.383 | | | | | | | | | | | |
| B7L74_06680 | NAD kinase | | -0.068 | 0.248 | 0.136 | -0.693 | -0.412 | -0.543 | -0.406 | -0.301 | -1.525 | -0.345 | -0.343 | | | | | | | | | | | |
| B7L74_06685 | DNA repair protein RecN | DNA repair and recombination pro | -1.537 | -0.779 | -2.515 | -2.636 | -0.389 | -0.503 | -2.074 | -1.628 | -1.194 | -1.987 | -2.417 | Early down | | | | | | | | | | |
| B7L74_06690 | hypothetical protein | | -0.482 | -0.009 | -0.130 | 0.201 | 0.421 | 0.551 | 0.214 | 1.029 | 0.901 | -0.398 | -1.030 | | | | | | | | | | | |
| B7L74_06695 | hypothetical protein | Hypothetical | -1.452 | 0.926 | -1.035 | -0.733 | 1.546 | 3.867 | -0.872 | 0.501 | 1.478 | -3.273 | -2.550 | Up in P21 | | | | | | | | | | |
| B7L74_06700 | hypothetical exported protein | | 1.052 | 0.312 | 1.703 | -0.698 | -0.519 | -0.601 | 0.767 | 0.112 | -0.715 | -0.080 | 0.501 | | | | | | | | | | | |
| B7L74_06705 | transcriptional repressor | | -0.040 | 0.268 | -0.829 | -0.054 | -0.808 | -0.773 | -1.019 | 0.060 | -1.720 | -0.771 | -0.443 | | | | | | | | | | | |
| B7L74_06710 | outer membrane protein assembly factor BamE | Membrane | -0.612 | -0.471 | -0.616 | 0.459 | -1.497 | -1.262 | 0.482 | 0.119 | -0.814 | -0.786 | -0.757 | | | | | | | | | Yes (Outer membrane/Lipoprotein) | | |
| B7L74_06715 | RnfH family protein | | 2.202 | 0.847 | -0.035 | -0.096 | 3.925 | 1.627 | 0.527 | 1.496 | 2.628 | -0.206 | -0.897 | | | | | | | | | | | |
| B7L74_06720 | ubiquinone-binding protein | | 2.043 | 0.848 | -0.035 | -0.083 | 5.004 | 3.605 | 1.508 | 1.496 | 2.628 | 0.917 | 0.090 | | | | | | | | | | | |
| B7L74_06725 | SsrA-binding protein | Protein processing | -1.132 | -0.872 | -1.676 | -0.881 | -2.622 | -2.149 | -1.127 | -0.423 | -1.160 | -0.755 | -0.750 | Early down | | | | | | | | | | |
| B7L74_06730 | methionine sulfoxide reductase MsrA | Stress Response | 0.068 | 0.843 | -0.034 | -1.781 | 4.295 | 5.378 | 3.626 | 4.314 | 4.264 | -1.264 | -0.458 | Early up | | | | | | | | | | |
| B7L74_06735 | hypothetical membrane associated protein | | 0.536 | 0.347 | -6.047 | -0.882 | -3.275 | 1.258 | -0.510 | 1.721 | 1.322 | -1.264 | -1.240 | | | | | | | | | | | |
| B7L74_06740 | phosphohydrolase | | -2.473 | -1.054 | -3.497 | -0.384 | -1.436 | -1.864 | -2.008 | -0.771 | -1.530 | -2.319 | -2.483 | Early down | | | | | | | | | | |
| B7L74_06745 | transfer-messenger RNA (gene=ssrA) | tRNA | 0.599 | 1.594 | 2.614 | 0.301 | 1.538 | 1.357 | 2.963 | 3.043 | 3.164 | 0.276 | 0.775 | Late up | | | | | | | | | | |
| B7L74_06750 | hypothetical protein | | 2.483 | 0.869 | 2.574 | 1.729 | 3.183 | 3.534 | 1.331 | 3.857 | 3.889 | 2.179 | 1.772 | Early up | | | | | | | | | | |
| B7L74_06755 | hypothetical protein | | 2.306 | 1.293 | -0.812 | 0.127 | 2.565 | 3.610 | 1.396 | 3.958 | 4.375 | -0.377 | -0.845 | Early up | | | | | | | | | | |
| B7L74_06760 | hypothetical cytosolic protein | | 3.853 | 1.491 | 3.058 | 0.941 | 5.431 | 4.851 | 3.676 | 5.035 | 5.654 | 2.331 | 1.612 | Continuous up | | | | | | | | | | |
| B7L74_06765 | hypothetical protein | | 0.248 | 1.712 | -0.609 | 1.924 | 1.490 | -0.428 | -0.785 | 5.128 | 4.553 | 0.415 | -0.799 | Late up | | | | | | | | | | |
| B7L74_06770 | hypothetical protein | | -0.476 | 0.552 | 0.380 | 1.974 | -1.291 | 1.609 | -3.501 | 3.987 | 2.746 | -0.255 | -0.197 | | | | | | | | | | | |
| B7L74_06775 | hypothetical cytosolic protein | Effector protein | 4.805 | 1.474 | 2.419 | 1.319 | 5.652 | 5.219 | 3.460 | 3.148 | 5.182 | 2.978 | 3.087 | Early up | | Yes | | | | | | | | |
| B7L74_06780 | hypothetical protein | Glycolysis / Gluconeogenesis | 0.741 | 0.589 | -3.756 | -0.469 | -0.081 | 1.890 | 1.309 | 4.538 | 4.736 | -0.972 | 0.073 | Late up | | | | Yes | | | | | | |
| B7L74_06785 | hypothetical cytosolic protein | | -0.798 | 0.619 | -1.219 | -3.068 | 0.719 | 1.783 | 0.432 | 3.195 | 3.578 | -1.654 | -0.307 | | | | | | | | | | | |
| B7L74_06790 | hypothetical protein | | 0.318 | 0.305 | 1.262 | -0.127 | 1.061 | 2.918 | -0.073 | 3.919 | 4.077 | 0.437 | 1.738 | Late up | | | | | | | | | | |
| B7L74_06795 | hypothetical protein | | 0.843 | 1.067 | 0.416 | 0.270 | 1.291 | 2.303 | 0.895 | 4.796 | 5.575 | -0.595 | 1.078 | Late up | | | | | | | | | | |
| B7L74_06800 | hypothetical cytosolic protein | | -0.584 | -0.574 | -0.478 | 1.604 | -1.435 | 0.231 | 0.643 | 2.926 | 2.356 | 1.730 | 0.699 | | | | | | | | | | | |
| B7L74_06805 | tRNA-Pro | tRNA | 0.115 | -1.549 | 0.575 | 0.285 | -1.351 | -4.206 | 0.923 | 0.150 | 0.863 | 0.501 | 0.742 | | | | | | | | | | | |
| B7L74_06810 | transcriptional regulator | | 0.375 | -0.717 | 0.574 | 1.463 | -0.334 | -1.070 | 0.479 | -0.162 | -0.393 | 1.071 | 0.968 | | | | | | | | | | | |
| B7L74_06815 | integration host factor subunit alpha | DNA replication proteins | 1.059 | -1.003 | 2.214 | 2.217 | -0.185 | -0.749 | 1.374 | -0.792 | -0.667 | 1.707 | 2.115 | Early up | | | | | | | | | | |
| B7L74_06820 | phenylalanine--tRNA ligase subunit beta | Aminoacyl-tRNA synthase | -1.304 | -1.827 | -0.972 | -1.040 | -2.353 | -2.000 | -1.223 | -1.675 | -1.496 | -0.934 | -0.679 | Early down | | | | | | | | | | |
| B7L74_06825 | phenylalanine--tRNA ligase subunit alpha | | 0.031 | 0.836 | -0.034 | -1.814 | 2.629 | 2.012 | 1.004 | 2.724 | 3.066 | -0.925 | -0.913 | | | | | | | | | | | |
| B7L74_06830 | 50S ribosomal protein L20 | Ribosomal proteins | -1.630 | -2.287 | -1.794 | -1.095 | -2.111 | -2.444 | -0.996 | -3.065 | -3.456 | -1.483 | -1.090 | Continuous down | | | | | | | | | | |
| B7L74_06835 | DUF1658 domain-containing protein | CB specific protein family | -0.273 | -0.577 | -0.722 | -1.050 | -0.960 | -0.731 | -0.836 | -1.360 | -3.182 | -2.151 | -1.285 | Late down | | | | | | | | | | |
| B7L74_06840 | DUF1658 domain-containing protein | CB specific protein family | -0.005 | -1.010 | 0.117 | -0.501 | -1.021 | -0.987 | -0.097 | -1.848 | -2.934 | -1.298 | -0.090 | Late down | | | | | | | | | | |
| B7L74_06845 | 50S ribosomal protein L35 | Ribosomal proteins | -1.371 | -1.759 | -0.911 | -0.514 | -2.625 | -1.844 | -0.957 | -2.236 | -1.899 | -1.468 | -0.692 | Early down | | | | | | | | | | |
| B7L74_06850 | translation initiation factor IF-3 | Translation factors | 2.704 | 0.837 | 0.966 | -1.810 | 4.137 | 2.673 | 1.837 | 2.153 | 3.857 | -0.310 | 0.107 | Early up | | | | | | | | | | |
| B7L74_06855 | threonine--tRNA ligase | Aminoacyl-tRNA synthase | -0.358 | -1.197 | -2.817 | 0.260 | -1.694 | -0.735 | -0.882 | -0.870 | -1.338 | -0.995 | -0.297 | Early down | | | | | | | | | | |

| | | | | | | | | | | | | | | | | | | | | | | | | |
|---|---|---|---|---|---|---|---|---|---|---|---|---|---|---|---|---|---|---|---|---|---|---|---|---|
| B7L74_06860 | hypothetical cytosolic protein | | -0.921 | -1.045 | 0.204 | 2.394 | 0.312 | -3.422 | 0.285 | 1.365 | 2.348 | 1.237 | 1.109 | | | | | | | | | | | |
| B7L74_06865 | radical SAM protein | | -1.277 | -0.681 | -0.118 | -0.260 | -0.068 | 0.777 | 0.925 | 0.837 | 2.083 | -0.376 | -0.272 | | | | | | | | | | | |
| B7L74_06870 | polyamine oxidase | | 0.817 | 0.343 | 1.918 | 0.463 | 0.305 | 0.741 | 1.242 | 1.302 | 1.602 | 1.698 | 1.193 | | | | | | | | | | | |
| B7L74_06875 | CBS domain-containing protein | | 2.675 | 1.164 | 2.258 | -0.160 | 0.862 | -0.842 | 3.029 | 3.200 | 2.863 | 1.555 | 2.775 | Late up | | | | | | | | | | |
| B7L74_06880 | anion permease | | 1.081 | -0.303 | 0.731 | 0.431 | -0.236 | 0.373 | 1.326 | 1.364 | 1.595 | 1.575 | 1.284 | | | | | | | | | | | |
| B7L74_06885 | FUSC family protein, Hypothetical membrane spannin | Membrane | -5.653 | 0.549 | 1.605 | -1.463 | -0.172 | 0.412 | -0.922 | -0.457 | -2.049 | -0.682 | -5.004 | | | | | | | | | | | |
| B7L74_06890 | hypothetical protein | | 0.248 | 0.783 | 0.645 | 0.414 | 1.706 | 2.170 | 0.769 | 2.034 | 1.996 | 0.145 | 0.356 | | | | | | | | | | | |
| B7L74_06895 | cyclin | | 1.975 | 0.090 | 1.895 | 1.921 | 2.798 | 2.908 | 2.098 | 0.554 | 2.110 | 2.040 | 3.113 | Early up | | | | | | | | | | |
| B7L74_06900 | EamA family transporter | Transporter | -0.038 | -0.584 | 0.976 | -2.220 | 3.802 | 1.157 | -0.905 | 0.065 | 4.580 | -0.952 | -0.564 | | | | | | | | | | | |
| B7L74_06905 | DNA polymerase III subunit alpha | DNA replication | -1.093 | -1.553 | -2.939 | -1.259 | -1.322 | -0.874 | -1.367 | -1.051 | -1.043 | -1.596 | -1.576 | Early down | | | | | | | | | | |
| B7L74_06910 | D-alanine--D-alanine ligase A | | -0.727 | -0.599 | -0.631 | -0.149 | -1.871 | -0.567 | -0.306 | -0.847 | -1.012 | -0.098 | 0.028 | | | | | | | | | | | Yes |
| B7L74_06915 | ribonuclease HII | | -1.008 | -0.894 | -1.209 | -0.920 | -1.667 | -0.379 | -2.547 | -0.232 | -0.354 | -1.606 | -1.198 | | | | | | | | | | | |
| B7L74_06920 | hypothetical protein | | -1.801 | -2.528 | -3.206 | -1.156 | -3.663 | -1.218 | -3.497 | -1.496 | -1.669 | -2.054 | -1.213 | Early down | | | | | | | | | | |
| B7L74_06925 | GMP synthase (glutamine-hydrolyzing) | | 0.666 | 1.094 | -0.274 | -2.669 | 1.840 | 0.971 | 2.211 | 1.545 | 2.349 | 0.125 | -0.426 | | | | | | | | | | | |
| B7L74_06930 | IMP dehydrogenase | | -1.058 | -1.583 | -2.319 | 0.276 | -2.545 | -1.410 | -1.788 | -1.161 | -1.313 | -1.186 | -0.237 | | | | | | | | | | | |
| B7L74_06935 | hypothetical protein | | 5.502 | 2.214 | 4.139 | 1.032 | 7.234 | 8.155 | 4.027 | 2.121 | 6.618 | 4.357 | 1.589 | Continuous up | | | | | | | | | | |
| B7L74_06940 | MFS transporter | Transporter | 1.335 | -1.011 | 2.009 | -0.118 | 1.986 | 1.731 | 2.982 | 1.204 | 2.947 | 1.634 | 2.389 | Late up | | | | | | | | Yes (Sugar/glucose/mannose:H+ symporter) | | |
| B7L74_06945 | trehalase | | -0.982 | -1.297 | -0.818 | 1.411 | -1.163 | -1.383 | -0.571 | -1.291 | -1.343 | -1.432 | -1.202 | | | | | | | | | | | |
| B7L74_06950 | amino acid permease, Glutamate/gammaaminobutyrate | Transporter | 0.090 | 0.848 | 2.408 | 3.903 | 2.737 | 3.398 | 1.590 | 1.496 | 6.565 | 0.461 | 0.851 | | | | | | | | | | | |
| B7L74_06955 | hypothetical protein | | 5.032 | 2.636 | 4.313 | -0.691 | 7.796 | 6.829 | 3.733 | 1.976 | 7.160 | 4.325 | 1.318 | Early up | | | | | | | | | | |
| B7L74_06960 | hypothetical cytosolic protein | Effector protein | 2.625 | -0.199 | 0.629 | 0.689 | 4.353 | 3.899 | 0.567 | 0.274 | 4.129 | 1.069 | -0.912 | Early up | | Yes | | | | | | | | |
| B7L74_06965 | phosphoglucosamine mutase | | -1.599 | -1.591 | -1.315 | 0.547 | -2.103 | -1.803 | -1.404 | -2.321 | -1.386 | -1.250 | -2.169 | | | | | | | | | | | |
| B7L74_06970 | dihydropteroate synthase | | -0.182 | -1.366 | -1.660 | -0.895 | -2.190 | -2.110 | -0.454 | -2.499 | -4.082 | -0.300 | -1.621 | | | | | | | | | | | |
| B7L74_06975 | ATP-dependent metalloprotease (gene=hflB) | Cell division | -0.637 | -1.150 | -0.561 | -1.153 | -1.034 | -0.882 | -0.792 | -1.834 | -2.000 | -0.770 | -1.139 | Late down | | | | | | | | | | |
| B7L74_06980 | ribosomal RNA large subunit methyltransferase E | Ribosome biogenesis | -1.534 | -1.544 | -1.999 | 0.296 | -2.547 | -1.869 | -1.039 | -2.353 | -2.946 | -0.301 | -0.160 | Early down | | | | | | | | | | |
| B7L74_06985 | putative Fe-S cluster assembly protein SufT | | -0.675 | -0.160 | -1.507 | -0.611 | -1.697 | -1.353 | -0.780 | -1.694 | -2.446 | -0.400 | -0.100 | Early down | | | | | | | | | | |
| B7L74_06990 | Fe-S cluster assembly scaffold SufA | Transfer RNA biogenesis | -1.048 | -0.516 | -2.641 | -1.137 | -2.246 | -1.417 | -1.225 | -2.179 | -2.882 | -1.363 | -0.874 | Early down | | | | | | | | | | |
| B7L74_06995 | iron-sulfur cluster assembly scaffold protein | Energy metabolism | -1.661 | -0.395 | -2.900 | -1.244 | -2.998 | -1.393 | -1.429 | -1.502 | -2.599 | -1.584 | -1.360 | Early down | | | | | | | | | | |
| B7L74_07000 | cysteine sulfinate desulfinase | Selenocompound metabolism | -1.315 | -0.469 | -2.750 | -0.932 | -3.361 | -0.915 | -1.295 | -1.432 | -1.884 | -2.166 | -2.028 | Early down | | | | | | | | | | |
| B7L74_07005 | Fe-S cluster assembly protein SufD | | -0.888 | -0.372 | -1.602 | 0.046 | -2.268 | -1.010 | -1.633 | -0.976 | -2.692 | -1.306 | -1.681 | Early down | | | | | | | | | | |
| B7L74_07010 | ABC transporter ATP-binding protein | Transporter | -0.026 | 0.265 | 0.975 | 1.041 | 2.083 | 2.117 | -0.114 | 1.285 | 1.986 | 0.567 | -1.482 | | | | | | | | | Yes (General) | | |
| B7L74_07015 | Fe-S cluster assembly protein SufB | | -1.371 | -0.126 | -0.364 | 0.522 | -2.797 | -1.123 | -0.741 | -0.830 | -1.824 | -0.111 | -0.590 | Early down | | | | | | | | | | |
| B7L74_07020 | transcriptional regulatorRrf2 family protein | Transcription factors | 2.088 | 0.923 | 4.865 | 3.347 | -0.241 | -0.304 | 1.781 | 1.861 | 0.838 | 4.313 | 4.125 | Early up | | | | | | | | | | |
| B7L74_07025 | cation diffusion facilitator family transporter | | 0.398 | 0.460 | -0.291 | 1.581 | 0.157 | 1.699 | 0.097 | 0.621 | 0.369 | -0.213 | -0.259 | | | | | | | | | | | |
| B7L74_07030 | exodeoxyribonuclease I | | 2.559 | 0.029 | 0.657 | -3.853 | 3.470 | 1.984 | 0.929 | -0.930 | 2.443 | 0.412 | -1.200 | | | | | | | | | | | |
| B7L74_07035 | hypothetical protein | | 0.043 | 0.081 | -1.302 | -0.993 | -0.402 | 0.989 | -0.337 | 1.222 | 0.975 | -1.274 | -0.847 | | | | | | | | | | | |
| B7L74_07040 | hypothetical exported protein | | 0.136 | -0.516 | -2.079 | -2.858 | -1.533 | -1.698 | -2.458 | -1.982 | -3.773 | -2.690 | -2.271 | Early down | | | | | | | | | | |
| B7L74_07045 | hypothetical protein | | 0.786 | 0.636 | 0.449 | 0.892 | 0.106 | 1.141 | 0.816 | 1.553 | 1.394 | 0.917 | 0.782 | | | | | | | | | | | |
| B7L74_07050 | hypothetical membrane associated protein | | 4.543 | 1.283 | 2.211 | 0.783 | 5.781 | 5.298 | 3.010 | 2.084 | 4.818 | 2.269 | 1.827 | Early up | | | | | | | | | | |
| B7L74_07055 | hypothetical membrane associated protein | Effector protein | 1.153 | 0.811 | -0.543 | -0.576 | 3.499 | 2.681 | 0.083 | 0.749 | 2.351 | 0.801 | -0.627 | Up in P16 | | Yes | | | | | | | | |
| B7L74_07060 | tRNA (adenosine(37)-N6)-threonylcarbamoyltransfera | | 0.820 | 0.170 | 0.910 | 1.389 | -0.052 | 0.338 | 0.909 | 0.282 | 0.217 | 0.908 | 1.115 | | | | | | | | | | | |
| B7L74_07065 | hypothetical protein | | 0.098 | 0.760 | -0.026 | 0.060 | 0.333 | 0.232 | -0.207 | 0.588 | -0.104 | -0.022 | -0.139 | | | | | | | | | | | |
| B7L74_07070 | hypothetical membrane associated protein | | 0.544 | 0.581 | 1.846 | 1.573 | 0.304 | 0.996 | 1.268 | 0.653 | 0.795 | 1.323 | 2.187 | Early up | | | | | | | | | | |
| B7L74_07075 | para-aminobenzoate synthetase component I | Folate biosynthesis | -1.640 | -1.374 | -2.686 | -0.852 | -2.820 | -0.696 | -1.552 | -0.438 | -1.172 | -1.862 | -1.687 | Early down | | | | | | | | | | |
| B7L74_07080 | tRNA-specific adenosine deaminase | | 0.712 | 0.319 | -2.739 | -3.072 | 0.226 | 0.052 | -0.769 | 0.140 | -0.330 | -2.097 | -3.255 | | | | | | | | | | | |
| B7L74_07085 | GTP pyrophosphokinase, relA | Signal transduction | -0.600 | -1.099 | 0.113 | 0.712 | -0.238 | 0.620 | 0.407 | 0.385 | 0.412 | 0.385 | 0.599 | | | | | | | | | | | |
| B7L74_07090 | zinc metalloprotease(coxK2 in Table 1 of RightonQ pa | Effector protein | -0.179 | -0.219 | 1.964 | 0.574 | 0.252 | 0.780 | 2.048 | 2.314 | 2.195 | 1.528 | 1.660 | | | Yes | | | | | | | | |
| B7L74_07095 | protein kinase | Endocytosis | 2.757 | -0.369 | 0.721 | 1.339 | 3.837 | 3.832 | 1.577 | 0.555 | 2.437 | 1.793 | 1.315 | Early up | | | | | | | | | | |
| B7L74_07100 | hypothetical proteinOmpA_membrane domain-contain | Membrane | -0.388 | -0.418 | -1.055 | 0.656 | -1.076 | -0.943 | -0.215 | -1.154 | -2.002 | -0.278 | 0.267 | | | | | | | | | | | |
| B7L74_07105 | quinol oxidase | Membrane | 1.075 | -0.249 | -0.584 | -0.408 | 2.938 | 2.869 | -0.173 | -1.217 | 2.869 | 0.579 | -0.570 | | | | | | | | | | | |
| B7L74_07110 | phosphatidate cytidylyltransferase | | 0.780 | -0.986 | -0.285 | 0.265 | 0.648 | 1.718 | 1.199 | -1.094 | 0.521 | 0.317 | 0.484 | | | | | | | | | | | |
| B7L74_07115 | ditrans%2Cpolycis-undecaprenyl-diphosphate synthase | Peptidoglycan syn | -1.869 | -1.859 | -3.166 | -0.391 | -2.616 | -2.818 | -1.807 | -2.708 | -3.746 | -1.354 | 0.335 | Early down | | | | | | | | | | Yes |
| B7L74_07120 | ribosome-recycling factor | | 1.414 | 0.822 | 0.235 | 0.524 | 1.523 | 1.621 | 1.652 | 1.551 | 1.250 | 1.962 | 1.926 | | | | | | | | | | | |
| B7L74_07125 | UMP kinase | Pyrimidine metabolism | -1.400 | -1.598 | -2.041 | 0.595 | -1.826 | -1.529 | -1.035 | -1.981 | -2.696 | -1.210 | -0.773 | Continuous down | | | | | | | | | | |
| B7L74_07130 | elongation factor Ts | | 1.774 | 0.766 | -0.034 | -1.770 | 3.983 | 1.841 | 0.825 | 1.748 | 2.469 | -1.473 | -0.899 | | | | | | | | | | | |
| B7L74_07135 | 30S ribosomal protein S2 | Ribosomal proteins | -0.637 | -0.985 | -0.400 | 0.943 | -1.533 | -1.054 | -0.112 | -0.830 | -1.826 | 0.256 | 0.736 | | | | | | | | | | | |
| B7L74_07140 | hypothetical cytosolic protein | Effector protein | 0.051 | -0.037 | 1.456 | 0.290 | -0.282 | -0.193 | 1.630 | -0.145 | 0.242 | 0.354 | 0.864 | | | Yes | | | | | | | | |
| B7L74_07145 | type I methionyl aminopeptidase | Peptidases and inhibitors | 0.483 | -0.186 | 1.020 | 2.196 | -0.498 | 0.622 | 0.720 | 0.922 | 0.964 | 1.597 | 1.806 | Early up | | | | | | | | | | |
| B7L74_07150 | pyridoxal-dependent decarboxylase | | 3.208 | 3.622 | -0.069 | -1.752 | 3.939 | 3.469 | 1.679 | 2.707 | 4.173 | -0.551 | -2.421 | | | | | | | | | | | |
| B7L74_07155 | phosphoadenosine phosphosulfate reductase | | 0.527 | 1.829 | -0.165 | -1.381 | 0.645 | 0.731 | 0.612 | 1.918 | 1.853 | 0.832 | 2.082 | | | | | | | | | | | |
| B7L74_07160 | hypothetical protein | | 1.748 | 1.828 | 2.345 | 1.286 | 0.280 | -0.273 | 2.112 | 2.638 | 1.471 | 1.776 | 3.277 | | | | | | | | | | | |
| B7L74_07165 | phosphosulfolactate synthase | | 0.285 | 0.449 | 1.033 | 0.705 | 2.362 | 2.132 | 2.359 | 2.510 | 3.903 | 1.114 | 1.733 | | | | | | | | | | | |
| B7L74_07170 | 2-phosphosulfolactate phosphatase | Methane metabolism | 1.439 | -0.116 | 0.374 | -1.023 | 5.109 | 4.473 | 4.234 | 3.874 | 4.809 | 0.561 | 1.603 | Early up | | | | | | | | | | |
| B7L74_07175 | non-ribosomal peptide synthetase | | 2.198 | 2.607 | 2.681 | -0.869 | 1.256 | 4.202 | 2.000 | 3.757 | 3.959 | 2.142 | 1.634 | | | | | | | | | | | |
| B7L74_07180 | hypothetical protein | | 0.083 | 0.847 | -0.034 | -2.484 | 1.206 | 1.059 | -0.930 | 0.897 | 2.593 | -1.256 | -0.897 | | | | | | | | | | | |
| B7L74_07185 | peptide synthetase | | 1.829 | 0.915 | 0.502 | -2.433 | 0.883 | 0.338 | 0.598 | 0.760 | 1.737 | 1.255 | 1.372 | | | | | | | | | | | |
| B7L74_07190 | L%2CD-transpeptidase, Enhanced entry protein | Entry | 1.921 | 2.175 | 3.193 | -1.739 | 3.396 | 2.193 | 4.097 | 2.261 | 5.731 | 1.358 | 1.542 | Early up | | | | | | | | | | |
| B7L74_07195 | hypothetical membrane spanning protein | | -0.544 | 0.520 | -0.717 | -1.093 | -0.172 | -0.431 | -0.739 | -0.201 | 0.728 | -0.609 | -0.999 | | | | | | | | | | | |
| B7L74_07200 | succinyl-CoA ligase subunit alpha | TCA cycle only | -0.749 | -1.703 | -3.536 | -1.976 | -1.830 | -2.220 | -1.672 | -3.938 | -3.292 | -1.554 | -0.723 | Early down | | | | | Yes | | | | | |
| B7L74_07205 | succinyl-CoA ligase subunit beta | Effector protein | 0.443 | -1.435 | -1.967 | -1.622 | -0.206 | -0.096 | -0.648 | -2.257 | -1.649 | 0.064 | 0.980 | | | Yes | | | Yes | | | | | |
| B7L74_07210 | dihydrolipoamide succinyltransferase | TCA cycle only | 0.801 | -1.763 | -2.037 | -0.852 | 0.549 | 0.787 | -0.884 | -3.962 | -0.722 | -0.759 | -0.678 | Down in P42 | | | | | Yes | | | | | |
| B7L74_07215 | 2-oxoglutarate dehydrogenase E1 component | TCA cycle only | -2.188 | -1.973 | -4.415 | -2.891 | -2.830 | -2.042 | -3.242 | -3.741 | -3.898 | -2.717 | -1.697 | Continuous down | | | | | Yes | | | | | |
| B7L74_07220 | succinate dehydrogenase iron-sulfur subunit (gene=sdh | Respiration/succinate | -2.400 | -1.516 | -4.392 | -1.900 | -2.715 | -2.928 | -3.054 | -2.539 | -3.706 | -2.751 | -2.725 | Continuous down | | | | | Yes | | | | | |
| B7L74_07225 | succinate dehydrogenase flavoprotein subunit | Respiration/succinate | -0.901 | -1.058 | -3.174 | -1.701 | -2.315 | -2.214 | -2.210 | -2.562 | -3.398 | -2.473 | -2.157 | Early down | | | | | Yes | | | | | |
| B7L74_07230 | succinate dehydrogenase hydrophobic membrane anch | Respiration/succinate | -1.252 | -0.508 | -2.021 | -1.299 | -1.250 | -1.237 | -1.669 | -0.995 | -1.693 | -1.324 | -1.649 | Early down | | | | | Yes | | | | | |
| B7L74_07235 | succinate dehydrogenase cytochrome b556 subunit | Respiration/succinate | 0.027 | 0.835 | -0.034 | -0.887 | 2.625 | 1.619 | 0.435 | 1.484 | 2.535 | -0.926 | -0.178 | | | | | | Yes | | | | | |
| B7L74_07240 | hypothetical exported protein | | 0.040 | 1.277 | 4.771 | -0.052 | 1.219 | 0.768 | 2.500 | 2.139 | 0.521 | 0.174 | -0.090 | Early up | | | | | | | | | | |
| B7L74_07245 | hypothetical protein | Pseudogenes | -0.327 | 1.196 | 1.967 | -0.713 | -0.119 | 0.477 | 0.156 | 1.076 | 0.278 | 1.014 | 0.391 | | | | | | | | | | | |
| B7L74_07250 | hypothetical protein | | 2.974 | -0.097 | 1.076 | 0.429 | 4.357 | 4.107 | 1.715 | -0.284 | 2.453 | 1.914 | 1.569 | Early up | | | | | | | | | | |
| B7L74_07255 | hypothetical protein | Effector protein | 4.398 | 0.759 | 2.200 | -0.586 | 5.768 | 5.539 | 3.115 | 0.268 | 3.749 | 3.454 | 2.870 | Early up | | Yes | | | | | | | | |
| B7L74_07260 | type II citrate synthase | | -0.555 | -0.165 | 0.539 | -1.495 | -0.289 | -0.132 | -0.508 | -0.383 | -0.255 | -0.871 | -1.280 | | | | | | Yes | | | | | |
| B7L74_07265 | hypothetical protein | | 0.561 | 0.120 | 0.944 | -0.036 | -0.074 | -0.081 | 0.939 | -0.226 | -1.090 | 0.397 | 0.584 | | | | | | | | | | | |
| B7L74_07270 | hypothetical exported protein | | 0.878 | -0.539 | 0.646 | 1.660 | 0.013 | 0.033 | 1.717 | 0.250 | -0.450 | 1.592 | 1.685 | | | | | | | | | Yes (General/Hypothetical) | | |
| B7L74_07275 | hypothetical exported membrane associated protein | | 0.530 | -0.608 | 0.605 | 1.999 | -0.598 | 0.488 | 1.886 | 1.520 | 1.356 | 2.155 | 1.541 | | | | | | | | | Yes (General/Hypothetical) | | |
| B7L74_07280 | hypothetical exported protein | | 2.730 | 0.408 | 2.043 | 3.612 | 3.458 | 4.438 | 2.843 | 3.108 | 3.871 | 3.741 | 2.703 | Early up | | | | | | | | Yes (General/Hypothetical) | | |
| B7L74_07285 | thiamine-phosphate kinase | | 0.115 | -0.379 | -0.650 | -0.139 | 0.376 | 0.855 | -0.162 | -0.210 | -0.702 | -1.038 | -1.190 | | | | | | | | | | | |
| B7L74_07290 | transcriptional regulator | Transcription factors | 0.550 | 0.289 | 1.667 | 2.321 | -0.229 | -0.641 | 1.059 | 0.820 | 0.126 | 2.384 | 1.963 | Early up | | | | | | | | | | |
| B7L74_07295 | N utilization substance protein B (nusB) | DNA, Transcription machinery | -0.208 | -0.934 | -1.048 | 1.630 | -0.599 | -0.599 | 1.018 | 0.135 | 1.106 | 1.359 | 2.836 | Up in P67 | | | | | | | | | | |
| B7L74_07300 | transcriptional regulator NrdR | | 0.575 | 0.240 | 0.586 | 1.795 | -0.257 | 0.631 | 0.373 | 0.364 | 0.566 | 0.758 | 1.349 | | | | | | | | | | | |
| B7L74_07305 | serine hydroxymethyltransferase (gene=glyA) | Glyoxylate and dicarboxylate meta | 0.227 | 0.133 | 0.057 | 2.796 | -0.042 | -0.525 | 0.805 | 0.803 | 0.108 | 1.305 | 2.358 | Early up | | | | | | | | | | |
| B7L74_07310 | hypothetical protein | | 0.210 | 1.194 | 1.870 | -0.161 | 1.151 | 0.655 | 1.883 | 1.980 | 1.651 | 2.448 | 2.106 | Late up | | | | | | | | | | |
| B7L74_07315 | DNA repair protein RadA | DNA | 0.258 | -0.622 | -0.486 | 1.148 | -1.580 | -0.566 | 0.353 | -0.303 | -0.878 | 1.269 | 1.062 | | | | | | | | | | | |
| B7L74_07320 | hypothetical protein | | -0.914 | -0.140 | -3.460 | -0.961 | 0.225 | 1.545 | 0.577 | 1.745 | 2.048 | -0.291 | -0.499 | | | | | | | | | | | |
| B7L74_07325 | delta-aminolevulinic acid dehydratase | Ion Binding | 0.175 | -0.789 | -0.401 | 0.639 | -1.436 | -1.535 | -0.002 | -1.554 | -1.914 | 0.263 | 0.200 | | | | | | | | | | | |
| B7L74_07330 | hypothetical proteinprotein=17 kDa common-antigen | Effector protein | 1.094 | 0.065 | 1.212 | 0.747 | -0.265 | -0.491 | 1.200 | -0.715 | -0.581 | 1.361 | 1.853 | | | Yes | | | | | | | | |
| B7L74_07335 | hypothetical protein | | 2.696 | 0.847 | -0.035 | -0.095 | 2.736 | 1.627 | 1.469 | 1.496 | 2.628 | 0.453 | 1.229 | | | | | | | | | | | |
| B7L74_07340 | hypothetical protein | | -0.399 | 0.280 | -0.282 | 0.010 | 0.485 | 1.977 | 0.563 | 1.802 | 1.372 | -0.338 | -0.178 | | | | | | | | | | | |
| B7L74_07345 | hypothetical protein | | 0.061 | 0.842 | -0.034 | -1.788 | 2.061 | 1.623 | 1.917 | 1.959 | 2.728 | -0.653 | -0.904 | | | | | | | | | | | |
| B7L74_07350 | hypothetical membrane spanning protein | | 1.527 | -0.165 | 3.001 | 2.172 | -0.965 | -0.426 | 1.976 | -0.404 | -0.490 | 3.357 | 3.671 | Late up | | | | | | | | | | |
| B7L74_07355 | hypothetical protein | | 3.119 | 1.312 | 2.083 | -1.627 | 2.936 | 3.830 | 1.576 | 0.850 | 2.686 | 1.430 | 0.244 | Early up | | | | | | | | | | |
| B7L74_07360 | tRNA pseudouridine(55) synthase TruB | | 1.027 | 0.823 | 1.485 | -0.215 | 0.540 | 1.000 | 0.749 | 1.708 | 1.317 | 0.335 | 0.720 | | | | | | | | | | | |
| B7L74_07365 | ribosome-binding factor A | | 0.060 | 0.842 | -0.034 | -1.789 | 1.973 | 1.623 | -0.220 | 1.491 | 1.136 | -1.269 | -0.904 | | | | | | | | | | | |
| B7L74_07370 | translation initiation factor IF-2 | | -0.391 | -0.141 | -2.571 | -0.969 | -1.304 | -1.589 | -1.043 | -1.765 | -2.738 | -0.954 | -0.293 | | | | | | | | | | | |
| B7L74_07375 | transcription termination/antitermination protein NusA | | -1.002 | -0.044 | -1.350 | 0.585 | -1.125 | -0.465 | -0.713 | -1.143 | -2.243 | -0.619 | -0.093 | | | | | | | | | | | |
| B7L74_07380 | ribosome maturation factor | Effector protein | 0.693 | -0.478 | 0.982 | 0.330 | 0.820 | 2.133 | 0.208 | -0.844 | 1.029 | 1.059 | 0.987 | | | Yes | | | | | | | | |
| B7L74_07385 | tRNA-Met | tRNA | 0.028 | 0.248 | 3.321 | 0.809 | 0.176 | -0.440 | 1.482 | 0.993 | -0.975 | 2.607 | 3.100 | Early up | | | | | | | | | | |
| B7L74_07390 | NADH-quinone oxidoreductase subunit N | Respiration/NADH | -1.864 | -1.800 | -2.837 | -2.276 | -2.979 | -1.835 | -2.467 | -2.999 | -3.969 | -2.244 | -2.241 | Continuous down | | | | | | Yes | | | | |
| B7L74_07395 | NADH-quinone oxidoreductase subunit M | Respiration/NADH | -2.367 | -1.646 | -4.996 | -3.466 | -2.973 | -2.347 | -3.172 | -3.299 | -3.921 | -2.861 | -3.103 | Early down | | | | | | Yes | | | | |
| B7L74_07400 | NADH-quinone oxidoreductase subunit L | Respiration/NADH | -2.603 | -2.107 | -5.562 | -3.407 | -2.866 | -2.124 | -3.057 | -3.235 | -2.668 | -3.139 | -3.159 | Early down | | | | | | Yes | | | | |
| B7L74_07405 | NADH-quinone oxidoreductase subunit K | Respiration/NADH | -2.564 | -1.791 | -5.297 | -4.334 | -2.176 | -2.112 | -3.269 | -3.553 | -2.965 | -2.946 | -2.598 | Early down | | | | | | Yes | | | | |
| B7L74_07410 | NADH:ubiquinone oxidoreductase subunit J | Respiration/NADH | -0.459 | 0.379 | -0.034 | -2.817 | 2.204 | 0.618 | -1.215 | 0.942 | 2.450 | -1.587 | -1.915 | | | | | | | Yes | | | | |
| B7L74_07415 | NADH-quinone oxidoreductase subunit I | Respiration/NADH | -2.911 | -1.488 | -4.510 | -3.886 | -2.545 | -1.721 | -2.715 | -3.832 | -3.526 | -2.871 | -2.864 | Early down | | | | | | Yes | | | | |
| B7L74_07420 | NADH-quinone oxidoreductase subunit H | Respiration/NADH | -2.185 | -1.711 | -3.652 | -3.208 | -3.239 | -2.353 | -3.308 | -3.669 | -3.827 | -2.306 | -2.188 | Continuous down | | | | | | Yes | | | | |
| B7L74_07425 | NADH-quinone oxidoreductase subunit G | Respiration/NADH | 0.073 | -0.385 | -2.762 | -0.812 | -0.479 | -1.228 | -0.262 | -1.929 | -3.098 | 0.441 | -0.611 | | | | | | | Yes | | | | |
| B7L74_07430 | NADH-quinone oxidoreductase subunit F | Respiration/NADH | -1.572 | -1.186 | -3.402 | -2.240 | -1.821 | -2.057 | -2.384 | -2.561 | -3.111 | -1.707 | -1.417 | Early down | | | | | | Yes | | | | |
| B7L74_07435 | NADH-quinone oxidoreductase subunit E | Respiration/NADH | -2.312 | -1.851 | -4.281 | -3.106 | -3.359 | -2.795 | -3.227 | -3.332 | -4.225 | -2.850 | -2.870 | Continuous down | | | | | | Yes | | | | |
| B7L74_07440 | NADH-quinone oxidoreductase subunit D | Respiration/NADH | -2.296 | -1.990 | -3.791 | -2.526 | -3.168 | -2.612 | -2.809 | -3.283 | -4.181 | -2.615 | -2.732 | Continuous down | | | | | | Yes | | | | |
| B7L74_07445 | NADH-quinone oxidoreductase subunit C | Respiration/NADH | 0.044 | 0.838 | -0.034 | 0.125 | 2.646 | 2.011 | 1.140 | 1.488 | 2.551 | -1.278 | -0.909 | | | | | | | Yes | | | | |
| B7L74_07450 | NADH-quinone oxidoreductase subunit B | Respiration/NADH | -0.245 | -0.739 | -1.648 | -2.366 | -1.092 | -1.724 | -1.401 | -2.266 | -2.210 | -2.377 | -2.562 | Early down | | | | | | Yes | | | | |
| B7L74_07455 | NADH-quinone oxidoreductase subunit A | Respiration/NADH | -1.730 | -2.250 | -3.750 | -2.594 | -0.564 | -0.200 | -2.151 | -2.670 | 1.047 | -1.432 | -2.521 | | | | | | | Yes | | | | |
| B7L74_07460 | tRNA-Leu | tRNA | 0.199 | -3.653 | -1.081 | -2.319 | -0.776 | -1.684 | -3.979 | -3.005 | -0.555 | -1.640 | -1.599 | | | | | | | | | | | |
| B7L74_07465 | preprotein translocase subunit SecG | Exported | -0.431 | -3.022 | 0.142 | -1.103 | -2.194 | -1.784 | -5.677 | -5.086 | -2.902 | -1.254 | -0.339 | Late down | | | | | | | | | | |
| B7L74_07470 | triose-phosphate isomerase | | -0.826 | -0.764 | 0.376 | -0.440 | -1.178 | -1.150 | -0.584 | -1.161 | -1.430 | -0.239 | 1.108 | | | | | Yes | | | | | | |
| B7L74_07475 | hypothetical protein | | 4.431 | 3.109 | 7.862 | 2.990 | 2.678 | 2.206 | 5.597 | 3.517 | 4.747 | 5.991 | 6.030 | Continuous up | | | | | | | | | | |
| B7L74_07480 | peptidylprolyl isomerase | | -0.947 | -0.764 | -1.106 | -0.127 | -0.233 | -0.266 | -0.513 | 0.110 | 0.507 | 0.108 | 0.459 | | | | Yes | | | | | | | |
| B7L74_07485 | hypothetical protein | | -0.449 | 0.621 | -1.237 | -0.188 | -0.245 | -0.065 | -1.188 | 0.561 | -0.613 | -0.952 | -1.334 | | | | | | | | | | | |
| B7L74_07490 | hypothetical protein | | -0.823 | 0.023 | 0.651 | 0.032 | -0.049 | -0.553 | 0.691 | -0.103 | 0.174 | 0.154 | 0.144 | | | | | | | | | | | |
| B7L74_07495 | hypothetical protein | | 2.883 | 0.528 | 0.668 | -0.516 | 1.199 | 4.060 | 1.799 | 0.098 | 3.882 | 1.277 | 0.476 | Early up | | | | | | | | | | |
| B7L74_07500 | hypothetical protein | | 4.325 | 0.011 | 0.957 | 0.717 | 6.013 | 4.934 | 1.700 | -0.236 | 2.676 | 2.592 | 2.730 | Early up | | | | | | | | | | |
| B7L74_07505 | hypothetical protein tetratricopeptide repeat family prot | Effector protein | 1.047 | 0.115 | 0.349 | -0.607 | 1.003 | 0.757 | 0.618 | 0.209 | 0.222 | 0.315 | 0.391 | | | Yes | | | | | | | | |
| B7L74_07510 | ATPase | ATP | -0.382 | -0.685 | 0.031 | 0.229 | -0.790 | 0.133 | -0.412 | 0.391 | 0.316 | 0.210 | -0.029 | | | | | | | | | | | |
| B7L74_07515 | hypothetical protein | | 1.411 | 0.454 | 0.416 | 0.333 | 2.904 | 3.551 | 0.494 | 0.757 | 1.389 | 1.184 | 0.318 | | | | | | | | | | | |
| B7L74_07520 | hypothetical exported protein | | 6.039 | 1.947 | 2.441 | 1.394 | 7.194 | 6.249 | 4.286 | 1.733 | 4.860 | 3.131 | 1.524 | Early up | | | | | | | | | | |
| B7L74_07525 | hypothetical protein | Effector protein | -0.842 | -0.153 | 2.143 | 2.441 | 0.123 | -0.145 | 0.629 | -0.075 | 0.642 | 0.617 | 1.074 | Up in P12 | | Yes | | | | | | | | |
| B7L74_07530 | hypothetical protein | | 3.454 | 0.598 | 2.108 | 0.080 | 4.970 | 4.655 | 2.604 | 1.887 | 3.586 | 2.494 | 2.275 | Early up | | | | | | | | | | |
| B7L74_07535 | hypothetical protein | | 3.144 | 1.027 | 2.877 | -0.205 | 4.650 | 4.659 | 3.153 | 1.825 | 2.749 | 1.819 | 1.918 | Early up | | | | | | | | | | |
| B7L74_07540 | tRNA-Asp | tRNA | 0.112 | 0.274 | 3.166 | -0.012 | 0.161 | -0.120 | 2.473 | 1.370 | 0.491 | 0.671 | 1.166 | Early up | | | | | | | | | | |
| B7L74_07545 | tRNA-Val | tRNA | 1.148 | 0.621 | -0.160 | -0.882 | 0.770 | 0.479 | 1.226 | 1.606 | -0.270 | -0.199 | -0.030 | | | | | | | | | | | |
| B7L74_07550 | hypothetical protein | | 0.145 | -0.298 | -0.920 | -0.022 | -0.695 | -0.772 | 0.017 | -0.636 | -1.587 | -0.007 | -0.176 | | | | | | | | | | | |
| B7L74_07555 | DNA-binding protein HU | DNA replication proteins | -0.688 | -0.879 | -0.107 | 2.383 | -1.198 | -1.531 | 0.200 | -1.075 | -1.370 | 0.205 | 0.455 | Up in P12 | | | | | | | | | | |
| B7L74_07560 | EVE domain-containing protein | | -0.572 | -0.829 | -1.248 | 0.448 | -0.965 | -1.228 | -0.496 | -1.222 | -1.654 | 0.392 | -0.002 | | | | | | | | | | | |
| B7L74_07565 | hypothetical protein | | -0.276 | -0.951 | -2.159 | -0.040 | -1.032 | -0.784 | -0.237 | -0.789 | -1.511 | -0.470 | -0.649 | | | | | | | | | | | |
| B7L74_07570 | metalloprotease TldD | | -1.511 | -3.308 | -1.932 | -1.072 | -2.469 | -2.708 | -2.604 | -7.405 | -1.368 | -0.783 | -0.874 | | | | | | | | | | | |
| B7L74_07575 | DUF3971 domain-containing protein | CB specific protein family | -1.064 | -1.143 | -1.775 | -0.822 | -1.635 | -0.814 | -1.158 | -1.478 | -1.536 | -0.486 | -0.387 | Early down | | | | | | | | | | |
| B7L74_07580 | rod shape-determining protein MreD | | 0.547 | 1.377 | 0.966 | -0.416 | 3.080 | 2.930 | 1.869 | 1.488 | 2.552 | 0.123 | 0.588 | | | | | | | | | | | |
| B7L74_07585 | rod shape-determining protein MreC | Chromosome and associated prote | -1.371 | -1.655 | -2.887 | -1.377 | -2.137 | -2.098 | -1.366 | -2.903 | -3.719 | -1.606 | -1.436 | Early down | | | | | | | | | | |
| B7L74_07590 | rod shape-determining protein | Chromosome and associated prote | -0.328 | -1.238 | -1.414 | -0.170 | -2.046 | -2.198 | -0.381 | -3.139 | -3.829 | -0.195 | -0.265 | Continuous down | | | | | | | | | | |
| B7L74_07595 | hypothetical protein | | 0.098 | 0.136 | 0.164 | -1.048 | -0.172 | -1.194 | 1.362 | -0.013 | -0.295 | 0.146 | 0.497 | | | | | | | | | | | |
| B7L74_07600 | aspartyl/glutamyl-tRNA(Asn/Gln) amidotransferase su | | -1.482 | -1.253 | -3.054 | 0.322 | -1.738 | -0.481 | -0.982 | -1.397 | -1.074 | -0.681 | -1.070 | | | | | | | | | | | |
| B7L74_07605 | glutamyl-tRNA(Gln) amidotransferase subunit A | | -0.421 | -0.925 | -2.077 | -1.301 | -1.667 | -0.610 | -1.242 | -1.909 | -1.923 | -1.661 | -1.829 | | | | | | | | | | | |
| B7L74_07610 | aspartyl/glutamyl-tRNA(Asn/Gln) amidotransferase su | Aminoacyl-tRNA synthase | -1.999 | -1.659 | -4.161 | -1.811 | -2.877 | -2.079 | -2.376 | -2.116 | -2.599 | -2.636 | -2.569 | Early down | | | | | | | | | | |
| B7L74_07615 | DNA-binding transcriptional regulator OxyR(Hydroge | Transcriptional regulators | 1.122 | 0.353 | 2.646 | 0.838 | 0.988 | 0.508 | 0.814 | 0.036 | 0.458 | 0.942 | 1.311 | | | | | | | | | | | |
| B7L74_07620 | alkyl hydroperoxide reductase, ahpC | PV detoxification | 2.264 | -0.164 | 5.971 | 2.637 | 1.687 | 2.466 | 7.286 | 3.427 | 3.670 | 2.457 | 3.157 | Early up | | | | | | | | Yes (Ions/iron) | | |
| B7L74_07625 | alkyl hydroperoxide reductase, aphD | PV detoxification | 0.902 | 0.435 | 1.536 | 0.449 | -0.295 | 1.240 | 2.717 | 1.728 | 1.151 | 1.099 | 1.040 | | | | | | | | | | | |
| B7L74_07630 | hypothetical protein | | 0.561 | 0.909 | -1.398 | 1.075 | 1.246 | 1.935 | 0.607 | 3.104 | 2.265 | 1.305 | 1.612 | | | | | | | | | | | |
| B7L74_07635 | pilus assembly protein PilW | | 1.856 | 0.720 | 2.097 | 1.156 | 1.592 | 1.152 | 1.657 | 1.986 | 2.096 | 1.339 | 1.792 | | | | | | | | | | | |
| B7L74_07640 | pilus assembly protein PilV | | -0.859 | 0.543 | -0.014 | 3.192 | 1.771 | 0.551 | 1.824 | 2.341 | 3.502 | -2.214 | -1.380 | | | | | | | | | | | |
| B7L74_07645 | hypothetical proteinmembrane protease family, stomati | Eucaryotic-like ORFs. | -1.849 | -0.678 | -2.670 | 0.625 | 1.021 | 1.128 | -0.436 | -0.188 | 0.766 | -0.338 | -1.670 | | | | | | | | | | | |
| B7L74_07650 | serine protease | Protein processing | 3.426 | -0.660 | -0.443 | 0.322 | 5.091 | 3.887 | 1.763 | 0.745 | 4.782 | 0.420 | -0.217 | Early up | | | | | | | | | | |
| B7L74_07655 | MFS transporter | Transporter | 0.141 | 0.124 | -0.153 | -0.136 | -0.920 | -0.410 | 0.061 | -0.720 | -2.038 | -0.243 | -0.375 | | | | | | | | | | | |
| B7L74_07660 | cyclin | | -1.081 | -0.250 | -0.069 | 4.107 | 2.291 | -0.850 | 1.270 | 0.785 | 4.878 | 3.208 | 1.664 | | | | | | | | | | | |
| B7L74_07665 | hypothetical protein | | 2.261 | 1.203 | 0.287 | -0.096 | 0.769 | 1.631 | 2.407 | 0.440 | 0.333 | -0.114 | 1.736 | | | | | | | | | | | |
| B7L74_07670 | cysteine--tRNA ligase | | -0.336 | 0.128 | -1.078 | 0.535 | -0.299 | -0.042 | -0.306 | -0.405 | -1.380 | -0.380 | 0.052 | | | | | | | | | | | |
| B7L74_07675 | glutamate--tRNA ligase | | 1.527 | 1.237 | 0.171 | 0.678 | 1.090 | 1.355 | 0.109 | -0.157 | 1.658 | 0.411 | -0.623 | | | | | | | | | | | |
| B7L74_07680 | UDP-2%2C3-diacylglucosamine hydrolase | Lipopolysaccharide biosynthesis | 0.163 | -0.095 | 1.604 | 1.059 | -0.264 | 0.495 | 0.479 | 1.203 | 1.007 | 0.659 | 0.467 | | | | | | | | | | Yes | |
| B7L74_07685 | transcriptional regulator | Virulence, Prokaryotic defense sys | 0.492 | 0.586 | 2.298 | 1.499 | 0.221 | -0.325 | 1.029 | 0.648 | -0.173 | 1.676 | 1.210 | Early up | | | | | | | | | | |
| B7L74_07690 | plasmid maintenance system killer protein | | -0.171 | 0.459 | 2.116 | 1.120 | 0.505 | -0.054 | 1.318 | 1.415 | 0.421 | 1.680 | 1.697 | | | | | | | | | | | |
| B7L74_07695 | ISAs1 family transposase | | 2.362 | 0.691 | -0.029 | 0.746 | 1.666 | 3.783 | 1.599 | 3.495 | 3.669 | 1.208 | 1.324 | Early up | | | | | | | | | | |
| B7L74_07700 | hypothetical protein | Effector protein | -0.886 | 0.513 | -0.425 | -0.285 | -0.725 | -0.400 | -1.206 | 0.021 | -0.954 | -0.409 | -0.655 | | | Yes | | | | | | | | |
| B7L74_07705 | pyridoxine 5'-phosphate synthase | Vitamin B6 metabolism | 0.341 | 0.693 | 3.585 | 0.745 | 1.369 | 0.561 | 1.863 | 0.623 | 2.558 | 1.931 | 2.764 | Early up | | | | | | | | | | |
| B7L74_07710 | hypothetical protein | | -0.130 | 0.388 | 0.254 | 0.388 | 0.704 | 1.065 | -0.720 | 1.430 | 0.922 | 0.405 | -0.388 | | | | | | | | | | | |
| B7L74_07715 | hypothetical protein | | -0.906 | -0.211 | -0.097 | -1.091 | -0.398 | 0.263 | -0.858 | -0.400 | -0.127 | -0.959 | -1.144 | | | | | | | | | | | |
| B7L74_07720 | hypothetical protein | | 0.634 | -0.504 | 1.585 | 2.247 | -0.671 | 0.766 | 1.258 | 0.856 | -0.717 | 2.293 | 2.724 | | | | | | | | | | | |
| B7L74_07725 | AAA family ATPase | | 1.550 | 1.347 | 1.246 | 0.367 | 3.424 | 3.228 | 2.760 | 3.596 | 2.748 | 1.729 | 3.615 | Early up | | | | | | | | | | |
| B7L74_07730 | DNA repair protein RecO | DNA | 1.098 | -0.030 | 0.773 | 1.513 | 1.366 | 0.824 | 1.586 | -0.109 | 0.974 | 1.204 | 1.013 | | | | | | | | | | | |
| B7L74_07735 | GTPase Era | | 0.594 | -1.563 | -1.397 | -0.370 | 0.499 | 0.735 | -0.609 | -2.188 | -0.864 | -0.911 | -0.541 | | | | | | | | | | | |
| B7L74_07740 | ribonuclease 3 | | -0.413 | 0.888 | 0.649 | -0.858 | 1.290 | 1.052 | 0.915 | 1.149 | 3.270 | -0.631 | -0.296 | | | | | | | | | | | |
| B7L74_07745 | S26 family signal peptidase | Protein export/signal peptidase | -0.542 | -0.618 | -0.980 | 0.332 | -1.787 | -0.754 | -0.065 | -1.178 | -2.023 | 0.103 | 0.450 | Down in P51 | | | | | | | | | | |
| B7L74_07750 | elongation factor 4 | | 0.601 | -0.379 | 2.706 | 0.511 | 0.352 | 0.070 | 0.847 | 0.078 | 0.146 | 0.689 | 1.472 | | | | | | | | | | | |
| B7L74_07755 | thioesterase | | -0.481 | -0.343 | -2.958 | -2.055 | -1.095 | -0.822 | -1.204 | -1.771 | -2.081 | -1.849 | -2.425 | Late down | | | | | | | | | | |
| B7L74_07760 | copper-translocating P-type ATPase | Transporter | -1.220 | -0.938 | -1.670 | -1.765 | -1.011 | -0.571 | -1.522 | -1.390 | -2.558 | -2.077 | -2.317 | Early down | | | | | | | | Yes (Ions/Copper) | | |
| B7L74_07765 | ATPase | ATP | -0.469 | -0.114 | 0.066 | -0.882 | -0.064 | 0.073 | -0.613 | -0.409 | -0.993 | -0.516 | -0.904 | | | | | | | | | | | |
| B7L74_07770 | tRNA(Ile)-lysidine synthetase | | 0.508 | 0.202 | -0.076 | -0.237 | 0.650 | 0.709 | 0.053 | -0.335 | -0.617 | -0.121 | -0.355 | | | | | | | | | | | |
| B7L74_07775 | acetyl-CoA carboxylase carboxyl transferase subunit a | | -0.033 | -0.440 | -0.335 | -0.557 | -1.817 | -0.682 | 0.011 | -2.554 | -0.179 | -0.112 | -0.316 | | | | | | | | | | | |
| B7L74_07780 | zinc transporter | Transporter | 0.119 | 0.609 | -1.451 | -0.125 | -0.175 | -0.779 | -0.819 | 0.057 | -1.888 | -1.285 | -1.402 | | | | | | | | | | | |
| B7L74_07785 | rhodanese-like domain-containing protein | | -1.116 | 0.460 | -2.843 | -2.554 | -0.414 | 0.010 | -2.191 | -1.535 | -1.545 | -2.295 | -2.911 | | | | | | | | | | | |
| B7L74_07790 | NAD(P)-dependent oxidoreductase | | -0.311 | 0.835 | -0.397 | -0.636 | 0.116 | 0.196 | -0.792 | 0.523 | 0.548 | -1.481 | -1.325 | | | | | | | | | | | |
| B7L74_07795 | short-chain dehydrogenase | | -1.008 | 0.822 | -0.698 | -0.515 | -1.269 | -0.042 | -1.214 | 0.313 | -0.896 | -1.377 | -1.709 | | | | | | | | | | | |
| B7L74_07800 | hypothetical protein | | -0.292 | 0.749 | -0.255 | -1.137 | 0.928 | 0.965 | -0.684 | 1.379 | 0.054 | 0.533 | -0.596 | | | | | | | | | | | |
| B7L74_07805 | hypothetical protein | | -0.882 | 1.656 | 0.659 | -0.874 | 1.646 | 0.762 | -0.659 | 1.885 | 2.990 | 0.393 | -0.211 | | | | | | | | | | | |
| B7L74_07810 | hypothetical protein | | 0.170 | 1.750 | 0.900 | -0.580 | 1.443 | 0.721 | -0.980 | 1.029 | 2.039 | 0.456 | -0.322 | | | | | | | | | | | |
| B7L74_07815 | tRNA (uridine(34)/cytosine(34)/5-carboxymethylamin | | 0.157 | 0.149 | 1.747 | 0.380 | 0.118 | -0.125 | 0.047 | 0.661 | 1.328 | -0.129 | -0.140 | | | | | | | | | | | |
| B7L74_07820 | glycerol-3-phosphate dehydrogenase (NAD(P)(+)) | Glycerophospholipid metabolism | -0.844 | -0.545 | -2.037 | -0.308 | -1.459 | -1.469 | -1.188 | -1.177 | -2.431 | -1.824 | -1.650 | Early down | | | | | | | | | | |
| B7L74_07825 | protein-export protein SecB | Protein export/sec pathway | -0.928 | -0.652 | -1.401 | 0.267 | -1.094 | -1.201 | -1.026 | -1.798 | -3.343 | -1.418 | -1.051 | Late down | | | | | | | | | | |
| B7L74_07830 | glutaredoxin 3 | PV detoxification<br>Chaperones and folding catalysts | -1.327 | -0.610 | -0.537 | -0.425 | -1.153 | -1.153 | -1.367 | -1.508 | -3.604 | -1.528 | -1.231 | Late down | | | Yes | | | | | | | |
| B7L74_07835 | sulfurtransferase | | -0.775 | -0.280 | -0.131 | 0.668 | -0.979 | -0.714 | -0.839 | -0.479 | -2.239 | -0.571 | -0.484 | Down in P51 | | | | | | | | | | |
| B7L74_07840 | hypothetical protein | | 4.895 | 0.432 | 2.279 | -0.937 | 5.609 | 5.845 | 2.489 | 1.529 | 5.881 | 4.024 | 1.213 | Early up | | | | | | | | | | |
| B7L74_07845 | hypothetical protein | Effector protein | 1.179 | -0.217 | -0.673 | -0.592 | 3.219 | 2.756 | -0.045 | 0.437 | 2.900 | 0.500 | -0.871 | Early up | | Yes | | | | | | | | |
| B7L74_07850 | hypothetical protein | Effector protein | -0.881 | -1.008 | -1.547 | -0.981 | -1.862 | -1.965 | -1.525 | -1.495 | -2.400 | -0.757 | -0.662 | Early down | | Yes | | | | | | | | |
| B7L74_07855 | hypothetical protein | | 0.055 | 0.789 | -1.619 | -2.089 | 1.022 | 1.775 | 0.100 | 1.104 | 0.935 | -2.175 | -2.067 | | | | | | | | | | | |
| B7L74_07860 | hypothetical protein | | -0.033 | 0.266 | -0.324 | -0.163 | -0.409 | -0.740 | -0.548 | -0.143 | -0.697 | -1.014 | -0.998 | | | | | | | | | | | |
| B7L74_07865 | peptidase | | -1.036 | 0.140 | -1.693 | -1.638 | -0.327 | -0.391 | -1.571 | -1.496 | -0.647 | -1.952 | -1.786 | Early down | | | | | | | | | | |
| B7L74_07870 | hypothetical membrane spanning protein | Effector protein | 0.408 | 1.167 | 0.150 | -0.766 | 0.431 | 0.119 | 1.129 | 0.076 | 0.548 | -1.145 | -0.774 | | | Yes | | | | | | | | |
| B7L74_07875 | hypothetical protein | | 0.229 | 1.099 | 2.232 | -0.021 | 0.536 | 0.884 | 0.657 | 1.786 | 0.784 | 0.504 | 0.468 | | | | | | | | | | | |
| B7L74_07880 | hypothetical protein | Effector protein | 1.452 | 0.513 | 2.179 | 0.370 | 2.459 | 2.226 | 1.158 | 1.642 | 1.856 | 0.694 | 0.695 | | | Yes | | | | | | | | |
| B7L74_07885 | hypothetical protein | | 1.501 | -0.832 | -0.921 | -1.331 | 1.775 | 2.311 | -3.918 | 1.496 | 2.785 | -0.604 | -1.514 | | | | | | | | | | | |
| B7L74_07890 | hypothetical protein | | -0.045 | 1.878 | 2.563 | -0.382 | 1.307 | 1.983 | 0.046 | 2.326 | 1.198 | 0.763 | 1.311 | | | | | | | | | | | |
| B7L74_07895 | hypothetical protein | | -0.642 | 0.553 | -1.280 | -1.741 | 0.157 | 0.629 | -0.450 | 0.511 | -0.775 | -0.700 | -2.215 | | | | | | | | | | | |
| B7L74_07900 | 2%2C3-bisphosphoglycerate-independent phosphoglyc | | -0.404 | -0.234 | -1.476 | 0.403 | -1.188 | -0.328 | -0.934 | -0.549 | -1.542 | -0.796 | -0.408 | | | | | Yes | | | | | | |
| B7L74_07905 | peptidase M23 | Cell growth | -0.689 | -0.359 | -0.772 | -0.527 | -0.718 | -0.217 | -0.924 | -1.333 | -3.073 | -0.982 | -1.067 | Down in P51 | | | | | | | | | | |
| B7L74_07910 | peptidase | | 0.009 | 0.751 | -1.836 | -3.519 | 4.390 | 1.196 | -0.736 | 0.078 | 3.376 | -2.885 | -2.580 | | | | | | | | | | | |
| B7L74_07915 | tryptophan/tyrosine permease | | 1.155 | 0.113 | -0.921 | -1.017 | 1.621 | 1.348 | -0.237 | -0.930 | -0.459 | -1.026 | -1.146 | | | | | | | | | Yes (Amino acid/tyrosine) | | |
| B7L74_07920 | hypothetical protein | | -0.824 | 0.827 | -0.464 | -0.893 | -0.046 | 0.336 | -0.935 | -0.356 | -0.093 | -1.739 | -1.337 | | | | | | | | | | | |
| B7L74_07925 | hypothetical protein | | -0.624 | 0.805 | 0.429 | -0.757 | 0.932 | -0.277 | -0.457 | -0.258 | -0.973 | -0.233 | -0.483 | | | | | | | | | | | |
| B7L74_07930 | hypothetical membrane associated protein | | -0.297 | 0.266 | 0.823 | 0.424 | 0.739 | 1.071 | -1.152 | -0.321 | 0.457 | -0.591 | -0.468 | | | | | | | | | | | |
| B7L74_07935 | hypothetical protein | Effector protein | 1.858 | 0.944 | 0.314 | -0.008 | 3.554 | 2.659 | 0.616 | 0.244 | 2.239 | 0.757 | -0.674 | | | Yes | | | | | | | | |
| B7L74_07940 | IS30 family transposase | Replication and repair | 3.531 | 1.061 | 0.804 | -0.453 | 4.007 | 4.883 | 1.846 | 2.171 | 2.963 | 1.029 | 0.073 | Early up | | | | | | | | | | |
| B7L74_07945 | hypothetical protein | | 7.348 | 2.545 | 2.592 | 1.983 | 6.910 | 7.983 | 3.824 | 4.372 | 7.036 | 4.149 | 1.338 | Early up | | | | | | | | | | |
| B7L74_07950 | thymidylate synthase | Pyrimidine metabolism | 0.207 | 0.668 | -1.863 | -0.847 | 0.446 | -0.105 | -0.632 | 0.075 | -2.040 | -1.157 | -1.497 | Early down | | | | | | | | | | |
| B7L74_07955 | hypothetical protein, Hypothetical membrane spanning | Membrane | 0.041 | 0.838 | -0.034 | -1.805 | 1.973 | 2.389 | 0.448 | 1.487 | 3.219 | -0.379 | -0.910 | | | | | | | | | | | |
| B7L74_07960 | prolipoprotein diacylglyceryl transferase | | -0.429 | 0.064 | -0.801 | 0.803 | -0.732 | -0.651 | -0.806 | -0.610 | -2.087 | -0.347 | -0.157 | | | | | | | | | | | |
| B7L74_07965 | phosphoenolpyruvate--protein phosphotransferase | | 0.215 | 0.774 | 1.054 | -1.326 | 0.054 | 0.508 | -0.385 | -0.732 | -0.202 | -0.771 | -0.402 | | | | | | | | | | | |
| B7L74_07970 | RNA pyrophosphohydrolase | RNA degradation | 1.662 | 2.144 | 4.724 | 0.767 | 4.042 | 2.301 | 3.847 | 2.388 | 2.828 | 1.743 | 3.290 | Early up | | | | | | | | | | |
| B7L74_07975 | hypothetical protein | | -1.367 | -0.205 | 3.597 | 2.239 | -0.157 | 0.823 | 1.414 | 0.463 | 0.021 | 2.001 | 1.518 | | | | | | | | | | | |
| B7L74_07980 | ribonucleoside-diphosphate reductase subunit alpha | | -0.592 | -0.255 | 1.792 | 1.040 | -0.614 | -0.620 | 0.603 | -0.593 | -1.284 | 0.552 | 0.511 | | | | | | | | | | | |
| B7L74_07985 | ribonucleotide-diphosphate reductase subunit beta | | -0.836 | -0.703 | 0.091 | 0.305 | -0.849 | -0.407 | -0.484 | -1.459 | -0.993 | -0.936 | -0.628 | | | | | | | | | | | |
| B7L74_07990 | hypothetical protein | | 2.178 | -0.771 | 2.651 | 0.867 | 2.320 | 2.801 | 1.425 | -0.965 | 1.988 | 1.579 | 1.320 | | | | | | | | | | | |
| B7L74_07995 | ISAs1 family transposase | | 2.772 | 0.713 | 0.389 | 0.316 | 2.634 | 4.265 | 1.829 | 3.326 | 3.534 | 1.217 | 1.205 | Early up | | | | | | | | | | |
| B7L74_08000 | hypothetical protein(cvpC in Table 1 of RightonQ pap | Effector protein | 0.170 | 0.074 | -0.495 | -1.028 | -0.045 | 1.733 | 0.197 | 0.759 | 0.231 | -0.566 | -0.424 | | | Yes | | | | | | | | |
| B7L74_08005 | hypothetical protein | | 0.578 | -0.142 | -1.018 | -1.633 | 0.307 | 1.397 | 0.351 | 1.293 | 1.027 | -1.485 | -0.500 | | | | | | | | | | | |
| B7L74_08010 | FmdB family transcriptional regulator | | 0.256 | 0.076 | 0.679 | 1.094 | -0.594 | -0.461 | 0.257 | -0.377 | -0.349 | 0.499 | 0.690 | | | | | | | | | | | |
| B7L74_08015 | hypothetical protein, Hypothetical membrane spanning | Membrane | 2.685 | 1.905 | -0.608 | -1.962 | 5.731 | 4.946 | 3.719 | 2.122 | 2.040 | -1.819 | -1.466 | Early up | | | | | | | | | | |
| B7L74_08020 | hypothetical protein | | 1.153 | 0.387 | 0.693 | 0.392 | 2.193 | 2.566 | 0.579 | 1.143 | 0.499 | 1.422 | 0.411 | | | | | | | | | | | |
| B7L74_08025 | hypothetical protein | | 2.717 | 0.731 | 1.443 | 0.914 | 3.782 | 3.834 | 2.032 | 1.154 | 3.017 | 3.535 | 1.891 | Early up | | | | | | | | | | |
| B7L74_08030 | hypothetical membrane spanning protein | | 2.622 | 0.934 | 1.762 | -0.587 | 3.912 | 4.206 | 1.257 | 1.446 | 3.293 | 1.473 | 1.197 | Early up | | | | | | | | | | |
| B7L74_08035 | hypothetical protein | | 5.624 | 1.091 | -14.577 | -2.738 | 7.839 | 9.610 | 0.435 | 0.887 | 8.975 | 1.972 | -0.542 | | | | | | | | | | | |
| B7L74_08040 | hypothetical protein | | 3.866 | 2.006 | 1.516 | 2.508 | 5.072 | 4.560 | 3.426 | 3.291 | 4.674 | 2.667 | 2.289 | Early up | | | | | | | | | | |
| B7L74_08045 | aspartate--tRNA ligase | | 0.115 | -0.122 | -1.042 | 1.377 | -0.008 | 0.417 | 0.537 | 0.526 | -0.859 | 0.095 | 0.567 | | | | | | | | | | | |
| B7L74_08050 | transcriptional regulator | Effector protein | 1.569 | 0.916 | -3.194 | 2.211 | 1.497 | 1.687 | 0.908 | 0.404 | -0.807 | 0.373 | 1.136 | | | Yes | | | | | | | | |
| B7L74_08055 | crossover junction endodeoxyribonuclease RuvC | Homologous recombination | 1.933 | 1.725 | 2.920 | 2.462 | 2.063 | 2.195 | 2.700 | 1.936 | 1.411 | 2.444 | 1.751 | Early up | | | | | | | | | | |
| B7L74_08060 | Holliday junction branch migration protein RuvA | Homologous recombination | 4.142 | 2.865 | 3.966 | 4.416 | 4.574 | 4.897 | 3.883 | 4.950 | 5.396 | 1.849 | 0.756 | Continuous up | | | | | | | | | | |
| B7L74_08065 | hypothetical protein(coxCC12 in Table 1 of RightonQ | Effector protein | -0.928 | -0.212 | -1.226 | -0.245 | -0.107 | 0.156 | -1.108 | 0.047 | 0.964 | -0.195 | -0.984 | | | Yes | | | | | | | | |
| B7L74_08070 | Holliday junction branch migration DNA helicase Ruv | | 0.766 | -0.033 | 0.252 | 0.751 | -0.786 | -0.038 | 0.511 | -0.819 | -1.637 | 0.362 | 0.320 | | | | | | | | | | | |
| B7L74_08075 | hypothetical protein | | 0.670 | -0.230 | 1.735 | -0.015 | -0.072 | 0.332 | 0.604 | -0.493 | -0.110 | 1.280 | 1.807 | | | | | | | | | | | |
| B7L74_08080 | IS110 family transposase | | -0.327 | 0.097 | 2.526 | 4.970 | -0.731 | -1.840 | 0.276 | 2.383 | 1.313 | 5.078 | 6.625 | | | | | | | | | | | |
| B7L74_08085 | MFS transporter | Transporter | 0.858 | -0.269 | 1.558 | -1.413 | 1.748 | 2.453 | 0.448 | 0.282 | 2.388 | 1.228 | 1.086 | Early up | | | | | | | | Yes (MFS) | | |
| B7L74_08090 | protein TolQ | | -0.603 | -0.112 | 0.308 | 1.830 | -0.618 | -0.206 | -0.182 | 0.580 | -0.865 | 0.719 | 0.579 | | | | | | | | | Yes (ion channels/mechanosensitive channel) | | |
| B7L74_08095 | protein TolR | | -1.267 | -0.500 | -1.379 | -3.117 | 2.944 | 2.142 | 0.238 | 1.754 | 3.015 | 0.858 | -2.244 | | | | | | | | | Yes (ion channels/mechanosensitive channel) | | |
| B7L74_08100 | hypothetical membrane spanning protein | Effector protein | -0.633 | 0.562 | 1.126 | 0.633 | 3.395 | 4.061 | 1.117 | 1.663 | 1.942 | 1.791 | 1.231 | Early up | | Yes | | | | | | | | |
| B7L74_08105 | peptidase M23 | | 8.467 | 0.698 | 5.875 | 0.431 | 8.824 | 7.916 | 6.345 | 3.432 | 9.314 | 3.840 | 2.507 | | | | | | | | | | | |
| B7L74_08110 | hypothetical protein protein=ribonuclease BN | | 2.130 | 1.002 | 1.787 | 1.261 | 2.780 | 2.842 | 1.154 | 2.362 | 3.295 | 2.435 | 1.822 | Early up | | | | | | | | | | |
| B7L74_08115 | NAD(P)H:quinone oxidoreductase%2C type IV | | -0.192 | 0.259 | 1.142 | 0.485 | -0.065 | 0.391 | 0.368 | 1.353 | 0.918 | 1.210 | 1.128 | | | | | | | | | | | |
| B7L74_08120 | cell division protein ZapE | | 0.091 | 0.848 | -0.034 | -1.759 | 3.371 | 4.217 | 3.600 | 1.497 | 2.602 | -1.251 | -0.895 | | | | | | | | | | | |
| B7L74_08125 | DUF1658 domain-containing protein | | -0.315 | 0.964 | -0.522 | -0.954 | 0.292 | 0.324 | -0.789 | 0.055 | -1.553 | -1.216 | -0.454 | | | | | | | | | | | |
| B7L74_08130 | DUF1658 domain-containing protein | | -0.508 | 0.011 | -1.541 | -1.223 | -0.463 | -0.234 | -1.230 | -0.316 | -1.985 | -1.205 | -1.121 | | | | | | | | | | | |
| B7L74_08135 | hypothetical protein | | 0.050 | 0.840 | -0.034 | -1.798 | 1.973 | 1.622 | 0.456 | 1.489 | 2.000 | -1.275 | -0.908 | | | | | | | | | | | |
| B7L74_08140 | Na+/H+ antiporter subunit G | Transporter | 2.248 | 0.102 | 0.598 | -0.389 | 3.589 | 4.777 | 0.356 | -0.003 | 3.816 | 1.261 | 0.431 | | | | | | | | | Yes (Na+:H+ antiporter ) | | |
| B7L74_08145 | cation:proton antiporter | Transporter | -1.273 | -1.321 | -4.455 | 0.025 | -0.131 | 1.318 | -1.554 | -0.434 | -0.163 | -1.622 | -0.757 | Down in P10 | | | | | | | | Yes (Na+:H+ antiporter ) | | |
| B7L74_08150 | sodium:proton antiporter | Transporter | -0.905 | -0.361 | -1.386 | -1.539 | 0.190 | 1.344 | -1.044 | 0.481 | 1.322 | -1.282 | -0.960 | | | | | | | | | Yes (Na+:H+ antiporter ) | | |
| B7L74_08155 | Na+/H+ antiporter subunit D | | -0.684 | 0.296 | -1.227 | -0.623 | 0.249 | 0.957 | -1.215 | 0.823 | 0.467 | -0.252 | -0.737 | | | | | | | | | Yes (Na+:H+ antiporter ) | | |
| B7L74_08160 | cation:proton antiporter | | -0.874 | -0.461 | -3.510 | -1.377 | -0.497 | -0.107 | -0.818 | -1.238 | -1.479 | -0.754 | -0.984 | | | | | | | | | Yes (Na+:H+ antiporter ) | | |
| B7L74_08165 | Na(+)/H(+) antiporter subunit B | Transporter | 2.009 | 0.313 | 2.008 | -0.928 | 3.143 | 4.090 | 2.348 | 1.162 | 4.047 | 3.039 | 2.702 | Early up | | | | | | | | Yes (Na+:H+ antiporter ) | | |
| B7L74_08170 | Na(+)/H(+) antiporter subunit A | Transporter | 2.250 | 0.895 | 7.282 | -2.622 | 7.740 | 8.687 | 7.245 | 5.700 | 9.258 | 7.737 | 8.147 | Continuous up | | | | | | | | Yes (Na+:H+ antiporter ) | | |
| B7L74_08175 | hypothetical protein, Hypothetical membrane associate | Membrane | 0.500 | -0.324 | 2.000 | 1.430 | -0.463 | -0.892 | 0.890 | -0.328 | -1.574 | 2.300 | 1.326 | | | | | | | | | | | |
| B7L74_08180 | sodium:proton antiporter | Transporter | -0.518 | -0.106 | -0.440 | -0.273 | 0.377 | 0.965 | -0.641 | 0.884 | 1.864 | 0.329 | 0.033 | | | | | | | | | Yes (Na+:H+ antiporter ) | | |
| B7L74_08185 | hypothetical protein | | 0.753 | -0.292 | 1.622 | -0.232 | -0.029 | 0.433 | 0.488 | -0.737 | -0.155 | 1.208 | 1.612 | | | | | | | | | | | |
| B7L74_08190 | IS110 family transposase | | -3.702 | 1.606 | 0.201 | -0.805 | -1.056 | -2.165 | -0.756 | 3.255 | 0.541 | 2.017 | 2.387 | | | | | | | | | | | |
| B7L74_08195 | 30S ribosomal protein S21 | Ribosomal proteins | -1.338 | -0.409 | -1.653 | 1.266 | -0.938 | -1.317 | -1.692 | 0.038 | -1.663 | -0.983 | -1.048 | | | | | | | | | | | |
| B7L74_08200 | glutamyl-tRNA amidotransferase | Effector protein | -1.164 | -0.593 | -3.636 | 0.243 | -1.308 | -1.778 | -1.272 | -0.818 | -2.225 | -1.261 | -0.710 | Early down | | Yes | | | | | | | | |
| B7L74_08205 | DNA primase | DNA | -0.736 | -0.768 | -0.308 | -0.935 | -0.191 | 0.262 | -0.316 | -0.241 | -0.102 | -0.530 | -0.240 | | | | | | | | | | | |
| B7L74_08210 | RNA polymerase sigma factor RpoD | | -1.362 | -0.024 | -0.691 | -0.802 | -0.075 | -0.101 | -1.134 | -0.041 | -0.998 | -1.417 | -1.273 | | | | | | | | | | | |
| B7L74_08215 | lipoprotein | | -0.650 | 0.910 | 0.193 | 0.049 | 0.123 | 0.336 | -0.275 | 1.154 | -0.567 | -0.627 | -0.467 | | | | | | | | | | | |
| B7L74_08220 | hypothetical protein | | 0.126 | -0.620 | 1.187 | 1.083 | -0.065 | 1.519 | 0.971 | 1.477 | 1.547 | 1.533 | 0.889 | | | | | | | | | | | |
| B7L74_08225 | hypothetical protein | Lysine degradation | 1.623 | -2.651 | 2.198 | 2.271 | -0.540 | 1.910 | 1.659 | 2.485 | -0.574 | 2.551 | 2.233 | Early up | | | | | | | | | | |
| B7L74_08230 | tRNA-Met | tRNA | 0.281 | -1.933 | 1.199 | 0.947 | -0.928 | -0.868 | 1.442 | -0.204 | -0.871 | 0.836 | 1.147 | | | | | | | | | | | |
| B7L74_08235 | hypothetical proteinprotein=heat resistant agglutinin | | -0.166 | -1.815 | 0.053 | 0.809 | 0.202 | -0.140 | 1.060 | -0.251 | -2.257 | 0.940 | 1.203 | Down in P51 | | | | | | | | | | |
| B7L74_08240 | ATP-dependent zinc protease | | 2.419 | -0.412 | 2.360 | 1.762 | 0.609 | 1.885 | 2.724 | 2.072 | 1.705 | 3.372 | 2.564 | Early up | | | | | | | | | | |

| | | | | | | | | | | | | | | | | | | | | | | | |
|---|---|---|---|---|---|---|---|---|---|---|---|---|---|---|---|---|---|---|---|---|---|---|---|
| B7L74_08245 | 30S ribosomal protein S6--L-glutamate ligase | | 0.302 | 1.895 | -0.630 | -0.974 | 1.377 | 4.200 | 0.260 | 3.185 | 2.357 | -0.470 | 0.011 | | | | | | | | | | |
| B7L74_08250 | succinylglutamate desuccinylase | | 4.440 | -0.906 | 1.192 | 1.412 | 4.732 | 6.398 | 3.646 | 1.976 | 4.721 | 3.998 | 3.103 | Early up | | | | | | | | | |
| B7L74_08255 | mechanosensitive ion channel protein MscS | Transporter | 7.599 | 4.920 | 5.805 | 2.047 | 10.023 | 10.069 | 5.679 | 4.544 | 8.859 | 6.144 | 3.909 | Early up | | | | | | | | Yes (ion channels/mechanosensitive channel) | |
| B7L74_08260 | hypothetical protein | Effector protein | 3.664 | 0.590 | 2.229 | 1.688 | 3.178 | 4.121 | 1.839 | 0.819 | 3.578 | 1.813 | 2.108 | Early up | | Yes | | | | | | | |
| B7L74_08265 | hypothetical protein | | 7.993 | 2.519 | 3.531 | 1.291 | 8.126 | 9.037 | 4.692 | 5.117 | 7.253 | 3.370 | 1.237 | Continuous up | | | | | | | | | |
| B7L74_08270 | ankryin | Effector protein | 2.621 | 0.503 | 1.252 | -0.324 | 3.983 | 4.703 | 1.984 | 3.291 | 3.763 | 1.833 | 1.694 | Early up | | Yes | | | | | | | |
| B7L74_08275 | hypothetical protein | | 1.020 | 0.765 | 2.672 | -0.174 | 2.226 | 2.671 | 1.961 | 2.806 | 2.532 | 0.563 | 0.884 | Early up | | | | | | | | | |
| B7L74_08280 | hypothetical protein | | 0.164 | 1.229 | 0.872 | -1.095 | 1.515 | 2.179 | 1.957 | 3.222 | 3.763 | 2.519 | 1.792 | Late up | | | | | | | | | |
| B7L74_08285 | hypothetical protein | | 8.764 | 2.907 | 3.784 | 2.846 | 6.303 | 9.169 | 5.287 | 5.534 | 6.260 | 4.633 | 3.675 | Early up | | | | | | | | | |
| B7L74_08290 | hypothetical protein | Effector protein | 9.261 | 5.986 | 5.221 | 2.730 | 11.649 | 10.660 | 6.905 | 4.487 | 9.360 | 6.312 | 3.195 | Early up | | Yes | | | | | | | |
| B7L74_08295 | hypothetical protein | | 8.558 | 2.965 | 3.147 | 0.734 | 9.463 | 10.171 | 3.935 | 3.997 | 8.095 | 4.167 | 2.003 | | | | | | | | | | |
| B7L74_08300 | hypothetical protein | | 1.776 | 1.619 | 0.443 | 1.410 | 3.263 | 3.986 | 1.979 | 3.751 | 4.156 | 1.545 | 0.793 | Continuous up | | | | | | | | | |
| B7L74_08305 | hypothetical protein | Peptidases and inhibitors | -0.260 | 0.122 | 1.949 | 1.428 | -1.048 | 0.974 | 3.365 | 1.753 | 2.496 | 2.072 | 2.864 | Late up | | | | | | | | | |
| B7L74_08310 | hypothetical protein | | 0.530 | 1.330 | 1.756 | 1.373 | 1.985 | 1.560 | 2.959 | 2.821 | 3.526 | 3.546 | 4.107 | Late up | | | | | | | | | |
| B7L74_08315 | hypothetical protein | | 2.595 | 1.531 | 4.489 | 3.278 | 3.480 | 2.983 | 3.880 | 1.506 | 4.773 | 0.186 | 5.631 | | | | | | | | | | |
| B7L74_08320 | hypothetical protein | | 0.713 | 0.683 | -0.833 | -1.696 | -1.654 | -1.284 | -1.631 | -1.351 | -1.637 | -1.391 | -1.383 | | | | | | | | | | |
| B7L74_08325 | Type IV secretion protein IcmB | T4BSS | 1.189 | -0.492 | -1.351 | -4.465 | 2.024 | 2.023 | -1.206 | -2.867 | 1.093 | -0.821 | -2.239 | | Yes | | | | | | | | |
| B7L74_08330 | HNH endonuclease, DotN/IcmJ | T4BSS | 0.572 | 0.123 | -1.472 | -2.593 | -1.417 | -2.053 | -1.847 | -2.761 | -4.057 | -1.348 | -1.408 | Continuous down | Yes | | | | | | | | |
| B7L74_08335 | Type IV secretion protein DotP/IcmD | T4BSS | 0.872 | 0.467 | -2.778 | -3.025 | -1.097 | -1.321 | -2.898 | -3.551 | -4.676 | -2.396 | -2.768 | Continuous down | Yes | | | | | | | | |
| B7L74_08340 | Type IV secretion protein DotE/IcmC | T4BSS | 0.056 | 0.387 | -6.228 | -5.211 | -2.317 | -2.662 | -4.005 | -3.302 | -6.244 | -4.093 | -4.395 | Early down | Yes | | | | | | | | |
| B7L74_08345 | Type IV secretion protein DotF/IcmG | T4BSS | -0.415 | 0.229 | -14.447 | -5.820 | -1.907 | -2.457 | -5.260 | -4.222 | -7.113 | -5.562 | -5.239 | Early down | Yes | | | | | | | | |
| B7L74_08350 | Type IV secretion protein DotG/IcmE | T4BSS | -0.010 | 0.167 | -5.508 | -6.199 | -2.136 | -2.744 | -4.678 | -3.817 | -6.639 | -4.805 | -4.523 | Early down | Yes | | | | | | | | |
| B7L74_08355 | Type IV secretion protein IcmK | T4BSS | 0.545 | 0.837 | 1.966 | -0.296 | 2.640 | 2.389 | 1.911 | 1.487 | 2.547 | -0.077 | 0.656 | | Yes | | | | | | | | |
| B7L74_08360 | Type IV secretion protein DotJ/IcmL | T4BSS | -0.925 | 0.266 | -9.768 | -6.887 | -2.689 | -2.589 | -5.303 | -4.431 | -5.828 | -5.927 | -5.696 | Early down | Yes | | | | | | | | |
| B7L74_08365 | Type IV secretion protein DotI/IcmL | T4BSS | -1.207 | -0.130 | -9.081 | -6.534 | -2.835 | -3.274 | -5.763 | -4.262 | -5.632 | -5.743 | -5.596 | Early down | Yes | | | | | | | | |
| B7L74_08370 | hypothetical protein, DotK/IcmN | T4BSS | -0.477 | 0.428 | -7.975 | -6.370 | -2.517 | -2.794 | -5.111 | -4.329 | -9.607 | -5.882 | -5.132 | Early down | Yes | | | | | | | | |
| B7L74_08375 | phosphoesterase, DotL/IcmO | T4BSS | -1.088 | -0.091 | -6.806 | -6.019 | -3.005 | -2.871 | -5.591 | -4.954 | -6.019 | -5.313 | -4.971 | Early down | Yes | | | | | | | | |
| B7L74_08380 | phosphoesterase DotM/IcmP | T4BSS | -1.445 | -0.978 | -5.799 | -5.320 | -3.883 | -5.145 | -5.390 | -4.304 | -5.873 | -5.447 | -5.064 | Early down | Yes | | | | | | | | |
| B7L74_08385 | Dot/Icm secretion system protein IcmQ | T4BSS | 0.134 | 0.316 | -5.978 | -4.577 | -2.415 | -2.842 | -4.596 | -4.962 | -5.186 | -4.014 | -3.389 | Early down | Yes | | | | | | | | |
| B7L74_08390 | hypothetical protein | Effector protein | 0.044 | 0.838 | -0.034 | -0.885 | 2.645 | 1.621 | 0.835 | 1.488 | 1.883 | -0.184 | -0.126 | | | Yes | | | | | | | |
| B7L74_08395 | hypothetical protein | | 1.173 | 1.042 | -0.644 | -2.545 | 0.405 | 0.354 | 0.184 | -1.163 | -1.982 | -2.887 | -2.252 | | | | | | | | | | |
| B7L74_08400 | hypothetical protein | Effector protein | -0.605 | -0.159 | -2.461 | -1.239 | -1.125 | -0.897 | -1.850 | -1.862 | -3.374 | -1.584 | -2.176 | Early down | | Yes | | | | | | | |
| B7L74_08405 | ATPase | ATP | -1.109 | 0.567 | -1.493 | -1.316 | -0.816 | -0.933 | -1.816 | -1.236 | -1.343 | -0.110 | -1.205 | | | | | | | | | | |
| B7L74_08410 | hypothetical protein | Effector protein | -0.784 | 0.112 | -1.372 | -1.932 | -1.491 | -0.421 | -1.465 | -1.323 | -1.374 | -0.903 | -0.925 | | | Yes | | | | | | | |
| B7L74_08415 | hypothetical protein | | 0.803 | -0.190 | 1.694 | -0.252 | 0.120 | 0.629 | 0.482 | -0.803 | -0.240 | 1.174 | 1.492 | | | | | | | | | | |
| B7L74_08420 | IS110 family transposase | | 0.282 | -0.233 | 0.853 | 0.175 | -0.573 | -1.183 | 0.844 | 0.759 | 0.584 | 1.547 | 2.481 | | | | | | | | | | |
| B7L74_08425 | hypothetical protein | | -0.560 | -0.006 | -2.125 | 0.108 | -0.454 | -0.721 | -0.707 | -1.927 | -2.051 | -0.438 | -0.718 | | | | | | | | | | |
| B7L74_08430 | phosphoesterase, IcmT | T4BSS | 0.441 | 0.114 | -0.468 | -0.125 | -0.818 | -1.370 | -0.648 | -2.390 | -2.933 | -0.346 | -0.474 | Late down | Yes | | | | | | | | |
| B7L74_08435 | Type IV secretion protein IcmS | T4BSS | 0.082 | 0.846 | -0.034 | -1.769 | 2.694 | 2.759 | 3.455 | 2.336 | 2.373 | -1.257 | -0.898 | | Yes | | | | | | | | |
| B7L74_08440 | LuxR family transcriptional regulator, DotD | T4BSS | 0.112 | -0.309 | -4.844 | -3.141 | -2.568 | -2.838 | -3.598 | -4.158 | -5.372 | -2.720 | -2.694 | Early down | Yes | | | | | | | | |
| B7L74_08445 | Type IV secretion system protein DotC | T4BSS | -0.578 | -0.506 | -5.697 | -4.412 | -3.085 | -3.069 | -3.954 | -4.345 | -5.380 | -3.911 | -3.398 | Early down | Yes | | | | | | | | |
| B7L74_08450 | Dot/Icm secretion system ATPase DotB | T4BSS | 0.219 | -0.023 | -1.708 | -1.914 | -1.583 | -1.908 | -1.310 | -2.481 | -3.072 | -1.493 | -1.616 | Continuous down | Yes | | | | | | | | |
| B7L74_08455 | transporter | Transporter | 0.543 | -0.111 | -0.277 | -0.763 | 0.380 | 0.916 | -0.317 | -1.771 | -0.226 | 0.804 | 1.545 | | | | | | | | | Yes (Glycine betaine) | |
| B7L74_08460 | hypothetical protein | | 1.346 | 1.748 | -3.324 | -4.544 | -0.648 | -1.669 | -2.841 | -0.862 | -0.741 | -1.427 | -2.592 | | | | | | | | | | |
| B7L74_08465 | Type IV secretion protein DotA | T4BSS | -7.933 | -7.351 | -7.833 | -3.990 | -6.044 | -5.641 | -6.286 | -4.322 | -3.536 | -2.824 | -1.169 | Early down | Yes | | | | | | | | |
| B7L74_08470 | Type IV secretion protein IcmV | T4BSS | 0.053 | 0.840 | -0.034 | -1.795 | 1.973 | 1.622 | 0.460 | 0.016 | 2.560 | -1.273 | -0.907 | | Yes | | | | | | | | |
| B7L74_08475 | phosphoesterase, IcmW | T4BSS | 0.609 | 0.641 | -3.667 | -5.416 | -1.820 | -1.861 | -4.317 | -4.193 | -4.050 | -4.192 | -3.830 | Early down | Yes | | | | | | | | |
| B7L74_08480 | hypothetical protein, | | -0.529 | 0.263 | -0.610 | -1.993 | 4.122 | 3.022 | 0.805 | 2.572 | 3.402 | -1.852 | -1.485 | Early up | | | | | | | | | |
| B7L74_08485 | phosphoesterase, IcmX | T4BSS | -1.007 | 0.284 | -6.130 | -6.118 | -2.387 | -2.218 | -5.285 | -4.300 | -4.628 | -5.261 | -5.068 | Early down | Yes | | | | | | | | |
| B7L74_08490 | hypothetical protein | | 1.011 | 1.166 | 1.006 | -0.306 | -0.259 | -0.578 | 0.021 | -1.014 | -3.162 | 0.143 | 0.492 | Down in P51 | | | | | | | | | |
| B7L74_08495 | virulence factor | | 2.576 | 0.805 | 3.558 | 4.279 | 3.582 | 0.192 | 0.771 | 2.700 | 3.364 | 2.795 | 1.987 | | | | | | | | | | |
| B7L74_08500 | bifunctional heptose 7-phosphate kinase/heptose 1-pho | Lipopolysaccharide biosynthesis | 0.566 | 0.908 | 0.460 | 1.964 | -0.057 | 0.663 | 0.830 | 0.828 | -0.737 | 0.916 | 1.667 | | | | | | | | | | Yes |
| B7L74_08505 | hypothetical protein | | -2.353 | -0.231 | -2.478 | 0.652 | 1.388 | -0.816 | -0.231 | 1.813 | 0.185 | 0.162 | 0.009 | | | | | | | | | | |
| B7L74_08510 | alpha-L-glycero-D-manno-heptose beta-1%2C4-glucos | O-antigen synthesis | 0.621 | 0.393 | 0.364 | 0.011 | 0.102 | 0.231 | 0.470 | -0.741 | -1.610 | 0.476 | 0.402 | | | | | | | | | | |
| B7L74_08515 | hypothetical protein | | -0.133 | 0.130 | 0.158 | 0.577 | -1.246 | 0.333 | 0.356 | -0.335 | -1.157 | 0.600 | 0.610 | | | | | | | | | | |
| B7L74_08520 | O-antigen ligase | Lipopolysaccharide biosynthesis | 0.062 | 0.160 | 0.294 | 1.893 | -0.734 | 0.334 | 0.204 | 0.210 | 0.993 | 1.679 | 2.304 | | | | | | | | | | Yes |
| B7L74_08525 | hypothetical cytosolic protein | | 1.663 | 2.350 | 1.882 | 2.389 | 4.572 | 2.683 | 2.647 | 2.042 | 3.806 | 0.860 | 1.584 | Early up | | | | | | | | | |
| B7L74_08530 | lipopolysaccharide heptosyltransferase II | Lipopolysaccharide biosynthesis | -0.016 | -0.698 | -0.590 | 0.951 | -1.351 | -1.602 | -0.182 | -1.553 | -3.005 | -0.298 | -1.110 | | | | | | | | | | Yes |
| B7L74_08535 | zinc-finger domain-containing protein | | -0.810 | -0.211 | 1.020 | 0.858 | -0.939 | -1.131 | 0.457 | -0.989 | -0.567 | 0.606 | 0.925 | | | | | | | | | | |
| B7L74_08540 | hypothetical protein | | 0.090 | 2.989 | -0.035 | 0.349 | 2.737 | 1.627 | 1.471 | 4.552 | 2.628 | 2.289 | 2.822 | | | | | | | | | | |
| B7L74_08545 | CBS domain-containing protein | Uncharacter ized | 0.550 | -0.378 | 0.565 | 1.501 | -0.652 | -0.203 | 0.447 | -0.324 | -0.625 | 1.167 | 0.925 | | | | | | | | | | |
| B7L74_08550 | hypothetical protein | Effector protein | 1.933 | 0.848 | -0.035 | 0.278 | 5.883 | 5.143 | 3.280 | 6.533 | 6.125 | 0.469 | 0.052 | | | Yes | | | | | | | |
| B7L74_08555 | hypothetical protein | | -0.469 | 1.278 | 2.385 | 0.974 | 2.176 | 0.645 | 2.284 | 1.943 | 2.370 | 1.705 | 0.932 | | | | | | | | | | |
| B7L74_08560 | hypothetical protein | | 0.608 | 0.516 | 1.014 | 2.040 | 0.767 | 1.837 | 1.088 | 1.374 | 1.352 | 1.425 | 1.224 | | | | | | | | | | |
| B7L74_08565 | hypothetical protein | | -0.392 | 0.089 | 0.183 | 0.054 | 1.189 | 1.793 | 0.416 | -1.549 | -0.808 | -0.099 | -0.288 | | | | | | | | | | |
| B7L74_08570 | RNA polymerase sigma factor RpoS | Biofilm formation - Vibrio cholera | 3.772 | 1.453 | 3.120 | 2.479 | 5.180 | 5.448 | 2.870 | -0.326 | 5.017 | 4.447 | 4.217 | Early up | | | | | | | | | |
| B7L74_08575 | peptidase | Structural proteins | 3.816 | 0.197 | 3.247 | 2.568 | 3.728 | 5.339 | 2.233 | -0.806 | 3.831 | 3.688 | 3.913 | Early up | | | | | | | | | |
| B7L74_08580 | 5'-nucleotidase SurE | | 0.074 | -0.724 | -0.058 | 0.271 | -1.152 | -0.404 | 0.375 | -1.435 | -1.856 | 0.370 | 0.548 | | | | | | | | | | |
| B7L74_08585 | cell division protein FtsB | Cell division | -0.306 | -0.392 | -0.632 | -0.536 | -2.014 | -0.624 | 0.016 | -1.572 | -1.717 | 0.234 | 0.150 | Early down | | | | | | | | | |
| B7L74_08590 | enolase | | 0.205 | 0.105 | 0.445 | -0.492 | -0.332 | -0.582 | 0.056 | -1.121 | -1.318 | 0.623 | 0.241 | | | | | Yes | | | | | |
| B7L74_08595 | 3-deoxy-8-phosphooctulonate synthase | Lipopolysaccharide biosynthesis | 0.509 | -0.913 | 0.236 | 0.939 | -0.278 | -0.356 | 0.503 | -0.711 | -1.249 | 0.877 | 1.190 | | | | | | | | | | Yes |
| B7L74_08600 | DUF1658 domain-containing protein | | -0.201 | -0.557 | -0.689 | 0.392 | 0.172 | 0.405 | -0.272 | -1.153 | -2.415 | -0.699 | -0.317 | | | | | | | | | | |
| B7L74_08605 | hypothetical cytosolic protein | Effector protein | 0.831 | 1.040 | 0.666 | 0.301 | 1.130 | 0.422 | 0.648 | 0.771 | 1.038 | 0.617 | 0.735 | | | Yes | | | | | | | |
| B7L74_08610 | hemerythrin | Effector protein | 1.265 | 1.376 | 1.269 | -0.260 | 1.718 | 0.433 | 1.728 | 1.602 | 2.457 | 1.671 | 1.880 | | | Yes | | | | | | | |
| B7L74_08615 | diamine N-acetyltransferase | | -0.544 | 1.168 | 0.833 | 0.546 | 0.746 | 0.163 | -0.408 | 0.622 | 1.064 | -1.121 | -1.354 | | | | | | | | | | |
| B7L74_08620 | hypothetical protein | | -0.917 | -0.033 | -5.879 | 2.276 | 1.100 | 0.252 | -1.261 | 0.571 | 0.788 | -0.531 | -0.833 | | | | | | | | | | |
| B7L74_08625 | hypothetical protein | | 1.060 | 1.740 | 2.477 | -0.028 | 0.885 | 0.369 | 0.777 | 1.385 | 1.651 | 0.454 | 0.757 | Early up | | | | | | | | | |
| B7L74_08630 | hypothetical exported protein | | 0.889 | 1.993 | 3.153 | -0.644 | 1.759 | 1.415 | 1.228 | 1.305 | 2.264 | 1.098 | 1.350 | Early up | | | | | | | | | |
| B7L74_08635 | CTP synthetase (gene=pyrG) | | 1.066 | 0.498 | -0.104 | 0.999 | 0.770 | 0.523 | 0.494 | -0.242 | -1.087 | 0.519 | 0.929 | | | | | | | | | | |
| B7L74_08640 | hypothetical protein | | 6.256 | 2.914 | 4.100 | 2.011 | 7.673 | 7.612 | 5.279 | 2.974 | 6.999 | 4.560 | 4.878 | Continuous up | | | | | | | | | |
| B7L74_08645 | hypothetical protein | | 5.072 | 1.866 | 0.976 | -1.485 | 3.907 | 5.787 | 1.255 | 2.514 | 4.619 | 0.409 | -0.561 | | | | | | | | | | |
| B7L74_08650 | hypothetical protein | Effector protein | -0.568 | 1.398 | -0.053 | -1.241 | 2.289 | 1.928 | 0.626 | 0.289 | 1.027 | -0.152 | -0.466 | | | Yes | | | | | | | |
| B7L74_08655 | hypothetical protein | Effector protein | -0.730 | 0.320 | -0.628 | -1.525 | 0.322 | -0.602 | 0.122 | -0.803 | 0.018 | -1.749 | -0.761 | | | Yes | | | | | | | |
| B7L74_08660 | hypothetical protein | | 0.598 | -0.246 | -0.560 | 0.273 | 0.289 | 0.828 | 0.604 | 1.124 | 1.133 | -0.455 | 0.267 | | | | | | | | | | |
| B7L74_08665 | hypothetical protein | | 0.062 | 0.842 | -0.034 | -1.787 | 2.511 | 0.975 | -0.071 | 1.491 | 1.879 | -1.268 | -0.904 | | | | | | | | | | |
| B7L74_08670 | hypothetical protein | | -1.326 | -0.214 | -0.930 | 0.973 | -0.600 | -0.303 | -0.484 | 1.770 | 0.285 | -0.640 | -0.905 | | | | | | | | | | |
| B7L74_08675 | dCTP deaminase | | -0.215 | 0.150 | -0.670 | 1.881 | -0.523 | -0.736 | -0.498 | -0.035 | -0.727 | -0.078 | 0.241 | | | | | | | | | | |
| B7L74_08680 | sodium:proton antiporter | Transporter | 1.197 | -1.234 | -0.804 | -2.968 | 2.270 | -0.246 | 0.695 | 0.510 | -0.371 | 0.589 | 1.726 | | | | | | | | | | |
| B7L74_08685 | hypothetical protein | | -0.938 | -0.226 | -0.838 | 0.156 | -0.207 | -0.650 | -0.921 | -1.425 | -1.856 | 0.507 | -0.130 | | | | | | | | | | |
| B7L74_08690 | transcriptional regulator | | -0.219 | -0.191 | 2.473 | 1.584 | 0.530 | 0.582 | 0.524 | -0.699 | 0.209 | 2.589 | 2.369 | Late up | | | | | | | | | |
| B7L74_08695 | hypothetical cytosolic protein | | -1.019 | -5.376 | -6.279 | 0.407 | 1.825 | 0.450 | -0.068 | 0.514 | 1.473 | 3.747 | 1.025 | | | | | | | | | | |
| B7L74_08700 | hypothetical protein | | 4.421 | 5.679 | -0.210 | 0.434 | 1.530 | 1.761 | 1.053 | 1.007 | 2.235 | 0.783 | 1.702 | | | | | | | | | | |
| B7L74_08705 | hypothetical protein | | 0.825 | 1.388 | 1.498 | -0.460 | 1.669 | 0.963 | 0.887 | 1.868 | 1.592 | 1.361 | 1.286 | | | | | | | | | | |
| B7L74_08710 | hypothetical protein | | 1.040 | 1.304 | 2.848 | 0.410 | 2.004 | 0.825 | 1.925 | 2.080 | 2.776 | 1.349 | 2.152 | Early up | | | | | | | | | |
| B7L74_08715 | methionine--tRNA ligase | | -0.875 | -0.252 | -1.321 | 0.742 | -6.825 | -1.187 | -1.216 | -0.022 | -1.385 | -0.356 | -0.503 | | | | | | | | | | |
| B7L74_08720 | 4Fe-4S ferredoxin | | -0.539 | -1.055 | -0.205 | -0.291 | -0.109 | 0.250 | -0.139 | -0.222 | 0.555 | -1.033 | -0.287 | | | | | | | | | | |
| B7L74_08725 | endonuclease III | | -1.012 | 0.584 | -0.452 | 0.820 | 0.650 | 1.020 | 0.129 | 0.051 | -0.208 | -0.346 | -0.068 | | | | | | | | | | |
| B7L74_08730 | ATPase | ATP | 2.790 | 0.804 | 3.558 | 5.108 | 4.578 | 4.194 | 3.875 | 4.463 | 4.481 | 5.781 | 4.968 | | | | | | | | | | |
| B7L74_08735 | hypothetical protein | | 0.687 | -0.688 | 1.276 | 1.370 | -1.403 | 0.566 | 1.370 | 1.839 | 1.534 | 1.371 | 1.567 | | | | | | | | | | |
| B7L74_08740 | IS110 family transposase | | 0.500 | 0.232 | 1.757 | -0.659 | -0.221 | -0.133 | 0.540 | -0.157 | -1.350 | 1.256 | 1.236 | | | | | | | | | | |
| B7L74_08745 | hypothetical protein | | -0.572 | -1.595 | -0.035 | -1.668 | 0.294 | -0.816 | -0.247 | -0.947 | 1.209 | -1.201 | -0.869 | | | | | | | | | | |
| B7L74_08750 | hypothetical membrane spanning protein | | -0.388 | -0.400 | -0.626 | 0.060 | -0.570 | 0.026 | 0.157 | 0.084 | -0.716 | 0.629 | 0.522 | | | | | | | | | | |
| B7L74_08755 | hypothetical cytosolic protein | | 0.558 | -0.167 | 0.966 | 1.518 | 0.973 | 1.575 | 1.251 | 1.956 | 0.976 | 1.827 | 1.344 | | | | | | | | | | |
| B7L74_08760 | dihydroorotase | | -1.226 | -0.794 | -0.929 | 0.978 | -1.318 | -0.774 | -0.763 | -1.702 | -2.777 | 0.900 | 0.955 | | | | | | | | | | |
| B7L74_08765 | ribonuclease T | | 0.589 | -1.206 | -0.464 | -1.370 | -1.006 | 0.326 | 1.736 | -1.643 | -2.367 | 0.053 | -0.389 | | | | | | | | | | |
| B7L74_08770 | hypothetical protein | | -3.647 | -0.387 | 2.220 | 2.058 | -1.083 | -5.347 | -0.721 | -1.086 | -3.313 | 0.660 | 0.124 | Early down | | | | | | | | | |
| B7L74_08775 | alkyl hydroperoxide reductase | PV detoxification | -0.540 | 2.030 | 2.321 | 2.590 | 0.844 | 0.374 | 1.336 | 2.053 | 2.241 | 0.209 | 0.112 | Early up | | | | | | | | | |
| B7L74_08780 | hypothetical protein | | -0.313 | 0.305 | -2.103 | 1.876 | 1.434 | -0.080 | -0.959 | 1.825 | -0.651 | 0.308 | 0.311 | | | | | | | | | | |
| B7L74_08785 | superoxide dismutase [Fe] | Antioxidant, PV detoxification | -1.098 | -0.945 | 0.704 | 2.836 | -0.812 | -1.402 | -0.226 | -1.146 | -2.241 | -0.659 | -0.648 | | | | | | | | | | |
| B7L74_08790 | 4-hydroxy-tetrahydrodipicolinate reductase | | 0.033 | -0.936 | -1.515 | 1.850 | -2.056 | -0.187 | 0.135 | -1.492 | -0.968 | -0.749 | -0.616 | | | | | | | | | | |
| B7L74_08795 | hypothetical protein | | -0.061 | 0.983 | 1.007 | -0.149 | 0.173 | 0.016 | 0.316 | 0.178 | -0.870 | 0.726 | 0.000 | | | | | | | | | | |
| B7L74_08800 | hypothetical protein | | 0.083 | 0.693 | 1.612 | 1.963 | 1.037 | 0.539 | 0.570 | 1.699 | 1.391 | 1.843 | 1.193 | | | | | | | | | | |
| B7L74_08805 | GIY-YIG nuclease | | -0.919 | -0.416 | -1.534 | -0.547 | -1.385 | -1.719 | -1.027 | -1.217 | -2.752 | -0.466 | -0.846 | | | | | | | | | | |
| B7L74_08810 | hypothetical protein | | 0.399 | -1.462 | -1.351 | -1.223 | -1.341 | -0.953 | -1.078 | -1.859 | -2.528 | -2.130 | -1.050 | | | | | | | | | | |
| B7L74_08815 | glycine dehydrogenase | Glyoxylate and dicarboxylate meta | -3.526 | -7.045 | -9.487 | -1.811 | -5.661 | -3.801 | -2.045 | -6.995 | -3.606 | -2.265 | -1.606 | Early down | | | | | | | | | |
| B7L74_08820 | glycine dehydrogenase | Glyoxylate and dicarboxylate meta | -0.317 | -1.635 | -1.419 | 0.099 | -1.448 | -1.332 | -0.212 | -2.248 | -2.777 | -0.291 | 0.594 | Late down | | | | | | | | | |
| B7L74_08825 | glycine cleavage system protein H | Glyoxylate and dicarboxylate meta | -0.804 | -1.086 | -1.842 | 0.460 | -2.391 | -2.035 | -1.240 | -2.973 | -3.162 | -0.813 | -0.597 | Early down | | | | | | | | | |
| B7L74_08830 | glycine cleavage system protein T | | -0.109 | -0.369 | -0.804 | 0.576 | -1.365 | -0.763 | 0.148 | -1.687 | -0.942 | 0.516 | 0.745 | | | | | | | | | | |
| B7L74_08835 | IS110 family transposase | | -0.239 | -1.101 | 2.239 | -5.742 | -1.340 | -0.882 | -0.006 | -0.502 | -1.127 | 1.115 | 1.895 | | | | | | | | | | |
| B7L74_08840 | hypothetical protein | | 1.328 | -0.584 | 3.024 | 0.097 | -1.291 | -2.401 | 2.115 | -2.532 | -1.400 | 0.921 | 1.200 | | | | | | | | | | |
| B7L74_08845 | molecular chaperone GroEL | RNA degradation | -3.803 | -5.000 | -2.898 | -1.230 | -4.207 | -3.762 | -3.130 | -5.914 | -6.405 | -3.232 | -3.092 | Continuous down | | | Yes | | | | | | |
| B7L74_08850 | co-chaperone GroES | Chaperones and folding catalysts | -4.105 | -5.423 | -2.974 | -0.744 | -4.313 | -3.882 | -3.460 | -6.157 | -7.813 | -3.635 | -3.571 | Continuous down | | | Yes | | | | | | |
| B7L74_08855 | aconitate hydratase | TCA cycle only | -1.099 | -3.007 | -3.621 | -1.690 | -1.482 | -1.010 | -2.157 | -4.233 | -2.879 | -2.153 | -1.494 | Late down | | | | | Yes | | | | |
| B7L74_08860 | hypothetical protein | | -0.511 | -2.141 | -1.450 | 0.080 | -0.907 | -0.834 | -0.546 | -2.669 | -1.170 | 0.096 | 0.981 | | | | | | | | | | |
| B7L74_08865 | AMP-binding protein | | -0.304 | 0.079 | -0.079 | -2.459 | 0.889 | 0.784 | -0.608 | -0.180 | 0.769 | -0.297 | -0.154 | | | | | | | | Yes | | |
| B7L74_08870 | thiol:disulfide interchange protein | | 1.488 | -0.088 | 0.117 | 0.158 | 1.753 | 2.269 | 0.810 | -0.660 | 1.954 | 0.590 | 0.658 | | | | Yes | | | | | | |
| B7L74_08875 | hypothetical protein | Effector protein | 4.170 | 0.555 | 1.740 | 0.502 | 4.906 | 5.551 | 2.070 | 0.700 | 3.963 | 2.179 | 1.819 | Early up | | Yes | | | | | | | |
| B7L74_08880 | acetyl-CoA carboxylase biotin carboxyl carrier protein | Effector protein | 0.114 | -1.133 | -0.773 | 0.568 | -1.685 | -1.434 | 0.426 | -1.580 | -1.446 | 0.170 | 0.413 | | | Yes | | | | | | | |
| B7L74_08885 | acetyl-CoA carboxylase biotin carboxylase subunit | Pyruvate metabolism | -0.442 | -1.473 | -2.134 | -0.420 | -2.232 | -1.709 | -0.862 | -2.097 | -1.299 | -1.202 | -0.855 | Early down | | | | | | | | | |
| B7L74_08890 | ornithine cyclodeaminase | Arginine and proline metabolism | -0.800 | -1.432 | -2.527 | -0.175 | -3.085 | -1.551 | -1.009 | -1.563 | -0.615 | -0.052 | -0.454 | Early down | | | | | | | | | |
| B7L74_08895 | nicotinamide mononucleotide transporter | Transporter | 3.266 | 3.448 | 4.136 | -1.834 | 4.138 | 4.558 | 4.507 | 1.931 | 3.659 | 2.602 | 3.113 | Early up | | | | | | | | Yes (Vitamins/thiamin) | |
| B7L74_08900 | coproporphyrinogen III oxidase | | 0.046 | -0.674 | -0.765 | -0.559 | -0.954 | -0.795 | -0.482 | -1.477 | -1.273 | -0.633 | -0.282 | | | | | | | | | | |
| B7L74_08905 | phosphoserine phosphatase | | -1.116 | -1.292 | -0.452 | 0.060 | -1.129 | -1.080 | -0.989 | -0.593 | -0.003 | -0.129 | -0.228 | | | | | | | | | | |
| B7L74_08910 | hypothetical protein | | 0.578 | -0.063 | -0.824 | 1.132 | 0.684 | 1.519 | 0.921 | 0.940 | 1.959 | 0.922 | 0.740 | | | | | | | | | Yes (Na+ symporter) | |
| B7L74_08915 | hypothetical protein | Transporter | 0.071 | 0.844 | -0.034 | -1.778 | 2.082 | 4.132 | 0.479 | 1.803 | 3.231 | -0.741 | -0.426 | Up in P21 | | | | | | | | Yes (Na+ symporter) | |
| B7L74_08920 | hypothetical protein | Transporter | 5.083 | 1.647 | 2.349 | 0.858 | 5.886 | 6.342 | 3.000 | 0.604 | 5.949 | 3.560 | 2.009 | Continuous up | | | | | | | | Yes (Na+ symporter) | |
| B7L74_08925 | D-3-phosphoglycerate dehydrogenase | | -0.777 | 0.081 | 0.558 | 0.232 | 2.555 | 0.589 | 0.358 | 1.452 | 1.316 | -0.735 | -1.082 | | | | | | | | | | |
| B7L74_08930 | hypothetical membrane spanning protein | | -0.081 | -0.583 | -0.821 | -0.769 | -0.789 | -0.735 | -0.611 | -1.342 | -0.789 | -0.179 | -0.093 | | | | | | | | | | |
| B7L74_08935 | DnaA regulatory inactivator Hda | DNA replication proteins | -1.406 | -1.686 | -3.291 | -1.326 | -2.411 | -2.281 | -1.369 | -2.662 | -2.531 | -1.190 | -1.576 | Early down | | | | | | | | | |
| B7L74_08940 | hypothetical exported protein | | -0.365 | -0.650 | 0.335 | 0.019 | -1.102 | -1.144 | -0.168 | -1.711 | -2.151 | 0.053 | -0.014 | Late down | | | | | | | | | |
| B7L74_08945 | phosphoribosylformylglycinamidine cyclo-ligase | Purine metabolism | 0.025 | -1.501 | -0.352 | 0.358 | -1.057 | -1.814 | -0.236 | -1.874 | -2.222 | 0.115 | 0.404 | Early down | | | | | | | | | |
| B7L74_08950 | phosphoribosylglycinamide formyltransferase | | 0.815 | 3.727 | -0.034 | -1.788 | 3.838 | 2.889 | 1.072 | 2.733 | 4.878 | -1.268 | -0.904 | | | | | | | | | | |
| B7L74_08955 | integration host factor subunit beta | DNA replication proteins | -0.155 | -0.023 | 1.688 | 2.061 | -0.181 | -0.541 | 0.013 | 0.233 | -1.380 | 0.384 | 0.069 | Up in P12 | | | | | | | | | |
| B7L74_08960 | rRNA (cytidine-2'-O-)-methyltransferase | | 0.090 | 0.847 | -0.035 | -0.096 | 2.736 | 1.627 | 0.527 | 1.496 | 3.748 | -0.206 | 0.843 | | | | | | | | | | |
| B7L74_08965 | penicillin-binding protein activator | | 0.225 | -0.849 | 0.055 | 0.178 | -1.043 | -1.259 | 0.099 | -0.507 | -1.244 | 0.479 | 0.467 | | | | | | | | | | |
| B7L74_08970 | YraN family protein | | -0.617 | -1.184 | -0.709 | 0.327 | -1.286 | 0.989 | 0.028 | 1.435 | 1.223 | 0.804 | 0.825 | | | | | | | | | | |
| B7L74_08975 | phosphoheptose isomerase | Lipopolysaccharide biosynthesis | -0.075 | -0.290 | -0.161 | 1.101 | -2.153 | -0.905 | 0.523 | -0.651 | -0.982 | 0.472 | 0.635 | | | | | | | | | | Yes |
| B7L74_08980 | phospholipid-binding protein | | -0.400 | -0.950 | -0.825 | 1.701 | 1.495 | 1.410 | 1.780 | 1.842 | 1.375 | -1.921 | -1.769 | | | | | | | | | | |
| B7L74_08985 | site-2 protease family protein, Membrane endopeptidas | Membrane | 0.166 | 0.152 | 1.074 | 1.205 | 0.241 | -0.373 | -0.516 | -0.263 | -0.863 | 0.263 | -0.003 | | | | | | | | | | |
| B7L74_08990 | stringent starvation protein B | Starvation, Transcription machiner | -0.728 | -0.644 | -1.778 | 0.151 | -1.399 | -1.660 | -1.391 | -1.938 | -2.349 | -1.503 | -1.872 | Early down | | | | | | | | | |
| B7L74_08995 | stringent starvation protein A (gene=sspA) | Starvation, Transcription machiner | -0.300 | -0.158 | -3.236 | -0.340 | -1.814 | -1.889 | -1.523 | -2.309 | -2.565 | -1.571 | -1.876 | Early down | | | | | | | | | |
| B7L74_09000 | 30S ribosomal protein S9 | Ribosomal proteins | 0.447 | 0.065 | -0.126 | 0.551 | -0.478 | -0.788 | -0.479 | -1.980 | -2.166 | -0.820 | -0.642 | Late down | | | | | | | | | |
| B7L74_09005 | 50S ribosomal protein L13 | Ribosomal proteins | -0.106 | 0.046 | -1.154 | 1.250 | -0.909 | -0.652 | -0.780 | -0.987 | -2.465 | -0.756 | -0.392 | Down in P51 | | | | | | | | | |
| B7L74_09010 | integration host factor | | -0.312 | -1.210 | 0.601 | 1.433 | -1.896 | -2.326 | 0.428 | -2.488 | -2.765 | 1.626 | 0.475 | Early down | | | | | | | | | |
| B7L74_09015 | hypothetical protein(cig57 in Table 1 of RightonQ pap | Effector protein | -0.738 | 0.062 | -3.881 | -2.795 | -1.921 | -2.597 | -3.302 | -3.141 | -3.603 | -2.886 | -2.989 | Early down | | Yes | | | | | | | |
| B7L74_09020 | hypothetical protein | Effector protein | -0.725 | 0.236 | -3.594 | -4.123 | -1.952 | -2.960 | -3.978 | -3.613 | -3.820 | -2.916 | -3.142 | Early down | | Yes | | | | | | | |
| B7L74_09025 | hypothetical cytosolic protein | | -0.506 | -0.557 | 1.989 | 1.493 | -1.520 | -0.743 | 0.832 | -1.079 | -0.030 | 0.467 | 1.083 | | | | | | | | | | |
| B7L74_09030 | hypothetical protein | Effector protein | 0.975 | -0.282 | 1.669 | 2.256 | -0.201 | -0.713 | 1.772 | -0.193 | -0.256 | 1.608 | 2.250 | Early up | | Yes | | | | | | | |
| B7L74_09035 | hypothetical protein | | 0.865 | -0.253 | 0.755 | -1.312 | 0.425 | 1.227 | 0.991 | 1.656 | 1.840 | 0.923 | 0.482 | | | | | | | | | | |
| B7L74_09040 | Bcr/CflA family drug resistance efflux transporter | Transporter | 0.439 | -0.234 | 0.517 | 0.883 | -0.731 | -0.238 | 0.597 | 0.407 | 0.636 | 1.490 | 1.425 | | | | | | | | | Yes (drug resistance) | |
| B7L74_09045 | hypothetical protein/AnkM | Effector protein | -0.222 | -0.362 | 0.281 | 0.556 | -0.179 | -0.758 | -0.203 | -1.329 | -0.683 | 1.096 | 1.467 | | | Yes | | | | | | | |
| B7L74_09050 | hypothetical protein | | 0.074 | 0.009 | 2.031 | 3.168 | 0.529 | -0.491 | 0.553 | -0.053 | 1.909 | 2.290 | 3.086 | Early up | | | | | | | | | |
| B7L74_09055 | IS110 family transposase | | 0.425 | 0.307 | 1.878 | -0.652 | -0.210 | -0.181 | 0.454 | -0.036 | -1.251 | 1.231 | 1.132 | | | | | | | | | | |
| B7L74_09060 | hypothetical protein | | -1.337 | 0.777 | -1.462 | -0.303 | 1.309 | 1.162 | 0.080 | 0.069 | 1.201 | -0.159 | 0.158 | | | | | | | | | | |
| B7L74_09065 | DUF1658 domain-containing protein | | -0.021 | 2.863 | 4.432 | 2.434 | 3.999 | 1.154 | 2.558 | 3.796 | 3.387 | 3.551 | 3.262 | | | | | | | | | | |
| B7L74_09070 | hybrid sensor histidine kinase/response regulator | Intracellular growth. | 0.197 | -0.594 | 0.252 | -0.885 | -0.831 | -0.608 | 0.637 | 0.032 | -0.734 | 0.697 | 0.393 | | | | | | | | | | |
| B7L74_09075 | FAD-dependent oxidoreductase | | 0.444 | -0.325 | 1.000 | -0.247 | -0.343 | -0.241 | 0.171 | 0.577 | 0.410 | 0.314 | 0.236 | | | | | | | | | | |
| B7L74_09080 | ATPase | ATP | 0.230 | -0.729 | -0.456 | 0.711 | -1.221 | 0.463 | 0.957 | 1.460 | 1.192 | 0.621 | 0.842 | | | | | | | | | | |
| B7L74_09085 | hypothetical protein | Hypothetical | 0.171 | -0.937 | 1.324 | 2.979 | 0.296 | -1.622 | 1.319 | -0.145 | -0.891 | 1.563 | 0.839 | Up in P12 | | | | | | | | | |
| B7L74_09090 | hypothetical protein | | -1.396 | -0.638 | -1.520 | -0.266 | 2.127 | 2.944 | 0.667 | 0.010 | 1.142 | -1.815 | -0.267 | | | | | | | | | | |
| B7L74_09095 | hypothetical protein | | -0.511 | 1.547 | 0.168 | -0.058 | 0.402 | 1.050 | 0.009 | 0.529 | -1.263 | -0.386 | -0.658 | | | | | | | | | | |
| B7L74_09100 | ferrous iron transporter B | Transporter | 4.027 | 5.864 | 6.278 | 5.117 | 5.815 | 6.482 | 5.692 | 5.755 | 3.950 | 4.665 | 4.829 | Early up | | | | | | | | Yes (Ions/iron) | |
| B7L74_09105 | ferrous iron transport protein A | | -0.409 | 1.714 | 2.530 | 1.966 | 0.622 | 1.055 | 1.176 | 0.621 | -0.656 | 1.950 | 1.613 | | | | | | | | | Yes (Ions/iron) | |
| B7L74_09110 | hypothetical exported protein | | -0.731 | 0.554 | 0.299 | -1.004 | 0.038 | -0.226 | -0.076 | -0.027 | -0.188 | -0.398 | -0.405 | | | | | | | | | | |
| B7L74_09115 | alpha/beta hydrolase | Effector protein | -0.237 | 0.712 | -0.168 | -0.094 | -0.566 | 0.094 | -0.174 | 0.271 | -0.953 | -0.332 | 0.021 | | | Yes | | | | | | | |
| B7L74_09120 | ABC transporter ATP-binding protein | Transporter | -1.636 | 0.728 | -1.055 | 0.654 | -0.398 | 0.327 | -0.969 | -0.034 | -1.017 | -1.285 | -0.740 | | | | | | | | | Yes (Ions/Nitrate/taurine) | |
| B7L74_09125 | ABC transporter permease | Transporter | 0.155 | 0.871 | -0.268 | -0.460 | 0.351 | 1.370 | -0.134 | 0.204 | -0.181 | 0.012 | -0.023 | | | | | | | | | Yes (Ions/Nitrate/taurine) | |
| B7L74_09130 | GTP-binding protein | Chromosome and associated prote | 0.415 | 1.205 | 0.751 | 3.347 | 0.406 | 0.237 | 0.619 | 0.472 | 0.835 | 1.449 | 1.603 | Up in P12 | | | | | | | | | |
| B7L74_09135 | hypothetical protein | | 0.273 | 1.128 | 0.789 | 0.909 | 0.598 | 0.110 | -0.659 | 0.859 | 0.572 | 1.009 | 0.822 | | | | | | | | | | |
| B7L74_09140 | hypothetical protein | | -0.069 | 1.750 | -1.519 | 0.366 | 0.304 | 0.503 | -0.444 | 0.701 | -0.418 | 0.764 | -0.398 | | | | | | | | | | |
| B7L74_09145 | hypothetical protein | | -0.159 | 0.982 | -0.699 | 0.835 | 0.254 | -0.680 | -0.529 | -0.643 | -0.653 | 0.737 | 0.215 | | | | | | | | | | |
| B7L74_09150 | fructose-1%2C6-bisphosphate aldolase | Glycolysis / Gluconeogenesis | -0.547 | 0.335 | -0.914 | 0.986 | -0.617 | -0.521 | -0.112 | -1.202 | -2.703 | -0.271 | -0.065 | Down in P51 | | | | Yes | | | | | |
| B7L74_09155 | hypothetical protein | Effector protein | 1.048 | 0.287 | 0.219 | -0.394 | 0.905 | 1.971 | -0.506 | -0.752 | 0.383 | 0.224 | 0.071 | | | Yes | | | | | | | |
| B7L74_09160 | pyruvate kinase | | -1.849 | -0.294 | -1.294 | -1.947 | -1.217 | -0.782 | -1.513 | -1.012 | -0.462 | -1.912 | -1.514 | | | | | Yes | | | | | |
| B7L74_09165 | phosphoglycerate kinase | | -1.532 | -0.727 | -1.597 | -2.246 | 1.066 | 0.056 | -1.125 | 0.300 | 0.975 | -2.848 | -2.476 | | | | | Yes | | | | | |
| B7L74_09170 | type I glyceraldehyde-3-phosphate dehydrogenase | Glycolysis / Gluconeogenesis | -2.740 | -0.817 | -3.337 | -1.557 | -2.151 | -2.020 | -2.610 | -2.535 | -3.290 | -2.256 | -1.644 | Early down | | | | Yes | | | | | |
| B7L74_09175 | transketolase | Pentose phosphate pathway | -1.773 | -0.037 | -3.560 | -0.520 | -2.396 | -1.316 | -1.827 | -1.692 | -2.437 | -1.953 | -1.366 | Early down | | | | | | | | | |
| B7L74_09180 | IS110 family transposase | | 0.197 | -0.284 | 1.580 | -0.626 | -0.877 | -0.642 | 0.377 | -0.761 | -1.040 | 1.152 | 1.531 | | | | | | | | | | |
| B7L74_09185 | hypothetical protein | | 0.096 | 0.849 | -2.477 | -1.391 | 1.203 | 0.964 | 0.507 | 0.016 | 1.161 | -1.248 | -0.894 | | | | | | | | | | |
| B7L74_09190 | ATPase | ATP | -0.262 | 0.675 | -0.563 | -0.291 | 0.782 | 0.078 | 0.073 | 0.581 | 0.763 | 0.198 | 0.066 | | | | | | | | | | |
| B7L74_09195 | glutamine--fructose-6-phosphate aminotransferase | | -0.992 | -0.193 | -0.389 | -0.563 | -1.290 | -1.142 | -1.436 | -0.710 | -1.433 | -1.038 | -0.878 | | | | | | | | | | |
| B7L74_09200 | DNA-binding protein | | 0.041 | 0.838 | -0.034 | -1.805 | 1.973 | 1.621 | 0.448 | 1.487 | 2.548 | -1.279 | -0.910 | | | | | | | | | | |
| B7L74_09205 | hypothetical protein | | -0.309 | 0.430 | 2.159 | 0.907 | -0.317 | -0.387 | -0.432 | 0.757 | -0.966 | -0.067 | -0.201 | | | | | | | | | | |
| B7L74_09210 | gluconolactonase | Effector protein | -0.565 | 0.736 | 0.460 | -0.418 | 0.315 | 0.512 | -0.676 | 0.176 | -0.066 | -1.479 | -1.151 | | | Yes | | | | | | | |
| B7L74_09215 | hypothetical protein | Effector protein | -0.662 | 1.268 | -1.358 | 0.048 | 0.869 | 0.846 | -0.977 | 0.905 | -0.392 | -1.235 | -1.149 | | | Yes | | | | | | | |
| B7L74_09220 | MFS transporter | Transporter | -1.045 | 0.856 | -0.035 | 2.721 | 1.970 | 0.146 | 1.101 | 0.015 | 2.126 | -0.135 | 1.180 | | | | | | | | | | |
| B7L74_09225 | hypothetical protein | Effector protein | 0.470 | 0.995 | -1.434 | -0.186 | -0.161 | 0.137 | -0.243 | 0.104 | 0.743 | -0.049 | 0.897 | | | Yes | | | | | | | |
| B7L74_09230 | DNA polymerase I | DNA | 1.069 | -0.534 | -0.491 | 0.266 | 1.756 | 2.091 | 0.110 | -1.318 | 0.254 | 0.491 | 1.055 | | | | | | | | | | |
| B7L74_09235 | amino acid permease | Transporter | -0.254 | 0.081 | -0.640 | 0.221 | -0.723 | -0.193 | 0.267 | -0.713 | -1.625 | -0.091 | 0.122 | | | | | | | | | | |
| B7L74_09240 | acyl-CoA thioesterase | | 0.235 | -0.225 | -1.305 | 1.461 | -0.378 | -0.502 | -0.266 | -0.808 | -0.458 | 0.104 | 0.177 | | | | | | | | | | |
| B7L74_09245 | amino acid transporter | Transporter | 0.030 | -0.340 | -0.041 | 0.114 | -0.883 | -0.367 | 0.185 | -0.286 | 0.293 | -0.163 | 0.268 | | | | | | | | | | |
| B7L74_09250 | hypothetical proteinprotein=cupin superfamily protein | | -0.497 | -0.117 | -1.495 | -0.421 | -1.086 | -1.147 | -0.699 | -1.275 | -1.376 | -0.921 | -0.981 | | | | | | | | | | |
| B7L74_09255 | N-acetyltransferase | | -0.825 | -0.638 | -2.494 | -0.533 | -0.949 | -1.056 | -1.318 | -1.671 | -2.662 | -1.290 | -1.242 | Early down | | | | | | | | | |
| B7L74_09260 | hypothetical membrane spanning protein | Transporter | -0.923 | -0.155 | -1.035 | -2.773 | 4.298 | 2.530 | -1.224 | 0.494 | 5.158 | -2.259 | -1.899 | Early up | | | | | | | | Yes (Vitamins) | |
| B7L74_09265 | hypothetical protein | | -2.504 | -0.834 | -1.513 | 0.652 | -1.417 | -1.520 | -1.836 | -1.278 | -1.984 | -2.912 | -3.201 | Early down | | | | | | | | | |
| B7L74_09270 | hypothetical protein | | -1.981 | -0.798 | -2.538 | -0.441 | -1.031 | -1.329 | -2.276 | -0.467 | -1.614 | -2.020 | -2.547 | Early down | | | | | | | | | |
| B7L74_09275 | hypothetical protein | | 0.177 | 0.789 | 3.774 | 1.105 | 0.623 | 0.036 | 2.634 | 2.202 | 3.893 | 2.777 | 3.580 | Early up | | | | | | | | | |
| B7L74_09280 | helix-turn-helix transcriptional regulator | Transcriptional regulators | -0.756 | -0.122 | 0.789 | 0.328 | 0.538 | -0.077 | -0.126 | 1.427 | 2.394 | 1.849 | 1.863 | | | | | | | | | | |
| B7L74_09285 | helix-turn-helix transcriptional regulator | Transcriptional regulators | -0.763 | -1.064 | -1.862 | -0.495 | 0.002 | -0.370 | -1.240 | -0.673 | -0.043 | -0.039 | -0.450 | | | | | | | | | | |
| B7L74_09290 | hypothetical protein | | -0.519 | 0.138 | -0.448 | 1.434 | 0.553 | -0.688 | -0.926 | 0.868 | 1.452 | 1.245 | 1.065 | | | | | | | | | | |
| B7L74_09295 | hypothetical protein | | 2.028 | 1.731 | 1.144 | 0.971 | 1.841 | 2.547 | 0.995 | 0.962 | 2.143 | 1.431 | 2.332 | | | | | | | | | | |
| B7L74_09300 | ABC transporter substrate-binding protein | Transporter | -0.193 | -1.595 | -0.698 | -0.300 | -1.474 | -0.956 | -0.324 | -0.316 | -0.544 | -0.324 | -0.910 | | | | | | | | | Yes (drug resistance) | |
| B7L74_09305 | ABC transporter ATP-binding protein | Transporter | -1.068 | -0.614 | -1.329 | -1.610 | -1.426 | -0.959 | -0.624 | -1.083 | -0.964 | -1.053 | -0.936 | | | | | | | | | Yes (drug resistance) | |
| B7L74_09310 | ABC transporter substrate-binding protein | Transporter | -0.102 | 1.640 | -0.044 | 2.655 | 2.655 | 2.679 | 0.032 | 1.183 | 0.929 | -0.234 | -0.862 | | | | | | | | | | |
| B7L74_09315 | type I secretion protein TolC | | 1.442 | -0.807 | 0.468 | 0.289 | 2.092 | 1.953 | 1.235 | 0.092 | 1.304 | 1.161 | 0.687 | | | | | | | | | | |
| B7L74_09320 | erythronate-4-phosphate dehydrogenase | | 0.877 | 1.332 | 1.127 | -0.883 | 0.534 | 0.761 | 0.464 | 0.860 | 0.482 | 0.629 | 0.483 | | | | | | | | | | |
| B7L74_09325 | hypothetical protein | | 0.056 | 0.841 | -0.034 | -1.792 | 1.973 | 1.623 | 0.463 | 1.490 | 1.136 | -1.271 | -0.906 | | | | | | | | | | |
| B7L74_09330 | hypothetical protein Outer membrane protein | Membrane | 1.520 | 0.717 | 1.206 | 2.994 | -0.117 | 1.248 | 1.915 | 1.936 | 1.277 | 2.595 | 1.924 | Early up | | | | | | | | Yes (Outer membrane/Membrane) | |
| B7L74_09335 | primosomal protein N' | | -0.646 | -0.549 | -0.613 | -0.035 | -1.520 | 0.396 | 0.246 | 0.281 | 0.351 | -0.196 | -0.072 | | | | | | | | | | |
| B7L74_09340 | elongation factor P | | 0.612 | 0.411 | 0.311 | 1.644 | -0.318 | 0.371 | 0.129 | 0.015 | -1.588 | 0.036 | 0.146 | | | | | | | | | | |
| B7L74_09345 | EF-P lysine aminoacylase GenX | | 0.283 | 0.061 | -1.233 | 1.205 | -0.823 | -0.103 | -0.133 | 0.431 | -0.342 | -0.570 | -0.009 | | | | | | | | | | |
| B7L74_09350 | hypothetical protein(cvpD in Table 1 of RightonQ pap | Effector protein | -0.425 | 0.280 | 0.295 | -0.191 | 1.126 | 1.580 | -0.698 | 0.759 | 1.847 | -0.184 | -0.389 | | | Yes | | | | | | | |
| B7L74_09355 | hypothetical membrane associated protein | Effector protein | 1.212 | 0.115 | 0.272 | 0.230 | 3.700 | 2.648 | 0.573 | 0.089 | 2.084 | 0.945 | 0.204 | Up in P16 | | Yes | | | | | | | |
| B7L74_09360 | hypothetical protein | | -0.107 | 0.497 | -0.289 | 1.537 | -0.121 | -0.710 | 0.162 | 0.423 | -0.016 | 0.506 | 1.253 | | | | | | | | | | |
| B7L74_09365 | hypothetical protein | | -0.172 | -0.001 | -0.603 | 0.615 | -1.193 | -1.491 | -0.566 | -0.154 | 0.289 | -0.329 | -0.392 | | | | | | | | | | |
| B7L74_09370 | superoxide dismutase | Antioxidant PV detoxification | 0.015 | 1.049 | 1.648 | 0.778 | 0.798 | 0.771 | 0.049 | 0.551 | 0.775 | 0.040 | -0.444 | | | | | | | | | | |
| B7L74_09375 | hypothetical protein (icaA in Table 1 of RightonQ pap | Effector protein | 0.112 | 1.611 | -1.140 | -0.537 | 0.915 | 0.034 | -0.835 | 0.858 | -0.011 | 0.966 | -0.077 | | | Yes | | | | | | | |
| B7L74_09380 | hypothetical protein | | 2.724 | 1.856 | -0.763 | -2.412 | 0.862 | 1.312 | 0.168 | 0.268 | 0.267 | -1.354 | -1.068 | | | | | | | | | | |
| B7L74_09385 | hypothetical protein | Effector protein | 0.200 | -1.008 | -1.071 | 0.685 | 0.138 | 0.487 | 0.247 | 0.596 | -1.906 | -1.084 | -0.891 | | | Yes | | | | | | | |
| B7L74_09390 | phosphatidylserine decarboxylase proenzyme | | -0.362 | 0.076 | -0.225 | 0.895 | -1.015 | -0.678 | -0.424 | 0.316 | 0.336 | -0.624 | -0.285 | | | | | | | | | | |
| B7L74_09395 | multifunctional CCA tRNA nucleotidyl transferase/2'3 | | -0.418 | -0.485 | -0.079 | 0.839 | -1.201 | -0.291 | -0.243 | -0.733 | -0.003 | -0.002 | 0.226 | | | | | | | | | | |
| B7L74_09400 | SAM-dependent methyltransferase | | 0.454 | 0.898 | 1.751 | 0.308 | 0.593 | 0.238 | 0.678 | 0.552 | 0.159 | 0.967 | 1.024 | | | | | | | | | | |
| B7L74_09405 | outer membrane lipoprotein LolB | Membrane | -0.233 | -0.180 | -0.011 | -0.477 | -0.509 | -0.593 | -0.215 | -0.092 | -1.138 | 0.314 | -0.108 | | | | | | | | | | |
| B7L74_09410 | tRNA-Gln | tRNA | 0.584 | -0.010 | 1.473 | 1.109 | 0.413 | -0.764 | 1.428 | 0.224 | -0.337 | 1.458 | 1.470 | | | | | | | | | | |
| B7L74_09415 | ribose-phosphate pyrophosphokinase | | 0.055 | -0.866 | -0.524 | 1.474 | -2.759 | -1.055 | -0.363 | -0.893 | -0.631 | 0.740 | 0.687 | | | | | | | | | | |
| B7L74_09420 | dolichyl-phosphate-mannose--protein mannosyltransfe | | 6.326 | 2.664 | 3.619 | 1.246 | 7.716 | 7.989 | 4.015 | 2.035 | 6.510 | 4.140 | 2.254 | Continuous up | | | | | | | | | |
| B7L74_09425 | hypothetical protein | | -2.166 | -3.456 | -4.338 | -6.053 | -0.415 | 2.309 | -3.777 | 2.018 | 1.172 | -3.848 | -5.201 | | | | | | | | | | |
| B7L74_09430 | hypothetical protein | | 0.711 | -0.644 | -1.526 | -3.241 | 1.245 | 0.136 | -0.964 | 0.005 | 1.137 | -1.822 | -1.443 | | | | | | | | | | |
| B7L74_09435 | glucose-1-phosphate thymidylyltransferase | O-Antigen nucleotide sugar biosy | 0.172 | -0.006 | 0.058 | 2.087 | -0.235 | -0.075 | 0.507 | 0.116 | 0.243 | 1.591 | 1.428 | | | | | | | | | | Yes |
| B7L74_09440 | amine oxidase | | -1.487 | -0.666 | -2.531 | -1.592 | -1.602 | -1.054 | -1.643 | -1.515 | -2.301 | -1.249 | -1.148 | Early down | | | | | | | | | |
| B7L74_09445 | homoserine dehydrogenase | | -1.438 | -0.168 | -0.634 | -0.160 | -0.796 | -0.444 | -0.939 | -0.471 | -0.813 | -0.758 | -0.510 | | | | | | | | | | |
| B7L74_09450 | NAD-dependent dehydratase | | -1.709 | -0.937 | -1.531 | -0.431 | -1.471 | -0.700 | -0.843 | -1.442 | -1.180 | -0.703 | -0.316 | | | | | | | | | | |
| B7L74_09455 | dTDP-4-dehydrorhamnose 3%2C5-epimerase | O-Antigen nucleotide sugar biosy | 0.478 | -0.973 | -0.263 | -0.694 | 0.159 | 0.213 | 0.368 | 0.143 | 0.947 | -0.223 | 0.525 | | | | | | | | | | Yes |
| B7L74_09460 | aminobutyraldehyde dehydrogenase | | 0.087 | 0.319 | 0.140 | 1.327 | -0.654 | -0.219 | 0.969 | 0.715 | 1.099 | 1.654 | 1.605 | | | | | | | | | | |
| B7L74_09465 | 50S ribosomal protein L25 | Ribosomal proteins | -1.031 | 0.709 | -1.584 | 1.700 | -0.475 | -0.666 | -1.126 | 0.579 | -2.163 | -1.148 | -0.349 | | | | | | | | | | |
| B7L74_09470 | peptidyl-tRNA hydrolase | | 0.566 | 0.673 | -0.878 | 1.860 | -0.781 | -0.495 | -0.597 | 0.580 | -0.471 | -1.613 | -0.298 | | | | | | | | | | |
| B7L74_09475 | GTP-binding protein YchF | | -0.212 | 0.098 | 0.241 | 0.045 | -0.947 | -0.461 | 0.170 | 0.087 | -1.272 | 0.777 | 0.453 | | | | | | | | | | |
| B7L74_09480 | hypothetical exported protein | Exported | -0.084 | 0.515 | 1.248 | 1.236 | 0.946 | 0.561 | -0.089 | 0.959 | 0.193 | 1.699 | 0.940 | | | | | | | | | | |
| B7L74_09485 | tRNA-Met | tRNA | 0.137 | 0.945 | 1.480 | 0.399 | -0.588 | -0.236 | 0.940 | 0.601 | 0.909 | 0.128 | 0.540 | | | | | | | | | | |
| B7L74_09490 | hypothetical protein | | 0.320 | 1.115 | 1.578 | -0.106 | 1.516 | 0.958 | 0.656 | 1.059 | -0.261 | 0.206 | 0.865 | | | | | | | | | | |
| B7L74_09495 | DUF1658 domain-containing protein | | 0.035 | 0.837 | -0.034 | -1.810 | 2.635 | 1.620 | 0.443 | 1.486 | 2.542 | -1.283 | -0.912 | | | | | | | | | | |
| B7L74_09500 | DUF1658 domain-containing protein | | -0.427 | 0.008 | 2.041 | 0.337 | 0.038 | 0.604 | 1.206 | 0.435 | 0.793 | 0.411 | 0.779 | | | | | | | | | | |
| B7L74_09505 | hypothetical membrane spanning protein | | 0.187 | 0.494 | 1.725 | 1.029 | 0.625 | 0.250 | 0.456 | 0.840 | 0.357 | 1.391 | 1.293 | | | | | | | | | | |
| B7L74_09510 | hypothetical protein | | -0.078 | 0.881 | 3.291 | 1.745 | 1.456 | 0.866 | 1.270 | 0.474 | 2.973 | 0.642 | 2.167 | Early up | | | | | | | | | |
| B7L74_09515 | hypothetical exported protein | | -0.500 | 1.219 | 3.631 | 1.202 | 2.275 | 1.798 | 2.450 | 1.284 | 4.056 | 0.997 | 2.091 | Early up | | | | | | | | | |
| B7L74_09520 | hypothetical protein | | 0.621 | 0.734 | 0.666 | 0.754 | 1.017 | 0.950 | 1.363 | 1.887 | 1.806 | 2.241 | 1.409 | Late up | | | | | | | | | |
| B7L74_09525 | hypothetical protein | | 0.045 | 0.839 | -0.034 | -1.802 | 2.646 | 1.955 | 0.845 | 1.911 | 1.882 | -0.770 | -0.909 | | | | | | | | | | |
| B7L74_09530 | hypothetical protein | | 0.309 | 1.734 | 1.439 | 0.108 | 0.781 | 0.594 | -0.056 | 0.889 | 1.329 | 0.376 | -0.093 | | | | | | | | | | |
| B7L74_09535 | hypothetical protein | | 1.345 | 1.905 | 0.828 | 0.466 | 0.297 | 1.788 | 0.115 | 0.564 | 0.844 | 0.494 | 1.048 | | | | | | | | | | |
| B7L74_09540 | hypothetical protein | | 5.074 | 1.678 | 2.976 | 2.321 | 6.198 | 6.392 | 3.181 | 0.634 | 5.399 | 3.240 | 3.774 | Early up | | | | | | | | | |
| B7L74_09545 | hypothetical protein, Hypothetical membrane spanning | Membrane | 4.304 | 1.189 | 2.303 | -0.399 | 5.631 | 5.413 | 2.535 | 1.257 | 4.733 | 2.891 | 2.454 | Continuous up | | | | | | | | | |
| B7L74_09550 | dolichyl-phosphate-mannose--protein mannosyltransfe | | -1.920 | -1.338 | -2.855 | -1.810 | 1.272 | 1.435 | -2.375 | 1.126 | -0.275 | -3.203 | -2.903 | | | | | | | | | | |
| B7L74_09555 | sugar translocase | | -0.923 | -1.183 | -1.018 | -1.868 | 0.989 | 0.639 | -0.136 | 2.271 | 0.152 | -0.587 | -1.888 | | | | | | | | | | |
| B7L74_09560 | hypothetical cytosolic protein | | 0.061 | 0.210 | 0.917 | -0.685 | -1.617 | -0.760 | 0.198 | 0.832 | 0.728 | 1.356 | 1.422 | | | | | | | | | | |
| B7L74_09565 | fimbrial protein, PilF | Pilus | -0.663 | 1.388 | 1.128 | -1.462 | 0.507 | 0.277 | -0.523 | 1.337 | 1.864 | 1.175 | 1.170 | | | | | | | | | | |
| B7L74_09570 | 3-hydroxyisobutyryl-CoA hydrolase | | -0.305 | 0.446 | 0.716 | 0.588 | -0.392 | 0.385 | 0.037 | 0.738 | 0.107 | 0.015 | 0.244 | | | | | | | | | | |
| B7L74_09575 | ABC transporter ATP-binding protein | Transporter | -0.065 | -0.532 | -0.363 | -1.404 | -1.678 | 1.326 | 1.032 | 1.326 | 0.349 | -0.664 | 0.170 | | | | | | | | | | |
| B7L74_09580 | peptide ABC transporter permease | Transporter | 1.841 | 1.143 | -0.737 | 0.072 | 2.630 | 2.813 | 1.274 | 2.107 | 3.773 | 0.961 | 0.697 | | | | | | | | | | |
| B7L74_09585 | peptide ABC transporter permease | Transporter | 0.576 | -0.014 | -1.340 | -0.179 | 0.788 | 2.264 | 1.921 | 2.421 | 2.797 | 0.942 | 1.034 | | | | | | | | | | |
| B7L74_09590 | peptide ABC transporter substrate-binding protein | Transporter | 0.334 | 0.477 | 1.085 | 0.289 | -1.079 | 0.979 | 1.633 | 2.175 | 1.690 | 1.254 | 1.093 | | | | | | | | | | |
| B7L74_09595 | EF-P beta-lysylation protein EpmB | Translation factors | 0.303 | 0.019 | 0.247 | 0.312 | 3.137 | 3.235 | 0.849 | -0.259 | 2.154 | 0.350 | -0.197 | Early up | | | | | | | | | |
| B7L74_09600 | hypothetical protein (cvpE in Table 1 of RightonQ pap | Effector protein | 6.174 | 1.032 | -21.214 | -7.864 | 5.420 | 4.397 | -0.652 | -17.897 | 2.566 | 1.106 | -1.089 | | | Yes | | | | | | | |
| B7L74_09605 | hypothetical protein | | 0.297 | 0.157 | -1.069 | 0.079 | 0.019 | -0.071 | -1.224 | -0.648 | -2.061 | -1.198 | -1.208 | | | | | | | | | | |
| B7L74_09610 | hypothetical protein, Hypothetical membrane associate | Membrane | -1.445 | -0.184 | -2.658 | -1.216 | -1.576 | -1.141 | -1.841 | -2.158 | -3.208 | -1.930 | -1.749 | Early down | | | | | | | | | |
| B7L74_09615 | DNA topoisomerase IV subunit A | DNA replication proteins | -0.562 | -0.510 | -0.799 | 0.094 | -1.479 | -1.342 | -0.494 | -1.400 | -1.782 | -0.695 | 0.033 | Early down | | | | | | | | | |
| B7L74_09620 | hypothetical protein | | -0.881 | -0.774 | 1.015 | -1.754 | -3.411 | -0.164 | 0.390 | 0.773 | -0.040 | 0.543 | 1.005 | | | | | | | | | | |
| B7L74_09625 | DUF1658 domain-containing protein | CB specific protein family | 0.790 | 2.153 | 3.841 | -1.316 | 3.285 | 2.581 | 2.242 | 1.874 | 1.040 | 0.711 | 0.680 | Early up | | | | | | | | | |
| B7L74_09630 | hypothetical exported protein | | -0.270 | 0.288 | 0.174 | 1.102 | 0.199 | -0.134 | -0.625 | -0.131 | -0.662 | 0.120 | -0.406 | | | | | | | | | | |
| B7L74_09635 | hypothetical protein | | 0.166 | -1.595 | -0.035 | 0.601 | 0.294 | 0.146 | 0.077 | 1.419 | 0.185 | 0.278 | -1.304 | | | | | | | | | | |

| | | | | | | | | | | | | | | | | | | | | | | | | |
|---|---|---|---|---|---|---|---|---|---|---|---|---|---|---|---|---|---|---|---|---|---|---|---|---|
| B7L74_09640 | demethoxyubiquinone hydroxylase family protein | | -0.540 | 0.148 | 0.613 | 0.442 | 0.285 | 0.072 | -0.252 | -0.592 | -0.145 | -0.093 | 0.284 | | | | | | | | | | | |
| B7L74_09645 | glutamine amidotransferase | | -0.099 | -0.625 | -1.508 | 5.108 | 1.264 | 1.116 | 2.040 | 0.023 | 2.090 | 3.222 | 0.109 | | | | | | | | | | | |
| B7L74_09650 | ribulose-phosphate 3-epimerase | | -0.104 | -0.627 | -1.510 | 4.102 | 1.262 | 1.114 | 2.030 | 0.021 | 2.086 | 2.213 | 0.109 | | | | | | | | | | | |
| B7L74_09655 | molecular chaperone DjlA | | -0.107 | -0.629 | -1.511 | 2.754 | 1.260 | 1.112 | 0.031 | 0.019 | 2.084 | 0.013 | 0.109 | | | | Yes | | | | | | | |
| B7L74_09660 | glutamate--cysteine ligase | | -0.567 | -0.513 | 0.099 | -0.865 | -0.293 | -0.398 | -0.816 | -1.375 | -1.020 | -1.251 | -1.057 | | | | | | | | | | | |
| B7L74_09665 | glutathione synthetase | ATP | -0.129 | -0.165 | 0.549 | 0.083 | -0.221 | -0.404 | -0.377 | -1.235 | -1.703 | -0.934 | -1.181 | | | | | | | | | | | |
| B7L74_09670 | hypothetical protein | | 5.838 | 0.847 | -0.035 | 9.657 | 6.529 | 5.821 | 6.960 | 1.495 | 2.627 | 6.675 | 7.211 | | | | | | | | | | | |
| B7L74_09675 | ribosomal RNA small subunit methyltransferase J | | -0.663 | 0.457 | 0.431 | -1.032 | 0.429 | 0.738 | -0.380 | -0.650 | -0.105 | 0.247 | -0.591 | | | | | | | | | | | |
| B7L74_09680 | ATPase | ATP | -1.699 | -5.463 | -6.365 | 1.066 | -4.958 | -4.384 | 0.210 | 2.348 | -1.367 | 0.004 | 0.567 | | | | | | | | | | | |
| B7L74_09685 | iron-sulfur cluster insertion protein ErpA | | 0.932 | 0.795 | 1.219 | 1.456 | -0.121 | -0.340 | 0.424 | 0.407 | -0.557 | 0.488 | 0.525 | | | | | | | | | | | |
| B7L74_09690 | peptide deformylase 2 | | -0.655 | -0.130 | -1.167 | 0.181 | -1.106 | -1.389 | -0.953 | -1.298 | -2.257 | -0.792 | -0.965 | Early down | | | | | | | | | | |
| B7L74_09695 | hypothetical membrane spanning protein | | 0.310 | 1.169 | 1.714 | 0.072 | 2.833 | 1.455 | 2.449 | 2.034 | 2.069 | -1.394 | 2.542 | | | | | | | | | | | |
| B7L74_09700 | rubredoxin | PV detoxification | 0.090 | 0.848 | -0.035 | -0.788 | 2.737 | 1.627 | 0.527 | 1.496 | 2.628 | -1.252 | 0.953 | | | | | | | | | | | |
| B7L74_09705 | glutamate-1-semialdehyde 2%2C1-aminomutase | Porphyrin and chlorophyll metabol | -0.551 | -0.294 | -1.012 | 0.086 | -1.653 | -1.443 | -0.737 | -1.584 | -2.267 | -0.436 | -0.463 | Early down | | | | | | | | | | |
| B7L74_09710 | queuosine biosynthesis protein | | -0.540 | 1.698 | -0.629 | 0.980 | 3.550 | 1.416 | -0.133 | 0.895 | 1.410 | -1.517 | -1.501 | | | | | | | | | | | |
| B7L74_09715 | hypothetical exported membrane associated protein | | -0.835 | 0.318 | -0.677 | 0.101 | -0.504 | -1.360 | -0.558 | -0.727 | -0.286 | 0.045 | -0.156 | | | | | | | | | | | |
| B7L74_09720 | hypothetical protein | | -1.074 | -1.592 | -2.929 | -1.614 | -3.600 | -0.904 | -2.379 | -1.398 | -0.278 | -2.182 | -1.056 | | | | | | | | | | | |
| B7L74_09725 | peptidase | Peptidoglycan synbeta-Lactam resi | -0.801 | -0.911 | -1.748 | -1.440 | -1.605 | -2.117 | -1.057 | -1.465 | -2.269 | -1.014 | -1.095 | Early down | | | | | | | | | | Yes |
| B7L74_09730 | pilus assembly protein PilM | | 0.885 | 1.358 | -1.005 | 0.744 | 2.224 | 2.971 | 1.398 | 1.302 | 4.194 | 0.464 | 0.637 | | | | | | | | | | | |
| B7L74_09735 | pilus assembly protein PilN | | 1.791 | 3.305 | 0.972 | -0.797 | 4.084 | 3.589 | 1.457 | 3.157 | 4.140 | 0.487 | 0.096 | | | | | | | | | | | |
| B7L74_09740 | hypothetical protein | | 0.030 | 0.835 | 0.966 | -0.352 | 2.628 | 1.619 | 0.437 | 1.485 | 2.537 | -1.286 | -0.914 | | | | | | | | | | | |
| B7L74_09745 | secretin, pilQ | Pilus | -1.949 | -0.220 | -2.016 | 1.692 | 0.650 | -0.362 | -1.542 | -0.496 | -0.097 | -0.911 | -0.661 | | | | | | | | | | | |
| B7L74_09750 | shikimate kinase (gene=aroK) | Phenylalanine, tyrosine and tryptop | 0.384 | 5.323 | 2.598 | 2.934 | 4.372 | 5.016 | 2.512 | 3.772 | 2.021 | 0.687 | 1.494 | Early up | | | | | | | | | | |
| B7L74_09755 | 3-dehydroquinate synthase | | -2.758 | 2.715 | 0.566 | 0.687 | 1.886 | 2.048 | -0.398 | 1.541 | -1.902 | -1.285 | 0.288 | | | | | | | | | | | |
| B7L74_09760 | DUF1658 domain-containing protein | | 1.117 | 2.073 | 0.679 | 1.123 | 1.844 | 1.518 | 1.059 | 0.081 | 0.103 | -0.619 | 1.126 | | | | | | | | | | | |
| B7L74_09765 | DUF1658 domain-containing protein | | 0.697 | 1.538 | 0.167 | -0.213 | 0.660 | 0.797 | -0.571 | 0.541 | -1.247 | -0.901 | 0.048 | | | | | | | | | | | |
| B7L74_09770 | hypothetical protein | | -1.379 | 0.875 | -0.362 | 1.220 | -0.099 | -1.117 | -1.772 | 0.313 | -1.821 | 0.717 | 0.084 | | | | | | | | | | | |
| B7L74_09775 | MFS transporter | Transporter | 0.090 | 0.822 | 0.764 | 1.478 | 0.500 | 0.698 | 0.484 | 1.203 | 1.385 | 0.754 | 1.213 | | | | | | | | | | | |
| B7L74_09780 | IS110 family transposase | | 0.201 | -1.238 | 0.941 | -0.275 | 0.674 | 2.321 | 1.768 | -2.709 | 0.935 | 0.953 | 0.779 | | | | | | | | | | | |
| B7L74_09785 | hypothetical protein | | 0.662 | -1.031 | 0.980 | -0.435 | -1.613 | -0.275 | 0.432 | -1.619 | -2.847 | 0.171 | 0.465 | | | | | | | | | | | |
| B7L74_09790 | methyltransferase | | 1.081 | -0.430 | -0.577 | -0.482 | 0.772 | 1.388 | 0.185 | -1.254 | -0.463 | 0.279 | 0.482 | | | | | | | | | | | |
| B7L74_09795 | DUF1658 domain-containing protein | CB specific protein family | -1.710 | -0.153 | -2.550 | -3.114 | -1.642 | -1.473 | -2.101 | -2.044 | -3.114 | -3.111 | -3.089 | Early down | | | | | | | | | | |
| B7L74_09800 | DUF1658 domain-containing protein | | 0.358 | -0.203 | -0.862 | -1.515 | -0.955 | -1.941 | -0.519 | -1.739 | -1.702 | -1.133 | -1.056 | | | | | | | | | | | |
| B7L74_09805 | peptidase M16 | Peptidases and inhibitors | -0.321 | -0.679 | -0.148 | -1.246 | -1.378 | -0.912 | -0.866 | -1.949 | -2.419 | -1.337 | -1.438 | Late down | | | | | | | | | | |
| B7L74_09810 | peptidase M16 | | 0.578 | -0.021 | -3.724 | -1.986 | 0.937 | -1.202 | -0.496 | -0.904 | -0.316 | -2.442 | -1.981 | | | | | | | | | | | |
| B7L74_09815 | signal recognition particle-docking protein FtsY | | -0.977 | -0.779 | -0.554 | -0.122 | -1.417 | -0.878 | -0.787 | -1.560 | -1.171 | -1.159 | -0.740 | | | | | | | | | | | |
| B7L74_09820 | cell division ATP-binding protein FtsE | ABC transporters | -1.187 | -0.856 | -2.002 | -1.153 | -1.204 | -0.497 | -1.653 | -1.695 | -2.118 | -1.491 | -1.439 | Early down | | | | | | | | Yes (Cell division) | | |
| B7L74_09825 | ABC transporter permease | Transporter | -0.027 | 0.580 | -1.710 | 1.168 | 3.744 | 0.748 | 0.612 | 1.075 | 0.308 | 0.182 | 0.783 | | | | | | | | | Yes (Cell division) | | |
| B7L74_09830 | hypothetical protein | | -1.437 | -0.797 | -3.226 | -2.068 | -0.960 | -1.590 | -1.017 | -1.796 | -2.762 | -1.816 | -2.033 | Early down | | | | | | | | | | |
| B7L74_09835 | N-acetyltransferase | | -0.483 | -0.177 | 1.506 | 1.077 | -0.507 | -0.931 | 0.684 | -0.317 | 0.542 | 0.590 | 1.550 | | | | | | | | | | | |
| B7L74_09840 | hypothetical protein | | -0.048 | -0.582 | 1.978 | -1.040 | 2.390 | 2.452 | 1.195 | 1.028 | 3.671 | 1.865 | -2.327 | | | | | | | | | | | |
| B7L74_09845 | RNA polymerase factor sigma-32 RpoH | Transcription machinery | -1.087 | -1.176 | -2.205 | -0.564 | -2.376 | -2.367 | -1.546 | -2.441 | -3.378 | -1.365 | -1.381 | Continuous down | | | | | | | | | | |
| B7L74_09850 | hypothetical protein outer membrane protein | surface antigen Coml | -2.086 | -1.836 | -3.636 | 0.555 | -2.342 | -2.774 | -1.837 | -2.793 | -4.023 | -2.435 | -2.330 | Early down | | | | | | | | | | |
| B7L74_09855 | DUF1658 domain-containing protein | | -0.104 | 0.163 | -0.038 | -0.761 | 0.700 | -0.143 | 0.463 | -0.144 | -0.403 | -1.282 | -0.367 | | | | | | | | | | | |
| B7L74_09860 | hypothetical protein | | 1.618 | 1.513 | 1.966 | -1.684 | 1.918 | 1.832 | 0.891 | 1.433 | 1.392 | 0.094 | 0.391 | | | | | | | | | | | |
| B7L74_09865 | hypothetical protein | | 0.062 | 0.842 | -0.034 | -1.120 | 1.973 | 1.053 | 0.469 | 1.491 | 1.135 | -0.433 | -0.432 | | | | | | | | | | | |
| B7L74_09870 | glycine--tRNA ligase subunit alpha | Aminoacyl-tRNA synthase | 5.459 | -1.057 | 0.597 | 0.216 | 4.502 | 5.562 | 1.328 | -1.375 | 4.787 | 3.355 | 1.417 | Early up | | | | | | | | | | |
| B7L74_09875 | glycine--tRNA ligase subunit beta | Aminoacyl-tRNA synthase | 4.111 | 1.180 | 0.911 | 0.450 | 4.977 | 5.757 | 0.596 | -0.368 | 0.599 | 1.611 | -0.111 | Early up | | | | | | | | | | |
| B7L74_09880 | 16S rRNA m5C967 methyltransferase | | -0.862 | 0.894 | -2.217 | 0.491 | 1.336 | -0.933 | -1.263 | -0.069 | -2.608 | 0.221 | 0.862 | | | | | | | | | | | |
| B7L74_09885 | universal stress protein A (UspA) | | 1.087 | 1.838 | 0.954 | -0.121 | 2.191 | 1.135 | 1.499 | 1.983 | 2.120 | -0.258 | 0.096 | | | | | | | | | | | |
| B7L74_09890 | 50S ribosomal protein L34 | Ribosomal proteins | 0.381 | 1.867 | -0.696 | 3.821 | 0.012 | 0.303 | 0.065 | 0.621 | -0.439 | -1.443 | -0.312 | Up in P12 | | | | | | | | | | |
| B7L74_09895 | ribonuclease P protein component | | -0.034 | 0.955 | -1.376 | -2.762 | 1.301 | 0.279 | -0.893 | 0.145 | 1.207 | -2.621 | -2.252 | | | | | | | | | | | |
| B7L74_09900 | membrane protein insertion efficiency factor YidD | | 0.075 | 0.845 | -0.034 | 1.071 | 1.298 | 0.971 | 0.483 | 1.494 | 1.877 | -1.260 | -0.900 | | | | | | | | | | | |
| B7L74_09905 | membrane protein insertase YidC | | 1.053 | 0.289 | -1.774 | 0.673 | -0.449 | -0.879 | -0.329 | -1.094 | -0.757 | -0.500 | -0.656 | | | | | | | | | | | |
| B7L74_09910 | hypothetical protein | | 1.192 | 0.912 | 2.771 | -1.187 | 1.484 | 0.211 | 2.424 | 2.014 | 2.073 | 1.286 | 3.273 | | | | | | | | | | | |
| B7L74_09915 | tRNA uridine-5-carboxymethylaminomethyl(34) synth | | 0.073 | 0.845 | -0.034 | 1.776 | 2.683 | 1.625 | 1.137 | 2.900 | 2.582 | 2.685 | 5.865 | | | | | | | | | | | |
| B7L74_09920 | tRNA uridine-5-carboxymethylaminomethyl(34) synth | Transfer RNA biogenesis | -0.266 | -0.691 | -1.135 | -1.019 | -1.396 | -1.853 | -0.976 | -1.062 | -1.863 | -1.359 | -1.363 | Early down | | | | | | | | | | |
| B7L74_09925 | ribosomal RNA small subunit methyltransferase G | Ribosome biogenesis | -1.548 | -1.341 | -0.249 | -0.251 | -2.050 | -2.265 | -0.875 | -1.944 | -2.258 | -0.752 | -0.625 | Continuous down | | | | | | | | | | |
| B7L74_09930 | chromosome partitioning protein | Chromosome and associated prote | -0.931 | -1.272 | -1.528 | -0.751 | -2.070 | -2.402 | -1.195 | -2.061 | -3.052 | -0.914 | -0.754 | Continuous down | | | | | | | | | | |
| B7L74_09935 | chromosome partitioning protein ParB | Cell division | 0.037 | 0.698 | 0.621 | -1.721 | 1.343 | 0.280 | 1.114 | 1.202 | 2.721 | 0.656 | -1.102 | | | | | | | | | | | |
| B7L74_09940 | acyl-CoA synthetase | | 3.062 | -0.828 | -0.323 | -0.144 | 3.810 | 3.034 | 1.372 | -1.720 | 2.696 | 1.013 | 0.692 | | | | | | | | | | | |
| B7L74_09945 | MFS transporter | Transporter | -0.544 | 0.259 | -0.611 | -2.389 | 2.055 | 1.738 | -0.136 | 1.332 | 1.964 | 0.164 | -1.490 | | | | | | | | | | | |
| B7L74_09950 | DUF1328 domain-containing protein | | 0.260 | 0.698 | 2.190 | 0.731 | 1.114 | 0.732 | 1.452 | 1.424 | 2.953 | 1.666 | 1.339 | | | | | | | | | | | |
| B7L74_09955 | hypothetical protein | | 0.552 | 0.403 | 2.419 | 0.755 | 0.394 | 0.157 | 1.270 | 0.208 | 1.535 | 2.021 | 2.598 | | | | | | | | | | | |
| B7L74_09960 | hypothetical protein | | 1.959 | 1.334 | 5.013 | 0.371 | 3.198 | 1.553 | 4.174 | 1.766 | 4.168 | 3.404 | 5.493 | Early up | | | | | | | | | | |
| B7L74_09965 | hypothetical protein | | 2.366 | 1.368 | 5.532 | 0.850 | 4.616 | 2.591 | 5.149 | 2.228 | 5.296 | 3.976 | 6.227 | Early up | | | | | | | | | | |
| B7L74_09970 | Ku proteinNon-homologous end joining protein Ku | DNA Non-homologous end-joinin | 5.781 | 1.877 | 3.953 | 2.407 | 6.981 | 7.214 | 4.299 | 2.436 | 6.331 | 5.111 | 4.728 | Continuous up | | | | | | | | | | |
| B7L74_09975 | ATP-dependent DNA ligase | Non-homologous end-joining | -0.888 | 0.067 | -0.836 | -2.679 | 7.321 | 3.811 | -0.352 | 5.015 | 9.272 | -3.032 | -2.686 | Early up | | | | | | | | | | |
| B7L74_09980 | DUF1658 domain-containing protein | | 0.998 | 1.078 | 1.097 | 0.030 | 0.621 | 1.134 | 0.634 | 0.653 | 0.552 | 0.319 | 0.696 | | | | | | | | | | | |
| B7L74_09985 | hypothetical protein | | 2.845 | 1.929 | 2.558 | -0.019 | 3.013 | 2.769 | 2.755 | 2.430 | 1.400 | 1.081 | 2.977 | | | | | | | | | | | |
| B7L74_09990 | DUF1658 domain-containing protein | | 1.335 | 0.266 | 0.969 | -0.306 | 0.424 | 1.017 | 1.243 | 1.130 | 0.724 | 0.534 | 1.441 | | | | | | | | | | | |
| B7L74_09995 | hypothetical protein | | 0.452 | 0.570 | -0.904 | 0.740 | -1.980 | 1.232 | 0.879 | 2.156 | 1.867 | 1.417 | 0.742 | | | | | | | | | | | |
| B7L74_10000 | hypothetical cytosolic protein | | 0.646 | -0.323 | -0.011 | -0.469 | 0.635 | 1.262 | 1.045 | 0.893 | -0.152 | -0.916 | 0.043 | | | | | | | | | | | |
| B7L74_10005 | hypothetical protein | | 1.152 | 0.794 | 1.976 | -0.699 | 3.128 | 0.195 | 0.068 | 2.517 | 4.435 | -0.052 | 0.130 | | | | | | | | | | | |
| B7L74_10010 | AAA family ATPase | | -0.591 | -0.506 | 0.340 | 0.831 | -0.272 | -0.228 | -0.025 | 0.078 | 0.040 | 0.868 | 0.274 | | | | | | | | | | | |
| B7L74_10015 | F0F1 ATP synthase subunit I | Photosynthesis proteins | 0.200 | -0.642 | -1.649 | 0.230 | -1.048 | -1.412 | -0.408 | -1.420 | -2.919 | -0.765 | -0.227 | Down in P51 | | | | | | | | | | |
| B7L74_10020 | ATP synthase subunit A | Respiration/ATPase | -1.820 | -1.428 | -3.956 | -0.503 | -1.892 | -1.827 | -1.611 | -1.665 | -2.284 | -1.788 | -1.442 | Early down | | | | | | Yes | | | | |
| B7L74_10025 | F0F1 ATP synthase subunit C | Respiration/ATPase | -1.883 | -2.514 | -3.562 | -1.398 | -1.999 | -2.583 | -2.295 | -2.012 | -3.078 | -2.147 | -1.891 | Early down | | | | | | Yes | | | | |
| B7L74_10030 | ATP synthase subunit B | Respiration/ATPase | -2.620 | -1.721 | -3.434 | -1.380 | -2.022 | -2.108 | -2.424 | -3.021 | -3.090 | -2.864 | -2.261 | Early down | | | | | | Yes | | | | |
| B7L74_10035 | F0F1 ATP synthase subunit delta | Respiration/ATPase | 0.254 | 0.592 | -2.905 | -0.295 | -0.900 | -0.862 | 1.791 | 0.220 | 0.595 | -2.035 | 0.014 | | | | | | | Yes | | | | |
| B7L74_10040 | ATP synthase subunit alpha | Respiration/ATPase | -0.551 | -1.633 | -3.223 | -1.913 | -1.436 | -0.951 | -1.667 | -2.875 | -3.180 | -1.961 | -2.070 | Early down | | | | | | Yes | | | | |
| B7L74_10045 | ATP synthase subunit gamma | Respiration/ATPase | -2.218 | -2.143 | -3.864 | -2.321 | -2.457 | -2.305 | -2.524 | -3.281 | -4.518 | -2.722 | -2.214 | Continuous down | | | | | | Yes | | | | |
| B7L74_10050 | ATP synthase subunit beta | Respiration/ATPase | -1.246 | -1.460 | -1.809 | -0.982 | -1.737 | -1.947 | -1.747 | -2.617 | -2.516 | -2.121 | -1.710 | Early down | | | | | | Yes | | | | |
| B7L74_10055 | ATP synthase epsilon chain | Respiration/ATPase | -0.136 | -0.644 | 0.321 | -1.191 | -1.184 | -1.508 | -0.251 | -0.971 | -1.613 | -0.515 | -0.408 | | | | | | | Yes | | | | |
| B7L74_10060 | bifunctional N-acetylglucosamine-1-phosphate uridyltr | O-Antigen nucleotide sugar biosyr | -1.267 | -1.694 | -0.460 | -1.046 | -1.926 | -1.556 | -1.116 | -2.102 | -2.067 | -1.050 | -0.865 | Early down | | | | | | | | | Yes | |
| B7L74_10065 | DUF1658 domain-containing protein | CB specific protein family | 1.134 | 1.974 | 4.596 | -1.285 | 2.957 | 2.607 | 2.805 | 1.945 | 2.820 | 1.499 | 1.543 | Early up | | | | | | | | | | |
| B7L74_10070 | DUF1658 domain-containing protein | CB specific protein family | 0.606 | 1.708 | 4.069 | -1.332 | 2.370 | 2.152 | 2.163 | 1.344 | 2.921 | 1.742 | 1.860 | Early up | | | | | | | | | | |
| B7L74_10075 | ATPase | ATP | 0.976 | 0.426 | 1.317 | 2.266 | 0.833 | 0.634 | 0.995 | 0.777 | 0.826 | 1.372 | 1.246 | Up in P12 | | | | | | | | | | |
| B7L74_10080 | hypothetical membrane spanning protein | | 1.184 | 1.115 | 2.802 | -0.860 | 1.142 | 0.906 | 1.076 | 0.560 | 2.407 | 0.597 | 0.550 | | | | | | | | | | | |
| B7L74_10085 | hypothetical protein | | -0.473 | 0.889 | 0.916 | 1.239 | -0.228 | 0.055 | -0.885 | 0.829 | -1.365 | 0.544 | 1.057 | | | | | | | | | | | |
| B7L74_10090 | ABC transporter ATP-binding protein | Transporter | 0.438 | -0.281 | -0.823 | -0.094 | -1.287 | -1.410 | 0.152 | -0.015 | -0.407 | 0.122 | -0.026 | | | | | | | | | | | |
| B7L74_10095 | hypothetical membrane spanning protein | | 1.396 | 0.848 | -0.034 | 1.333 | 5.626 | 5.259 | 8.502 | 1.496 | 7.532 | 6.193 | 0.078 | Late up | | | | | | | | | | |
| B7L74_10100 | phosphosulfolactate synthase | Lyase | 1.155 | -0.605 | 1.119 | 2.149 | -0.309 | 0.476 | 1.746 | 0.363 | 0.880 | 2.120 | 2.016 | Early up | | | | | | | | | | |
| B7L74_10105 | NAD(P) transhydrogenase subunit alpha | | -0.891 | 0.347 | -3.352 | 0.986 | 0.546 | 0.147 | -1.890 | -2.022 | -1.061 | -2.734 | -0.871 | | | | | | | | | | | |
| B7L74_10110 | DUF1658 domain-containing protein | | -0.381 | -0.255 | -1.354 | -0.707 | -0.425 | -0.365 | -0.377 | -1.015 | -1.111 | -0.967 | -0.889 | | | | | | | | | | | |
| B7L74_10115 | DUF1658 domain-containing protein | | 0.213 | 0.604 | -1.859 | 0.084 | 0.298 | 0.589 | -0.491 | -0.961 | -0.865 | -0.480 | -0.156 | | | | | | | | | | | |
| B7L74_10120 | NAD(P) transhydrogenase subunit alpha | | 1.058 | 0.389 | -3.444 | 1.591 | -1.050 | -1.101 | 0.891 | 0.847 | -0.665 | 0.886 | 1.298 | | | | | | | | | | | |
| B7L74_10125 | NAD(P) transhydrogenase subunit beta | Effector protein | -1.065 | -1.728 | -2.545 | -4.602 | 0.112 | 0.989 | 0.027 | -0.918 | 0.136 | -1.156 | -3.261 | | | Yes | | | | | | | | |
| B7L74_10130 | hypothetical protein | | 0.096 | 0.134 | 0.709 | -0.153 | -0.786 | -0.398 | 0.830 | 0.830 | -0.673 | 2.051 | 1.738 | | | | | | | | | | | |
| B7L74_10135 | hypothetical protein | | -0.351 | 0.342 | -1.748 | -0.545 | 0.122 | 1.662 | -0.563 | 1.286 | 2.640 | -0.540 | -0.008 | | | | | | | | | | | |
| B7L74_10140 | 16S rRNA (uracil(1498)-N(3))-methyltransferase | | -1.335 | -0.577 | -1.459 | -3.174 | 1.312 | 0.202 | -0.898 | 0.071 | 1.203 | -1.756 | -0.194 | | | | | | | | | | | |
| B7L74_10145 | hypothetical protein | | 0.944 | -0.267 | 1.660 | -0.141 | 0.178 | 0.737 | 0.556 | -0.648 | 0.172 | 1.233 | 1.669 | | | | | | | | | | | |
| B7L74_10150 | IS110 family transposase | Transposases harbored by IS1111 | -1.335 | -0.577 | -1.459 | -3.174 | 1.312 | 0.202 | -0.898 | 0.071 | 1.203 | 0.985 | 0.753 | | | | | | | | | | | |
| B7L74_10155 | hypothetical protein | | 0.704 | 0.222 | -0.102 | 0.354 | 0.380 | 0.014 | 0.997 | 0.481 | 0.060 | 1.025 | 1.116 | | | | | | | | | | | |
| B7L74_10160 | protein-(glutamine-N5) methyltransferase%2C release | | -1.521 | -6.274 | -0.466 | 0.606 | -1.170 | 0.454 | -2.147 | -2.584 | -1.471 | -1.261 | -1.421 | | | | | | | | | | | |
| B7L74_10165 | peptide chain release factor 1 | | -0.623 | -0.616 | -0.532 | 0.065 | -1.921 | -0.360 | -0.416 | -0.799 | -0.625 | -0.083 | 0.190 | | | | | | | | | | | |
| B7L74_10170 | glutamyl-tRNA reductase | Porphyrin and chlorophyll metabol | -0.951 | 1.934 | -0.060 | 3.858 | 1.544 | 1.730 | 2.476 | 1.232 | 1.118 | -0.148 | 1.649 | Up in P12 | | | | | | | | | | |
| B7L74_10175 | Bcr/CflA family drug resistance efflux transporter | Transporter | 5.057 | 1.681 | 2.806 | 0.261 | 5.978 | 6.890 | 2.768 | 0.898 | 5.474 | 3.176 | 2.118 | Continuous up | | | | | | | | Yes (drug resistance) | | |
| B7L74_10180 | diguanylate cyclase | Folate biosynthesis | -0.724 | -1.469 | -7.315 | 0.304 | -5.935 | -2.063 | -1.621 | -2.121 | -6.154 | -7.418 | -3.046 | Early down | | | | | | | | | | |
| B7L74_10185 | RNA polymerase-binding protein DksA | | 0.139 | -0.802 | -4.734 | 0.254 | -3.376 | -0.952 | -1.433 | 0.056 | -3.569 | -4.687 | -3.149 | | | | | | | | | | | |
| B7L74_10190 | diaminopimelate epimerase | | 0.061 | -0.347 | 0.460 | 2.155 | -0.304 | -0.656 | 0.522 | -1.107 | -1.840 | 0.890 | 1.442 | | | | | | | | | | | |
| B7L74_10195 | hypothetical protein | | 0.041 | 0.838 | -0.034 | -1.805 | 1.973 | 1.621 | 0.448 | 1.487 | 2.548 | -1.279 | -0.910 | | | | | | | | | | | |
| B7L74_10200 | LuxR family transcriptional regulator | Transcriptional regulators | -0.403 | -1.309 | -2.176 | -0.275 | 0.472 | 1.143 | -0.699 | 0.432 | -0.169 | 0.468 | -0.343 | | | | | | | | | | | |
| B7L74_10205 | hypothetical cytosolic protein | | 0.286 | -3.964 | -0.082 | 0.159 | 1.198 | 0.236 | 1.645 | 0.281 | -2.183 | -0.312 | -0.273 | | | | | | | | | | | |
| B7L74_10210 | hypothetical protein | | 0.158 | -0.074 | 0.309 | 1.545 | -0.020 | -0.528 | 0.209 | -1.581 | -4.374 | 0.983 | 1.009 | Down in P51 | | | | | | | | | | |
| B7L74_10215 | phospholipase/carboxylesterase | | 0.828 | 1.706 | 1.654 | -0.036 | 1.044 | 1.797 | 1.322 | 2.038 | 1.981 | 0.580 | 1.198 | | | | | | | | | | | |
| B7L74_10220 | mannose-1-phosphate guanylyltransferase | Effector protein / Transferase | 0.550 | 0.841 | -0.034 | 0.480 | 1.973 | 1.074 | 0.464 | 1.490 | 2.215 | -0.842 | 0.369 | | | Yes | | | | | | | | |
| B7L74_10225 | phosphotransferase | Amino sugar and nucleotide sugar | -0.014 | -0.698 | -0.560 | 0.702 | 1.574 | 3.758 | -0.739 | -0.457 | 1.523 | -0.056 | 0.189 | Up in P21 | | | | | | | | | | |
| B7L74_10230 | LPS biosynthesis protein | | 1.848 | -0.444 | 2.070 | 1.445 | 1.009 | -2.647 | 0.908 | -0.925 | -0.647 | 0.393 | 0.640 | | | | | | | | | Yes (Outer membrane/LPS) | | |
| B7L74_10235 | peptidylprolyl isomerase | Chaperones and folding catalysts | -1.693 | -0.699 | -0.551 | 0.003 | -0.716 | -0.656 | -1.213 | -1.523 | -2.613 | -1.689 | -1.285 | Late down | | | Yes | | | | | | | |
| B7L74_10240 | 4-hydroxythreonine-4-phosphate dehydrogenase PdxA | | 2.042 | 0.839 | -0.034 | -1.244 | 2.649 | 2.571 | 1.040 | 1.861 | 2.554 | 0.163 | 0.133 | | | | | | | | | | | |
| B7L74_10245 | ribosomal RNA small subunit methyltransferase A | | -0.220 | -0.536 | 1.167 | 1.540 | -0.164 | -0.389 | 0.651 | -0.372 | -0.719 | 1.161 | 0.703 | | | | | | | | | | | |
| B7L74_10250 | hypothetical protein | | 0.478 | 0.255 | 1.179 | 1.181 | 0.398 | -0.169 | 1.365 | -0.510 | -0.110 | 1.003 | 1.718 | | | | | | | | | | | |
| B7L74_10255 | universal stress protein A | | 0.326 | 0.230 | 1.068 | 1.857 | 0.197 | 0.182 | 1.014 | -0.203 | 0.289 | 1.381 | 1.878 | | | | | | | | | | | |
| B7L74_10260 | hypothetical exported protein | Exported | -0.375 | -0.815 | 0.133 | 3.390 | 0.025 | -0.740 | 0.599 | -0.662 | -1.833 | 2.006 | 0.502 | Up in P12 | | | | | | | | | | |
| B7L74_10265 | hypothetical exported protein | Exported | 0.914 | 0.229 | 2.104 | 2.040 | 0.368 | 0.415 | 1.036 | 1.171 | -0.069 | 2.339 | 1.897 | Early up | | | | | | | | | | |
| B7L74_10270 | peptidase U7 | | 1.152 | -0.243 | 0.772 | 0.880 | 1.007 | 0.612 | 1.122 | -1.016 | 0.189 | 1.109 | 0.989 | | | | | | | | | | | |
| B7L74_10275 | bis(5'-nucleosyl)-tetraphosphatase (symmetrical) | Eucaryotic-like ORFs. | 0.816 | 0.165 | 0.519 | 1.294 | -1.553 | 0.342 | 0.805 | 1.148 | 0.904 | 0.749 | 1.107 | | | | | | | | | | | |
| B7L74_10280 | hypothetical protein | | -0.060 | 2.353 | -1.493 | 1.043 | 1.279 | 1.131 | 0.041 | 2.574 | 2.850 | 0.686 | 1.946 | | | | | | | | | | | |
| B7L74_10285 | hypothetical protein | | 0.724 | 1.279 | 0.686 | -0.899 | 0.350 | 1.568 | 0.325 | 1.867 | 1.618 | 1.495 | 1.491 | | | | | | | | | | | |
| B7L74_10290 | IS110 family transposase | | 0.219 | -0.910 | 2.481 | 0.023 | -0.842 | -0.010 | 1.123 | -0.467 | -0.666 | 1.240 | 1.530 | | | | | | | | | | | |
| B7L74_10295 | hypothetical protein | AnkM. Eucaryotic-like ORFs. | 0.090 | 2.464 | -0.034 | 0.381 | 6.375 | 5.062 | 0.527 | 2.458 | 5.515 | -1.252 | -0.897 | | | | | | | | | | | |
| B7L74_10300 | translation repressor RelB | Prokaryotic defense system | -0.253 | -0.496 | 0.761 | 0.528 | -0.620 | -0.630 | -0.142 | -0.365 | -2.030 | 0.386 | 0.384 | Down in P51 | | | | | | | | | | |
| B7L74_10305 | Txe/YoeB family addiction module toxin | | 0.096 | 2.752 | -0.921 | -1.544 | 2.805 | 1.962 | 1.551 | 2.051 | 2.208 | -1.482 | -0.762 | | | | | | | | | | | |
| B7L74_10310 | dihydrofolate reductase | | 2.837 | 9.292 | -0.036 | 7.439 | 9.540 | 8.033 | 7.808 | 1.495 | 8.177 | 3.086 | -0.898 | | | | | | | | | | | |
| B7L74_10315 | hypothetical cytosolic protein | | 4.623 | 8.785 | -0.035 | 2.319 | 8.901 | 7.770 | 7.610 | 1.495 | 7.317 | 3.085 | -0.898 | | | | | | | | | | | |
| B7L74_10320 | acylphosphatase | | 0.712 | 0.656 | 2.247 | 1.330 | 0.771 | 0.325 | 0.646 | 1.115 | 0.003 | 0.425 | 0.699 | | | | | | | | | | | |
| B7L74_10325 | D-glycero-beta-D-manno-heptose-1%2C7-bisphosphat | Lipopolysaccharide biosynthesis | 0.320 | -1.628 | -1.498 | 0.639 | 0.081 | -0.845 | -0.727 | -0.979 | -0.579 | -0.250 | -0.234 | | | | | | | | | | Yes | |
| B7L74_10330 | methionyl-tRNA formyltransferase | | -0.385 | -0.319 | -0.140 | 0.344 | -0.533 | -0.258 | -0.440 | -0.590 | -0.907 | -0.306 | -0.015 | | | | | | | | | | | |
| B7L74_10335 | DNA protecting protein DprA | DNA | 0.764 | -0.306 | 0.758 | 0.579 | -0.853 | -0.293 | 0.633 | 0.052 | 0.136 | 1.469 | 1.362 | | | | | | | | | | | |
| B7L74_10340 | hypothetical protein protein=Smg protein | | -0.459 | 3.449 | -1.053 | -2.413 | 4.557 | 0.990 | 2.622 | 3.682 | 3.559 | 0.754 | -1.570 | | | | | | | | | | | |
| B7L74_10345 | DNA topoisomerase I | DNA | -1.129 | -1.444 | -2.016 | -0.473 | -1.468 | -1.423 | -0.785 | -1.213 | -1.934 | -0.779 | -0.070 | | | | | | | | | | | |
| B7L74_10350 | hypothetical protein | | -0.377 | -0.172 | -1.926 | -2.020 | -0.061 | -0.636 | -1.408 | -1.305 | -2.856 | -3.170 | -3.312 | Continuous down | | | | | | | | | | |
| B7L74_10355 | 5-(carboxyamino)imidazole ribonucleotide mutase | | -0.651 | -1.167 | -0.062 | -0.267 | -1.667 | -0.872 | -0.648 | -0.680 | -2.621 | -0.739 | -0.647 | | | | | | | | | | | |
| B7L74_10360 | threonylcarbamoyl-AMP synthase | | -0.308 | -0.769 | 1.118 | 0.718 | -1.472 | -0.419 | -0.075 | 0.014 | -1.141 | 0.541 | 0.274 | | | | | | | | | | | |
| B7L74_10365 | DUF1658 domain-containing protein | | 0.125 | 0.657 | 0.368 | -1.023 | 0.782 | 0.492 | -0.177 | 0.261 | -0.618 | -1.310 | -0.843 | | | | | | | | | | | |
| B7L74_10370 | DUF1658 domain-containing protein | | 0.478 | 0.221 | 0.443 | -0.447 | 0.270 | 0.761 | 0.120 | 1.211 | -0.198 | -0.008 | 0.318 | | | | | | | | | | | |
| B7L74_10375 | hypothetical protein | | 0.084 | 0.847 | -0.034 | -1.767 | 1.206 | 1.626 | -0.046 | 0.897 | 1.171 | -1.256 | -0.897 | | | | | | | | | | | |
| B7L74_10380 | ADP-heptose--LPS heptosyltransferase | | -0.875 | -1.300 | -0.996 | 0.475 | -0.754 | -0.571 | -0.017 | -1.474 | -1.307 | -0.491 | 0.773 | | | | | | | | | | | |
| B7L74_10385 | two-component sensor histidine kinase, RstB | RstB-like system two-comp system | -0.431 | -0.679 | -0.682 | 0.581 | -0.455 | -0.960 | -0.313 | 0.181 | -0.386 | -0.296 | 0.079 | | | | | | | | | | | |
| B7L74_10390 | DNA-binding response regulator(OmpR in RightonQ | Intracellular growthRstB-like syste | -0.666 | 1.157 | 1.655 | 1.523 | 0.162 | 0.924 | 0.342 | 1.395 | 0.449 | -3.255 | -0.312 | | | | | | | | | | | |
| B7L74_10395 | hypothetical protein | Effector protein | 4.426 | 1.445 | 1.662 | 0.777 | 5.627 | 5.488 | 2.341 | 1.391 | 4.462 | 2.744 | 2.143 | Early up | | Yes | | | | | | | | |
| B7L74_10400 | arginine--tRNA ligase | Aminoacyl-tRNA synthase | -1.427 | -1.632 | -1.993 | -0.082 | -2.126 | -2.562 | -1.343 | -1.787 | -2.130 | -1.149 | -0.978 | Early down | | | | | | | | | | |
| B7L74_10405 | hypothetical protein | | -1.988 | -0.347 | -1.600 | -0.918 | -1.193 | 0.305 | -1.342 | -0.831 | -1.316 | -0.757 | -0.746 | | | | | | | | | | | |
| B7L74_10410 | hypothetical exported protein | | 0.553 | 0.843 | -0.034 | 1.591 | 4.631 | 2.339 | 1.349 | 1.492 | 3.181 | 1.572 | 2.696 | | | | | | | | | | | |
| B7L74_10415 | ATP-dependent protease subunit HslV | Peptidases and inhibitors | -1.596 | -3.564 | -1.695 | -0.594 | -2.538 | -2.326 | -1.008 | -4.013 | -4.389 | -0.485 | -0.730 | Continuous down | | | | | | | | | | |
| B7L74_10420 | ATP-dependent protease ATP-binding subunit HslU | Chaperones and folding catalysts | -0.722 | -1.441 | -1.504 | 0.263 | -1.894 | -2.093 | -1.237 | -2.942 | -3.966 | -1.101 | -1.035 | Continuous down | | | Yes | | | | | | | |
| B7L74_10425 | hypothetical protein | Effector protein | -12.577 | -11.443 | -8.136 | 9.151 | 16.858 | 16.509 | 18.556 | 16.917 | 18.165 | -18.185 | -17.650 | | | Yes | | | | | | | | |
| B7L74_10430 | DUF421 domain-containing protein | | -0.584 | 1.425 | -0.620 | 0.131 | 1.500 | 0.803 | -0.428 | 2.751 | 2.261 | 0.987 | 0.456 | | | | | | | | | | | |
| B7L74_10435 | MFS transporter | Transporter | 0.835 | 0.866 | 1.460 | -0.053 | 1.240 | 1.011 | 0.429 | 0.479 | 0.250 | -0.413 | -0.922 | | | | | | | | | Yes (oligopeptide) | | |
| B7L74_10440 | hypothetical cytosolic protein | Effector protein | 0.085 | 0.765 | 0.655 | 1.892 | 0.446 | -0.392 | -0.134 | 0.129 | -1.405 | 1.653 | 0.704 | | | Yes | | | | | | | | |
| B7L74_10445 | ubiquinone/menaquinone biosynthesis C-methyltransfe | | 1.046 | 0.038 | 0.562 | 0.809 | -0.346 | -0.872 | 0.373 | -0.410 | -0.850 | 1.229 | 0.698 | | | | | | | | | | | |
| B7L74_10450 | hypothetical protein | | -0.490 | -0.350 | 0.157 | -1.334 | -1.618 | -1.462 | 0.215 | 0.211 | -0.977 | -0.577 | -0.768 | | | | | | | | | | | |
| B7L74_10455 | ubiquinone biosynthesis regulatory protein kinase UbiB | Kinase in Cofactor metabolism | 0.034 | 2.541 | 2.806 | 2.670 | 3.289 | 3.991 | 3.710 | 3.105 | 5.134 | 3.949 | 4.800 | Continuous up | | | | | | | | | | |
| B7L74_10460 | DUF1658 domain-containing protein | | 5.075 | 3.634 | -0.609 | -0.943 | 2.781 | 2.839 | 1.448 | 0.919 | 2.009 | -0.814 | 0.005 | | | | | | | | | | | |
| B7L74_10465 | DUF1658 domain-containing protein | CB specific protein family | 1.529 | 0.766 | -1.099 | -1.655 | 0.317 | -0.275 | -0.852 | -1.062 | -3.096 | -1.693 | -0.968 | Down in P51 | | | | | | | | | | |
| B7L74_10470 | amino acid permease | Transporter | 1.788 | 0.107 | 0.752 | 0.517 | 1.516 | 1.701 | 0.831 | 0.472 | 2.632 | 1.516 | 1.434 | | | | | | | | | | | |
| B7L74_10475 | ATPase | ATP | -0.772 | -0.732 | -0.516 | -1.369 | -0.054 | -0.851 | -0.481 | -0.281 | -0.522 | -0.480 | -0.488 | | | | | | | | | | | |
| B7L74_10480 | ATP-dependent protease | Protein processing | 0.988 | 0.664 | 3.177 | 2.291 | 0.665 | 0.107 | 2.380 | 0.703 | 1.908 | 1.584 | 3.530 | Early up | | | | | | | | | | |
| B7L74_10485 | hypothetical protein | | 0.599 | 1.060 | 3.507 | 2.041 | 2.042 | 0.962 | 4.249 | 1.879 | 4.432 | 2.362 | 3.240 | Continuous up | | | | | | | | | | |
| B7L74_10490 | cystathionine beta-synthase | | 1.603 | -0.932 | 1.101 | 1.343 | 1.333 | 2.046 | 1.857 | -0.059 | 0.887 | 1.516 | 1.300 | | | | | | | | | | | |
| B7L74_10495 | cystathionine beta-lyase, MetC | | -0.267 | -0.252 | -0.267 | -0.524 | -0.646 | -1.265 | 0.480 | -0.012 | 0.092 | 0.199 | 0.180 | | | | | | | | | | | |
| B7L74_10500 | hypothetical protein | | 0.026 | -0.835 | 0.229 | 1.602 | -1.203 | -1.351 | 0.488 | -0.030 | -1.622 | 0.841 | 1.265 | | | | | | | | | | | |
| B7L74_10505 | hypothetical exported protein | | -0.469 | -0.587 | -0.283 | 0.684 | 0.284 | -1.520 | 0.037 | 0.727 | 0.055 | 0.609 | 0.024 | | | | | | | | | | | |
| B7L74_10510 | hypothetical protein | Effector protein | 1.985 | 0.068 | 0.281 | 0.729 | 0.743 | 1.196 | 2.234 | 1.562 | -0.575 | 0.401 | 0.469 | | | Yes | | | | | | | | |
| B7L74_10515 | SH3 domain-containing protein | | -1.340 | 0.622 | 1.238 | 0.582 | 0.474 | 0.336 | -0.924 | 0.043 | -1.993 | -1.134 | -1.007 | | | | | | | | | | | |
| B7L74_10520 | S-adenosylmethionine synthase | Cysteine and methionine metabolis | 0.294 | -0.598 | -0.553 | 1.018 | -1.035 | -1.726 | -0.257 | -0.768 | -2.841 | 0.408 | 0.285 | Early down | | | | | | | | | | |
| B7L74_10525 | adenosylhomocysteinase | Cysteine and methionine metabolis | -0.691 | -1.190 | -2.332 | -0.200 | -1.428 | -2.128 | -0.904 | -1.353 | -2.360 | -1.058 | -0.911 | Early down | | | | | | | | | | |
| B7L74_10530 | sensor domain-containing diguanylate cyclase | | -0.467 | -0.796 | -0.079 | 0.934 | -0.247 | -1.635 | -0.597 | 0.258 | -0.650 | 0.611 | -0.054 | | | | | | | | | | | |
| B7L74_10535 | hypothetical cytosolic protein | | 0.559 | 0.847 | -0.034 | -1.764 | 3.359 | 0.967 | 1.819 | 1.496 | 0.186 | -1.254 | -0.418 | | | | | | | | | | | |
| B7L74_10540 | hypothetical protein | | 1.207 | 1.572 | 3.548 | -0.768 | 1.232 | 0.813 | 2.697 | 1.997 | 2.814 | 1.181 | 3.261 | Early up | | | | | | | | | | |
| B7L74_10545 | hypothetical protein | | 1.606 | 0.918 | 3.318 | 0.771 | 1.646 | 0.766 | 3.592 | 2.310 | 3.635 | 1.790 | 5.608 | | | | | | | | | | | |
| B7L74_10550 | hypothetical protein | | 0.979 | 1.134 | 2.026 | -0.601 | 1.625 | 0.840 | 1.414 | 1.052 | 1.799 | 0.430 | 0.802 | | | | | | | | | | | |
| B7L74_10555 | hypothetical protein | | 5.048 | -1.680 | 2.034 | 2.369 | 1.153 | -1.485 | 2.072 | 2.770 | 3.099 | 4.584 | 3.174 | | | | | | | | | Yes (toxins) | | |
| B7L74_10560 | peptide ABC transporter ATP-binding protein | Transporter | 0.174 | -0.596 | 1.333 | 0.094 | -0.214 | 0.232 | 1.795 | 1.640 | 0.343 | 1.558 | 1.981 | | | | | | | | | | | |
| B7L74_10565 | hypothetical protein | | -0.515 | 0.266 | -0.610 | -2.364 | 1.485 | 1.451 | -0.108 | 0.915 | 3.443 | -1.844 | -0.095 | | | | | | | | | | | |
| B7L74_10570 | hypothetical protein | ABC transporters | 1.177 | 1.480 | 1.087 | 0.732 | 1.993 | 2.219 | 1.542 | 3.834 | 3.698 | 1.495 | 2.770 | Early up | | | | | | | | Yes (toxins) | | |
| B7L74_10575 | HlyD family type I secretion periplasmic adaptor subun | | 2.293 | 0.778 | 2.370 | 1.433 | 2.744 | 2.187 | 2.370 | 1.322 | 1.760 | -0.184 | 2.559 | | | | | | | | | Yes (Outer membrane/adhesin) | | |
| B7L74_10580 | diguanylate cyclase | Transcriptional regulators | 0.436 | -1.365 | -0.253 | 0.020 | -0.752 | 2.650 | -2.487 | 2.165 | 2.202 | 1.297 | 2.144 | | | | | | | | | | | |
| B7L74_10585 | GGDEF-domain containing protein | | 2.534 | 2.265 | 3.296 | 0.908 | 2.418 | 1.435 | 3.009 | 1.653 | 5.254 | -1.501 | 0.099 | | | | | | | | | | | |
| B7L74_10590 | hypothetical protein | | 10.453 | 4.313 | 5.404 | -0.277 | 10.476 | 10.685 | 6.999 | 3.867 | 9.589 | 6.130 | 3.447 | Early up | | | | | | | | | | |
| B7L74_10595 | chitinase | | 1.308 | -0.308 | 2.162 | 0.965 | 1.226 | 1.523 | 0.876 | 3.123 | 3.195 | 2.075 | 2.745 | Late up | | | | | | | | | | |
| B7L74_10600 | hypothetical protein | | 0.125 | 1.738 | -2.477 | -1.098 | 0.294 | 1.555 | 0.544 | 2.075 | 0.185 | -0.777 | -0.156 | | | | | | | | | | | |
| B7L74_10605 | methylenetetrahydrofolate reductase [NAD(P)H] | | 1.706 | 1.300 | 0.167 | 0.159 | 0.372 | -0.075 | 0.309 | 0.278 | -0.950 | 0.780 | 0.337 | | | | | | | | | | | |
| B7L74_10610 | 5-methyltetrahydropteroyltriglutamate--homocysteine m | | -0.882 | 0.102 | -1.018 | -0.404 | -0.399 | -0.351 | -1.207 | 0.148 | -0.433 | -1.089 | -0.923 | | | | | | | | | | | |
| B7L74_10615 | tryptophan--tRNA ligase | Aminoacyl-tRNA synthase | 0.889 | 1.325 | 2.231 | 1.654 | 0.483 | 0.082 | 1.348 | 1.145 | 0.285 | 2.050 | 2.030 | Late up | | | | | | | | | | |
| B7L74_10620 | hypothetical protein | | 0.151 | 0.721 | 0.839 | 0.005 | -0.529 | -0.388 | 0.266 | -1.154 | -0.239 | -0.200 | 0.411 | | | | | | | | | | | |
| B7L74_10625 | hypothetical protein | | 2.068 | 2.396 | 2.628 | 0.338 | 0.387 | 3.607 | 1.904 | 3.455 | 3.462 | 2.172 | 2.661 | Early up | | | | | | | | | | |
| B7L74_10630 | hypothetical protein | | 2.466 | 1.828 | 0.950 | 2.633 | 0.279 | 3.392 | 1.617 | 2.384 | 5.116 | 2.590 | 1.519 | | | | | | | | | | | |
| B7L74_10635 | hypothetical protein(cirD in Table 1 of RightonQ pape | Effector protein | 0.343 | 0.476 | -0.569 | -0.250 | 0.042 | 0.422 | -0.665 | -0.333 | 0.526 | -0.169 | -0.533 | | | Yes | | | | | | | | |
| B7L74_10640 | DNA-dependent helicase II | | -0.227 | -0.828 | -0.939 | 0.216 | -1.532 | -0.764 | -0.103 | -1.242 | -1.157 | 0.129 | 0.364 | | | | | | | | | | | |
| B7L74_10645 | zinc transporter | Transporter | 0.901 | 0.837 | -0.034 | -1.810 | 1.973 | 1.620 | 0.443 | 1.862 | 2.543 | -1.282 | -0.912 | | | | | | | | | Yes (Ions/Zinc) | | |
| B7L74_10650 | hypothetical protein | Effector protein | 1.006 | 0.495 | 1.254 | 2.643 | 0.532 | 0.472 | 1.006 | 0.211 | -0.767 | 2.000 | 2.003 | Up in P12 | | Yes | | | | | | | | |
| B7L74_10655 | hypothetical cytosolic protein | | -0.605 | -0.008 | 2.041 | 2.002 | 0.290 | 0.194 | 1.985 | 0.154 | 1.947 | 1.440 | 1.732 | | | | | | | | | | | |
| B7L74_10660 | hypothetical protein | | -0.098 | -0.201 | 0.308 | -0.023 | -0.166 | -0.818 | -0.022 | -1.106 | -0.745 | -0.254 | 0.203 | | | | | | | | | | | |
| B7L74_10665 | proline/betaine transporter | | 0.239 | -0.781 | 0.527 | -0.452 | -0.070 | 0.296 | 0.197 | 0.669 | 0.943 | 0.800 | 1.159 | | | | | | | | | | | |
| B7L74_10670 | hypothetical protein | Effector protein | 0.752 | 0.231 | 2.502 | 1.343 | 0.382 | -0.474 | 1.232 | 1.566 | 1.399 | 2.462 | 2.761 | Late up | | Yes | | | | | | | | |
| B7L74_10675 | hypothetical protein | | -0.266 | 0.611 | 0.141 | 1.698 | 1.463 | 0.359 | -0.117 | 0.531 | 0.284 | 1.289 | 1.211 | | | | | | | | | | | |
| B7L74_10680 | hypothetical protein | | -0.150 | -0.170 | 0.531 | 1.257 | 1.589 | 1.497 | -1.982 | 2.391 | 0.155 | 0.764 | 0.204 | | | | | | | | | | | |
| B7L74_10685 | hypothetical protein | | 1.469 | -0.123 | -0.461 | 1.811 | -1.212 | 0.509 | 0.886 | 1.311 | 1.299 | 1.388 | 1.493 | | | | | | | | | | | |
| B7L74_10690 | hypothetical protein | | 1.545 | 1.118 | 1.112 | 0.562 | 3.013 | 2.482 | 0.368 | 2.281 | 3.898 | 1.337 | 1.077 | | | | | | | | | | | |
| B7L74_10695 | hypothetical protein | | 0.546 | 0.838 | 0.966 | -1.805 | 2.642 | 2.026 | 0.448 | 1.487 | 1.137 | -0.426 | -0.910 | | | | | | | | | | | |
| B7L74_10700 | hypothetical exported protein | | 0.369 | -0.527 | -1.334 | -0.084 | 0.183 | 0.321 | -0.392 | -1.381 | -0.100 | 0.279 | 0.660 | | | | | | | | | | | |
| B7L74_10705 | permease | | 0.038 | 0.837 | -0.034 | -1.261 | 2.995 | 1.620 | 0.445 | 1.486 | 2.545 | -1.281 | -0.446 | | | | | | | | | | | |
| B7L74_10710 | MFS transporter | Transporter | 2.384 | -0.613 | 0.235 | -0.025 | 3.473 | 4.196 | 0.366 | -1.449 | 3.158 | 1.225 | 0.610 | | | | | | | | | | | |
| B7L74_10715 | MFS transporter | Transporter | 0.780 | -1.092 | -2.361 | 2.385 | -0.095 | -0.674 | 0.107 | -1.131 | -1.314 | 1.480 | -0.163 | | | | | | | | | | | |
| B7L74_10720 | thymidine kinase | | -0.712 | -0.266 | 0.314 | 0.667 | -0.606 | -0.808 | -0.296 | 0.075 | -0.241 | -0.002 | 0.286 | | | | | | | | | | | |
| B7L74_10725 | chorismate mutase | | -0.144 | -0.225 | -0.583 | 0.922 | -1.109 | -0.962 | -0.435 | -1.381 | -2.692 | -0.083 | -0.078 | Down in P51 | | | | | | | | | | |
| B7L74_10730 | hypothetical membrane spanning protein | | 0.193 | -0.012 | 1.324 | -0.476 | 0.010 | -0.214 | 0.350 | -0.447 | 1.207 | 0.818 | 0.163 | | | | | | | | | | | |
| B7L74_10735 | hypothetical exported protein | | -0.372 | -0.428 | 0.836 | 0.736 | -1.132 | -1.029 | 0.232 | -0.045 | 0.127 | 0.964 | 1.166 | | | | | | | | | | | |
| B7L74_10740 | acyltransferase | | -1.477 | -1.370 | -1.563 | -1.049 | -2.118 | -2.433 | -1.065 | -1.887 | -3.387 | -1.122 | -0.978 | Early down | | | | | | | | | | |
| B7L74_10745 | hydroxymethylbilane synthase | | 1.116 | -0.704 | -0.412 | 0.037 | 2.231 | 2.750 | -0.529 | -2.223 | -1.391 | 0.204 | -0.420 | | | | | | | | | | | |
| B7L74_10750 | 3-dehydroquinase | Phenylalanine, tyrosine and tryptop | 3.878 | 2.728 | 1.550 | 1.540 | 6.798 | 6.931 | 3.166 | 1.924 | 8.263 | 4.794 | 3.975 | Early up | | | | | | | | | | |
| B7L74_10755 | NGG1p interacting factor NIF3 | Effector protein | -1.597 | -1.078 | -1.335 | -0.479 | -2.289 | -1.951 | -1.761 | -1.461 | -2.664 | -1.239 | -1.462 | | | Yes | | | | | | | | |
| B7L74_10760 | uroporphyrinogen-III synthase | | 0.897 | 0.376 | -1.035 | -1.381 | 1.512 | 3.009 | 2.415 | 3.304 | 2.994 | 1.848 | 0.019 | | | | | | | | | | | |
| B7L74_10765 | cell filamentation protein Fic | Effector protein | -0.548 | -0.096 | -0.159 | 0.261 | -0.214 | -0.135 | 0.080 | 0.581 | -0.551 | 0.526 | 0.381 | | | Yes | | | | | | | | |
| B7L74_10770 | hypothetical protein | | 0.805 | 1.189 | 3.183 | -1.222 | 0.662 | 0.303 | 1.198 | 0.811 | 1.140 | 0.876 | 0.688 | | | | | | | | | | | |
| B7L74_10775 | hypothetical protein uroporphyrin-III C-methyltransfera | | 0.164 | -0.119 | 0.395 | -0.290 | -0.317 | -0.097 | -0.338 | -1.331 | -0.708 | -0.255 | -0.558 | | | | | | | | | | | |
| B7L74_10780 | heme biosynthesis protein HemY | Cofactor metabolism | 2.453 | -0.073 | -0.319 | -1.717 | 3.449 | 3.408 | 0.431 | -1.258 | 2.581 | 0.275 | -0.649 | Early up | | | | | | | | | | |
| B7L74_10785 | hypothetical exported protein | Effector protein | 5.168 | 1.613 | 3.807 | 0.784 | 6.165 | 5.990 | 3.740 | 1.652 | 5.051 | 3.713 | 3.595 | Early up | | Yes | | | | | | | | |
| B7L74_10790 | 7-cyano-7-deazaguanine synthase | | 0.116 | -0.400 | 0.251 | 0.574 | -1.076 | -0.040 | 0.164 | -0.235 | -1.457 | 0.304 | 0.204 | | | | | | | | | | | |
| B7L74_10795 | hypothetical protein | | 4.633 | 0.623 | 0.171 | -0.781 | 4.632 | 5.716 | 0.748 | -0.477 | 4.079 | 0.205 | 0.147 | Early up | | | | | | | | | | |
| B7L74_10800 | hypothetical protein | | 2.386 | -0.040 | 0.962 | 1.913 | 3.310 | 3.269 | 1.246 | 0.875 | 3.104 | 3.588 | 2.768 | Early up | | | | | | | | | | |
| B7L74_10805 | hypothetical protein | | 1.660 | -0.228 | 1.662 | 2.097 | 2.647 | 2.433 | 1.222 | 0.691 | 2.734 | 3.340 | 2.662 | Late up | | | | | | | | | | |
| B7L74_10810 | transcription termination factor Rho | RNA degradation | -1.629 | -0.933 | -1.720 | 0.903 | -2.333 | -2.419 | -1.590 | -1.054 | -1.541 | 0.016 | -0.075 | Early down | | | | | | | | | | |
| B7L74_10815 | thiol reductase thioredoxin | PV detoxification | -0.247 | 0.119 | 2.871 | 0.699 | -0.724 | -0.785 | 1.108 | -0.124 | 0.771 | 0.528 | 0.807 | | | | | | | | | | | |
| B7L74_10820 | sodium-dependent transporter | | -0.314 | -0.142 | -0.387 | -1.183 | -0.194 | -0.268 | -0.945 | -0.885 | 0.021 | -0.323 | -0.451 | | | | | | | | | Yes (Na+ symporter) | | |
| B7L74_10825 | YggT family protein | Secretion system | -2.319 | -1.489 | -3.773 | -1.402 | -3.010 | -2.989 | -2.300 | -2.994 | -3.586 | -2.483 | -2.469 | Early down | | | | | | | | | | |
| B7L74_10830 | pyrroline-5-carboxylate reductase | Arginine and proline metabolism | -2.815 | -1.853 | -4.438 | -1.325 | -2.685 | -2.499 | -3.418 | -3.734 | -4.397 | -3.333 | -2.930 | Continuous down | | | | | | | | | | |
| B7L74_10835 | YggS family pyridoxal phosphate enzyme | Amino acid metabolism | -1.398 | -1.499 | -3.040 | 0.357 | -1.777 | -3.011 | -2.016 | -2.389 | -4.265 | -1.845 | -1.722 | Early down | | | | | | | | | | |
| B7L74_10840 | phosphoenolpyruvate carboxykinase (ATP) | Glycolysis / Gluconeogenesis | -0.993 | -0.817 | -3.126 | 1.108 | -2.253 | -1.016 | -1.483 | -1.984 | -3.552 | -1.039 | -0.843 | Early down | | | | Yes | Yes | | | | | |
| B7L74_10845 | DUF179 domain-containing protein | Transcription factors | 0.785 | -0.773 | -1.443 | 0.025 | -3.346 | -3.330 | -1.463 | -2.278 | -4.900 | -1.670 | -1.373 | Early down | | | | | | | | | | |
| B7L74_10850 | Holliday junction resolvase RuvX | | -1.360 | -0.959 | -0.926 | -0.380 | -1.038 | -1.119 | -1.062 | -2.103 | -1.971 | -1.305 | -1.477 | Late down | | | | | | | | | | |
| B7L74_10855 | aspartate carbamoyltransferase | | 0.233 | 0.866 | 0.012 | 1.283 | 1.344 | 0.505 | 0.120 | 1.190 | 2.768 | 0.075 | 1.262 | | | | | | | | | | | |
| B7L74_10860 | transcriptional regulator | | -0.560 | -0.197 | 0.716 | -0.307 | -0.953 | -0.752 | -0.211 | -0.983 | 0.235 | 0.112 | 0.106 | | | | | | | | | | | |
| B7L74_10865 | mobile mystery protein B | | -0.057 | 0.767 | 1.394 | 1.177 | -0.282 | 0.592 | 1.529 | 1.920 | 1.750 | 1.747 | 1.217 | | | | | | | | | | | |
| B7L74_10870 | hypothetical protein | | 2.568 | -0.640 | 0.025 | -1.346 | 2.413 | 3.328 | 1.637 | -0.217 | 1.889 | -0.031 | -0.861 | | | | | | | | | | | |
| B7L74_10875 | integrase | | -0.464 | -0.317 | -0.595 | -0.564 | -0.888 | -1.373 | -0.525 | -0.111 | -1.633 | -0.483 | -0.738 | | | | | | | | | | | |
| B7L74_10880 | hypothetical protein | Effector protein | 0.109 | -0.050 | -0.541 | -0.111 | -0.742 | -1.178 | -0.662 | -0.787 | -1.320 | -0.221 | -0.261 | | | Yes | | | | | | | | |
| B7L74_10885 | hypothetical protein | | -0.081 | 1.886 | 0.326 | -1.177 | 0.737 | 0.750 | -1.876 | 1.580 | 1.017 | -1.363 | -1.580 | | | | | | | | | | | |
| B7L74_10890 | phage tail protein | | -0.415 | 0.580 | 0.686 | 1.672 | 0.030 | -0.693 | -0.744 | 2.213 | 1.017 | 0.435 | 1.477 | | | | | | | | | | | |
| B7L74_10895 | hypothetical protein | | 1.501 | 0.359 | -1.277 | 0.701 | -0.751 | 2.051 | -0.779 | 0.357 | 1.724 | -0.603 | 0.097 | | | | | | | | | | | |
| B7L74_10900 | hypothetical protein | | 1.916 | 0.442 | -0.164 | -0.517 | 4.130 | 2.310 | -0.033 | 0.151 | 1.651 | -0.123 | 0.060 | Up in P16 | | | | | | | | | | |
| B7L74_10905 | hypothetical protein | | -0.095 | 1.051 | 0.194 | 0.389 | -0.191 | -0.906 | 0.079 | 0.823 | 0.331 | -0.279 | -1.360 | | | | | | | | | | | |
| B7L74_10910 | hypothetical protein | | 0.805 | 0.434 | 1.638 | 1.368 | 0.508 | -0.532 | 0.965 | 1.096 | 0.948 | 1.236 | 0.983 | | | | | | | | | | | |
| B7L74_10915 | integrase | | -0.561 | 0.277 | -0.837 | -0.380 | 0.475 | 0.553 | -1.083 | -0.426 | 0.738 | -0.886 | -1.220 | | | | | | | | | | | |
| B7L74_10920 | hypothetical protein | | -1.024 | 0.706 | -0.652 | -0.232 | -0.465 | -0.224 | -2.051 | 0.447 | 0.508 | -0.221 | -0.903 | | | | | | | | | | | |
| B7L74_10925 | recombinase | | 2.165 | 2.576 | -0.035 | -1.750 | 4.401 | 2.589 | 1.469 | 2.458 | 2.628 | 1.252 | -0.897 | | | | | | | | | | | |
| B7L74_10930 | hypothetical protein | | -0.332 | -0.330 | -2.227 | -0.094 | -0.484 | -0.771 | -1.295 | -1.863 | -2.041 | -1.953 | -1.511 | Early down | | | | | | | | | | |
| B7L74_10935 | hypothetical protein | Effector protein | -0.845 | 0.431 | -1.060 | -0.364 | -0.355 | -0.356 | -1.119 | 0.498 | -1.879 | -1.921 | -1.618 | | | Yes | | | | | | | | |
| B7L74_10940 | hypothetical protein | Effector protein | 8.362 | 4.806 | 3.993 | 2.588 | 8.947 | 8.827 | 4.413 | 4.227 | 7.502 | 4.061 | 2.887 | Early up | | Yes | | | | | | | | |
| B7L74_10945 | hypothetical protein | Effector protein | 0.375 | 1.478 | 2.842 | -0.613 | 1.631 | 1.053 | 2.990 | 1.482 | 3.092 | 0.223 | 0.793 | Early up | | Yes | | | | | | | | |
| B7L74_10950 | protein CbhE | Effector protein | 6.586 | 2.278 | 1.955 | 0.580 | 7.379 | 7.634 | 2.630 | 0.870 | 6.831 | 3.228 | 1.285 | Early up | | Yes | | | | | | | | |
| B7L74_10955 | hypothetical protein | | 7.857 | 4.520 | 3.789 | 1.774 | 9.490 | 8.936 | 5.271 | 2.237 | 8.461 | 5.030 | 3.010 | Early up | | | | | | | | | | |
| B7L74_10960 | hypothetical protein | | 6.868 | 2.664 | 2.274 | 1.149 | 7.730 | 8.306 | 4.754 | 2.945 | 7.222 | 4.322 | 2.158 | Continuous up | | | | | | | | | | |
| B7L74_10965 | hypothetical protein | | -0.857 | -0.196 | -0.034 | -1.800 | 2.512 | 2.011 | -0.218 | 1.861 | 2.935 | -1.276 | -0.908 | | | | | | | | | | | |
| B7L74_10970 | hypothetical protein | Effector protein | 0.957 | 0.124 | 1.624 | 0.479 | 1.535 | 1.505 | 1.383 | 1.412 | 1.679 | 0.283 | 0.843 | | | Yes | | | | | | | | |
| B7L74_10975 | hypothetical protein | | -0.673 | 0.697 | 1.364 | -0.511 | -0.557 | 0.053 | 2.604 | 2.159 | 1.433 | 0.071 | 0.251 | | | | | | | | | | | |
| B7L74_10980 | hypothetical protein | | -0.159 | -0.606 | -3.432 | 2.043 | 0.350 | -1.503 | -1.503 | 0.077 | -0.428 | -1.533 | -1.804 | | | | | | | | | | | |
| B7L74_10985 | hypothetical protein(cpeF in Table 1 of RightonQ pape | Effector protein | -0.749 | 0.226 | -2.924 | 0.011 | -0.771 | -0.532 | -2.020 | -0.741 | -2.037 | -1.595 | -2.682 | Early down | | Yes | | | | | | | | |
| B7L74_10990 | hypothetical protein | | 0.372 | 0.790 | -0.547 | 0.715 | 0.031 | -0.473 | -0.362 | 0.778 | -1.230 | -1.057 | -0.930 | | | | | | | | | | | |
| B7L74_10995 | hypothetical protein | Effector protein | -0.193 | 0.722 | 0.116 | 0.214 | 0.246 | -0.568 | -0.722 | 0.635 | -0.958 | -0.759 | -1.182 | | | Yes | | | | | | | | |
| B7L74_11000 | hypothetical protein | | 0.674 | 0.821 | -0.750 | 0.126 | 0.004 | -0.390 | -0.359 | -0.049 | -0.882 | 0.357 | -0.656 | | | | | | | | | | | |
| B7L74_11005 | hypothetical protein | | -1.123 | -0.274 | -0.205 | 1.169 | -0.515 | -1.828 | -1.473 | -0.643 | -1.927 | 0.085 | 0.342 | | | | | | | | | | | |
| B7L74_11010 | transcriptional regulator | Transcription factors | 0.880 | -0.049 | 3.543 | 1.997 | 2.247 | 1.858 | 1.470 | 0.326 | 1.577 | 3.697 | 4.322 | Late up | | | | | | | | | | |
| B7L74_11015 | hypothetical protein | | 4.348 | 2.222 | 5.673 | 4.847 | 5.177 | 6.011 | 4.604 | 2.916 | 4.542 | 6.646 | 6.393 | Early up | | | | | | | | | | |
| B7L74_11020 | hypothetical protein | | -0.635 | 0.889 | 0.597 | -0.676 | -0.640 | 0.067 | 0.867 | 0.507 | 0.109 | 0.461 | 0.821 | | | | | | | | | | | |
| B7L74_11025 | hypothetical protein | | -0.722 | 1.307 | -0.564 | -0.365 | 0.556 | 0.165 | -1.805 | 1.140 | 0.294 | -0.670 | -1.436 | | | | | | | | | | | |

| | | | | | | | | | | | | | | | | | | | | | | |
|---|---|---|---|---|---|---|---|---|---|---|---|---|---|---|---|---|---|---|---|---|---|---|
| B7L74_11030 | hypothetical membrane spanning protein | | -1.591 | 2.457 | -2.866 | -1.484 | 4.622 | 0.965 | 2.425 | 4.540 | -2.646 | -2.393 | -4.083 | | | | | | | | | Yes (Vitamins) |
| B7L74_11035 | phosphodiesterase | | 0.167 | 0.864 | -0.035 | -2.506 | 5.453 | 0.146 | 3.758 | 1.607 | 0.185 | -2.761 | -1.303 | | | | | | | | | |
| B7L74_11040 | hypothetical protein | | -1.250 | 1.253 | -0.525 | 0.518 | 0.637 | -0.515 | -2.689 | 2.118 | -0.023 | 0.425 | -1.216 | | | | | | | | | |
| B7L74_11045 | hypothetical protein | | 0.223 | 0.881 | 0.371 | 0.963 | 0.165 | -0.654 | 0.919 | 1.658 | 1.999 | 0.323 | 0.424 | | | | | | | | | |
| B7L74_11050 | hypothetical protein | | -0.235 | -0.247 | -1.693 | 1.014 | -1.797 | -1.442 | -0.507 | 0.771 | 0.488 | 0.838 | -0.478 | | | | | | | | | |
| B7L74_11055 | hypothetical protein | Effector protein | -1.870 | 0.086 | -2.894 | -3.607 | -3.237 | -2.819 | -3.049 | -2.585 | -3.310 | -1.810 | -4.499 | Early down | | Yes | | | | | | |
| B7L74_11060 | gamma carbonic anhydrase family protein | | -0.069 | 0.024 | -0.418 | 0.215 | -1.043 | -1.315 | -0.956 | -0.621 | -2.385 | 0.367 | 0.061 | | | | | | | | | |
| B7L74_11065 | chromosome partitioning protein ParB | | -1.519 | -3.214 | -4.096 | -0.603 | -0.435 | -2.434 | -1.889 | -1.604 | -1.434 | -1.083 | -1.406 | | | | | | | | | |
| B7L74_11070 | chromosome partitioning protein ParA | Chromosome and associated protei | 1.868 | 0.754 | 1.071 | 5.894 | 4.860 | 2.267 | 4.223 | 3.195 | 6.362 | 2.295 | 2.507 | Early up | | | | | | | | |
| B7L74_11075 | chromosome partitioning protein ParB | Transcription factors | -0.998 | -0.484 | -1.704 | -1.705 | -2.131 | -0.958 | -1.191 | -1.609 | -1.352 | -1.551 | -2.111 | Early down | | | | | | | | |
| B7L74_11080 | repA protein | | -0.781 | -0.761 | -0.985 | -0.248 | -1.045 | -2.532 | -0.588 | -1.404 | -0.585 | 0.668 | 1.241 | | | | | | | | | |
| B7L74_11085 | hypothetical protein | | 3.800 | 1.430 | 1.296 | -1.624 | -0.647 | -0.728 | 2.251 | 1.872 | 0.765 | 0.370 | -0.356 | | | | | | | | | |

Table S2: Detailed information on 14 downregulated effector proteins. The cells with significant L2fc (Log 2 fold change) values (pvalue<0.05) are color coded, where the orange cells represents downregulation (L2fc<0).

| Genes | Corresppondïng CBU(Cb NM I RSA493) annotation | Gene annotation | Verdict from transcriptomics (Number of passages with significant expression) | COG categories | First verified of its potential effector functions based on | More information | LOG2FOLD CHANGE COMPARED TO P1 (The significant L2FC values are highlighted in orange) | | | | | | | | | | |
|---|---|---|---|---|---|---|---|---|---|---|---|---|---|---|---|---|---|
| | | | | | | | P03 | P05 | P10 | P12 | P16 | P21 | P31 | P42 | P51 | P61 | P67 |
| B7L74_08200 | Cbu1594 | Hypothetical protein | Early down (3) | General function prediction only | T4BSS dependent translocation. PMC3754607 | Using proteomics it is confirmed to be localized in mitochondria. PMC7950127. | -1.164 | -0.593 | -3.636 | 0.243 | -1.308 | -1.778 | -1.272 | -0.818 | -2.225 | -1.261 | -0.710 |
| B7L74_03065 | Cbu0590 | Hypothetical protein | Early down (2) | No hits on COG | Bioinformatic screening and experimental translocation assay. PMC3754607 | | 0.519 | 0.254 | -1.577 | -1.935 | -0.646 | -1.170 | -0.816 | -0.990 | -4.509 | -1.213 | -1.430 |
| B7L74_11055 | Cbua0034 | Plasmid effector protein **CpeH** | Early down (2) | Posttranslational modification, protein turnover, chaperones | CyaA translocation assays of the plasmid. PMC3697647 | | -1.870 | 0.086 | -2.894 | -3.607 | -3.237 | -2.819 | -3.049 | -2.585 | -3.310 | -1.810 | -4.499 |
| B7L74_03970 | Cbu0781 | **AnkG** | Early down (3) | Signal transduction mechanisms | T4SS translocation experment. PMC2698476 | PMC2698476 also shows how it localizes at host microtubule and thus might be very important for infection. PMID 20944063 showed AnkG interfers with mammalian apoptosis pathway. | -1.342 | -0.419 | -2.295 | -0.094 | -1.106 | -0.941 | -1.254 | -0.258 | -0.156 | -1.663 | -1.512 |
| B7L74_10985 | Cbua0023 | **CpeF** | Early down (3) | General function prediction only | β-lactamase and adenylate cyclase translocation assays. PMC3067651 | It is a member of plasmid effector family and was first detected in PMC3067651 where it was identified as effecotr based on β-lactamase and adenylate cyclase translocation assays, and identification of a C-terminal secretion signal Translocated to host cytosol. Mutation= Growth defect | -0.749 | 0.226 | -2.924 | 0.011 | -0.771 | -0.532 | -2.020 | -0.741 | -2.037 | -1.595 | -2.682 |
| B7L74_06660 | Cbu1292 | **AnkK** | Early down (3) | Signal transduction mechanisms | Presence of Eukaryotic like ORF or domain i.e. ankyrin repeats. PMC2698476 | Predicted to be a effector protein because of the eukaryotic like ORF or domain i.e. ankyrin repeats. AnkK proteins are shown to be not secreted by T4SS inside the host cell (PMC2698476) but it can work without being delivered to host cell. PMID: 18279343 showed its importance for growth inside macrophages | -0.992 | -0.515 | -1.675 | -1.932 | -1.172 | -0.579 | -2.089 | -0.683 | -0.893 | -1.168 | -1.410 |
| B7L74_03275 | Cbu0635 | Hypothetical membrane spanning protein | Early down (4) | Lipid transport and metabolism | BlaM translocation assay. PMC3102713 | First reported in PMC3102713 where it was experimentally shown to be translocated using dot system using BlaM translocation assay. Expressed in mammalian Hela229 cells shows its distribution around golgi vesicles and potentially disturb mammalian secretory trafficking. It can interfere with host cell secretion(PMID: 21637816). | -0.831 | -1.109 | -2.043 | -0.356 | -0.864 | -0.643 | -1.108 | -1.529 | -1.654 | -1.168 | -1.626 |
| B7L74_04780 | Cbu0937 | **CirC** | Early down (4) | No hits on COG | Localization and transposon insertion mutation studies. PMC3754607 | PMC3754607 identified it as an effector by localization and transposon insertion mutation studies for intracellular replication and CCV formation. They showed that the mutant of this protein produced small vacuoles, which validated screens conducted by other groups that identified Cbu0937 as being an effector important for CCV biogenesis. | -1.284 | -1.404 | -2.237 | -0.906 | -2.118 | -2.035 | -1.249 | -1.740 | -1.841 | -1.635 | -0.970 |
| B7L74_07850 | Cbu1525 | Hypothetical protein | Early down (5) | No hits on COG | Cya translocation studies and T4 translocation signals. PMC3102713 | PMC3102713 identified it in cya translocation studies and also by looking for T4 translocation signals. This is a frameshifted ORF. | -0.881 | -1.008 | -1.547 | -0.981 | -1.862 | -1.965 | -1.525 | -1.495 | -2.400 | -0.757 | -0.662 |
| B7L74_00115 | Cbu0021 | **CvpB/Cig2** | Early down (7) | No hits on COG | Translocation via L. pneumophila Dot/Icm system.PMC3581968 | First reported in PMC3581968 where it is postulated to be an effector based on its translocation via L. pneumophila Dot/Icm system. PMCID: PMC4117601 reports its mutation(by transposon insertion) displayed multi-vacuolar phenotype without an overall replication defect.. Mutation thus causes growth defect and CCV fusion defect. | -0.841 | 0.586 | -4.204 | -3.919 | -0.179 | -0.732 | -3.980 | -2.729 | -2.967 | -3.686 | -3.805 |
| B7L74_08400 | Cbu1636 | Hypothetical protein | Early down (8) | Mobilome: prophages, transposons | PmrA like domain containing protein and Dot/Icm dependent translocation. PMC3003115. | Reported in article PMC3003115( Table S03). Identified by Dot/Icm substrate homologus/putative PmrA regulated eukaryotic-like domain containing protein. It also has dot/Icm dependent translocation. 45 KDa protein with coiled coil structure. | -0.605 | -0.159 | -2.461 | -1.239 | -1.125 | -0.897 | -1.850 | -1.862 | -3.374 | -1.584 | -2.176 |
| B7L74_01850 | Cbu0355 | **AnkD** | Early down (8) | Signal transduction mechanisms | Presence of ankyrin repreats as well as F-box domains. PMC2698476 | This protein has both ankyrin repreats as well as F-box domains. (PMC2698476) | -1.384 | -0.285 | -3.084 | -2.290 | -1.038 | -0.635 | -2.036 | -1.180 | -1.227 | -3.052 | -3.115 |
| B7L74_09015 | Cbu1751 | **Cig57** | Early down (9) | Replication, recombination and repair | PmrA like domain containing protein and Dot/Icm dependent translocation. PMC3003115. | From table S03 of PMC3003115, it is 49 KdA coiled coil protein. Identified by DotF binding protein/putative PmrA regulated eukaryotic-like domain containing protein. It also has dot/Icm dependent translocation. Can cause intracellular replication defects. | -0.738 | 0.062 | -3.881 | -2.795 | -1.921 | -2.597 | -3.302 | -3.141 | -3.603 | -2.886 | -2.989 |
| B7L74_09020 | Cbu1752 | Hypothetical protein | Early down (9) | Carbohydrate transport and metabolism | Loss-of-function mutations. PMC5865027 | Was found to be important for vacuole biogenesis and its mutation resulted in a small-vacuole (CCV) phenotype. PMC5865027 | -0.725 | 0.236 | -3.594 | -4.123 | -1.952 | -2.960 | -3.978 | -3.613 | -3.820 | -2.916 | -3.142 |

Table S3: Different types of SNPs and total DIP mutations occurring in genes. The heatmap illustrates percentage of population mutated in each passage. Column "Verdict from transcriptomics" lists the gene expression verdicts made only for the mutations in coding regions of DEGs. Column "Classification by their occurences" in each sheet classifies the mutation according to the pattern of its occurence in the passages.

SNPs_Missense: Single nucleotide polymorphisms; Missense mutations
SNPs_Intergenic: Single nucleotide polymorphisms; Intergenic mutations
SNPs_Non-sense: Single nucleotide polymorphisms; Nonsense mutations
SNPs_Silent: Single nucleotide polymorphisms; Silent mutations
DIPs: Deletion, Insertion polymorphisms

| Gene name from Breseq | Gene name from reference genome | Description/Annotation | Amino acid and basepair change | P03 | P05 | P10 | P13 | P16 | P21 | P31 | P42 | P51 | P61 | P67 | Verdict from transcriptomics | Classification by their occurences |
|---|---|---|---|---|---|---|---|---|---|---|---|---|---|---|---|---|
| LOBOINFK_00627 → | B7L74_03250 | bifunctional proline dehydrogenase/L-glutamate g | P521R (CCT→CGT) ‡ | | | | | | | | | | | | Continuous down | Maintained in later passages |
| LOBOINFK_00627 → | B7L74_03250 | bifunctional proline dehydrogenase/L-glutamate g | P521A (CCT→GCT) ‡ | | | | | | | | | | | | Continuous down | Maintained in later passages |
| LOBOINFK_00586 → | B7L74_03055 | quinolinate synthetase | L66M (TTG→ATG) | | | | | | | | | | | | Continuous down | Transient |
| LOBOINFK_00643 → | B7L74_03330 | riboflavin synthase | K111N (AAA→AAC) | | | | | | | | | | | | Continuous down | Transient |
| LOBOINFK_01867 ← | B7L74_01615 | molecular chaperone HtpG | F130V (TTT→GTT) | | | | | | | | | | | | Continuous down | Transient |
| LOBOINFK_02016 ← | B7L74_08450 | Dot/Icm secretion system ATPase DotB | E341D (GAA→GAC) | | | | | | | | | | | | Continuous down | Transient |
| LOBOINFK_02016 ← | B7L74_08450 | Dot/Icm secretion system ATPase DotB | G343V (GGT→GTT) | | | | | | | | | | | | Continuous down | Transient |
| LOBOINFK_00674 ← | B7L74_09545 | hypothetical protein | S52T (AGT→ACT) | | | | | | | | | | | | Continuous up | Transient |
| LOBOINFK_00674 ← | B7L74_09545 | hypothetical protein | E53D (GAG→GAT) | | | | | | | | | | | | Continuous up | Transient |
| LOBOINFK_00674 ← | B7L74_09545 | hypothetical protein | Y363N (TAT→AAT) | | | | | | | | | | | | Continuous up | Transient |
| LOBOINFK_01090 ← | B7L74_01865 | hypothetical protein | R92L (CGA→CTA) ‡ | | | | | | | | | | | | Continuous up | Transient |
| LOBOINFK_01656 → | B7L74_02020 | hypothetical protein | G1183V (GGG→GTG) | | | | | | | | | | | | Continuous up | Transient |
| LOBOINFK_01656 → | B7L74_02020 | hypothetical protein | T607A (ACG→GCG) | | | | | | | | | | | | Continuous up | Transient |
| LOBOINFK_01656 → | B7L74_02020 | hypothetical protein | H1179P (CAC→CCC) | | | | | | | | | | | | Continuous up | Transient |
| LOBOINFK_01824 → | B7L74_10175 | Bcr/CflA family drug resistance efflux transporter | F53S (TTT→TCT) | | | | | | | | | | | | Continuous up | Transient |
| LOBOINFK_01817 → | B7L74_03805 | alpha/beta hydrolase | F15V (TTT→GTT) | | | | | | | | | | | | Continuous up | Transient |
| LOBOINFK_01992 → | B7L74_08170 | Na(+)/H(+) antiporter subunit A | A155S (GCC→TCC) | | | | | | | | | | | | Continuous up | Transient |
| LOBOINFK_01992 → | B7L74_08170 | Na(+)/H(+) antiporter subunit A | K158Q (AAG→CAG) | | | | | | | | | | | | Continuous up | Transient |
| LOBOINFK_01557 ← | B7L74_03505 | NAD-dependent dehydratase | P264Q (CCA→CAA) | | | | | | | | | | | | Down in P10 | Transient |
| LOBOINFK_00727 → | B7L74_09255 | N-acetyltransferase | Y5N (TAT→AAT) | | | | | | | | | | | | Early down | Transient |
| LOBOINFK_00727 → | B7L74_09255 | N-acetyltransferase | Y4N (TAT→AAT) | | | | | | | | | | | | Early down | Transient |
| LOBOINFK_00025 → | B7L74_05345 | cytochrome o ubiquinol oxidase subunit III | W94L (TGG→TTG) | | | | | | | | | | | | Early down | Transient |
| LOBOINFK_01345 → | B7L74_01330 | 30S ribosomal protein S5 | V32G (GTG→GGG) | | | | | | | | | | | | Early down | Transient |
| LOBOINFK_00941 → | B7L74_03970 | hypothetical protein | V314F (GTT→TTT) | | | | | | | | | | | | Early down | Transient |
| LOBOINFK_00941 → | B7L74_03970 | hypothetical protein | K311N (AAA→AAT) ‡ | | | | | | | | | | | | Early down | Transient |
| LOBOINFK_00941 → | B7L74_03970 | hypothetical protein | K311Q (AAA→CAA) ‡ | | | | | | | | | | | | Early down | Transient |
| LOBOINFK_01872 ← | B7L74_01590 | GTP pyrophosphokinase SpoT | T262A (ACT→GCT) | | | | | | | | | | | | Early down | Present in all passages |
| LOBOINFK_00517 → | B7L74_07075 | para-aminobenzoate synthetase component I | S200A (TCA→GCA) ‡ | | | | | | | | | | | | Early down | Transient |
| LOBOINFK_01087 → | B7L74_01850 | hypothetical protein | Q76K (CAA→AAA) | | | | | | | | | | | | Early down | Transient |
| LOBOINFK_00963 → | B7L74_04100 | multidrug transporter AcrB | Q745H (CAG→CAT) | | | | | | | | | | | | Early down | Transient |
| LOBOINFK_00963 → | B7L74_04100 | multidrug transporter AcrB | M746L (ATG→CTG) | | | | | | | | | | | | Early down | Transient |
| LOBOINFK_00403 ← | B7L74_00420 | sulfatase | Q557H (CAA→CAC) ‡ | | | | | | | | | | | | Early down | Transient |
| LOBOINFK_00403 ← | B7L74_00420 | sulfatase | Q557E (CAA→GAA) ‡ | | | | | | | | | | | | Early down | Transient |
| LOBOINFK_00403 ← | B7L74_00420 | sulfatase | C556F (TGT→TTT) | | | | | | | | | | | | Early down | Transient |
| LOBOINFK_00403 ← | B7L74_00420 | sulfatase | L103R (CTT→CGT) | | | | | | | | | | | | Early down | Transient |
| LOBOINFK_00976 ← | B7L74_02550 | acyl carrier protein | Q15K (CAA→AAA) | | | | | | | | | | | | Early down | Transient |
| LOBOINFK_01417 ← | B7L74_06135 | DNA translocase FtsK | P276R (CCG→CGG) | | | | | | | | | | | | Early down | Transient |
| ligA → | B7L74_02820 | DNA ligase (NAD(+)) LigA | N577Y (AAT→TAT) | | | | | | | | | | | | Early down | Transient |
| LOBOINFK_01633 ← | B7L74_06270 | hypothetical protein | N356S (AAT→AGT) | | | | | | | | | | | | Early down | Present in all passages |
| LOBOINFK_01633 ← | B7L74_06270 | hypothetical protein | Q359E (CAA→GAA) | | | | | | | | | | | | Early down | Present in all passages |
| LOBOINFK_01633 ← | B7L74_06270 | hypothetical protein | S363N (AGT→AAT) | | | | | | | | | | | | Early down | Transient |
| LOBOINFK_01507 ← | B7L74_08890 | ornithine cyclodeaminase | N212H (AAC→CAC) | | | | | | | | | | | | Early down | Transient |
| LOBOINFK_01507 ← | B7L74_08890 | ornithine cyclodeaminase | Q208K (CAA→AAA) | | | | | | | | | | | | Early down | Transient |
| LOBOINFK_01549 ← | B7L74_03545 | ABC transporter ATP-binding protein | N196D (AAC→GAC) | | | | | | | | | | | | Early down | Present in all passages |
| LOBOINFK_01288 → | B7L74_10985 | hypothetical protein | M115K (ATG→AAG) | | | | | | | | | | | | Early down | Transient |
| LOBOINFK_01288 → | B7L74_10985 | hypothetical protein | L114H (CTT→CAT) | | | | | | | | | | | | Early down | Transient |
| LOBOINFK_01288 → | B7L74_10985 | hypothetical protein | V117I (GTC→ATC) | | | | | | | | | | | | Early down | Transient |
| LOBOINFK_00980 ← | B7L74_02530 | phosphate acyltransferase | M10I (ATG→ATT) | | | | | | | | | | | | Early down | Transient |
| LOBOINFK_01451 → | B7L74_08350 | type IV secretion protein IcmE | L535F (TTA→TTT) | | | | | | | | | | | | Early down | Transient |
| LOBOINFK_01451 → | B7L74_08350 | type IV secretion protein IcmE | K238N (AAA→AAC) | | | | | | | | | | | | Early down | Transient |
| LOBOINFK_00002 → | B7L74_05460 | SMC-Scp complex subunit ScpB | K85N (AAA→AAC) | | | | | | | | | | | | Early down | Transient |
| LOBOINFK_01415 ← | B7L74_06125 | replication-associated recombination protein A | K442N (AAA→AAC) | | | | | | | | | | | | Early down | Transient |
| LOBOINFK_01333 → | B7L74_01270 | 50S ribosomal protein L22 | K42N (AAA→AAC) | | | | | | | | | | | | Early down | Transient |
| LOBOINFK_01479 ← | B7L74_06660 | hypothetical protein | K304R (AAA→AGA) | | | | | | | | | | | | Early down | Transient |
| LOBOINFK_01891 → | B7L74_01215 | DNA-directed RNA polymerase subunit beta' | K280N (AAG→AAT) | | | | | | | | | | | | Early down | Maintained in later passages |
| LOBOINFK_01891 → | B7L74_01215 | DNA-directed RNA polymerase subunit beta' | L279F (TTG→TTC) | | | | | | | | | | | | Early down | Maintained in later passages |
| LOBOINFK_01506 → | B7L74_08885 | acetyl-CoA carboxylase biotin carboxylase subun | K159N (AAA→AAC) | | | | | | | | | | | | Early down | Transient |
| LOBOINFK_00482 ← | B7L74_06905 | DNA polymerase III subunit alpha | I88F (ATT→TTT) | | | | | | | | | | | | Early down | Transient |
| LOBOINFK_00482 ← | B7L74_06905 | DNA polymerase III subunit alpha | R713H (CGT→CAT) ‡ | | | | | | | | | | | | Early down | Transient |
| LOBOINFK_00510 ← | B7L74_07040 | hypothetical protein | I78L (ATA→TTA) | | | | | | | | | | | | Early down | Maintained in later passages |
| LOBOINFK_00510 ← | B7L74_07040 | hypothetical protein | T79S (ACT→TCT) | | | | | | | | | | | | Early down | Maintained in later passages |
| LOBOINFK_01847 → | B7L74_09725 | peptidase | I356T (ATT→ACT) | | | | | | | | | | | | Early down | Transient |
| LOBOINFK_01847 → | B7L74_09725 | peptidase | S358N (AGT→AAT) | | | | | | | | | | | | Early down | Transient |
| LOBOINFK_01847 → | B7L74_09725 | peptidase | N357I (AAT→ATT) | | | | | | | | | | | | Early down | Transient |
| LOBOINFK_00069 → | B7L74_05115 | adenylosuccinate synthetase | H419L (CAT→CTT) | | | | | | | | | | | | Early down | Transient |
| LOBOINFK_00069 → | B7L74_05115 | adenylosuccinate synthetase | K418M (AAG→ATG) | | | | | | | | | | | | Early down | Transient |
| LOBOINFK_01640 ← | B7L74_06310 | haloalkane dehalogenase | G88E (GGG→GAG) | | | | | | | | | | | | Early down | Transient |
| LOBOINFK_01041 → | B7L74_02170 | pantothenate synthetase | G85V (GGA→GTA) | | | | | | | | | | | | Early down | Transient |
| LOBOINFK_01041 → | B7L74_02170 | pantothenate synthetase | Y88S (TAC→TCC) | | | | | | | | | | | | Early down | Transient |
| LOBOINFK_00549 ← | B7L74_07225 | succinate dehydrogenase flavoprotein subunit | E548D (GAA→GAC) | | | | | | | | | | | | Early down | Transient |
| LOBOINFK_00344 ← | B7L74_00755 | preprotein translocase subunit SecA | E340D (GAA→GAC) | | | | | | | | | | | | Early down | Transient |
| LOBOINFK_00344 ← | B7L74_00755 | preprotein translocase subunit SecA | V341F (GTC→TTC) | | | | | | | | | | | | Early down | Transient |
| bdbD → | B7L74_09850 | Disulfide bond formation protein D | E246Q (GAA→CAA) | | | | | | | | | | | | Early down | Transient |
| bdbD → | B7L74_09850 | Disulfide bond formation protein D | I247L (ATC→CTC) | | | | | | | | | | | | Early down | Transient |
| LOBOINFK_00245 ← | B7L74_04245 | ABC transporter permease | C169W (TGT→TGG) | | | | | | | | | | | | Early down | Transient |
| LOBOINFK_00470 ← | B7L74_06855 | threonine--tRNA ligase | A80P (GCG→CCG) | | | | | | | | | | | | Early down | Transient |
| LOBOINFK_00470 ← | B7L74_06855 | threonine--tRNA ligase | V83L (GTA→CTA) | | | | | | | | | | | | Early down | Transient |
| LOBOINFK_01606 → | B7L74_00190 | beta-ketoacyl-ACP synthase | A296P (GCG→CCG) | | | | | | | | | | | | Early down | Present in all passages |
| LOBOINFK_01606 → | B7L74_00190 | beta-ketoacyl-ACP synthase | A190P (GCT→CCT) ‡ | | | | | | | | | | | | Early down | Transient |
| LOBOINFK_01606 → | B7L74_00190 | beta-ketoacyl-ACP synthase | R189K (AGG→AAG) | | | | | | | | | | | | Early down | Transient |
| LOBOINFK_01447 → | B7L74_08370 | lipoprotein | A19T (GCA→ACA) | | | | | | | | | | | | Early down | Transient |
| LOBOINFK_00338 → | B7L74_00795 | type 4 fimbrial biosynthesis protein | Y314N (TAC→AAC) | | | | | | | | | | | | Early up | Transient |
| LOBOINFK_00338 → | B7L74_00795 | type 4 fimbrial biosynthesis protein | E312D (GAG→GAT) | | | | | | | | | | | | Early up | Transient |

| |
|---|
| 5-10 |
| 10-30 |
| 30-50 |
| 50-70 |
| 70-90 |
| 90-99 |
| 100 |

| | | | | | |
|---|---|---|---|---|---|
| LOBOINFK_00338 → | B7L74_00795 | type 4 fimbrial biosynthesis protein | D315A (GAT→GCT) | Early up | Transient |
| LOBOINFK_00142 → | B7L74_04750 | glycerol-3-phosphate dehydrogenase | W102R (TGG→CGG) | Early up | Transient |
| LOBOINFK_00142 → | B7L74_04750 | glycerol-3-phosphate dehydrogenase | M103I (ATG→ATC) | Early up | Transient |
| LOBOINFK_00142 → | B7L74_04750 | glycerol-3-phosphate dehydrogenase | R105Q (CGA→CAA) | Early up | Transient |
| LOBOINFK_01924 ← | B7L74_05470 | hypothetical protein | V8I (GTT→ATT) | Early up | Maintained in later passages |
| LOBOINFK_01924 ← | B7L74_05470 | hypothetical protein | Y7D (TAT→GAT) | Early up | Maintained in later passages |
| LOBOINFK_01629 ← | B7L74_00060 | D-alanyl-D-alanine carboxypeptidase / D-alanyl-D | V55G (GTT→GGT) | Early up | Transient |
| LOBOINFK_00906 → | B7L74_05720 | hypothetical protein | V32A (GTA→GCA) ‡ | Early up | Transient |
| LOBOINFK_00906 → | B7L74_05720 | hypothetical protein | V32I (GTA→ATA) ‡ | Early up | Transient |
| LOBOINFK_01318 → | B7L74_02825 | hypothetical protein | V198G (GTT→GGT) | Early up | Transient |
| LOBOINFK_02003 → | B7L74_03640 | hypothetical protein | T7P (ACC→CCC) | Early up | Transient |
| LOBOINFK_00197 → | B7L74_04490 | hypothetical protein | S36T (AGC→ACC) | Early up | Transient |
| LOBOINFK_00400 ← | B7L74_00435 | phosphoglycerol transferase | Q431H (CAA→CAC) | Early up | Transient |
| LOBOINFK_01021 → | B7L74_02295 | hypothetical protein | Q39K (CAA→AAA) | Early up | Transient |
| LOBOINFK_01021 → | B7L74_02295 | hypothetical protein | T45P (ACA→CCA) | Early up | Transient |
| LOBOINFK_01053 ← | B7L74_01680 | hypothetical protein | N183Y (AAT→TAT) | Early up | Transient |
| LOBOINFK_01952 → | B7L74_08775 | alkyl hydroperoxide reductase | N143H (AAC→CAC) | Early up | Transient |
| LOBOINFK_01570 ← | B7L74_03430 | hypothetical protein | N103H (AAT→CAT) | Early up | Transient |
| LOBOINFK_01570 ← | B7L74_03430 | hypothetical protein | M101I (ATG→ATT) | Early up | Transient |
| LOBOINFK_01570 ← | B7L74_03430 | hypothetical protein | I102N (ATT→AAT) | Early up | Transient |
| LOBOINFK_01222 → | B7L74_09875 | glycine--tRNA ligase subunit beta | L54V (TTG→GTG) | Early up | Transient |
| LOBOINFK_01974 → | B7L74_08085 | MFS transporter | L356V (TTG→GTG) | Early up | Transient |
| LOBOINFK_01974 → | B7L74_08085 | MFS transporter | E116K (GAA→AAA) | Early up | Maintained in later passages |
| LOBOINFK_01160 ← | B7L74_07495 | hypothetical protein | L281F (TTA→TTT) | Early up | Transient |
| LOBOINFK_00173 ← | B7L74_04610 | hypothetical protein | L246Q (CTA→CAA) | Early up | Transient |
| LOBOINFK_01916 ← | B7L74_05510 | intracellular signaling protein | L204I (TTA→ATA) ‡ | Early up | Transient |
| LOBOINFK_01695 → | B7L74_09100 | ferrous iron transporter B | K811N (AAA→AAC) | Early up | Transient |
| LOBOINFK_01205 → | B7L74_07725 | hypothetical protein | K51N (AAA→AAC) | Early up | Transient |
| LOBOINFK_01468 → | B7L74_08260 | hypothetical protein | K19N (AAA→AAC) | Early up | Transient |
| LOBOINFK_01297 ← | B7L74_10940 | hypothetical protein | H33N (CAT→AAT) | Early up | Transient |
| LOBOINFK_01297 ← | B7L74_10940 | hypothetical protein | I32M (ATT→ATG) | Early up | Transient |
| LOBOINFK_01838 → | B7L74_10265 | hypothetical protein | E99K (GAA→AAA) | Early up | Maintained in later passages |
| LOBOINFK_01838 → | B7L74_10265 | hypothetical protein | P205L (CCT→CTT) | Early up | Transient |
| esiB → | B7L74_05845 | Secretory immunoglobulin A-binding protein EsiI | D891E (GAT→GAG) | Early up | Transient |
| esiB → | B7L74_05845 | Secretory immunoglobulin A-binding protein EsiI | A895D (GCT→GAT) | Early up | Transient |
| esiB → | B7L74_05845 | Secretory immunoglobulin A-binding protein EsiI | N893Y (AAT→TAT) | Early up | Transient |
| LOBOINFK_00638 ← | B7L74_03305 | leucine dehydrogenase | D116V (GAC→GTC) | Early up | Transient |
| lgrD ← | B7L74_06750 | Linear gramicidin synthase subunit D | C140Y (TGT→TAT) ‡ | Early up | Transient |
| lgrD ← | B7L74_06750 | Linear gramicidin synthase subunit D | C140R (TGT→CGT) ‡ | Early up | Transient |
| LOBOINFK_01084 ← | B7L74_01835 | cation transporter | P284A (CCT→GCT) | Late down | Transient |
| LOBOINFK_01940 ← | B7L74_08855 | aconitate hydratase | N817Y (AAT→TAT) | Late down | Transient |
| LOBOINFK_01940 ← | B7L74_08855 | aconitate hydratase | V208G (GTC→GGC) | Late down | Transient |
| hflB ← | B7L74_06975 | ATP-dependent metalloprotease | A164V (GCC→GTC) | Late down | Transient |
| LOBOINFK_00108 → | B7L74_04900 | putative glycosyl hydrolase | Y282N (TAT→AAT) | Late up | Transient |
| LOBOINFK_01573 ← | B7L74_06390 | long-chain fatty acid transporter | N239I (AAC→ATC) | Late up | Transient |
| LOBOINFK_01573 ← | B7L74_06390 | long-chain fatty acid transporter | F236L (TTT→TTA) | Late up | Transient |
| LOBOINFK_01372 → | B7L74_01505 | 2-amino-4-hydroxy-6- hydroxymethyldihydropter | L71S (TTG→TCG) ‡ | Late up | Transient |
| iucA → | B7L74_02160 | N(2)-citryl-N(6)-acetyl-N(6)-hydroxylysine synth | I455M (ATT→ATG) | Late up | Transient |
| iucA → | B7L74_02160 | N(2)-citryl-N(6)-acetyl-N(6)-hydroxylysine synth | F454L (TTC→TTA) | Late up | Transient |
| LOBOINFK_01545 ← | B7L74_03575 | transposase | I3N (ATT→AAT) | Late up | Transient |
| fadE_2 → | B7L74_03875 | acyl-CoA dehydrogenase | I174S (ATT→AGT) | Late up | Transient |
| fadE_2 → | B7L74_03875 | acyl-CoA dehydrogenase | L172I (CTA→ATA) | Late up | Transient |
| LOBOINFK_01285 → | B7L74_11010 | transcriptional regulator | F42I (TTT→ATT) | Late up | Transient |
| LOBOINFK_01285 → | B7L74_11010 | transcriptional regulator | D43H (GAT→CAT) | Late up | Transient |
| LOBOINFK_01285 → | B7L74_11010 | transcriptional regulator | E41Q (GAA→CAA) | Late up | Transient |
| LOBOINFK_00801 ← | B7L74_10615 | tryptophan--tRNA ligase | D257V (GAT→GTT) | Late up | Transient |
| LOBOINFK_00801 ← | B7L74_10615 | tryptophan--tRNA ligase | T260S (ACG→TCG) | Late up | Transient |
| LOBOINFK_01410 → | B7L74_07525 | hypothetical protein | H247Y (CAT→TAT) | Up in P12 | Transient |
| LOBOINFK_01410 → | B7L74_07525 | hypothetical protein | N248Y (AAT→TAT) | Up in P12 | Transient |
| LOBOINFK_01410 → | B7L74_07525 | hypothetical protein | N249D (AAT→GAT) | Up in P12 | Transient |
| LOBOINFK_01342 → | B7L74_01315 | 30S ribosomal protein S8 | R117P (CGG→CCG) | Up in P16 | Transient |
| LOBOINFK_00886 → | B7L74_05810 | IscS subfamily cysteine desulfurase | L176I (TTA→ATA) | Up in P21 | Transient |
| LOBOINFK_01831 ← | B7L74_10225 | phosphotransferase | K108N (AAA→AAC) | Up in P21 | Transient |
| LOBOINFK_00298 → | B7L74_00990 | hypothetical protein | *219K (TAA→AAA) | Up in P21 | Transient |
| LOBOINFK_00298 → | B7L74_00990 | hypothetical protein | F218L (TTT→TTA) | Up in P21 | Transient |
| ybhS → | B7L74_00255 | putative multidrug ABC transporter permease Ybl | L254V (TTA→GTA) | | Transient |
| traI → | B7L74_11025 | Multifunctional conjugation protein TraI | E482Q (GAA→CAA) | | Transient |
| traI → | B7L74_11025 | Multifunctional conjugation protein TraI | V481I (GTT→ATT) | | Transient |
| LOBOINFK_02034 ← | B7L74_10150 | IS110 family transposase | Q110R (CAG→CGG) | | Present in all passages |
| LOBOINFK_02012 → | B7L74_10080 | hypothetical protein | M32I (ATG→ATC) | | Maintained in later passages |
| LOBOINFK_01985 → | B7L74_08210 | RNA polymerase sigma factor RpoD | K14N (AAG→AAC) | | Transient |
| LOBOINFK_01973 → | B7L74_05555 | beta-N-acetylhexosaminidase | E220Q (GAA→CAA) | | Transient |
| LOBOINFK_01973 → | B7L74_05555 | beta-N-acetylhexosaminidase | D219H (GAT→CAT) | | Transient |
| LOBOINFK_01961 ← | B7L74_07965 | phosphoenolpyruvate--protein phosphotransferase | L567I (TTA→ATA) | | Transient |
| LOBOINFK_01961 ← | B7L74_07965 | phosphoenolpyruvate--protein phosphotransferase | L568I (TTA→ATA) | | Transient |
| LOBOINFK_01937 ← | B7L74_08870 | thiol:disulfide interchange protein | L425M (CTG→ATG) | | Transient |
| LOBOINFK_01935 ← | B7L74_05685 | hypothetical protein | K79N (AAA→AAC) | | Transient |
| LOBOINFK_01907 → | B7L74_08065 | hypothetical protein | K315N (AAA→AAC) | | Transient |
| LOBOINFK_01859 → | B7L74_09655 | molecular chaperone DjlA | Q43E (CAA→GAA) | | Transient |
| LOBOINFK_01811 ← | B7L74_03775 | ABC transporter ATP-binding protein | G213S (GGC→AGC) | | Transient |
| LOBOINFK_01800 → | B7L74_03715 | uridine phosphorylase | A105G (GCC→GGC) | | Transient |
| LOBOINFK_01762 ← | B7L74_08675 | dCTP deaminase | K147N (AAA→AAC) | | Transient |
| LOBOINFK_01761 ← | B7L74_08670 | hypothetical protein | K185N (AAG→AAC) | | Transient |
| LOBOINFK_01746 ← | B7L74_08580 | 5'-nucleotidase SurE | E192A (GAG→GCG) | | Transient |
| LOBOINFK_01714 → | B7L74_02650 | fructose 1,6-bisphosphatase | E24A (GAA→GCA) ‡ | | Transient |

| | | | | |
|---|---|---|---|---|
| LOBOINFK_01701 → | B7L74_02580 | TatD family deoxyribonuclease | T76S (ACG→TCG) | Transient |
| LOBOINFK_01645 → | B7L74_06335 | ATP-dependent helicase | H263Q (CAC→CAA) | Transient |
| LOBOINFK_01644 → | B7L74_06330 | ATP-dependent helicase | K690T (AAA→ACA) | Transient |
| LOBOINFK_01643 → | B7L74_06325 | two-component sensor histidine kinase QseC | T258I (ACT→ATT) | Maintained in later passages |
| LOBOINFK_01642 → | B7L74_06320 | DNA-binding response regulator | R119H (CGT→CAT) | Transient |
| LOBOINFK_01642 → | B7L74_06320 | DNA-binding response regulator | A82T (GCT→ACT) | Transient |
| LOBOINFK_01642 → | B7L74_06320 | DNA-binding response regulator | R69H (CGC→CAC) | Transient |
| LOBOINFK_01642 → | B7L74_06320 | DNA-binding response regulator | E45K (GAA→AAA) | Transient |
| LOBOINFK_01642 → | B7L74_06320 | DNA-binding response regulator | L93F (CTC→TTC) | Transient |
| LOBOINFK_01642 → | B7L74_06320 | DNA-binding response regulator | D48G (GAT→GGT) | Transient |
| LOBOINFK_01641 ← | B7L74_06315 | glutamate dehydrogenase | C425W (TGT→TGG) | Transient |
| LOBOINFK_01637 ← | B7L74_06295 | 2-dehydro-3-deoxygluconokinase | D87G (GAC→GGC) | Transient |
| LOBOINFK_01637 ← | B7L74_06295 | 2-dehydro-3-deoxygluconokinase | P90S (CCG→TCG) | Transient |
| LOBOINFK_01637 ← | B7L74_06295 | 2-dehydro-3-deoxygluconokinase | K88N (AAG→AAC) | Transient |
| LOBOINFK_01628 ← | B7L74_00065 | shikimate dehydrogenase | K37N (AAA→AAC) | Transient |
| LOBOINFK_01625 → | B7L74_00080 | anion permease | G32A (GGC→GCC) | Transient |
| LOBOINFK_01625 → | B7L74_00080 | anion permease | A28P (GCC→CCC) | Transient |
| LOBOINFK_01625 → | B7L74_00080 | anion permease | M31L (ATG→TTG) | Transient |
| LOBOINFK_01625 → | B7L74_00080 | anion permease | N29I (AAC→ATC) | Transient |
| LOBOINFK_01589 → | B7L74_06480 | Na+/H+ antiporter | I127M (ATT→ATG) | Transient |
| LOBOINFK_01586 ← | B7L74_06460 | polyprenyl synthetase | Y50S (TAC→TCC) | Transient |
| LOBOINFK_01574 → | B7L74_06395 | exodeoxyribonuclease VII large subunit | I93F (ATC→TTC) | Transient |
| LOBOINFK_01574 → | B7L74_06395 | exodeoxyribonuclease VII large subunit | E92D (GAA→GAT) | Present in all passages |
| LOBOINFK_01553 → | B7L74_03525 | adenylyltransferase | V449A (GTA→GCA) | Transient |
| LOBOINFK_01553 → | B7L74_03525 | adenylyltransferase | A446T (GCT→ACT) | Transient |
| LOBOINFK_01540 → | B7L74_03605 | MFS transporter | L65R (CTG→CGG) | Transient |
| LOBOINFK_01514 → | B7L74_08925 | D-3-phosphoglycerate dehydrogenase | N79T (AAC→ACC) | Transient |
| LOBOINFK_01499 → | B7L74_06570 | mandelate racemase | L223V (TTG→GTG) | Transient |
| LOBOINFK_01498 → | B7L74_06575 | short-chain dehydrogenase | L258F (TTA→TTC) | Transient |
| LOBOINFK_01437 ← | B7L74_06240 | hypothetical protein | G176R (GGG→AGG) | Transient |
| LOBOINFK_01437 ← | B7L74_06240 | hypothetical protein | N175K (AAT→AAA) | Transient |
| LOBOINFK_01437 ← | B7L74_06240 | hypothetical protein | V174G (GTC→GGC) | Transient |
| LOBOINFK_01413 → | B7L74_06115 | hypothetical protein | K209N (AAG→AAC) | Transient |
| LOBOINFK_01377 → | B7L74_01530 | hypothetical protein | V27G (GTC→GGC) | Transient |
| LOBOINFK_01371 → | B7L74_01500 | polynucleotide adenylyltransferase PcnB | T282P (ACA→CCA) | Transient |
| LOBOINFK_01362 → | B7L74_01430 | excinuclease ABC subunit A | G411V (GGT→GTT) | Transient |
| LOBOINFK_01362 → | B7L74_01430 | excinuclease ABC subunit A | T408P (ACT→CCT) | Transient |
| LOBOINFK_01344 → | B7L74_01325 | 50S ribosomal protein L18 | K63N (AAA→AAC) | Transient |
| LOBOINFK_01324 ← | B7L74_02860 | penicillin-binding protein | Q464H (CAA→CAC) | Transient |
| LOBOINFK_01310 → | B7L74_02770 | undecaprenyl-phosphate alpha-N-acetylglucosami | F103I (TTT→ATT) ‡ | Transient |
| LOBOINFK_01310 → | B7L74_02770 | undecaprenyl-phosphate alpha-N-acetylglucosami | L102I (TTA→ATA) | Transient |
| LOBOINFK_01310 → | B7L74_02770 | undecaprenyl-phosphate alpha-N-acetylglucosami | F99I (TTT→ATT) | Transient |
| LOBOINFK_01310 → | B7L74_02770 | undecaprenyl-phosphate alpha-N-acetylglucosami | L98I (TTA→ATA) | Transient |
| LOBOINFK_01310 → | B7L74_02770 | undecaprenyl-phosphate alpha-N-acetylglucosami | R97Q (CGA→CAA) | Transient |
| LOBOINFK_01307 → | B7L74_02755 | lipopolysaccharide assembly protein LapB | S314P (TCC→CCC) | Transient |
| LOBOINFK_01307 → | B7L74_02755 | lipopolysaccharide assembly protein LapB | C373F (TGC→TTC) | Transient |
| LOBOINFK_01307 → | B7L74_02755 | lipopolysaccharide assembly protein LapB | R215P (CGT→CCT) | Transient |
| LOBOINFK_01307 → | B7L74_02755 | lipopolysaccharide assembly protein LapB | A20P (GCA→CCA) | Transient |
| LOBOINFK_01307 → | B7L74_02755 | lipopolysaccharide assembly protein LapB | G221E (GGG→GAG) | Transient |
| LOBOINFK_01307 → | B7L74_02755 | lipopolysaccharide assembly protein LapB | S144R (AGC→AGA) | Transient |
| LOBOINFK_01307 → | B7L74_02755 | lipopolysaccharide assembly protein LapB | D125E (GAC→GAA) | Maintained in later passages |
| LOBOINFK_01307 → | B7L74_02755 | lipopolysaccharide assembly protein LapB | E155G (GAA→GGA) | Transient |
| LOBOINFK_01272 ← | B7L74_11080 | repA protein | N402K (AAT→AAA) | Transient |
| LOBOINFK_01272 ← | B7L74_11080 | repA protein | F403I (TTT→ATT) | Transient |
| LOBOINFK_01269 ← | B7L74_10865 | mobile mystery protein B | D7Y (GAT→TAT) | Transient |
| LOBOINFK_01224 ← | B7L74_09885 | universal stress protein A | L45H (CTT→CAT) | Transient |
| LOBOINFK_01212 ← | B7L74_09790 | methyltransferase | E191G (GAA→GGA) | Transient |
| LOBOINFK_01147 ← | B7L74_07425 | NADH-quinone oxidoreductase subunit G | N423T (AAC→ACC) | Transient |
| LOBOINFK_01147 ← | B7L74_07425 | NADH-quinone oxidoreductase subunit G | I422N (ATT→AAT) | Transient |
| LOBOINFK_01112 ← | B7L74_01975 | octaprenyl diphosphate synthase | F209V (TTC→GTC) | Transient |
| LOBOINFK_01079 ← | B7L74_01810 | xylulokinase | G233A (GGC→GCC) | Transient |
| LOBOINFK_01079 ← | B7L74_01810 | xylulokinase | G234A (GGC→GCC) | Transient |
| LOBOINFK_01063 → | B7L74_01725 | phosphomethylpyrimidine synthase | D468V (GAT→GTT) | Transient |
| LOBOINFK_01060 → | B7L74_01715 | hypothetical protein | I317S (ATC→AGC) ‡ | Transient |
| LOBOINFK_01060 → | B7L74_01715 | hypothetical protein | N316Y (AAT→TAT) | Transient |
| LOBOINFK_01050 → | B7L74_01655 | ribonuclease HI | H118Q (CAC→CAA) | Transient |
| LOBOINFK_01048 ← | B7L74_01645 | hydroxyacylglutathione hydrolase | Q25H (CAA→CAC) | Transient |
| LOBOINFK_01017 ← | B7L74_02315 | cytochrome C assembly protein | F97V (TTT→GTT) | Transient |
| LOBOINFK_01012 → | B7L74_02340 | histone-like protein Hq1 | R6P (CGA→CCA) | Transient |
| LOBOINFK_00985 ← | B7L74_02505 | 23S rRNA pseudouridine(955/2504/2580) syntha | L85M (TTG→ATG) ‡ | Transient |
| LOBOINFK_00970 → | B7L74_04140 | MATE family efflux transporter | T52S (ACA→TCA) | Transient |
| LOBOINFK_00966 ← | B7L74_04120 | valine--tRNA ligase | G341E (GGG→GAG) | Transient |
| LOBOINFK_00931 → | B7L74_03920 | methylisocitrate lyase | K250N (AAA→AAC) | Transient |
| LOBOINFK_00920 → | B7L74_03865 | transcriptional regulator | S121I (AGT→ATT) ‡ | Transient |
| LOBOINFK_00920 → | B7L74_03865 | transcriptional regulator | S121R (AGT→CGT) ‡ | Transient |
| LOBOINFK_00903 → | B7L74_05725 | murein transglycosylase | K340N (AAA→AAC) | Transient |
| LOBOINFK_00884 → | B7L74_05820 | tRNA (cytosine(32)/uridine(32)-2'-O)-methyltran | S56A (TCG→GCG) | Transient |
| LOBOINFK_00884 → | B7L74_05820 | tRNA (cytosine(32)/uridine(32)-2'-O)-methyltran | A148P (GCA→CCA) | Transient |
| LOBOINFK_00884 → | B7L74_05820 | tRNA (cytosine(32)/uridine(32)-2'-O)-methyltran | Q146P (CAG→CCG) | Transient |
| LOBOINFK_00879 → | B7L74_05850 | enhanced entry protein | I70N (ATT→AAT) | Transient |
| LOBOINFK_00879 → | B7L74_05850 | enhanced entry protein | S71R (AGC→CGC) | Transient |
| LOBOINFK_00874 → | B7L74_05885 | protein translocase subunit SecD | V254A (GTT→GCT) | Transient |
| LOBOINFK_00874 → | B7L74_05885 | protein translocase subunit SecD | S253N (AGC→AAC) | Transient |
| LOBOINFK_00868 ← | B7L74_05910 | transcription-repair coupling factor | V397G (GTT→GGT) | Transient |
| LOBOINFK_00853 → | B7L74_05990 | hypothetical protein | L71F (TTA→TTT) | Transient |

| | | | | |
|---|---|---|---|---|
| LOBOINFK_00815 → | B7L74_10690 | hypothetical protein | P181R (CCA→CGA) ‡ | Transient |
| LOBOINFK_00815 → | B7L74_10690 | hypothetical protein | P181A (CCA→GCA) ‡ | Transient |
| LOBOINFK_00810 → | B7L74_10660 | hypothetical protein | Q55R (CAA→CGA) | Transient |
| LOBOINFK_00770 → | B7L74_10445 | ubiquinone/menaquinone biosynthesis C-methyltr | L107R (CTT→CGT) | Transient |
| LOBOINFK_00769 → | B7L74_10440 | hypothetical protein | F78L (TTC→CTC) | Transient |
| LOBOINFK_00760 → | B7L74_10405 | hypothetical protein | V428A (GTT→GCT) ‡ | Transient |
| LOBOINFK_00760 → | B7L74_10405 | hypothetical protein | V428I (GTT→ATT) ‡ | Transient |
| LOBOINFK_00756 ← | B7L74_10385 | two-component sensor histidine kinase | G154C (GGT→TGT) | Transient |
| LOBOINFK_00739 ← | B7L74_09200 | DNA-binding protein | F71I (TTT→ATT) | Maintained in later passages |
| LOBOINFK_00733 ← | B7L74_09225 | hypothetical protein | N229T (AAC→ACC) | Transient |
| LOBOINFK_00732 → | B7L74_09230 | DNA polymerase I | L281F (CTT→TTT) | Transient |
| LOBOINFK_00732 → | B7L74_09230 | DNA polymerase I | L284V (CTT→GTT) | Transient |
| LOBOINFK_00732 → | B7L74_09230 | DNA polymerase I | N287S (AAC→AGC) | Transient |
| LOBOINFK_00730 ← | B7L74_09240 | acyl-CoA thioesterase | M85I (ATG→ATT) | Transient |
| LOBOINFK_00729 ← | B7L74_09245 | amino acid transporter | S352G (AGT→GGT) | Maintained in later passages |
| LOBOINFK_00729 ← | B7L74_09245 | amino acid transporter | C428Y (TGC→TAC) | Transient |
| LOBOINFK_00729 ← | B7L74_09245 | amino acid transporter | D129Y (GAT→TAT) | Transient |
| LOBOINFK_00729 ← | B7L74_09245 | amino acid transporter | V389L (GTG→TTG) | Transient |
| LOBOINFK_00729 ← | B7L74_09245 | amino acid transporter | F86L (TTT→CTT) | Transient |
| LOBOINFK_00729 ← | B7L74_09245 | amino acid transporter | L558P (CTG→CCG) | Transient |
| LOBOINFK_00717 → | B7L74_09300 | ABC transporter substrate-binding protein | K89N (AAA→AAC) | Transient |
| LOBOINFK_00642 ← | B7L74_03325 | prevent-host-death protein | K67N (AAA→AAC) | Transient |
| LOBOINFK_00642 ← | B7L74_03325 | prevent-host-death protein | R75L (CGT→CTT) | Transient |
| LOBOINFK_00618 ← | B7L74_03200 | hypothetical protein | N81H (AAC→CAC) | Transient |
| LOBOINFK_00574 → | B7L74_02995 | crotonase | V286L (GTG→TTG) | Transient |
| LOBOINFK_00574 → | B7L74_02995 | crotonase | N284T (AAC→ACC) | Transient |
| LOBOINFK_00519 ← | B7L74_07085 | GTP pyrophosphokinase | C658S (TGT→TCT) ‡ | Transient |
| LOBOINFK_00519 ← | B7L74_07085 | GTP pyrophosphokinase | C658S (TGT→AGT) ‡ | Transient |
| LOBOINFK_00494 ← | B7L74_06965 | phosphoglucosamine mutase | N275T (AAC→ACC) | Transient |
| LOBOINFK_00492 ← | B7L74_06950 | amino acid permease | M295R (ATG→AGG) | Transient |
| LOBOINFK_00486 ← | B7L74_06925 | GMP synthase (glutamine-hydrolyzing) | Q132R (CAA→CGA) ‡ | Transient |
| LOBOINFK_00481 ← | B7L74_06900 | EamA family transporter | D121A (GAT→GCT) | Transient |
| LOBOINFK_00481 ← | B7L74_06900 | EamA family transporter | S115I (AGC→ATC) | Transient |
| LOBOINFK_00478 ← | B7L74_06885 | FUSC family protein | K141N (AAA→AAC) | Transient |
| LOBOINFK_00478 ← | B7L74_06885 | FUSC family protein | S109C (TCT→TGT) | Transient |
| LOBOINFK_00431 → | B7L74_00275 | hypothetical protein | Q51H (CAA→CAC) | Transient |
| LOBOINFK_00346 ← | B7L74_00745 | hypothetical protein | L256I (CTT→ATT) | Transient |
| LOBOINFK_00346 ← | B7L74_00745 | hypothetical protein | N254K (AAT→AAA) | Transient |
| LOBOINFK_00339 → | B7L74_00790 | prepilin peptidase | F198V (TTT→GTT) | Transient |
| LOBOINFK_00339 → | B7L74_00790 | prepilin peptidase | Q200K (CAA→AAA) | Transient |
| LOBOINFK_00337 → | B7L74_00800 | type 4 pili biogenesis protein (nuleotide-binding p | K287T (AAA→ACA) | Transient |
| LOBOINFK_00318 ← | B7L74_00890 | hypothetical protein | N127Y (AAT→TAT) | Transient |
| LOBOINFK_00300 → | B7L74_00980 | membrane-associated protein | A153G (GCC→GGC) | Transient |
| LOBOINFK_00291 → | B7L74_01025 | hypothetical protein | E270D (GAG→GAC) | Transient |
| LOBOINFK_00290 → | B7L74_01030 | outer membrane protein assembly factor | K59N (AAA→AAC) | Transient |
| LOBOINFK_00288 ← | B7L74_01040 | aldose epimerase | F69C (TTT→TGT) | Transient |
| LOBOINFK_00287 → | B7L74_01045 | hypothetical protein | F71L (TTC→TTA) | Transient |
| LOBOINFK_00271 ← | B7L74_01110+B7L74_01115 | hypothetical protein | V21I (GTT→ATT) ‡ | Maintained in later passages |
| LOBOINFK_00271 ← | B7L74_01110+B7L74_01115 | hypothetical protein | V21D (GTT→GAT) ‡ | Maintained in later passages |
| LOBOINFK_00255 → | B7L74_04195 | NAD-dependent malic enzyme | A401G (GCC→GGC) | Transient |
| LOBOINFK_00255 → | B7L74_04195 | NAD-dependent malic enzyme | S399R (AGC→AGG) | Transient |
| LOBOINFK_00238 ← | B7L74_04280 | asparagine synthetase B | D244H (GAC→CAC) | Transient |
| LOBOINFK_00238 ← | B7L74_04280 | asparagine synthetase B | K236N (AAG→AAT) | Transient |
| LOBOINFK_00218 → | B7L74_04385 | hypothetical protein | G113C (GGC→TGC) | Transient |
| LOBOINFK_00202 ← | B7L74_04460 | aspartate-semialdehyde dehydrogenase | F196L (TTT→TTA) | Transient |
| LOBOINFK_00193 ← | B7L74_04515 | translational GTPase TypA | P299L (CCG→CTG) | Transient |
| LOBOINFK_00153 ← | B7L74_04695 | fatty acid desaturase | N333T (AAC→ACC) | Transient |
| LOBOINFK_00137 → | B7L74_04775 | hypothetical protein | V87G (GTT→GGT) | Transient |
| LOBOINFK_00133 ← | B7L74_04795 | AsmA family protein | M413I (ATG→ATC) | Transient |
| LOBOINFK_00094 → | B7L74_04980 | dihydroorotate dehydrogenase (quinone) | Q341L (CAG→CTG) | Present in all passages |
| LOBOINFK_00094 → | B7L74_04980 | dihydroorotate dehydrogenase (quinone) | T342A (ACG→GCG) | Present in all passages |
| LOBOINFK_00087 ← | B7L74_05015 | hypothetical protein | S109F (TCT→TTT) | Transient |
| LOBOINFK_00076 → | B7L74_05075 | hypothetical protein | M1T (ATG→ACG) † | Maintained in later passages |
| LOBOINFK_00076 → | B7L74_05075 | hypothetical protein | R4H (CGT→CAT) | Transient |
| LOBOINFK_00071 ← | B7L74_05105 | protoheme IX farnesyltransferase | R97P (CGA→CCA) | Transient |
| LOBOINFK_00071 ← | B7L74_05105 | protoheme IX farnesyltransferase | V52G (GTT→GGT) | Transient |
| LOBOINFK_00068 → | B7L74_05120 | GTP-binding protein | V346G (GTG→GGG) | Transient |
| LOBOINFK_00040 ← | B7L74_05280 | ATPase | K420T (AAG→ACG) | Transient |
| LOBOINFK_00031 ← | B7L74_05320 | nicotinate phosphoribosyltransferase | K411Q (AAA→CAA) | Transient |
| LOBOINFK_00017 ← | B7L74_05385 | hypothetical protein | N11T (AAC→ACC) | Transient |
| LOBOINFK_00017 ← | B7L74_05385 | hypothetical protein | T273S (ACC→TCC) | Transient |
| LOBOINFK_00017 ← | B7L74_05385 | hypothetical protein | V271D (GTC→GAC) | Transient |
| LOBOINFK_00006 ← | B7L74_05440 | DNA mismatch repair protein MutS | F162V (TTT→GTT) | Transient |
| damX → | B7L74_00270 | Cell division protein DamX | W224L (TGG→TTG) | Transient |
| carB → | B7L74_06610 | carbamoyl phosphate synthase large subunit | F933L (TTT→TTG) | Transient |
| carB → | B7L74_06610 | carbamoyl phosphate synthase large subunit | A932G (GCC→GGC) | Transient |
| carB → | B7L74_06610 | carbamoyl phosphate synthase large subunit | Q931K (CAA→AAA) | Transient |

| Gene name from Breseq | Gene name from reference genome | Description/Annotation | Amino acid and basepair change | P03 | P05 | P10 | P13 | P16 | P21 | P31 | P42 | P51 | P61 | P67 | Verdict from transcript | Classification by their occurences | Does the intergenic mutation (occuring in most passages) falls in a promoter |
|---|---|---|---|---|---|---|---|---|---|---|---|---|---|---|---|---|---|
| LOBOINFK_01313 ← / → tppB_2 | B7L74_02785 / B7L74_02805 | hypothetical protein/MFS transpo | intergenic (-283/-338) | | | | | | | | | | | | | Maintained in later passages | None |
| LOBOINFK_00892 ← / ← LOBOINFK_00893 | B7L74_05780 / B7L74_05775 | curved DNA-binding protein/L,D | intergenic (-173/+63) | | | | | | | | | | | | | Maintained in later passages | None |
| LOBOINFK_01572 ← / ← LOBOINFK_01573 | B7L74_06385 / B7L74_06390 | malate dehydrogenase/long-chain | intergenic (-177/+18) | | | | | | | | | | | | | Maintained in later passages | None |
| LOBOINFK_01572 ← / ← LOBOINFK_01573 | B7L74_06385 / B7L74_06390 | malate dehydrogenase/long-chain | intergenic (-176/+19) | | | | | | | | | | | | | Maintained in later passages | None |
| – / → LOBOINFK_01440 | / B7L74_08410 | –/hypothetical protein | intergenic (–/-197) | | | | | | | | | | | | | Maintained in later passages | None |
| – / ← LOBOINFK_01633 | / B7L74_06270 | –/hypothetical protein | intergenic (–/+57) | | | | | | | | | | | | | Maintained in later passages | None |
| LOBOINFK_00877 ← / → LOBOINFK_00878 | B7L74_05870 / B7L74_05855 | hypothetical protein/L,D-transpep | intergenic (-645/-272) | | | | | | | | | | | | | Present in all passages | None |
| LOBOINFK_01964 → / – | B7L74_07985 / | ribonucleotide-diphosphate reduc | intergenic (+52/–) | | | | | | | | | | | | | Present in all passages | None |
| LOBOINFK_01942 → / – | B7L74_08845 / | molecular chaperone GroEL/– | intergenic (+36/–) | | | | | | | | | | | | | Present in all passages | None |
| LOBOINFK_01139 ← / ← LOBOINFK_01140 | B7L74_07385 / B7L74_07390 | tRNA-Met/NADH-quinone oxido | intergenic (-169/+72) | | | | | | | | | | | | | Present in all passages | None |
| LOBOINFK_00410 ← / → LOBOINFK_00411 | B7L74_00390 / B7L74_00380 | 2-octaprenyl-3-methyl-6-methox | intergenic (-161/-147) | | | | | | | | | | | | | Present in all passages | None |
| LOBOINFK_00892 ← / ← LOBOINFK_00893 | B7L74_05780 / B7L74_05775 | curved DNA-binding protein/L,D | intergenic (-157/+79) | | | | | | | | | | | | | Present in all passages | None |
| LOBOINFK_00877 ← / → LOBOINFK_00878 | B7L74_05870 / B7L74_05855 | hypothetical protein/L,D-transpep | intergenic (-639/-278) | | | | | | | | | | | | | Present in all passages | None |
| LOBOINFK_01964 → / – | B7L74_07985 / | ribonucleotide-diphosphate reduc | intergenic (+51/–) | | | | | | | | | | | | | Present in all passages | None |
| – / → LOBOINFK_00660 | / B7L74_09615 | –/DNA topoisomerase IV subunit | intergenic (–/-340) | | | | | | | | | | | | | Present in all passages | None |
| LOBOINFK_01724 ← / – | B7L74_02705 / | N-ethylammeline chlorohydrolas | intergenic (-386/–) | | | | | | | | | | | | | Present in all passages | None |
| LOBOINFK_01313 ← / → tppB_2 | B7L74_02785 / B7L74_02805 | hypothetical protein/MFS transpo | intergenic (-244/-377) | | | | | | | | | | | | | Present in all passages | None |
| – / → LOBOINFK_00660 | / B7L74_09615 | –/DNA topoisomerase IV subunit | intergenic (–/-311) | | | | | | | | | | | | | Present in all passages | None |
| LOBOINFK_01817 → / – | B7L74_03805 / | alpha/beta hydrolase/– | intergenic (+169/–) | | | | | | | | | | | | | Present in all passages | None |
| LOBOINFK_00892 ← / ← LOBOINFK_00893 | B7L74_05780 / B7L74_05775 | curved DNA-binding protein/L,D | intergenic (-190/+46) | | | | | | | | | | | | | Present in all passages | None |
| LOBOINFK_00877 ← / → LOBOINFK_00878 | B7L74_05870 / B7L74_05855 | hypothetical protein/L,D-transpep | intergenic (-654/-263) | | | | | | | | | | | | | Present in all passages | None |
| – / ← LOBOINFK_01633 | / B7L74_06270 | –/hypothetical protein | intergenic (–/+86) | | | | | | | | | | | | | Present in all passages | None |
| LOBOINFK_00839 ← / ← phrA | B7L74_06060 / B7L74_06050 | hypothetical protein/Deoxyribod | intergenic (-309/+455) | | | | | | | | | | | | | Present in all passages | None |
| LOBOINFK_01942 → / – | B7L74_08845 / | molecular chaperone GroEL/– | intergenic (+103/–) | | | | | | | | | | | | | Present in all passages | None |
| LOBOINFK_01724 ← / – | B7L74_02705 / | N-ethylammeline chlorohydrolas | intergenic (-470/–) | | | | | | | | | | | | | Present in all passages | None |
| – / ← LOBOINFK_01633 | / B7L74_06270 | –/hypothetical protein | intergenic (–/+80) | | | | | | | | | | | | | Present in all passages | None |
| – / → LOBOINFK_01440 | / B7L74_08410 | –/hypothetical protein | intergenic (–/-191) | | | | | | | | | | | | | Present in all passages | None |
| ybhL ← / ← LOBOINFK_00275 | B7L74_01100 / B7L74_01091 | Inner membrane protein YbhL/hy | intergenic (-164/+517) | | | | | | | | | | | | | Present in all passages | None |
| LOBOINFK_00877 ← / → LOBOINFK_00878 | B7L74_05870 / B7L74_05855 | hypothetical protein/L,D-transpep | intergenic (-648/-269) | | | | | | | | | | | | | Present in all passages | None |
| LOBOINFK_01964 → / – | B7L74_07985 / | ribonucleotide-diphosphate reduc | intergenic (+50/–) | | | | | | | | | | | | | Present in all passages | None |
| LOBOINFK_01942 → / – | B7L74_08845 / | molecular chaperone GroEL/– | intergenic (+33/–) | | | | | | | | | | | | | Present in all passages | None |
| – / ← LOBOINFK_01633 | / B7L74_06270 | –/hypothetical protein | intergenic (–/+122) | | | | | | | | | | | | | Present in all passages | None |
| LOBOINFK_01998 → / – | B7L74_08140 / | Na+/H+ antiporter subunit G/– | intergenic (+87/–) | | | | | | | | | | | | | Present in all passages | None |
| LOBOINFK_01962 ← / → LOBOINFK_01963 | B7L74_07970 / B7L74_07980 | RNA pyrophosphohydrolase/ribo | intergenic (-213/-304) | | | | | | | | | | | | | Present in all passages | None |
| LOBOINFK_01724 ← / – | B7L74_02705 / | N-ethylammeline chlorohydrolas | intergenic (-389/–) | | | | | | | | | | | | | Present in all passages | None |
| LOBOINFK_00056 → / ← LOBOINFK_00057 | B7L74_05190 / B7L74_05185 | beta-carotene 15,15'-monooxyge | intergenic (+21/+30) | | | | | | | | | | | | | Maintained in later passages | Promoter: LOBOINFK_00057, No gene expression change in this gene |
| LOBOINFK_00284 ← / ← LOBOINFK_00285 | B7L74_01055 / B7L74_01050 | glutamate--tRNA ligase/hypothet | intergenic (-58/+251) | | | | | | | | | | | | | Present in all passages | Promoter: LOBOINFK_00284, No gene expression change in this gene |
| LOBOINFK_00284 ← / ← LOBOINFK_00285 | B7L74_01055 / B7L74_01050 | glutamate--tRNA ligase/hypothet | intergenic (-57/+252) | | | | | | | | | | | | | Present in all passages | Promoter: LOBOINFK_00284, No gene expression change in this gene |
| LOBOINFK_00284 ← / ← LOBOINFK_00285 | B7L74_01055 / B7L74_01050 | glutamate--tRNA ligase/hypothet | intergenic (-56/+253) | | | | | | | | | | | | | Present in all passages | Promoter: LOBOINFK_00284, No gene expression change in this gene |
| LOBOINFK_00303 ← / ← LOBOINFK_00304 | B7L74_00970 / B7L74_00968 | hypothetical protein/5S ribosoma | intergenic (-138/+32) | | | | | | | | | | | | | Present in all passages | Promoter: LOBOINFK_00303, No gene expression change in this gene |
| LOBOINFK_00303 ← / ← LOBOINFK_00304 | B7L74_00970 / B7L74_00967 | hypothetical protein/5S ribosoma | intergenic (-141/+29) | | | | | | | | | | | | | Present in all passages | Promoter: LOBOINFK_00303, No gene expression change in this gene |
| LOBOINFK_00303 ← / ← LOBOINFK_00304 | B7L74_00970 / B7L74_00966 | hypothetical protein/5S ribosoma | intergenic (-140/+30) | | | | | | | | | | | | | Present in all passages | Promoter: LOBOINFK_00303, No gene expression change in this gene |
| LOBOINFK_00303 ← / ← LOBOINFK_00304 | B7L74_00970 / B7L74_00965 | hypothetical protein/5S ribosoma | intergenic (-143/+27) | | | | | | | | | | | | | Present in all passages | Promoter: LOBOINFK_00303, No gene expression change in this gene |
| LOBOINFK_00399 ← / ← LOBOINFK_00400 | B7L74_00440 / B7L74_00435 | protein TolA/phosphoglycerol tra | intergenic (-87/+82) | | | | | | | | | | | | | Maintained in later passages | Promoter: LOBOINFK_00399, No gene expression change in this gene |
| LOBOINFK_00399 ← / ← LOBOINFK_00400 | B7L74_00440 / B7L74_00435 | protein TolA/phosphoglycerol tra | intergenic (-86/+83) | | | | | | | | | | | | | Maintained in later passages | Promoter: LOBOINFK_00399, No gene expression change in this gene |
| LOBOINFK_00636 ← / ← LOBOINFK_00637 | B7L74_03295 / B7L74_03300 | alpha-ketoacid dehydrogenase su | intergenic (-73/+132) | | | | | | | | | | | | | Maintained in later passages | Promoter: LOBOINFK_00636, No gene expression change in this gene |
| LOBOINFK_00772 → / ← LOBOINFK_00773 | B7L74_10455 / B7L74_10470 | ubiquinone biosynthesis regulato | intergenic (+130/+275) | | | | | | | | | | | | | Present in all passages | Promoter: LOBOINFK_00773, No gene expression change in this gene |
| LOBOINFK_01127 ← / ← LOBOINFK_01128 | B7L74_07315 / B7L74_02145 | DNA repair protein RadA/tRNA- | intergenic (-109/+126) | | | | | | | | | | | | | Present in all passages | Promoter: LOBOINFK_01127, No gene expression change in this gene |
| LOBOINFK_01309 → / → LOBOINFK_01310 | B7L74_02765 / B7L74_02770 | competence protein ComEA/unde | intergenic (+24/-188) | | | | | | | | | | | | | Present in all passages | Promoter: LOBOINFK_01310, No gene expression change in this gene |
| LOBOINFK_01309 → / → LOBOINFK_01310 | B7L74_02765 / B7L74_02770 | competence protein ComEA/unde | intergenic (+23/-189) | | | | | | | | | | | | | Present in all passages | Promoter: LOBOINFK_01310, No gene expression change in this gene |
| LOBOINFK_01478 → / ← LOBOINFK_01479 | B7L74_06665 / B7L74_06660 | nucleotide exchange factor GrpE | intergenic (+76/+54) | | | | | | | | | | | | | Present in all passages | Promoter: LOBOINFK_01479, Early down |
| LOBOINFK_01478 → / ← LOBOINFK_01479 | B7L74_06665 / B7L74_06660 | nucleotide exchange factor GrpE | intergenic (+73/+57) | | | | | | | | | | | | | Present in all passages | Promoter: LOBOINFK_01479, Early down |
| LOBOINFK_01478 → / ← LOBOINFK_01479 | B7L74_06665 / B7L74_06660 | nucleotide exchange factor GrpE | intergenic (+70/+60) | | | | | | | | | | | | | Present in all passages | Promoter: LOBOINFK_01479, Early down |
| LOBOINFK_01485 ← / ← LOBOINFK_01486 | B7L74_06635 / B7L74_06630 | ATPase/MFS transporter | intergenic (-39/+32) | | | | | | | | | | | | | Maintained in later passages | Promoter: LOBOINFK_01485, No gene expression change in this gene |
| LOBOINFK_01485 ← / ← LOBOINFK_01486 | B7L74_06635 / B7L74_06630 | ATPase/MFS transporter | intergenic (-38/+33) | | | | | | | | | | | | | Maintained in later passages | Promoter: LOBOINFK_01485, No gene expression change in this gene |
| LOBOINFK_01485 ← / ← LOBOINFK_01486 | B7L74_06635 / B7L74_06630 | ATPase/MFS transporter | intergenic (-33/+38) | | | | | | | | | | | | | Present in all passages | Promoter: LOBOINFK_01485, No gene expression change in this gene |
| LOBOINFK_01485 ← / ← LOBOINFK_01486 | B7L74_06635 / B7L74_06630 | ATPase/MFS transporter | intergenic (-44/+27) | | | | | | | | | | | | | Present in all passages | Promoter: LOBOINFK_01485, No gene expression change in this gene |
| LOBOINFK_01487 ← / ← LOBOINFK_01488 | B7L74_06625 / B7L74_06620 | ATPase/hypothetical protein | intergenic (-34/+72) | | | | | | | | | | | | | Present in all passages | Promoter: LOBOINFK_01487, No gene expression change in this gene |
| LOBOINFK_01487 ← / ← LOBOINFK_01488 | B7L74_06625 / B7L74_06620 | ATPase/hypothetical protein | intergenic (-45/+61) | | | | | | | | | | | | | Present in all passages | Promoter: LOBOINFK_01487, No gene expression change in this gene |
| LOBOINFK_01602 → / → LOBOINFK_01603 | B7L74_00215 / B7L74_00210 | oligopeptidase A/ketoacyl-ACP s | intergenic (+29/-52) | | | | | | | | | | | | | Maintained in later passages | Promoter: LOBOINFK_01603, Early down |
| LOBOINFK_01602 → / → LOBOINFK_01603 | B7L74_00215 / B7L74_00210 | oligopeptidase A/ketoacyl-ACP s | intergenic (+28/-53) | | | | | | | | | | | | | Maintained in later passages | Promoter: LOBOINFK_01603, Early down |
| LOBOINFK_01627 → / ← LOBOINFK_01628 | B7L74_00075 / B7L74_00065 | hypothetical protein/shikimate de | intergenic (+234/+110) | | | | | | | | | | | | | Maintained in later passages | Promoter: LOBOINFK_01628, No gene expression change in this gene |
| LOBOINFK_01681 → / → LOBOINFK_01682 | B7L74_09170 / B7L74_09165 | type I glyceraldehyde-3-phospha | intergenic (+24/-63) | | | | | | | | | | | | | Maintained in later passages | Promoter: LOBOINFK_01682, No gene expression change in this gene |
| LOBOINFK_01770 ← / ← LOBOINFK_01771 | B7L74_08730 / B7L74_08735 | ATPase/hypothetical protein | intergenic (-32/+35) | | | | | | | | | | | | | Present in all passages | Promoter: LOBOINFK_01770, No gene expression change in this gene |
| LOBOINFK_01770 ← / ← LOBOINFK_01771 | B7L74_08730 / B7L74_08735 | ATPase/hypothetical protein | intergenic (-36/+31) | | | | | | | | | | | | | Present in all passages | Promoter: LOBOINFK_01770, No gene expression change in this gene |
| LOBOINFK_01770 ← / ← LOBOINFK_01771 | B7L74_08730 / B7L74_08735 | ATPase/hypothetical protein | intergenic (-26/+41) | | | | | | | | | | | | | Present in all passages | Promoter: LOBOINFK_01770, No gene expression change in this gene |
| LOBOINFK_01770 ← / ← LOBOINFK_01771 | B7L74_08730 / B7L74_08735 | ATPase/hypothetical protein | intergenic (-16/+51) | | | | | | | | | | | | | Present in all passages | Promoter: LOBOINFK_01770, No gene expression change in this gene |
| LOBOINFK_01770 ← / ← LOBOINFK_01771 | B7L74_08730 / B7L74_08735 | ATPase/hypothetical protein | intergenic (-20/+47) | | | | | | | | | | | | | Present in all passages | Promoter: LOBOINFK_01770, No gene expression change in this gene |
| dnaA → / → LOBOINFK_01790 | B7L74_00010 / B7L74_00015 | chromosomal replication initiator | intergenic (+41/-103) | | | | | | | | | | | | | Maintained in later passages | Promoter: LOBOINFK_01790, Early down |
| dnaA → / → LOBOINFK_01790 | B7L74_00010 / B7L74_00015 | chromosomal replication initiator | intergenic (+40/-104) | | | | | | | | | | | | | Maintained in later passages | Promoter: LOBOINFK_01790, Early down |
| ybhL ← / ← LOBOINFK_00275 | B7L74_01100 / B7L74_01090 | Inner membrane protein YbhL/hy | intergenic (-130/+551) | | | | | | | | | | | | | Present in all passages | Promoter: YbhL, No gene expression change in this gene |
| LOBOINFK_00866 → / ← LOBOINFK_00867 | B7L74_05920 / B7L74_05915 | HAD family hydrolase/hypotheti | intergenic (+63/+143) | | | | | | | | | | | | | Transient | |
| – / → LOBOINFK_02021 | / B7L74_10120 | –/NAD(P) transhydrogenase subu | intergenic (–/-197) | | | | | | | | | | | | | Transient | |
| LOBOINFK_00702 ← / ← LOBOINFK_00703 | B7L74_09385 / B7L74_09375 | hypothetical protein/hypothetical | intergenic (-270/+32) | | | | | | | | | | | | | Transient | |
| – / → LOBOINFK_01936 | / B7L74_08875 | –/hypothetical protein | intergenic (–/-328) | | | | | | | | | | | | | Transient | |
| – / → LOBOINFK_01936 | / B7L74_08875 | –/hypothetical protein | intergenic (–/-438) | | | | | | | | | | | | | Transient | |
| – / → LOBOINFK_01865 | / B7L74_01625 | –/bifunctional methylenetetrahyd | intergenic (–/-213) | | | | | | | | | | | | | Transient | |
| LOBOINFK_00636 ← / ← LOBOINFK_00637 | B7L74_03295 / B7L74_03300 | alpha-ketoacid dehydrogenase su | intergenic (-131/+74) | | | | | | | | | | | | | Transient | |
| fadE_1 → / → LOBOINFK_00573 | B7L74_02980 / B7L74_02996 | acyl-CoA dehydrogenase/acetyl-C | intergenic (+194/-351) | | | | | | | | | | | | | Transient | |
| LOBOINFK_01700 → / – | B7L74_09070 / | hybrid sensor histidine kinase/res | intergenic (+312/–) | | | | | | | | | | | | | Transient | |
| LOBOINFK_01599 → / ← LOBOINFK_01600 | B7L74_06525 / B7L74_06530 | protein ScvA/pyridine nucleotide | intergenic (+41/+15) | | | | | | | | | | | | | Transient | |
| LOBOINFK_00492 ← / → LOBOINFK_00493 | B7L74_06950 / B7L74_06960 | amino acid permease/hypothetica | intergenic (-339/-169) | | | | | | | | | | | | | Transient | |
| LOBOINFK_01481 → / → LOBOINFK_01482 | B7L74_06650 / B7L74_06645 | molecular chaperone DnaK/mole | intergenic (+91/-52) | | | | | | | | | | | | | Transient | |
| LOBOINFK_01285 → / → LOBOINFK_01286 | B7L74_11010 / B7L74_10995 | transcriptional regulator/hypothet | intergenic (+385/-485) | | | | | | | | | | | | | Transient | |
| LOBOINFK_01212 ← / ← LOBOINFK_01213 | B7L74_09790 / B7L74_09805 | methyltransferase/peptidase M16 | intergenic (-245/+251) | | | | | | | | | | | | | Transient | |
| LOBOINFK_01175 → / ← LOBOINFK_01176 | B7L74_07865 / B7L74_07860 | peptidase/hypothetical protein | intergenic (+119/+53) | | | | | | | | | | | | | Transient | |
| LOBOINFK_00425 ← / ← LOBOINFK_00426 | B7L74_00310 / B7L74_00300 | hypothetical protein/channel prot | intergenic (-447/+107) | | | | | | | | | | | | | Transient | |
| LOBOINFK_00402 ← / ← LOBOINFK_00403 | B7L74_00425 / B7L74_00420 | entericidin B/sulfatase | intergenic (-104/+150) | | | | | | | | | | | | | Transient | |
| LOBOINFK_00342 → / ← LOBOINFK_00343 | B7L74_00775 / B7L74_00760 | cell division protein ZapD/7,8-di | intergenic (+300/+68) | | | | | | | | | | | | | Transient | |
| LOBOINFK_00305 ← | B7L74_00960 | 23S ribosomal RNA | oncoding (1715/4342 nt | | | | | | | | | | | | | Transient | |
| LOBOINFK_00305 ← | B7L74_00960 | 23S ribosomal RNA | oncoding (2697/4342 nt | | | | | | | | | | | | | Transient | |
| LOBOINFK_00259 ← / → LOBOINFK_00260 | B7L74_04170 / B7L74_04165 | TetR family transcriptional regula | intergenic (-22/-228) | | | | | | | | | | | | | Transient | |
| LOBOINFK_01085 ← / → LOBOINFK_01086 | B7L74_01840 / B7L74_01845 | protease SohB/amino acid perme | intergenic (-147/-83) | | | | | | | | | | | | | Transient | |
| LOBOINFK_01031 → / ← LOBOINFK_01032 | B7L74_02240 / B7L74_02220 | hypothetical protein/mechanosen | intergenic (+642/+189) | | | | | | | | | | | | | Transient | |
| LOBOINFK_01031 → / ← LOBOINFK_01032 | B7L74_02240 / B7L74_02220 | hypothetical protein/mechanosen | intergenic (+601/+230) | | | | | | | | | | | | | Transient | |
| LOBOINFK_01031 → / ← LOBOINFK_01032 | B7L74_02240 / B7L74_02220 | hypothetical protein/mechanosen | intergenic (+596/+235) | | | | | | | | | | | | | Transient | |
| LOBOINFK_01011 ← / → LOBOINFK_01012 | B7L74_02345 / B7L74_02340 | hypothetical protein/histone-like | intergenic (-114/-177) | | | | | | | | | | | | | Transient | |
| – / → LOBOINFK_02021 | / B7L74_10120 | –/NAD(P) transhydrogenase subu | intergenic (–/-230) | | | | | | | | | | | | | Transient | |
| – / → LOBOINFK_02021 | / B7L74_10120 | –/NAD(P) transhydrogenase subu | intergenic (–/-232) | | | | | | | | | | | | | Transient | |
| yjdF → / ← LOBOINFK_00676 | B7L74_09540 / B7L74_09530 | putative protein YjdF/hypothetica | intergenic (+413/+143) | | | | | | | | | | | | | Transient | |
| LOBOINFK_01785 ← / → LOBOINFK_01786 | B7L74_10840 / B7L74_10845 | phosphoenolpyruvate carboxykin | intergenic (-96/-80) | | | | | | | | | | | | | Transient | |
| LOBOINFK_00636 ← / ← LOBOINFK_00637 | B7L74_03295 / B7L74_03300 | alpha-ketoacid dehydrogenase su | intergenic (-124/+81) | | | | | | | | | | | | | Transient | |
| fadE_1 → / → LOBOINFK_00573 | B7L74_02980 / B7L74_02995 | acyl-CoA dehydrogenase/acetyl-C | intergenic (+195/-350) | | | | | | | | | | | | | Transient | |
| – / → LOBOINFK_01680 | / B7L74_09175 | –/transketolase | intergenic (–/-121) | | | | | | | | | | | | | Transient | |
| LOBOINFK_01632 → / – | B7L74_00040 / | antitoxin/– | intergenic (+97/–) | | | | | | | | | | | | | Transient | |
| LOBOINFK_01599 → / ← LOBOINFK_01600 | B7L74_06525 / B7L74_06530 | protein ScvA/pyridine nucleotide | intergenic (+42/+14) | | | | | | | | | | | | | Transient | |
| LOBOINFK_00467 ← / ← LOBOINFK_00468 | B7L74_06830 / B7L74_06845 | 50S ribosomal protein L20/50S r | intergenic (-477/+163) | | | | | | | | | | | | | Transient | |
| LOBOINFK_00467 ← / ← LOBOINFK_00468 | B7L74_06830 / B7L74_06845 | 50S ribosomal protein L20/50S r | intergenic (-420/+220) | | | | | | | | | | | | | Transient | |
| LOBOINFK_00467 ← / ← LOBOINFK_00468 | B7L74_06830 / B7L74_06845 | 50S ribosomal protein L20/50S r | intergenic (-248/+392) | | | | | | | | | | | | | Transient | |
| LOBOINFK_01491 → / → carB | B7L74_06615 / B7L74_06610 | carbamoyl phosphate synthase sm | intergenic (+36/-46) | | | | | | | | | | | | | Transient | |
| LOBOINFK_01485 ← / ← LOBOINFK_01486 | B7L74_06635 / B7L74_06630 | ATPase/MFS transporter | intergenic (-23/+48) | | | | | | | | | | | | | Transient | |
| – / ← LOBOINFK_01474 | / B7L74_06685 | –/DNA repair protein RecN | intergenic (–/+94) | | | | | | | | | | | | | Transient | |
| LOBOINFK_01433 ← / → LOBOINFK_01434 | B7L74_06220 / B7L74_06225 | hypothetical protein/hypothetical | intergenic (-198/-134) | | | | | | | | | | | | | Transient | |
| LOBOINFK_01359 ← / ← LOBOINFK_01360 | B7L74_01415 / B7L74_01420 | single-stranded DNA-binding pro | intergenic (-48/+2) | | | | | | | | | | | | | Transient | |
| LOBOINFK_01313 ← / → tppB_2 | B7L74_02785 / B7L74_02805 | hypothetical protein/MFS transpo | intergenic (-219/-402) | | | | | | | | | | | | | Transient | |
| LOBOINFK_01212 ← / ← LOBOINFK_01213 | B7L74_09790 / B7L74_09805 | methyltransferase/peptidase M16 | intergenic (-204/+292) | | | | | | | | | | | | | Transient | |
| LOBOINFK_01114 → / → LOBOINFK_01115 | B7L74_07255 / B7L74_07260 | hypothetical protein/type II citrat | intergenic (+104/-130) | | | | | | | | | | | | | Transient | |
| LOBOINFK_00410 ← / → LOBOINFK_00411 | B7L74_00390 / B7L74_00380 | 2-octaprenyl-3-methyl-6-methox | intergenic (-97/-211) | | | | | | | | | | | | | Transient | |
| LOBOINFK_00342 → / ← LOBOINFK_00343 | B7L74_00775 / B7L74_00760 | cell division protein ZapD/7,8-di | intergenic (+221/+147) | | | | | | | | | | | | | Transient | |
| LOBOINFK_00305 ← | B7L74_00960 | 23S ribosomal RNA | oncoding (3293/4342 nt) | | | | | | | | | | | | | Transient | |
| LOBOINFK_00291 → / ← LOBOINFK_00292 | B7L74_01025 / B7L74_01020 | hypothetical protein/hypothetical | intergenic (+31/+39) | | | | | | | | | | | | | Transient | |
| LOBOINFK_00225 → / ← LOBOINFK_00226 | B7L74_04345 / B7L74_04340 | methylglyoxal synthase/polyribo | intergenic (+66/+26) | | | | | | | | | | | | | Transient | |
| LOBOINFK_01088 → / ← LOBOINFK_01089 | B7L74_01850 / B7L74_01860 | hypothetical protein/hypothetical | intergenic (+661/+54) | | | | | | | | | | | | | Transient | |
| LOBOINFK_01058 → / ← LOBOINFK_01059 | B7L74_01705 / B7L74_01710 | acyl-CoA synthetase/phosphoribo | intergenic (+74/+188) | | | | | | | | | | | | | Transient | |
| LOBOINFK_01058 → / ← LOBOINFK_01059 | B7L74_01705 / B7L74_01710 | acyl-CoA synthetase/phosphoribo | intergenic (+69/+193) | | | | | | | | | | | | | Transient | |
| LOBOINFK_00929 → / → LOBOINFK_00930 | B7L74_03910 / B7L74_03915 | GMP synthase/ATPase | intergenic (+184/-25) | | | | | | | | | | | | | Transient | |
| LOBOINFK_00772 → / ← LOBOINFK_00773 | B7L74_10455 / B7L74_10470 | ubiquinone biosynthesis regulato | intergenic (+146/+259) | | | | | | | | | | | | | Transient | |
| LOBOINFK_00704 → / → LOBOINFK_00705 | B7L74_09370 / B7L74_09365 | superoxide dismutase/hypothetica | intergenic (+75/-41) | | | | | | | | | | | | | Transient | |
| LOBOINFK_00704 → / → LOBOINFK_00705 | B7L74_09370 / B7L74_09365 | superoxide dismutase/hypothetica | intergenic (+74/-42) | | | | | | | | | | | | | Transient | |
| LOBOINFK_01531 → / → LOBOINFK_01532 | B7L74_09010 / B7L74_09015 | integration host factor/hypothetic | intergenic (+45/-63) | | | | | | | | | | | | | Transient | |
| – / ← LOBOINFK_00443 | / B7L74_06690 | –/hypothetical protein | intergenic (–/+116) | | | | | | | | | | | | | Transient | |
| LOBOINFK_01459 ← / → LOBOINFK_01460 | B7L74_08305 / B7L74_08300 | hypothetical protein/hypothetical | intergenic (-121/-172) | | | | | | | | | | | | | Transient | |
| LOBOINFK_01368 → / → LOBOINFK_01369 | B7L74_01470 / B7L74_01490 | catalase/hypothetical protein | intergenic (+296/-567) | | | | | | | | | | | | | Transient | |
| LOBOINFK_01359 ← / ← LOBOINFK_01360 | B7L74_01415 / B7L74_01420 | single-stranded DNA-binding pro | intergenic (-46/+4) | | | | | | | | | | | | | Transient | |
| LOBOINFK_01058 → / ← LOBOINFK_01059 | B7L74_01705 / B7L74_01710 | acyl-CoA synthetase/phosphoribo | intergenic (+75/+187) | | | | | | | | | | | | | Transient | |
| LOBOINFK_01862 ← / → LOBOINFK_01863 | B7L74_09640 / B7L74_09635 | demethoxyubiquinone hydroxyla | intergenic (-140/-29) | | | | | | | | | | | | | Transient | |
| LOBOINFK_01856 ← / ← LOBOINFK_01857 | B7L74_09675 / B7L74_09665 | ribosomal RNA small subunit me | intergenic (-84/+323) | | | | | | | | | | | | | Transient | |
| LOBOINFK_01761 ← / ← LOBOINFK_01762 | B7L74_08670 / B7L74_08675 | hypothetical protein/dCTP deami | intergenic (-44/+110) | | | | | | | | | | | | | Transient | |
| LOBOINFK_01467 → / → LOBOINFK_01468 | B7L74_08265 / B7L74_08260 | hypothetical protein/hypothetical | intergenic (+297/-42) | | | | | | | | | | | | | Transient | |
| LOBOINFK_01378 ← / – | B7L74_01530 / | hypothetical protein/– | intergenic (-182/–) | | | | | | | | | | | | | Transient | |
| LOBOINFK_01378 ← / – | B7L74_01530 / | hypothetical protein/– | intergenic (-175/–) | | | | | | | | | | | | | Transient | |
| LOBOINFK_01378 ← / – | B7L74_01530 / | hypothetical protein/– | intergenic (-168/–) | | | | | | | | | | | | | Transient | |
| LOBOINFK_01378 ← / – | B7L74_01530 / | hypothetical protein/– | intergenic (-161/–) | | | | | | | | | | | | | Transient | |
| LOBOINFK_01378 ← / – | B7L74_01530 / | hypothetical protein/– | intergenic (-147/–) | | | | | | | | | | | | | Transient | |
| LOBOINFK_01304 → / → LOBOINFK_01305 | B7L74_02740 / B7L74_02745 | cytidylate kinase/30S ribosomal p | intergenic (+67/-77) | | | | | | | | | | | | | Transient | |
| LOBOINFK_01212 ← / ← LOBOINFK_01213 | B7L74_09790 / B7L74_09805 | methyltransferase/peptidase M16 | intergenic (-212/+284) | | | | | | | | | | | | | Transient | |
| LOBOINFK_00347 ← / ← LOBOINFK_00348 | B7L74_00740 / B7L74_00735 | hypothetical protein/UDP-3-O-[3 | intergenic (-421/+149) | | | | | | | | | | | | | Transient | |
| LOBOINFK_00342 → / ← LOBOINFK_00343 | B7L74_00775 / B7L74_00760 | cell division protein ZapD/7,8-di | intergenic (+232/+136) | | | | | | | | | | | | | Transient | |

| |
|---|
| 5-10 |
| 10-30 |
| 30-50 |
| 50-70 |
| 70-90 |
| 90-99 |
| 100 |

| | | | | | | | | | | | | | | | | |
|---|---|---|---|---|---|---|---|---|---|---|---|---|---|---|---|---|
| LOBOINFK_00305 ← / ← LOBOINFK_00306 | B7L74_00960 / B7L74_00957 | 23S ribosomal RNA/tRNA-Ala | intergenic (-138/+61) | | | | | | | | | | | | | Transient |
| LOBOINFK_00225 → / ← LOBOINFK_00226 | B7L74_04345 / B7L74_04340 | methylglyoxal synthase/polyribo | intergenic (+41/+51) | | | | | | | | | | | | | Transient |
| LOBOINFK_00099 ← / ← LOBOINFK_00100 | B7L74_04955 / B7L74_04940 | cytochrome d terminal oxidase su | intergenic (-132/+338) | | | | | | | | | | | | | Transient |
| coaE → / ← LOBOINFK_01110 | B7L74_01960 / B7L74_01966 | Dephospho-CoA kinase/hypothet | intergenic (+12/+37) | | | | | | | | | | | | | Transient |
| LOBOINFK_01031 → / ← LOBOINFK_01032 | B7L74_02240 / B7L74_02220 | hypothetical protein/mechanosen | intergenic (+561/+270) | | | | | | | | | | | | | Transient |
| LOBOINFK_00871 → / → mvl | B7L74_05900 / B7L74_05896 | hypothetical protein/Lectin MVL | intergenic (+77/-151) | | | | | | | | | | | | | Transient |
| – / ← LOBOINFK_02015 | / B7L74_08455 | –/transporter | intergenic (–/+76) | | | | | | | | | | | | | Transient |
| LOBOINFK_02012 → / ← LOBOINFK_02013 | B7L74_10080 / B7L74_10075 | hypothetical protein/ATPase | intergenic (+39/+58) | | | | | | | | | | | | | Transient |
| LOBOINFK_02012 → / ← LOBOINFK_02013 | B7L74_10080 / B7L74_10075 | hypothetical protein/ATPase | intergenic (+31/+66) | | | | | | | | | | | | | Transient |
| LOBOINFK_01998 → / – | B7L74_08140 / | Na+/H+ antiporter subunit G/– | intergenic (+297/–) | | | | | | | | | | | | | Transient |
| – / ← LOBOINFK_01990 | / B7L74_08180 | –/sodium:proton antiporter | intergenic (–/+89) | | | | | | | | | | | | | Transient |
| – / → LOBOINFK_00660 | / B7L74_09615 | –/DNA topoisomerase IV subunit | intergenic (–/-350) | | | | | | | | | | | | | Transient |
| LOBOINFK_01788 → / → dnaA | B7L74_10855 / B7L74_00010 | aspartate carbamoyltransferase/ch | intergenic (+69/-153) | | | | | | | | | | | | | Transient |
| LOBOINFK_01788 → / → dnaA | B7L74_10855 / B7L74_00010 | aspartate carbamoyltransferase/ch | intergenic (+25/-197) | | | | | | | | | | | | | Transient |
| LOBOINFK_01724 ← / – | B7L74_02705 / | N-ethylammeline chlorohydrolas | intergenic (-454/–) | | | | | | | | | | | | | Transient |
| LOBOINFK_01636 ← / ← LOBOINFK_01637 | B7L74_06285 / B7L74_06295 | 4-hydroxy-tetrahydrodipicolinate | intergenic (-169/+68) | | | | | | | | | | | | | Transient |
| LOBOINFK_01627 → / ← LOBOINFK_01628 | B7L74_00075 / B7L74_00065 | hypothetical protein/shikimate de | intergenic (+232/+112) | | | | | | | | | | | | | Transient |
| LOBOINFK_01627 → / ← LOBOINFK_01628 | B7L74_00075 / B7L74_00065 | hypothetical protein/shikimate de | intergenic (+230/+114) | | | | | | | | | | | | | Transient |
| LOBOINFK_01467 → / → LOBOINFK_01468 | B7L74_08265 / B7L74_08260 | hypothetical protein/hypothetical | intergenic (+308/-31) | | | | | | | | | | | | | Transient |
| LOBOINFK_01467 → / → LOBOINFK_01468 | B7L74_08265 / B7L74_08260 | hypothetical protein/hypothetical | intergenic (+289/-50) | | | | | | | | | | | | | Transient |
| – / → LOBOINFK_01440 | / B7L74_08410 | –/hypothetical protein | intergenic (–/-152) | | | | | | | | | | | | | Transient |
| LOBOINFK_01313 ← / → tppB_2 | B7L74_02785 / B7L74_02805 | hypothetical protein/MFS transpo | intergenic (-234/-387) | | | | | | | | | | | | | Transient |
| LOBOINFK_01128 ← | B7L74_02145 | tRNA-Lys | noncoding (70/78 nt) | | | | | | | | | | | | | Transient |
| – / → LOBOINFK_00001 | / B7L74_05465 | –/segregation/condensation prote | intergenic (–/-89) | | | | | | | | | | | | | Transient |
| LOBOINFK_01085 ← / → LOBOINFK_01086 | B7L74_01840 / B7L74_01845 | protease SohB/amino acid perme | intergenic (-152/-78) | | | | | | | | | | | | | Transient |
| LOBOINFK_01084 ← / ← LOBOINFK_01085 | B7L74_01835 / B7L74_01840 | cation transporter/protease SohB | intergenic (-136/+1) | | | | | | | | | | | | | Transient |
| LOBOINFK_01031 → / ← LOBOINFK_01032 | B7L74_02240 / B7L74_02220 | hypothetical protein/mechanosen | intergenic (+604/+227) | | | | | | | | | | | | | Transient |
| LOBOINFK_01031 → / ← LOBOINFK_01032 | B7L74_02240 / B7L74_02220 | hypothetical protein/mechanosen | intergenic (+600/+231) | | | | | | | | | | | | | Transient |
| LOBOINFK_01031 → / ← LOBOINFK_01032 | B7L74_02240 / B7L74_02220 | hypothetical protein/mechanosen | intergenic (+576/+255) | | | | | | | | | | | | | Transient |
| LOBOINFK_00871 → / → mvl | B7L74_05900 / B7L74_05895 | hypothetical protein/Lectin MVL | intergenic (+68/-160) | | | | | | | | | | | | | Transient |
| LOBOINFK_00867 ← / ← LOBOINFK_00868 | B7L74_05915 / B7L74_05910 | hypothetical protein/transcription | intergenic (-114/+30) | | | | | | | | | | | | | Transient |
| LOBOINFK_00839 ← / ← phrA | B7L74_06060 / B7L74_06050 | hypothetical protein/Deoxyribod | intergenic (-351/+413) | | | | | | | | | | | | | Transient |
| LOBOINFK_00772 → / ← LOBOINFK_00773 | B7L74_10455 / B7L74_10470 | ubiquinone biosynthesis regulato | intergenic (+140/+265) | | | | | | | | | | | | | Transient |
| LOBOINFK_00754 → / ← LOBOINFK_00755 | B7L74_10360 / B7L74_10380 | threonylcarbamoyl-AMP synthas | intergenic (+277/+384) | | | | | | | | | | | | | Transient |
| LOBOINFK_00754 → / ← LOBOINFK_00755 | B7L74_10360 / B7L74_10380 | threonylcarbamoyl-AMP synthas | intergenic (+276/+385) | | | | | | | | | | | | | Transient |
| – / → LOBOINFK_02021 | / B7L74_10120 | –/NAD(P) transhydrogenase subu | intergenic (–/-200) | | | | | | | | | | | | | Transient |
| yjdF → / ← LOBOINFK_00676 | B7L74_09540 / B7L74_09530 | putative protein YjdF/hypothetica | intergenic (+414/+142) | | | | | | | | | | | | | Transient |
| – / → LOBOINFK_00660 | / B7L74_09615 | –/DNA topoisomerase IV subunit | intergenic (–/-366) | | | | | | | | | | | | | Transient |
| LOBOINFK_01817 → / – | B7L74_03805 / | alpha/beta hydrolase/– | intergenic (+75/–) | | | | | | | | | | | | | Transient |
| LOBOINFK_01785 ← / → LOBOINFK_01786 | B7L74_10840 / B7L74_10845 | phosphoenolpyruvate carboxykin | intergenic (-99/-77) | | | | | | | | | | | | | Transient |
| fadE_1 → / → LOBOINFK_00573 | B7L74_02980 / B7L74_02994 | acyl-CoA dehydrogenase/acetyl-C | intergenic (+187/-358) | | | | | | | | | | | | | Transient |
| LOBOINFK_01761 ← / ← LOBOINFK_01762 | B7L74_08670 / B7L74_08675 | hypothetical protein/dCTP deami | intergenic (-30/+124) | | | | | | | | | | | | | Transient |
| LOBOINFK_01684 ← / → LOBOINFK_01685 | B7L74_09155 / B7L74_09150 | hypothetical protein/fructose-1,6 | intergenic (-153/-138) | | | | | | | | | | | | | Transient |
| LOBOINFK_01608 → / → LOBOINFK_01609 | B7L74_00180 / B7L74_00165 | 4'-phosphopantetheinyl transfera | intergenic (+228/-375) | | | | | | | | | | | | | Transient |
| LOBOINFK_01608 → / → LOBOINFK_01609 | B7L74_00180 / B7L74_00165 | 4'-phosphopantetheinyl transfera | intergenic (+223/-380) | | | | | | | | | | | | | Transient |
| LOBOINFK_01599 → / ← LOBOINFK_01600 | B7L74_06525 / B7L74_06530 | protein ScvA/pyridine nucleotide | intergenic (+39/+17) | | | | | | | | | | | | | Transient |
| LOBOINFK_01585 → / ← LOBOINFK_01586 | B7L74_06450 / B7L74_06460 | hypothetical protein/polyprenyl s | intergenic (+263/+80) | | | | | | | | | | | | | Transient |
| LOBOINFK_01585 → / ← LOBOINFK_01586 | B7L74_06450 / B7L74_06460 | hypothetical protein/polyprenyl s | intergenic (+249/+94) | | | | | | | | | | | | | Transient |
| LOBOINFK_00467 ← / ← LOBOINFK_00468 | B7L74_06830 / B7L74_06845 | 50S ribosomal protein L20/50S r | intergenic (-450/+190) | | | | | | | | | | | | | Transient |
| LOBOINFK_01491 → / → carB | B7L74_06615 / B7L74_06610 | carbamoyl phosphate synthase sm | intergenic (+49/-33) | | | | | | | | | | | | | Transient |
| LOBOINFK_01485 ← / ← LOBOINFK_01486 | B7L74_06635 / B7L74_06630 | ATPase/MFS transporter | intergenic (-54/+17) | | | | | | | | | | | | | Transient |
| – / ← LOBOINFK_01474 | / B7L74_06685 | –/DNA repair protein RecN | intergenic (–/+100) | | | | | | | | | | | | | Transient |
| LOBOINFK_01359 ← / ← LOBOINFK_01360 | B7L74_01415 / B7L74_01420 | single-stranded DNA-binding pro | intergenic (-44/+6) | | | | | | | | | | | | | Transient |
| LOBOINFK_01313 ← / → tppB_2 | B7L74_02785 / B7L74_02805 | hypothetical protein/MFS transpo | intergenic (-245/-376) | | | | | | | | | | | | | Transient |
| LOBOINFK_01313 ← / → tppB_2 | B7L74_02785 / B7L74_02805 | hypothetical protein/MFS transpo | intergenic (-234/-387) | | | | | | | | | | | | | Transient |
| LOBOINFK_01313 ← / → tppB_2 | B7L74_02785 / B7L74_02805 | hypothetical protein/MFS transpo | intergenic (-217/-404) | | | | | | | | | | | | | Transient |
| LOBOINFK_01212 ← / ← LOBOINFK_01213 | B7L74_09790 / B7L74_09805 | methyltransferase/peptidase M16 | intergenic (-213/+283) | | | | | | | | | | | | | Transient |
| LOBOINFK_00379 ← / → LOBOINFK_00380 | B7L74_00565 / B7L74_00550 | methionine import ATP-binding | intergenic (-111/-418) | | | | | | | | | | | | | Transient |
| LOBOINFK_00347 ← / ← LOBOINFK_00348 | B7L74_00740 / B7L74_00735 | hypothetical protein/UDP-3-O-[3 | intergenic (-466/+104) | | | | | | | | | | | | | Transient |
| LOBOINFK_00291 → / ← LOBOINFK_00292 | B7L74_01025 / B7L74_01020 | hypothetical protein/hypothetical | intergenic (+30/+40) | | | | | | | | | | | | | Transient |
| LOBOINFK_00015 → / ← LOBOINFK_00016 | B7L74_05395 / B7L74_05385 | hypothetical protein/hypothetical | intergenic (+178/+6) | | | | | | | | | | | | | Transient |
| LOBOINFK_01031 → / ← LOBOINFK_01032 | B7L74_02240 / B7L74_02220 | hypothetical protein/mechanosen | intergenic (+602/+229) | | | | | | | | | | | | | Transient |
| fabG ← / ← LOBOINFK_00978 | B7L74_02545 / B7L74_02541 | beta-ketoacyl-ACP reductase/mal | intergenic (-42/+34) | | | | | | | | | | | | | Transient |
| LOBOINFK_00754 → / ← LOBOINFK_00755 | B7L74_10360 / B7L74_10380 | threonylcarbamoyl-AMP synthas | intergenic (+300/+361) | | | | | | | | | | | | | Transient |
| LOBOINFK_00754 → / ← LOBOINFK_00755 | B7L74_10360 / B7L74_10380 | threonylcarbamoyl-AMP synthas | intergenic (+298/+363) | | | | | | | | | | | | | Transient |
| LOBOINFK_00754 → / ← LOBOINFK_00755 | B7L74_10360 / B7L74_10380 | threonylcarbamoyl-AMP synthas | intergenic (+265/+396) | | | | | | | | | | | | | Transient |
| – / → LOBOINFK_02021 | / B7L74_10120 | –/NAD(P) transhydrogenase subu | intergenic (–/-231) | | | | | | | | | | | | | Transient |
| – / → LOBOINFK_01957 | / B7L74_07945 | –/hypothetical protein | intergenic (–/-153) | | | | | | | | | | | | | Transient |
| – / → LOBOINFK_01936 | / B7L74_08875 | –/hypothetical protein | intergenic (–/-415) | | | | | | | | | | | | | Transient |
| LOBOINFK_00636 ← / ← LOBOINFK_00637 | B7L74_03295 / B7L74_03300 | alpha-ketoacid dehydrogenase su | intergenic (-65/+140) | | | | | | | | | | | | | Transient |
| LOBOINFK_00581 → / → LOBOINFK_00582 | B7L74_03030 / B7L74_03035 | monothiol glutaredoxin, Grx4 fa | intergenic (+83/-114) | | | | | | | | | | | | | Transient |
| LOBOINFK_00581 → / → LOBOINFK_00582 | B7L74_03030 / B7L74_03035 | monothiol glutaredoxin, Grx4 fa | intergenic (+80/-117) | | | | | | | | | | | | | Transient |
| fadE_1 → / → LOBOINFK_00573 | B7L74_02980 / B7L74_02993 | acyl-CoA dehydrogenase/acetyl-C | intergenic (+197/-348) | | | | | | | | | | | | | Transient |
| LOBOINFK_01763 ← / ← LOBOINFK_01764 | B7L74_08680 / B7L74_08690 | sodium:proton antiporter/transcri | intergenic (-106/+328) | | | | | | | | | | | | | Transient |
| LOBOINFK_01709 → / ← LOBOINFK_01710 | B7L74_02625 / B7L74_02640 | hypothetical protein/MFS transpo | intergenic (+160/+239) | | | | | | | | | | | | | Transient |
| LOBOINFK_01709 → / ← LOBOINFK_01710 | B7L74_02625 / B7L74_02640 | hypothetical protein/MFS transpo | intergenic (+155/+244) | | | | | | | | | | | | | Transient |
| LOBOINFK_01701 → / ← LOBOINFK_01702 | B7L74_02580 / B7L74_02590 | TatD family deoxyribonuclease/g | intergenic (+80/+105) | | | | | | | | | | | | | Transient |
| LOBOINFK_01701 → / ← LOBOINFK_01702 | B7L74_02580 / B7L74_02590 | TatD family deoxyribonuclease/g | intergenic (+72/+113) | | | | | | | | | | | | | Transient |
| LOBOINFK_01636 ← / ← LOBOINFK_01637 | B7L74_06285 / B7L74_06295 | 4-hydroxy-tetrahydrodipicolinate | intergenic (-158/+79) | | | | | | | | | | | | | Transient |
| – / ← LOBOINFK_01633 | / B7L74_06270 | –/hypothetical protein | intergenic (–/+46) | | | | | | | | | | | | | Transient |
| LOBOINFK_01608 → / → LOBOINFK_01609 | B7L74_00180 / B7L74_00165 | 4'-phosphopantetheinyl transfera | intergenic (+220/-383) | | | | | | | | | | | | | Transient |
| LOBOINFK_01602 → / → LOBOINFK_01603 | B7L74_00215 / B7L74_00210 | oligopeptidase A/ketoacyl-ACP sy | intergenic (+21/-60) | | | | | | | | | | | | | Transient |
| LOBOINFK_01585 → / ← LOBOINFK_01586 | B7L74_06450 / B7L74_06460 | hypothetical protein/polyprenyl s | intergenic (+240/+103) | | | | | | | | | | | | | Transient |
| LOBOINFK_01585 → / ← LOBOINFK_01586 | B7L74_06450 / B7L74_06460 | hypothetical protein/polyprenyl s | intergenic (+221/+122) | | | | | | | | | | | | | Transient |
| LOBOINFK_01544 → / ← LOBOINFK_01545 | B7L74_03585 / B7L74_03575 | hypothetical protein/transposase | intergenic (+81/+61) | | | | | | | | | | | | | Transient |
| LOBOINFK_00467 ← / ← LOBOINFK_00468 | B7L74_06830 / B7L74_06845 | 50S ribosomal protein L20/50S r | intergenic (-448/+192) | | | | | | | | | | | | | Transient |
| LOBOINFK_00467 ← / ← LOBOINFK_00468 | B7L74_06830 / B7L74_06845 | 50S ribosomal protein L20/50S r | intergenic (-447/+193) | | | | | | | | | | | | | Transient |
| LOBOINFK_00467 ← / ← LOBOINFK_00468 | B7L74_06830 / B7L74_06845 | 50S ribosomal protein L20/50S r | intergenic (-246/+394) | | | | | | | | | | | | | Transient |
| LOBOINFK_00467 ← / ← LOBOINFK_00468 | B7L74_06830 / B7L74_06845 | 50S ribosomal protein L20/50S r | intergenic (-234/+406) | | | | | | | | | | | | | Transient |
| LOBOINFK_00467 ← / ← LOBOINFK_00468 | B7L74_06830 / B7L74_06845 | 50S ribosomal protein L20/50S r | intergenic (-211/+429) | | | | | | | | | | | | | Transient |
| – / ← LOBOINFK_00443 | / B7L74_06690 | –/hypothetical protein | intergenic (–/+113) | | | | | | | | | | | | | Transient |
| – / ← LOBOINFK_00443 | / B7L74_06690 | –/hypothetical protein | intergenic (–/+120) | | | | | | | | | | | | | Transient |
| LOBOINFK_01428 ← / → LOBOINFK_01429 | B7L74_06200 / B7L74_06210 | delta(24(24(1)))-sterol reductase/ | intergenic (-251/-206) | | | | | | | | | | | | | Transient |
| LOBOINFK_01313 ← / → tppB_2 | B7L74_02785 / B7L74_02805 | hypothetical protein/MFS transpo | intergenic (-272/-349) | | | | | | | | | | | | | Transient |
| LOBOINFK_01313 ← / → tppB_2 | B7L74_02785 / B7L74_02805 | hypothetical protein/MFS transpo | intergenic (-242/-379) | | | | | | | | | | | | | Transient |
| LOBOINFK_01309 → / → LOBOINFK_01310 | B7L74_02765 / B7L74_02770 | competence protein ComEA/unde | intergenic (+92/-120) | | | | | | | | | | | | | Transient |
| LOBOINFK_01268 → / ← LOBOINFK_01269 | B7L74_10870 / B7L74_10865 | hypothetical protein/mobile myst | intergenic (+56/+205) | | | | | | | | | | | | | Transient |
| LOBOINFK_01240 → / ← LOBOINFK_01241 | B7L74_09975 / B7L74_09995 | ATP-dependent DNA ligase/hypo | intergenic (+147/+207) | | | | | | | | | | | | | Transient |
| LOBOINFK_01212 ← / ← LOBOINFK_01213 | B7L74_09790 / B7L74_09805 | methyltransferase/peptidase M16 | intergenic (-240/+256) | | | | | | | | | | | | | Transient |
| LOBOINFK_01212 ← / ← LOBOINFK_01213 | B7L74_09790 / B7L74_09805 | methyltransferase/peptidase M16 | intergenic (-210/+286) | | | | | | | | | | | | | Transient |
| LOBOINFK_01139 ← / ← LOBOINFK_01140 | B7L74_07385 / B7L74_07390 | tRNA-Met/NADH-quinone oxido | intergenic (-156/+85) | | | | | | | | | | | | | Transient |
| LOBOINFK_01127 ← / ← LOBOINFK_01128 | B7L74_07315 /B7L74_02145 | DNA repair protein RadA/tRNA- | intergenic (-120/+115) | | | | | | | | | | | | | Transient |
| LOBOINFK_01114 → / → LOBOINFK_01115 | B7L74_07255 / B7L74_07260 | hypothetical protein/type II citrat | intergenic (+82/-152) | | | | | | | | | | | | | Transient |
| LOBOINFK_00438 → / ← LOBOINFK_00439 | B7L74_00250 / B7L74_00245 | lipoprotein/hypothetical protein | intergenic (+15/+82) | | | | | | | | | | | | | Transient |
| LOBOINFK_00347 ← / ← LOBOINFK_00348 | B7L74_00740 / B7L74_00735 | hypothetical protein/UDP-3-O-[3 | intergenic (-438/+132) | | | | | | | | | | | | | Transient |
| LOBOINFK_00347 ← / ← LOBOINFK_00348 | B7L74_00740 / B7L74_00735 | hypothetical protein/UDP-3-O-[3 | intergenic (-410/+160) | | | | | | | | | | | | | Transient |
| LOBOINFK_00342 → / ← LOBOINFK_00343 | B7L74_00775 / B7L74_00760 | cell division protein ZapD/7,8-di | intergenic (+249/+119) | | | | | | | | | | | | | Transient |
| LOBOINFK_00305 ← / ← LOBOINFK_00306 | B7L74_00960 / B7L74_00956 | 23S ribosomal RNA/tRNA-Ala | intergenic (-134/+65) | | | | | | | | | | | | | Transient |
| LOBOINFK_00270 ← / ← LOBOINFK_00271 | _01125 / B7L74_01110+B7L74 | cytochrome d terminal oxidase su | intergenic (-331/+63) | | | | | | | | | | | | | Transient |
| LOBOINFK_00225 → / ← LOBOINFK_00226 | B7L74_04345 / B7L74_04340 | methylglyoxal synthase/polyribo | intergenic (+16/+76) | | | | | | | | | | | | | Transient |
| LOBOINFK_00193 ← / → LOBOINFK_00194 | B7L74_04515 / B7L74_04510 | translational GTPase TypA/hypot | intergenic (-250/-19) | | | | | | | | | | | | | Transient |
| LOBOINFK_00056 → / ← LOBOINFK_00057 | B7L74_05190 / B7L74_05185 | beta-carotene 15,15'-monooxyge | intergenic (+32/+19) | | | | | | | | | | | | | Transient |
| LOBOINFK_01088 → / ← LOBOINFK_01089 | B7L74_01850 / B7L74_01860 | hypothetical protein/hypothetical | intergenic (+668/+47) | | | | | | | | | | | | | Transient |
| LOBOINFK_01088 → / ← LOBOINFK_01089 | B7L74_01850 / B7L74_01860 | hypothetical protein/hypothetical | intergenic (+654/+61) | | | | | | | | | | | | | Transient |
| LOBOINFK_01088 → / ← LOBOINFK_01089 | B7L74_01850 / B7L74_01860 | hypothetical protein/hypothetical | intergenic (+56/+659) | | | | | | | | | | | | | Transient |
| LOBOINFK_01088 → / ← LOBOINFK_01089 | B7L74_01850 / B7L74_01860 | hypothetical protein/hypothetical | intergenic (+53/+662) | | | | | | | | | | | | | Transient |
| LOBOINFK_00883 ← / → LOBOINFK_00884 | B7L74_05830 / B7L74_05820 | inositol monophosphatase/tRNA | intergenic (-146/-128) | | | | | | | | | | | | | Transient |
| LOBOINFK_00883 ← / → LOBOINFK_00884 | B7L74_05830 / B7L74_05820 | inositol monophosphatase/tRNA | intergenic (-139/-135) | | | | | | | | | | | | | Transient |
| LOBOINFK_01935 ← / – | B7L74_05685 / | hypothetical protein/– | intergenic (-116/–) | | | | | | | | | | | | | Transient |
| fadE_1 → / → LOBOINFK_00573 | B7L74_02980 / B7L74_02992 | acyl-CoA dehydrogenase/acetyl-C | intergenic (+207/-338) | | | | | | | | | | | | | Transient |
| LOBOINFK_01681 → / → LOBOINFK_01682 | B7L74_09170 / B7L74_09165 | type I glyceraldehyde-3-phospha | intergenic (+21/-66) | | | | | | | | | | | | | Transient |
| LOBOINFK_01585 → / ← LOBOINFK_01586 | B7L74_06450 / B7L74_06460 | hypothetical protein/polyprenyl s | intergenic (+262/+81) | | | | | | | | | | | | | Transient |
| LOBOINFK_01585 → / ← LOBOINFK_01586 | B7L74_06450 / B7L74_06460 | hypothetical protein/polyprenyl s | intergenic (+236/+107) | | | | | | | | | | | | | Transient |
| LOBOINFK_00492 ← / → LOBOINFK_00493 | B7L74_06950 / B7L74_06960 | amino acid permease/hypothetica | intergenic (-352/-156) | | | | | | | | | | | | | Transient |
| LOBOINFK_01491 → / → carB | B7L74_06615 / B7L74_06610 | carbamoyl phosphate synthase sm | intergenic (+44/-38) | | | | | | | | | | | | | Transient |
| LOBOINFK_01491 → / → carB | B7L74_06615 / B7L74_06610 | carbamoyl phosphate synthase sm | intergenic (+41/-41) | | | | | | | | | | | | | Transient |
| – / → LOBOINFK_01440 | / B7L74_08410 | –/hypothetical protein | intergenic (–/-216) | | | | | | | | | | | | | Transient |
| LOBOINFK_01407 → / → LOBOINFK_01408 | B7L74_07540 / B7L74_07530 | tRNA-Asp/hypothetical protein | intergenic (+41/-252) | | | | | | | | | | | | | Transient |
| LOBOINFK_01405 → / → LOBOINFK_01406 | B7L74_07555 / B7L74_07545 | DNA-binding protein HU/tRNA- | intergenic (+136/-102) | | | | | | | | | | | | | Transient |
| LOBOINFK_01393 → / ← gatB | B7L74_07615 / B7L74_07610 | DNA-binding transcriptional regu | intergenic (+60/+84) | | | | | | | | | | | | | Transient |
| LOBOINFK_01212 ← / ← LOBOINFK_01213 | B7L74_09790 / B7L74_09805 | methyltransferase/peptidase M16 | intergenic (-202/+294) | | | | | | | | | | | | | Transient |
| LOBOINFK_00347 ← / ← LOBOINFK_00348 | B7L74_00740 / B7L74_00735 | hypothetical protein/UDP-3-O-[3 | intergenic (-453/+117) | | | | | | | | | | | | | Transient |
| LOBOINFK_00342 → / ← LOBOINFK_00343 | B7L74_00775 / B7L74_00760 | cell division protein ZapD/7,8-di | intergenic (+264/+104) | | | | | | | | | | | | | Transient |
| coaE → / ← LOBOINFK_01110 | B7L74_01960 / B7L74_01965 | Dephospho-CoA kinase/hypothet | intergenic (+38/+11) | | | | | | | | | | | | | Transient |
| LOBOINFK_01031 → / ← LOBOINFK_01032 | B7L74_02240 / B7L74_02220 | hypothetical protein/mechanosen | intergenic (+593/+238) | | | | | | | | | | | | | Transient |
| LOBOINFK_01011 ← / → LOBOINFK_01012 | B7L74_02345 / B7L74_02340 | hypothetical protein/histone-like | intergenic (-108/-183) | | | | | | | | | | | | | Transient |
| LOBOINFK_00986 → / ← LOBOINFK_00987 | B7L74_02500 / B7L74_02495 | ribonuclease E/G/hypothetical pr | intergenic (+40/+20) | | | | | | | | | | | | | Transient |
| LOBOINFK_00867 ← / ← LOBOINFK_00868 | B7L74_05915 / B7L74_05910 | hypothetical protein/transcription | intergenic (-113/+31) | | | | | | | | | | | | | Transient |
| LOBOINFK_01608 → / → LOBOINFK_01609 | B7L74_00180 / B7L74_00165 | 4'-phosphopantetheinyl transfera | intergenic (+309/-294) | | | | | | | | | | | | | Transient |
| LOBOINFK_01599 → / ← LOBOINFK_01600 | B7L74_06525 / B7L74_06530 | protein ScvA/pyridine nucleotide | intergenic (+40/+16) | | | | | | | | | | | | | Transient |
| LOBOINFK_00467 ← / ← LOBOINFK_00468 | B7L74_06830 / B7L74_06845 | 50S ribosomal protein L20/50S r | intergenic (-439/+201) | | | | | | | | | | | | | Transient |
| LOBOINFK_01481 → / → LOBOINFK_01482 | B7L74_06650 / B7L74_06645 | molecular chaperone DnaK/mole | intergenic (+33/-110) | | | | | | | | | | | | | Transient |
| LOBOINFK_01224 ← / → LOBOINFK_01225 | B7L74_09885 / B7L74_09895 | universal stress protein A/ribonuc | intergenic (-10/-650) | | | | | | | | | | | | | Transient |
| LOBOINFK_01133 → / ← LOBOINFK_01134 | B7L74_07350 / B7L74_07360 | hypothetical protein/tRNA pseud | intergenic (+281/+528) | | | | | | | | | | | | | Transient |
| LOBOINFK_00347 ← / ← LOBOINFK_00348 | B7L74_00740 / B7L74_00735 | hypothetical protein/UDP-3-O-[3 | intergenic (-411/+159) | | | | | | | | | | | | | Transient |
| LOBOINFK_00305 ← | B7L74_00960 | 23S ribosomal RNA | oncoding (2694/4342 nt | | | | | | | | | | | | | Transient |
| LOBOINFK_00305 ← | B7L74_00960 | 23S ribosomal RNA | oncoding (2832/4342 nt) | | | | | | | | | | | | | Transient |
| LOBOINFK_00259 ← / → LOBOINFK_00260 | B7L74_04170 / B7L74_04165 | TetR family transcriptional regula | intergenic (-23/-227) | | | | | | | | | | | | | Transient |
| LOBOINFK_01096 ← / → LOBOINFK_01097 | B7L74_01890 / B7L74_01895 | DNA-binding response regulator | intergenic (-27/-25) | | | | | | | | | | | | | Transient |
| LOBOINFK_01096 ← / → LOBOINFK_01097 | B7L74_01890 / B7L74_01895 | DNA-binding response regulator | intergenic (-25/-27) | | | | | | | | | | | | | Transient |
| LOBOINFK_01096 ← / → LOBOINFK_01097 | B7L74_01890 / B7L74_01895 | DNA-binding response regulator | intergenic (-23/-29) | | | | | | | | | | | | | Transient |

| | | | | | | | | | | | | | | | | |
|---|---|---|---|---|---|---|---|---|---|---|---|---|---|---|---|---|
| LOBOINFK_01058 → / ← LOBOINFK_01059 | B7L74_01705 / B7L74_01710 | acyl-CoA synthetase/phosphoribo | intergenic (+116/+146) | | | | | | | | | | | | | Transient |
| LOBOINFK_01058 → / ← LOBOINFK_01059 | B7L74_01705 / B7L74_01710 | acyl-CoA synthetase/phosphoribo | intergenic (+115/+147) | | | | | | | | | | | | | Transient |
| LOBOINFK_01976 → / → LOBOINFK_01977 | B7L74_08095 / B7L74_08100 | protein TolR/hypothetical protein | intergenic (+51/-92) | | | | | | | | | | | | | Transient |
| LOBOINFK_01854 → / → LOBOINFK_01855 | B7L74_09685 / B7L74_09680 | iron-sulfur cluster insertion prote | intergenic (+104/-41) | | | | | | | | | | | | | Transient |
| LOBOINFK_01781 → / ← LOBOINFK_01782 | B7L74_10820 / B7L74_10825 | sodium-dependent transporter/Yg | intergenic (+13/+82) | | | | | | | | | | | | | Transient |
| LOBOINFK_01781 → / ← LOBOINFK_01782 | B7L74_10820 / B7L74_10825 | sodium-dependent transporter/Yg | intergenic (+12/+83) | | | | | | | | | | | | | Transient |
| LOBOINFK_01778 → / ← LOBOINFK_01779 | B7L74_10805 / B7L74_10810 | hypothetical protein/transcription | intergenic (+117/+70) | | | | | | | | | | | | | Transient |
| LOBOINFK_01777 → / → LOBOINFK_01778 | B7L74_10790 / B7L74_10805 | 7-cyano-7-deazaguanine synthase | intergenic (+20/-387) | | | | | | | | | | | | | Transient |
| LOBOINFK_00625 → / → LOBOINFK_00626 | B7L74_03235 / B7L74_03245 | hypothetical protein/inorganic py | intergenic (+189/-194) | | | | | | | | | | | | | Transient |
| LOBOINFK_00625 → / → LOBOINFK_00626 | B7L74_03235 / B7L74_03245 | hypothetical protein/inorganic py | intergenic (+188/-195) | | | | | | | | | | | | | Transient |
| LOBOINFK_00601 → / ← LOBOINFK_00602 | B7L74_03115 / B7L74_03125 | hypothetical protein/hypothetical | intergenic (+292/+655) | | | | | | | | | | | | | Transient |
| LOBOINFK_01638 → / ← LOBOINFK_01639 | B7L74_06300 / B7L74_06305 | DNA recombination protein Rmu | intergenic (+73/+32) | | | | | | | | | | | | | Transient |
| LOBOINFK_01638 → / ← LOBOINFK_01639 | B7L74_06300 / B7L74_06305 | DNA recombination protein Rmu | intergenic (+72/+33) | | | | | | | | | | | | | Transient |
| LOBOINFK_01638 → / ← LOBOINFK_01639 | B7L74_06300 / B7L74_06305 | DNA recombination protein Rmu | intergenic (+71/+34) | | | | | | | | | | | | | Transient |
| LOBOINFK_01632 → / – | B7L74_00040 / | antitoxin/– | intergenic (+99/–) | | | | | | | | | | | | | Transient |
| LOBOINFK_01615 → / → LOBOINFK_01616 | B7L74_00140 / B7L74_00130 | ribose-5-phosphate isomerase/car | intergenic (+8/-357) | | | | | | | | | | | | | Transient |
| LOBOINFK_01615 → / → LOBOINFK_01616 | B7L74_00140 / B7L74_00130 | ribose-5-phosphate isomerase/car | intergenic (+7/-358) | | | | | | | | | | | | | Transient |
| LOBOINFK_01405 → / → LOBOINFK_01406 | B7L74_07555 / B7L74_07545 | DNA-binding protein HU/tRNA- | intergenic (+137/-101) | | | | | | | | | | | | | Transient |
| LOBOINFK_01405 → / → LOBOINFK_01406 | B7L74_07555 / B7L74_07545 | DNA-binding protein HU/tRNA- | intergenic (+135/-103) | | | | | | | | | | | | | Transient |
| LOBOINFK_01162 → / → LOBOINFK_01163 | B7L74_07935 / B7L74_07930 | hypothetical protein/hypothetical | intergenic (+180/-134) | | | | | | | | | | | | | Transient |
| LOBOINFK_01153 ← / ← LOBOINFK_01154 | B7L74_07455 / B7L74_07460 | NADH-quinone oxidoreductase s | intergenic (-111/+95) | | | | | | | | | | | | | Transient |
| LOBOINFK_01153 ← / ← LOBOINFK_01154 | B7L74_07455 / B7L74_07460 | NADH-quinone oxidoreductase s | intergenic (-110/+96) | | | | | | | | | | | | | Transient |
| LOBOINFK_00438 → / ← LOBOINFK_00439 | B7L74_00250 / B7L74_00245 | lipoprotein/hypothetical protein | intergenic (+20/+77) | | | | | | | | | | | | | Transient |
| LOBOINFK_00438 → / ← LOBOINFK_00439 | B7L74_00250 / B7L74_00245 | lipoprotein/hypothetical protein | intergenic (+19/+78) | | | | | | | | | | | | | Transient |
| LOBOINFK_00362 ← / ← LOBOINFK_00363 | B7L74_00665 / B7L74_00655 | hypothetical protein/phospho-N- | intergenic (-303/+71) | | | | | | | | | | | | | Transient |
| LOBOINFK_00362 ← / ← LOBOINFK_00363 | B7L74_00665 / B7L74_00655 | hypothetical protein/phospho-N- | intergenic (-302/+72) | | | | | | | | | | | | | Transient |
| LOBOINFK_01050 → / → LOBOINFK_01051 | B7L74_01655 / B7L74_01665 | ribonuclease HI/DNA polymerase | intergenic (+105/-325) | | | | | | | | | | | | | Transient |
| LOBOINFK_01031 → / ← LOBOINFK_01032 | B7L74_02240 / B7L74_02220 | hypothetical protein/mechanosen | intergenic (+648/+183) | | | | | | | | | | | | | Transient |
| fabG ← / ← LOBOINFK_00978 | B7L74_02545 / B7L74_02540 | beta-ketoacyl-ACP reductase/mal | intergenic (-43/+33) | | | | | | | | | | | | | Transient |
| LOBOINFK_00754 → / ← LOBOINFK_00755 | B7L74_10360 / B7L74_10380 | threonylcarbamoyl-AMP synthas | intergenic (+290/+371) | | | | | | | | | | | | | Transient |
| LOBOINFK_00754 → / ← LOBOINFK_00755 | B7L74_10360 / B7L74_10380 | threonylcarbamoyl-AMP synthas | intergenic (+288/+373) | | | | | | | | | | | | | Transient |
| LOBOINFK_00754 → / ← LOBOINFK_00755 | B7L74_10360 / B7L74_10380 | threonylcarbamoyl-AMP synthas | intergenic (+274/+387) | | | | | | | | | | | | | Transient |
| – / → LOBOINFK_02021 | / B7L74_10122 | –/NAD(P) transhydrogenase subu | intergenic (–/-221) | | | | | | | | | | | | | Transient |
| – / → LOBOINFK_02021 | / B7L74_10121 | –/NAD(P) transhydrogenase subu | intergenic (–/-227) | | | | | | | | | | | | | Transient |
| – / → LOBOINFK_02021 | / B7L74_10120 | –/NAD(P) transhydrogenase subu | intergenic (–/-268) | | | | | | | | | | | | | Transient |
| LOBOINFK_01998 → / – | B7L74_08140 / – | Na+/H+ antiporter subunit G/– | intergenic (+362/–) | | | | | | | | | | | | | Transient |
| LOBOINFK_01998 → / – | B7L74_08140 / – | Na+/H+ antiporter subunit G/– | intergenic (+295/–) | | | | | | | | | | | | | Transient |
| LOBOINFK_01935 ← / – | B7L74_05685 / – | hypothetical protein/– | intergenic (-132/–) | | | | | | | | | | | | | Transient |
| – / → LOBOINFK_01793 | / B7L74_03680 | –/ABC transporter permease | intergenic (–/-79) | | | | | | | | | | | | | Transient |
| LOBOINFK_01778 → / ← LOBOINFK_01779 | B7L74_10805 / B7L74_10810 | hypothetical protein/transcription | intergenic (+113/+74) | | | | | | | | | | | | | Transient |
| LOBOINFK_01777 → / → LOBOINFK_01778 | B7L74_10790 / B7L74_10805 | 7-cyano-7-deazaguanine synthase | intergenic (+16/-391) | | | | | | | | | | | | | Transient |
| LOBOINFK_00601 → / ← LOBOINFK_00602 | B7L74_03115 / B7L74_03125 | hypothetical protein/hypothetical | intergenic (+287/+660) | | | | | | | | | | | | | Transient |
| LOBOINFK_00581 → / → LOBOINFK_00582 | B7L74_03030 / B7L74_03035 | monothiol glutaredoxin, Grx4 fa | intergenic (+88/-109) | | | | | | | | | | | | | Transient |
| LOBOINFK_01749 ← / – | B7L74_08595 / | 3-deoxy-8-phosphooctulonate sy | intergenic (-224/–) | | | | | | | | | | | | | Transient |
| – / → LOBOINFK_01680 | / B7L74_09175 | –/transketolase | intergenic (–/-105) | | | | | | | | | | | | | Transient |
| LOBOINFK_01632 → / – | B7L74_00040 / | antitoxin/– | intergenic (+91/–) | | | | | | | | | | | | | Transient |
| LOBOINFK_01602 → / → LOBOINFK_01603 | B7L74_00215 / B7L74_00210 | oligopeptidase A/ketoacyl-ACP s | intergenic (+36/-45) | | | | | | | | | | | | | Transient |
| LOBOINFK_00467 ← / ← LOBOINFK_00468 | B7L74_06830 / B7L74_06845 | 50S ribosomal protein L20/50S r | intergenic (-218/+422) | | | | | | | | | | | | | Transient |
| LOBOINFK_00467 ← / ← LOBOINFK_00468 | B7L74_06830 / B7L74_06845 | 50S ribosomal protein L20/50S r | intergenic (-209/+431) | | | | | | | | | | | | | Transient |
| – / ← LOBOINFK_00443 | / B7L74_06690 | –/hypothetical protein | intergenic (–/+119) | | | | | | | | | | | | | Transient |
| – / ← LOBOINFK_00443 | / B7L74_06690 | –/hypothetical protein | intergenic (–/+124) | | | | | | | | | | | | | Transient |
| LOBOINFK_01428 ← / → LOBOINFK_01429 | B7L74_06200 / B7L74_06210 | delta(24(24(1)))-sterol reductase/ | intergenic (-218/-239) | | | | | | | | | | | | | Transient |
| LOBOINFK_01309 → / → LOBOINFK_01310 | B7L74_02765 / B7L74_02770 | competence protein ComEA/und | intergenic (+93/-119) | | | | | | | | | | | | | Transient |
| LOBOINFK_01268 → / ← LOBOINFK_01269 | B7L74_10870 / B7L74_10865 | hypothetical protein/mobile myst | intergenic (+59/+202) | | | | | | | | | | | | | Transient |
| LOBOINFK_01212 ← / ← LOBOINFK_01213 | B7L74_09790 / B7L74_09805 | methyltransferase/peptidase M16 | intergenic (-260/+236) | | | | | | | | | | | | | Transient |
| LOBOINFK_01212 ← / ← LOBOINFK_01213 | B7L74_09790 / B7L74_09805 | methyltransferase/peptidase M16 | intergenic (-257/+239) | | | | | | | | | | | | | Transient |
| LOBOINFK_01139 ← / ← LOBOINFK_01140 | B7L74_07385 / B7L74_07390 | tRNA-Met/NADH-quinone oxido | intergenic (-160/+81) | | | | | | | | | | | | | Transient |
| LOBOINFK_01133 → / ← LOBOINFK_01134 | B7L74_07350 / B7L74_07360 | hypothetical protein/tRNA pseud | intergenic (+280/+529) | | | | | | | | | | | | | Transient |
| LOBOINFK_01127 ← / ← LOBOINFK_01128 | B7L74_07315 / B7L74_02145 | DNA repair protein RadA/tRNA- | intergenic (-98/+137) | | | | | | | | | | | | | Transient |
| LOBOINFK_00438 → / ← LOBOINFK_00439 | B7L74_00250 / B7L74_00245 | lipoprotein/hypothetical protein | intergenic (+24/+73) | | | | | | | | | | | | | Transient |
| LOBOINFK_00410 ← / → LOBOINFK_00411 | B7L74_00390 / B7L74_00380 | 2-octaprenyl-3-methyl-6-methox | intergenic (-112/-196) | | | | | | | | | | | | | Transient |
| LOBOINFK_00342 → / ← LOBOINFK_00343 | B7L74_00775 / B7L74_00760 | cell division protein ZapD/7,8-di | intergenic (+297/+71) | | | | | | | | | | | | | Transient |
| LOBOINFK_00342 → / ← LOBOINFK_00343 | B7L74_00775 / B7L74_00760 | cell division protein ZapD/7,8-di | intergenic (+230/+138) | | | | | | | | | | | | | Transient |
| LOBOINFK_00305 ← / ← LOBOINFK_00306 | B7L74_00960 / B7L74_00955 | 23S ribosomal RNA/tRNA-Ala | intergenic (-143/+56) | | | | | | | | | | | | | Transient |
| LOBOINFK_00225 → / ← LOBOINFK_00226 | B7L74_04345 / B7L74_04340 | methylglyoxal synthase/polyribo | intergenic (+84/+8) | | | | | | | | | | | | | Transient |
| LOBOINFK_00225 → / ← LOBOINFK_00226 | B7L74_04345 / B7L74_04340 | methylglyoxal synthase/polyribo | intergenic (+55/+37) | | | | | | | | | | | | | Transient |
| LOBOINFK_00225 → / ← LOBOINFK_00226 | B7L74_04345 / B7L74_04340 | methylglyoxal synthase/polyribo | intergenic (+54/+38) | | | | | | | | | | | | | Transient |
| LOBOINFK_00193 ← / → LOBOINFK_00194 | B7L74_04515 / B7L74_04510 | translational GTPase TypA/hypot | intergenic (-247/-22) | | | | | | | | | | | | | Transient |
| coaE → / ← LOBOINFK_01110 | B7L74_01960 / B7L74_01966 | Dephospho-CoA kinase/hypothet | intergenic (+40/+9) | | | | | | | | | | | | | Transient |
| coaE → / ← LOBOINFK_01110 | B7L74_01960 / B7L74_01965 | Dephospho-CoA kinase/hypothet | intergenic (+25/+24) | | | | | | | | | | | | | Transient |
| LOBOINFK_01058 → / ← LOBOINFK_01059 | B7L74_01705 / B7L74_01710 | acyl-CoA synthetase/phosphoribo | intergenic (+68/+194) | | | | | | | | | | | | | Transient |
| LOBOINFK_01046 ← | B7L74_01495 | tRNA-Glu | noncoding (52/82 nt) | | | | | | | | | | | | | Transient |
| LOBOINFK_00867 ← / ← LOBOINFK_00868 | B7L74_05915 / B7L74_05910 | hypothetical protein/transcription | intergenic (-112/+32) | | | | | | | | | | | | | Transient |
| LOBOINFK_01998 → / – | B7L74_08140 / | Na+/H+ antiporter subunit G/– | intergenic (+95/–) | | | | | | | | | | | | | Transient |
| LOBOINFK_00702 ← / ← LOBOINFK_00703 | B7L74_09385 / B7L74_09375 | hypothetical protein/hypothetical | intergenic (-250/+52) | | | | | | | | | | | | | Transient |
| – / → LOBOINFK_01936 | / B7L74_08875 | –/hypothetical protein | intergenic (–/-462) | | | | | | | | | | | | | Transient |
| fadE_1 → / → LOBOINFK_00573 | B7L74_02980 / B7L74_02991 | acyl-CoA dehydrogenase/acetyl- | intergenic (+222/-323) | | | | | | | | | | | | | Transient |
| fadE_1 → / → LOBOINFK_00573 | B7L74_02980 / B7L74_02990 | acyl-CoA dehydrogenase/acetyl- | intergenic (+188/-357) | | | | | | | | | | | | | Transient |
| LOBOINFK_01761 ← / ← LOBOINFK_01762 | B7L74_08670 / B7L74_08675 | hypothetical protein/dCTP deami | intergenic (-123/+31) | | | | | | | | | | | | | Transient |
| LOBOINFK_01608 → / → LOBOINFK_01609 | B7L74_00180 / B7L74_00165 | 4'-phosphopantetheinyl transfera | intergenic (+221/-382) | | | | | | | | | | | | | Transient |
| – / → LOBOINFK_01505 | / B7L74_08880 | –/acetyl-CoA carboxylase biotin | intergenic (–/-140) | | | | | | | | | | | | | Transient |
| LOBOINFK_00483 ← / ← LOBOINFK_00484 | B7L74_06910 / B7L74_06915 | D-alanine--D-alanine ligase A/rib | intergenic (-105/+45) | | | | | | | | | | | | | Transient |
| LOBOINFK_01467 → / → LOBOINFK_01468 | B7L74_08265 / B7L74_08260 | hypothetical protein/hypothetical | intergenic (+300/-39) | | | | | | | | | | | | | Transient |
| LOBOINFK_01304 → / → LOBOINFK_01305 | B7L74_02740 / B7L74_02745 | cytidylate kinase/30S ribosomal p | intergenic (+68/-76) | | | | | | | | | | | | | Transient |
| LOBOINFK_01128 ← | B7L74_02145 | tRNA-Lys | noncoding (69/78 nt) | | | | | | | | | | | | | Transient |
| LOBOINFK_00410 ← / → LOBOINFK_00411 | B7L74_00390 / B7L74_00380 | 2-octaprenyl-3-methyl-6-methox | intergenic (-105/-203) | | | | | | | | | | | | | Transient |
| LOBOINFK_00305 ← | B7L74_00960 | 23S ribosomal RNA | oncoding (3628/4342 nt) | | | | | | | | | | | | | Transient |
| LOBOINFK_00270 ← / ← LOBOINFK_00271 | 01125 / B7L74_01110+B7L74 | cytochrome d terminal oxidase su | intergenic (-188/+206) | | | | | | | | | | | | | Transient |
| LOBOINFK_00270 ← / ← LOBOINFK_00271 | 01125 / B7L74_01110+B7L74 | cytochrome d terminal oxidase su | intergenic (-159/+235) | | | | | | | | | | | | | Transient |
| LOBOINFK_00270 ← / ← LOBOINFK_00271 | 01125 / B7L74_01110+B7L74 | cytochrome d terminal oxidase su | intergenic (-151/+243) | | | | | | | | | | | | | Transient |
| LOBOINFK_00099 ← / ← LOBOINFK_00100 | B7L74_04955 / B7L74_04940 | cytochrome d terminal oxidase su | intergenic (-135/+335) | | | | | | | | | | | | | Transient |
| LOBOINFK_00059 ← / → LOBOINFK_00060 | B7L74_05170 / B7L74_05160 | adenosylmethionine--8-amino-7- | intergenic (-184/-185) | | | | | | | | | | | | | Transient |

| Gene name from Breseq | Gene name from reference genome | Description/ Annotation | Amino acid and basepair change | P03 | P05 | P10 | P13 | P16 | P21 | P31 | P42 | P51 | P61 | P67 | Verdict from transcriptomics | Classification by their occurences |
|---|---|---|---|---|---|---|---|---|---|---|---|---|---|---|---|---|
| LOBOINFK_00800 ← | B7L74_10610 | 5-methyltetrahydropteroyl | K550* (AAG→TAG) | | | | | | | | | | | | | Maintained in later passages |
| LOBOINFK_01645 → | B7L74_06335 | ATP-dependent helicase | C262* (TGT→TGA) | | | | | | | | | | | | | Transient |
| LOBOINFK_00518 → | B7L74_07080 | tRNA-specific adenosine | L16* (TTA→TAA) | | | | | | | | | | | | | Transient |
| LOBOINFK_01434 → | B7L74_06225 | hypothetical protein | L425* (TTA→TAA) | | | | | | | | | | | | | Transient |
| LOBOINFK_01434 → | B7L74_06225 | hypothetical protein | L424* (TTA→TAA) | | | | | | | | | | | | | Transient |
| LOBOINFK_01362 → | B7L74_01430 | excinuclease ABC subuni | C409* (TGT→TGA) | | | | | | | | | | | | | Transient |
| LOBOINFK_00318 ← | B7L74_00890 | hypothetical protein | R128* (AGA→TGA) | | | | | | | | | | | | | Transient |
| LOBOINFK_01935 ← | B7L74_05685 | hypothetical protein | S77* (TCA→TAA) | | | | | | | | | | | | | Transient |
| LOBOINFK_01714 → | B7L74_02650 | fructose 1,6-bisphosphata | E24* (GAA→TAA) ‡ | | | | | | | | | | | | | Transient |
| LOBOINFK_01164 → | B7L74_07920 | hypothetical protein | G43* (GGA→TGA) | | | | | | | | | | | | | Transient |
| LOBOINFK_00486 ← | B7L74_06925 | GMP synthase (glutamine | Q132* (CAA→TAA) ‡ | | | | | | | | | | | | | Transient |
| LOBOINFK_00251 ← | B7L74_04215 | hypothetical protein | Q127* (CAA→TAA) | | | | | | | | | | | | | Transient |
| lapA → | B7L74_02750 | Lipopolysaccharide assem | Q85* (CAA→TAA) | | | | | | | | | | | | | Transient |
| LOBOINFK_00251 ← | B7L74_04215 | hypothetical protein | E111* (GAG→TAG) | | | | | | | | | | | | | Transient |
| LOBOINFK_01060 → | B7L74_01715 | hypothetical protein | K319* (AAA→TAA) | | | | | | | | | | | | | Transient |
| LOBOINFK_00985 ← | B7L74_02505 | 23S rRNA pseudouridine | L85* (TTG→TAG) ‡ | | | | | | | | | | | | | Transient |
| LOBOINFK_01990 ← | B7L74_08180 | sodium:proton antiporter | Y247* (TAT→TAA) | | | | | | | | | | | | | Transient |
| LOBOINFK_00519 ← | B7L74_07085 | GTP pyrophosphokinase | C658* (TGT→TGA) ‡ | | | | | | | | | | | | | Transient |
| LOBOINFK_01187 ← | B7L74_07805 | hypothetical protein | L16* (TTA→TAA) | | | | | | | | | | | | | Transient |
| LOBOINFK_01187 ← | B7L74_07805 | hypothetical protein | L17* (TTA→TAA) | | | | | | | | | | | | | Transient |
| LOBOINFK_00195 → | B7L74_04505 | helix-turn-helix domain-c | K27* (AAA→TAA) | | | | | | | | | | | | | Transient |
| LOBOINFK_00812 ← | B7L74_10670 | hypothetical protein | Y250* (TAT→TAG) | | | | | | | | | | | | Late up | Transient |
| LOBOINFK_01838 → | B7L74_10265 | hypothetical protein | Y180* (TAT→TAA) | | | | | | | | | | | | Early up | Transient |
| LOBOINFK_00517 → | B7L74_07075 | para-aminobenzoate synth | W202* (TGG→TGA) | | | | | | | | | | | | Early down | Transient |
| LOBOINFK_00517 → | B7L74_07075 | para-aminobenzoate synth | S200* (TCA→TAA) ‡ | | | | | | | | | | | | Early down | Transient |
| LOBOINFK_01916 ← | B7L74_05510 | intracellular signaling prot | L204* (TTA→TAA) ‡ | | | | | | | | | | | | Early up | Transient |
| LOBOINFK_01414 ← | B7L74_06120 | serine--tRNA ligase | L18* (TTA→TAA) | | | | | | | | | | | | Late down | Transient |
| esiB → | B7L74_05845 | Secretory immunoglobuli | K894* (AAA→TAA) | | | | | | | | | | | | Early up | Transient |
| LOBOINFK_01817 → | B7L74_03805 | alpha/beta hydrolase | K171* (AAA→TAA) | | | | | | | | | | | | Continuous up | Transient |
| LOBOINFK_00270 ← | B7L74_01125 | cytochrome d terminal oxi | E515* (GAA→TAA) | | | | | | | | | | | | Early up | Transient |
| LOBOINFK_00573 → | B7L74_02990 | acetyl-CoA acetyltransfera | C412* (TGC→TGA) | | | | | | | | | | | | Early down | Transient |

| |
|---|
| 5-10 |
| 10-30 |
| 30-50 |
| 50-70 |
| 70-90 |
| 90-99 |
| 100 |

| Gene name from Breseq | Gene name from reference genome | Description/Annotation | Amino acid and basepair change | P03 | P05 | P10 | P13 | P16 | P21 | P31 | P42 | P51 | P61 | P67 | Verdict from transcriptomics | Classification by their occurences |
|---|---|---|---|---|---|---|---|---|---|---|---|---|---|---|---|---|
| LOBOINFK_01090 ← | B7L74_01865 | hypothetical protein | R92R (CGA→CGC) ‡ | | | | | | | | | | | | | Transient |
| LOBOINFK_01050 → | B7L74_01655 | ribonuclease HI | V121V (GTT→GTG) | | | | | | | | | | | | | Transient |
| LOBOINFK_01013 ← | B7L74_02335 | thiol reductase thioredoxin | G89G (GGA→GGC) | | | | | | | | | | | | | Present in all passages |
| LOBOINFK_00952 ← | B7L74_04040 | hypothetical protein | A20A (GCA→GCC) | | | | | | | | | | | | | Maintained in later passages |
| LOBOINFK_00830 → | B7L74_10760 | uroporphyrinogen-III synthase | A115A (GCT→GCG) | | | | | | | | | | | | | Transient |
| LOBOINFK_00807 → | B7L74_10645 | zinc transporter | A225A (GCT→GCG) | | | | | | | | | | | | | Transient |
| LOBOINFK_00590 → | B7L74_03075 | hypothetical protein | V18V (GTT→GTG) | | | | | | | | | | | | | Transient |
| LOBOINFK_01656 → | B7L74_02020 | hypothetical protein | R1110R (CGT→CGG) | | | | | | | | | | | | | Transient |
| LOBOINFK_01645 → | B7L74_06335 | ATP-dependent helicase | A261A (GCT→GCG) | | | | | | | | | | | | | Transient |
| LOBOINFK_01610 → | B7L74_00160 | hydroxymethylglutaryl-CoA r | A297A (GCT→GCG) | | | | | | | | | | | | | Transient |
| LOBOINFK_00490 → | B7L74_06940 | MFS transporter | A83A (GCT→GCG) | | | | | | | | | | | | | Transient |
| LOBOINFK_01453 → | B7L74_08340 | type IV secretion protein IcmC | A117A (GCT→GCG) | | | | | | | | | | | | | Transient |
| LOBOINFK_01344 → | B7L74_01325 | 50S ribosomal protein L18 | R12R (CGT→CGG) | | | | | | | | | | | | | Transient |
| ligA → | B7L74_02820 | DNA ligase (NAD(+)) LigA | R513R (CGT→CGG) | | | | | | | | | | | | | Transient |
| damX → | B7L74_00270 | Cell division protein DamX | G222G (GGT→GGG) | | | | | | | | | | | | | Transient |
| LOBOINFK_01034 → | B7L74_02210 | glycine cleavage system protei | F57F (TTT→TTC) | | | | | | | | | | | | | Transient |
| LOBOINFK_01849 ← | B7L74_09710 | queuosine biosynthesis protein | R112R (CGA→CGG) | | | | | | | | | | | | | Maintained in later passages |
| LOBOINFK_01606 → | B7L74_00190 | beta-ketoacyl-ACP synthase | A190A (GCT→GCC) ‡ | | | | | | | | | | | | | Transient |
| LOBOINFK_00518 → | B7L74_07080 | tRNA-specific adenosine deam | N19N (AAT→AAC) | | | | | | | | | | | | | Transient |
| LOBOINFK_00517 → | B7L74_07075 | para-aminobenzoate synthetase | Y197Y (TAT→TAC) | | | | | | | | | | | | | Transient |
| LOBOINFK_01310 → | B7L74_02770 | undecaprenyl-phosphate alpha- | F103F (TTT→TTC) ‡ | | | | | | | | | | | | | Transient |
| LOBOINFK_01288 → | B7L74_10985 | hypothetical protein | D112D (GAT→GAC) | | | | | | | | | | | | | Transient |
| traI → | B7L74_11025 | Multifunctional conjugation pr | L483L (TTG→CTG) | | | | | | | | | | | | | Transient |
| iucA → | B7L74_02160 | N(2)-citryl-N(6)-acetyl-N(6)-h | S330S (TCT→TCA) | | | | | | | | | | | | | Transient |
| LOBOINFK_00800 ← | B7L74_10610 | 5-methyltetrahydropteroyltrigl | G558G (GGA→GGT) | | | | | | | | | | | | | Transient |
| LOBOINFK_02011 ← | B7L74_10090 | ABC transporter ATP-binding | L395L (CTA→CTT) | | | | | | | | | | | | | Transient |
| LOBOINFK_00717 → | B7L74_09300 | ABC transporter substrate-bin | I88I (ATT→ATA) | | | | | | | | | | | | | Transient |
| LOBOINFK_00627 → | B7L74_03250 | bifunctional proline dehydroge | G523G (GGT→GGA) | | | | | | | | | | | | | Transient |
| LOBOINFK_00627 → | B7L74_03250 | bifunctional proline dehydroge | I518I (ATT→ATA) | | | | | | | | | | | | | Transient |
| LOBOINFK_01553 → | B7L74_03525 | adenylyltransferase | S447S (TCT→TCA) | | | | | | | | | | | | | Transient |
| LOBOINFK_00542 → | B7L74_07190 | L,D-transpeptidase | R101R (CGT→CGA) | | | | | | | | | | | | | Transient |
| LOBOINFK_01115 → | B7L74_07260 | type II citrate synthase | L135L (CTT→CTA) | | | | | | | | | | | | | Transient |
| LOBOINFK_00017 ← | B7L74_05385 | hypothetical protein | G269G (GGA→GGT) | | | | | | | | | | | | | Maintained in later passages |
| LOBOINFK_00884 → | B7L74_05820 | tRNA (cytosine(32)/uridine(32 | T149T (ACG→ACT) | | | | | | | | | | | | | Transient |
| LOBOINFK_00879 → | B7L74_05850 | enhanced entry protein | A69A (GCG→GCT) | | | | | | | | | | | | | Transient |
| LOBOINFK_00812 ← | B7L74_10670 | hypothetical protein | A249A (GCC→GCA) | | | | | | | | | | | | | Transient |
| LOBOINFK_01952 → | B7L74_08775 | alkyl hydroperoxide reductase | V142V (GTG→GTT) | | | | | | | | | | | | | Transient |
| LOBOINFK_01431 ← | B7L74_06220 | hypothetical protein | A235A (GCC→GCA) | | | | | | | | | | | | | Transient |
| LOBOINFK_01360 ← | B7L74_01420 | MFS transporter | R445R (CGA→AGA) | | | | | | | | | | | | | Transient |
| LOBOINFK_00344 ← | B7L74_00755 | preprotein translocase subunit | R80R (CGC→CGA) | | | | | | | | | | | | | Transient |
| LOBOINFK_00131 ← | B7L74_04805 | Fe(2+)-trafficking protein | A41A (GCC→GCA) | | | | | | | | | | | | | Transient |
| LOBOINFK_00547 ← | B7L74_07215 | 2-oxoglutarate dehydrogenase | T815T (ACC→ACG) | | | | | | | | | | | | | Transient |
| LOBOINFK_00547 ← | B7L74_07215 | 2-oxoglutarate dehydrogenase | R816R (CGC→CGG) | | | | | | | | | | | | | Transient |
| LOBOINFK_01363 → | B7L74_01440 | uroporphyrinogen decarboxyla | A348A (GCG→GCC) | | | | | | | | | | | | | Transient |
| LOBOINFK_01247 → | B7L74_10020 | ATP synthase subunit A | R61R (CGG→CGC) | | | | | | | | | | | | | Transient |
| LOBOINFK_00942 → | B7L74_03975 | hypothetical protein | A391A (GCG→GCA) | | | | | | | | | | | | | Transient |
| LOBOINFK_01849 ← | B7L74_09710 | queuosine biosynthesis protein | S111S (TCC→TCT) | | | | | | | | | | | | | Maintained in later passages |
| LOBOINFK_01633 ← | B7L74_06270 | hypothetical protein | N361N (AAC→AAT) | | | | | | | | | | | | | Present in all passages |
| LOBOINFK_01549 ← | B7L74_03545 | ABC transporter ATP-binding | L195L (CTA→TTA) | | | | | | | | | | | | | Present in all passages |
| LOBOINFK_01437 ← | B7L74_06240 | hypothetical protein | H177H (CAC→CAT) | | | | | | | | | | | | | Transient |
| LOBOINFK_01372 → | B7L74_01505 | 2-amino-4-hydroxy-6- hydrox | L71L (TTG→TTA) ‡ | | | | | | | | | | | | | Transient |
| LOBOINFK_00970 → | B7L74_04140 | MATE family efflux transporte | R376R (CGC→CGT) | | | | | | | | | | | | | Transient |
| LOBOINFK_02034 ← | B7L74_10150 | IS110 family transposase | K34K (AAG→AAA) | | | | | | | | | | | | | Present in all passages |
| LOBOINFK_01656 → | B7L74_02020 | hypothetical protein | L606L (CTA→TTA) | | | | | | | | | | | | | Transient |
| LOBOINFK_01363 → | B7L74_01440 | uroporphyrinogen decarboxyla | A349A (GCC→GCT) | | | | | | | | | | | | | Transient |
| LOBOINFK_01250 → | B7L74_10035 | F0F1 ATP synthase subunit d | A15A (GCC→GCT) | | | | | | | | | | | | | Transient |
| LOBOINFK_00094 → | B7L74_04980 | dihydroorotate dehydrogenase | R340R (CGC→CGT) | | | | | | | | | | | | | Present in all passages |
| LOBOINFK_01060 → | B7L74_01715 | hypothetical protein | I317I (ATC→ATA) ‡ | | | | | | | | | | | | | Transient |
| LOBOINFK_01886 → | B7L74_01185 | 50S ribosomal protein L11 | T71T (ACC→ACA) | | | | | | | | | | | | | Transient |
| LOBOINFK_01589 → | B7L74_06480 | Na+/H+ antiporter | A126A (GCC→GCA) | | | | | | | | | | | | | Transient |
| LOBOINFK_00910 ← | B7L74_03815 | multidrug transporter AcrB | A422A (GCT→GCA) | | | | | | | | | | | | | Present in all passages |
| LOBOINFK_00801 ← | B7L74_10615 | tryptophan--tRNA ligase | P258P (CCT→CCA) | | | | | | | | | | | | | Transient |
| LOBOINFK_00674 ← | B7L74_09545 | hypothetical protein | I362I (ATT→ATA) | | | | | | | | | | | | | Transient |
| LOBOINFK_00638 ← | B7L74_03305 | leucine dehydrogenase | G269G (GGT→GGA) | | | | | | | | | | | | | Transient |
| LOBOINFK_01573 ← | B7L74_06390 | long-chain fatty acid transporte | I237I (ATT→ATA) | | | | | | | | | | | | | Transient |
| LOBOINFK_00542 → | B7L74_07190 | L,D-transpeptidase | I99I (ATA→ATT) | | | | | | | | | | | | | Transient |
| rhsC ← | B7L74_10905 | Putative deoxyribonuclease Rh | S354S (TCT→TCA) | | | | | | | | | | | | | Transient |
| LOBOINFK_01250 → | B7L74_10035 | F0F1 ATP synthase subunit d | A14A (GCA→GCT) | | | | | | | | | | | | | Transient |
| LOBOINFK_00970 → | B7L74_04140 | MATE family efflux transporte | L375L (TTA→TTG) | | | | | | | | | | | | | Transient |
| LOBOINFK_00966 ← | B7L74_04120 | valine--tRNA ligase | P339P (CCT→CCC) | | | | | | | | | | | | | Transient |
| LOBOINFK_01811 ← | B7L74_03775 | ABC transporter ATP-binding | A214A (GCT→GCC) | | | | | | | | | | | | | Transient |
| LOBOINFK_01673 ← | B7L74_02110 | hypothetical protein | D353D (GAT→GAC) | | | | | | | | | | | | | Transient |
| LOBOINFK_01640 ← | B7L74_06310 | haloalkane dehalogenase | A86A (GCT→GCC) | | | | | | | | | | | | | Transient |
| LOBOINFK_00482 ← | B7L74_06905 | DNA polymerase III subunit a | R713R (CGT→CGC) ‡ | | | | | | | | | | | | | Transient |
| LOBOINFK_01434 → | B7L74_06225 | hypothetical protein | L422L (CTA→CTG) | | | | | | | | | | | | | Transient |
| LOBOINFK_01363 → | B7L74_01440 | uroporphyrinogen decarboxyla | T347T (ACA→ACG) | | | | | | | | | | | | | Transient |
| LOBOINFK_01250 → | B7L74_10035 | F0F1 ATP synthase subunit d | K13K (AAA→AAG) | | | | | | | | | | | | | Transient |
| LOBOINFK_00414 ← | B7L74_00370 | hypothetical protein | F107F (TTT→TTC) | | | | | | | | | | | | | Transient |
| LOBOINFK_00271 ← | L74_01110+B7L74_011 | hypothetical protein | V21V (GTT→GTC) ‡ | | | | | | | | | | | | | Maintained in later passages |
| LOBOINFK_01036 ← | B7L74_02200 | aminoacyl-tRNA hydrolase | R12R (CGT→CGG) | | | | | | | | | | | | | Transient |
| LOBOINFK_01891 → | B7L74_01215 | DNA-directed RNA polymera | R278R (CGA→CGC) | | | | | | | | | | | | | Maintained in later passages |
| LOBOINFK_00629 → | B7L74_03260 | phosphoribosylformylglycinar | G703G (GGA→GGC) | | | | | | | | | | | | | Transient |
| LOBOINFK_01760 ← | B7L74_08665 | hypothetical protein | G2G (GGT→GGG) | | | | | | | | | | | | | Transient |
| LOBOINFK_01421 ← | B7L74_06155 | ATP-dependent Clp protease A | R368R (CGT→CGG) | | | | | | | | | | | | | Transient |
| LOBOINFK_01344 → | B7L74_01325 | 50S ribosomal protein L18 | R9R (CGA→CGC) | | | | | | | | | | | | | Transient |
| LOBOINFK_01155 ← | B7L74_07465 | preprotein translocase subunit | A62A (GCT→GCG) | | | | | | | | | | | | | Transient |
| LOBOINFK_00344 ← | B7L74_00755 | preprotein translocase subunit | V79V (GTT→GTG) | | | | | | | | | | | | | Transient |
| LOBOINFK_00292 ← | B7L74_01020 | hypothetical protein | A13A (GCT→GCG) | | | | | | | | | | | | | Transient |
| LOBOINFK_00229 ← | B7L74_04325 | UTP--glucose-1-phosphate uri | G261G (GGT→GGG) | | | | | | | | | | | | | Transient |
| LOBOINFK_00215 ← | B7L74_04395 | hypothetical protein | P207P (CCT→CCG) | | | | | | | | | | | | | Present in all passages |

| |
|---|
| 5-10 |
| 10-30 |
| 30-50 |
| 50-70 |
| 70-90 |
| 90-99 |
| 100 |

| Gene name from Breseq | Gene name from reference genome | Description/Annotation | Position | Deletion/insertion Mutation | P03 | P05 | P10 | P13 | P16 | P21 | P31 | P42 | P51 | P61 | P67 | Verdict from transcriptomics | Classification by their occurences |
|---|---|---|---|---|---|---|---|---|---|---|---|---|---|---|---|---|---|
| LOBOINFK_00638 ← | B7L74_03305 | leucine dehydrogenase | coding (803/1095 nt) | plusC | | | | | | | | | | | | Early up | Transient |
| LOBOINFK_00638 ← | B7L74_03305 | leucine dehydrogenase | coding (813/1095 nt) | Δ1 bp | | | | | | | | | | | | Early up | Transient |
| LOBOINFK_01455 → | B7L74_08330 | HNH endonuclease | coding (73/636 nt) | +T | | | | | | | | | | | | Continuous down | Maintained in later passages |
| LOBOINFK_01455 → | B7L74_08330 | HNH endonuclease | coding (71/636 nt) | Δ1 bp | | | | | | | | | | | | Continuous down | Maintained in later passages |
| LOBOINFK_01831 ← | B7L74_10225 | phosphotransferase | coding (607/987 nt) | +T | | | | | | | | | | | | Up in P21 | Transient |
| LOBOINFK_01831 ← | B7L74_10225 | phosphotransferase | coding (609/987 nt) | Δ1 bp | | | | | | | | | | | | Up in P21 | Transient |
| LOBOINFK_01916 ← | B7L74_05510 | intracellular signaling protein | coding (617/1230 nt) | plusC | | | | | | | | | | | | Early up | Transient |
| LOBOINFK_01916 ← | B7L74_05510 | intracellular signaling protein | coding (604/1230 nt) | Δ1 bp | | | | | | | | | | | | Early up | Transient |
| LOBOINFK_00838 ← | B7L74_06065 | Bcr/CflA family drug resistance efflux trans | coding (54/1197 nt) | +G | | | | | | | | | | | | Late up | Transient |
| LOBOINFK_00838 ← | B7L74_06065 | Bcr/CflA family drug resistance efflux trans | coding (45/1197 nt) | Δ1 bp | | | | | | | | | | | | Late up | Maintained in later passages |
| hflB ← | B7L74_06975 | ATP-dependent metalloprotease | coding (382-393/1944 nt) | Δ12 bp | | | | | | | | | | | | Late down | Transient |
| LOBOINFK_01656 → | B7L74_02020 | hypothetical protein | coding (3544/4179 nt) | +G | | | | | | | | | | | | Continuous up | Transient |
| LOBOINFK_01656 → | B7L74_02020 | hypothetical protein | coding (3540/4179 nt) | Δ1 bp | | | | | | | | | | | | Continuous up | Transient |
| LOBOINFK_01891 → | B7L74_01215 | DNA-directed RNA polymerase subunit bet | coding (3016/4245 nt) | +G | | | | | | | | | | | | Early down | Transient |
| LOBOINFK_01891 → | B7L74_01215 | DNA-directed RNA polymerase subunit bet | coding (3011/4245 nt) | Δ1 bp | | | | | | | | | | | | Early down | Transient |
| LOBOINFK_01987 → | B7L74_08225 | hypothetical protein | coding (247/330 nt) | +G | | | | | | | | | | | | Early up | Present in all passages |
| LOBOINFK_01987 → | B7L74_08225 | hypothetical protein | coding (241/330 nt) | Δ1 bp | | | | | | | | | | | | Early up | Present in all passages |
| LOBOINFK_01838 → | B7L74_10265 | hypothetical protein | coding (127/675 nt) | CGCACCCATCCATTCTA | | | | | | | | | | | | Early up | Transient |
| LOBOINFK_01838 → | B7L74_10265 | hypothetical protein | coding (337/675 nt) | +A | | | | | | | | | | | | Early up | Transient |
| LOBOINFK_01838 → | B7L74_10265 | hypothetical protein | coding (327/675 nt) | +G | | | | | | | | | | | | Early up | Transient |
| LOBOINFK_01838 → | B7L74_10265 | hypothetical protein | coding (339/675 nt) | Δ1 bp | | | | | | | | | | | | Early up | Transient |
| LOBOINFK_00247 ← | B7L74_04235 | asparagine synthetase B | coding (1188/1899 nt) | +A | | | | | | | | | | | | Down in P10 | Transient |
| LOBOINFK_00247 ← | B7L74_04235 | asparagine synthetase B | coding (1190/1899 nt) | Δ1 bp | | | | | | | | | | | | Down in P10 | Transient |
| INFK_01359 ← / ← LOBOINFK | B7L74_01415 / B7L74_0142 | single-stranded DNA-binding protein/MFS | intergenic (-13/+37) | $(AATCTCCTA)_{1\to2}$ | | | | | | | | | | | | | Transient |
| LOBOINFK_00291 → | B7L74_01025 | hypothetical protein | coding (804/2790 nt) | +A | | | | | | | | | | | | | Transient |
| LOBOINFK_01307 → | B7L74_02755 | lipopolysaccharide assembly protein LapB | coding (168/1167 nt) | +A | | | | | | | | | | | | | Transient |
| INFK_01383 → / → LOBOINFK | B7L74_07670 / B7L74_0766 | cysteine--tRNA ligase/cyclin | intergenic (+155/-181) | +A | | | | | | | | | | | | | Transient |
| LOBOINFK_01503 → | B7L74_06550 | hypothetical protein | coding (80/1221 nt) | +A | | | | | | | | | | | | | Maintained in later passages |
| – / ← LOBOINFK_00443 | / B7L74_06690 | –/hypothetical protein | intergenic (–/+151) | +A | | | | | | | | | | | | | Transient |
| LOBOINFK_01551 ← | B7L74_03535 | short-chain dehydrogenase | coding (163/750 nt) | +A | | | | | | | | | | | | | Present in all passages |
| LOBOINFK_01724 ← / – | B7L74_02705 | N-ethylammeline chlorohydrolase/– | intergenic (-451/–) | +A | | | | | | | | | | | | | Transient |
| LOBOINFK_01724 ← / – | B7L74_02705 | N-ethylammeline chlorohydrolase/– | intergenic (-451/–) | +A | | | | | | | | | | | | | Transient |
| LOBOINFK_00732 → | B7L74_09230 | DNA polymerase I | coding (846/2688 nt) | +A | | | | | | | | | | | | | Transient |
| LOBOINFK_00858 ← | B7L74_05960 | tryptophan synthase subunit alpha | coding (228/804 nt) | +A | | | | | | | | | | | | | Transient |
| INFK_01011 ← / → LOBOINFK | B7L74_02345 / B7L74_0234 | hypothetical protein/histone-like protein Hq1 | intergenic (-281/-10) | +A | | | | | | | | | | | | | Transient |
| LOBOINFK_00104 ← | B7L74_04920 | XRE family transcriptional regulator | coding (75/552 nt) | +G | | | | | | | | | | | | | Transient |
| LOBOINFK_00114 → | B7L74_04875 | NADH pyrophosphatase | coding (736/747 nt) | +G | | | | | | | | | | | | | Maintained in later passages |
| LOBOINFK_00146 → | B7L74_04730 | methylmalonate-semialdehyde dehydrogenas | coding (19/1497 nt) | +G | | | | | | | | | | | | | Maintained in later passages |
| nusA ← | B7L74_07375 | transcription termination/antitermination prot | coding (316/1512 nt) | +G | | | | | | | | | | | | | Present in all passages |
| LOBOINFK_01307 → | B7L74_02755 | lipopolysaccharide assembly protein LapB | coding (168/1167 nt) | +G | | | | | | | | | | | | | Transient |
| fadE_1 → / → LOBOINFK_00573 | B7L74_02980 / B7L74_0299 | acyl-CoA dehydrogenase/acetyl-CoA acetylt | intergenic (+189/-356) | +G | | | | | | | | | | | | | Transient |
| – / → LOBOINFK_00660 | / B7L74_09615 | –/DNA topoisomerase IV subunit A | intergenic (–/-363) | +G | | | | | | | | | | | | | Transient |
| LOBOINFK_00800 ← | B7L74_10610 | 5-methyltetrahydropteroyltriglutamate-- hom | coding (1644/2328 nt) | +G | | | | | | | | | | | | | Transient |
| LOBOINFK_02025 → | B7L74_05700 | lytic transglycosylase | coding (374/1278 nt) | +G | | | | | | | | | | | | | Transient |
| LOBOINFK_00281 ← | B7L74_01070 | transposase | coding (90/276 nt) | +T | | | | | | | | | | | | | Transient |
| INFK_00379 ← / → LOBOINFK | B7L74_00565 / B7L74_0055 | methionine import ATP-binding protein Met | intergenic (-251/-278) | +T | | | | | | | | | | | | | Maintained in later passages |
| LOBOINFK_01307 → | B7L74_02755 | lipopolysaccharide assembly protein LapB | coding (168/1167 nt) | +T | | | | | | | | | | | | | Transient |
| LOBOINFK_01307 → | B7L74_02755 | lipopolysaccharide assembly protein LapB | coding (826/1167 nt) | +T | | | | | | | | | | | | | Transient |
| – / ← LOBOINFK_00443 | / B7L74_06690 | –/hypothetical protein | intergenic (–/+92) | +T | | | | | | | | | | | | | Transient |
| LOBOINFK_00481 ← | B7L74_06900 | EamA family transporter | coding (351/918 nt) | +T | | | | | | | | | | | | | Transient |
| LOBOINFK_01610 → | B7L74_00160 | hydroxymethylglutaryl-CoA reductase | coding (561/1188 nt) | +T | | | | | | | | | | | | | Transient |
| INFK_01761 ← / ← LOBOINFK | B7L74_08670 / B7L74_0867 | hypothetical protein/dCTP deaminase | intergenic (-29/+125) | +T | | | | | | | | | | | | | Transient |
| INFK_01761 ← / ← LOBOINFK | B7L74_08670 / B7L74_0867 | hypothetical protein/dCTP deaminase | intergenic (-127/+27) | +T | | | | | | | | | | | | | Transient |
| INFK_01777 → / → LOBOINFK | B7L74_10790 / B7L74_1080 | 7-cyano-7-deazaguanine synthase/hypothetic | intergenic (+19/-388) | +T | | | | | | | | | | | | | Transient |
| INFK_01778 → / ← LOBOINFK | B7L74_10805 / B7L74_1081 | hypothetical protein/transcription termination | intergenic (+117/+70) | +T | | | | | | | | | | | | | Transient |
| LOBOINFK_01794 → | B7L74_03685 | ABC transporter ATP-binding protein | coding (654/807 nt) | +T | | | | | | | | | | | | | Transient |
| LOBOINFK_00714 → | B7L74_09315 | type I secretion protein TolC | coding (373/1524 nt) | +T | | | | | | | | | | | | | Transient |
| LOBOINFK_02025 → | B7L74_05700 | lytic transglycosylase | coding (374/1278 nt) | +T | | | | | | | | | | | | | Transient |
| – / → LOBOINFK_01680 | / B7L74_09175 | –/transketolase | intergenic (–/-123) | plusC | | | | | | | | | | | | | Transient |
| INFK_00347 ← / ← LOBOINFK | B7L74_00740 / B7L74_0073 | hypothetical protein/UDP-3-O-[3-hydroxym | intergenic (-468/+102) | plusC | | | | | | | | | | | | | Transient |
| INFK_01272 ← / ← LOBOINFK | B7L74_11080 / B7L74_1107 | repA protein/chromosome partitioning prote | intergenic (-44/+13) | plusC | | | | | | | | | | | | | Transient |
| LOBOINFK_01305 → / → lapA | B7L74_02745 / B7L74_0275 | 30S ribosomal protein S1/Lipopolysaccharid | intergenic (+13/-142) | plusC | | | | | | | | | | | | | Present in all passages |
| LOBOINFK_01724 ← / – | B7L74_02705 | N-ethylammeline chlorohydrolase/– | intergenic (-391/–) | plusC | | | | | | | | | | | | | Present in all passages |
| LOBOINFK_00574 → | B7L74_02995 | crotonase | coding (1587/2028 nt) | plusC | | | | | | | | | | | | | Transient |
| INFK_02012 → / ← LOBOINFK | B7L74_10080 / B7L74_1007 | hypothetical protein/ATPase | intergenic (+30/+67) | plusC | | | | | | | | | | | | | Transient |
| LOBOINFK_02014 ← / – | B7L74_10075 | ATPase/– | intergenic (-45/–) | plusC | | | | | | | | | | | | | Transient |
| LOBOINFK_00104 ← | B7L74_04920 | XRE family transcriptional regulator | coding (80/552 nt) | Δ1 bp | | | | | | | | | | | | | Transient |
| LOBOINFK_00114 → | B7L74_04875 | NADH pyrophosphatase | coding (732/747 nt) | Δ1 bp | | | | | | | | | | | | | Maintained in later passages |
| LOBOINFK_00146 → | B7L74_04730 | methylmalonate-semialdehyde dehydrogenas | coding (24/1497 nt) | Δ1 bp | | | | | | | | | | | | | Present in all passages |
| INFK_00225 → / ← LOBOINFK | B7L74_04345 / B7L74_0434 | methylglyoxal synthase/polyribonucleotide n | intergenic (+52/+40) | Δ1 bp | | | | | | | | | | | | | Transient |
| INFK_00225 → / ← LOBOINFK | B7L74_04345 / B7L74_0434 | methylglyoxal synthase/polyribonucleotide n | intergenic (+54/+38) | Δ1 bp | | | | | | | | | | | | | Transient |
| INFK_00225 → / ← LOBOINFK | B7L74_04345 / B7L74_0434 | methylglyoxal synthase/polyribonucleotide n | intergenic (+55/+37) | Δ1 bp | | | | | | | | | | | | | Transient |
| LOBOINFK_00281 ← | B7L74_01070 | transposase | coding (93/276 nt) | Δ1 bp | | | | | | | | | | | | | Transient |
| LOBOINFK_00291 → | B7L74_01025 | hypothetical protein | coding (816/2790 nt) | Δ1 bp | | | | | | | | | | | | | Transient |
| LOBOINFK_00373 → | B7L74_00600 | hypothetical protein | coding (93/213 nt) | Δ1 bp | | | | | | | | | | | | | Transient |
| INFK_00379 ← / → LOBOINFK | B7L74_00565 / B7L74_0055 | methionine import ATP-binding protein Met | intergenic (-254/-275) | Δ1 bp | | | | | | | | | | | | | Maintained in later passages |
| INFK_00407 ← / ← LOBOINFK | B7L74_00405 / B7L74_0040 | hypothetical protein/hypothetical protein | intergenic (-102/+74) | Δ1 bp | | | | | | | | | | | | | Transient |
| nusA ← | B7L74_07375 | transcription termination/antitermination prot | coding (323/1512 nt) | Δ1 bp | | | | | | | | | | | | | Present in all passages |
| LOBOINFK_01255 → / – | B7L74_10060 | bifunctional N-acetylglucosamine-1-phospha | intergenic (+175/–) | Δ1 bp | | | | | | | | | | | | | Transient |
| LOBOINFK_01255 → / – | B7L74_10060 | bifunctional N-acetylglucosamine-1-phospha | intergenic (+176/–) | Δ1 bp | | | | | | | | | | | | | Transient |
| LOBOINFK_01255 → / – | B7L74_10060 | bifunctional N-acetylglucosamine-1-phospha | intergenic (+179/–) | Δ1 bp | | | | | | | | | | | | | Transient |
| LOBOINFK_01305 → / → lapA | B7L74_02745 / B7L74_0275 | 30S ribosomal protein S1/Lipopolysaccharid | intergenic (+17/-138) | Δ1 bp | | | | | | | | | | | | | Transient |
| LOBOINFK_01307 → | B7L74_02755 | lipopolysaccharide assembly protein LapB | coding (169/1167 nt) | Δ1 bp | | | | | | | | | | | | | Transient |
| LOBOINFK_01307 → | B7L74_02755 | lipopolysaccharide assembly protein LapB | coding (170/1167 nt) | Δ1 bp | | | | | | | | | | | | | Transient |
| LOBOINFK_01307 → | B7L74_02755 | lipopolysaccharide assembly protein LapB | coding (171/1167 nt) | Δ1 bp | | | | | | | | | | | | | Transient |
| LOBOINFK_01307 → | B7L74_02755 | lipopolysaccharide assembly protein LapB | coding (584/1167 nt) | Δ1 bp | | | | | | | | | | | | | Transient |
| LOBOINFK_01307 → | B7L74_02755 | lipopolysaccharide assembly protein LapB | coding (939/1167 nt) | Δ1 bp | | | | | | | | | | | | | Transient |
| LOBOINFK_01307 → | B7L74_02755 | lipopolysaccharide assembly protein LapB | coding (1115/1167 nt) | Δ1 bp | | | | | | | | | | | | | Transient |
| LOBOINFK_01313 ← / → tppB_2 | B7L74_02785 / B7L74_0280 | hypothetical protein/MFS transporter | intergenic (-220/-401) | Δ1 bp | | | | | | | | | | | | | Transient |
| – / → LOBOINFK_01440 | / B7L74_08410 | –/hypothetical protein | intergenic (–/-219) | Δ1 bp | | | | | | | | | | | | | Transient |
| LOBOINFK_01503 → | B7L74_06550 | hypothetical protein | coding (83/1221 nt) | Δ1 bp | | | | | | | | | | | | | Transient |
| LOBOINFK_00481 ← | B7L74_06900 | EamA family transporter | coding (355/918 nt) | Δ1 bp | | | | | | | | | | | | | Transient |
| LOBOINFK_01551 ← | B7L74_03535 | short-chain dehydrogenase | coding (166/750 nt) | Δ1 bp | | | | | | | | | | | | | Maintained in later passages |
| INFK_01585 → / ← LOBOINFK | B7L74_06450 / B7L74_0646 | hypothetical protein/polyprenyl synthetase | intergenic (+252/+91) | Δ1 bp | | | | | | | | | | | | | Transient |
| LOBOINFK_01610 → | B7L74_00160 | hydroxymethylglutaryl-CoA reductase | coding (556/1188 nt) | Δ1 bp | | | | | | | | | | | | | Transient |
| INFK_01709 → / ← LOBOINFK | B7L74_02625 / B7L74_0264 | hypothetical protein/MFS transporter | intergenic (+115/+284) | Δ1 bp | | | | | | | | | | | | | Transient |
| LOBOINFK_01724 ← / – | B7L74_02705 | N-ethylammeline chlorohydrolase/– | intergenic (-382/–) | Δ1 bp | | | | | | | | | | | | | Transient |
| INFK_01761 ← / ← LOBOINFK | B7L74_08670 / B7L74_0867 | hypothetical protein/dCTP deaminase | intergenic (-39/+115) | Δ1 bp | | | | | | | | | | | | | Transient |
| INFK_01761 ← / ← LOBOINFK | B7L74_08670 / B7L74_0867 | hypothetical protein/dCTP deaminase | intergenic (-136/+18) | Δ1 bp | | | | | | | | | | | | | Transient |
| fadE_1 → / → LOBOINFK_00573 | B7L74_02980 / B7L74_0299 | acyl-CoA dehydrogenase/acetyl-CoA acetylt | intergenic (+192/-353) | Δ1 bp | | | | | | | | | | | | | Transient |
| LOBOINFK_00574 → | B7L74_02995 | crotonase | coding (1579/2028 nt) | Δ1 bp | | | | | | | | | | | | | Transient |
| LOBOINFK_01942 → / – | B7L74_08845 | molecular chaperone GroEL/– | intergenic (+113/–) | Δ1 bp | | | | | | | | | | | | | Transient |
| – / → LOBOINFK_00660 | / B7L74_09615 | –/DNA topoisomerase IV subunit A | intergenic (–/-368) | Δ1 bp | | | | | | | | | | | | | Transient |
| LOBOINFK_00714 → | B7L74_09315 | type I secretion protein TolC | coding (364/1524 nt) | Δ1 bp | | | | | | | | | | | | | Transient |
| LOBOINFK_00732 → | B7L74_09230 | DNA polymerase I | coding (854/2688 nt) | Δ1 bp | | | | | | | | | | | | | Transient |
| LOBOINFK_02014 ← / – | B7L74_10075 | ATPase/– | intergenic (-54/–) | Δ1 bp | | | | | | | | | | | | | Transient |
| LOBOINFK_00800 ← | B7L74_10610 | 5-methyltetrahydropteroyltriglutamate-- hom | coding (1654/2328 nt) | Δ1 bp | | | | | | | | | | | | | Maintained in later passages |
| LOBOINFK_02025 → | B7L74_05700 | lytic transglycosylase | coding (363/1278 nt) | Δ1 bp | | | | | | | | | | | | | Transient |
| LOBOINFK_02025 → | B7L74_05700 | lytic transglycosylase | coding (364/1278 nt) | Δ1 bp | | | | | | | | | | | | | Transient |
| LOBOINFK_00839 ← / ← phrA | B7L74_06060 / B7L74_0605 | hypothetical protein/Deoxyribodipyrimidine | intergenic (-321/+443) | Δ1 bp | | | | | | | | | | | | | Transient |
| LOBOINFK_00858 ← | B7L74_05960 | tryptophan synthase subunit alpha | coding (241/804 nt) | Δ1 bp | | | | | | | | | | | | | Transient |
| LOBOINFK_00869 ← | B7L74_05905 | tRNA 2-thiouridine(34) synthase MnmA | coding (941/1089 nt) | Δ1 bp | | | | | | | | | | | | | Transient |
| INFK_00892 ← / ← LOBOINFK | B7L74_05780 / B7L74_0577 | curved DNA-binding protein/L,D-transpepti | intergenic (-155/+81) | Δ1 bp | | | | | | | | | | | | | Maintained in later passages |
| LOBOINFK_00373 → | B7L74_00600 | hypothetical protein | coding (49-50/213 nt) | Δ2 bp | | | | | | | | | | | | | Transient |
| LOBOINFK_01307 → | B7L74_02755 | lipopolysaccharide assembly protein LapB | coding (170-172/1167 nt) | Δ3 bp | | | | | | | | | | | | | Transient |
| – / ← LOBOINFK_02034 | / B7L74_10150 | –/IS110 family transposase | intergenic (–/+3) | Δ32 bp | | | | | | | | | | | | | Transient |
| LOBOINFK_00373 → | B7L74_00600 | hypothetical protein | coding (40-46/213 nt) | Δ7 bp | | | | | | | | | | | | | Transient |
| tppB_2 → | B7L74_02805 | MFS transporter | oding (1490-1496/1500 n | Δ7 bp | | | | | | | | | | | | | Maintained in later passages |

| |
|---|
| 5-10 |
| 10-30 |
| 30-50 |
| 50-70 |
| 70-90 |
| 90-99 |
| 100 |

Table S4: Detailed information on 30 hypothetical proteins that are downregulated and could be potential effector proteins. The subcellular localization of these proteins were identified using Psortb. The protein family of the genes were identified using pfam classification and only top hit is listed here. More information on these genes were mined from published and unpublished research articles with full references below.

| Genes | Corresponding CBU(*Cb* NM I RSA493) annotation | Gene annotation | Verdict from transcriptomics | Number of passages with signficant L2FC values | Subcellular localization | Protein family | More information | | | |
|---|---|---|---|---|---|---|---|---|---|---|
| B7L74_04285 | CBU_0841 | Glycosyltransferase family 1 protein | Early down | 5 | Cytoplasmic | Glycos_transf_1, Glycosyl transferases group 1 | This protein was found to be one of the gene that accumulated mutations over a long period of culture and thus was related to change in LPS [1]. | Cytoplasmic | Cytoplasmic | 13 |
| B7L74_01595 | CBU_0304 | Reactive intermediate/imine deaminase | Early down | 9 | Cytoplasmic | Ribonuc_L-PSP, Endoribonuclease L-PSP | This protein was found to be upregulated in ACCM-1 as compared to growth in ACCM-2 after 14 days of growth [2]. | Cytoplasmic | Unknown | 7 |
| B7L74_06350 | CBU_1234 | Uncharacterized protein | Early down | 6 | Cytoplasmic | AAA_14, AAA domain | It has been found as one of the *C. burnetii* T4BSS candidate effector proteins that possesses an E block motif (see result table [3]). | Unknown | Cytoplasmic Membran | 8 |
| B7L74_05675 | CBU_1098 | Hypothetical cytosolic protein (Cig28) | Late down | 4 | Cytoplasmic | DUF762, Coxiella burnetii protein of unknown function (DUF762) | It was found to be one of the immunoreactive proteins [4]. This protein have Pmr motif but shown to be not translocated by Dot system [5]. Has been named as Cig28. | Cytoplasmic Membrane | Periplasmic | 1 |
| B7L74_04275 | CBU_0839 | Glycosyl transferase | Early down | 5 | Cytoplasmic | Glycos_transf_1, Glycosyl transferases group 1 | The mutation in this protein has been shown to changed the LPS from phase 1 to Phase 2 [1]. | Cytoplasmic | | |
| B7L74_04000 | CBU_0787 | AMP-binding protein | Early down | 6 | Cytoplasmic | AMP-binding, AMP-binding enzyme | This protein was shown to be downregulated in the mutated PmrA regulatroy sequence experiment [5]. It was suspected to be regulated by PmrA regulatory sequence but could not be confirmed if it is an effector protein. | Cytoplasmic | | |
| B7L74_03070 | CBU_0591 | Hypothetical cytosolic protein | Early down | 10 | Cytoplasmic | DUF2797, Protein of unknown function (DUF2797) | No information. | Cytoplasmic Membrane | | |
| B7L74_03060 | CBU_0589 | Ferredoxin | Early down | 7 | Cytoplasmic | No hits found. | No information. | Unknown | | |
| B7L74_03480 | CBU_0672 | Hypothetical protein | Early down | 5 | Cytoplasmic | No hits found. | No information. | Cytoplasmic | | |
| B7L74_09270 | CBU_1802 | Hypothetical protein | Early down | 4 | Cytoplasmic | No hits found. | No information. | Cytoplasmic | | |
| B7L74_06740 | CBU_1308 | Phosphohydrolase | Early down | 4 | Cytoplasmic | No hits found. | No information. | Cytoplasmic | | |
| B7L74_01575 | CBU_0300 | YicC family protein | Early down | 4 | Cytoplasmic | YicC_N, YicC-like family, N-terminal region | No information on this protein in Cb. But in general YicC seems to be involved in stress induced mutation in *E.coli*. | Cytoplasmic Membrane | | |
| B7L74_04315 | CBU_0847 | 3-hydroxyacyl-CoA dehydrogenase | Early down | 4 | Cytoplasmic | adh_short, short chain dehydrogenase | No information. | Cytoplasmic Membrane | | |
| B7L74_01105 | CBU_0215 | NlpC-P60 family protein. Peptidase | Early down | 7 | Cytoplasmic Membrane | SH3_6, SH3 domain (SH3b1 type) | It is a peptidase and one of the immunoreactive protein found in [6]. It is predicted to be regulated by PmrA sequences [3]. PmrA has been associated with the effector proteins. | Cytoplasmic Membrane | | |
| B7L74_07575 | CBU_1468 | DUF3971 domain-containing protein | Early down | 4 | Cytoplasmic Membrane | DUF3971, Protein of unknown function | The transposon mutation in this gene has been shown to cause moderate intracellular replication [7]. | Unknown | | |
| B7L74_09610 | CBU_1865 | Hypothetical membrane associated protein | Early down | 7 | Cytoplasmic Membrane | bPH_2, Bacterial PH domain | In a study [4], this protein has been found to be one of the immunoreactive proteins that could have implications in candidate vaccines. | Cytoplasmic | | |
| B7L74_00420 | CBU_0084 | K07014; Uncharacterized protein (Cig3) | Early down | 5 | Cytoplasmic Membrane | DUF3413, Domain of unknown function (DUF3413) | This protein is found to be one of the proteins that are affected by Pmr mutation (and thus predicted effector protein) but was not translocated by T4SS [5]. It has been named as Phosphoglycerol transferase MdoB, Cig3. | Cytoplasmic | | |
| B7L74_03075 | CBU_0592 | Hypothetical membrane spanning protein | Continuous down | 7 | Cytoplasmic Membrane | No hits found. | No information. | Cytoplasmic | | |
| B7L74_10740 | CBU_2073 | Acyltransferase | Early down | 4 | Cytoplasmic Membrane | Acyltransferase, Acyltransferase | No information. | Cytoplasmic Membrane | | |
| B7L74_10350 | CBU_2001 | Hypothetical protein | Continuous down | 4 | Cytoplasmic Membrane | No hits found. | No information. | Cytoplasmic Membrane | | |
| B7L74_01205 | CBU_0230 | Hypothetical protein | Early down | 4 | Cytoplasmic Membrane | No hits found. | No information. | Extracellular | | |
| B7L74_04930 | CBU_0962 | Short-chain dehydrogenase | Continuous down | 4 | Extracellular | adh_short, short chain dehydrogenase | This protein have been found in immunoprecipitation methods in study [8]. | Cytoplasmic Membrane | | |
| B7L74_07865 | CBU_1529 | Peptidase | Early down | 4 | Periplasmic | Peptidase_S9, Prolyl oligopeptidase family | No information. | Periplasmic | | |
| B7L74_10930 | CBUA0012 | Hypothetical protein | Early down | 5 | Unknown | DUF807, Coxiella burnetii protein of unknown function (DUF807) | This protein was found to be present on the same ORFs (with other 5 hypothetical proteins) as other plasmid effectors (which are translocated by Dot system). However, it has been shown to be not translocated by Dot/Icm system [9]. | Cytoplasmic | Extracellular | 1 |
| B7L74_07040 | CBU_1366 | Hypothetical exported protein (Cig40) | Early down | 6 | Unknown | No hits found. | This coiled-coil domain containing protein was found to be regulated by PmrA in *L. pneumophila* [5][10] but was not secreted outside *C.burnetii* [11]. It is predicted to be somehow working as a effector protein. | Unknown | | |
| B7L74_09850 | CBU_1910 | Hypothetical outer membrane protein surface antigen Com1 | Early down | 7 | Unknown | DSBA, DSBA-like thioredoxin domain | This protein has been mentioned in these two studies [12] [13] but no function has been predicted. It is surface antigen Com1 and surface antigens in general helps bacteria to evade into the epithelial cells making this protein to have probable roles in pathogenesis. | Cytoplasmic | | |
| B7L74_01355 | | Hypothetical protein | Continuous down | 8 | Unknown | No hits found. | No information. | Unknown | | |
| B7L74_09265 | CBU_1801 | Hypothetical protein | Early down | 5 | Unknown | No hits found. | Doesn't have much information on this protein except that a lot of mutations has been seen in this operon in the study [14]. | Cytoplasmic | | |
| B7L74_09795 | | DUF1658 domain-containing protein | Early down | 4 | Unknown | No hits found. | No information. | Unknown | | |
| B7L74_09830 | CBU_1906 | Hypothetical protein | Early down | 4 | Unknown | No hits found. | No information. | Unknown | | |

Table S5: Sequencing and assembly statistics for genomes of all passages of *Coxiella burnetii* in this study.

| Passage | Completeness | Contamination | Number of contigs | N50 | L50 | N90 | Total length |
|---|---|---|---|---|---|---|---|
| Passage 10 | 100 | 0 | 64 | 49903 | 11 | 15966 | 1975973 |
| Passage 21 | 100 | 0 | 64 | 75629 | 9 | 16820 | 1980957 |
| Passage 61 | 100 | 0 | 63 | 63550 | 11 | 16844 | 1975487 |
| Passage 51 | 100 | 0 | 63 | 51753 | 10 | 18488 | 1981816 |
| Passage 3 | 100 | 0 | 58 | 66036 | 9 | 19293 | 1975591 |
| Passage 42 | 100 | 0 | 60 | 66036 | 10 | 19293 | 1975994 |
| Passage 67 | 100 | 0 | 59 | 66036 | 10 | 19293 | 1976118 |
| Passage 16 | 100 | 0 | 56 | 71520 | 9 | 19389 | 1976105 |
| Passage 1 | 100 | 0 | 59 | 69758 | 9 | 20406 | 1975592 |
| Passage 5 | 100 | 0 | 57 | 69778 | 9 | 20406 | 1976018 |
| Passage 13 | 100 | 0 | 59 | 71520 | 9 | 20406 | 1976303 |
| Passage 31 | 100 | 0 | 57 | 71520 | 9 | 20406 | 1976112 |
